# Innovative GenExpA software for selecting suitable reference genes for reliable normalization of gene expression in melanoma

**DOI:** 10.1101/2021.05.10.443386

**Authors:** Dorota Hoja-Łukowicz, Dawid Maciążek, Piotr Kościelniak, Marcelina E. Janik

**Affiliations:** Institute of Zoology and Biomedical Research, Jagiellonian University, Gronostajowa 9, 30-387 Krakow, Poland; Smoluchowski Institute of Physics, Jagiellonian University, Łojasiewicza 11, 30-348 Krakow, Poland; Institute of Mathematics, Jagiellonian University, Łojasiewicza 6, 30-348 Kraków, Poland

**Keywords:** *B4GALT*, gene stability, melanoma, NormFinder, RT-qPCR

## Abstract

The algorithms commonly used to select the best stable reference gene in RT-qPCR data analysis have their limitations. We showed that simple selection of the reference gene or pair of genes with the lowest stability value from the pool of potential reference genes - a commonly used approach - is not sufficient to accurately and reliably normalize the target gene transcript and can lead to biologically incorrect conclusions. For reliable assessment of changes in a target gene expression level, we propose our innovative GenExpA software, which works in a manner independent of the experimental model and the normalizer used. GenExpA software selects the best reference by combining the NormFinder algorithm with progressive removal of the least stable gene from the candidate genes in a given experimental model and in the set of daughter models assigned to it. The reliability of references is validated based on the consistency of the statistical analyses of normalized target gene expression levels through all models, described by the coherence score (CS). The use of the CS value imparts a new quality to qPCR analysis because it clarifies how low the stability value of reference must be in order for biologically correct conclusions to be drawn. We tested our method on qPCR data for the *B4GALT* genes family in melanoma, which is characterized by a high mutation rate, and in melanocytes. GenExpA is available at https://github.com/DorotaHojaLukowicz/GenExpA or https://www.sciencemarket.pl/baza-programow-open-source#oferty.

**Highlights:** GenExpA – next-generation software for normalizer selection and validation

The GenExpA tool defines how low the stability value of the reference should be

The GenExpA tool determines the level of gene expression analysis robustness

The GenExpA tool increases the speed of analysis and reduces cost

## Introduction

Achieving reliable results for normalization of the transcript level of a target gene requires an internal reference gene or pair of genes (housekeeping genes, HKGs) with stable expression between the analyzed samples and under different experimental conditions. Moreover, it is believed that the transcript levels of the reference and target genes should be expressed at roughly the same level [1]. However, the transcript levels of many of the HKGs typically used as references may change significantly under physiological or pathological conditions as well as across tissue types and experimental conditions [1, 2, 3, 4, 5, 6, 7, 8]. Therefore the selection and validation of the normalizer are the most critical stages in qPCR data analysis.

Typically, one or more of five algorithms (geNorm [3], NormFinder [9], BestKeeper [10], the comparative ΔCt method [11] and RefFinder; http://150.216.56.64/referencegene.php) are used to select the best stable single or combination of reference genes from a panel of candidate genes, although there are also other proposed algorithms [12] The normalizer with the lowest stability value is considered the best and is then used for calculation of target gene expression in a group of samples. We have recently shown that this commonly accepted approach seems not enough for accurate and reliable target gene transcript normalization and that it may lead to biologically incorrect conclusions [13]. We demonstrated that beyond the parent model of samples of interest (experimental model), it is necessary to construct daughter models (auxiliary models built from the same samples as the experimental model but with fewer samples – combinations of samples of interest without repetition), followed by selection and validation of an appropriate normalizer for each model. Here we have shown that validation of the normalizer is done by determining the coherence score for target gene expression analyses performed for all given models (experimental and daughter models). If the statistical analysis produces inconsistent results for the expression level of a target gene based on a comparison of individual sets of two samples through all models, we have to search for new references with improved stability values via progressive removal of the least stable candidate reference gene in each sample, followed by re-selection and re-validation of new normalizers each time [13]. However, this proposed approach of RT-qPCR data analysis has been a very time-consuming task, requiring the use of several specialist software packages. This problem led us to automate these calculations by developing the GenExpA tool (Gene Expression Analyzer). It is a comprehensive tool based on a previously described workflow for quantified as well as raw qPCR data [13]. Based on the selected reference gene or pair of genes, GenExpA calculates the relative target gene expression level, performs statistical analyses and generates results in the form of graphs and tables. The innovative aspect of GenExpA is that it estimates the coherence of results for each target gene in all designed models. It determines the robustness of normalization and works out the coherence score for the individual target gene. The advantage of this strategy is its ability to determine relative target gene expression based on the consistency of the results in all tested samples, regardless of the experimental model and the reference gene or pair of genes used. Here, based on qPCR data for B4GalT1–B4GalT7 transcripts in melanocytes and melanoma cells, we show how to perform the analysis step by step, and how to solve problems using this new methodological approach.

## Results

A flow chart of target gene expression and statistical analysis of qPCR data using the GenExpA tool is shown in Fig. 1. At the beginning, we uploaded raw qPCR data for candidate reference (*HPRT1, PGK1, RPS23, SNRPA*; set 1) and target (*B4GALT1–B4GALT7*) genes (Supplementary Table S1) as well as quantified qPCR data for candidate reference genes (Supplementary Table S2) obtained for five human cell lines (melanocytes and melanoma cells from different stages of oncogenic progression) comprising the experimental model. It is important to note that here “cell line” means “sample”. Since all PCR reactions were performed in three technical repeats for every one of the three biological replicates, therefore a set of three mean Ct values (for each of the genes) and a set of three means quantified values (for each of the candidate reference genes) were used as inputs for a given cell line. The uploaded quantified values were calculated based on the calibration curve previously prepared for all candidate reference genes in all analyzed cell lines. Working with the GenExpA interface, the potential reference genes were selected from the pool of all analyzed genes presented in the ‘Available reference genes’ window, and a panel of 26 possible models (the experimental model and 25 daughter models which are combinations, without repetition, of 5 cell lines taken 2 or 3 or 4 or 5 at a time) was automatically designed using the ‘Generate combinations’ option. Because of the normal data distribution (data not shown), the Pairwise t-test, Holm adjustment test was chosen from the ‘Statistical model’ bar. Finally, 0 was set in the ‘Remove repetitions’ box that means all four potential reference genes (set 1) were used by NormFinder. The ‘Confidence’ box was set by default at 0.05. The ‘Select best remove for model’ option was left unselected. After clicking on ‘Run calculation’, the GenExpA tool started the analysis. First, the implemented NormFinder algorithm determined the most stable reference gene or pair of genes from four candidate reference genes in each of the 26 models (Table 1, part A; remove repetition level 0). The stability value for the selected reference gene pair *RPS23*/*SNRPA* in the experimental model (model No. 26) was 0.364, and the reference stability values for auxiliary models ranged from 0.077 for reference gene pair *HPRT1*/*RPS23* in model No. 7 to 0.504 for reference gene pair *RPS23*/*SNARPA* in model No. 16 (Table 1, part A). Then the relative expression levels of target genes *B4GALT1–B4GALT7* were determined as relative quantification (RQ) values in each of the 26 models via normalization to the reference gene/genes assigned for these models. After clicking on ‘Export results’, the GenExpA software generated tables summarizing the calculated reference genes’ stability values, the obtained RQ values and the statistical test results, as well as attributed coherence values (Supplementary Table S3). By choosing the ‘Export graph’ option, box-plots representing the medians of the obtained RQ values with statistical significance bars were generated as .png files, each representing a set of target genes in one of the analyzed models (graphs for all models are compiled in Supplementary Fig. S1). In this analysis, GenExpA calculated the coherence score at level 0.90 for four target genes (*B4GALT3, B4GALT5, B4GALT6, B4GALT7*). This value resulted from unreliable/uncertain normalization. GenExpA calculated also the average coherence score at level 0.94.

**Table 1.** Reference genes and their stability values in a panel of the experimental model and daughter models selected by NormFinder software on the base of 4 (part A) and 5 (part B) potential reference genes before (0) and after removing one (1) or two (2) the least stable genes from a given set of 4 and 5 reference genes.

| No. | model | 4 potential reference genes - **part A** |  | 5 potential reference genes - **part B** |  |  |
| --- | --- | --- | --- | --- | --- | --- |
|  |  | remove repetition level |  | remove repetition level |  |  |
|  |  | 0 | 1 | 0 | 1 | 2 |
| 1 | HEMa-LP Mel202 | *HPRT1*<br>0.212 | *HPRT1 SNRPA*<br>**0.090** | *RPS23 SNRPA*<br>0.074 | *GUSB SNRPA*<br>**0.063** | *HPRT1*<br>0.076 |
| 2 | HEMa-LP WM35 | *HPRT1 PGK1*<br>0.248 | *PGK1*<br>**0.157** | *RPS23 PGK1*<br>**0.113** | *HPRT1 RPS23*<br>0.208 | *HPRT1 GUSB*<br>0.225 |
| 3 | HEMa-LP WM793 | *HPRT1*<br>0.310 | *HPRT1 SNRPA*<br>**0.084** | *GUSB*<br>0.093 | *GUSB SNRPA*<br>**0.037** | *GUSB PGK1*<br>0.101 |
| 4 | HEMa-LP WM266-4 | *RPS23 PGK1*<br>**0.252** | *HPRT1 PGK1*<br>0.374 | *RPS23 PGK1*<br>0.145 | *GUSB*<br>**0.131** | *GUSB RPS23*<br>0.177 |
| 5 | Mel202 WM35 | *PGK1*<br>**0.105** | *HPRT1 RPS23*<br>0.121 | *GUSB RPS23*<br>**0.088** | *PGK1*<br>0.105 | *HPRT1 RPS23*<br>0.121 |
| 6 | Mel202 WM793 | *HPRT1 RPS23*<br>0.092 | *HPRT1 SNRPA*<br>**0.056** | *GUSB RPS23*<br>**0.031** | *GUSB SNRPA*<br>0.077 | *HPRT1 SNRPA*<br>0.056 |
| 7 | Mel202 WM266-4 | *HPRT1 RPS23*<br>**0.077** | *HPRT1 RPS23*<br>0.171 | *HPRT1 RPS23*<br>0.091 | *GUSB*<br>0.171 | *RPS23*<br>**0.010** |
| 8 | WM35 WM793 | *SNRPA*<br>0.192 | *PGK1*<br>**0.099** | *GUSB RPS23*<br>**0.085** | *SNRPA*<br>0.192 | *PGK1*<br>0.099 |
| 9 | WM35 WM266-4 | *RPS23*<br>0.112 | *RPS23*<br>**0.078** | *RPS23 PGK1*<br>**0.078** | *RPS23*<br>0.112 | *RPS23*<br>0.078 |
| 10 | WM793 WM266-4 | *HPRT1 RPS23*<br>**0.125** | *HPRT1 PGK1*<br>0.145 | *PGK1 SNRPA*<br>0.159 | *HPRT1 PGK1*<br>0.088 | *GUSB*<br>**0.051** |
| 11 | HEMa-LP Mel202 WM35 | *RPS23 SNRPA*<br>0.251 | *HPRT1 SNRPA*<br>**0.146** | *HPRT1*<br>**0.138** | *HPRT1 GUSB*<br>0.272 | *HPRT1 RPS23*<br>0.333 |
| 12 | HEMa-LP Mel202 WM793 | *HPRT1*<br>0.341 | *HPRT1 SNRPA*<br>**0.084** | *HPRT1*<br>0.257 | *GUSB SNRPA*<br>**0.061** | *HPRT1 SNRPA*<br>0.084 |
| 13 | HEMa-LP Mel202 WM266-4 | *RPS23 SNRPA*<br>**0.261** | *HPRT1 SNRPA*<br>0.356 | *RPS23 SNRPA*<br>**0.128** | *HPRT1 GUSB*<br>0.203 | *GUSB*<br>0.172 |
| 14 | HEMa-LP WM35 WM793 | *HPRT1*<br>0.402 | *PGK1*<br>**0.134** | *HPRT1*<br>0.264 | *HPRT1*<br>0.235 | *PGK1*<br>**0.134** |
| 15 | HEMa-LP WM35 WM266-4 | *HPRT1 PGK1*<br>**0.253** | *HPRT1 SNRPA*<br>0.376 | *RPS23 PGK1*<br>**0.123** | *HPRT1 GUSB*<br>0.241 | *HPRT1 GUSB*<br>0.256 |
| 16 | HEMa-LP WM793 WM266-4 | *RPS23 SNRPA*<br>0.504 | *HPRT1 PGK1*<br>**0.300** | *GUSB*<br>0.331 | *HPRT1 PGK1*<br>**0.085** | *GUSB SNRPA*<br>0.266 |
| 17 | Mel202 WM35 WM793 | *HPRT1 RPS23*<br>0.181 | *HPRT1 SNRPA*<br>**0.106** | *GUSB RPS23*<br>**0.077** | *HPRT1 RPS23*<br>0.181 | *HPRT1 SNRPA*<br>0.106 |
| 18 | Mel202 WM35 WM266-4 | *HPRT1 RPS23*<br>**0.147** | *HPRT1 RPS23*<br>0.185 | *HPRT1 PGK1*<br>0.207 | *HPRT1 RPS23*<br>**0.147** | *HPRT1 RPS23*<br>0.185 |
| 19 | Mel202 WM793 WM266-4 | *HPRT1 RPS23*<br>**0.106** | *HPRT1 PGK1*<br>0.222 | *PGK1 SNRPA*<br>0.200 | *GUSB SNRPA*<br>**0.100** | *GUSB*<br>0.249 |
| 20 | WM35 WM793 WM266-4 | *HPRT1 RPS23*<br>**0.177** | *HPRT1 PGK1*<br>0.187 | *PGK1*<br>0.219 | *HPRT1 PGK1*<br>0.272 | *HPRT1 PGK1*<br>**0.187** |
| 21 | HEMa-LP Mel202 WM35 WM793 | *HPRT1*<br>0.370 | *HPRT1 SNRPA*<br>**0.122** | *HPRT1*<br>0.236 | *HPRT1 PGK1*<br>0.229 | *HPRT1 SNRPA*<br>**0.122** |
| 22 | HEMa-LP Mel202 WM35 WM266-4 | *HPRT1 PGK1*<br>**0.237** | *HPRT1 SNRPA*<br>0.316 | *RPS23 SNRPA*<br>**0.220** | *HPRT1 GUSB*<br>0.264 | *HPRT1 GUSB*<br>0.323 |
| 23 | HEMa-LP Mel202 WM793 WM266-4 | *RPS23 SNRPA*<br>0.403 | *HPRT1 SNRPA*<br>**0.292** | *HPRT1*<br>0.362 | *GUSB SNRPA*<br>**0.086** | *GUSB PGK1*<br>0.284 |
| 24 | HEMa-LP WM35 WM793 WM266-4 | *RPS23 SNRPA*<br>0.436 | *HPRT1 PGK1*<br>**0.299** | *HPRT1*<br>0.394 | *HPRT1 PGK1*<br>**0.239** | *HPRT1 PGK1*<br>0.299 |
| 25 | Mel202 WM35 WM793 WM266-4 | *HPRT1 RPS23*<br>**0.153** | *HPRT1 PGK1*<br>0.219 | *PGK1*<br>0.233 | *HPRT1 RPS23*<br>**0.153** | *HPRT1 PGK1*<br>0.219 |
| 26 | HEMa-LP Mel202 WM35 WM793 WM266-4 | *RPS23 SNRPA*<br>0.364 | *HPRT1 SNRPA*<br>**0.267** | *HPRT1*<br>0.330 | *HPRT1 PGK1*<br>**0.215** | *HPRT1 SNRPA*<br>0.267 |

The differences in the choice of the normalizer or obtained stability values are written in **bold**.

**Figure 1.**
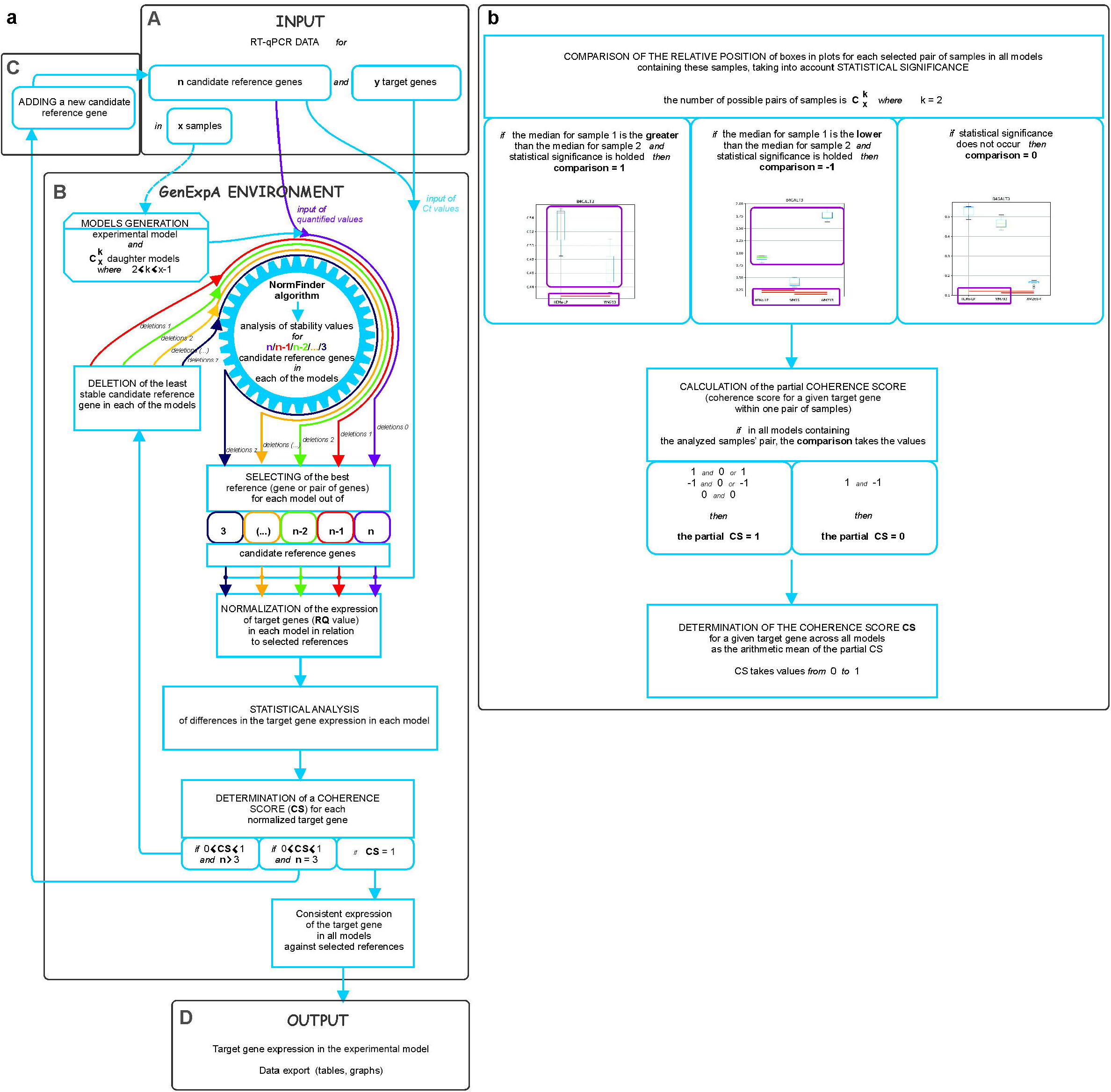
Overview of the GenExpA tool workflow. ***(a)*** (A) The user uploads a table with Ct values for ‘n’ candidate reference genes and **‘**y’ target genes measured in ‘x’ samples. The user uploads a table with calculated quantified values for ‘n’ candidate reference genes. (B) In the setting mode (not shown in this paper; see the GenExpA manual), the user separates ‘n’ potential reference genes from a list of all genes. After clicking on ‘Generate combinations’, the GenExpA tool automatically creates a set of possible ‘z’ models, i.e., ‘z’ models composed of samples of interest, which are combinations without repetitions (order does not matter) of two or more samples from the pool of ‘x’ samples. The ‘z’ models consist of the experimental model and its ‘z-1’ daughter models (auxiliary models); for example, if the experimental model consists of 5 samples, the number of daughter models is 25 (10 models of 2 samples, 10 models of 3 samples, and 5 models of 4 samples). The user unchecks the statistical model, wherein the pairwise t-test with Holm adjustment is recommended as a statistical model for multiple pairs of observations in the case of a normal distribution (to determine the significance of a difference in gene expression between two or more samples). Alternatively, the non-parametric Mann-Whitney test or the ANOVA Kruskal-Wallis test with subsequent post hoc Dunn’s test are recommended for models composed of two unmatched samples or three or more unmatched samples, respectively, in the case of a non-normal distribution [23]. In the setting mode, the user determines the P value indicating the level of statistical significance (default P<0.05;not shown in this paper; see the GenExpA manual). The user starts the automated data analysis by clicking on ‘Run calculations’; it causes the implemented NormFinder algorithm to select, in each of the analyzed models, the best references between ‘n’ potential reference genes (purple line) according to the stability values of their expression in the experimental model and its auxiliary models. The best stability value is referred to the minimal combined inter- and intrasample expression variation of the gene [9]. NormFinder works on Ct values converted to quantified values, which were calculated based on the calibration curve previously prepared for all candidate reference genes. Then the GenExpA tool normalizes the expression of target genes (RQ value) in each model in relation to selected references, conducts a statistical analysis and calculates the coherence score (CS) separately for each target gene (details of how GenExpA determines CS are shown in ***(b)***). If the CS value is equal to 1, the target gene expression analysis is completed. In the case of inconsistent results (CS <1), the user can change the setting parameters by choosing 1 in the ‘Remove repetitions’ box and by marking the option ‘Select best remove for models’ (not shown in this paper; see the GenExpA manual). These new settings cause the least stable HKG in each model to be removed from the pool of ‘n’ candidate reference genes. The user starts the improved analysis by clicking on ‘Run calculations’. The GenExpA tool chooses new references based on a reduced pool of HKGs to ‘n-1’ in each model and conducts a reanalysis (statistic and CS calculation) of the normalized target gene(s) (red line). This approach can be repeated serially (number in ‘Remove repetitions’ box should be increased by 1 each time) when the CS value is below 1 and the pool of reference genes is not less than three (from the green line to the navy blue line). It is important to note that after marking the option 'Select best remove for models’, GenExpA selects the reference for a given model from the level at which the stability value was the lowest. (C) If the gene removal process does not yield consistency of analysis, the user can upload a new input extended with data for an additional HKG or HKGs and perform an improved analysis based on the strategy of removing the least stable HKG until consistency of results is reached. (D) The user can export the results and graphs of all analyses in publication-ready form. The results show coherence scores, the best reference gene or gene pair results, RQ data and statistics. ***(b)*** Flow chart of determination of the coherence score (CS) for a given normalized target gene. First, the GenExpA tool calculates the comparison values for a given target gene in each pair of samples in each model built from these samples. To do this, the software downloads the p-value attributed to a given two samples, and if p≥ 0.05 it assigns the value 0 to the comparison. If p<0.05 and the median of RQ for sample 1 is greater or lower than the median of RQ for sample 2, then comparison = 1 or −1, respectively. The order of samples is fixed. In the aforementioned experimental model, consisting of five samples, where the number of all generated models is 26 (1 experimental model and 25 daughter models), a given pair of samples is present in 8 out of 26 models, so eight comparison values are generated. Next, the software analyzes the obtained comparison values to determine the partial CS value for a given target gene in a given two samples. If the eight comparison values are composed of 1 and −1, then the partial CS=0; in other cases (1 and 1; −1 and −1; 0 and 0; 1 and 0; −1 and 0) the partial CS=1. Finally, the software calculates the CS value for a given target gene as the arithmetic mean of the partial CSs across all pairs of samples (in the aforementioned 26 models there are 10 different pairs of two samples). The resulting CS takes a value between 0 to 1.

(Supplementary Fig. S2). Next, to improve the estimation of the expression of these target genes, the least stable reference gene in each model was removed by setting 1 in the ‘Remove repetitions’ box. Also, marking ‘Select best remove for models’ caused GenExpA to choose the reference from the level of removal for which the stability value was lower in a given model.

This approach raised the average coherence score of the analyses of all *B4GALT* target genes from 0.94 to 0.99 and gave lower stability values of the normalizers in 15 of the 26 models. Now these stability values ranged between 0.056 for *HPRT1/SNRPA* in model No. 6 and 0.300 for *HPRT1/PGK1* in model No. 16 (Table 1, part A; remove repetition level 1). The stability value of the normalizer (*HPRT1/SNRPA*) for the experimental model (model No. 26) also was improved: 0.267. The coherence scores for *B4GALT1, B4GALT2* and *B4GALT4* were kept at 1, and for genes *B4GALT3, B4GALT5* and *B4GALT7* reached the value 1, confirming the consistency of the analysis of the expression levels of these genes. At this point of the analysis, the coherence score for gene *B4GALT6* was below 1 (Supplementary Fig. S3), suggesting that the selected references are not suitable for accurate and reliable normalization of the B4GALT6 transcript level (Supplementary Table S4 contains the tables and Supplementary Fig. S4 the box-plots generated in this analysis). Executed removal of the least stable gene reduced the pool of potential reference genes to three, that is, the minimum number of genes required for NormFinder selection [9], thus ending the possibility of removing the next-weakest reference gene. To continue the analysis, we enlarged the pool of candidate reference genes by adding a new HKG, *GUSB* (inputs with mean Ct and quantified qPCR data are presented in Supplementary Tables S5 and S6). Then the best reference gene or pair of genes from five candidate HKGs was selected (digit 0 entered in the ‘Remove repetitions’ box and without choosing ‘Select best remove for models’ box). The selected references possessed stability values ranging from 0.031 for *GUSB/RPS23* (auxiliary model No. 6) to 0.394 for *HPRT1* (auxiliary model No. 24), and 0.330 in the experimental model (model No. 26) (Table 1, part B; remove repetition level 0). Validation of the selected references confirmed the consistency of expression analysis for genes *B4GALT1– B4GALT3* and *B4GALT5* (Supplementary Fig. S5, Supplementary Table S7, Supplementary Fig. S6). Removing the least stable gene from five candidate reference genes (digit 1 entered in ‘Remove repetitions’ box and choosing ‘Select best remove for models’ box) (Table 1, part B; remove repetition level 1; Supplementary Fig. S7) allowed us to assign better stability values to 11 of the 26 models (Table 1, part B; remove repetition level 1), however, the coherence score for *B4GALT4* was still below 1 (Supplementary Table S8; Supplementary Fig. S8). Sequential removal of the two least stable potential reference genes (digit 2 entered in ‘Remove repetitions’ box and choosing ‘Select best remove for models’ box) did not complete the analysis for *B4GALT4* (Supplementary Fig. S9). Supplementary Table S9 contains the tables and Supplementary Fig. S10 the box-plots corresponding to this analysis. At this point of the analysis, however, the coherence score dropped to 0.9 for gene *B4GALT6*.

Having the Ct values for five potential reference genes, we conducted similar selections of references from four other possible combinations of 4 candidate reference genes: *RPS23, SNRPA, HPRT1* and *GUSB* (set 2); *SNRPA, HPRT1, GUSB* and *PGK1* (set 3); *HPRT1, GUSB, PGK1* and *RPS23* (set 4); and *GUSB, PGK1, RPS23* and *SNRPA* (set 5), before and after rejection of the weakest reference genes. The results obtained from normalization of target gene expression in the 26 models were exported from the GenExpA tool as Supplementary Tables S10–S17 and corresponding Supplementary Fig. S11–S18. A summary of all target gene expression analyses in the experimental model of interest, based on the selection of references from four (sets 1–5) and five candidate reference genes, is presented in Supplementary Fig. S19. Based on it, we concluded that if, at different levels of rejection of the weakest reference gene from sets of four or five candidate reference genes, the analysis of the expression of a given target gene ended with CS = 1, the obtained results do not contradict each other (for example, compare the expression levels of *B4GALT1* in sets 1–5 and the set of five candidate reference genes – box-plots 1–13 in Supplementary Fig. S19). However, starting the analysis with a higher number of potential reference genes creates an opportunity to select the reference gene with a lower stability value in the experimental model as well as auxiliary models, and therefore gives a more robust analysis of target gene expression. For example, the use of a set of five candidate reference genes (*PGK1, RPS23, SNRPA, HPRT1, GUSB*) with sequential removal of the two least stable potential reference genes gave the lowest stability value of the selected reference gene for the experimental model (0.215) and auxiliary models (0.010–0.239). On the other hand, we noticed that box-plots 13 and 91 show exactly the same results as, respectively, box-plots 2, 5, 6, 10, 12 and 80, 83, 84, 90 (compare the medians of relative quantity (RQ) and statistical significance between samples in Supplementary Fig. S19). It means that in the case of *B4GALT1* and *B4GALT7* gene expression analysis, the use of four HKGs from sets 2 or 5 with rejection of the most unstable HKG, or from set 3, is sufficient. The same is true for genes *B4GALT2* and *B4GALT3* and set 3 (compare box-plots 26 and 39 with 18, 19and 31, 32, respectively, in Supplementary Fig. S19). Similar relationships occur for gene *B4GALT5* and sets 3 and 1 after rejection of the most unstable HKG (compare box-plots 78 with 57, 58 and 54 in Supplementary Fig. S19). Therefore, adding a fifth housekeeping gene is mandatory to confirm the robustness of an analysis conducted with the above-mentioned sets of four candidate reference genes. Interestingly, the lower the HKG stability value, the better the CS value, that is, the more robust the analysis, potentially much more correct biologically.

On the other hand, selection of the normalizer based on five potential reference genes and with ‘Remove repetition’ set at 2 led to an inconsistent analysis for the relative expression of gene *B4GALT6* (box-plot 78 in Supplementary Fig. S19), although the same box-plots were generated with ‘Remove repetition’ set at 1 as well as in the case of set 3 with ‘Remove repetition’ set at 0 or 1 (compare box-plots 70, 71, 77 and 78 in Supplementary Fig. S19), and the reference stability values for the experimental model remained the same (0.215). It means that if a more robust analysis is needed, to improve its consistency another HKG/s should be introduced into the pool of candidate reference genes. Another approach may be to complete the analysis with ‘Remove repetition’ set at 1 (box-plot 77 in Supplementary Fig. S19) with indication of the CS values and stability parameters for the reference genes selected in the experimental and auxiliary models.

We want to emphasize that simple application of the NormFinder algorithm (‘Remove repetition’ 0) may lead to a biological misinterpretation of the obtained results, as it strongly depends on the choice of a set of potential reference genes. For example, sets 1 and 5 showed significantly higher relative expression of gene *B4GALT6* in WM793 cells than in WM266-4 cells. In turn, sets 2, 4 and the set of five HKGs showed, in these two cell lines, an inverse relationship between the *B4GALT6* gene expression that was also statistically significant (compare box-plots 66 and 74 with box-plots 68, 72 and 76 in Supplementary Fig. S19). Moreover, the relative expression levels of *B4GALT5, B4GALT6* and *B4GALT* 7 were significantly lower in cell line WM35 than in cell line WM266-4, as shown in sets 2 and 5 of candidate reference genes (compare box-plots 55, 63 and 68, 76 and 81, 89 with 59 and 72 and 85 in Supplementary Fig. S19), or significantly higher as shown in set 4 (box-plot 72 in Supplementary Fig. S19).

Confusing results are also observed for gene *B4GALT3*: in sets 1 (box-plot 27 in Supplementary Fig. S19), 2 (box-plot 29 in Supplementary Fig. S19), 3 (box-plot 31 in Supplementary Fig. S19), 5 (box-plot 35 in Supplementary Fig. S19) and in set of five candidate reference genes (box-plot 37 in Supplementary Fig. S19) its relative expression level was significantly lower in cell line WM35 than in cell line WM793, but in set 4 the relationship was the reverse (box-plot 33 in Supplementary Fig. S19). Similar for the gene *B4GALT6*: in sets 1 (box-plot 66 in Supplementary Fig. S19) and 5 (box-plot 74 in Supplementary Fig. S19) its relative expression level was significantly lower in cell line WM35 than in cell line WM793, but in set 4 the relationship was the reverse (box-plot 72 in Supplementary Fig. S19). In addition, a larger pool of candidate reference genes does not always lead to selecting the better normalizer in simple application of the NormFinder algorithm (compare the reference stability values for the experimental model and auxiliary models in set 3 and the set of five candidate reference genes; ‘Remove repetition’ 0). In contrast, our method based on gradual removal of the gene with the highest variability of expression leads to selection of a normalizer with a lower stability value (compare the reference stability values for the experimental model and auxiliary models in sets 1–5 with ‘Remove repetition’ 1 and the set of five candidate reference genes with ‘Remove repetition’ 2).

Figure 2 shows the results of GenExpA analysis for the expression levels of target genes *B4GALT1–B4GALT7* in the analyzed experimental model. We obtained consistent and complete analyses for all target genes under the given stability values for the experimental and auxiliary models. These presented results demonstrate that in melanoma cells, in particular metastatic melanoma cell line WM266-4, the expression of all *B4GALT* genes decreases relative to that of melanocytes. An exception is melanoma cell line WM793, in which the expression levels of genes *B4GALT2* and *B4GALT4* show no significant differences as compared to those for melanocytes.

**Figure 2.**
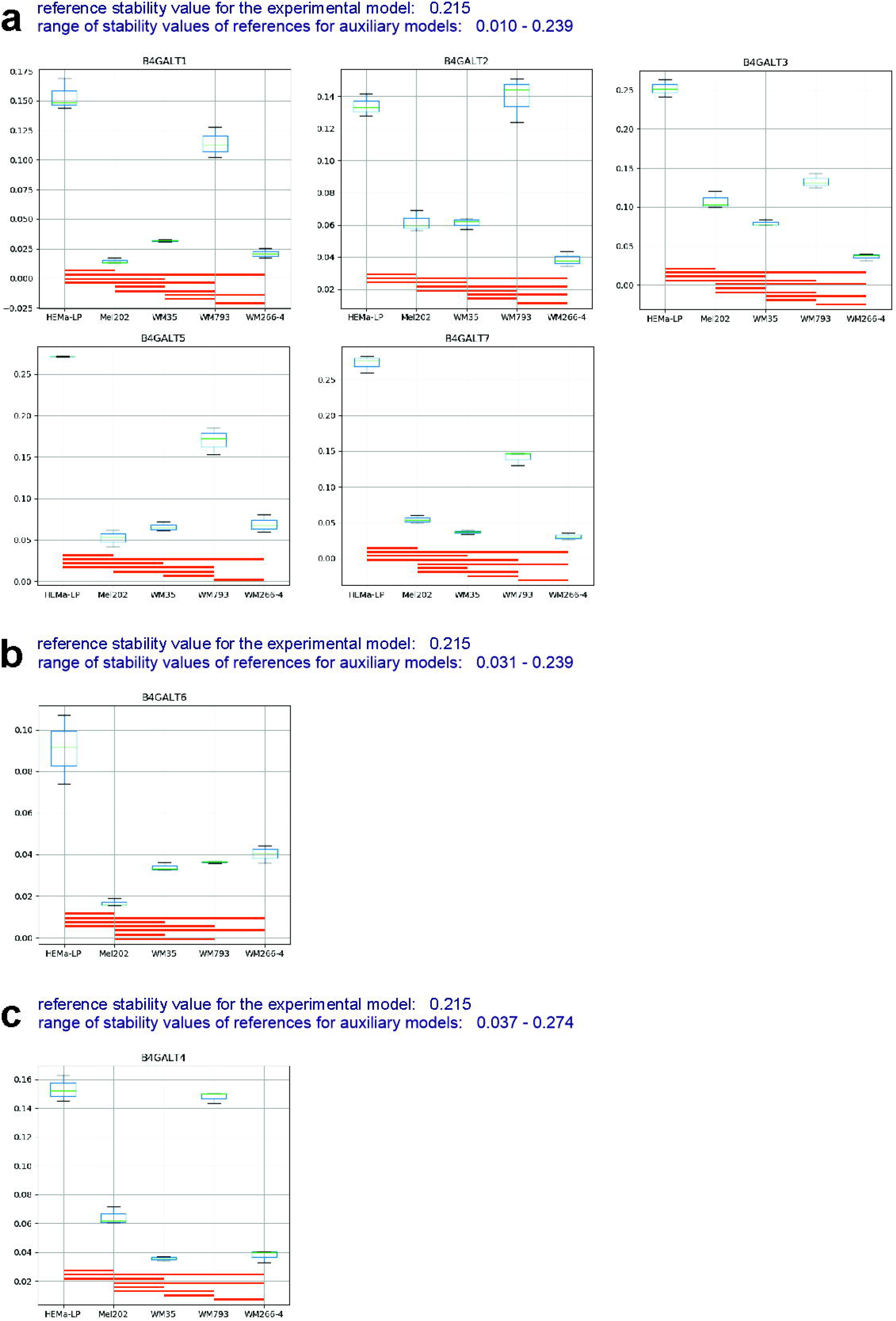
Medians of relative quantity (RQ) for target genes *B4GALT1*–*B4GALT7* in the experimental model normalized to a reference gene or pair of reference genes with the best stability values resulting from GenExpA analysis based on a set of five candidate reference genes and remove repetition level 2 (**a**) or remove repetition level 1 (**b**) or based on a set 3 (**c**) (see Table 1, part B). The statistical analyses used the pairwise t-test with Holm adjustment. Red line represents statistical significance, P<0.05.

## Discussion

Obtaining reliable results from the PCR reaction is the outcome of many factors related to handling of the material, beginning with the step of RNA/DNA isolation and ending in analysis of the results. That is why the MIQE guidelines were proposed for transparent reporting of experimental details in all publication prepared with the use of qPCR technique [14]. The reaction needs to be standardized in molecular analyses of all kinds of biological materials [15, 16, 17, 18]. Over the past decade the MIQE guidelines have not come into common use. Dijkstra et al. [15] made a critical analysis of qPCR result normalization in 179 publications regarding colorectal cancer. They showed that only 3% of these studies applied qPCR methodology based on the use and validation of multiple reference genes. Several statistical are used to select the best stable single gene or combination of reference genes from a panel of candidate genes. It should be stressed that each of these algorithms has its limitations which affect the ranking of candidate reference genes and finally the selection of the best normalizer [19, 20]. Recent studies have proposed the use of at least three different methods, but the problem of how to interpret conflicting results obtained from the use of different statistical methods still puzzles researchers. They try to integrate a few algorithms to average the stability ranks; for example, the RefFinder algorithm calculates the geometric mean of ranking values obtained by four other algorithms [19]. But these statistical methods differ from each other; averaging the ranks can lead to a suboptimal assessment of stable reference genes [20]. It is commonly accepted that the lower the stability value of the reference, the greater the certainty of correct determination of the relative expression of the target gene. Yet it has not been clarified how low the stability value must be. In this paper we showed that the procedure for choosing a suitable reference involves two steps: applying an algorithm for best reference gene selection (here, NormFinder), and checking whether the stability level of the selected reference is low enough to lead to a correct biological interpretation of relative target gene expression (here, determination of CS for the analysis). Through the application of these two steps, our method validates the chosen reference based on determination of the coherence score for the normalized target gene. In our method we used the model-dependent NormFinder algorithm, and by simulating the removal of the least stable gene from a set of candidate genes in the experimental as well as daughter models we were able to obtain more stable normalizers for each model. Previously, this approach helped us to exclude incorrect normalizations and biological misinterpretations concerning the alteration of *ERAP1* gene expression during melanoma progression [13]. We chose the NormFinder algorithm because it calculates the stability of a reference gene or pair of genes by analysing the variation of its expression within and between samples. However, genes with high overall variation influence the stability ranking of candidate reference genes. Herein we have demonstrated that simple application of NormFinder (‘Remove repetition’ 0) may lead to biological misinterpretation of the results, because it depends on the HKG set used to select the reference. Our method eliminates the influence of the most variable gene or genes on the stability rankings of all other candidate reference genes. In addition, breaking down the experimental model into daughter models (auxiliary models) allows us to check whether a uniform trend of target gene expression among particular cell lines is maintained. This trend decides the consistency of the analysis and is described by coherence score CS=1. Obtaining full consistency of the analysis is tantamount to its completion. As we suggest here, based on our *B4GALT* gene expression analysis, the lower the stability value, the better the coherence of the obtained results, which, we propose, is related to their biological correctness. If, despite the application of our strategy, a uniform trend of target gene expression among particular cell lines has not been achieved, introducing a successive HKG/s, along with repeated rejection of the weakest gene, can lead to selection of the better reference and, in turn, biologically correct conclusions. It is important to note that once selected, a reference for a given model will not necessarily be selected once again in an experiment repeated for this model after some time, even if the pool of candidate reference genes used is exactly the same. This is because the gene expression pattern at the time point of sample collection depends on the current rate of cell mass growth and on microenvironmental conditions [21, 22].

## Conclusions

Here we have presented an expanded version of our already published method for RT-qPCR data analysis implemented in the GeneExpA tool – software for reference gene validation and automatic calculation of target gene expression. GenExpA will assign a CS value ranging from 0 to 1 to each analysis; a fully coherent analysis is described by a CS value of 1. We suggest starting the analysis with a set of at least four potential HKGs. If the results are not satisfactory – if the CS value is below 1 – introducing another candidate HKG/s to the set of candidate reference genes may improve them. However, even if CS is equal to 1 for a given target gene, enlarging the pool of candidate reference genes can lead to an analysis that is more robust in terms of between-sample statistical significance in the experimental model. Interestingly, our workflow, including full target gene analysis in order to validate normalizer, shows that the lower stability value of the selected reference gene or pair of genes is related to the biological correctness of the results. Moreover, our workflow is the first one that allows the user to define the required stability value needed to draw biologically correct conclusions. For clarity of the biological conclusions drawn, the description of the results should give stability values assigned to the experimental model and auxiliary models; this may help other researchers to compare their results and understand any differences between their outcomes and yours. GenExpA software can carry out an analysis of many genes independently at the same time.

## Materials and methods

### Cell culture

Human epidermal melanocytes (adult, HEMa-LP) were obtained from Gibco (Life Technologies). Human melanoma cell lines Mel202 (uveal primary), WM793 (cutaneous primary, vertical growth phase) and WM266-4 (skin metastasis) were from the ESTDAB Melanoma Cell Bank (Tübingen, Germany). Human melanoma cell line WM35 (cutaneous primary, radial/vertical growth phase) was kindly donated by Prof. A. Mackiewicz of the Department of Cancer Immunology, University School of Medical Sciences, Greater Poland Cancer Center, Poznań, Poland. Cell lines were cultured as described previously [12]. All cultures were verified mycoplasma-free using the MycoAlert mycoplasma detection kit (Lonza).

### Reverse transcription qPCR

Total RNA extraction, the reverse-transcription reaction and real-time qPCR reactions were performed as previously described [12]. RNA and cDNA concentrations as well as their quality were assessed with a NanoDrop 2000 spectrophotometer (Supplementary Data). Housekeeping (*GUSB, HPRT1, PGK1, RPS23, SNRPA*) and target (*B4GALT1–7*) gene-specific mRNAs were amplified with the use of TaqMan Gene Expression Assays, as listed in Table 2. All reactions were performed in three biological and three technical replicates. The reaction results were analyzed with StepOne Software ver. 2.0. Raw Ct values were calculated using StepOne ver. 2.0, applying automatic threshold and baseline settings.

**Table 2.** Summary of candidate reference and target genes

| Symbol | Gene name<br>(Assay ID, TaqMan probes;<br>Amplicon size bp) | Location<br>(GeneCards) | Description |
| --- | --- | --- | --- |
| **Candidate reference genes** |  |  |  |
| *GUSB* | Glucuronidase Beta (Hs00939627_m1;<br>96 bp) | 7q11.21 | Glycosaminoglycan metabolism |
| *HPRT1* | Hypoxanthine Phosphoribosyltransferase<br>1 (Hs99999909_m1; 100 bp) | Xq26.2-q26.3 | Generation of purine nucleotides |
| *PGK1* | Phosphoglycerate Kinase 1<br>(Hs00943178_g1; 73 bp) | Xq21.1 | Metabolic pathway of glycolysis |
| *RPS23* | Ribosomal Protein S23 (Hs01922548_s1;<br>90 bp) | 5q14.2 | poly(A) RNA binding and<br>structural constituent of ribosome |
| *SNRPA* | Small Nuclear Ribonucleoprotein<br>Polypeptide A (Hs00190231_m1; 123 bp) | 19q13.2 | Splicing of cellular pre-mRNAs |
| **Target genes** |  |  |  |
| *B4GALT1* | Beta-1,4-Galactosyltransferase 1<br>(Hs00155245_m1; 70 bp) | 9p21.1 | Biosynthesis of different<br>glycoconjugates and saccharide<br>structures |
| *B4GALT2* | Beta-1,4-Galactosyltransferase 2<br>(Hs00243566_m1; 57 bp) | 1p34.1 |  |
| *B4GALT3* | Beta-1,4-Galactosyltransferase 3<br>(Hs00937515_g1; 68 bp) | 1q23.3 |  |
| *B4GALT4* | Beta-1,4-Galactosyltransferase 4<br>(Hs00186850_m1; 70 bp) | 3q13.32 |  |
| *B4GALT5* | Beta-1,4-Galactosyltransferase 5<br>(Hs00941041_m1; 66 bp) | 20q13.13 |  |
| *B4GALT6* | Beta-1,4-Galactosyltransferase 6<br>(Hs00999574_m1; 69 bp) | 18q11.1 |  |
| *B4GALT7* | Beta-1,4-Galactosyltransferase 7<br>(Hs01011260_g1; 74 bp) | 5q35.3 |  |

### RT-qPCR data analysis

The GenExpA tool was used for fully automated qPCR data analysis. The gene or combination of two HKGs with the lowest stability value were selected as the best reference by combining the model-based variance estimation with progressive removal of the least stable reference gene. For statistical analysis, the pairwise t-test with Holm adjustment was used. P values less than 0.05 were taken to indicate statistical significance (P<0.05).

## Data availability

The data used to support the findings of this study are included within the article.

## Acknowledgments

Michael Jacobs line-edited the paper for submission. This work was supported by the National Science Centre, Poland (grant number 2016/21/B/NZ3/00348) to D.H.-Ł; the Institute of Zoology and Biomedical Research, Jagiellonian University (grant number N18/DBS/000007). We thank Małgorzata Dutka (Department of Molecular Biophysics, Faculty of Biochemistry, Biophysics and Biotechnology of the Jagiellonian University) for reading the manuscript and valuable comments.

## Author Contributions

D.H-Ł and M.E.J wrote the manuscript, supervised the writing of the software, tested the software and performed experiments. D.M wrote the software and the user manual. D.H-Ł prepared figures and acquired the financial support. P.K designed the statistical method. All authors have given approval to the final version of the manuscript.

