## Supplementary material for "Innovative GenExpA software for selecting suitable reference genes for reliable normalization of gene expression in melanoma": Supplementary Data.pdf

Supplementary Data. NanoDrop spectrophotometer data of RNA purity. The number in parentheses refers to the biological replicate.

| sample | RNA Conc. [ng/µl] | A260 | A280 | 260/280 | spectra |
|---|---|---|---|---|---|
| HEMa-LP (1) | 162.6 | 4.065 | 1.925 | 2.11 | 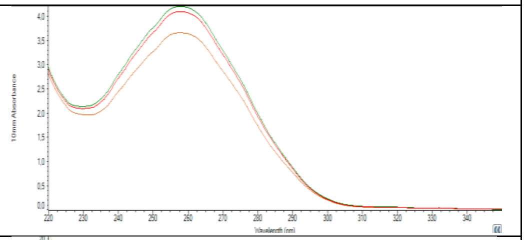 |
| HEMa-LP (2) | 166.8 | 4.169 | 1.984 | 2.10 | |
| HEMa-LP (3) | 145.2 | 3.630 | 1.726 | 2.10 | |
| Mel202 (1) | 582 | 14.551 | 6.973 | 2.09 | 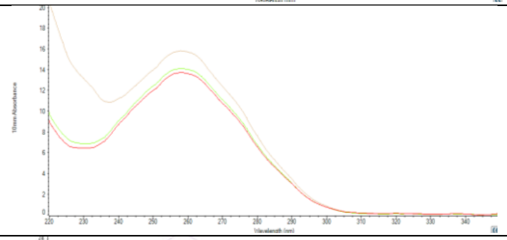 |
| Mel202 (2) | 680.7 | 17.017 | 7.966 | 2.14 | |
| Mel202 (3) | 586.2 | 14.656 | 7.014 | 2.09 | |
| WM35 (1) | 695.8 | 21.085 | 11.375 | 1.85 | 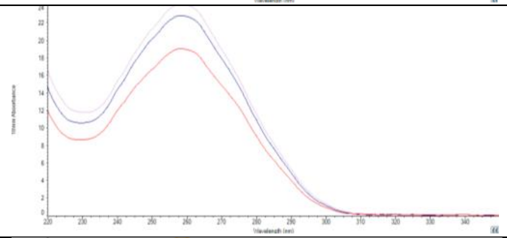 |
| WM35 (2) | 729.3 | 22.101 | 11.874 | 1.86 | |
| WM35 (3) | 930.1 | 28.185 | 15.223 | 1.85 | |
| WM793 (1) | 544.8 | 13.619 | 6.556 | 2.08 | 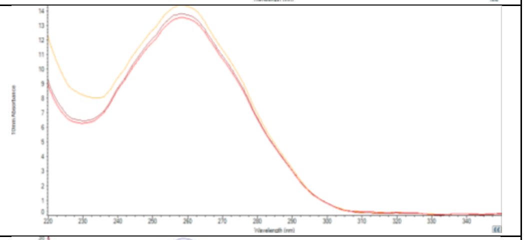 |
| WM793 (2) | 569.7 | 14.242 | 6.812 | 2.09 | |
| WM793 (3) | 590.6 | 14.766 | 7.103 | 2.08 | |
| WM266-4 (1) | 835.4 | 20.886 | 9.953 | 2.10 | 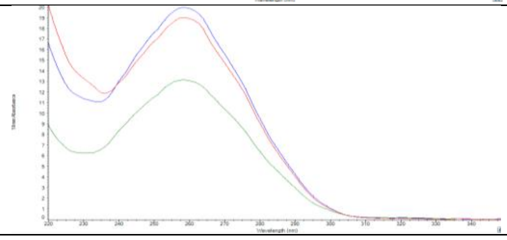 |
| WM266-4 (2) | 543.8 | 13.595 | 6.524 | 2.08 | |
| WM266-4 (3) | 782.2 | 19.554 | 9.326 | 2.10 | |
