## Supplementary material for "Innovative GenExpA software for selecting suitable reference genes for reliable normalization of gene expression in melanoma": Supplementary Figures S1-S19.pdf

### Model No. 1

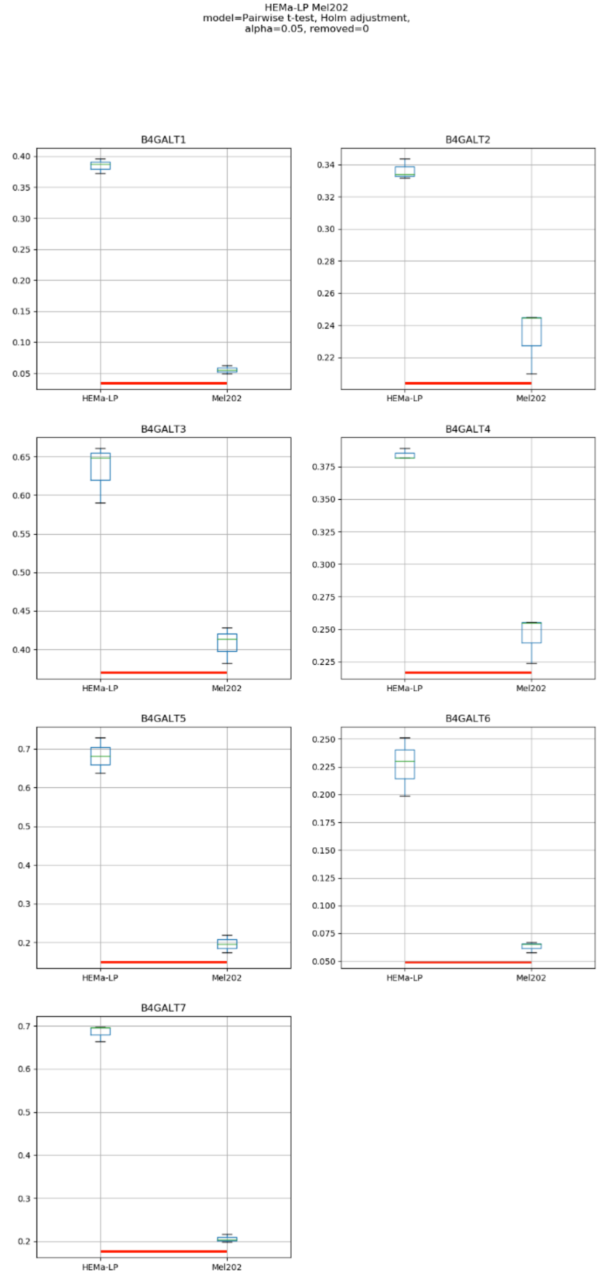

HEMa-LP Mel202
model=Pairwise t-test, Holm adjustment,
alpha=0.05, removed=0
B4GALT1
B4GALT2
B4GALT3
B4GALT4
B4GALT5
B4GALT6
B4GALT7
HEMa-LP
Mel202

### Model No. 2

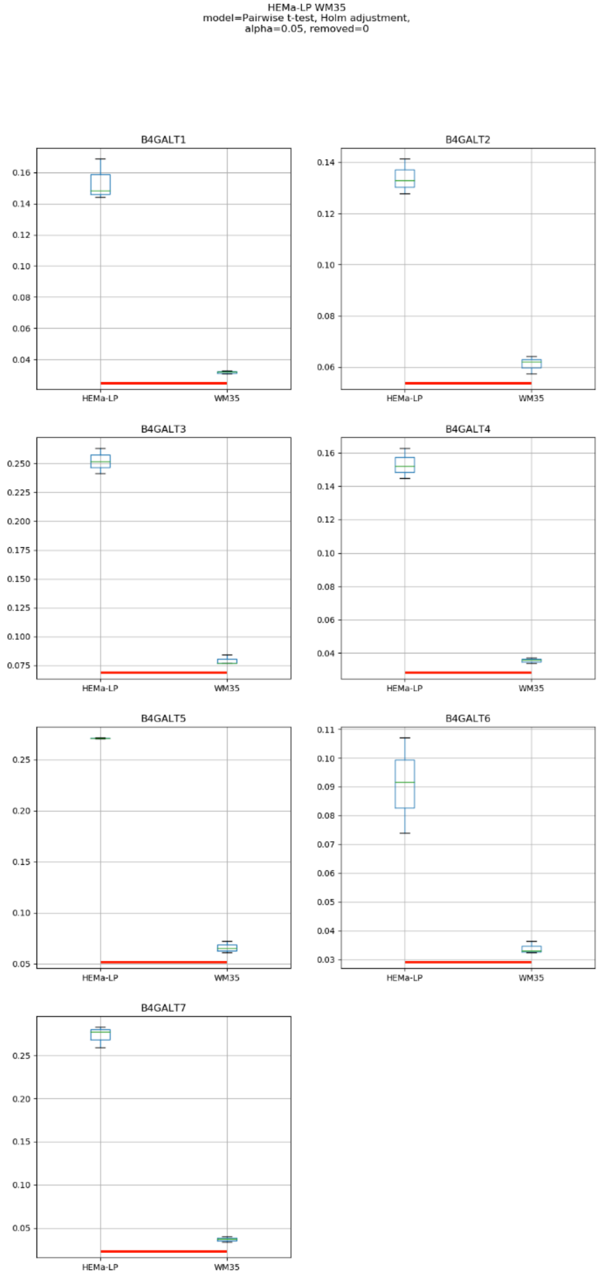

HEMa-LP WM35
model=Pairwise t-test, Holm adjustment,
alpha=0.05, removed=0
B4GALT1
B4GALT2
B4GALT3
B4GALT4
B4GALT5
B4GALT6
B4GALT7
HEMa-LP
WM35

Model No. 3

HEMa-LP WM793
Mann-Whitney/Kruska-Wallis, post: Dunn's test,
alpha=0.05, removed=0

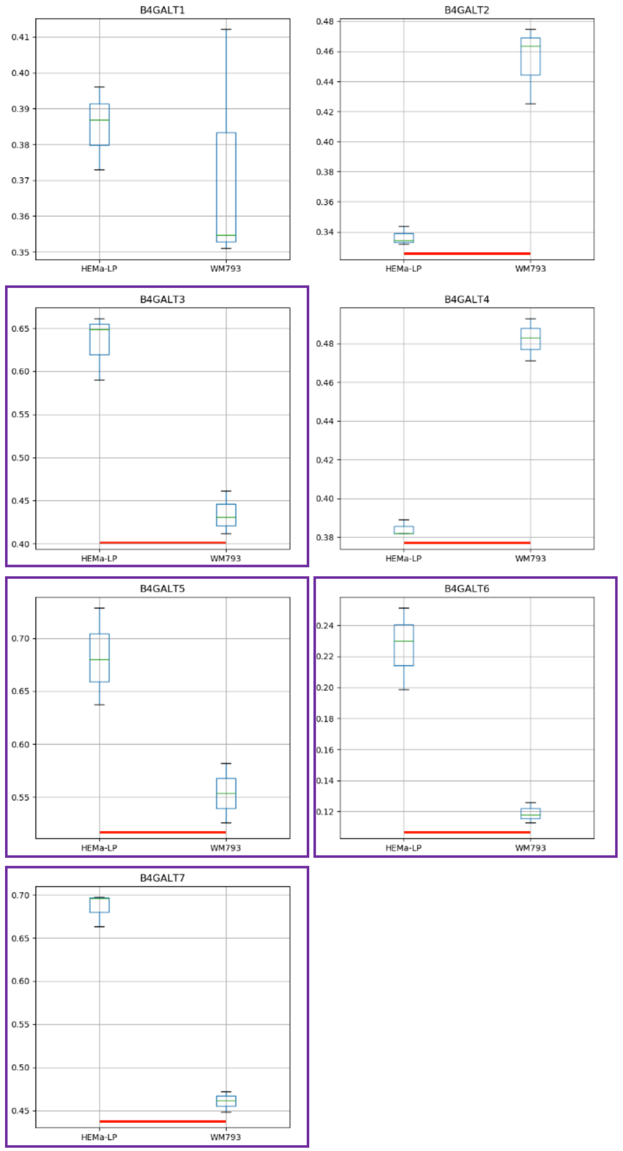

### Model No. 4

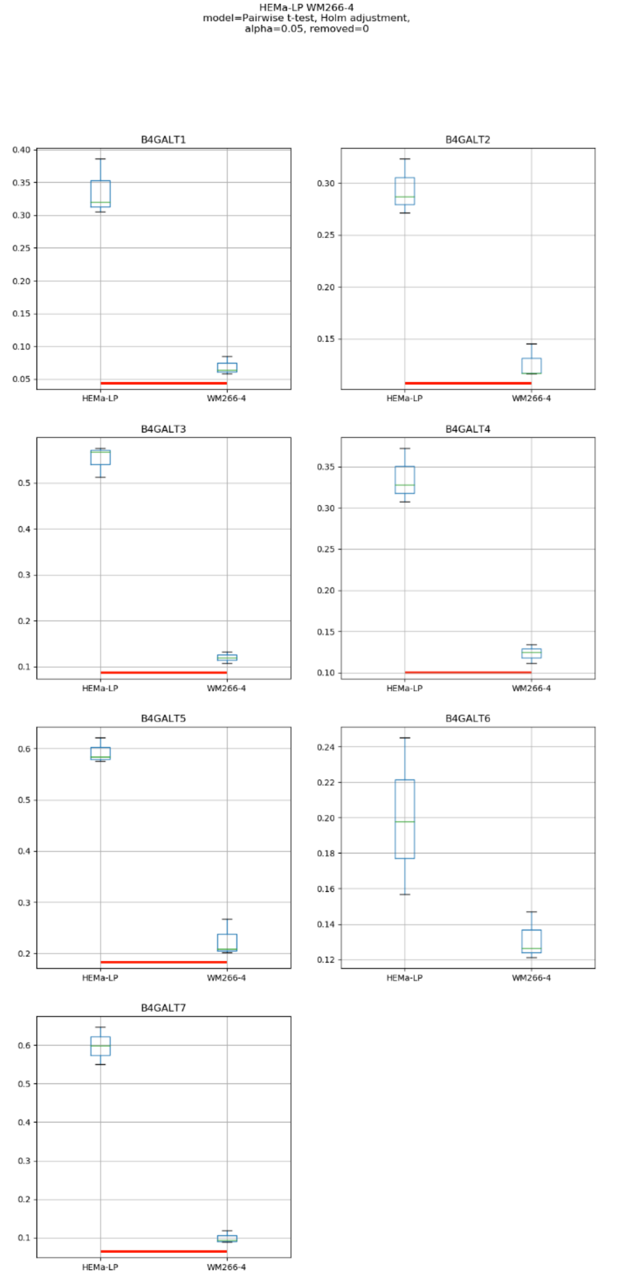

HEMa-LP WM266-4
model=Pairwise t-test, Holm adjustment,
alpha=0.05, removed=0
B4GALT1
B4GALT2
B4GALT3
B4GALT4
B4GALT5
B4GALT6
B4GALT7
HEMa-LP
WM266-4

### Model No. 5

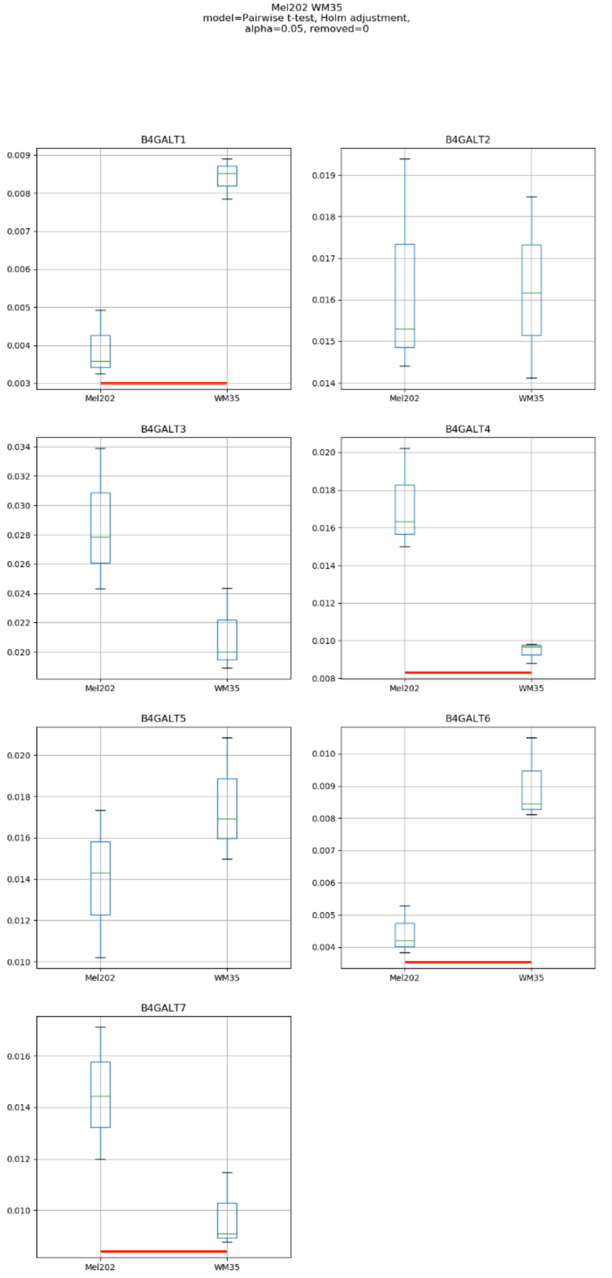

Mel202 WM35
model=Pairwise t-test, Holm adjustment,
alpha=0.05, removed=0
B4GALT1
B4GALT2
B4GALT3
B4GALT4
B4GALT5
B4GALT6
B4GALT7
Mel202
WM35

### Model No. 6

Mel202 WM793
model=Pairwise t-test, Holm adjustment,
alpha=0.05, removed=0

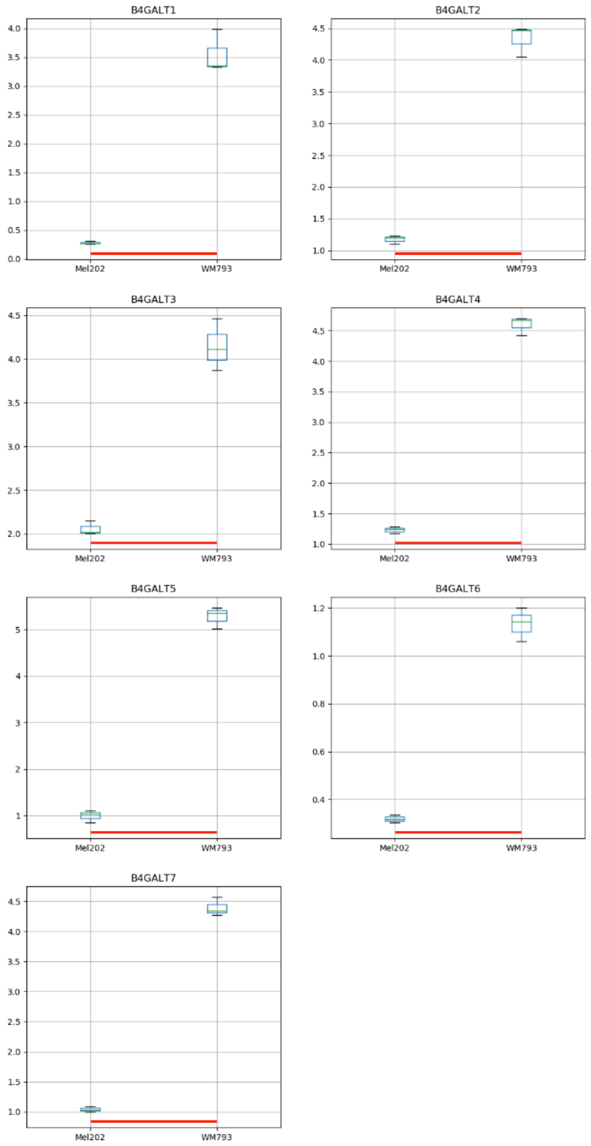

Model No. 7

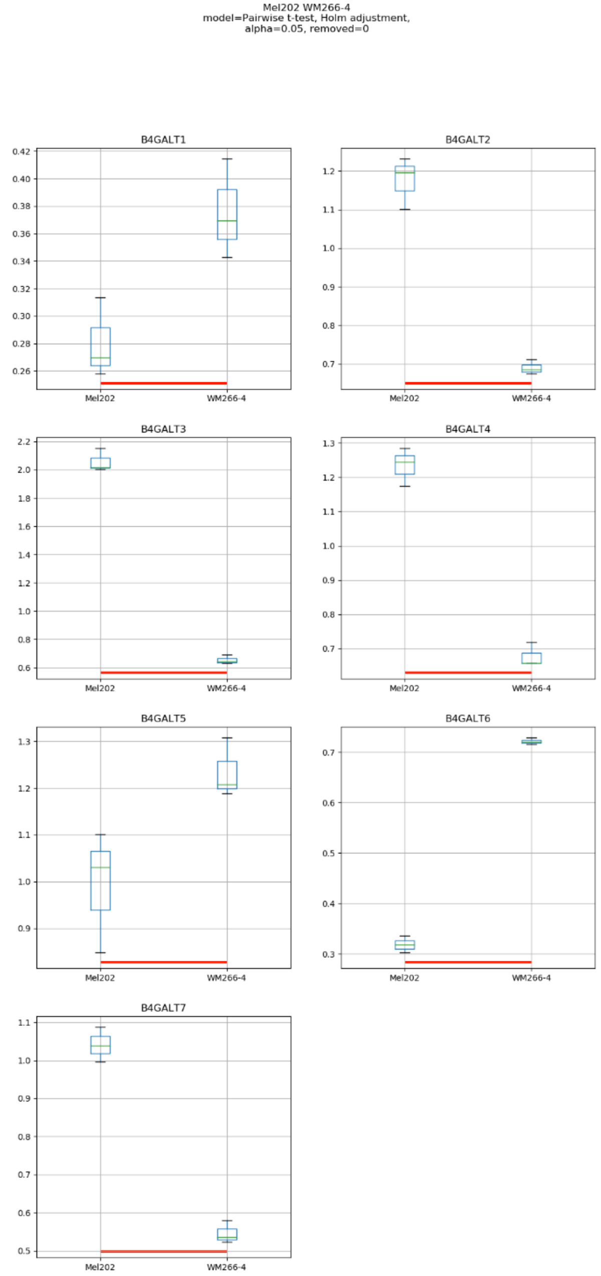
Mel202 WM266-4
model=Pairwise t-test, Holm adjustment,
alpha=0.05, removed=0
B4GALT1
B4GALT2
B4GALT3
B4GALT4
B4GALT5
B4GALT6
B4GALT7
Mel202
WM266-4

Model No. 8

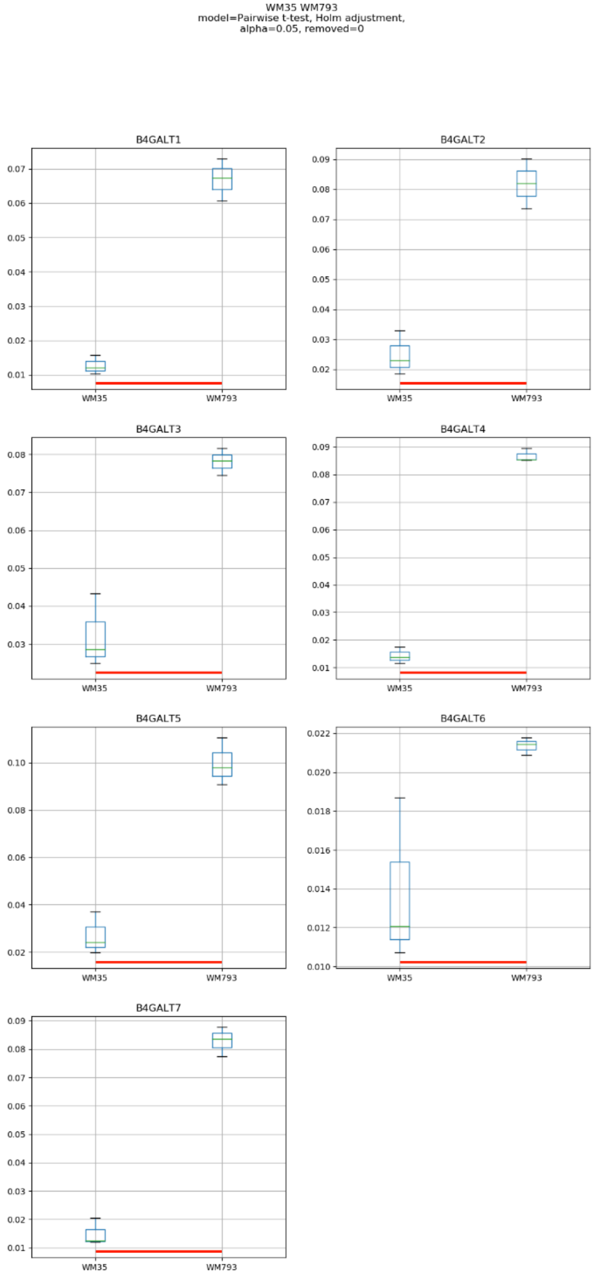
WM35 WM793
model=Pairwise t-test, Holm adjustment,
alpha=0.05, removed=0
B4GALT1
B4GALT2
B4GALT3
B4GALT4
B4GALT5
B4GALT6
B4GALT7
WM35
WM793

Model No. 9

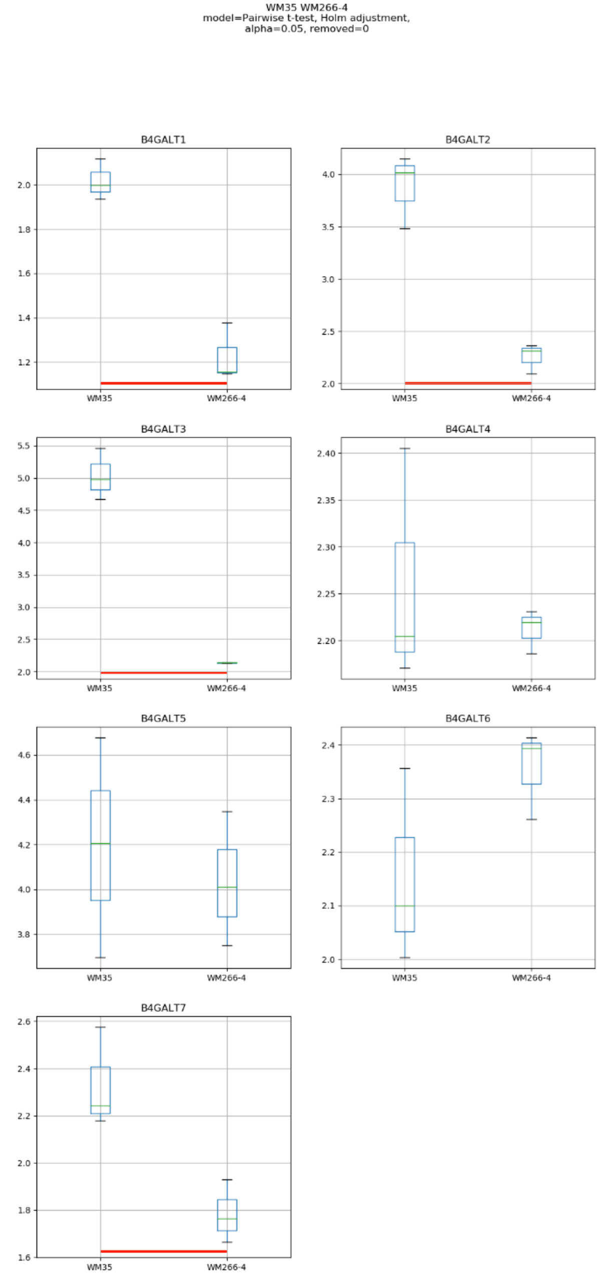
WM35 WM266-4
model=Pairwise t-test, Holm adjustment,
alpha=0.05, removed=0
B4GALT1
B4GALT2
B4GALT3
B4GALT4
B4GALT5
B4GALT6
B4GALT7
WM35
WM266-4

### Model No. 10

WM793 WM266-4
model=Pairwise t-test, Holm adjustment,
alpha=0.05, removed=0

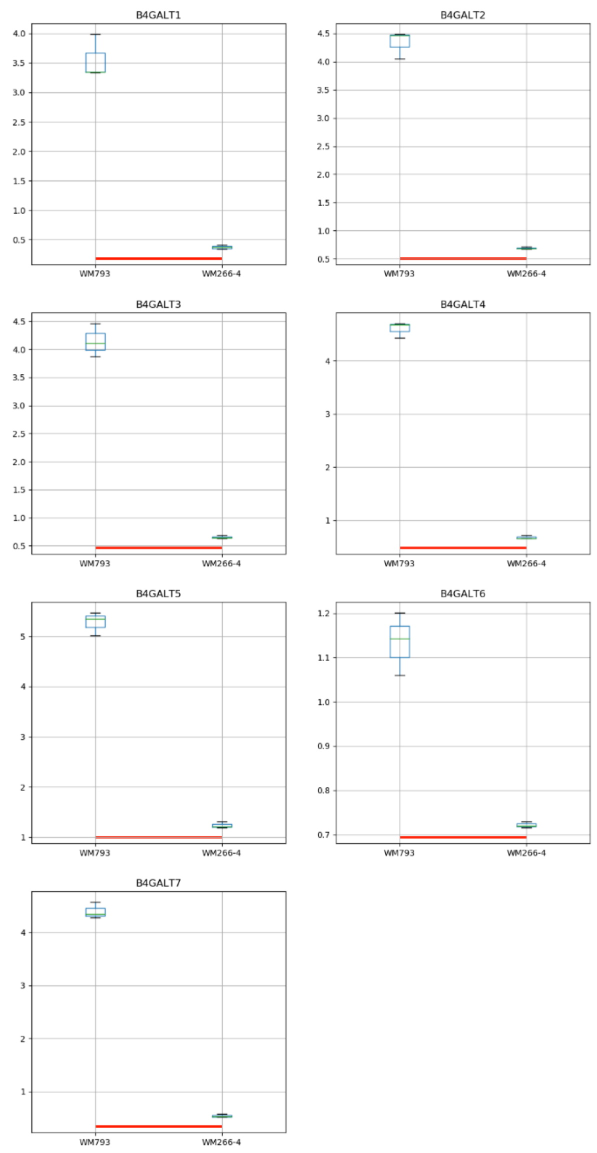

### Model No. 11

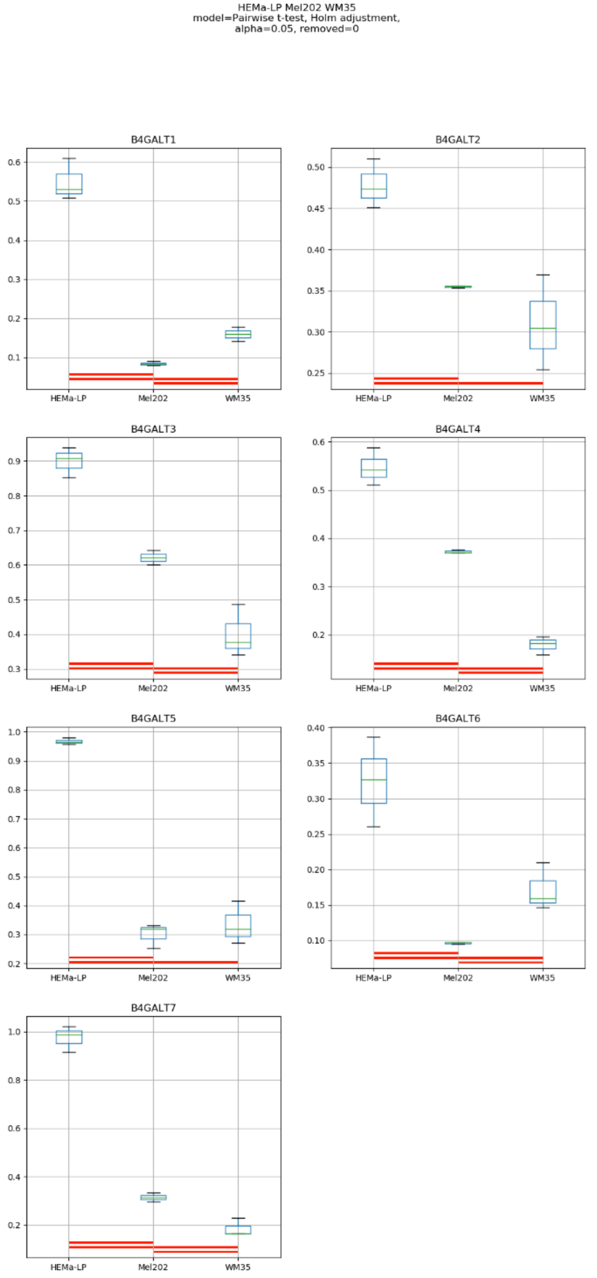
HEMa-LP Mel202 WM35
model=Pairwise t-test, Holm adjustment,
alpha=0.05, removed=0
B4GALT1
B4GALT2
B4GALT3
B4GALT4
B4GALT5
B4GALT6
B4GALT7
HEMa-LP
Mel202
WM35

Model No. 12

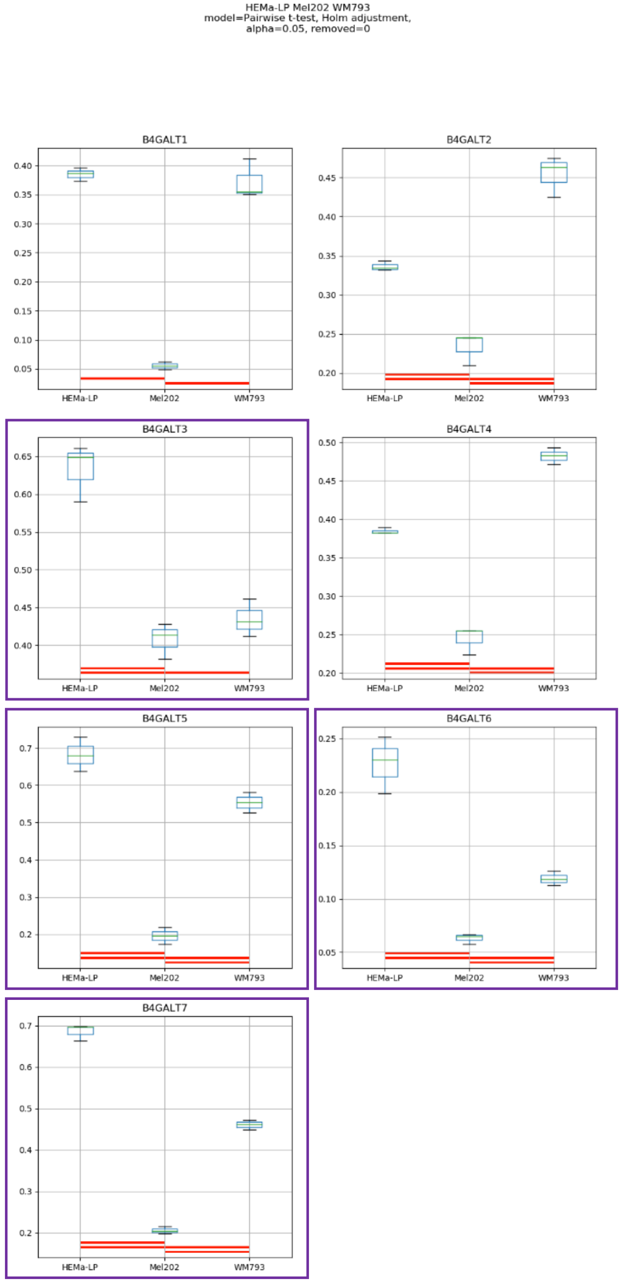
HEMa-LP Mel202 WM793
model=Pairwise t-test, Holm adjustment,
alpha=0.05, removed=0
B4GALT1
B4GALT2
B4GALT3
B4GALT4
B4GALT5
B4GALT6
B4GALT7
HEMa-LP
Mel202
WM793

### Model No. 13

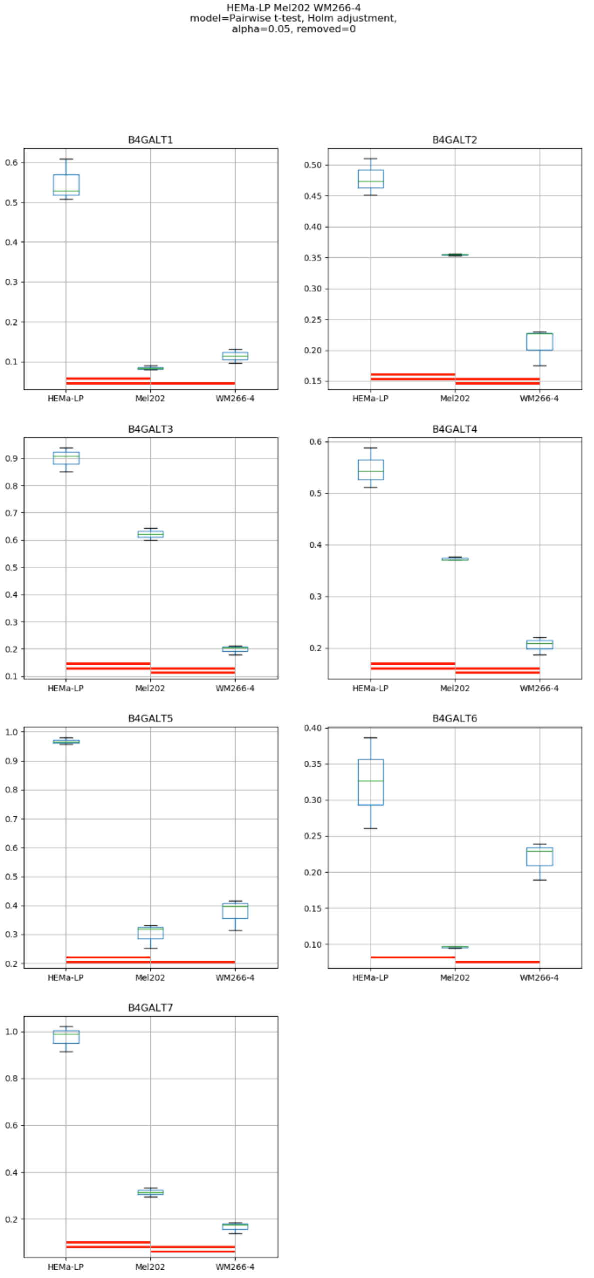

HEMa-LP Mel202 WM266-4
model=Pairwise t-test, Holm adjustment,
alpha=0.05, removed=0
B4GALT1
B4GALT2
B4GALT3
B4GALT4
B4GALT5
B4GALT6
B4GALT7
HEMa-LP
Mel202
WM266-4

Model No. 14

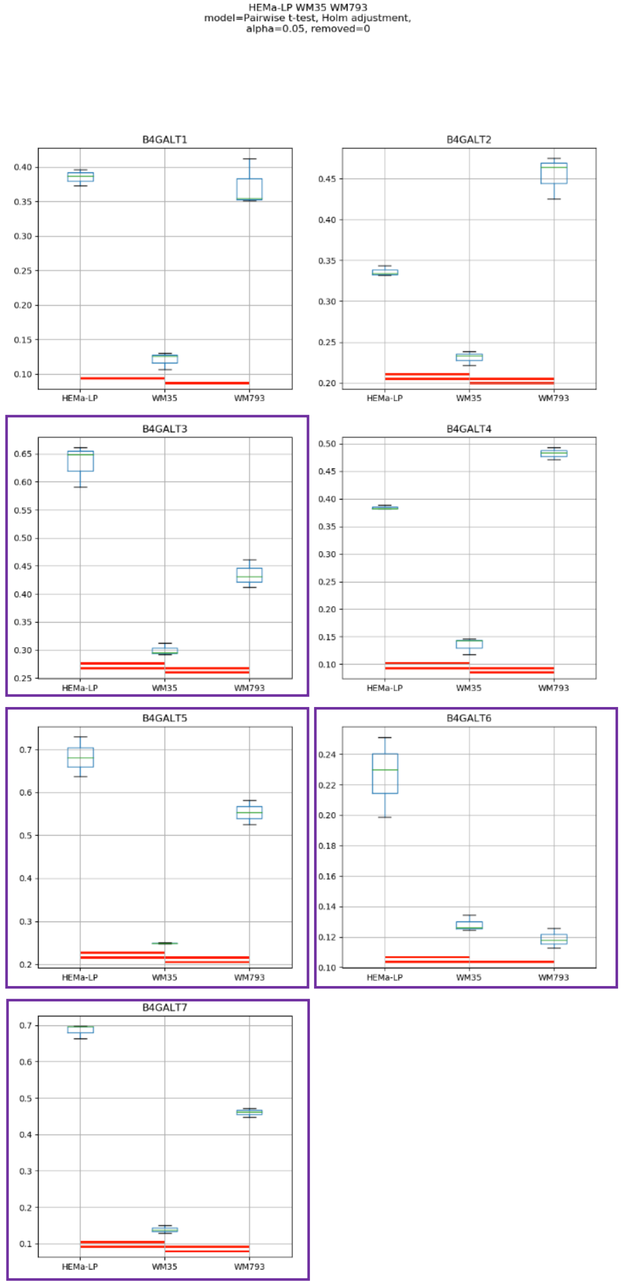
HEMa-LP WM35 WM793
model=Pairwise t-test, Holm adjustment,
alpha=0.05, removed=0
B4GALT1
B4GALT2
B4GALT3
B4GALT4
B4GALT5
B4GALT6
B4GALT7
HEMa-LP
WM35
WM793

Model No. 15

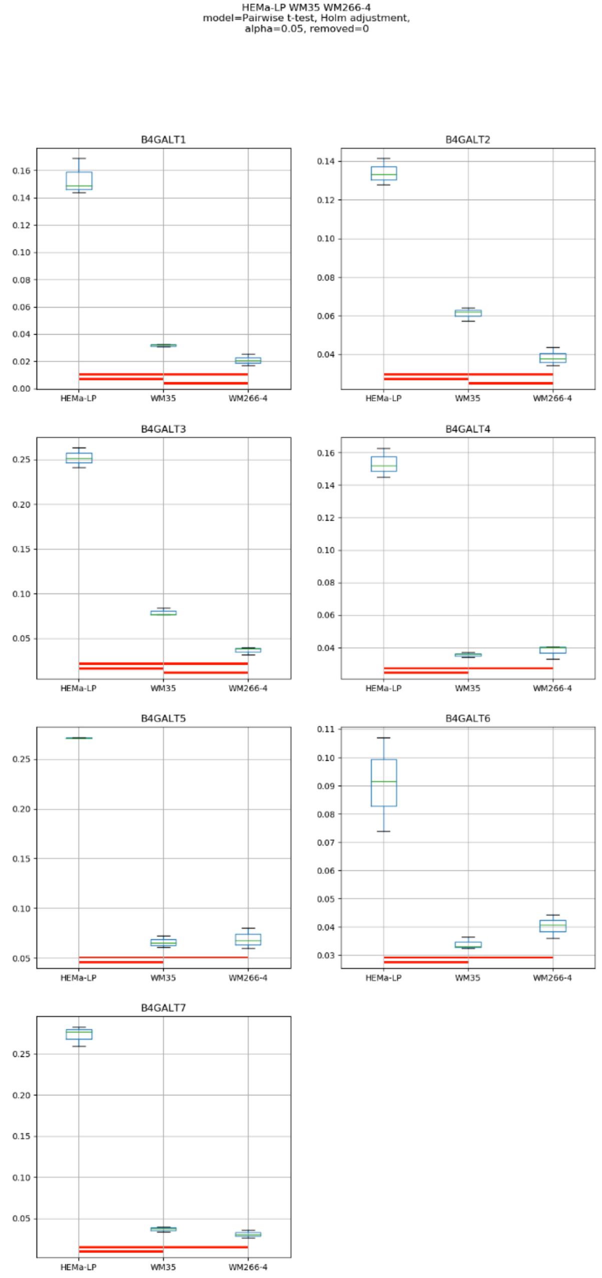
HEMa-LP WM35 WM266-4
model=Pairwise t-test, Holm adjustment,
alpha=0.05, removed=0
B4GALT1
B4GALT2
B4GALT3
B4GALT4
B4GALT5
B4GALT6
B4GALT7
HEMa-LP
WM35
WM266-4

### Model No. 16

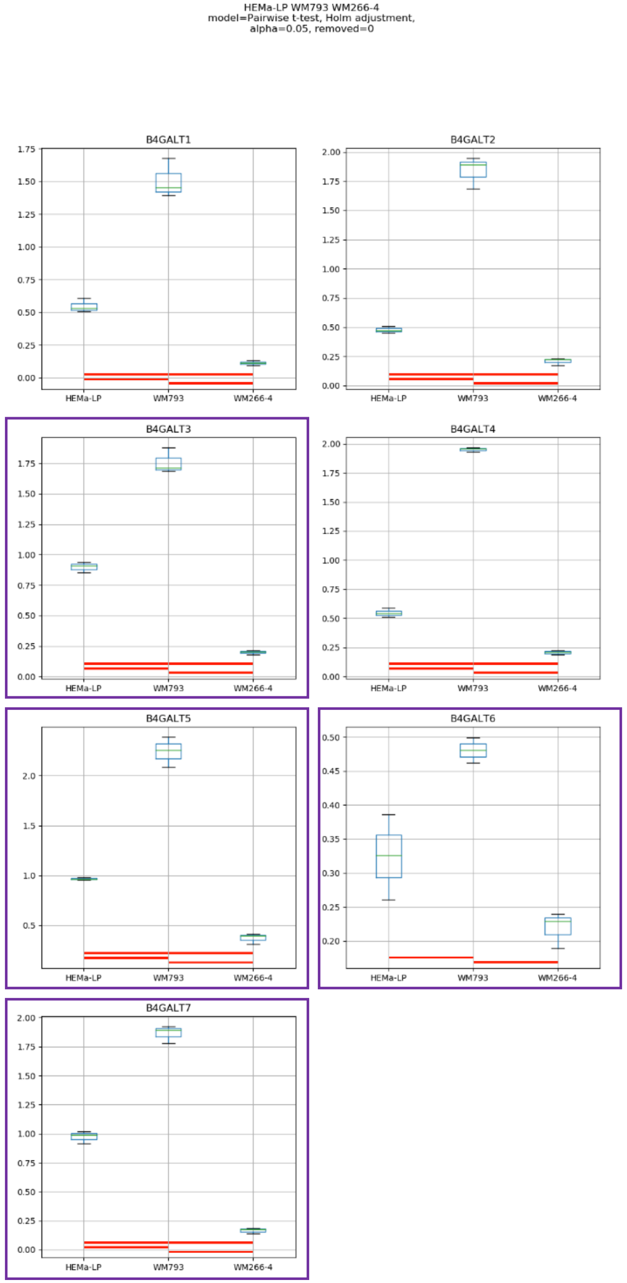
HEMa-LP WM793 WM266-4
model=Pairwise t-test, Holm adjustment,
alpha=0.05, removed=0
B4GALT1
B4GALT2
B4GALT3
B4GALT4
B4GALT5
B4GALT6
B4GALT7
HEMa-LP
WM793
WM266-4

### Model No. 17

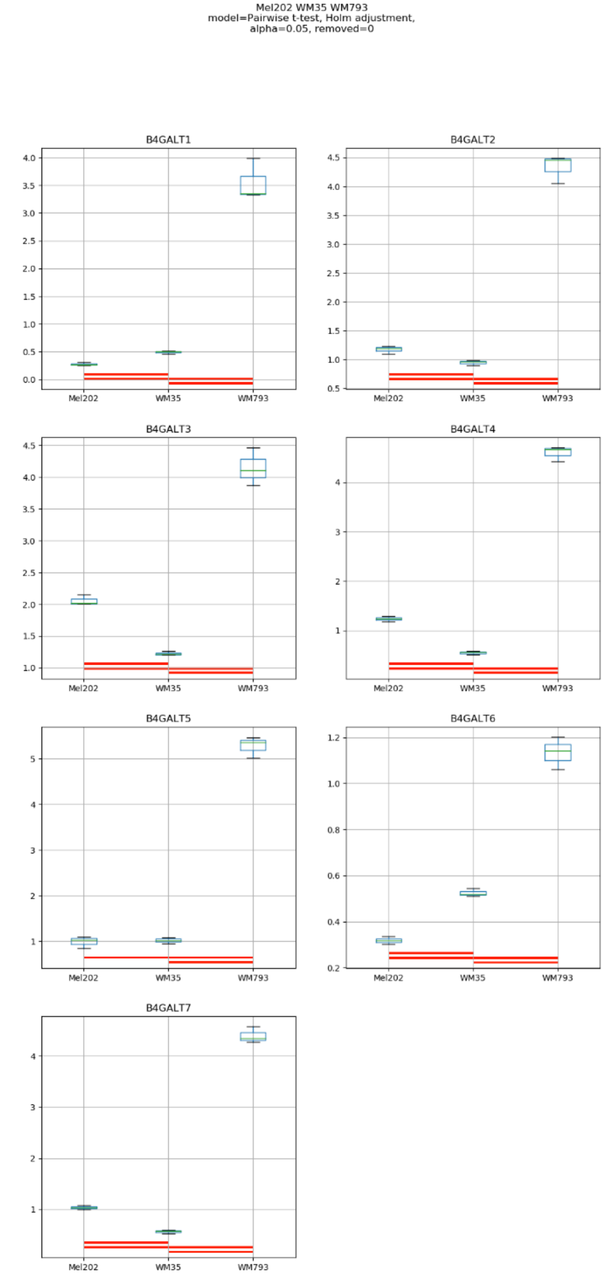
Mel202 WM35 WM793
model=Pairwise t-test, Holm adjustment,
alpha=0.05, removed=0
B4GALT1
B4GALT2
B4GALT3
B4GALT4
B4GALT5
B4GALT6
B4GALT7
Mel202
WM35
WM793

#### Model No. 18

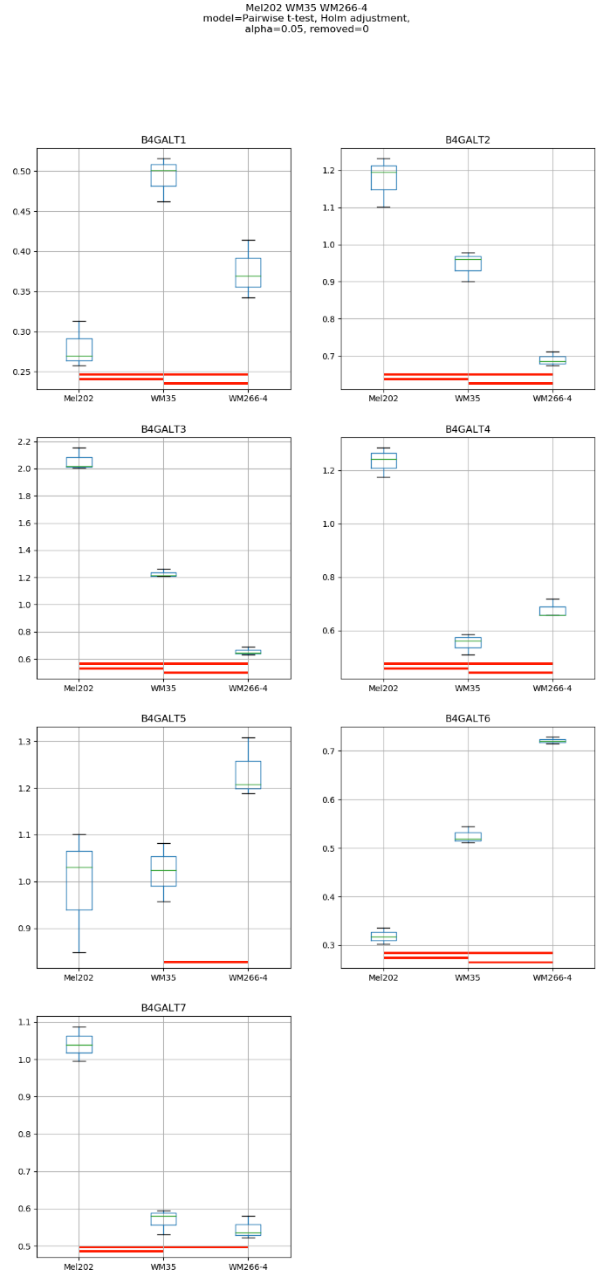
Mel202 WM35 WM266-4
model=Pairwise t-test, Holm adjustment,
alpha=0.05, removed=0
B4GALT1
B4GALT2
B4GALT3
B4GALT4
B4GALT5
B4GALT6
B4GALT7
Mel202
WM35
WM266-4

### Model No. 19

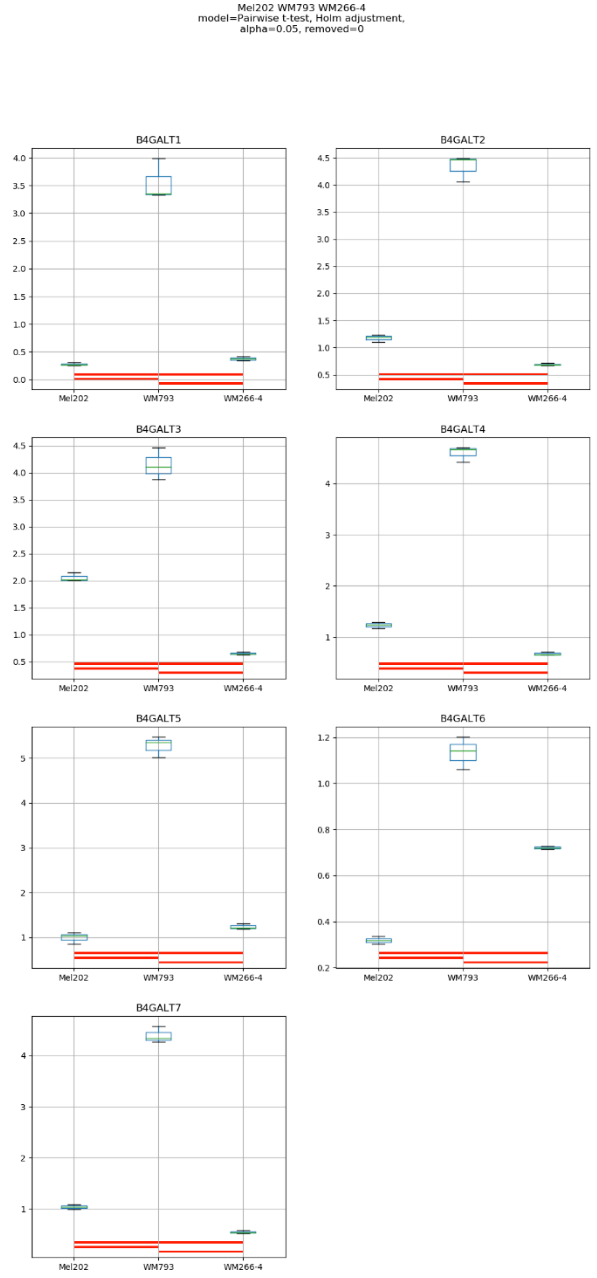
Mel202 WM793 WM266-4
model=Pairwise t-test, Holm adjustment,
alpha=0.05, removed=0
B4GALT1
B4GALT2
B4GALT3
B4GALT4
B4GALT5
B4GALT6
B4GALT7
Mel202
WM793
WM266-4

### Model No. 20

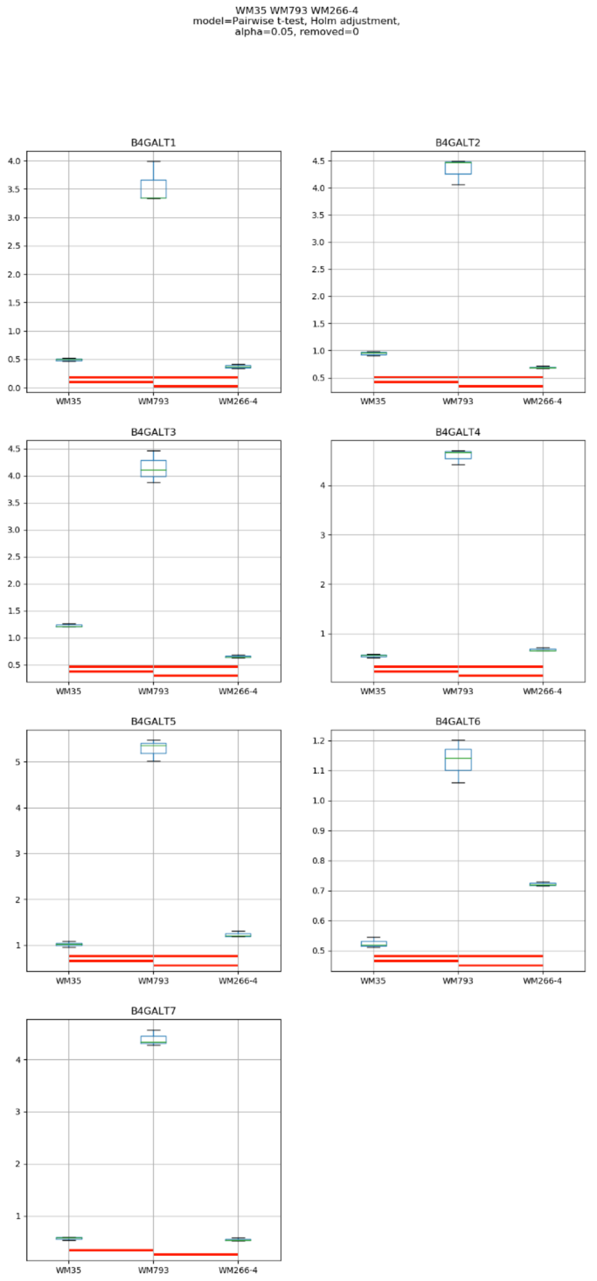
WM35 WM793 WM266-4
model=Pairwise t-test, Holm adjustment,
alpha=0.05, removed=0
B4GALT1
B4GALT2
B4GALT3
B4GALT4
B4GALT5
B4GALT6
B4GALT7
WM35
WM793
WM266-4

### Model No. 21

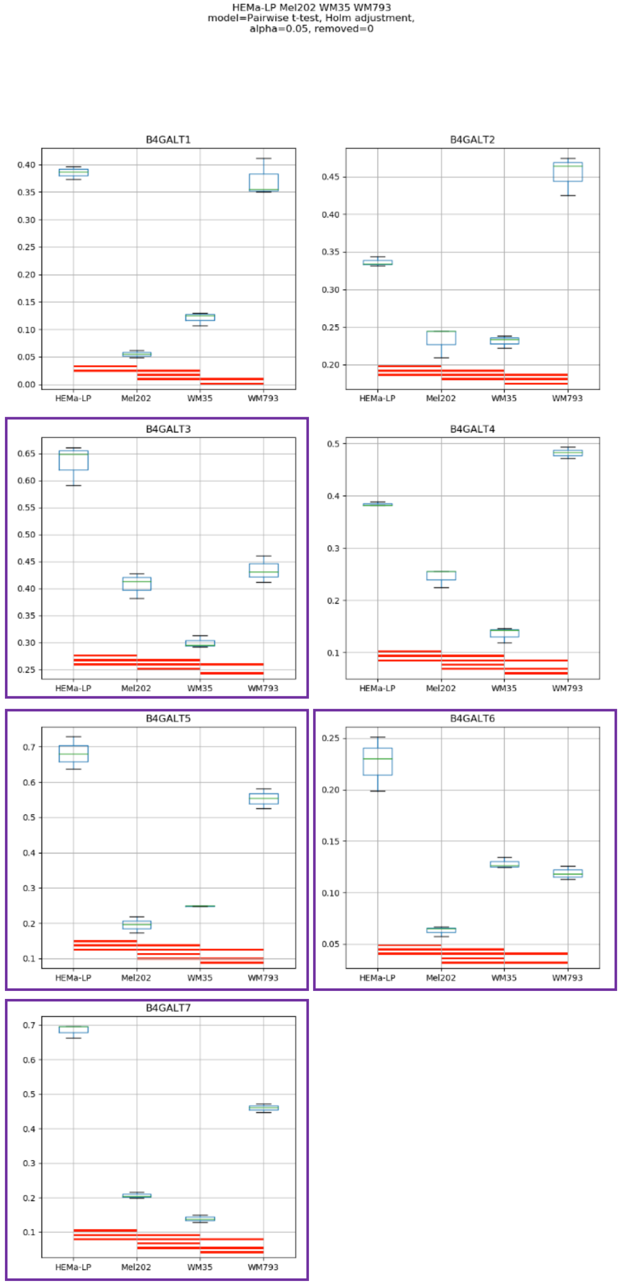
HEMa-LP Mel202 WM35 WM793
model=Pairwise t-test, Holm adjustment,
alpha=0.05, removed=0
B4GALT1
B4GALT2
B4GALT3
B4GALT4
B4GALT5
B4GALT6
B4GALT7
HEMa-LP
Mel202
WM35
WM793

### Model No. 22

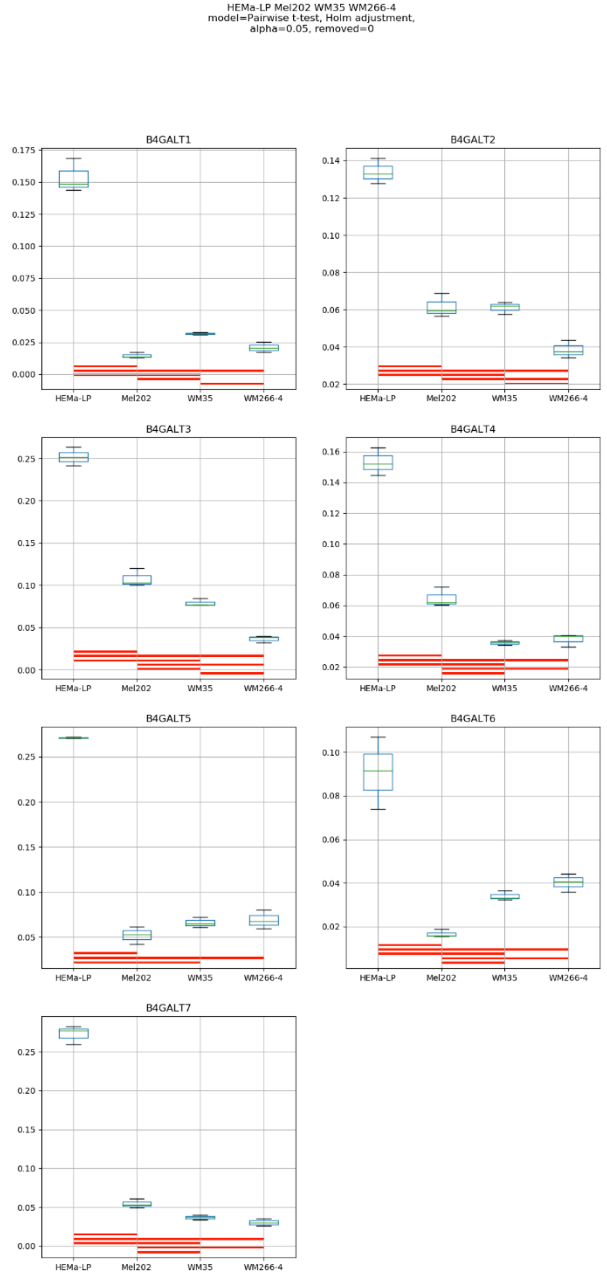
HEMa-LP Mel202 WM35 WM266-4
model=Pairwise t-test, Holm adjustment,
alpha=0.05, removed=0
B4GALT1
B4GALT2
B4GALT3
B4GALT4
B4GALT5
B4GALT6
B4GALT7
HEMa-LP
Mel202
WM35
WM266-4

### Model No. 23

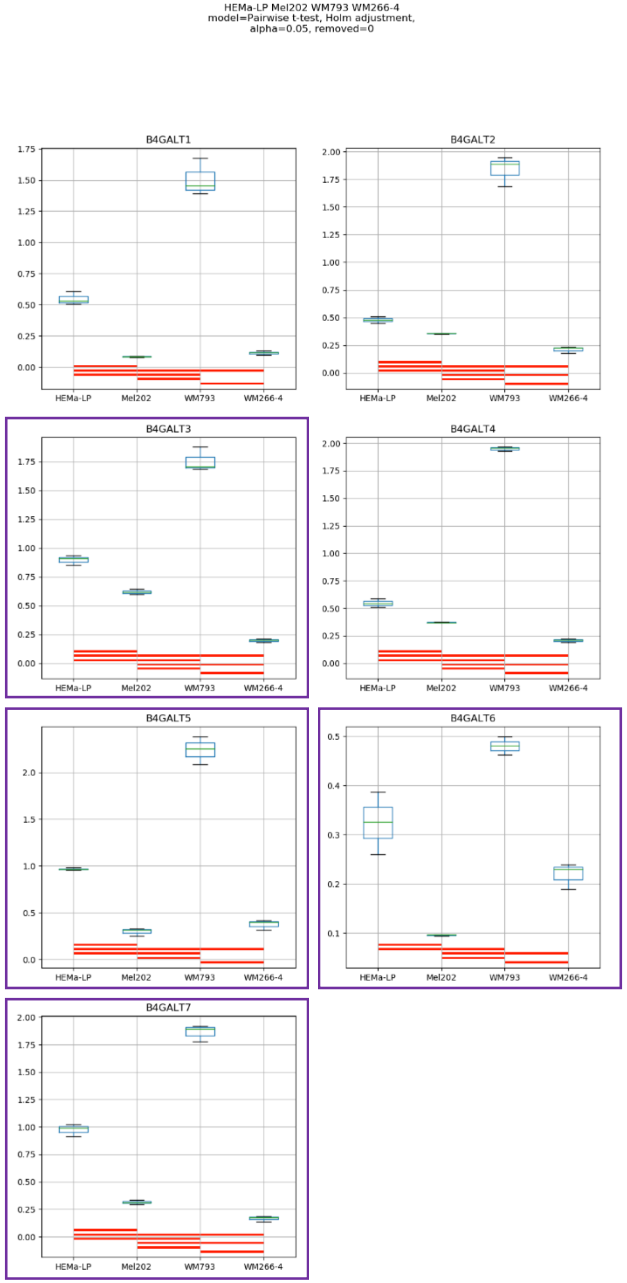
HEMa-LP Mel202 WM793 WM266-4
model=Pairwise t-test, Holm adjustment,
alpha=0.05, removed=0
B4GALT1
B4GALT2
B4GALT3
B4GALT4
B4GALT5
B4GALT6
B4GALT7
HEMa-LP
Mel202
WM793
WM266-4

### Model No. 24

HEMa-LP WM35 WM793 WM266-4
model=Pairwise t-test, Holm adjustment,
alpha=0.05, removed=0

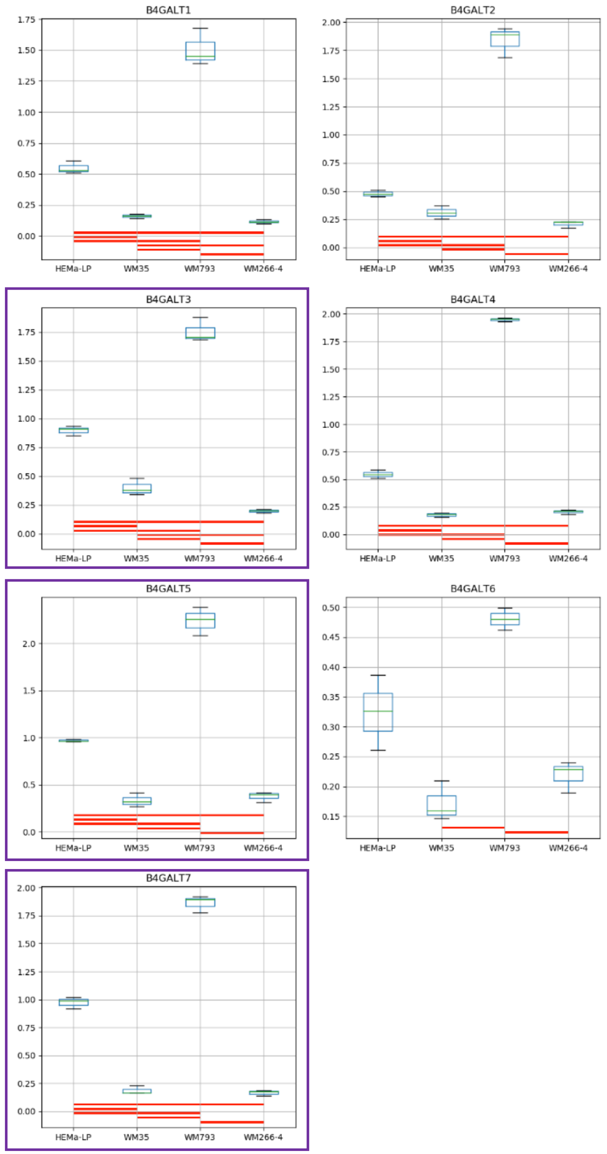

Model No. 25

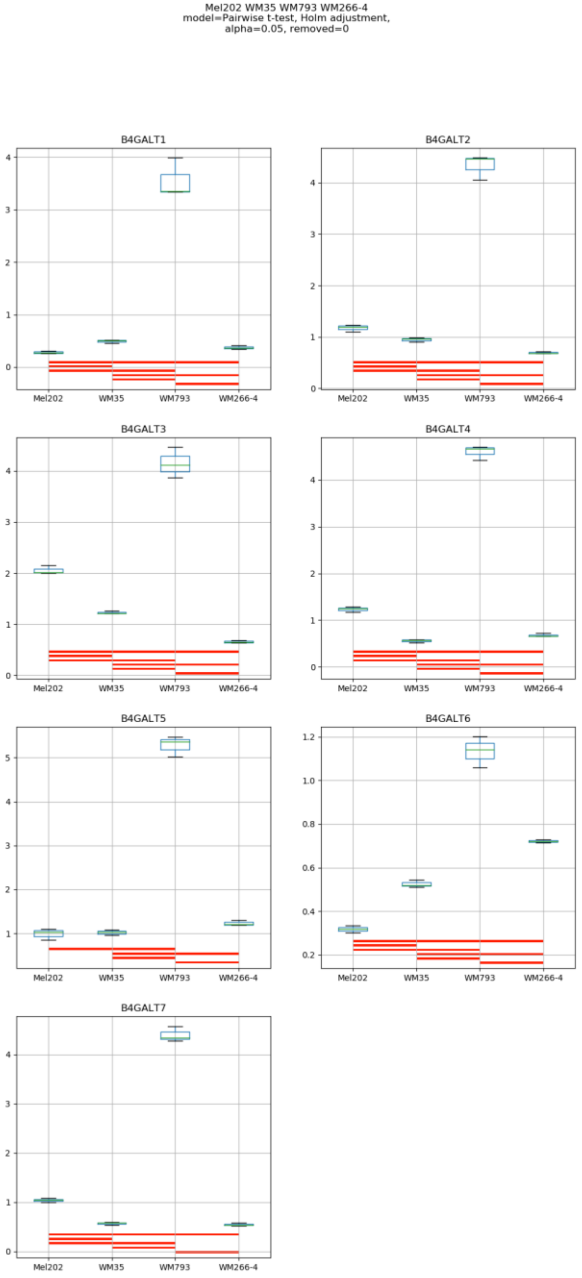
Mel202 WM35 WM793 WM266-4
model=Pairwise t-test, Holm adjustment,
alpha=0.05, removed=0
B4GALT1
B4GALT2
B4GALT3
B4GALT4
B4GALT5
B4GALT6
B4GALT7
Mel202
WM35
WM793
WM266-4

### Model No. 26

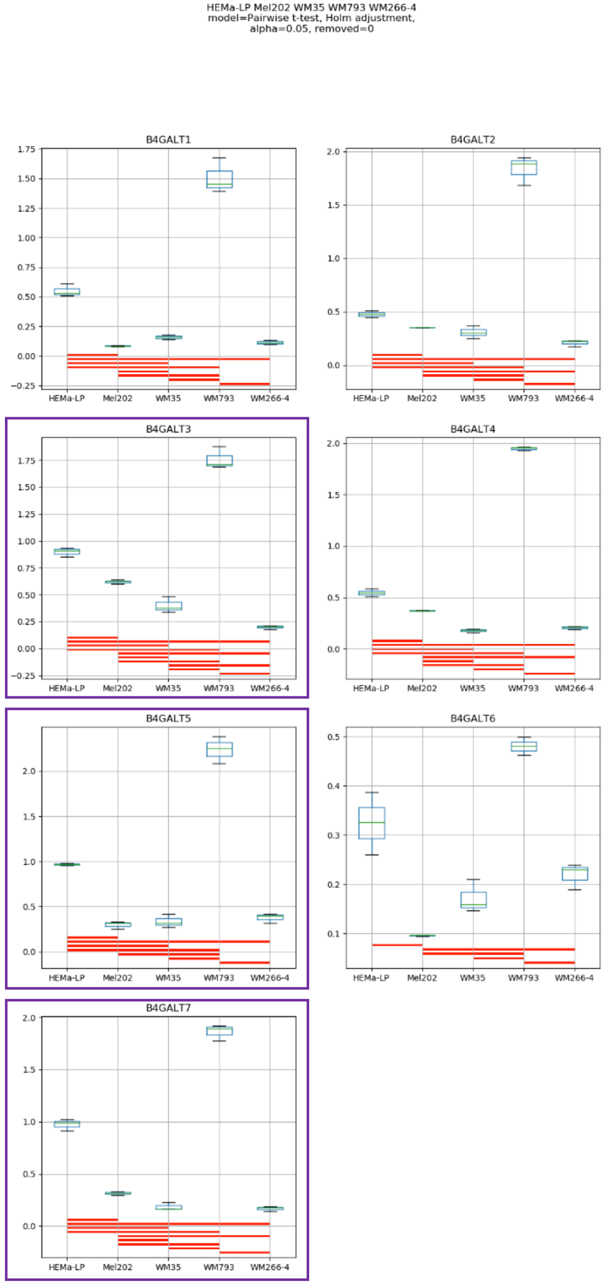
HEMa-LP Mel202 WM35 WM793 WM266-4
model=Pairwise t-test, Holm adjustment,
alpha=0.05, removed=0
B4GALT1
B4GALT2
B4GALT3
B4GALT4
B4GALT5
B4GALT6
B4GALT7
HEMa-LP
Mel202
WM35
WM793
WM266-4

**Supplementary Figure S1:** Medians of relative quantity (RQ) for target genes *B4GALT1–B4GALT7* normalized to the reference gene or pair of reference genes with the best stability values resulting from GenExpA analysis based on set 1 of four candidate reference genes (*HPRT1*, *PGK1*, *RPS23*, *SNRPA*) and remove repetition level 0 (see Table 1, part A). The statistical analyses used the pairwise t-test with Holm adjustment. Red line represents statistical significance, $P<0.05$. Frames denote box-plots representing the medians of the RQ values for a given target gene without a repeatable pattern in the same cell lines under different models (the 'Remove repetitions' box set to 0). Detailed results are listed in Supplementary Table 3.

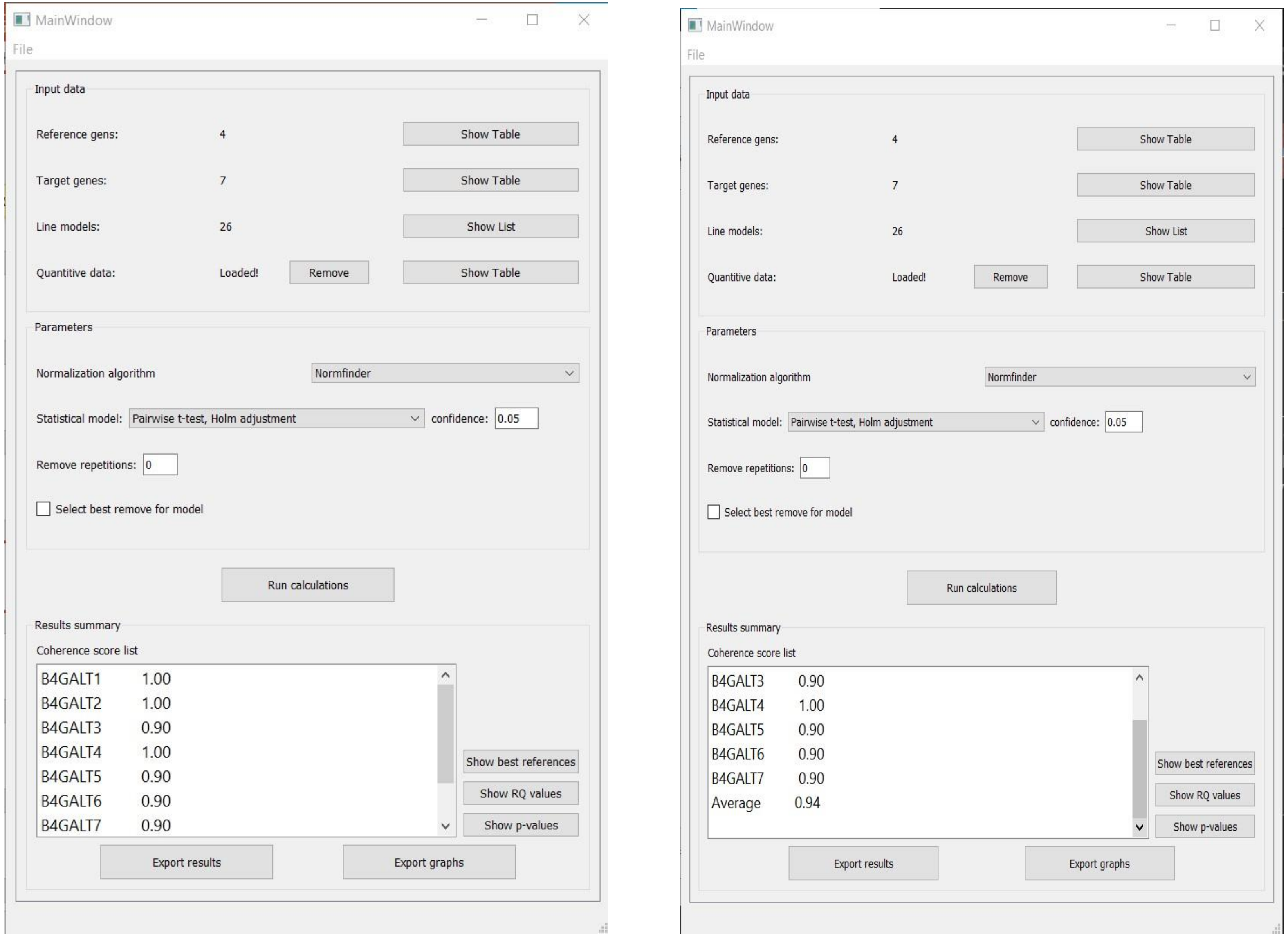


**Supplementary Figure S2:** Print screen of a GenExpA main window with selected parameters of the analysis and coherence score (CS) values for target genes *B4GALT1–B4GALT7* and average CS value of the analysis.

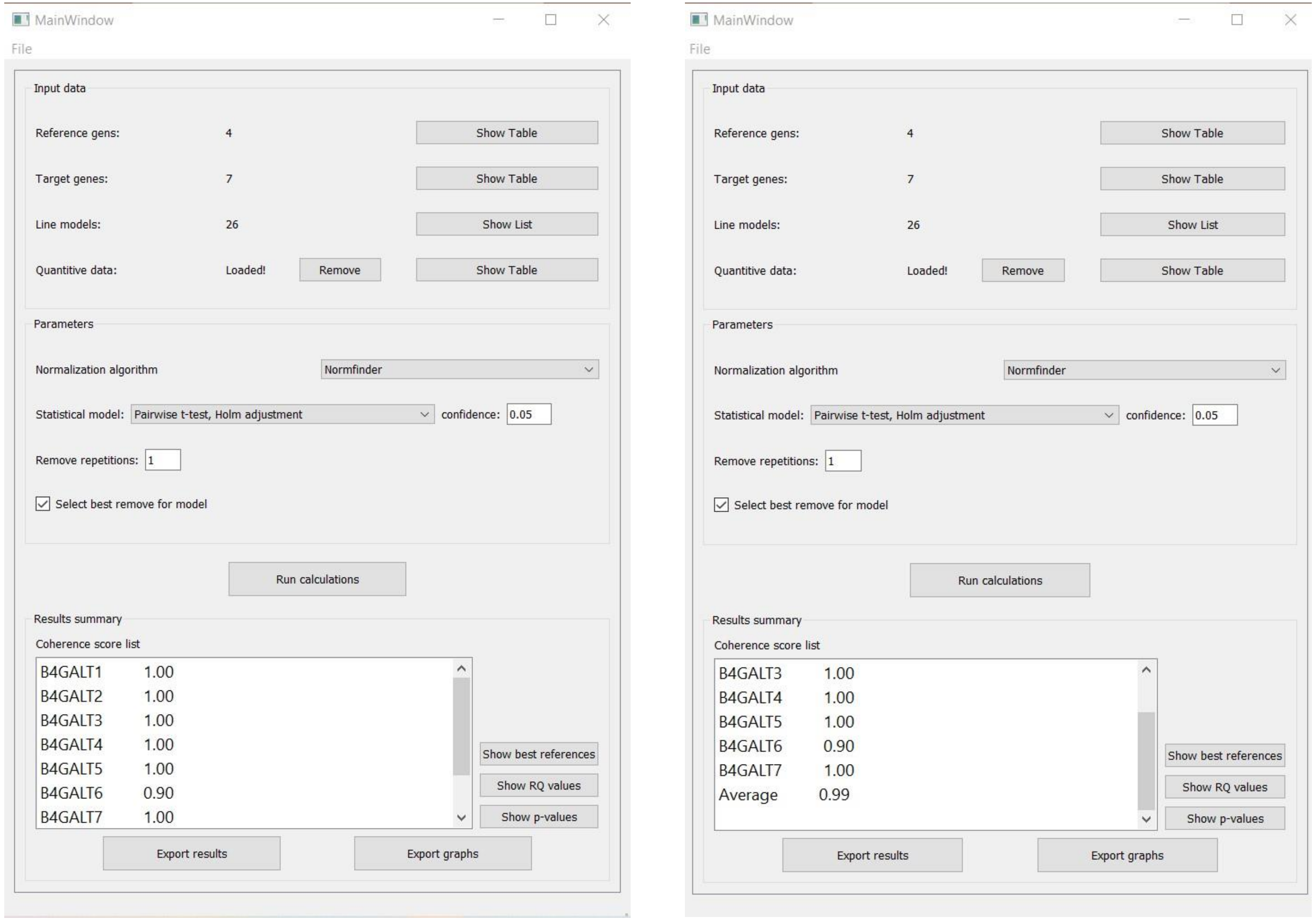


**Supplementary Figure S3:** Print screen of a GenExpA main window with selected parameters of the analysis and coherence score (CS) values for target genes *B4GALT1–B4GALT7* and average CS value of the analysis.

### Model No. 1

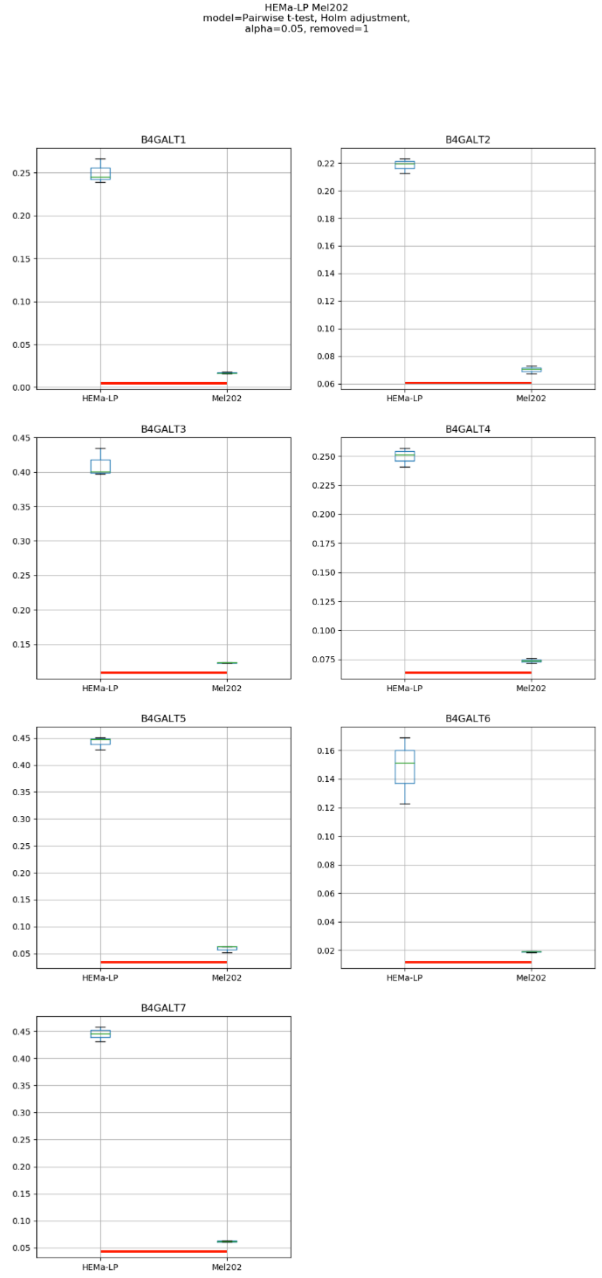
HEMa-LP Mel202
model=Pairwise t-test, Holm adjustment,
alpha=0.05, removed=1
B4GALT1
B4GALT2
B4GALT3
B4GALT4
B4GALT5
B4GALT6
B4GALT7
HEMa-LP
Mel202

### Model No. 2

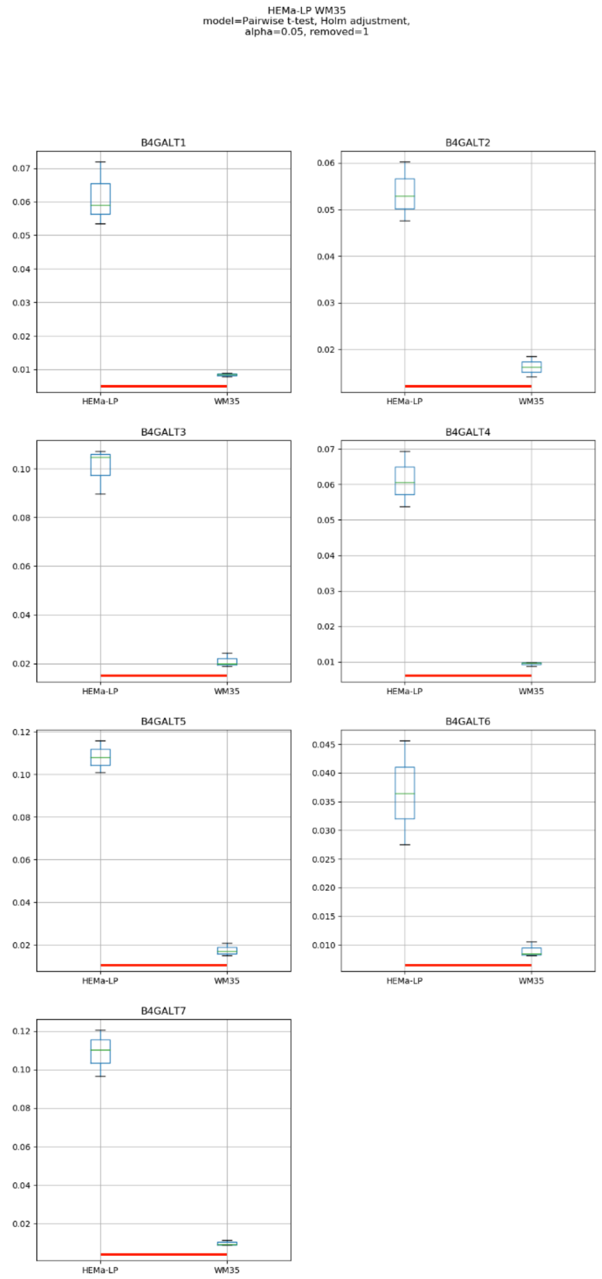
HEMa-LP WM35
model=Pairwise t-test, Holm adjustment,
alpha=0.05, removed=1
B4GALT1
B4GALT2
B4GALT3
B4GALT4
B4GALT5
B4GALT6
B4GALT7
HEMa-LP
WM35

### Model No. 3

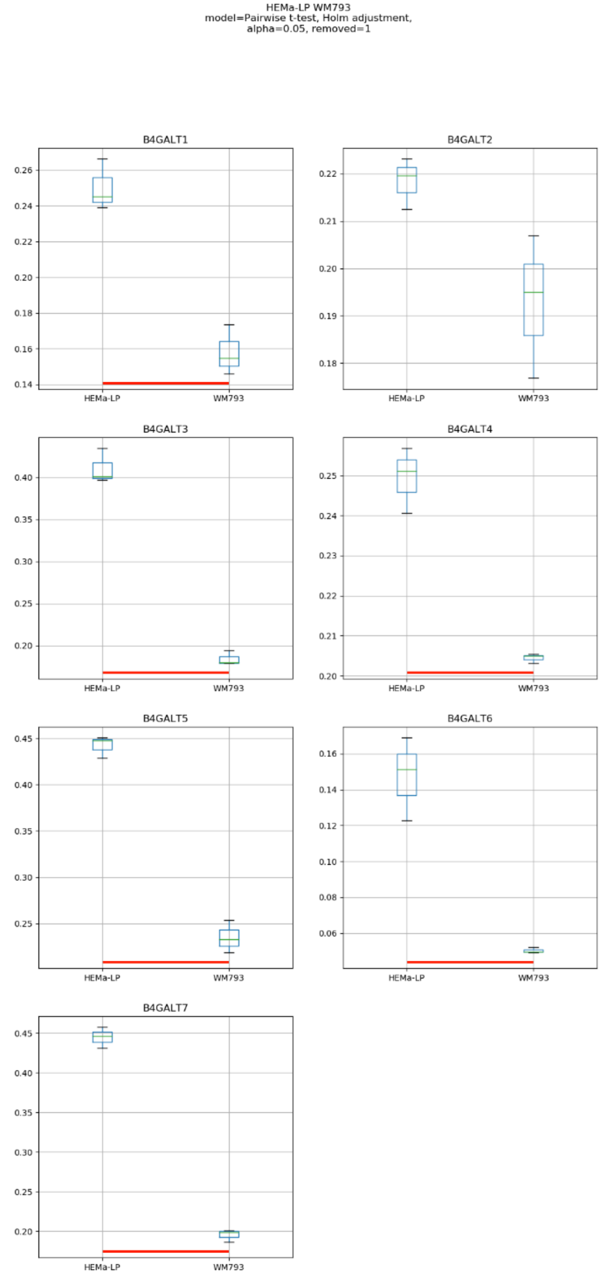
HEMa-LP WM793
model=Pairwise t-test, Holm adjustment,
alpha=0.05, removed=1
B4GALT1
B4GALT2
B4GALT3
B4GALT4
B4GALT5
B4GALT6
B4GALT7
HEMa-LP
WM793

### Model No. 4

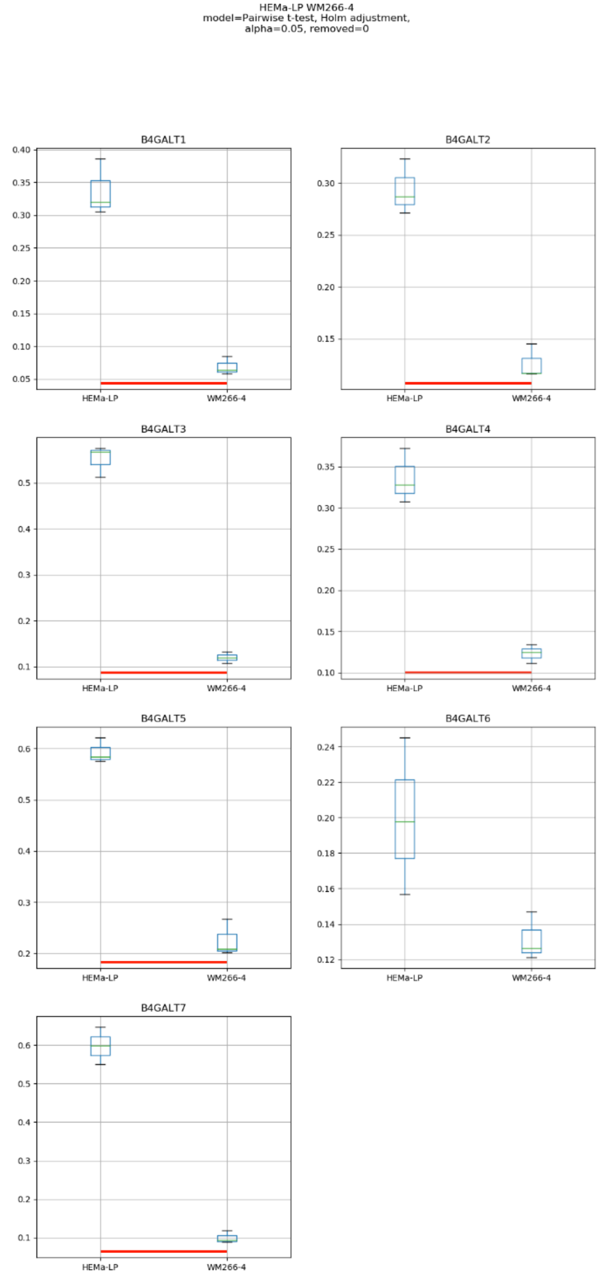
HEMa-LP WM266-4
model=Pairwise t-test, Holm adjustment,
alpha=0.05, removed=0
B4GALT1
B4GALT2
B4GALT3
B4GALT4
B4GALT5
B4GALT6
B4GALT7
HEMa-LP
WM266-4

### Model No. 5

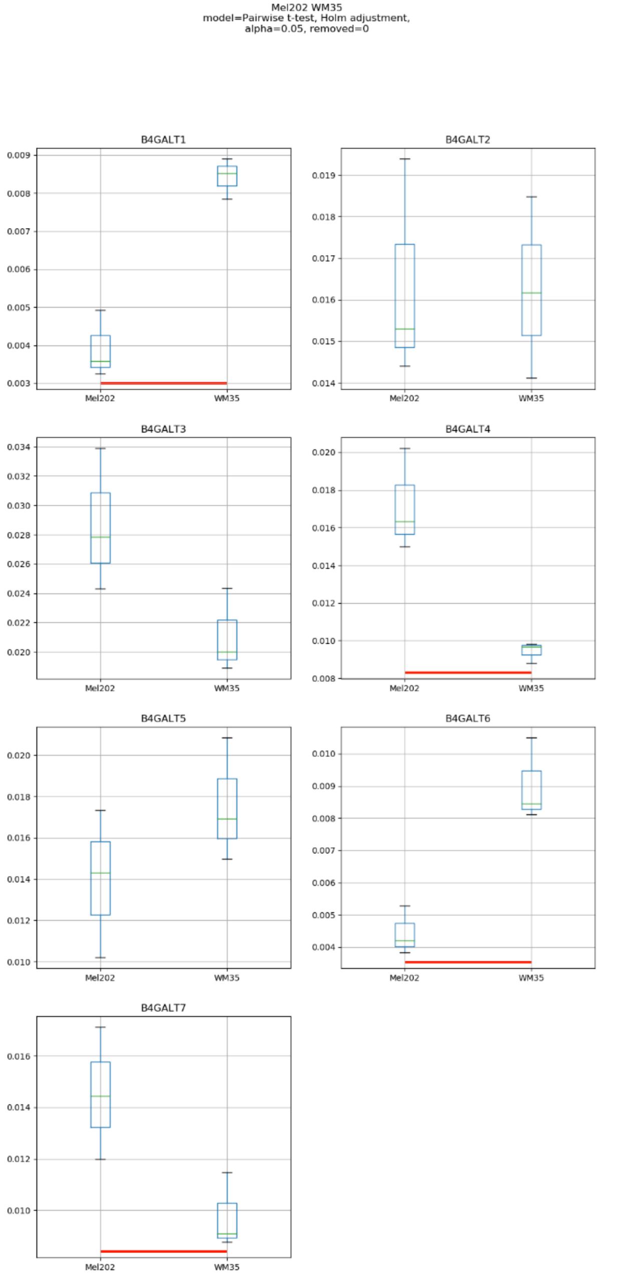

Mel202 WM35
model=Pairwise t-test, Holm adjustment,
alpha=0.05, removed=0
B4GALT1
B4GALT2
B4GALT3
B4GALT4
B4GALT5
B4GALT6
B4GALT7
Mel202
WM35

### Model No. 6

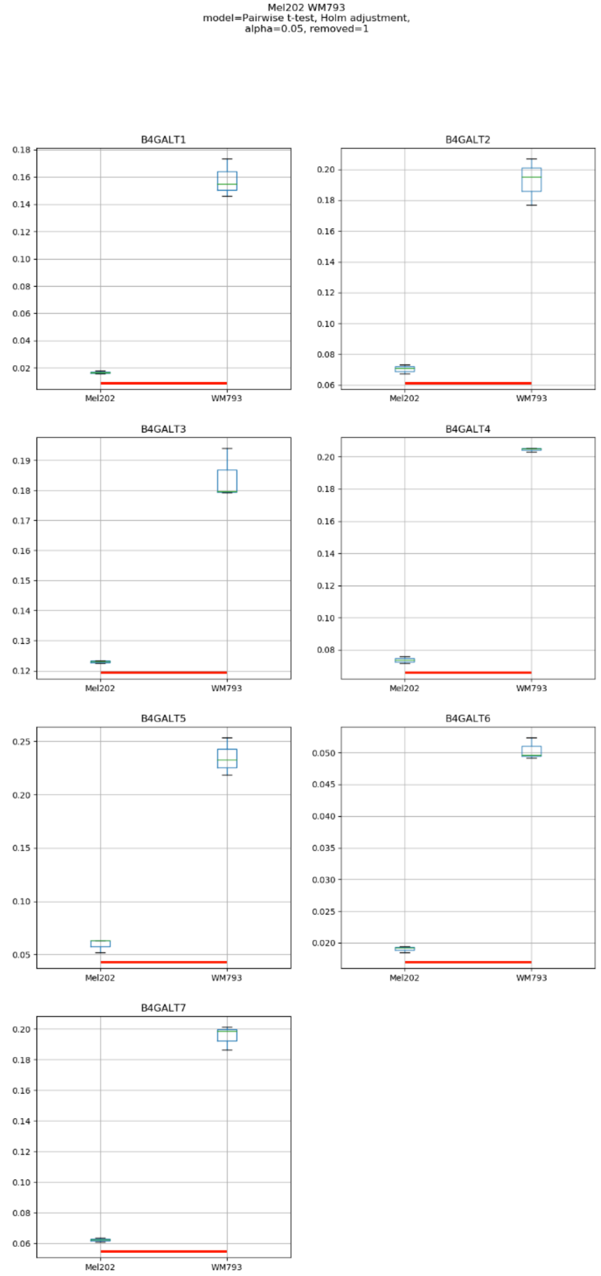
Mel202 WM793
model=Pairwise t-test, Holm adjustment,
alpha=0.05, removed=1
B4GALT1
B4GALT2
B4GALT3
B4GALT4
B4GALT5
B4GALT6
B4GALT7
Mel202
WM793

### Model No. 7

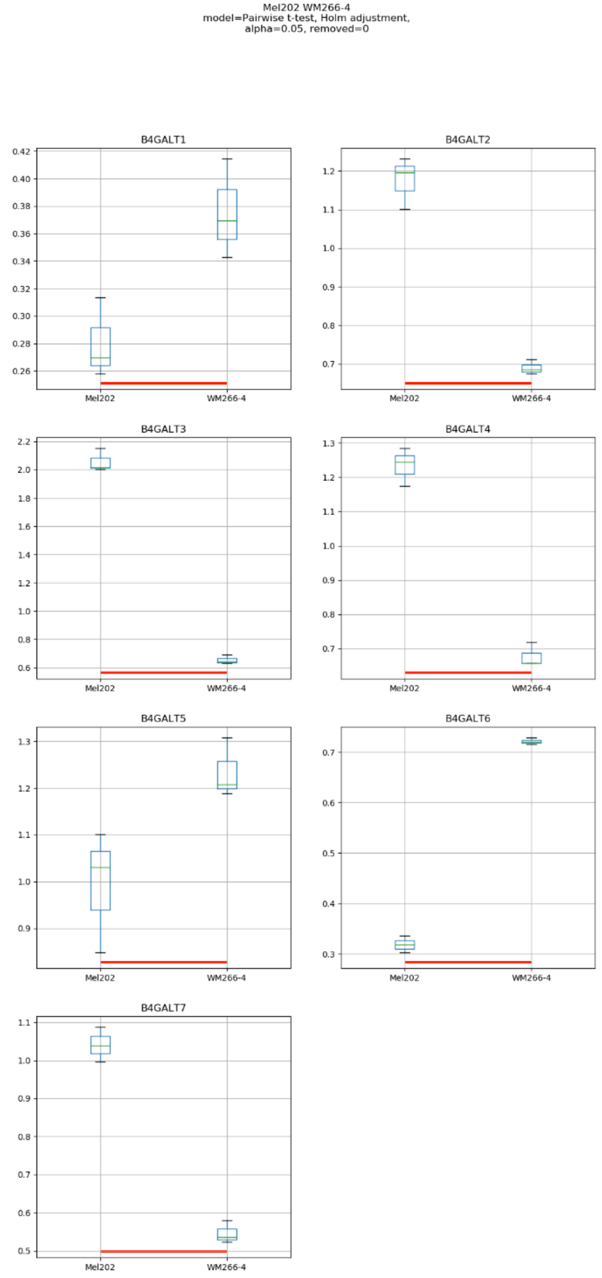
Mel202 WM266-4
model=Pairwise t-test, Holm adjustment,
alpha=0.05, removed=0
B4GALT1
B4GALT2
B4GALT3
B4GALT4
B4GALT5
B4GALT6
B4GALT7
Mel202
WM266-4

### Model No. 8

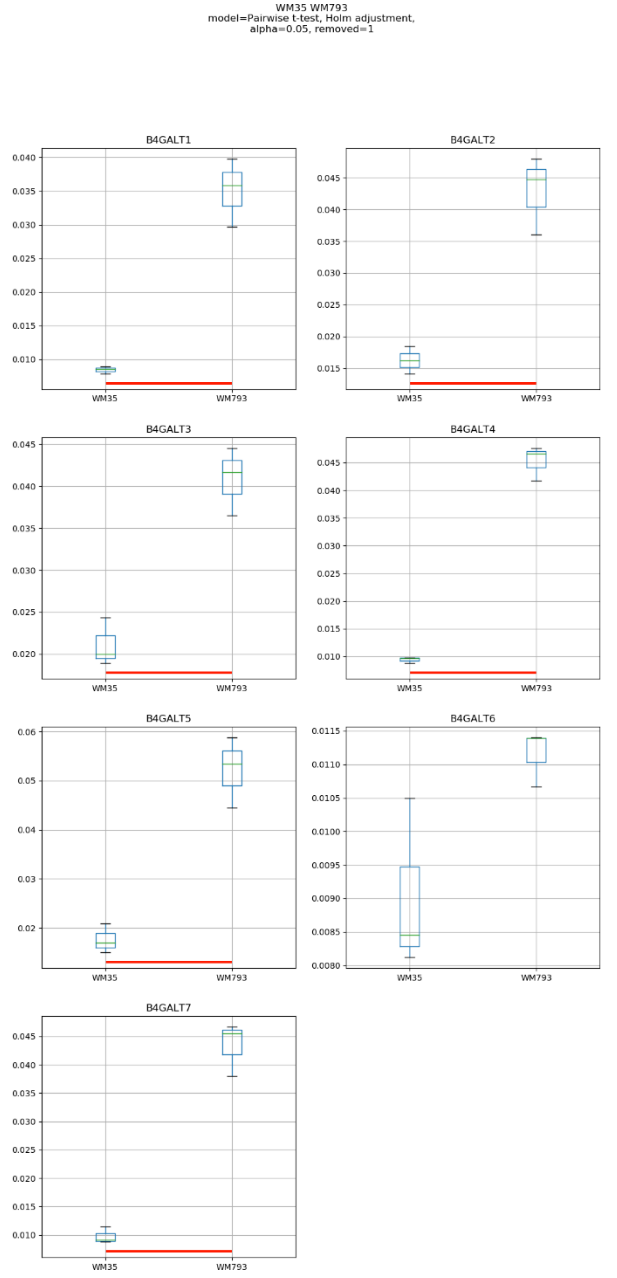
WM35 WM793
model=Pairwise t-test, Holm adjustment,
alpha=0.05, removed=1
B4GALT1
B4GALT2
B4GALT3
B4GALT4
B4GALT5
B4GALT6
B4GALT7
WM35
WM793

### Model No. 9

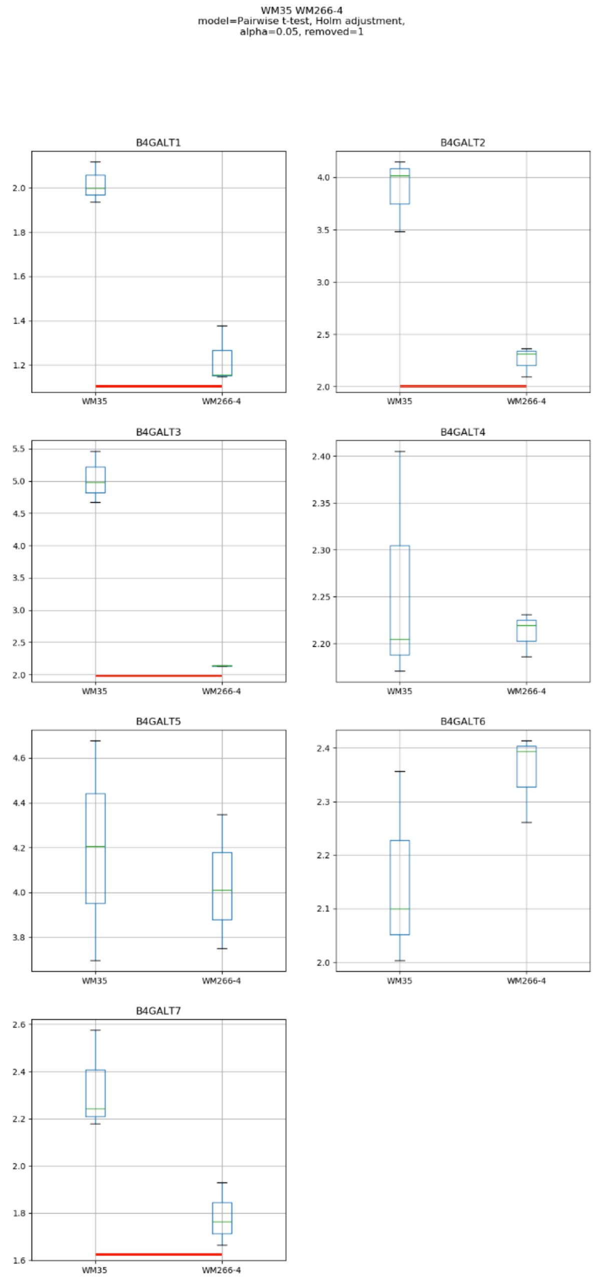
WM35 WM266-4
model=Pairwise t-test, Holm adjustment,
alpha=0.05, removed=1
B4GALT1
B4GALT2
B4GALT3
B4GALT4
B4GALT5
B4GALT6
B4GALT7
WM35
WM266-4

### Model No. 10

WM793 WM266-4
Mann-Whitney/Kruska-Wallis, post: Dunn's test,
alpha=0.05, removed=1

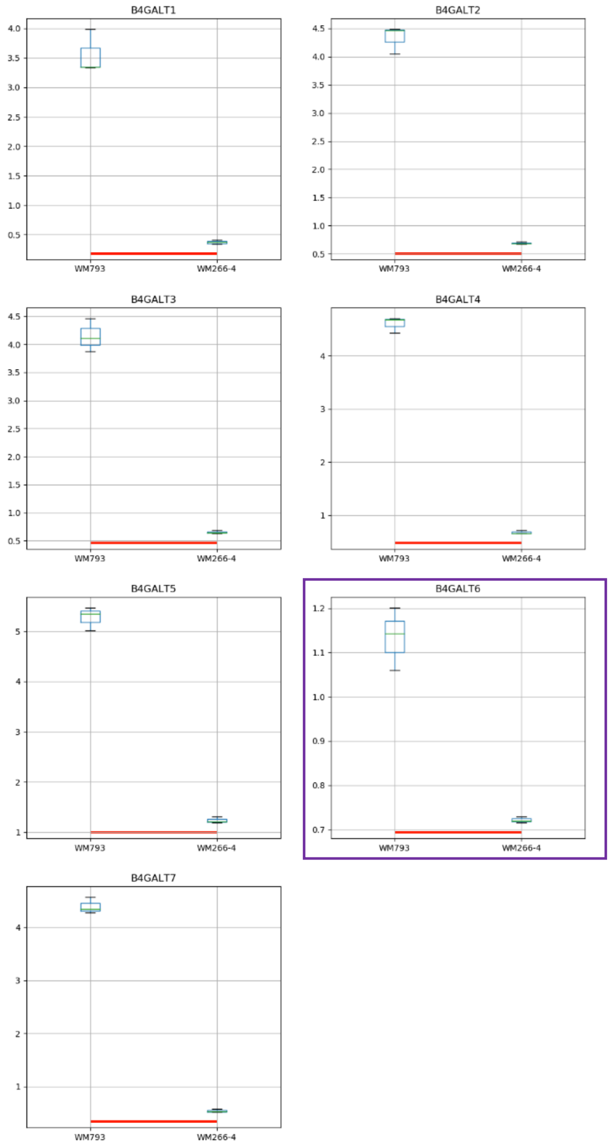

Model No. 11

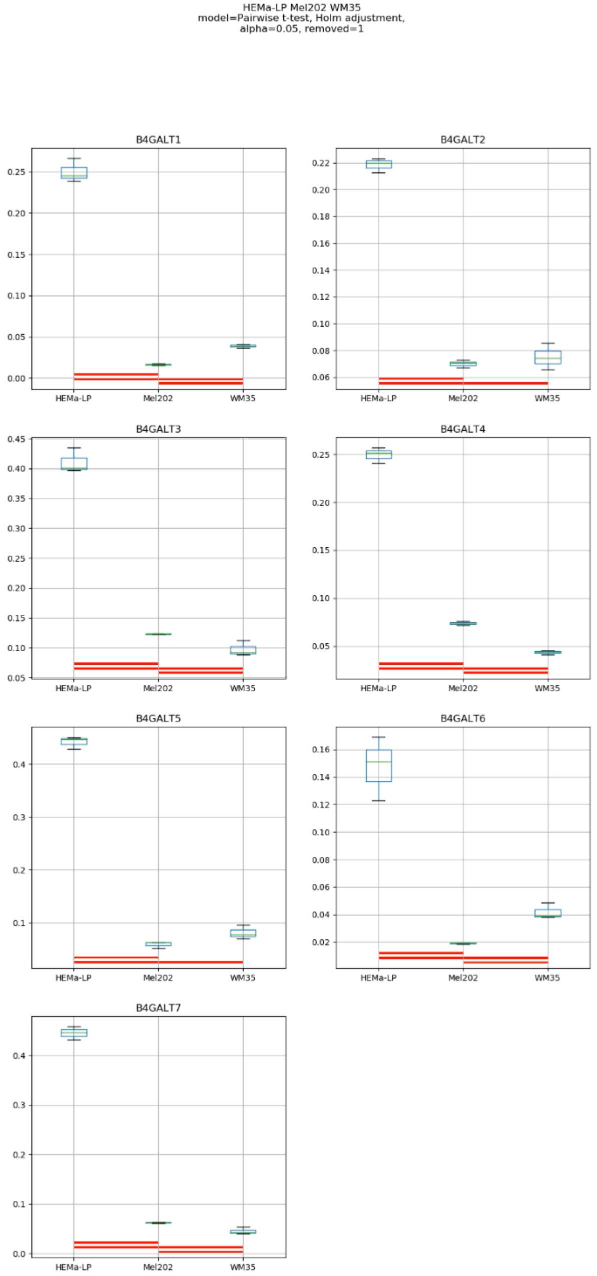
HEMa-LP Mel202 WM35
model=Pairwise t-test, Holm adjustment,
alpha=0.05, removed=1
B4GALT1
B4GALT2
B4GALT3
B4GALT4
B4GALT5
B4GALT6
B4GALT7
HEMa-LP
Mel202
WM35

### Model No. 12

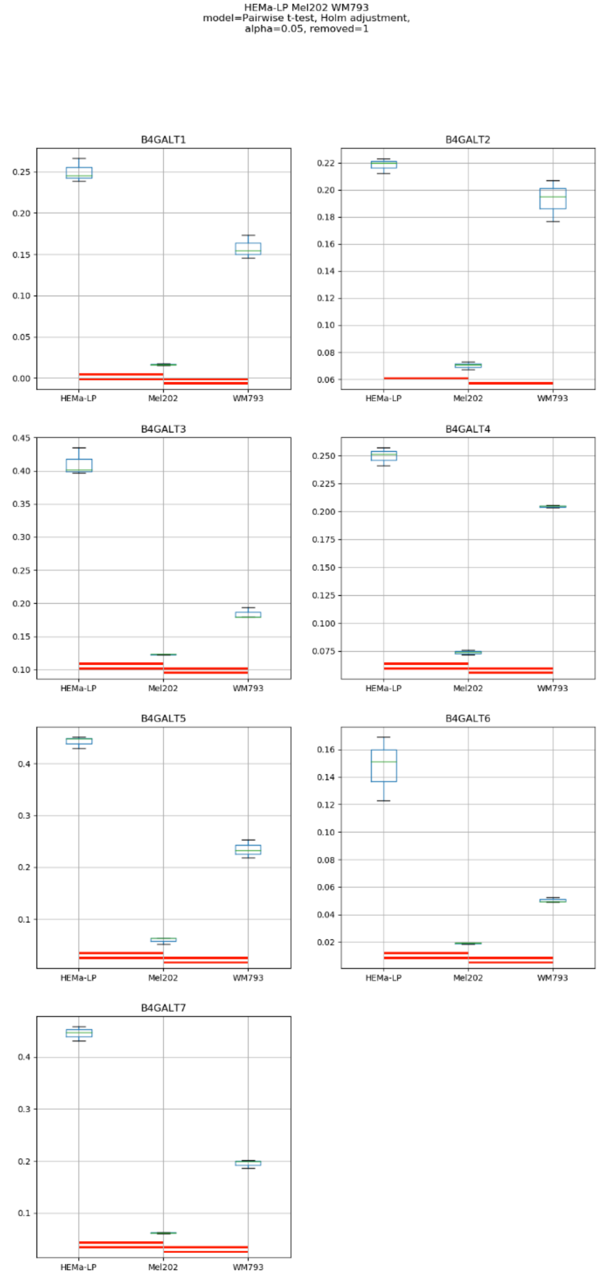
HEMa-LP Mel202 WM793
model=Pairwise t-test, Holm adjustment,
alpha=0.05, removed=1
B4GALT1
B4GALT2
B4GALT3
B4GALT4
B4GALT5
B4GALT6
B4GALT7
HEMa-LP
Mel202
WM793

### Model No. 13

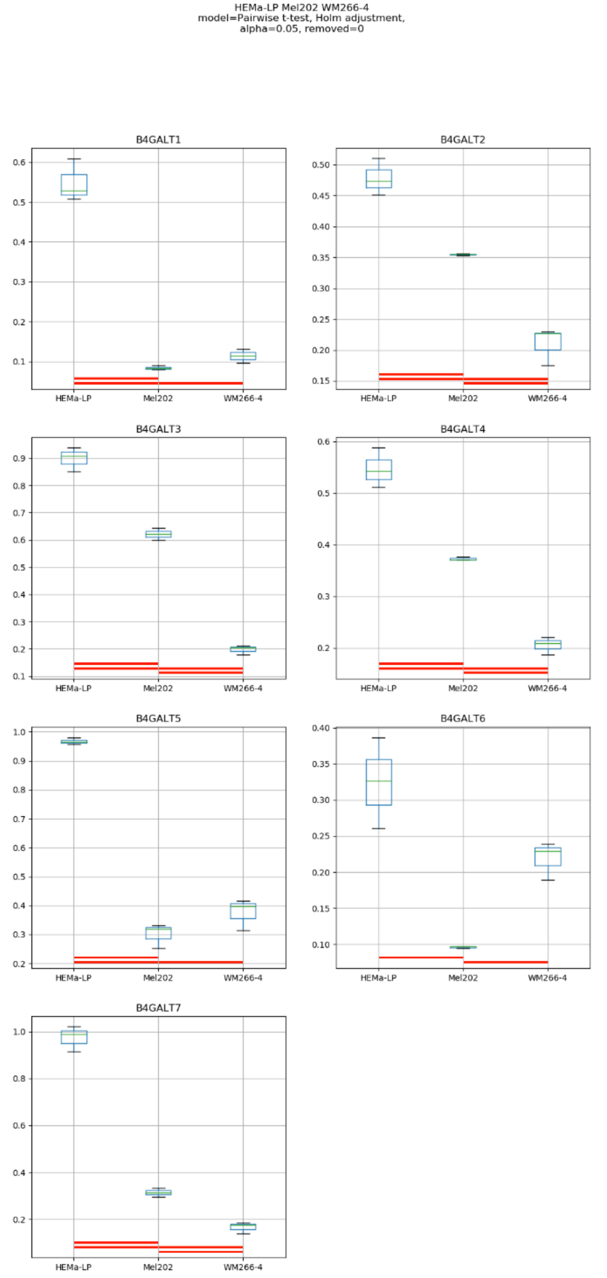

HEMa-LP Mel202 WM266-4
model=Pairwise t-test, Holm adjustment,
alpha=0.05, removed=0
B4GALT1
B4GALT2
B4GALT3
B4GALT4
B4GALT5
B4GALT6
B4GALT7
HEMa-LP
Mel202
WM266-4

### Model No. 14

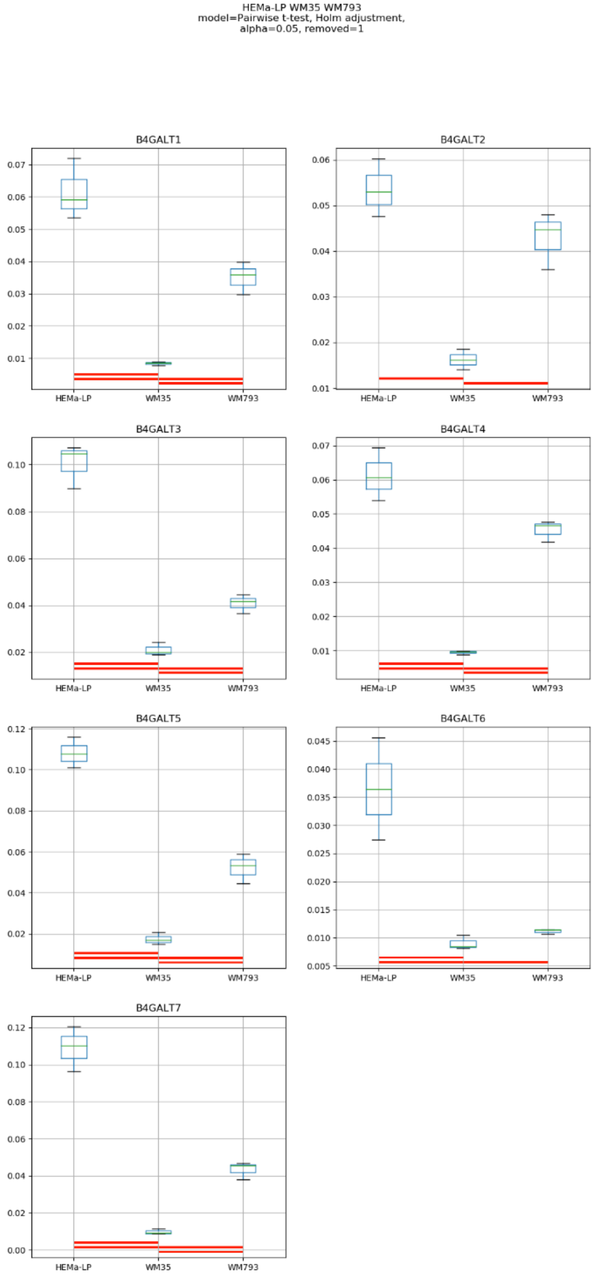
HEMa-LP WM35 WM793
model=Pairwise t-test, Holm adjustment,
alpha=0.05, removed=1
B4GALT1
B4GALT2
B4GALT3
B4GALT4
B4GALT5
B4GALT6
B4GALT7
HEMa-LP
WM35
WM793

### Model No. 15

HEMa-LP WM35 WM266-4
model=Pairwise t-test, Holm adjustment,
alpha=0.05, removed=0

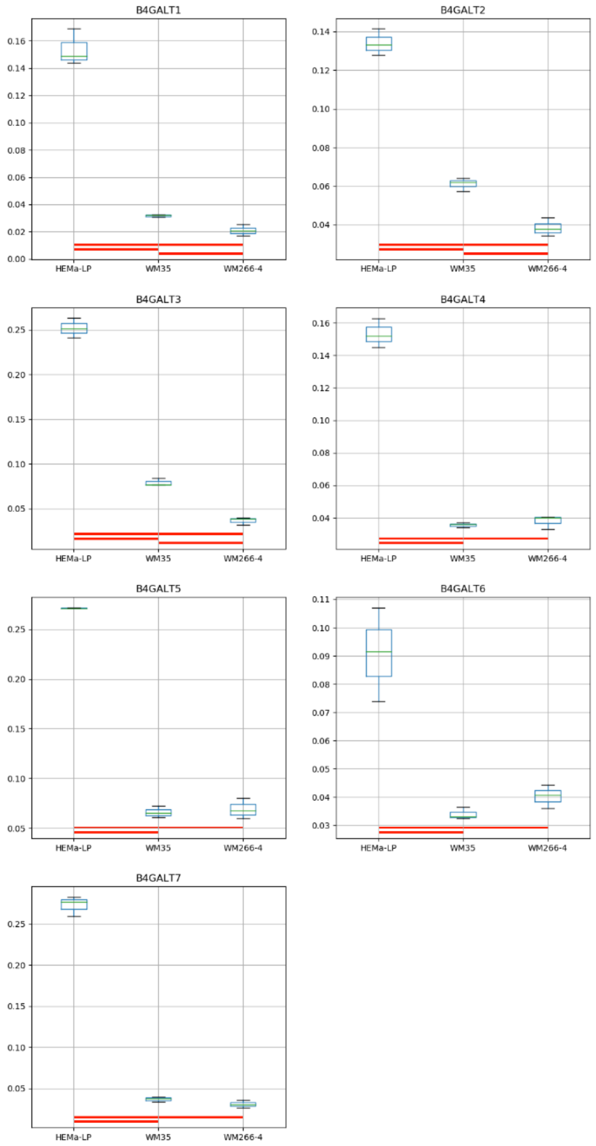

### Model No. 16

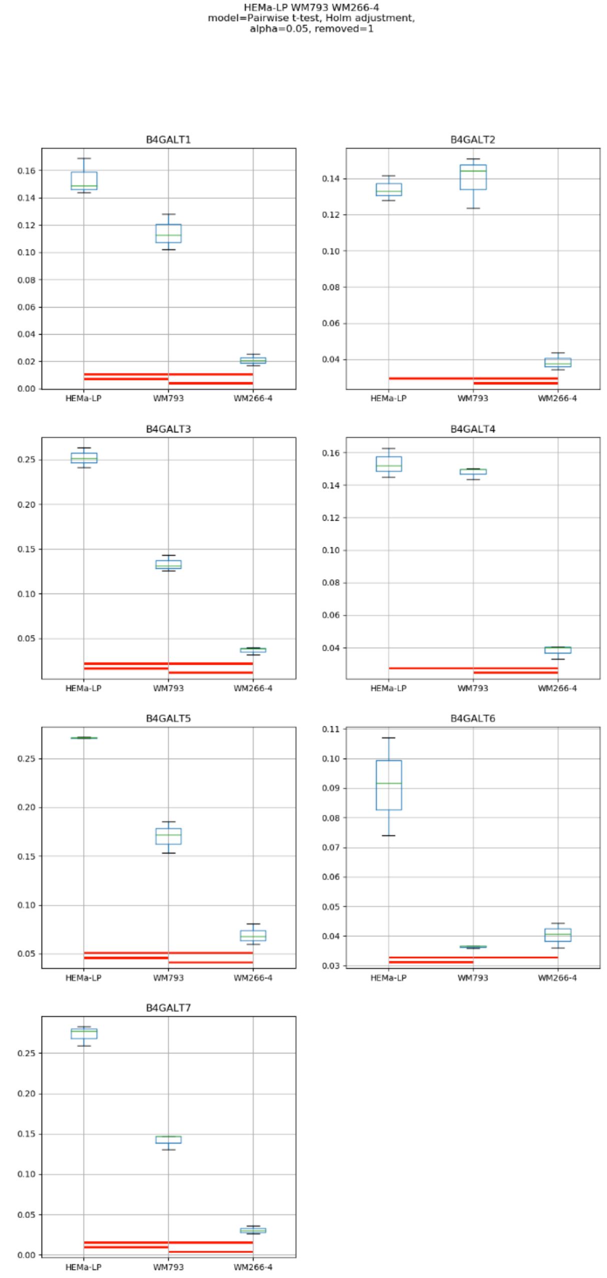
HEMa-LP WM793 WM266-4
model=Pairwise t-test, Holm adjustment,
alpha=0.05, removed=1
B4GALT1
B4GALT2
B4GALT3
B4GALT4
B4GALT5
B4GALT6
B4GALT7
HEMa-LP
WM793
WM266-4

### Model No. 17

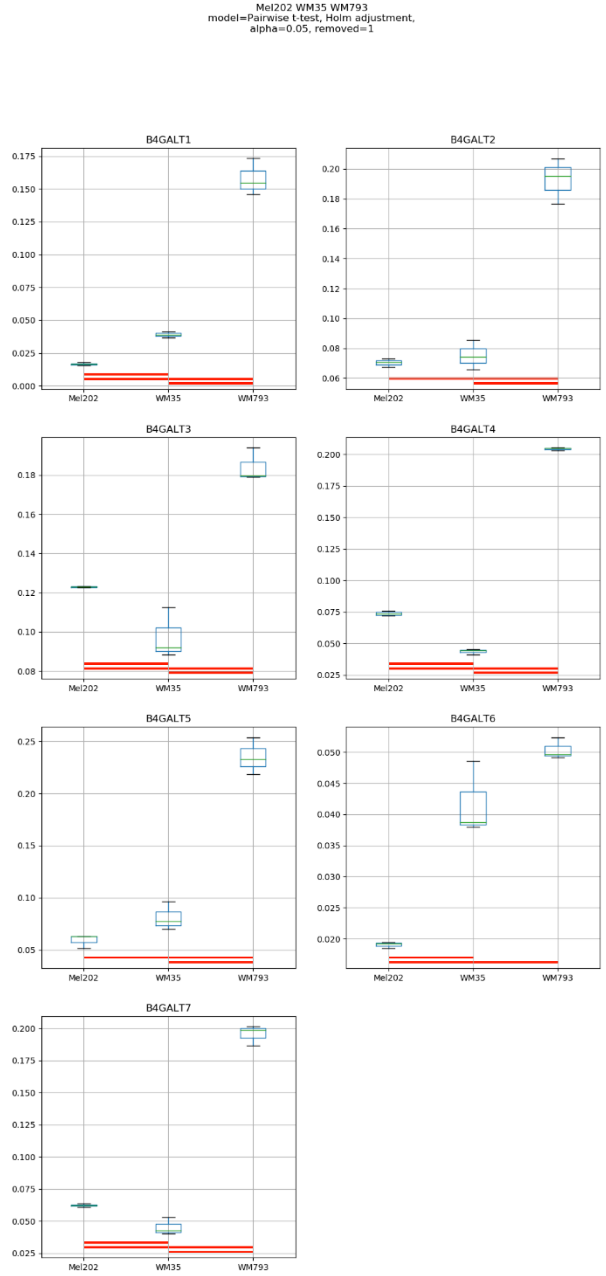
Mel202 WM35 WM793
model=Pairwise t-test, Holm adjustment,
alpha=0.05, removed=1
B4GALT1
B4GALT2
B4GALT3
B4GALT4
B4GALT5
B4GALT6
B4GALT7
Mel202
WM35
WM793

Model No. 18

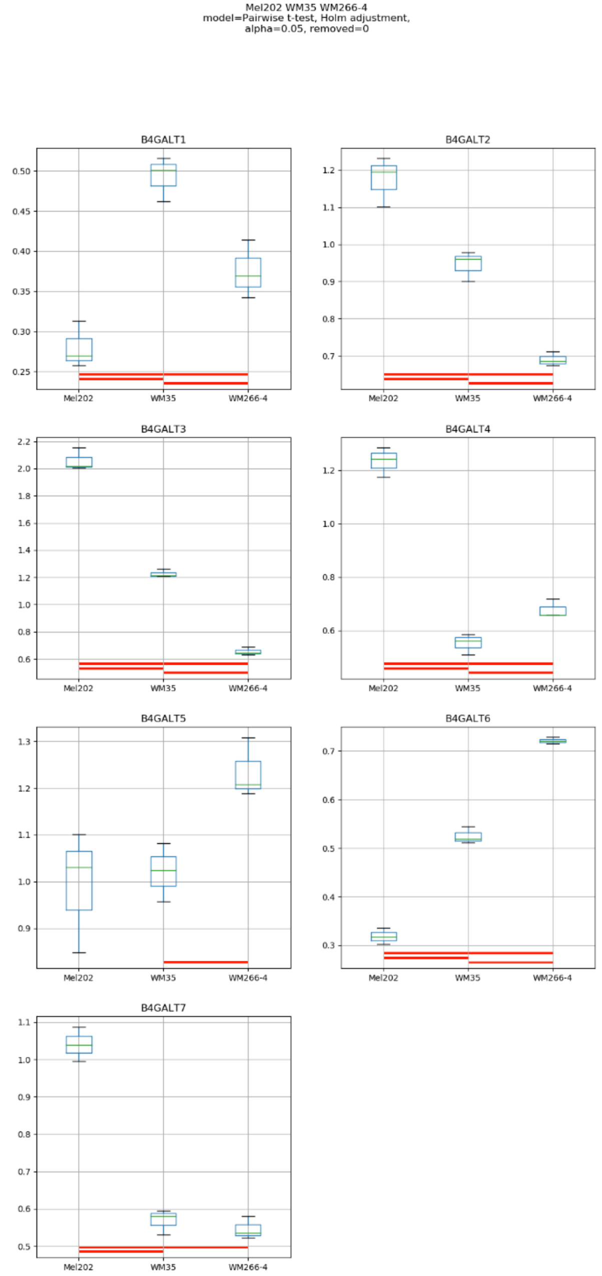
Mel202 WM35 WM266-4
model=Pairwise t-test, Holm adjustment,
alpha=0.05, removed=0
B4GALT1
B4GALT2
B4GALT3
B4GALT4
B4GALT5
B4GALT6
B4GALT7
Mel202
WM35
WM266-4

### Model No. 19

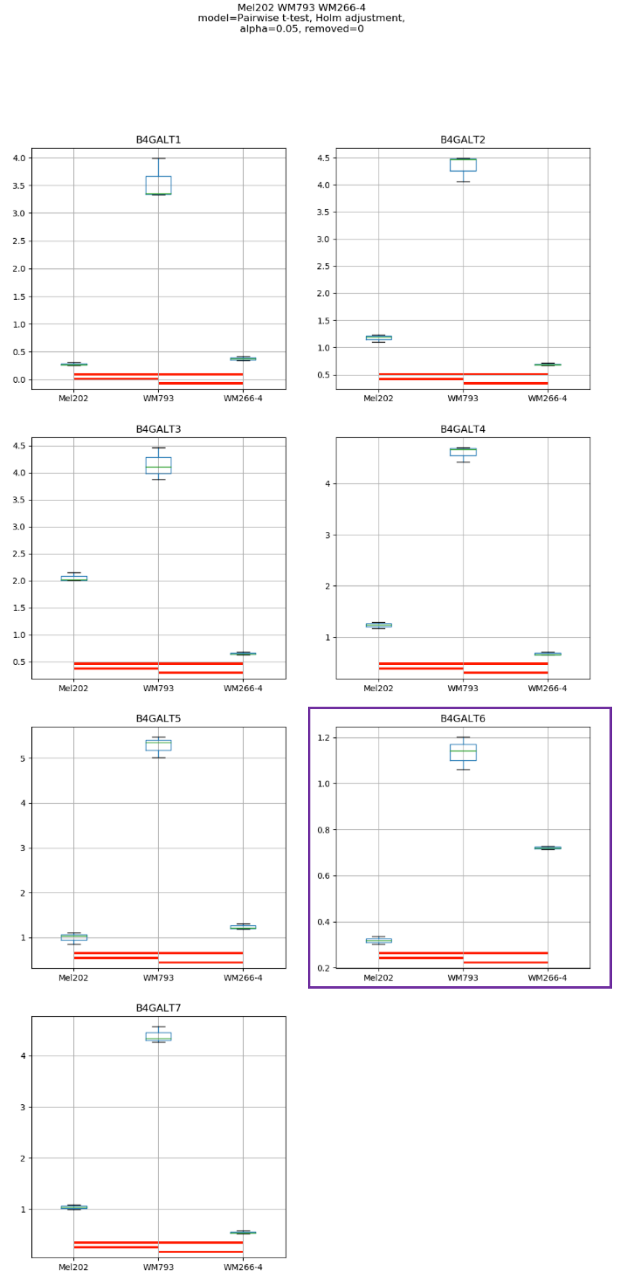
Mel202 WM793 WM266-4
model=Pairwise t-test, Holm adjustment,
alpha=0.05, removed=0
B4GALT1
B4GALT2
B4GALT3
B4GALT4
B4GALT5
B4GALT6
B4GALT7
Mel202
WM793
WM266-4

### Model No. 20

WM35 WM793 WM266-4
Mann-Whitney/Kruska-Wallis, post: Dunn's test,
alpha=0.05, removed=0

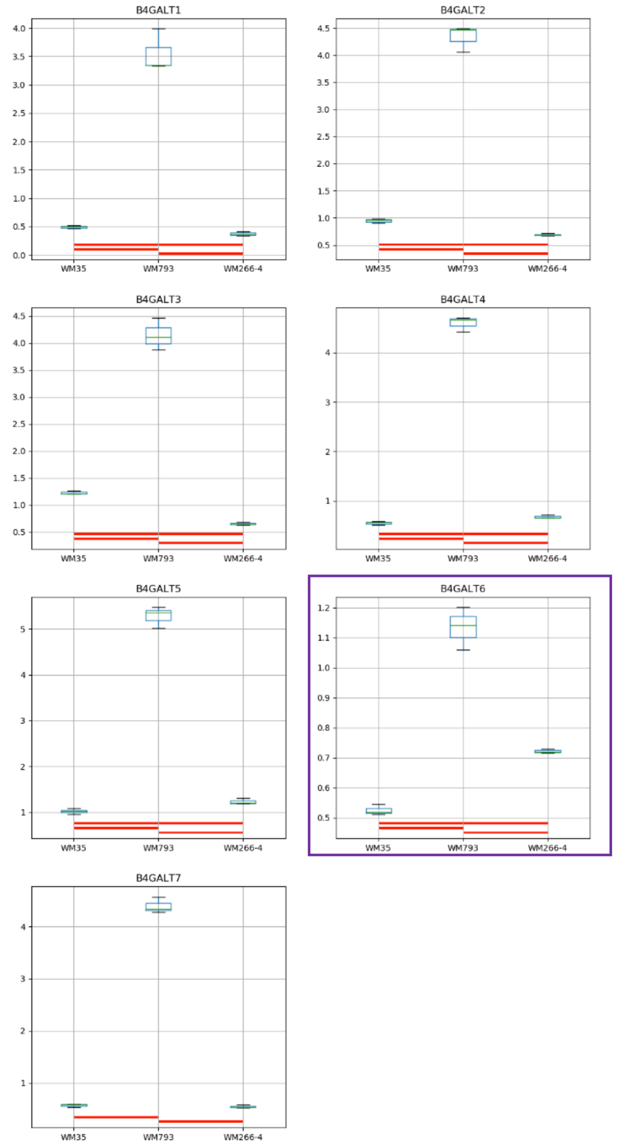

### Model No. 21

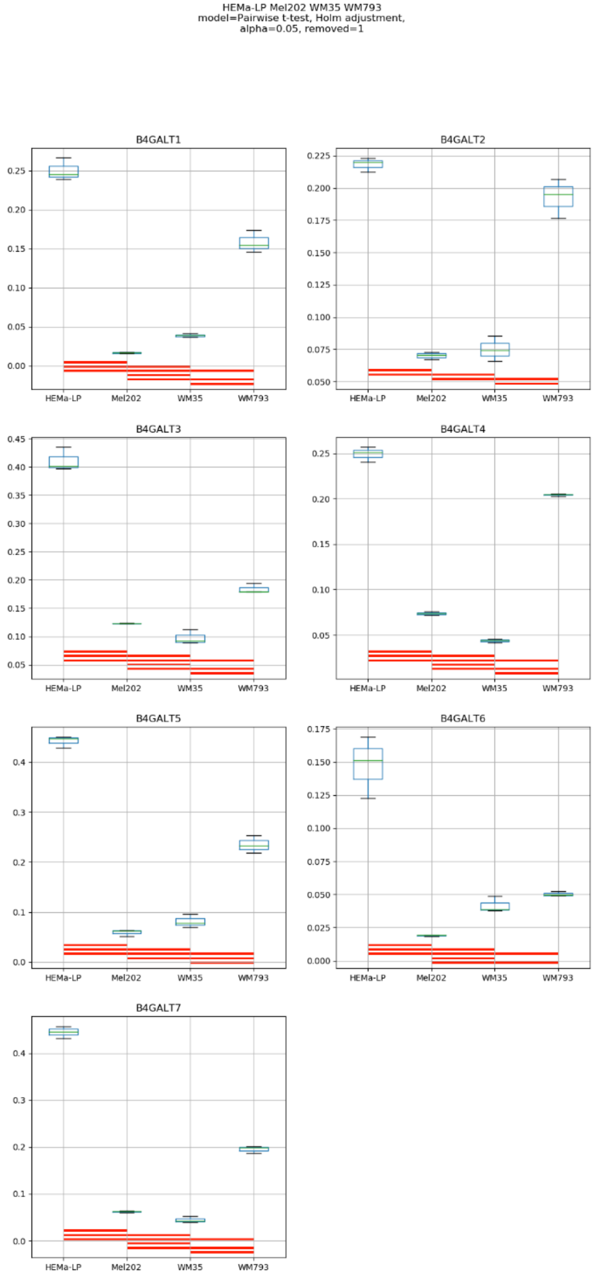

HEMa-LP Mel202 WM35 WM793
model=Pairwise t-test, Holm adjustment,
alpha=0.05, removed=1
B4GALT1
B4GALT2
B4GALT3
B4GALT4
B4GALT5
B4GALT6
B4GALT7
HEMa-LP
Mel202
WM35
WM793

### Model No. 22

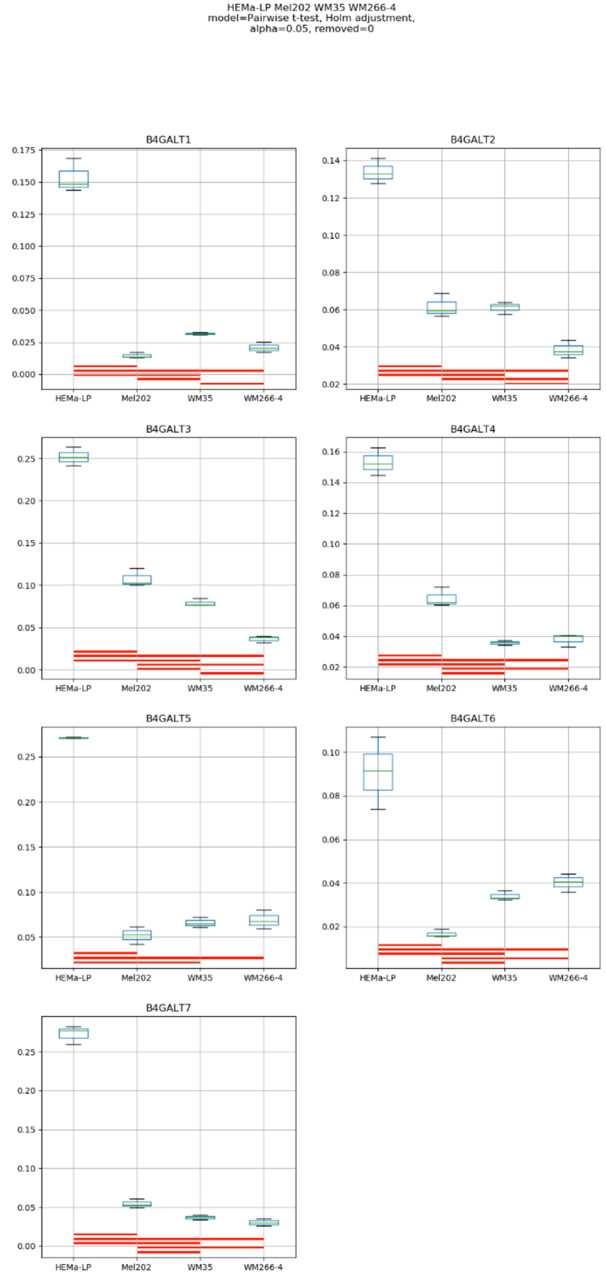
HEMa-LP Mel202 WM35 WM266-4
model=Pairwise t-test, Holm adjustment,
alpha=0.05, removed=0
B4GALT1
B4GALT2
B4GALT3
B4GALT4
B4GALT5
B4GALT6
B4GALT7
HEMa-LP
Mel202
WM35
WM266-4

### Model No. 23

HEMa-LP Mel202 WM793 WM266-4
model=Pairwise t-test, Holm adjustment,
alpha=0.05, removed=1

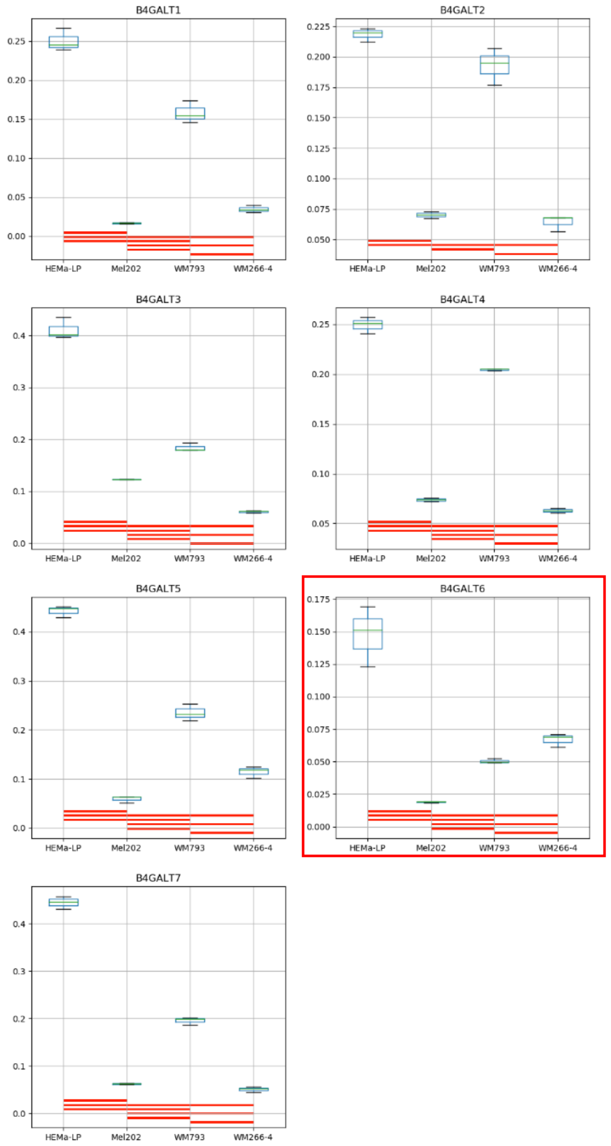

### Model No. 24

HEMa-LP WM35 WM793 WM266-4
model=Pairwise t-test, Holm adjustment,
alpha=0.05, removed=1

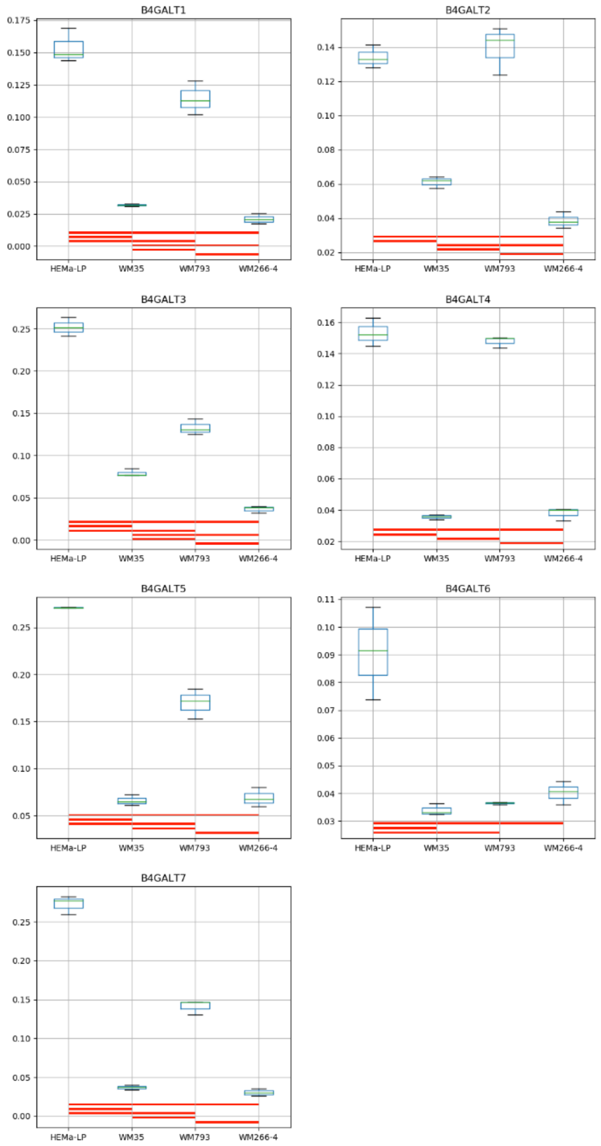

### Model No. 25

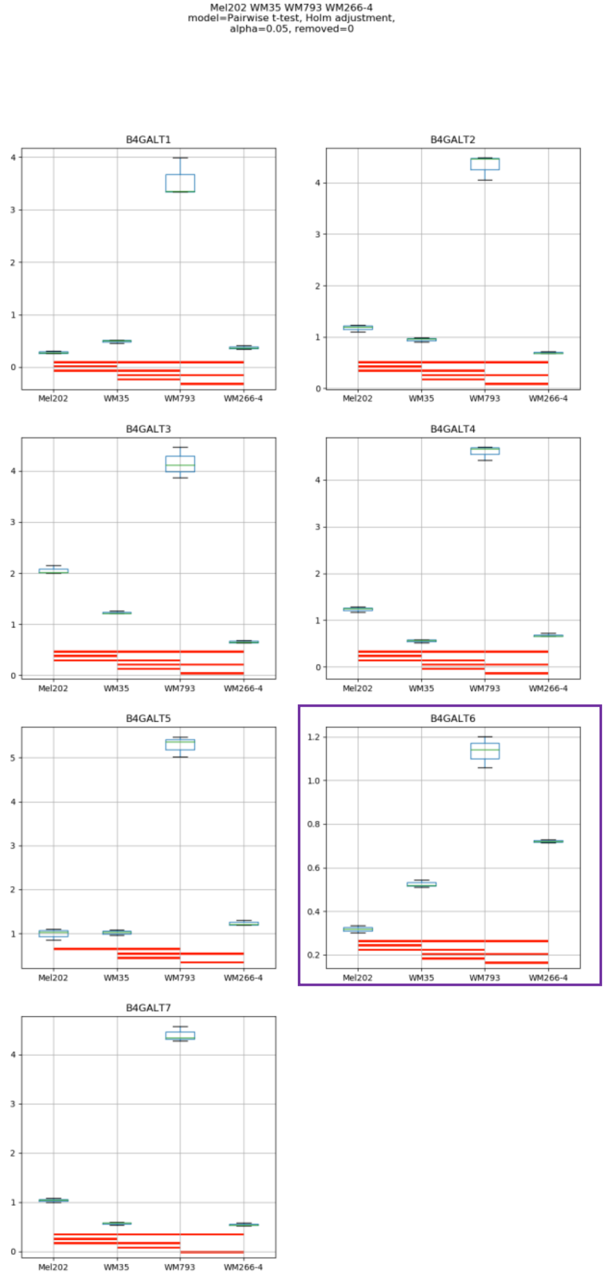

Mel202 WM35 WM793 WM266-4
model=Pairwise t-test, Holm adjustment,
alpha=0.05, removed=0
B4GALT1
B4GALT2
B4GALT3
B4GALT4
B4GALT5
B4GALT6
B4GALT7
Mel202
WM35
WM793
WM266-4

Model No. 26

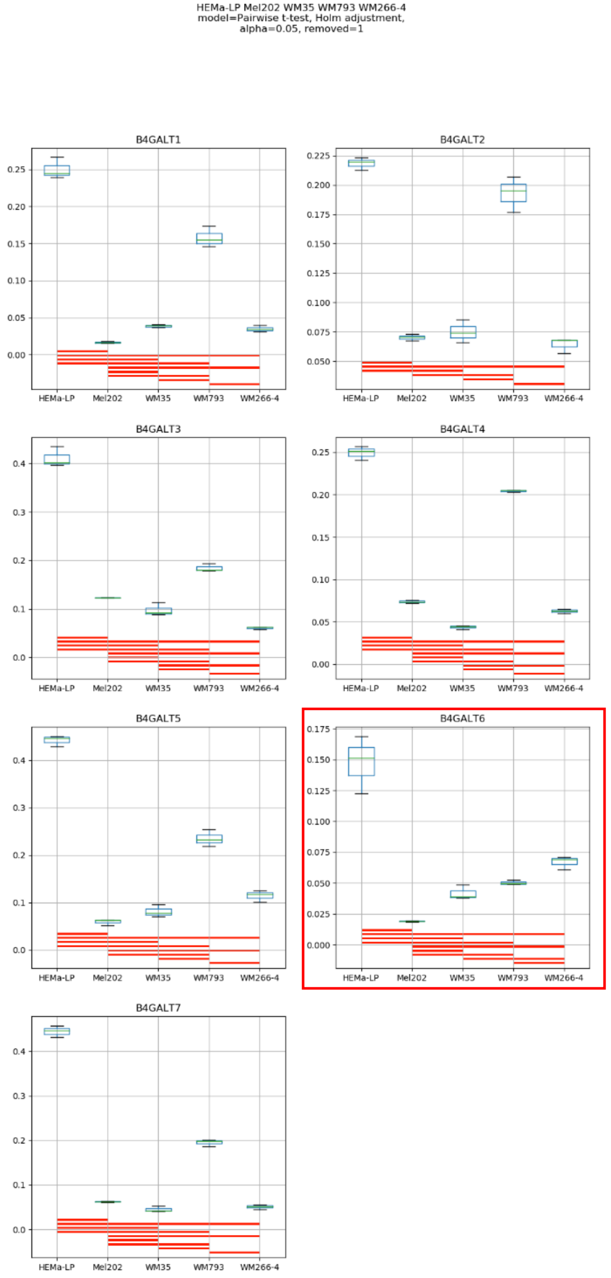

HEMa-LP Mel202 WM35 WM793 WM266-4
model=Pairwise t-test, Holm adjustment,
alpha=0.05, removed=1
B4GALT1
B4GALT2
B4GALT3
B4GALT4
B4GALT5
B4GALT6
B4GALT7
HEMa-LP
Mel202
WM35
WM793
WM266-4

**Supplementary Figure S4:** Medians of relative quantity (RQ) for target genes B4GALT1–B4GALT7 normalized to the reference gene or pair of reference genes with the best stability values resulting from GenExpA analysis based on set 1 of four candidate reference genes (HPRT1, PGK1, RPS23, SNRPA) and remove repetition level 1 (see Table 1, part A). The statistical analyses used the pairwise t-test with Holm adjustment. Red line represents statistical significance, $P < 0.05$. Frames denote box-plots representing the medians of the RQ values for a given target gene without a repeatable pattern in the same cell lines under different models (the 'Remove repetitions' box set to 0 – violet frame,  or to 1 – red frame). Detailed results are listed in Supplementary Table 4.

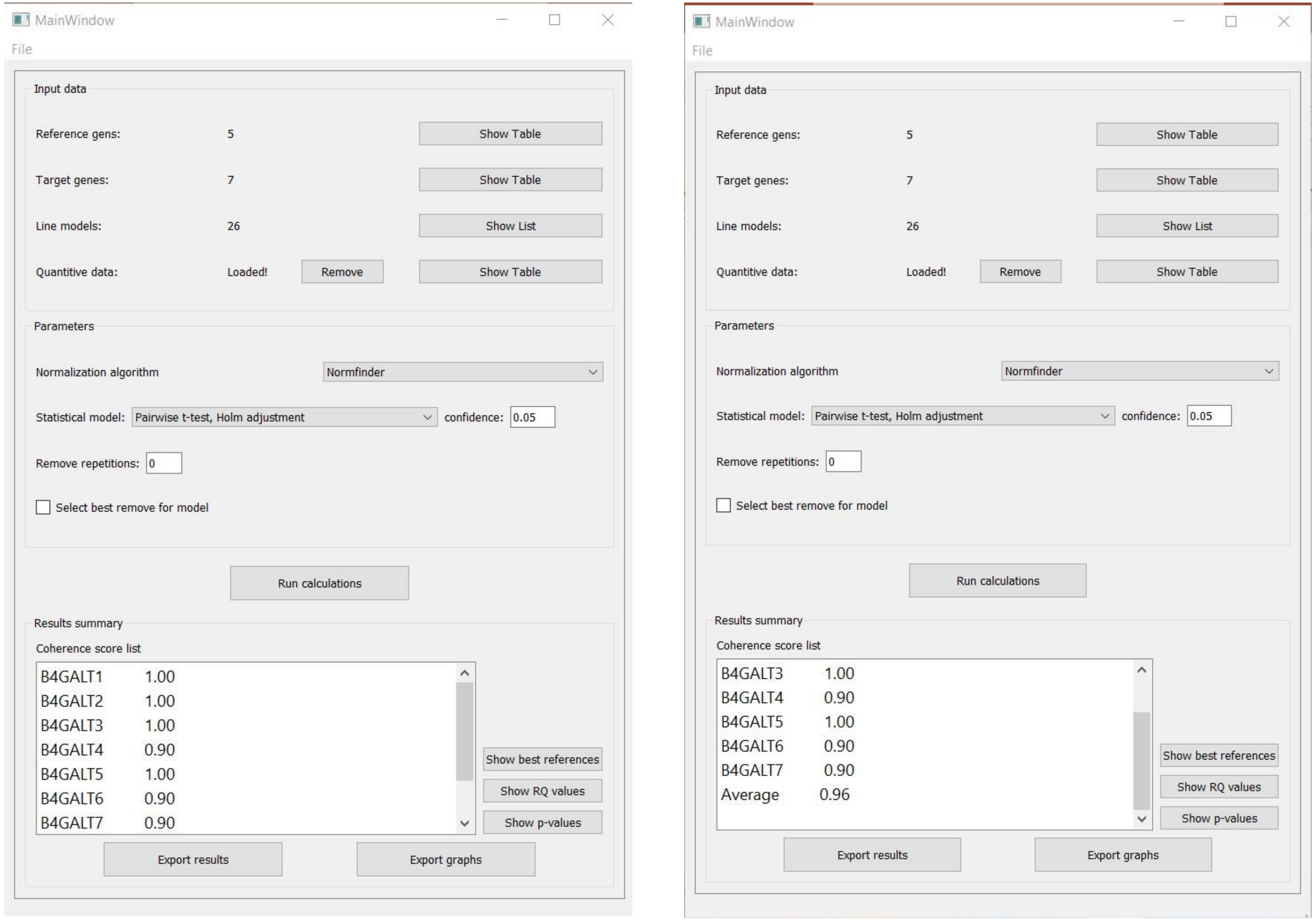


**Supplementary Figure S5:** Print screen of a GenExpA main window with selected parameters of the analysis and coherence score (CS) values for target genes *B4GALT1–B4GALT7* and average CS value of the analysis.

### Model No. 1

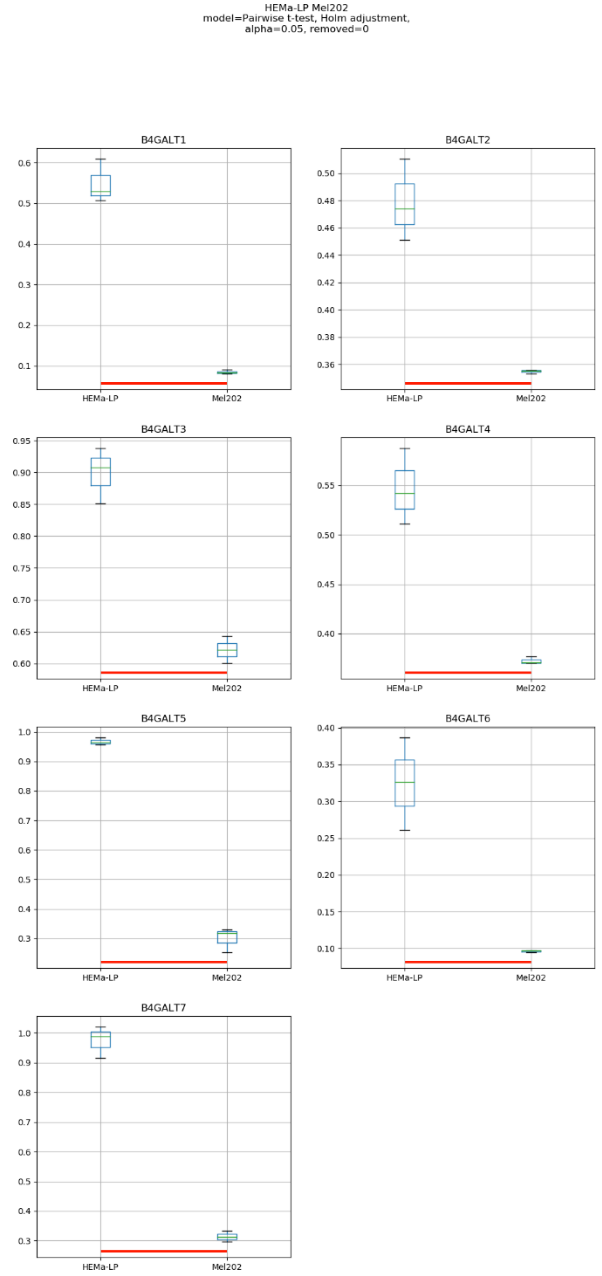
HEMa-LP Mel202
model=Pairwise t-test, Holm adjustment,
alpha=0.05, removed=0
B4GALT1
B4GALT2
B4GALT3
B4GALT4
B4GALT5
B4GALT6
B4GALT7
HEMa-LP
Mel202

### Model No. 2

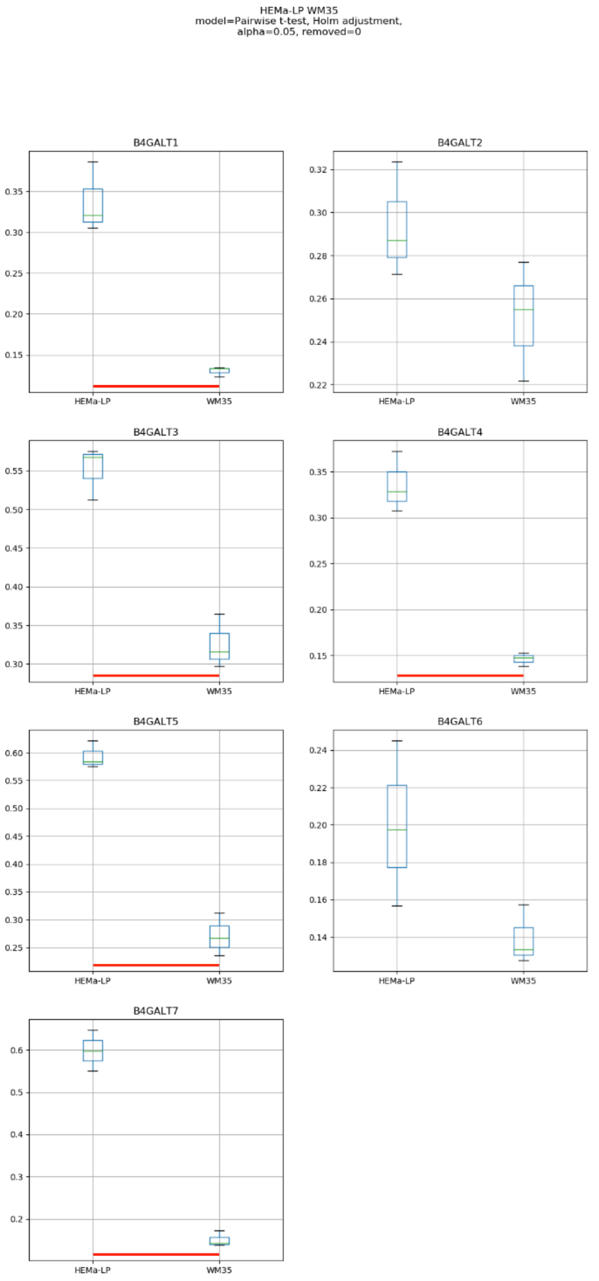

HEMa-LP WM35
model=Pairwise t-test, Holm adjustment,
alpha=0.05, removed=0
B4GALT1
B4GALT2
B4GALT3
B4GALT4
B4GALT5
B4GALT6
B4GALT7
HEMa-LP
WM35

### Model No. 3

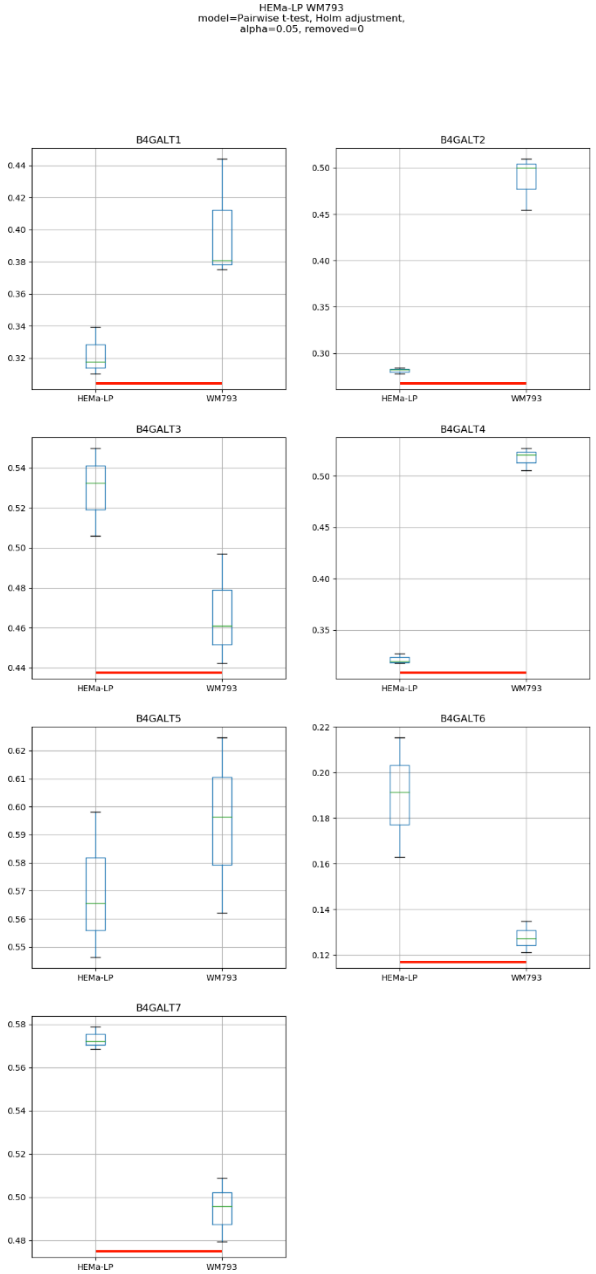
HEMa-LP WM793
model=Pairwise t-test, Holm adjustment,
alpha=0.05, removed=0
B4GALT1
B4GALT2
B4GALT3
B4GALT4
B4GALT5
B4GALT6
B4GALT7
HEMa-LP
WM793

### Model No. 4

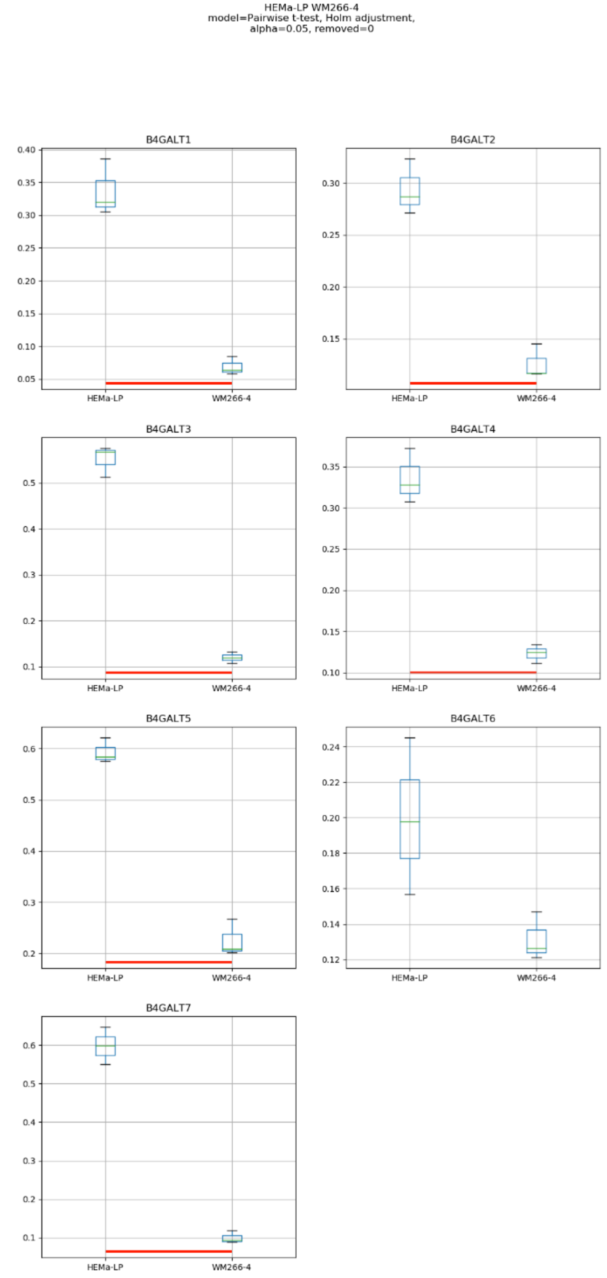
HEMa-LP WM266-4
model=Pairwise t-test, Holm adjustment,
alpha=0.05, removed=0
B4GALT1
B4GALT2
B4GALT3
B4GALT4
B4GALT5
B4GALT6
B4GALT7
HEMa-LP
WM266-4

### Model No. 5

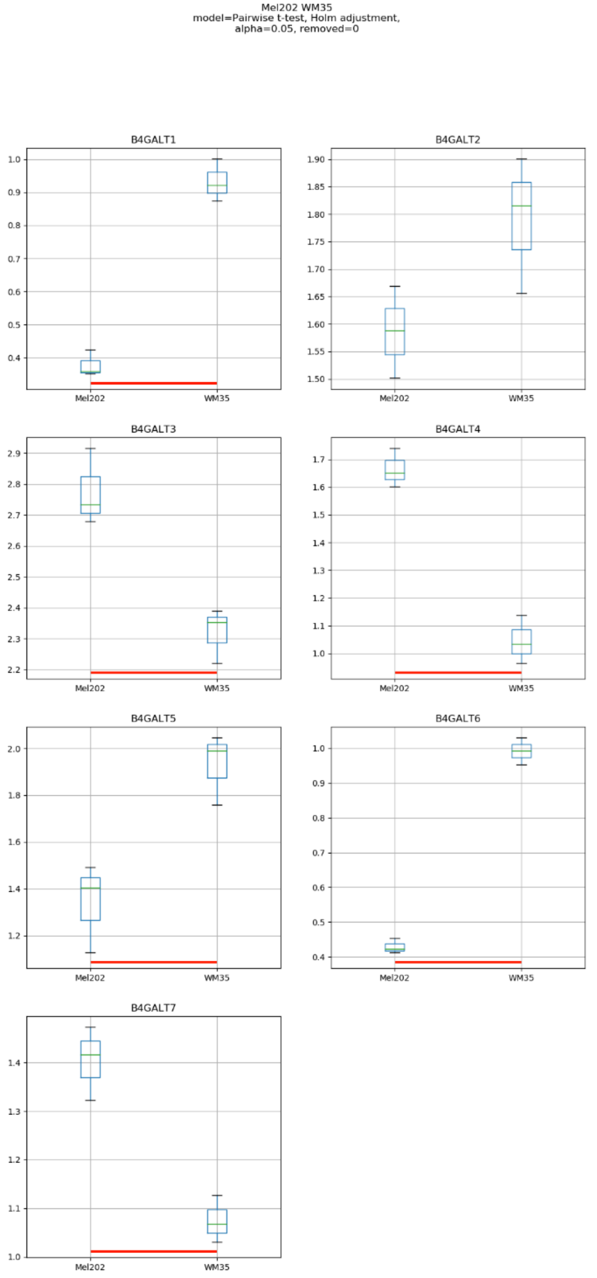
Mel202 WM35
model=Pairwise t-test, Holm adjustment,
alpha=0.05, removed=0
B4GALT1
B4GALT2
B4GALT3
B4GALT4
B4GALT5
B4GALT6
B4GALT7
Mel202
WM35

### Model No. 6

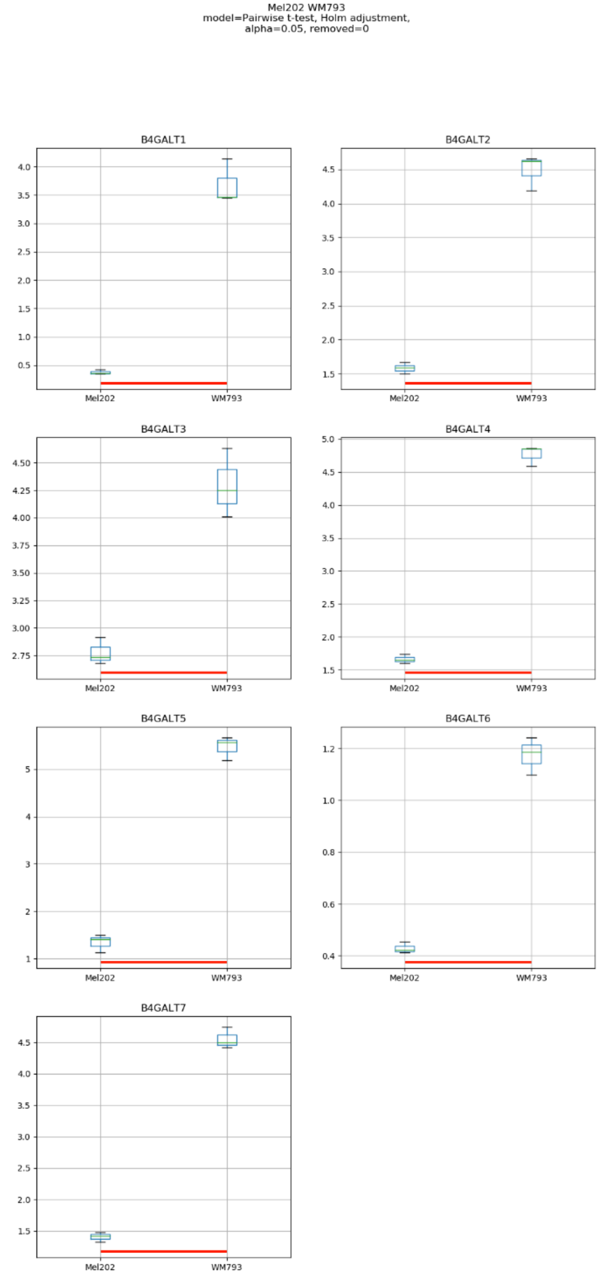

Mel202 WM793
model=Pairwise t-test, Holm adjustment,
alpha=0.05, removed=0
B4GALT1
B4GALT2
B4GALT3
B4GALT4
B4GALT5
B4GALT6
B4GALT7
Mel202
WM793

### Model No. 7

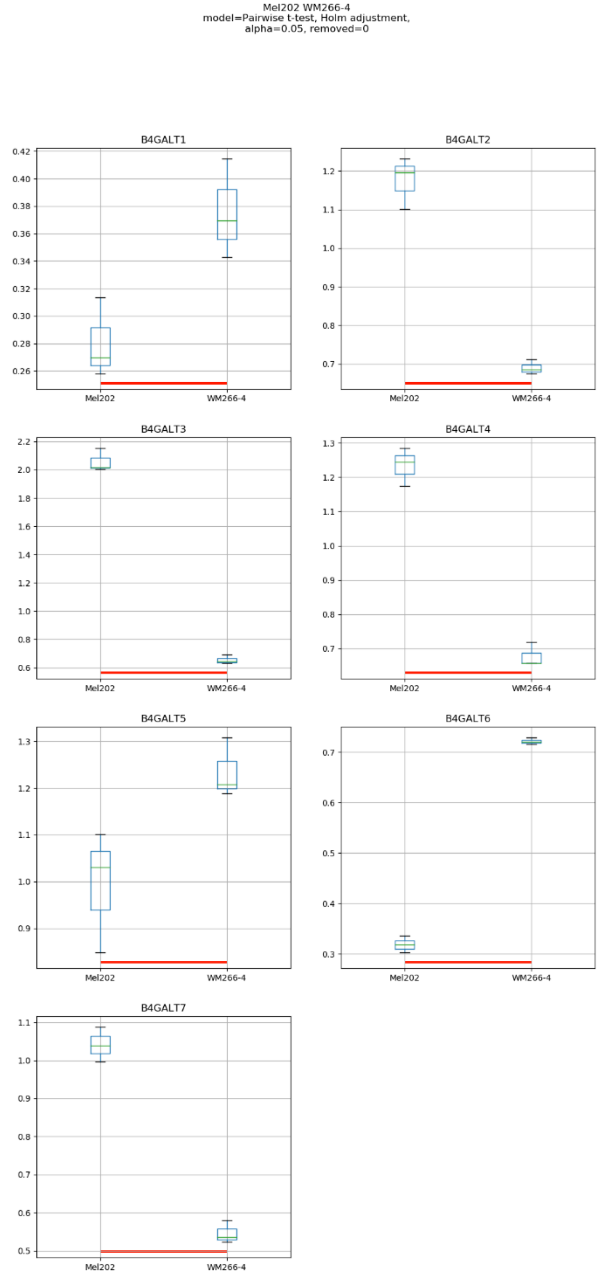

Mel202 WM266-4
model=Pairwise t-test, Holm adjustment,
alpha=0.05, removed=0
B4GALT1
0.42
0.40
0.38
0.36
0.34
0.32
0.30
0.28
0.26
Mel202
WM266-4
B4GALT2
1.2
1.1
1.0
0.9
0.8
0.7
Mel202
WM266-4
B4GALT3
2.2
2.0
1.8
1.6
1.4
1.2
1.0
0.8
0.6
Mel202
WM266-4
B4GALT4
1.3
1.2
1.1
1.0
0.9
0.8
0.7
Mel202
WM266-4
B4GALT5
1.3
1.2
1.1
1.0
0.9
Mel202
WM266-4
B4GALT6
0.7
0.6
0.5
0.4
0.3
Mel202
WM266-4
B4GALT7
1.1
1.0
0.9
0.8
0.7
0.6
0.5
Mel202
WM266-4

### Model No. 8

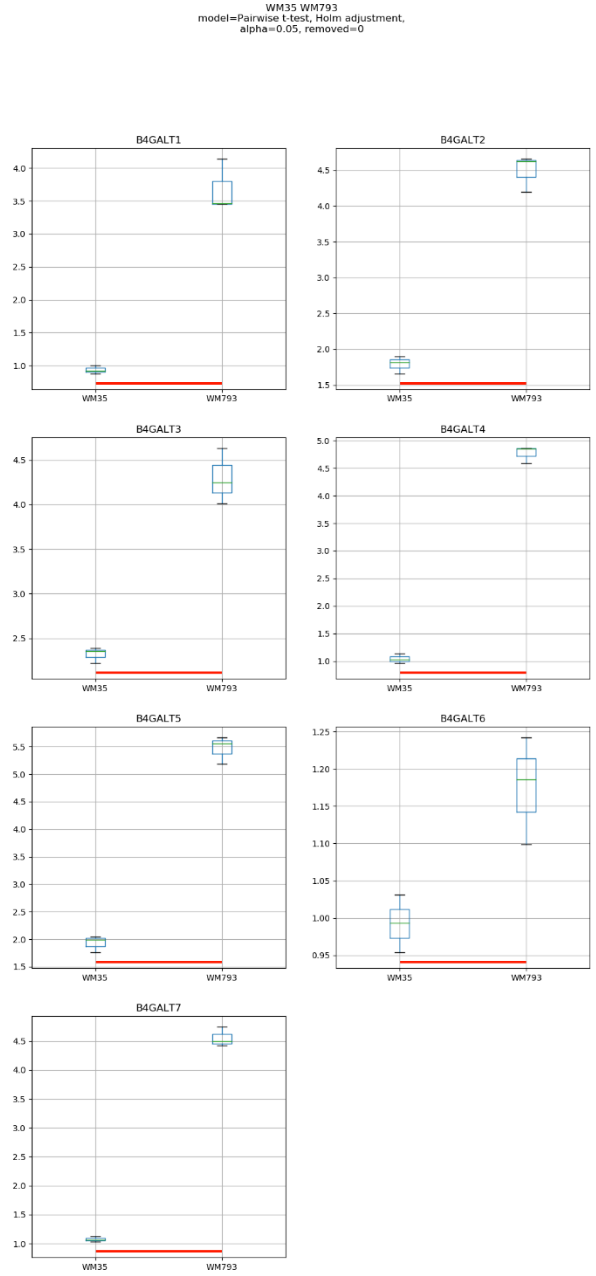
WM35 WM793
model=Pairwise t-test, Holm adjustment,
alpha=0.05, removed=0
B4GALT1
B4GALT2
B4GALT3
B4GALT4
B4GALT5
B4GALT6
B4GALT7
WM35
WM793

### Model No. 9

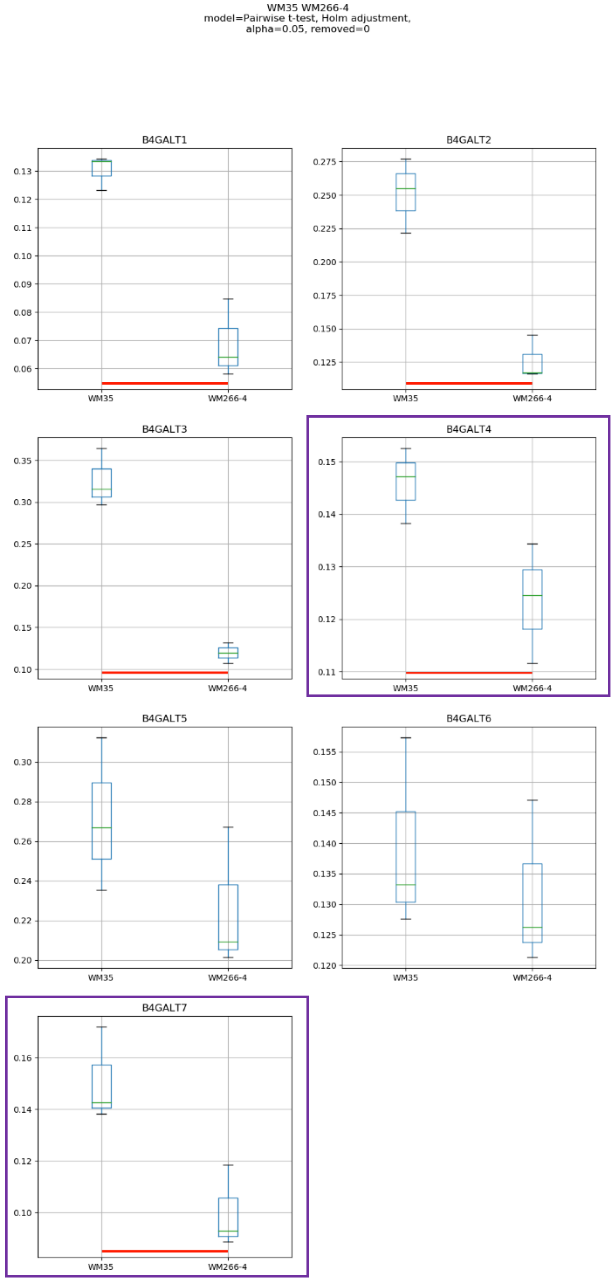
WM35 WM266-4
model=Pairwise t-test, Holm adjustment,
alpha=0.05, removed=0
B4GALT1
B4GALT2
B4GALT3
B4GALT4
B4GALT5
B4GALT6
B4GALT7
WM35
WM266-4

Model No. 10

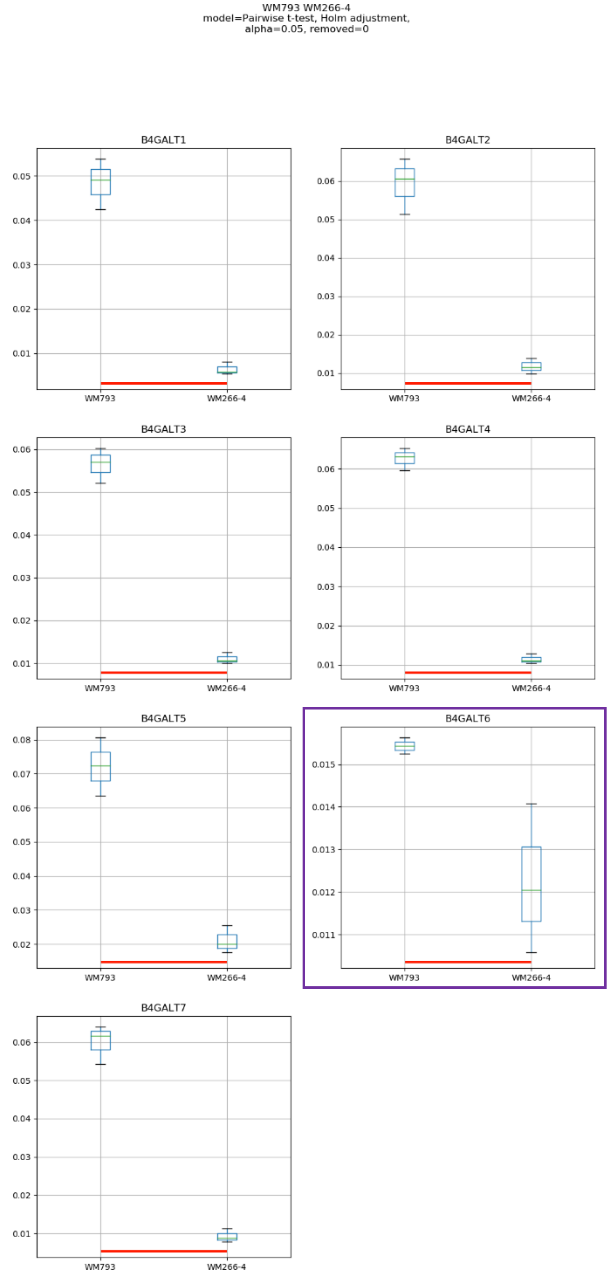
WM793 WM266-4
model=Pairwise t-test, Holm adjustment,
alpha=0.05, removed=0
B4GALT1
B4GALT2
B4GALT3
B4GALT4
B4GALT5
B4GALT6
B4GALT7
WM793
WM266-4

### Model No. 11

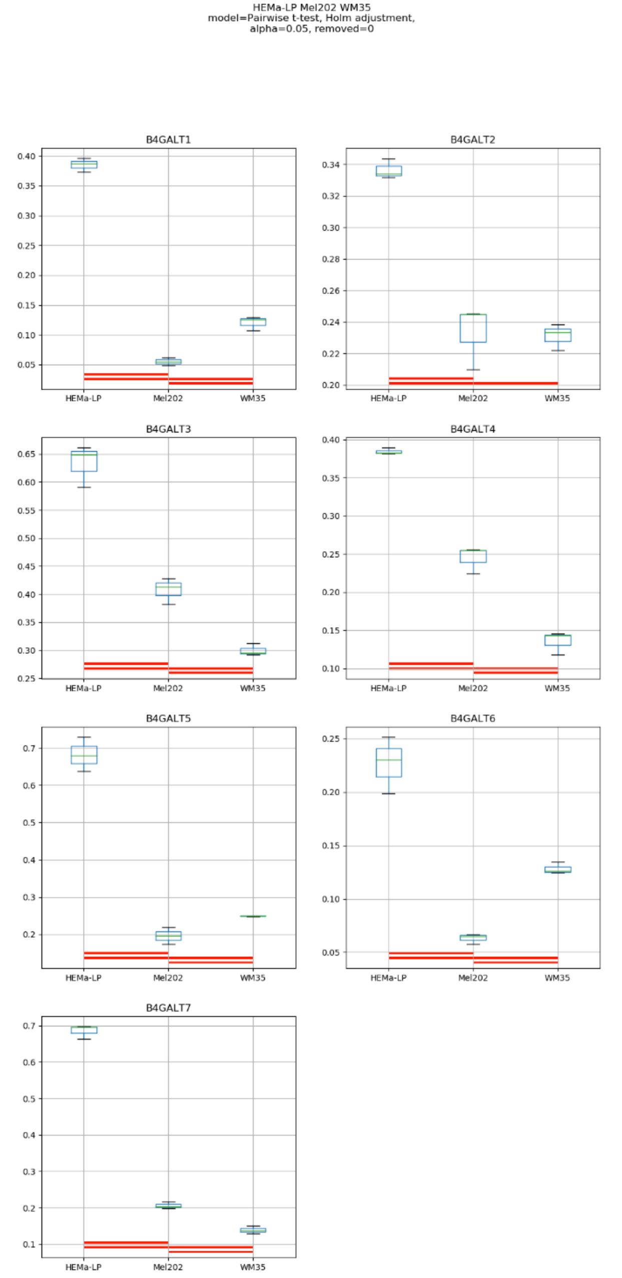
HEMa-LP Mel202 WM35
model=Pairwise t-test, Holm adjustment,
alpha=0.05, removed=0
B4GALT1
B4GALT2
B4GALT3
B4GALT4
B4GALT5
B4GALT6
B4GALT7
HEMa-LP
Mel202
WM35

### Model No. 12

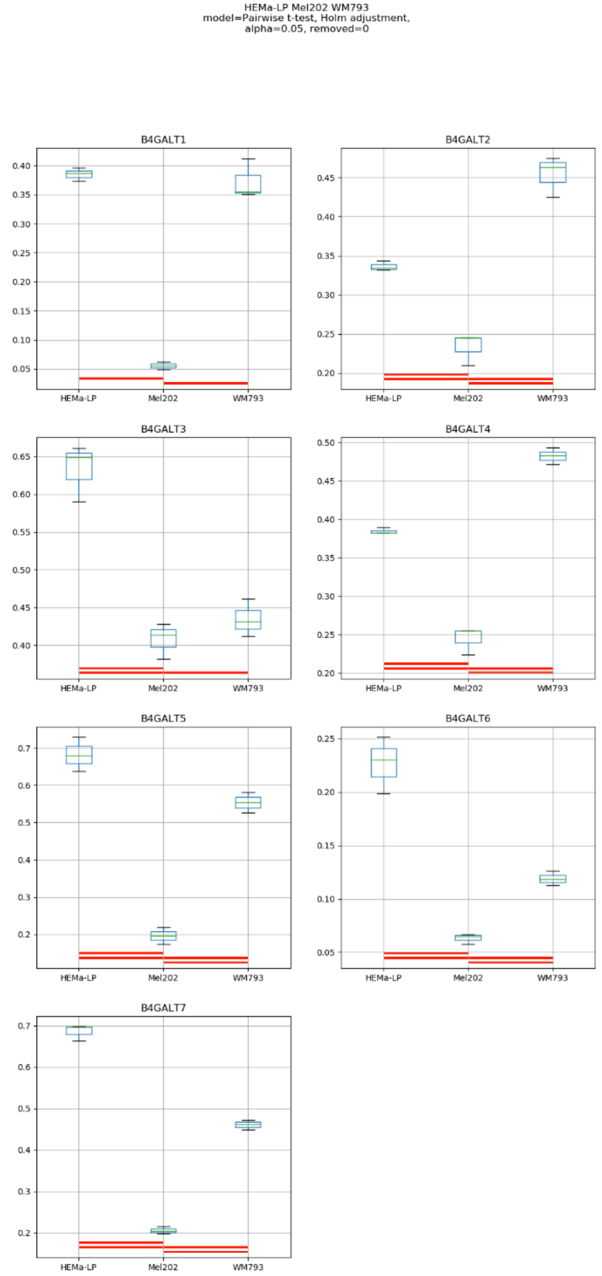
HEMa-LP Mel202 WM793
model=Pairwise t-test, Holm adjustment,
alpha=0.05, removed=0
B4GALT1
B4GALT2
B4GALT3
B4GALT4
B4GALT5
B4GALT6
B4GALT7
HEMa-LP
Mel202
WM793

### Model No. 13

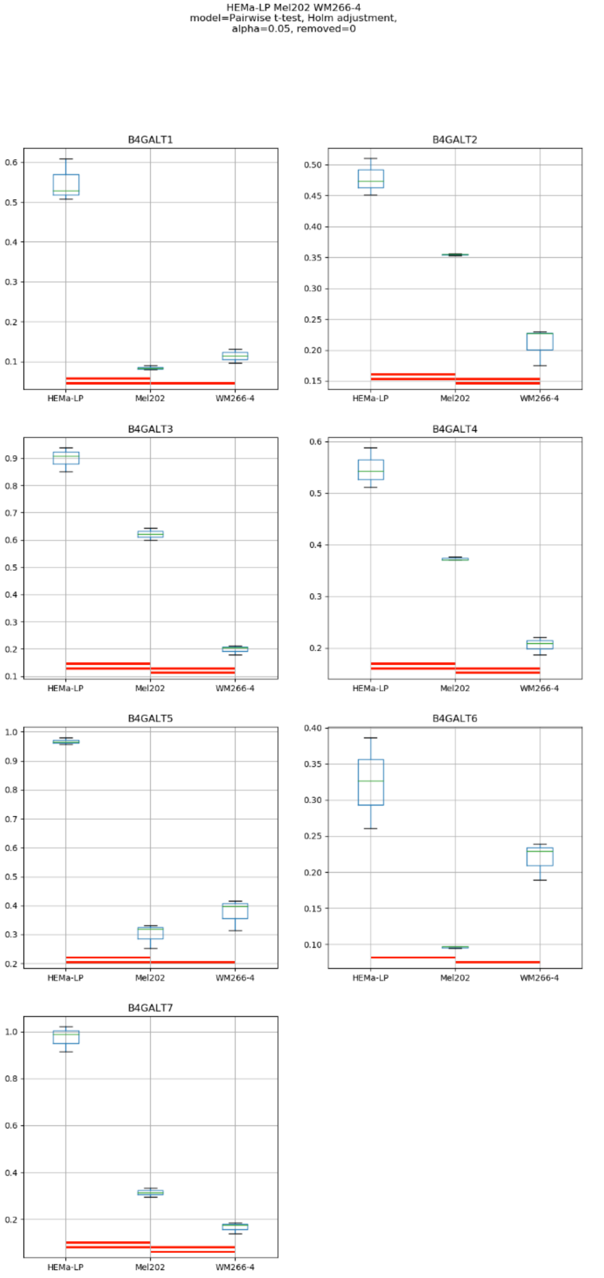

HEMa-LP Mel202 WM266-4
model=Pairwise t-test, Holm adjustment,
alpha=0.05, removed=0
B4GALT1
B4GALT2
B4GALT3
B4GALT4
B4GALT5
B4GALT6
B4GALT7
HEMa-LP
Mel202
WM266-4

Model No. 14

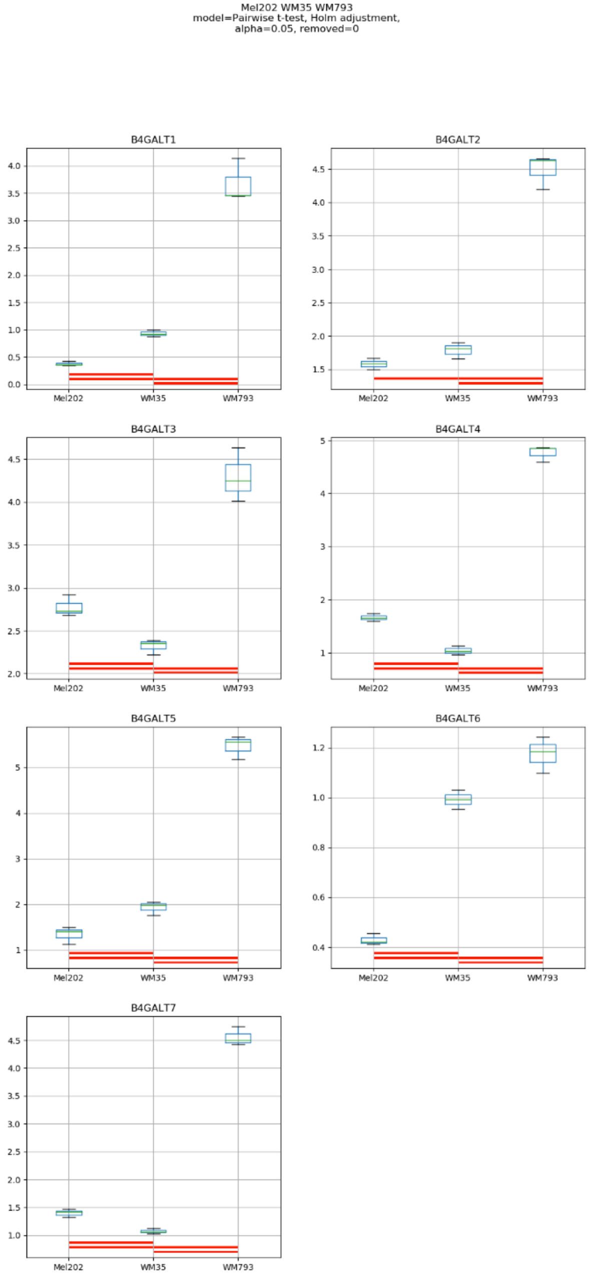
Mel202 WM35 WM793
model=Pairwise t-test, Holm adjustment,
alpha=0.05, removed=0
B4GALT1
B4GALT2
B4GALT3
B4GALT4
B4GALT5
B4GALT6
B4GALT7
Mel202
WM35
WM793

Model No. 15

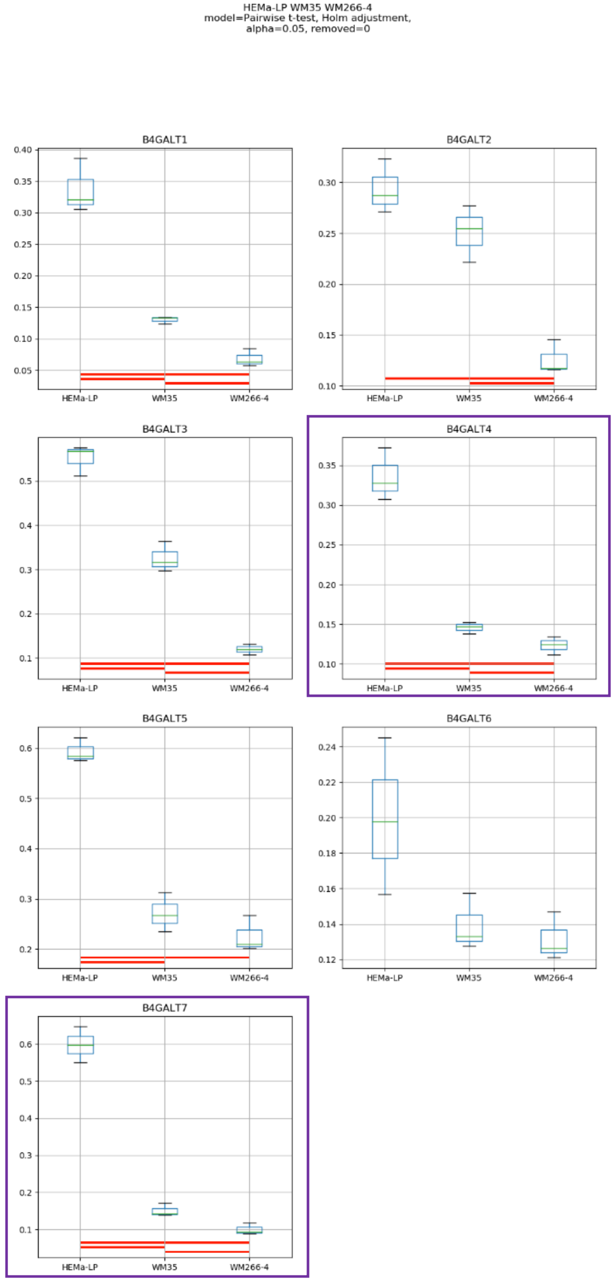
HEMa-LP WM35 WM266-4
model=Pairwise t-test, Holm adjustment,
alpha=0.05, removed=0
B4GALT1
B4GALT2
B4GALT3
B4GALT4
B4GALT5
B4GALT6
B4GALT7
HEMa-LP
WM35
WM266-4

### Model No. 16

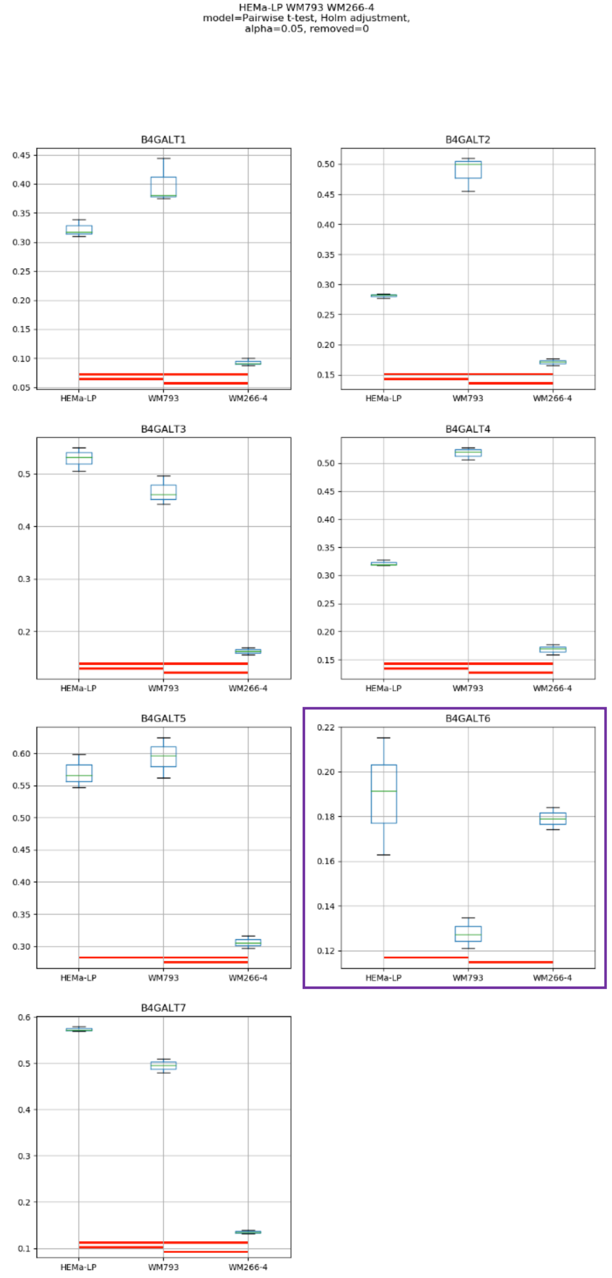
HEMa-LP WM793 WM266-4
model=Pairwise t-test, Holm adjustment,
alpha=0.05, removed=0
B4GALT1
B4GALT2
B4GALT3
B4GALT4
B4GALT5
B4GALT6
B4GALT7
HEMa-LP
WM793
WM266-4

### Model No. 17

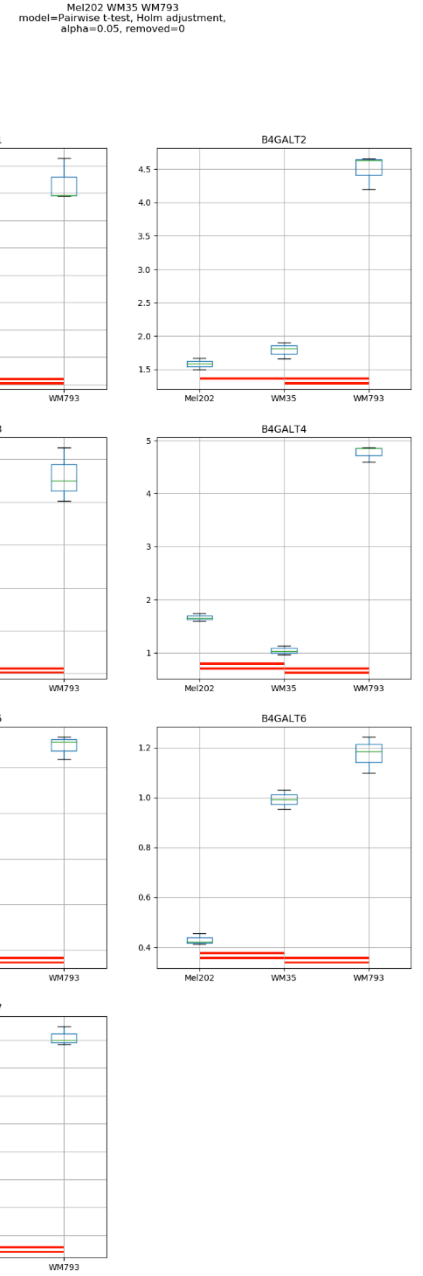
Mel202 WM35 WM793
model=Pairwise t-test, Holm adjustment,
alpha=0.05, removed=0
B4GALT1
B4GALT2
B4GALT3
B4GALT4
B4GALT5
B4GALT6
B4GALT7
Mel202
WM35
WM793

### Model No. 18

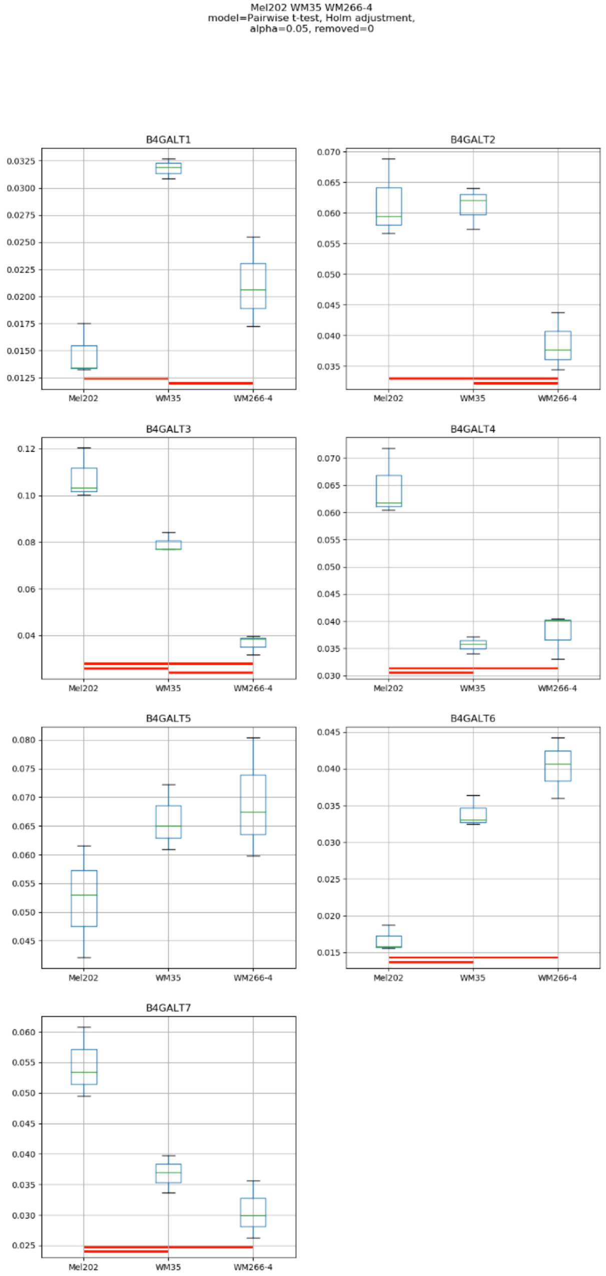
Mel202 WM35 WM266-4
model=Pairwise t-test, Holm adjustment,
alpha=0.05, removed=0
B4GALT1
B4GALT2
B4GALT3
B4GALT4
B4GALT5
B4GALT6
B4GALT7
Mel202
WM35
WM266-4

### Model No. 19

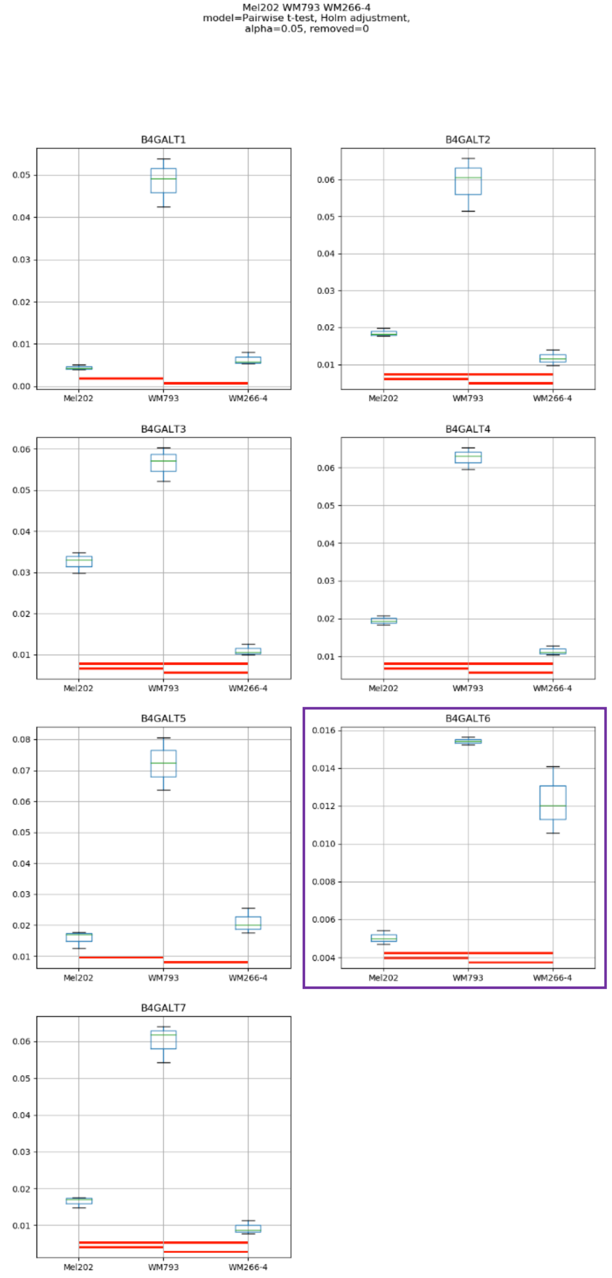
Mel202 WM793 WM266-4
model=Pairwise t-test, Holm adjustment,
alpha=0.05, removed=0
B4GALT1
B4GALT2
B4GALT3
B4GALT4
B4GALT5
B4GALT6
B4GALT7
Mel202
WM793
WM266-4

Model No. 20

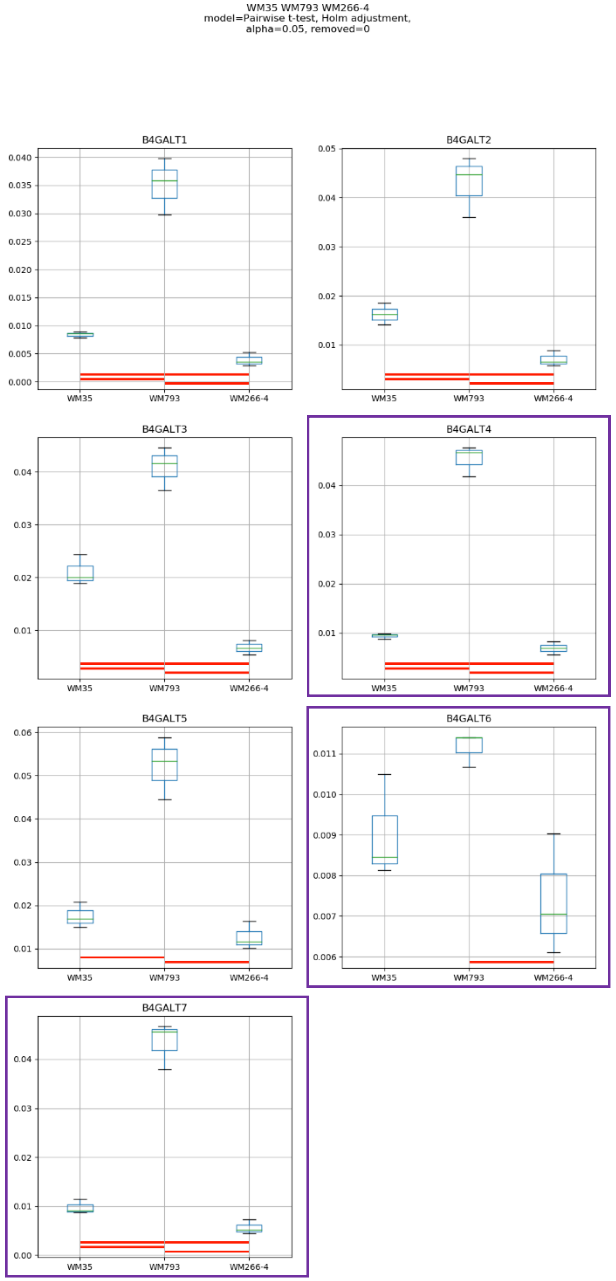
WM35 WM793 WM266-4
model=Pairwise t-test, Holm adjustment,
alpha=0.05, removed=0
B4GALT1
B4GALT2
B4GALT3
B4GALT4
B4GALT5
B4GALT6
B4GALT7
WM35
WM793
WM266-4

### Model No. 21

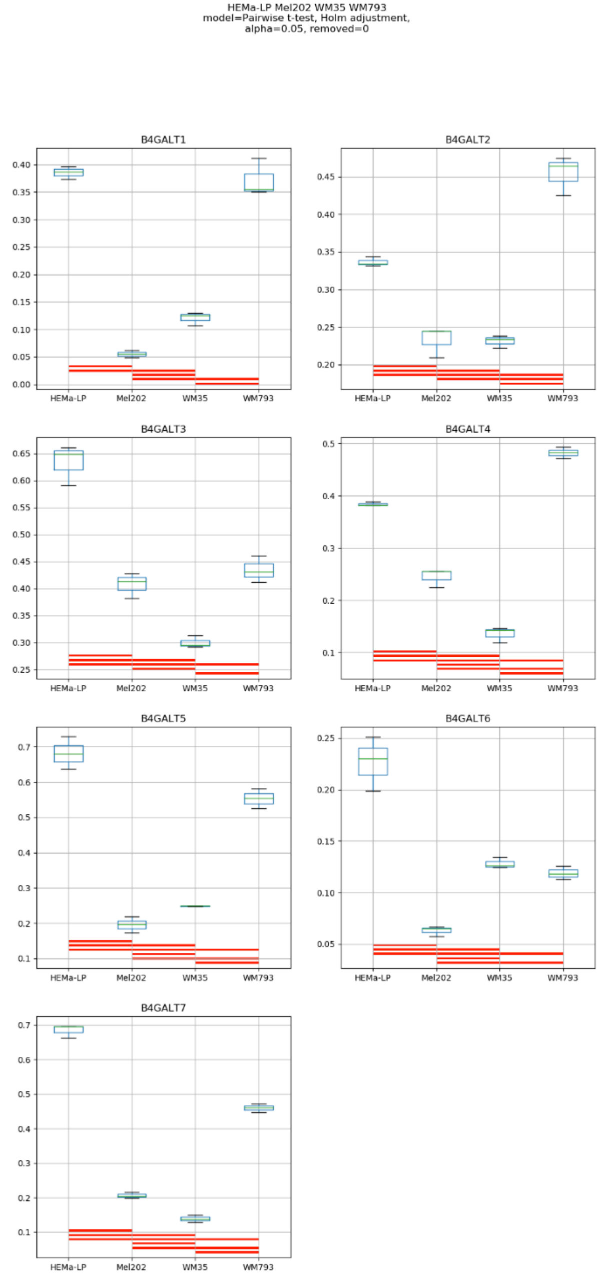
HEMa-LP Mel202 WM35 WM793
model=Pairwise t-test, Holm adjustment,
alpha=0.05, removed=0
B4GALT1
B4GALT2
B4GALT3
B4GALT4
B4GALT5
B4GALT6
B4GALT7
HEMa-LP
Mel202
WM35
WM793

Model No. 22

HEMa-LP Mel202 WM35 WM266-4
model=Pairwise t-test, Holm adjustment,
alpha=0.05, removed=0

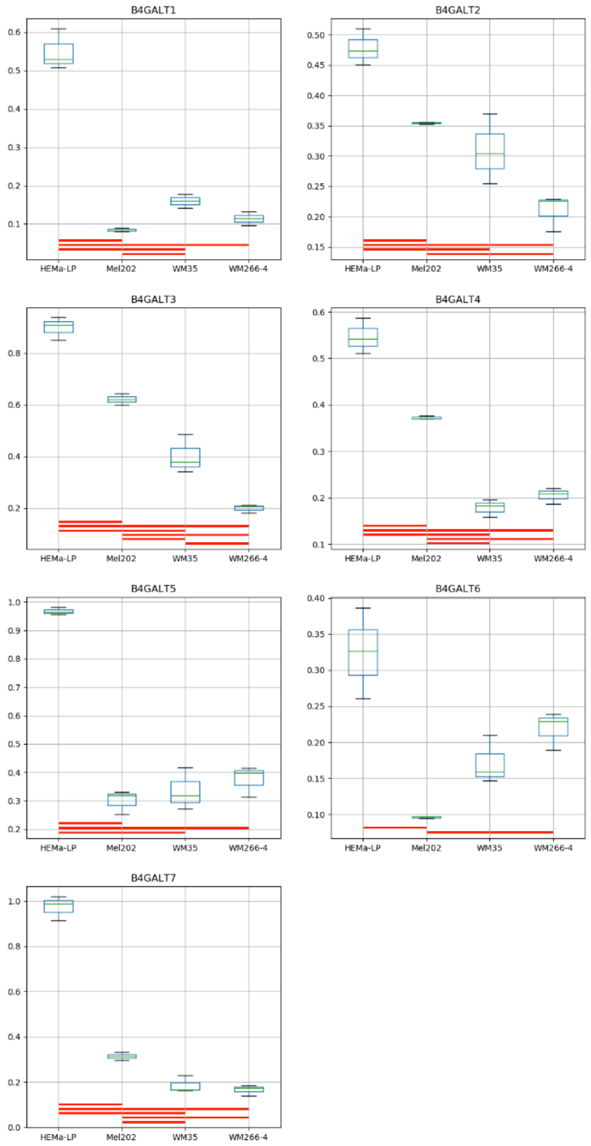

### Model No. 23

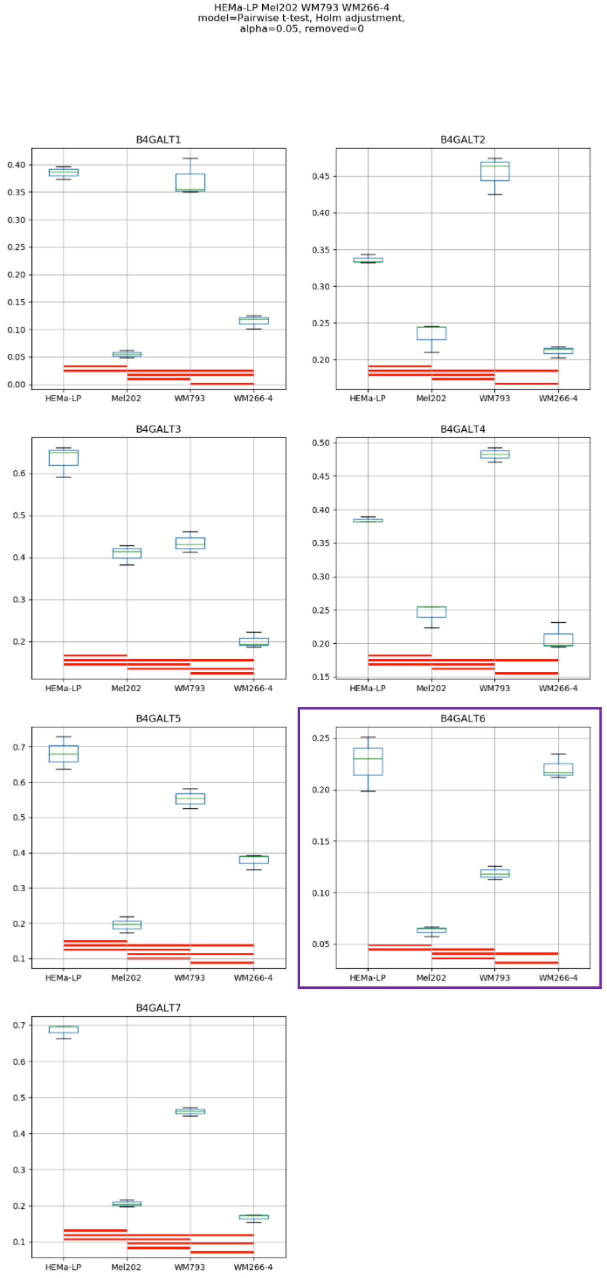
HEMa-LP Mel202 WM793 WM266-4
model=Pairwise t-test, Holm adjustment,
alpha=0.05, removed=0
B4GALT1
B4GALT2
B4GALT3
B4GALT4
B4GALT5
B4GALT6
B4GALT7
HEMa-LP
Mel202
WM793
WM266-4

Model No. 24

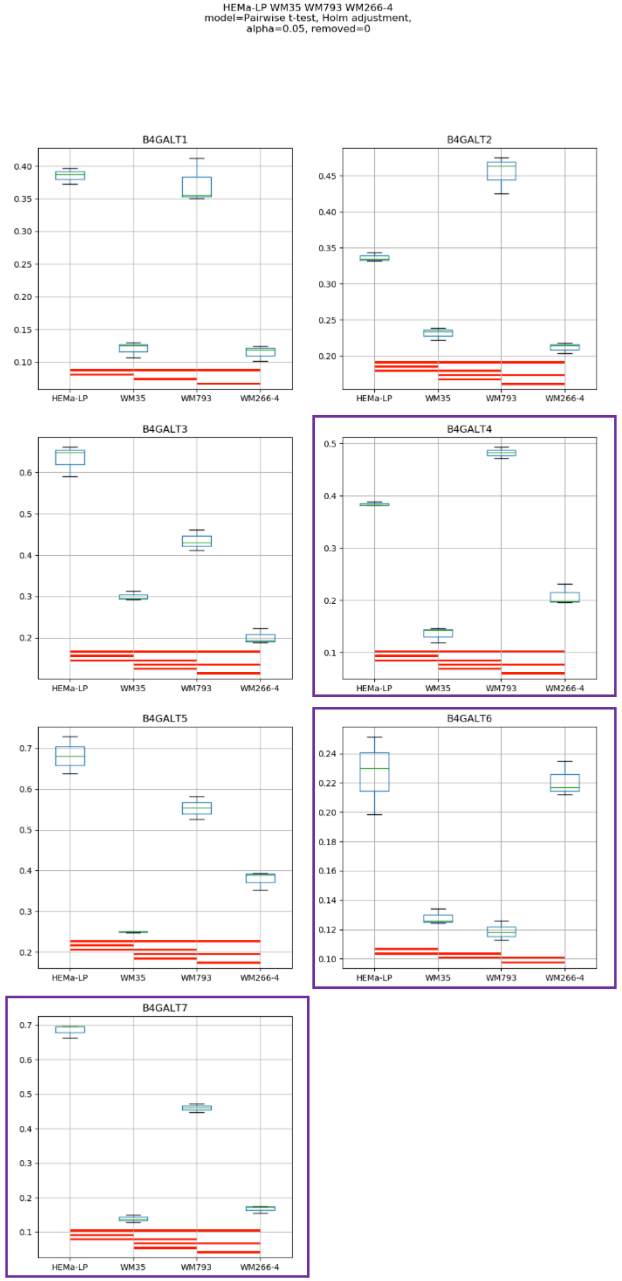
HEMa-LP WM35 WM793 WM266-4
model=Pairwise t-test, Holm adjustment,
alpha=0.05, removed=0
B4GALT1
B4GALT2
B4GALT3
B4GALT4
B4GALT5
B4GALT6
B4GALT7
HEMa-LP
WM35
WM793
WM266-4

Model No. 25

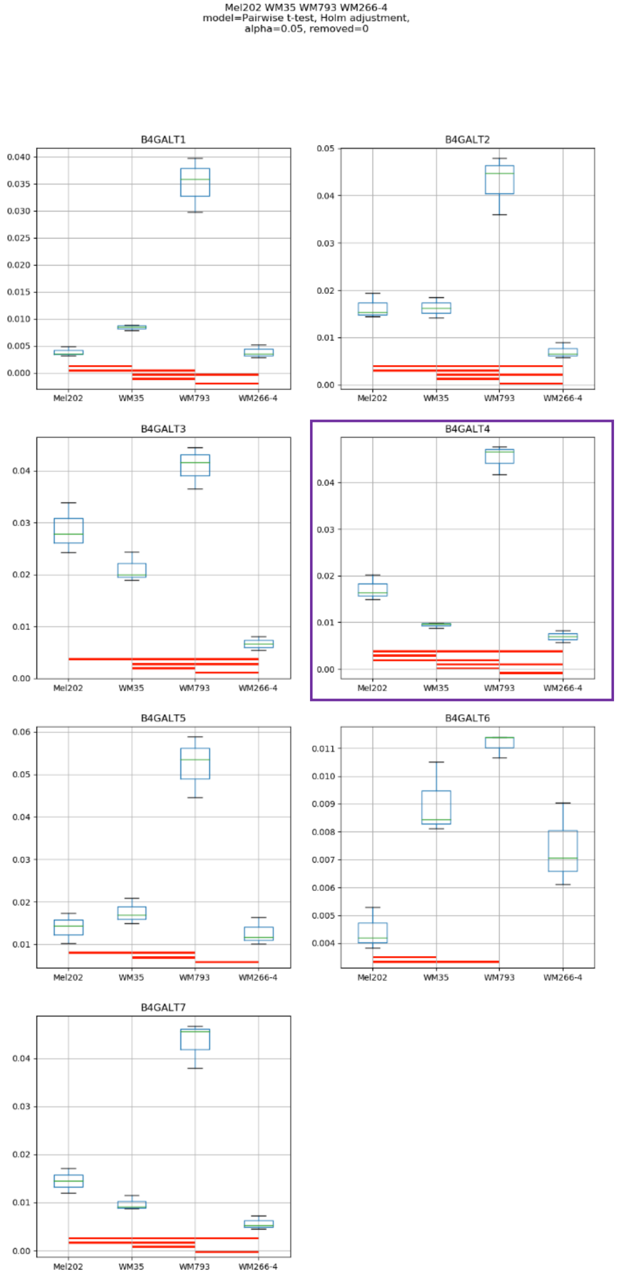
Mel202 WM35 WM793 WM266-4
model=Pairwise t-test, Holm adjustment,
alpha=0.05, removed=0
B4GALT1
B4GALT2
B4GALT3
B4GALT4
B4GALT5
B4GALT6
B4GALT7
Mel202
WM35
WM793
WM266-4

### Model No. 26

HEMa-LP Mel202 WM35 WM793 WM266-4
model=Pairwise t-test, Holm adjustment,
alpha=0.05, removed=0

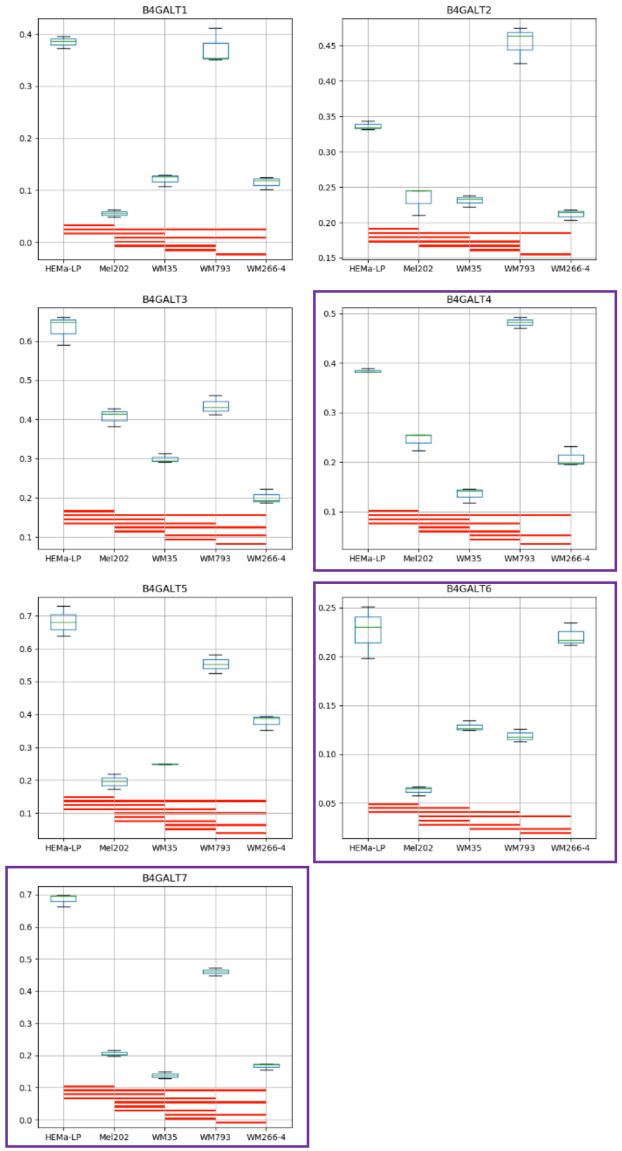

**Supplementary Figure S6:** Medians of relative quantity (RQ) for target genes *B4GALT1–B4GALT7* normalized to the reference gene or pair of reference genes with the best stability values resulting from GenExpA analysis based on set of five candidate reference genes (*PGK1*, *RPS23*, *SNRPA*, *HPRT1*, *GUSB*) and remove repetition level 0 (see Table 1, part B). The statistical analyses used the pairwise t-test with Holm adjustment. Red line represents statistical significance, $P < 0.05$. Frames denote box-plots representing the medians of the RQ values for a given target gene without a repeatable pattern in the same cell lines under different models (the 'Remove repetitions' box set to 0). Detailed results are listed in Supplementary Table 7.

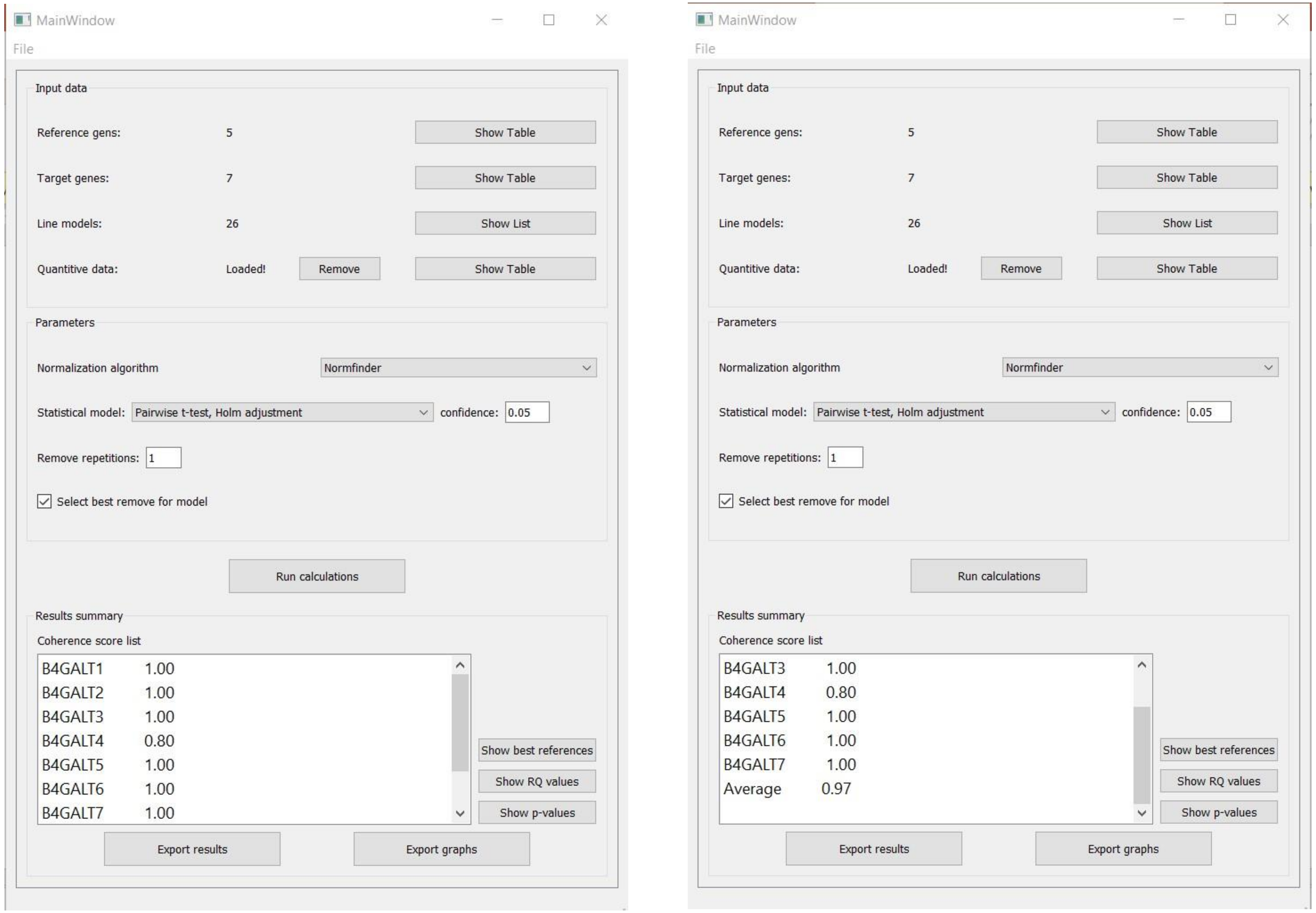


**Supplementary Figure S7:** Print screen of a GenExpA main window with selected parameters of the analysis and coherence score (CS) values for target genes *B4GALT1–B4GALT7* and average CS value of the analysis.

### Model No. 1

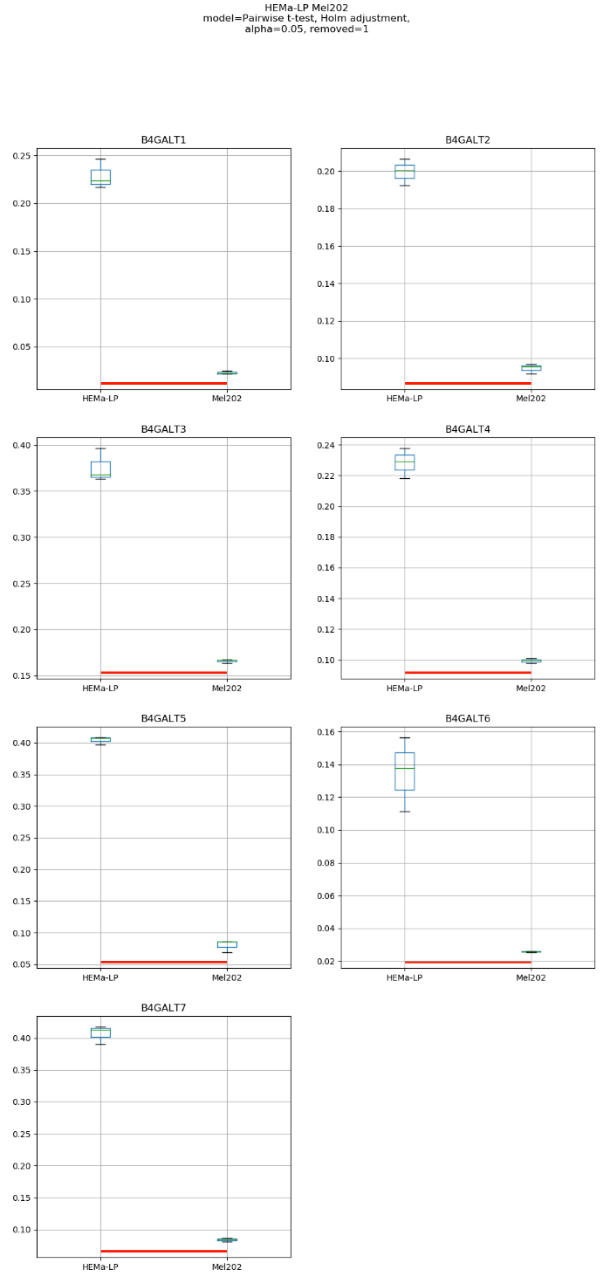
HEMa-LP Mel202
model=Pairwise t-test, Holm adjustment,
alpha=0.05, removed=1
B4GALT1
B4GALT2
B4GALT3
B4GALT4
B4GALT5
B4GALT6
B4GALT7
HEMa-LP
Mel202

### Model No. 2

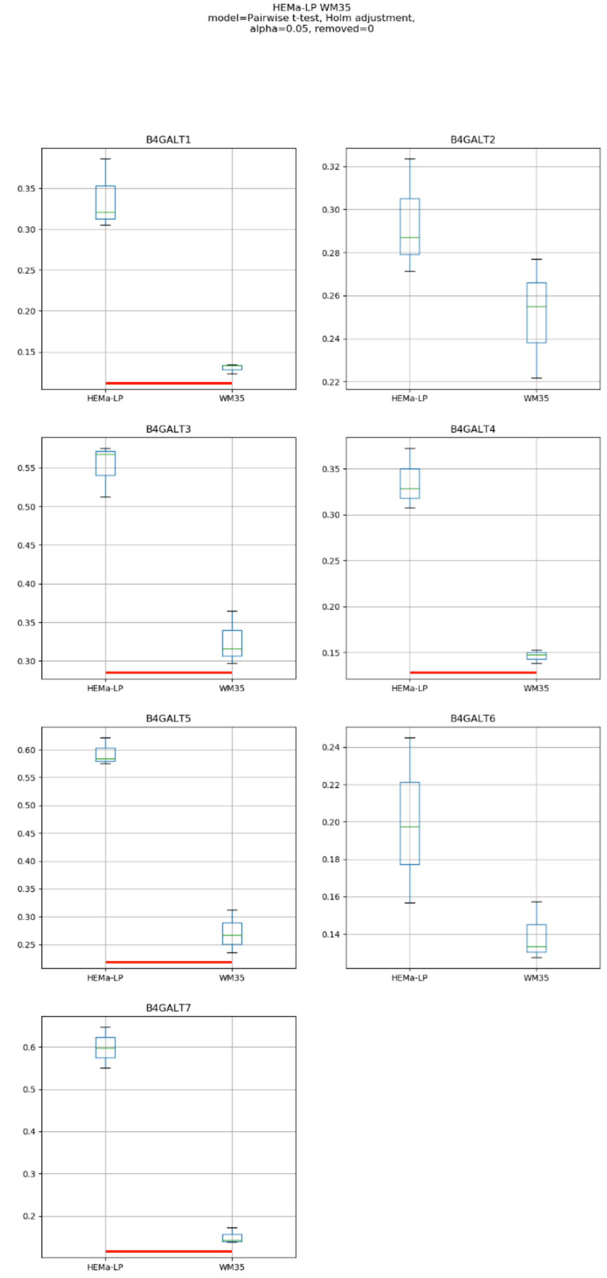
HEMa-LP WM35
model=Pairwise t-test, Holm adjustment,
alpha=0.05, removed=0
B4GALT1
B4GALT2
B4GALT3
B4GALT4
B4GALT5
B4GALT6
B4GALT7
HEMa-LP
WM35

### Model No. 3

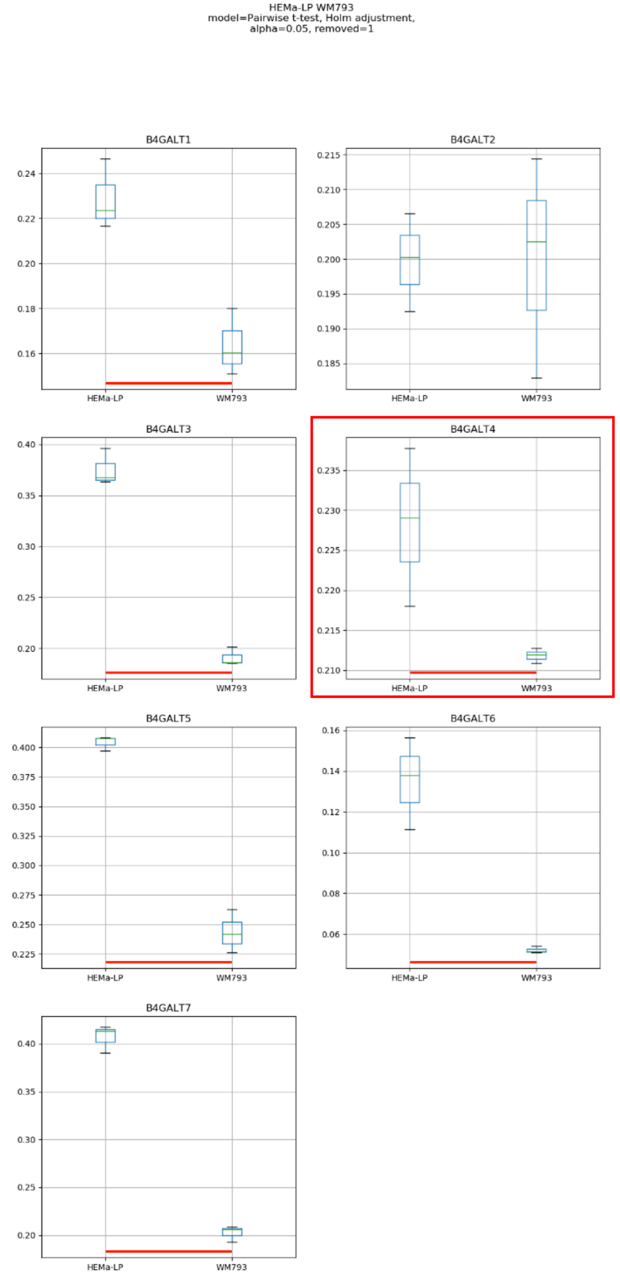
HEMa-LP WM793
model=Pairwise t-test, Holm adjustment,
alpha=0.05, removed=1
B4GALT1
B4GALT2
B4GALT3
B4GALT4
B4GALT5
B4GALT6
B4GALT7
HEMa-LP
WM793

### Model No. 4

HEMa-LP WM266-4
model=Pairwise t-test, Holm adjustment,
alpha=0.05, removed=1

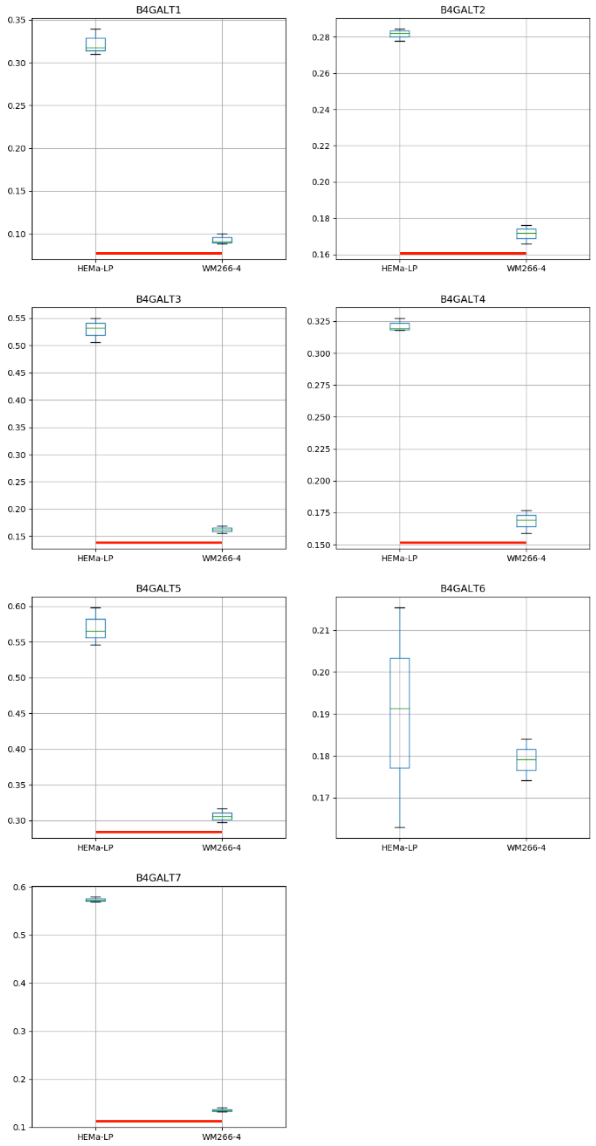

### Model No. 5

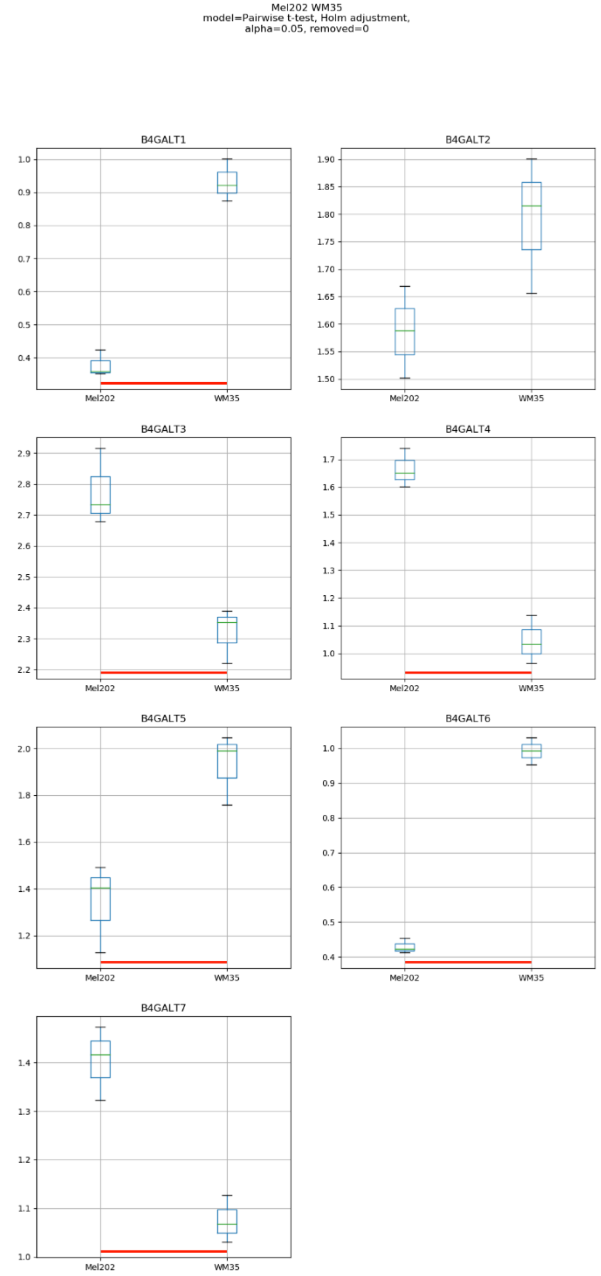
Mel202 WM35
model=Pairwise t-test, Holm adjustment,
alpha=0.05, removed=0
B4GALT1
B4GALT2
B4GALT3
B4GALT4
B4GALT5
B4GALT6
B4GALT7
Mel202
WM35

### Model No. 6

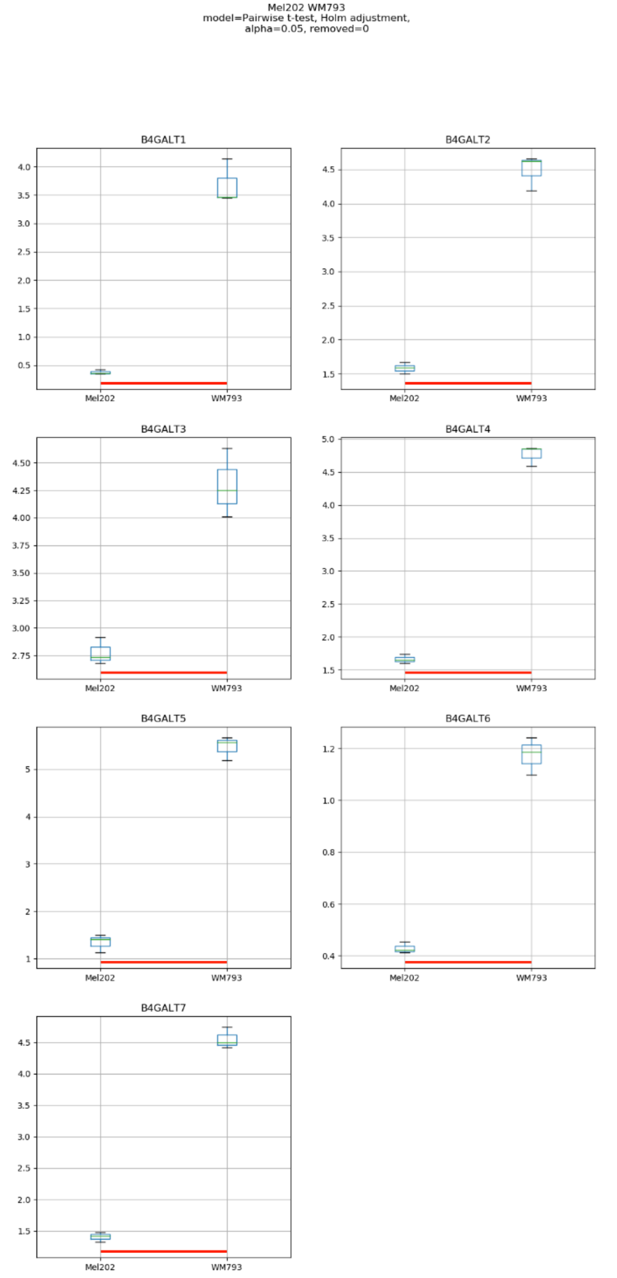
Mel202 WM793
model=Pairwise t-test, Holm adjustment,
alpha=0.05, removed=0
B4GALT1
B4GALT2
B4GALT3
B4GALT4
B4GALT5
B4GALT6
B4GALT7
Mel202
WM793

### Model No. 7

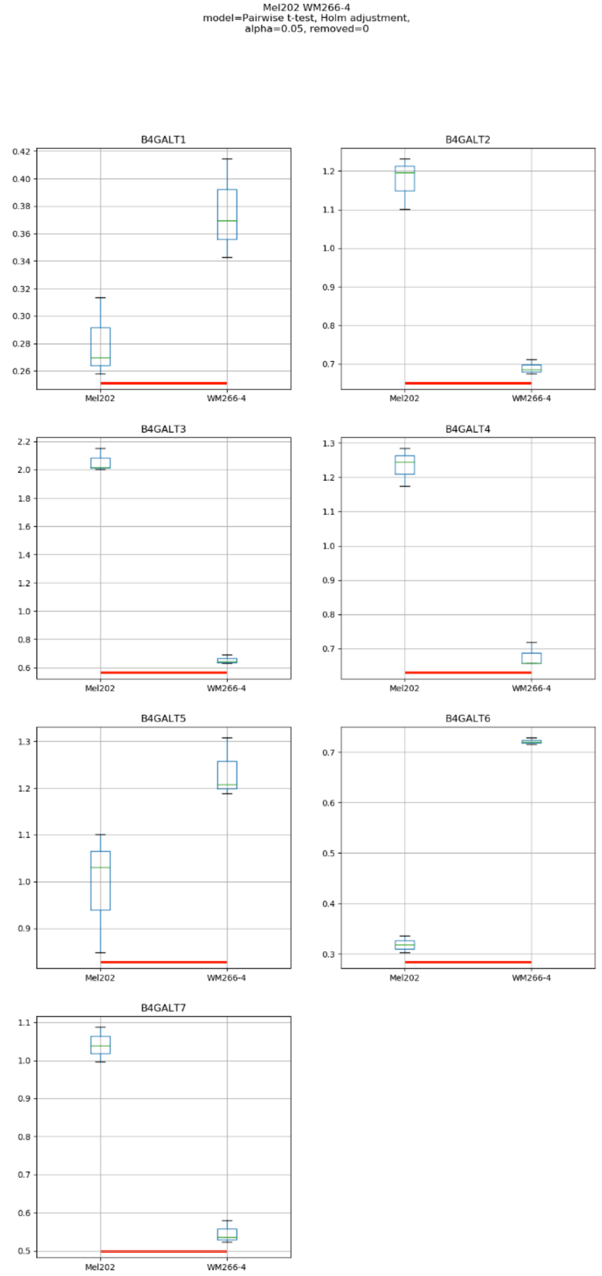
Mel202 WM266-4
model=Pairwise t-test, Holm adjustment,
alpha=0.05, removed=0
B4GALT1
B4GALT2
B4GALT3
B4GALT4
B4GALT5
B4GALT6
B4GALT7
Mel202
WM266-4

### Model No. 8

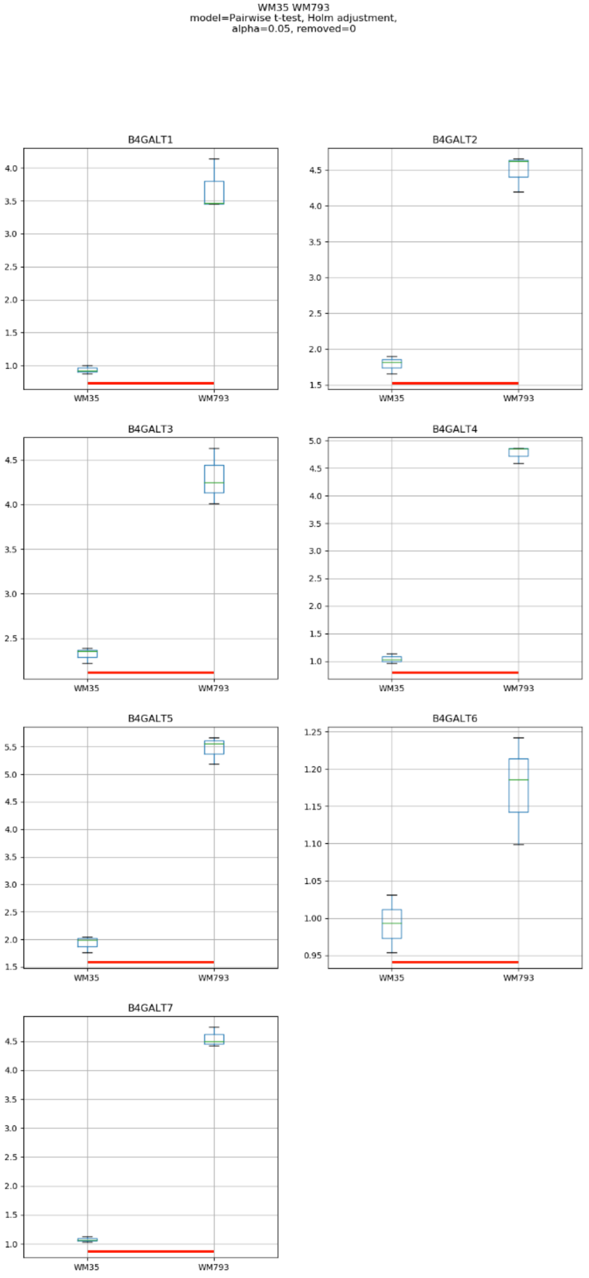

WM35 WM793
model=Pairwise t-test, Holm adjustment,
alpha=0.05, removed=0
B4GALT1
B4GALT2
B4GALT3
B4GALT4
B4GALT5
B4GALT6
B4GALT7
WM35
WM793

### Model No. 9

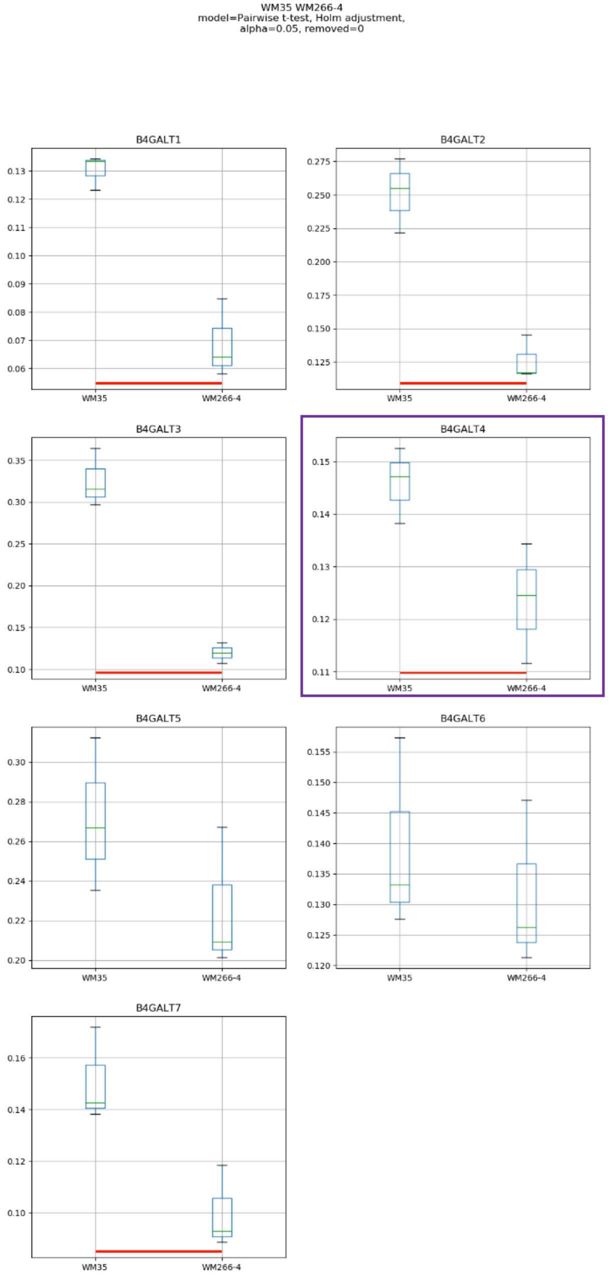
WM35 WM266-4
model=Pairwise t-test, Holm adjustment,
alpha=0.05, removed=0
B4GALT1
B4GALT2
B4GALT3
B4GALT4
B4GALT5
B4GALT6
B4GALT7
WM35
WM266-4

### Model No. 10

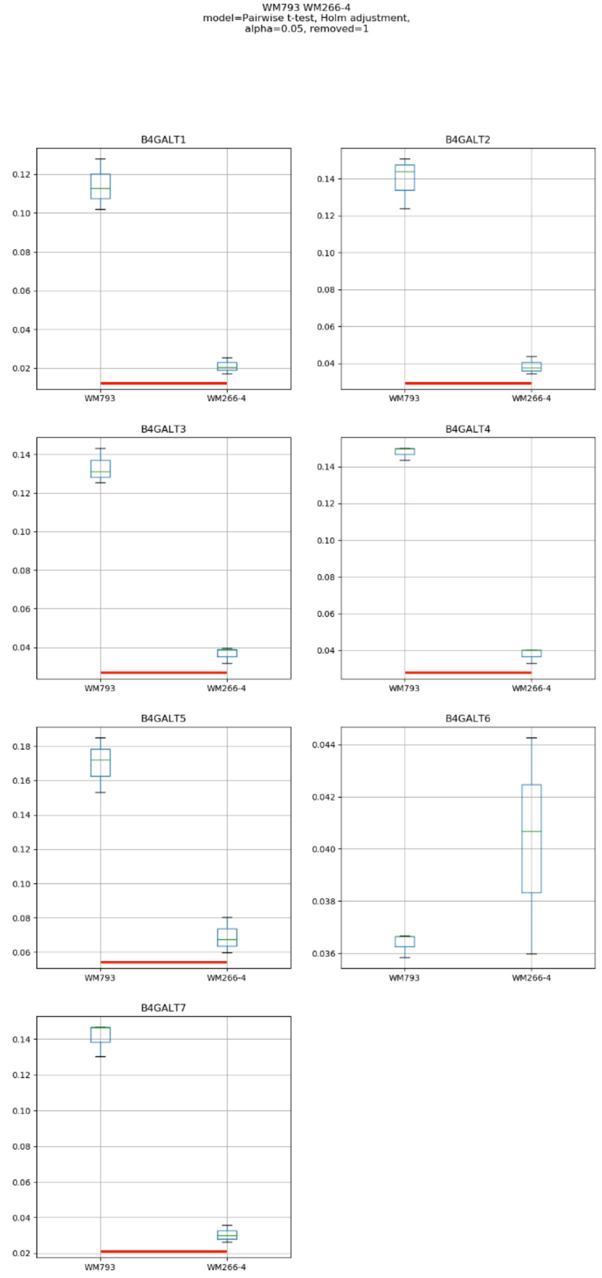
WM793 WM266-4
model=Pairwise t-test, Holm adjustment,
alpha=0.05, removed=1
B4GALT1
B4GALT2
B4GALT3
B4GALT4
B4GALT5
B4GALT6
B4GALT7
WM793
WM266-4

Model No. 11

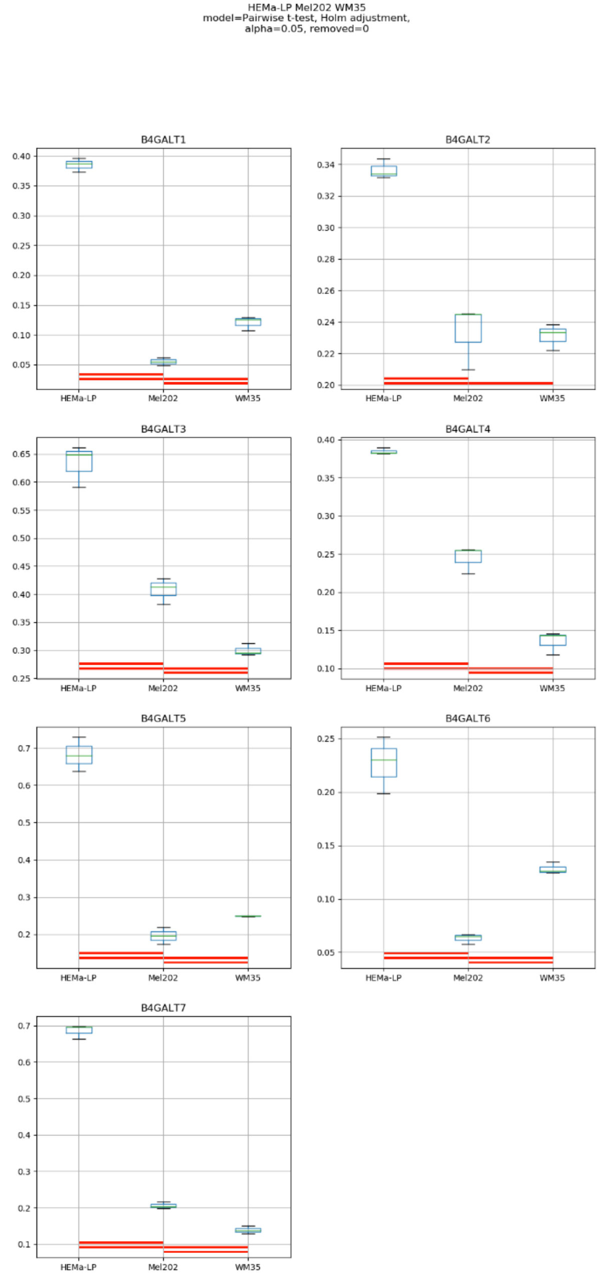
HEMa-LP Mel202 WM35
model=Pairwise t-test, Holm adjustment,
alpha=0.05, removed=0
B4GALT1
B4GALT2
B4GALT3
B4GALT4
B4GALT5
B4GALT6
B4GALT7
HEMa-LP
Mel202
WM35

### Model No. 12

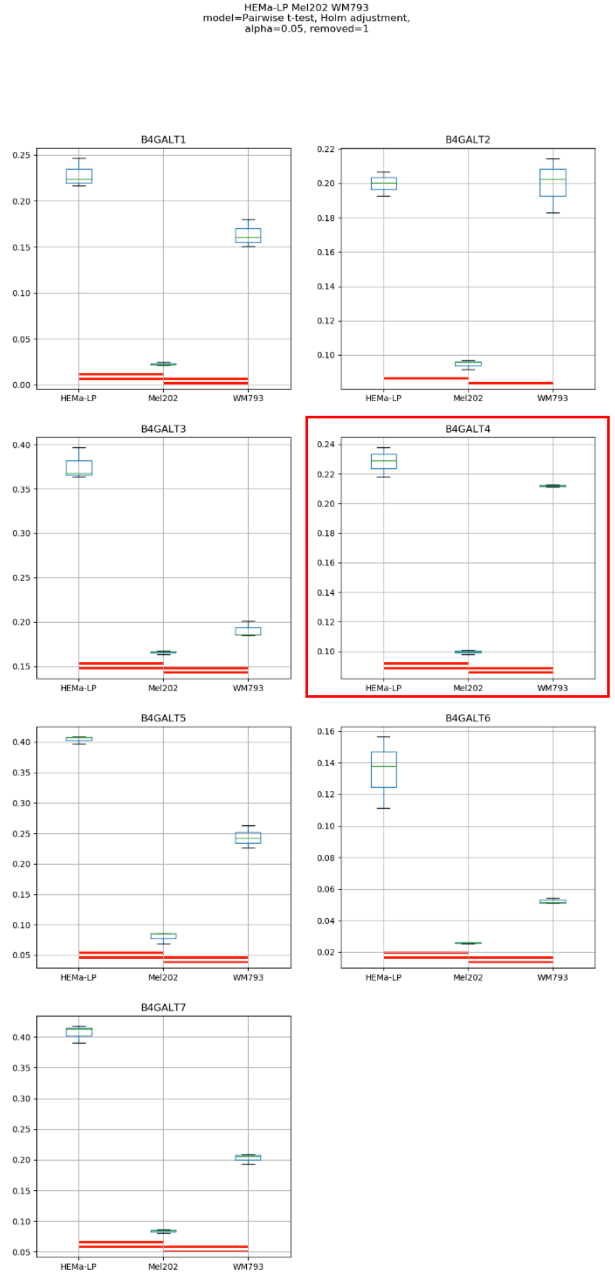
HEMa-LP Mel202 WM793
model=Pairwise t-test, Holm adjustment,
alpha=0.05, removed=1
B4GALT1
B4GALT2
B4GALT3
B4GALT4
B4GALT5
B4GALT6
B4GALT7
HEMa-LP
Mel202
WM793

### Model No. 13

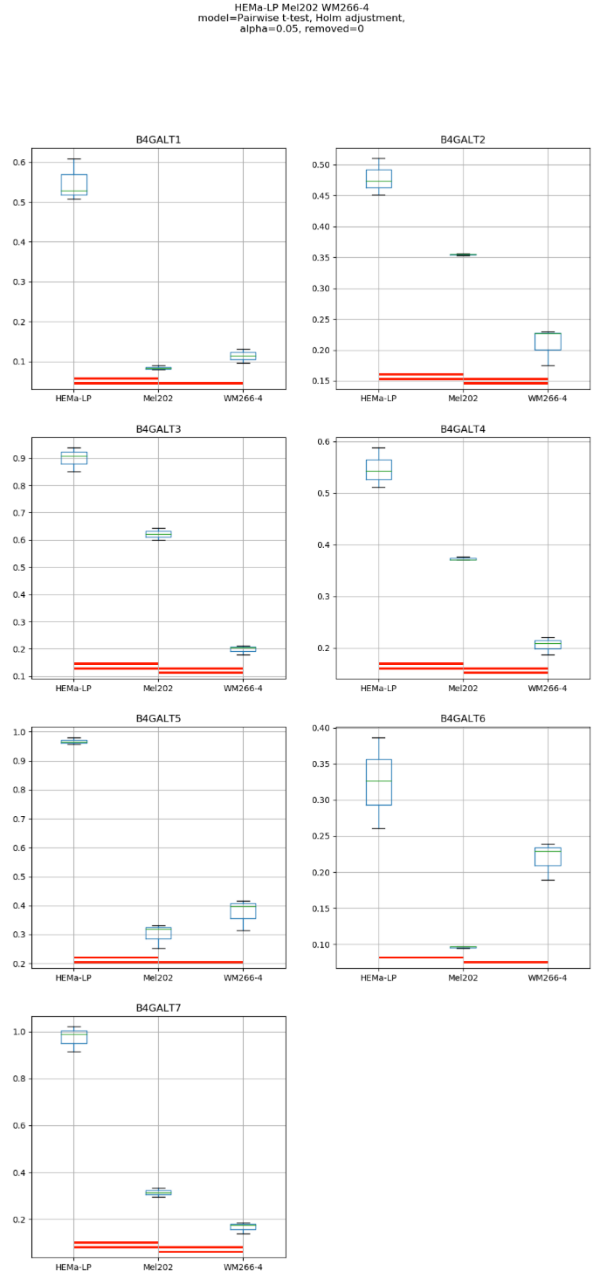

HEMa-LP Mel202 WM266-4
model=Pairwise t-test, Holm adjustment,
alpha=0.05, removed=0
B4GALT1
B4GALT2
B4GALT3
B4GALT4
B4GALT5
B4GALT6
B4GALT7
HEMa-LP
Mel202
WM266-4

Model No. 14

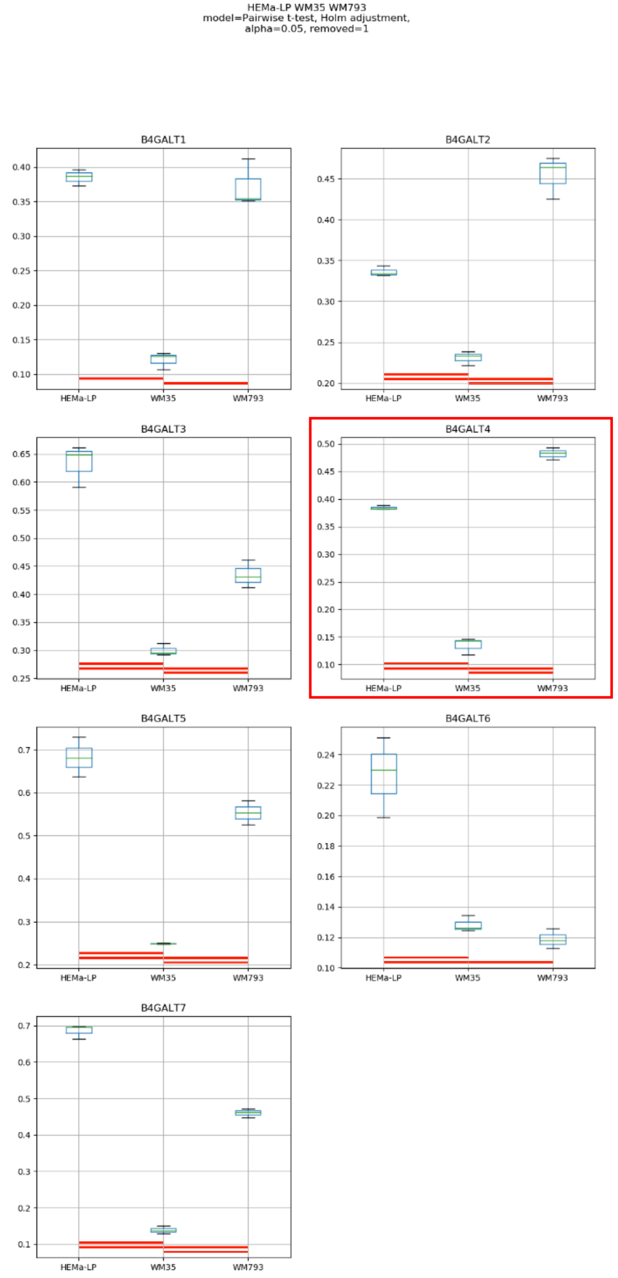
HEMa-LP WM35 WM793
model=Pairwise t-test, Holm adjustment,
alpha=0.05, removed=1
B4GALT1
B4GALT2
B4GALT3
B4GALT4
B4GALT5
B4GALT6
B4GALT7
HEMa-LP
WM35
WM793

### Model No. 15

HEMa-LP WM35 WM266-4
model=Pairwise t-test, Holm adjustment,
alpha=0.05, removed=0

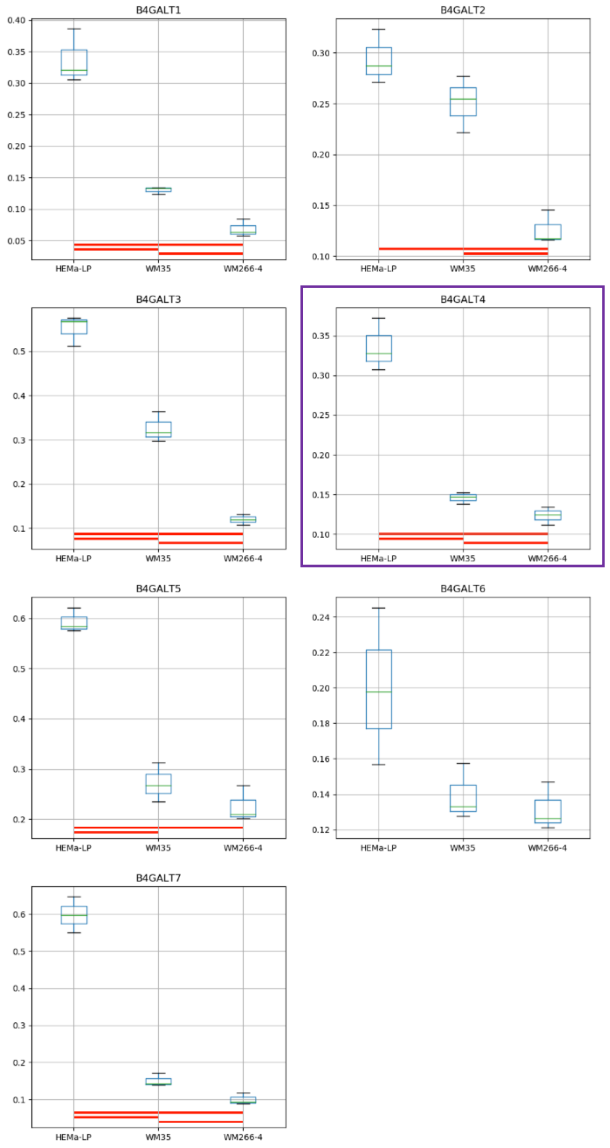

### Model No. 16

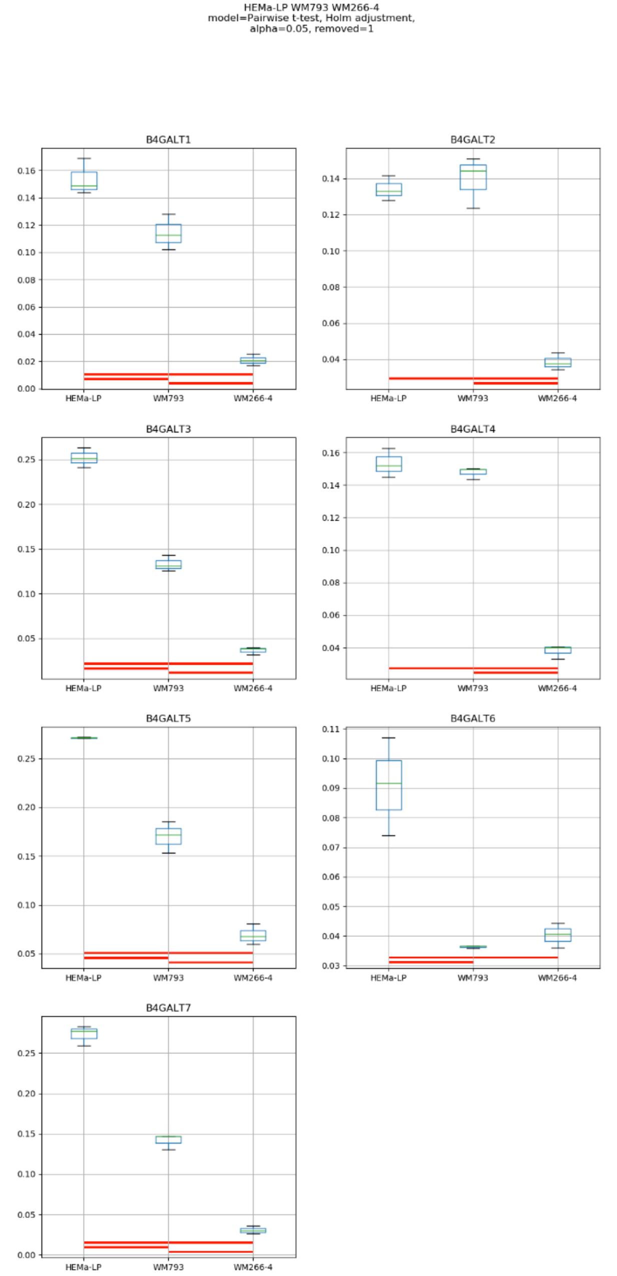
HEMa-LP WM793 WM266-4
model=Pairwise t-test, Holm adjustment,
alpha=0.05, removed=1
B4GALT1
B4GALT2
B4GALT3
B4GALT4
B4GALT5
B4GALT6
B4GALT7
HEMa-LP
WM793
WM266-4

Model No. 17

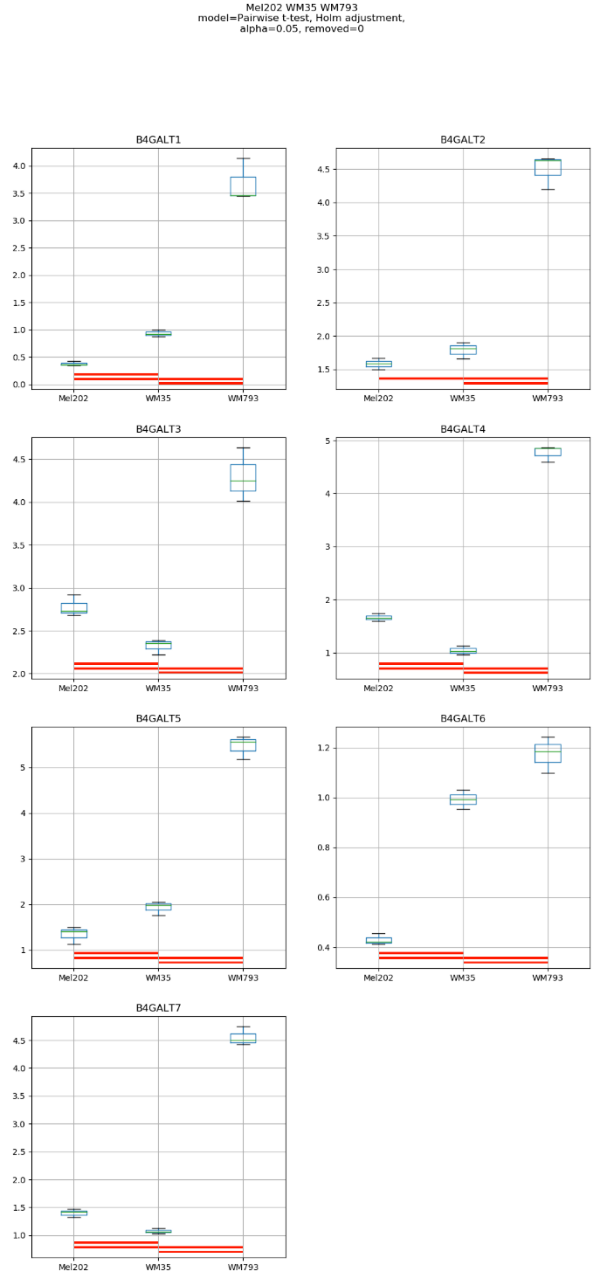
Mel202 WM35 WM793
model=Pairwise t-test, Holm adjustment,
alpha=0.05, removed=0
B4GALT1
B4GALT2
B4GALT3
B4GALT4
B4GALT5
B4GALT6
B4GALT7
Mel202
WM35
WM793

### Model No. 18

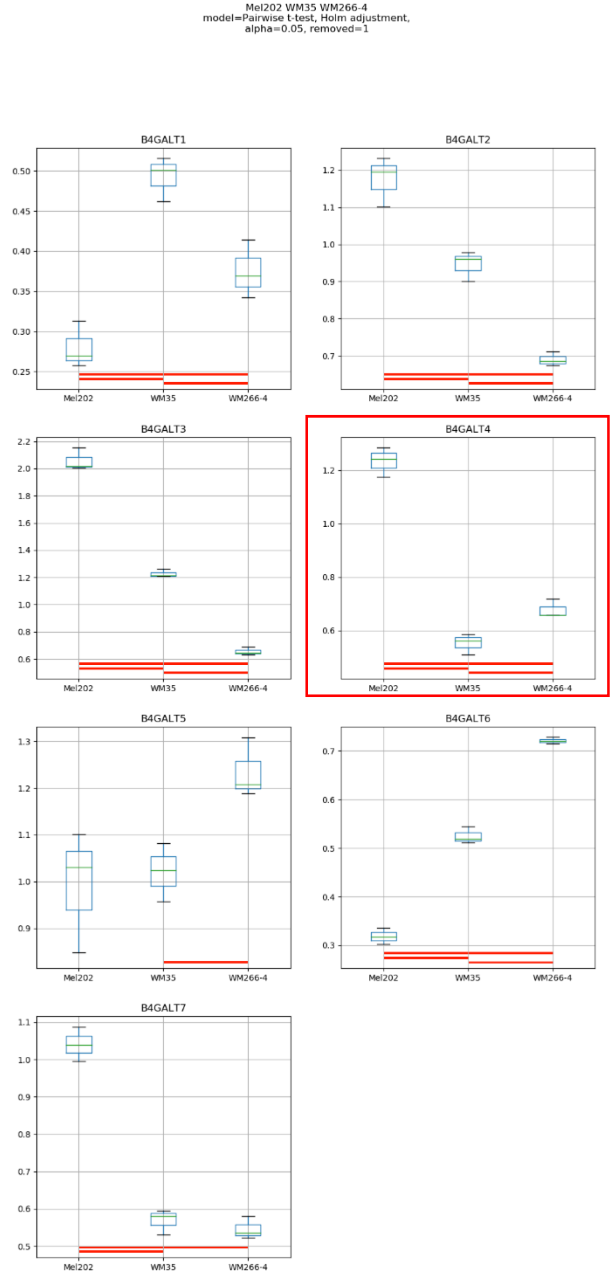
Mel202 WM35 WM266-4
model=Pairwise t-test, Holm adjustment,
alpha=0.05, removed=1
B4GALT1
B4GALT2
B4GALT3
B4GALT4
B4GALT5
B4GALT6
B4GALT7
Mel202
WM35
WM266-4

Model No. 19

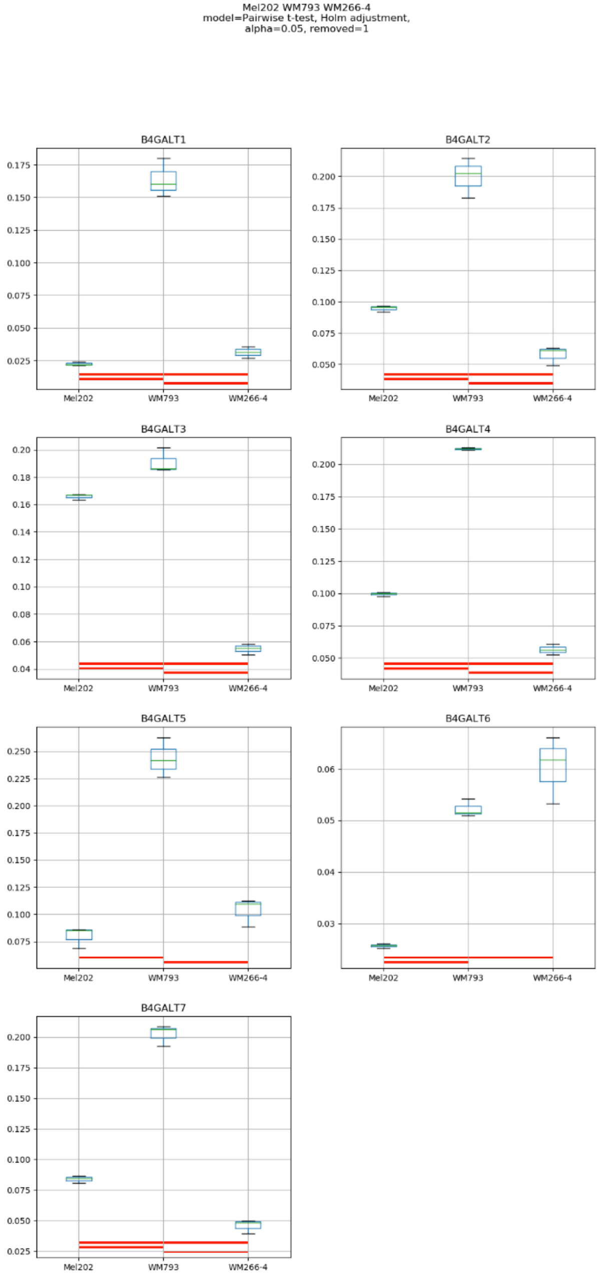
Mel202 WM793 WM266-4
model=Pairwise t-test, Holm adjustment,
alpha=0.05, removed=1
B4GALT1
B4GALT2
B4GALT3
B4GALT4
B4GALT5
B4GALT6
B4GALT7
Mel202
WM793
WM266-4

### Model No. 20

WM35 WM793 WM266-4
model=Pairwise t-test, Holm adjustment,
alpha=0.05, removed=0

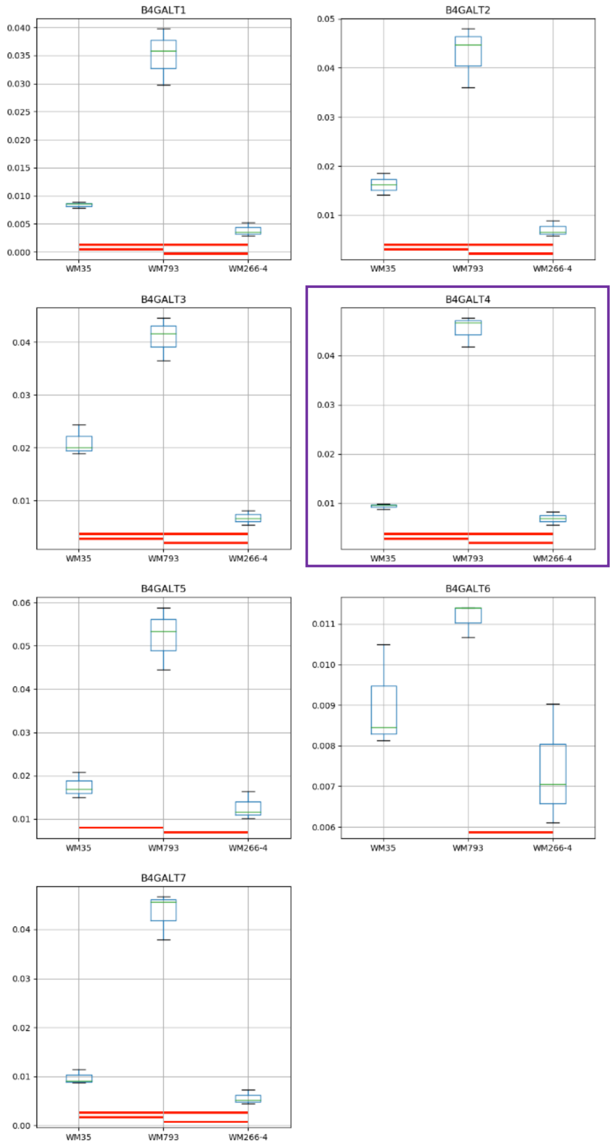

### Model No. 21

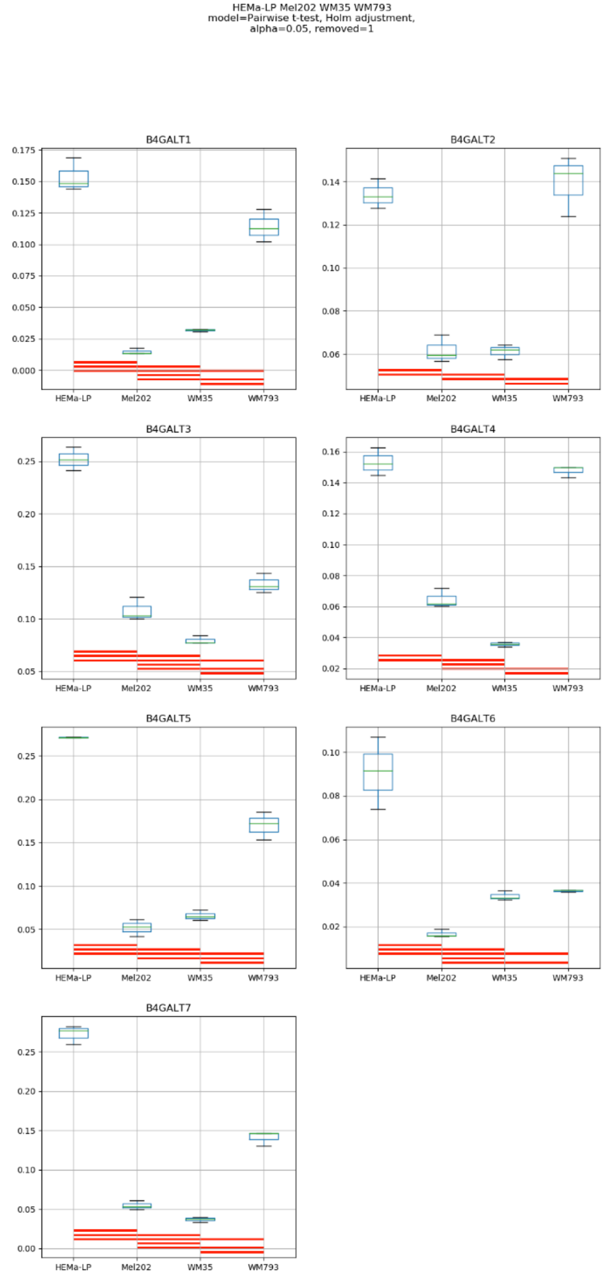
HEMa-LP Mel202 WM35 WM793
model=Pairwise t-test, Holm adjustment,
alpha=0.05, removed=1
B4GALT1
B4GALT2
B4GALT3
B4GALT4
B4GALT5
B4GALT6
B4GALT7
HEMa-LP
Mel202
WM35
WM793

### Model No. 22

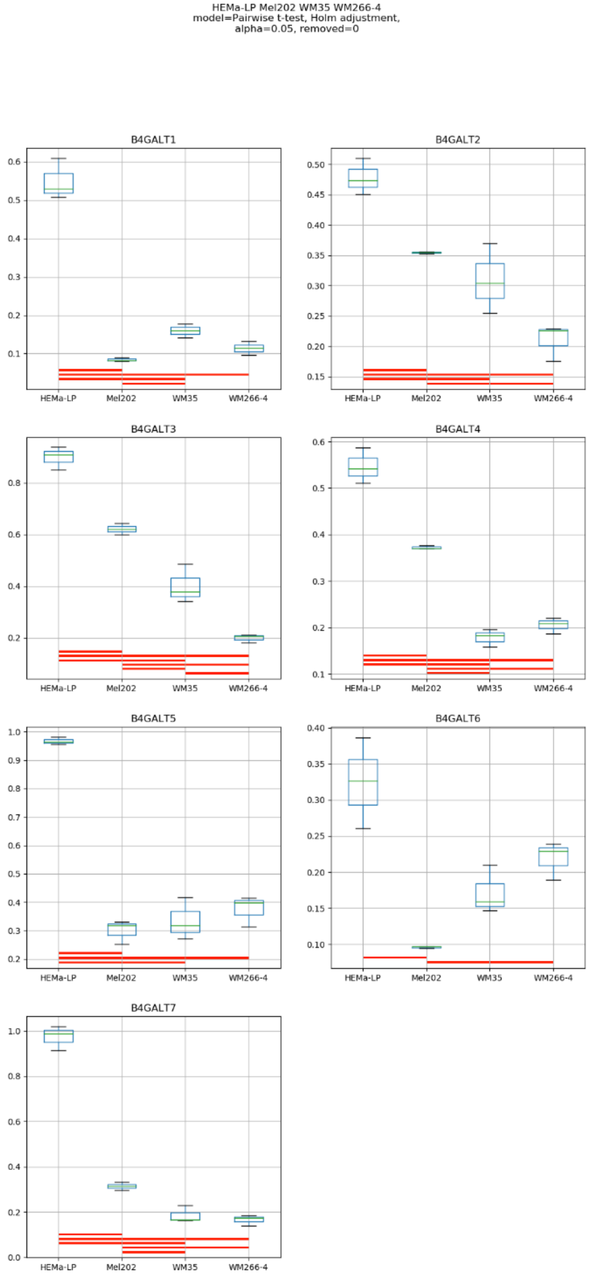
HEMa-LP Mel202 WM35 WM266-4
model=Pairwise t-test, Holm adjustment,
alpha=0.05, removed=0
B4GALT1
B4GALT2
B4GALT3
B4GALT4
B4GALT5
B4GALT6
B4GALT7
HEMa-LP
Mel202
WM35
WM266-4

### Model No. 23

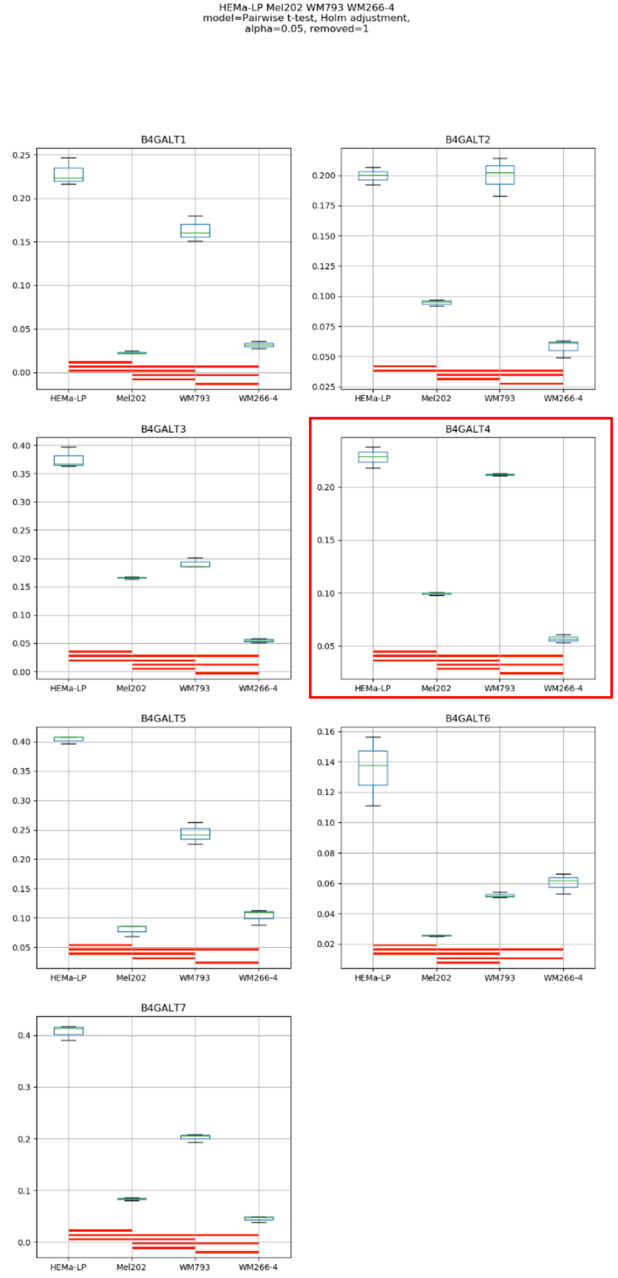

HEMa-LP Mel202 WM793 WM266-4
model=Pairwise t-test, Holm adjustment,
alpha=0.05, removed=1
B4GALT1
B4GALT2
B4GALT3
B4GALT4
B4GALT5
B4GALT6
B4GALT7
HEMa-LP
Mel202
WM793
WM266-4

Model No. 24

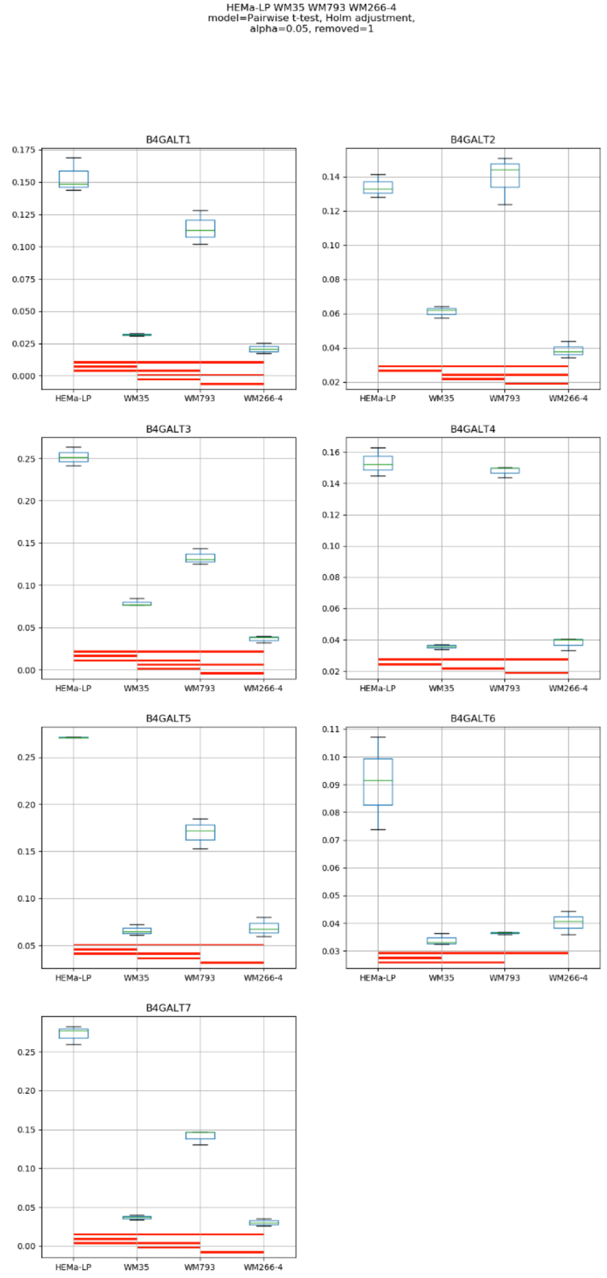
HEMa-LP WM35 WM793 WM266-4
model=Pairwise t-test, Holm adjustment,
alpha=0.05, removed=1
B4GALT1
B4GALT2
B4GALT3
B4GALT4
B4GALT5
B4GALT6
B4GALT7
HEMa-LP
WM35
WM793
WM266-4

### Model No. 25

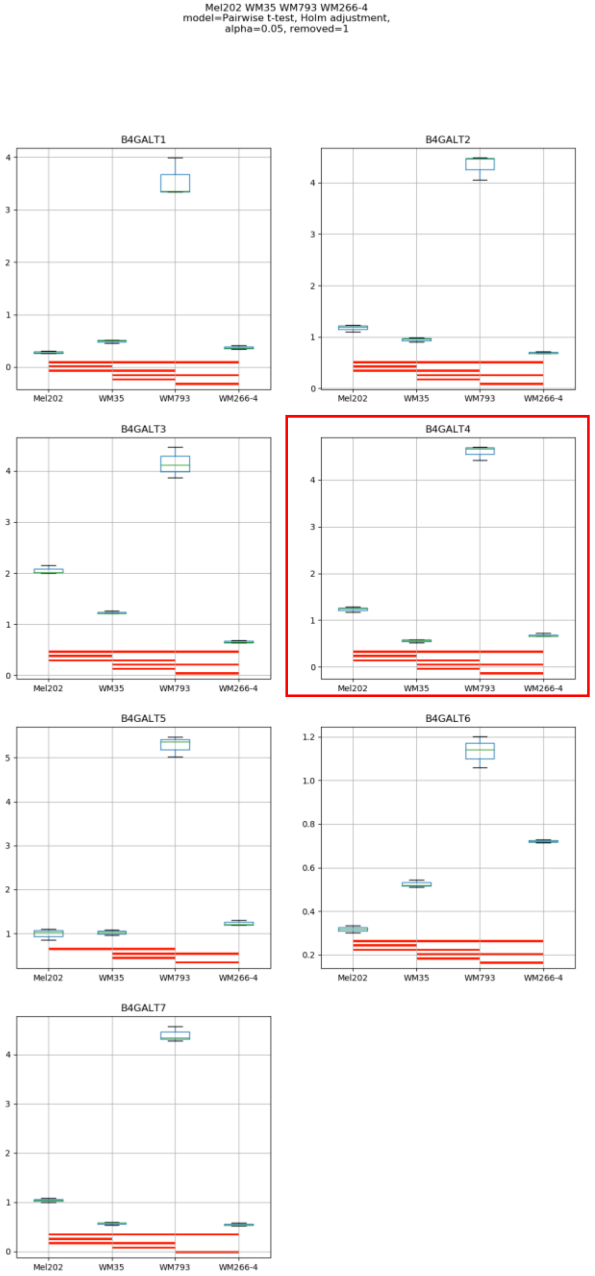
Mel202 WM35 WM793 WM266-4
model=Pairwise t-test, Holm adjustment,
alpha=0.05, removed=1
B4GALT1
B4GALT2
B4GALT3
B4GALT4
B4GALT5
B4GALT6
B4GALT7
Mel202
WM35
WM793
WM266-4

### Model No. 26

HEMa-LP Mel202 WM35 WM793 WM266-4
model=Pairwise t-test, Holm adjustment,
alpha=0.05, removed=1

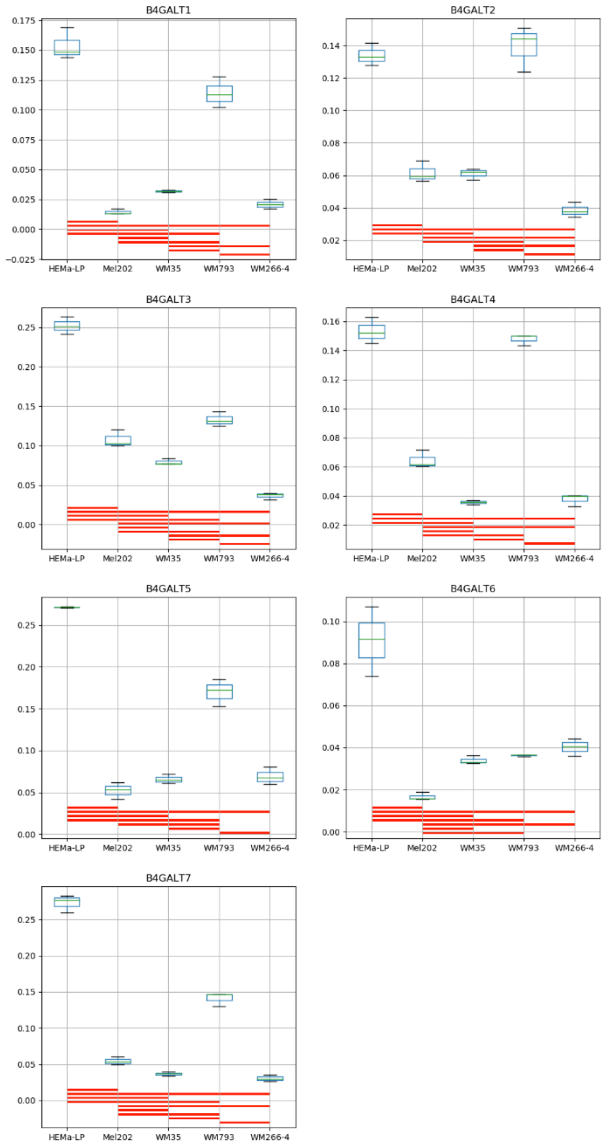

**Supplementary Figure S8:** Medians of relative quantity (RQ) for target genes *B4GALT1–B4GALT7* normalized to the reference gene or pair of reference genes with the best stability values resulting from GenExpA analysis based on set of five candidate reference genes (*PGK1*, *RPS23*, *SNRPA*, *HPRT1*, *GUSB*) and remove repetition level 1 (see Table 1, part B). The statistical analyses used the pairwise t-test with Holm adjustment. Red line represents statistical significance, $P < 0.05$. Frames denote box-plots representing the medians of the RQ values for a given target gene without a repeatable pattern in the same cell lines under different models (the 'Remove repetitions' box set to 0 – violet frame, or to 1 – red frame). Detailed results are listed in Supplementary Table 8.

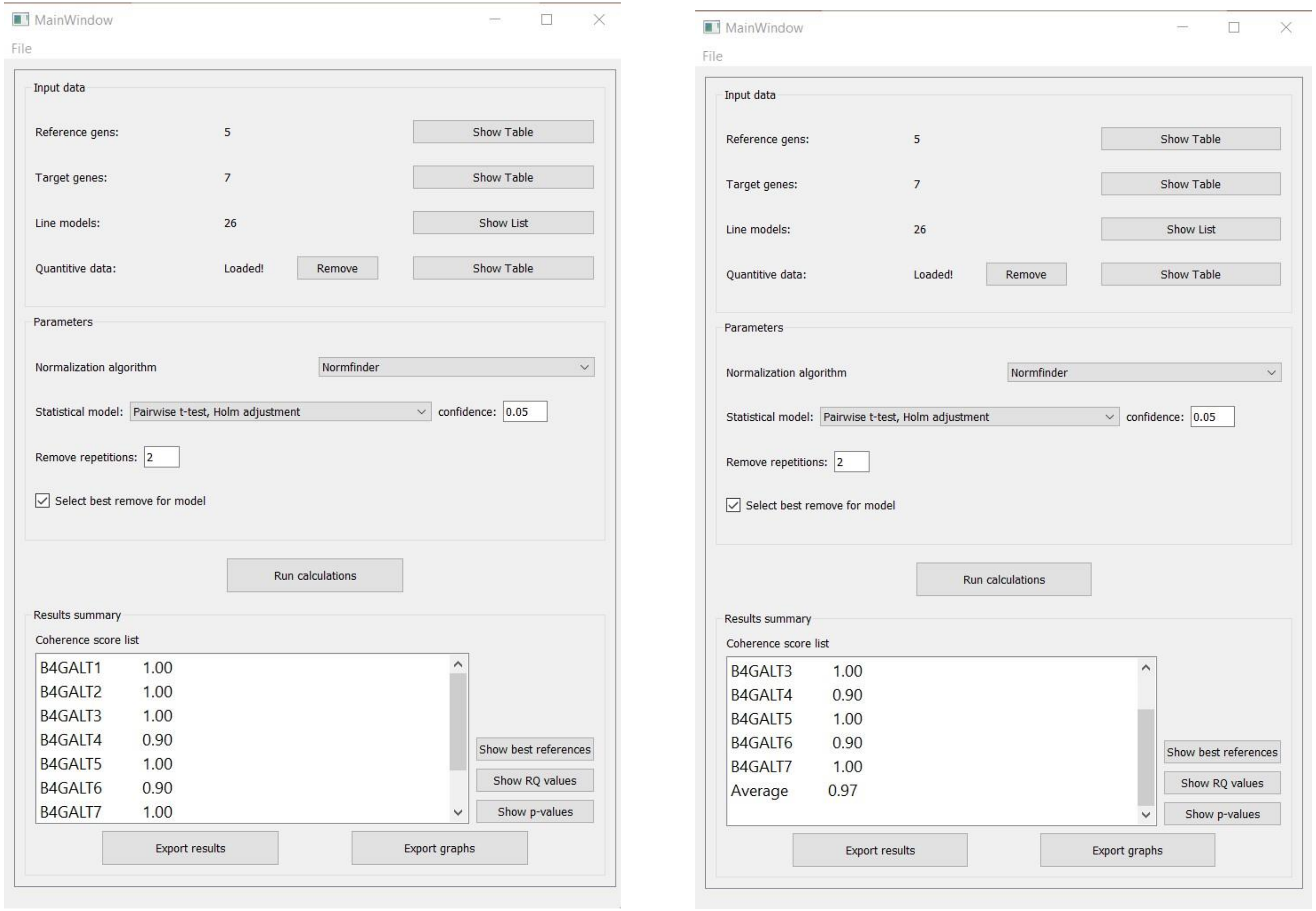


**Supplementary Figure S9:** Print screen of a GenExpA main window with selected parameters of the analysis and coherence score (CS) values for target genes *B4GALT1–B4GALT7* and average CS value of the analysis.

### Model No. 1

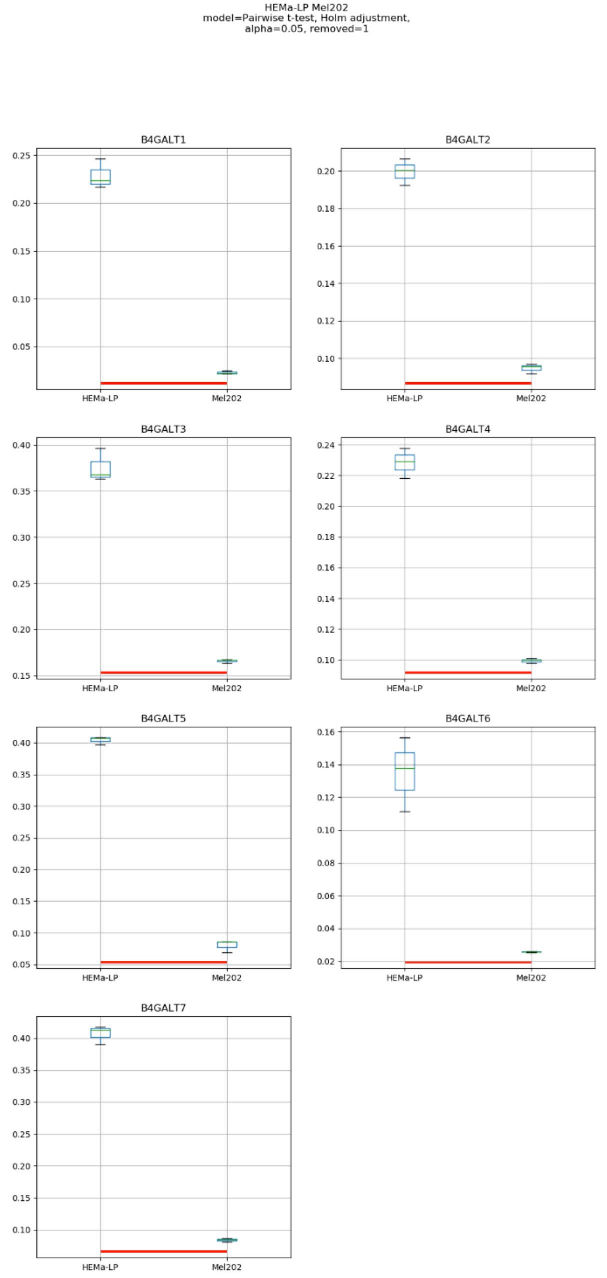
HEMa-LP Mel202
model=Pairwise t-test, Holm adjustment,
alpha=0.05, removed=1
B4GALT1
B4GALT2
B4GALT3
B4GALT4
B4GALT5
B4GALT6
B4GALT7
HEMa-LP
Mel202

### Model No. 2

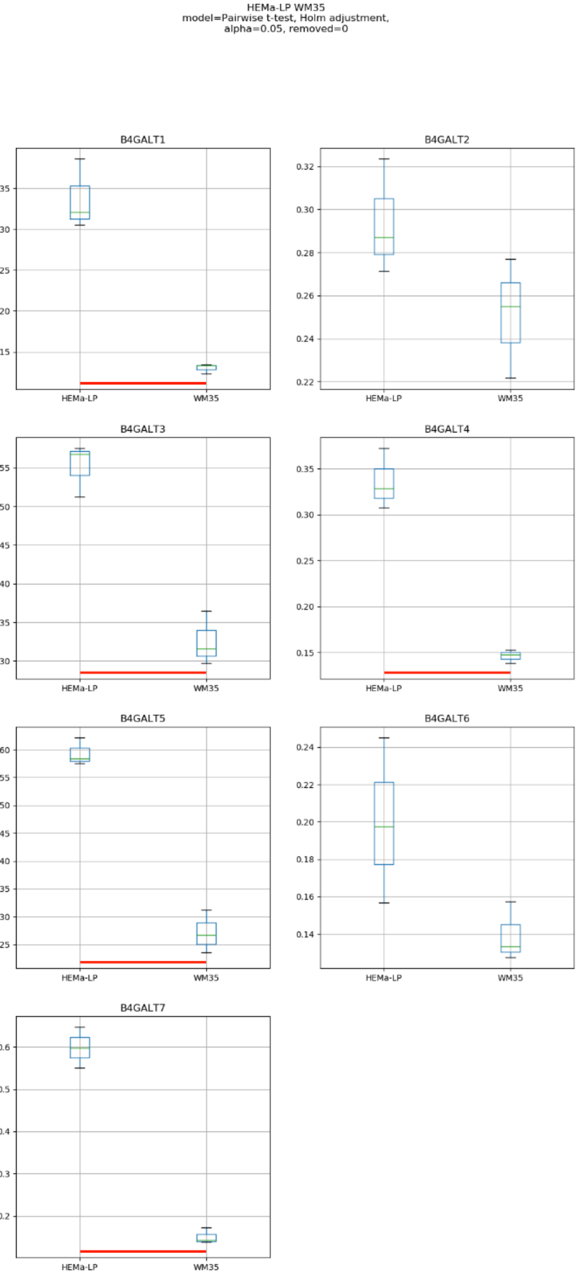
HEMa-LP WM35
model=Pairwise t-test, Holm adjustment,
alpha=0.05, removed=0
B4GALT1
B4GALT2
B4GALT3
B4GALT4
B4GALT5
B4GALT6
B4GALT7
HEMa-LP
WM35

### Model No. 3

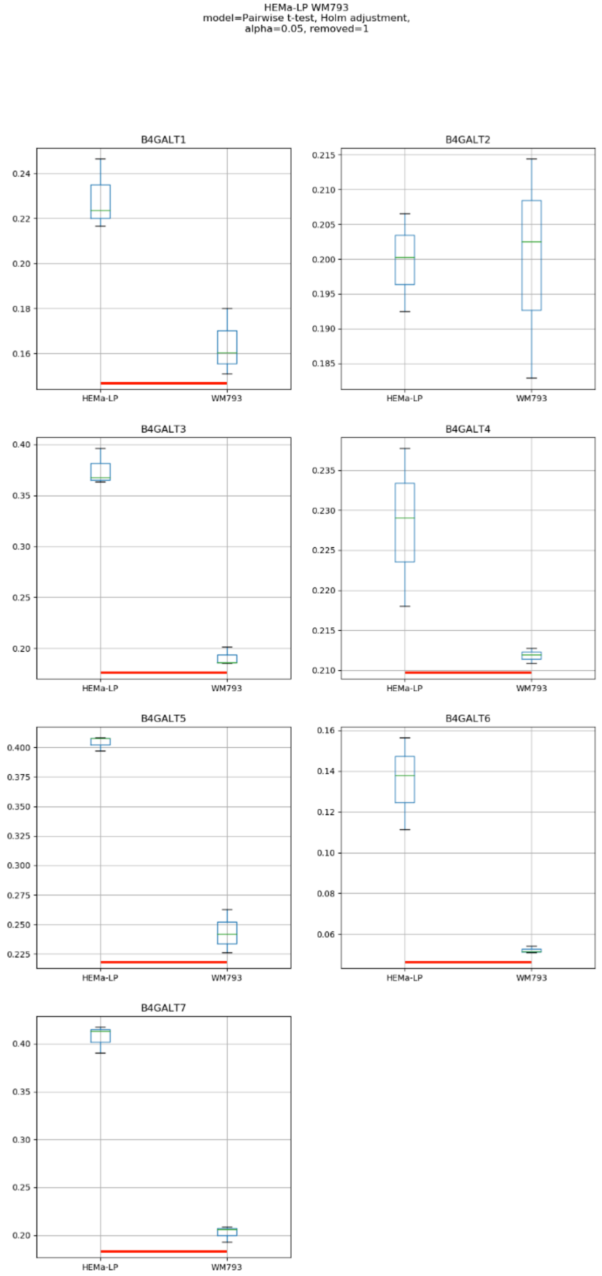
HEMa-LP WM793
model=Pairwise t-test, Holm adjustment,
alpha=0.05, removed=1
B4GALT1
B4GALT2
B4GALT3
B4GALT4
B4GALT5
B4GALT6
B4GALT7
HEMa-LP
WM793

### Model No. 4

HEMa-LP WM266-4
model=Pairwise t-test, Holm adjustment,
alpha=0.05, removed=1

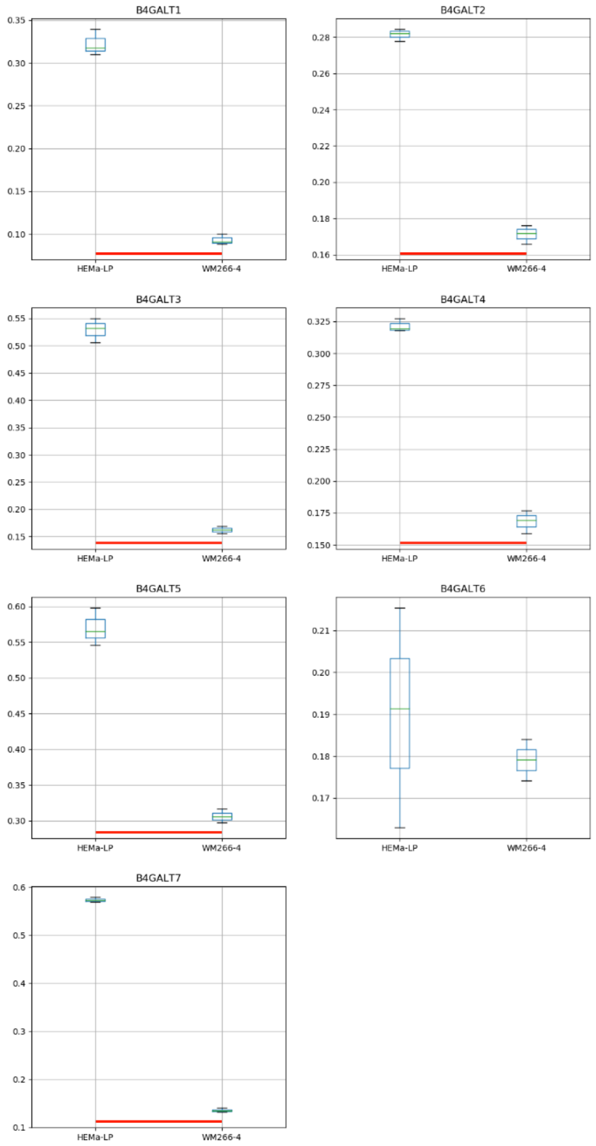

### Model No. 5

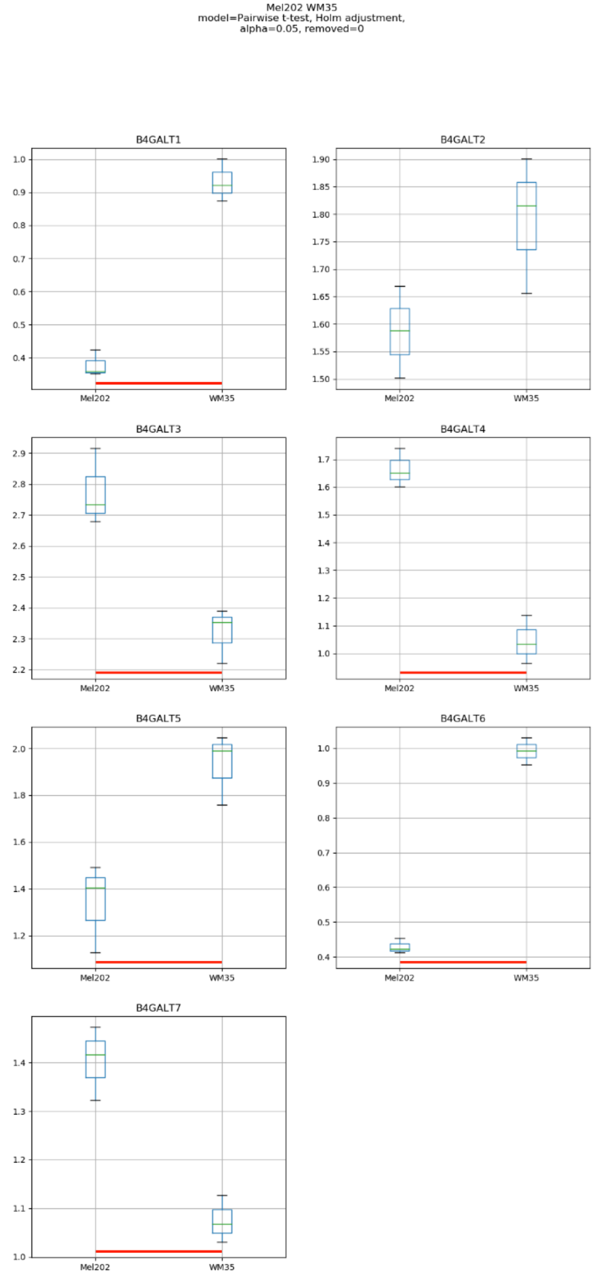
Mel202 WM35
model=Pairwise t-test, Holm adjustment,
alpha=0.05, removed=0
B4GALT1
B4GALT2
B4GALT3
B4GALT4
B4GALT5
B4GALT6
B4GALT7
Mel202
WM35

### Model No. 6

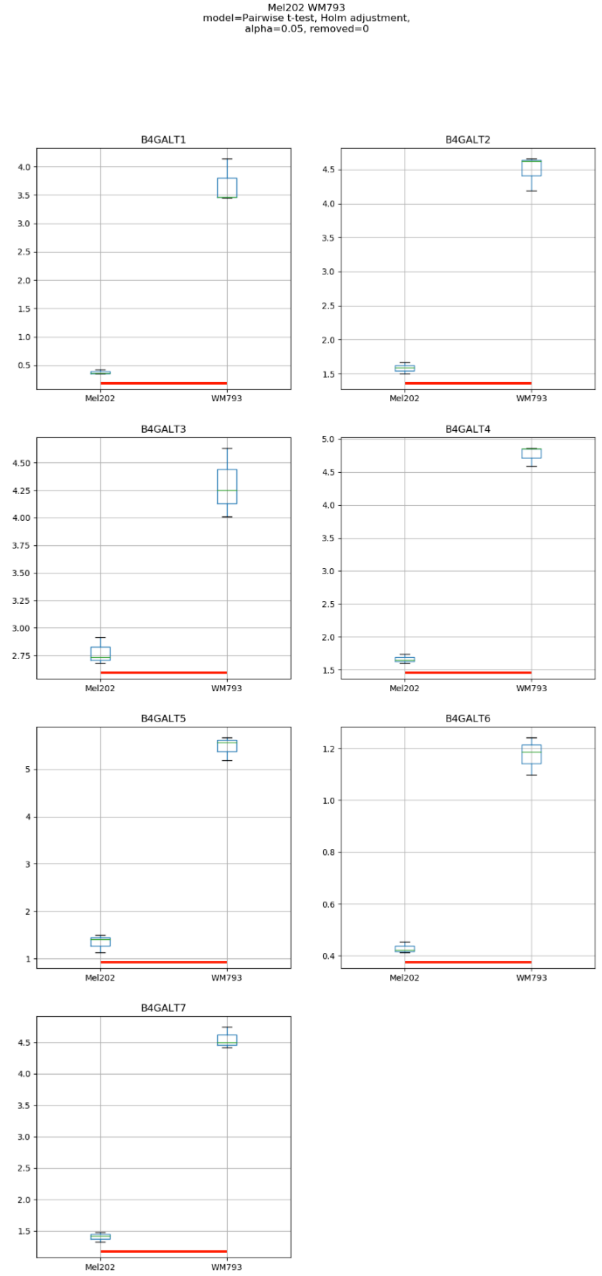
Mel202 WM793
model=Pairwise t-test, Holm adjustment,
alpha=0.05, removed=0
B4GALT1
B4GALT2
B4GALT3
B4GALT4
B4GALT5
B4GALT6
B4GALT7
Mel202
WM793

### Model No. 7

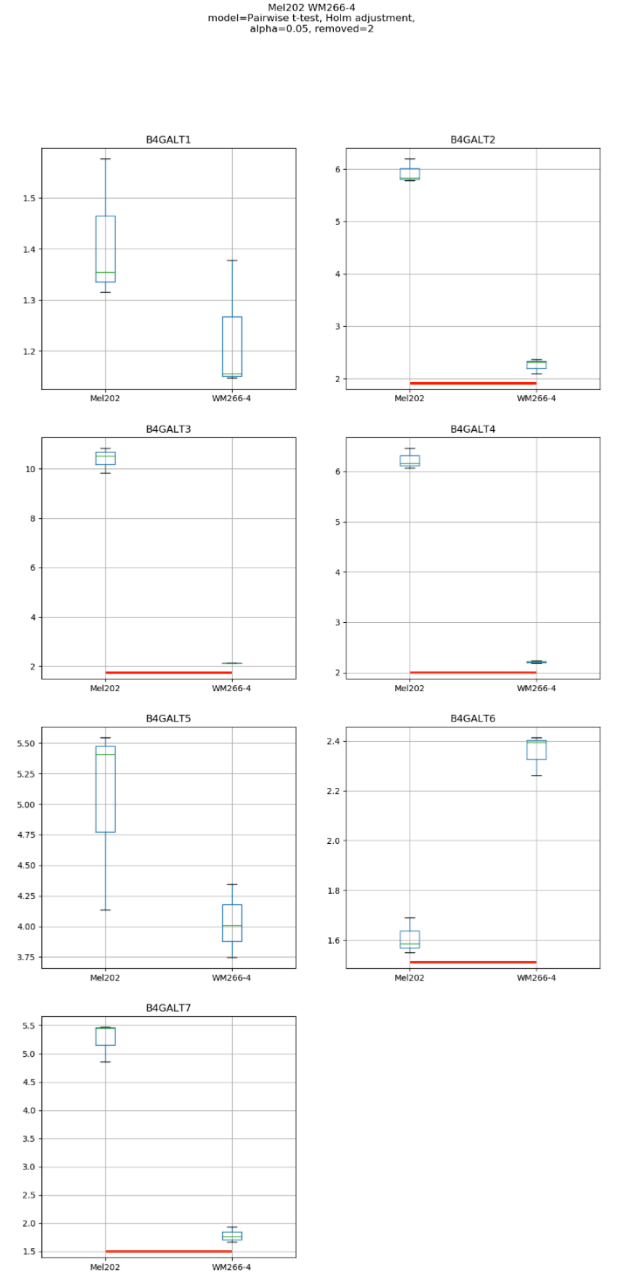
Mel202 WM266-4
model=Pairwise t-test, Holm adjustment,
alpha=0.05, removed=2
B4GALT1
B4GALT2
B4GALT3
B4GALT4
B4GALT5
B4GALT6
B4GALT7
Mel202
WM266-4

### Model No. 8

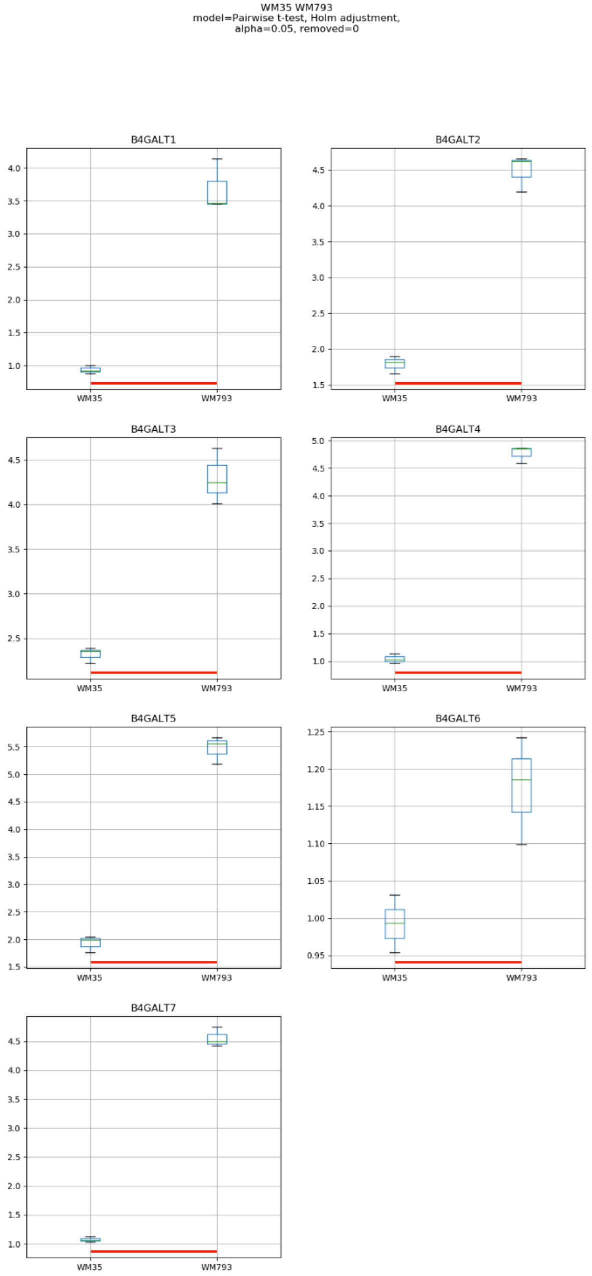
WM35 WM793
model=Pairwise t-test, Holm adjustment,
alpha=0.05, removed=0
B4GALT1
B4GALT2
B4GALT3
B4GALT4
B4GALT5
B4GALT6
B4GALT7
WM35
WM793

Model No. 9

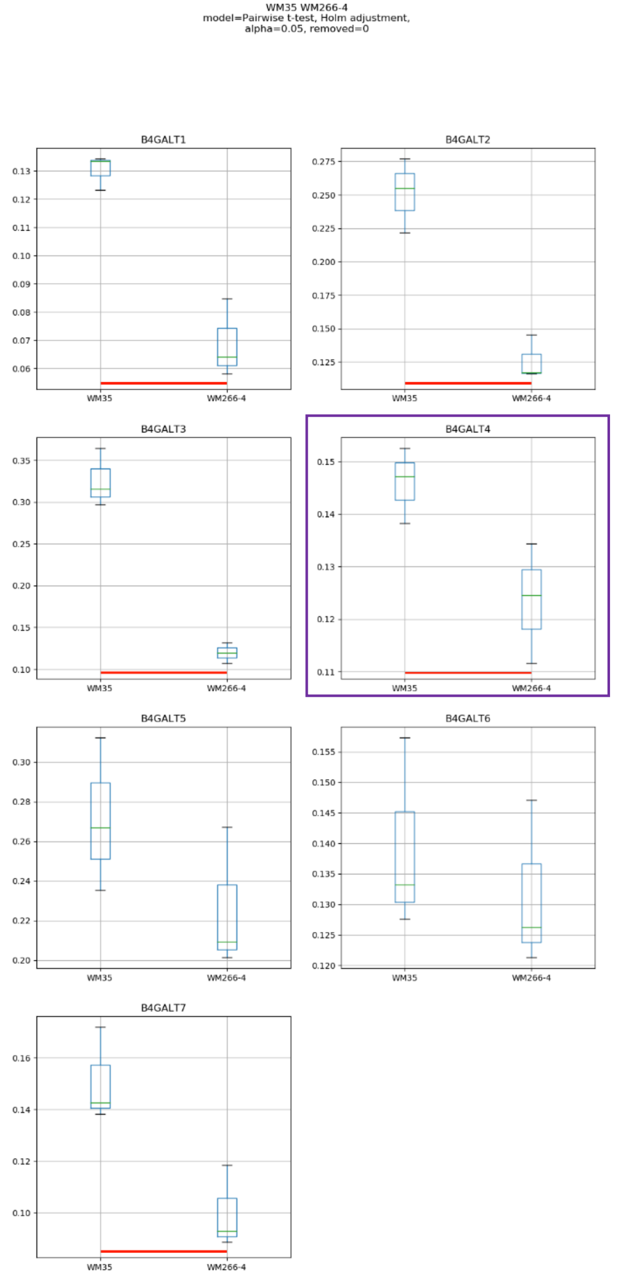
WM35 WM266-4
model=Pairwise t-test, Holm adjustment,
alpha=0.05, removed=0
B4GALT1
B4GALT2
B4GALT3
B4GALT4
B4GALT5
B4GALT6
B4GALT7
WM35
WM266-4

Model No. 10

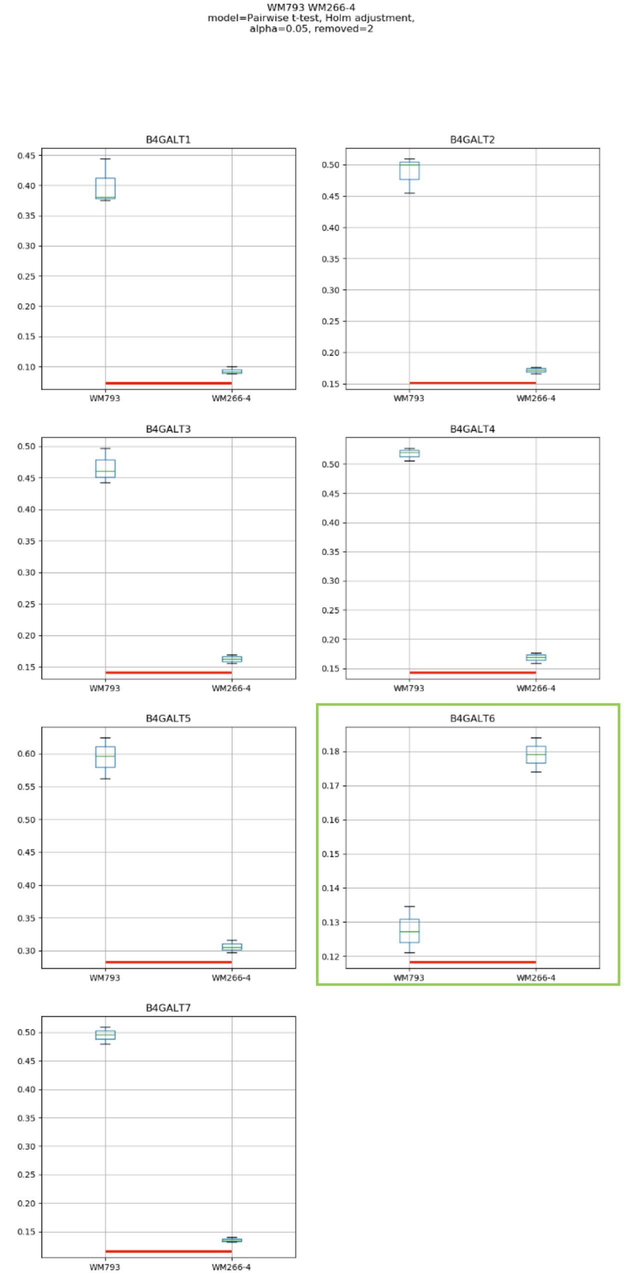
WM793 WM266-4
model=Pairwise t-test, Holm adjustment,
alpha=0.05, removed=2
B4GALT1
B4GALT2
B4GALT3
B4GALT4
B4GALT5
B4GALT6
B4GALT7
WM793
WM266-4

### Model No. 11

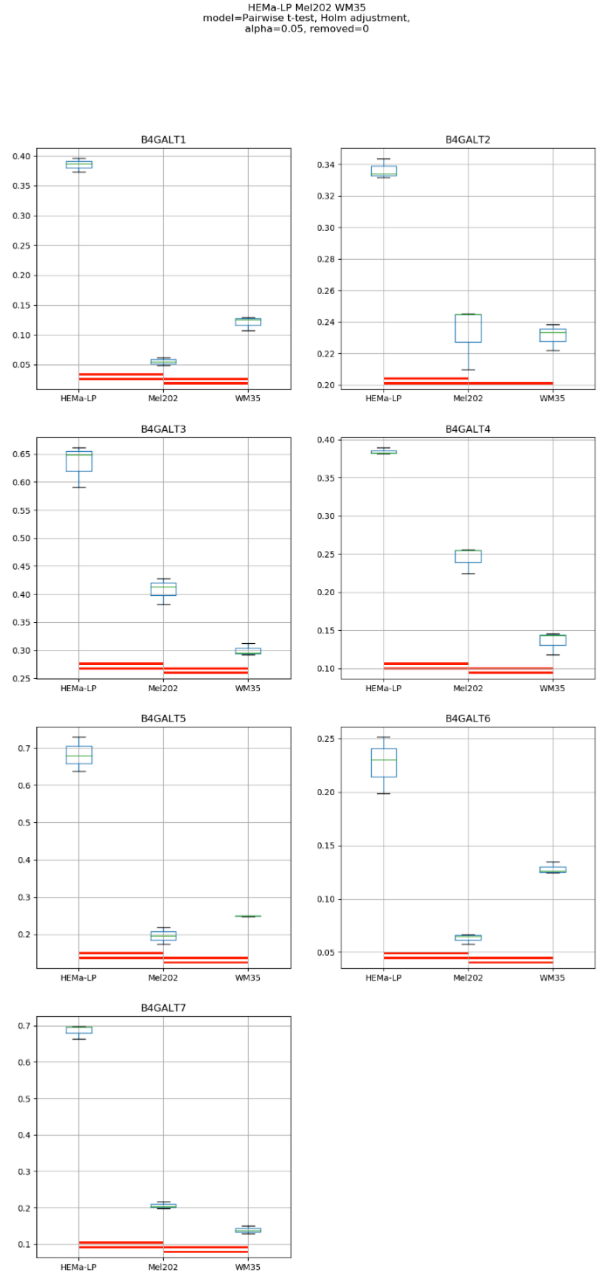
HEMa-LP Mel202 WM35
model=Pairwise t-test, Holm adjustment,
alpha=0.05, removed=0
B4GALT1
B4GALT2
B4GALT3
B4GALT4
B4GALT5
B4GALT6
B4GALT7
HEMa-LP
Mel202
WM35

### Model No. 12

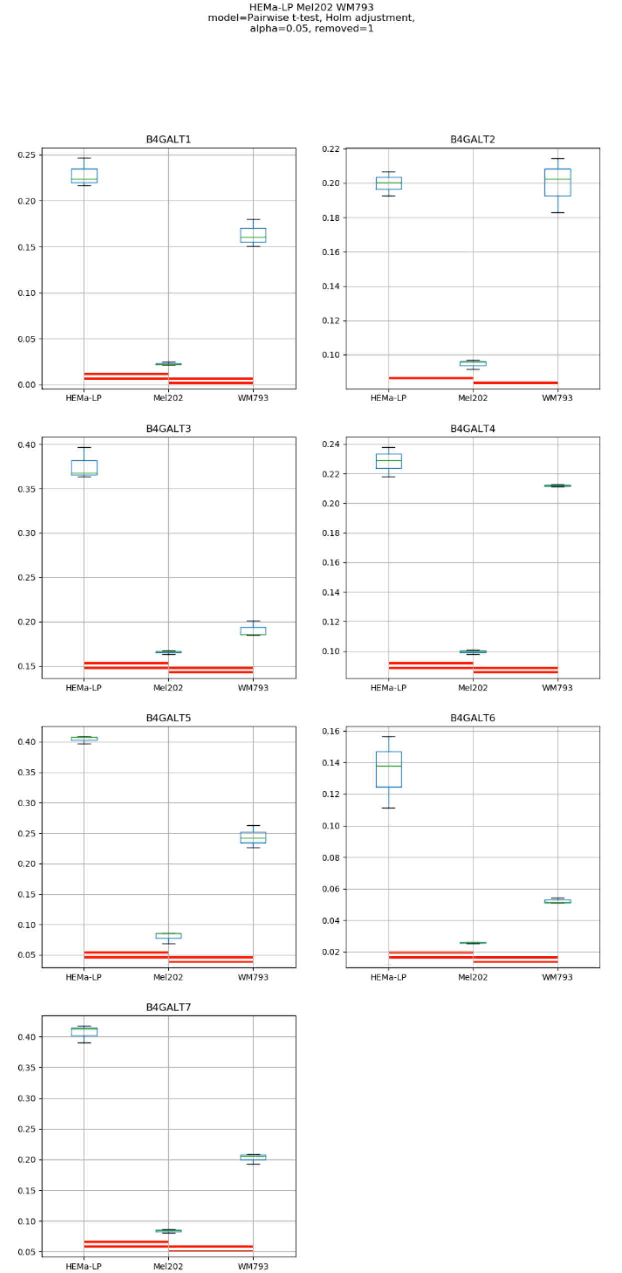

HEMa-LP Mel202 WM793
model=Pairwise t-test, Holm adjustment,
alpha=0.05, removed=1
B4GALT1
B4GALT2
B4GALT3
B4GALT4
B4GALT5
B4GALT6
B4GALT7
HEMa-LP
Mel202
WM793

### Model No. 13

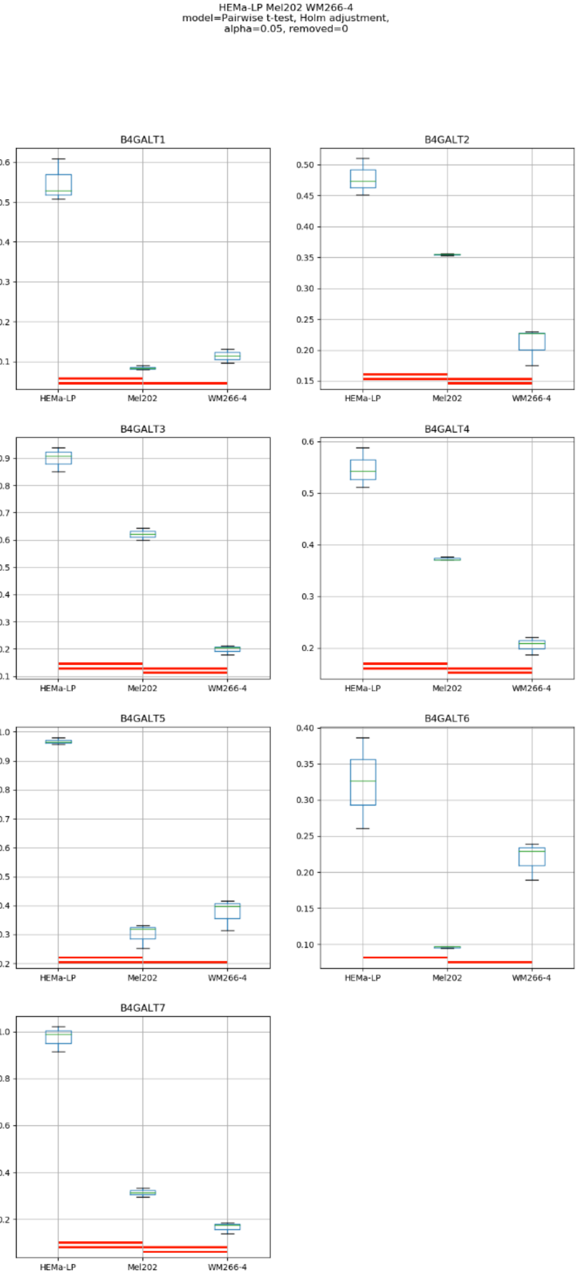
HEMa-LP Mel202 WM266-4
model=Pairwise t-test, Holm adjustment,
alpha=0.05, removed=0
B4GALT1
B4GALT2
B4GALT3
B4GALT4
B4GALT5
B4GALT6
B4GALT7
HEMa-LP
Mel202
WM266-4

Model No. 14

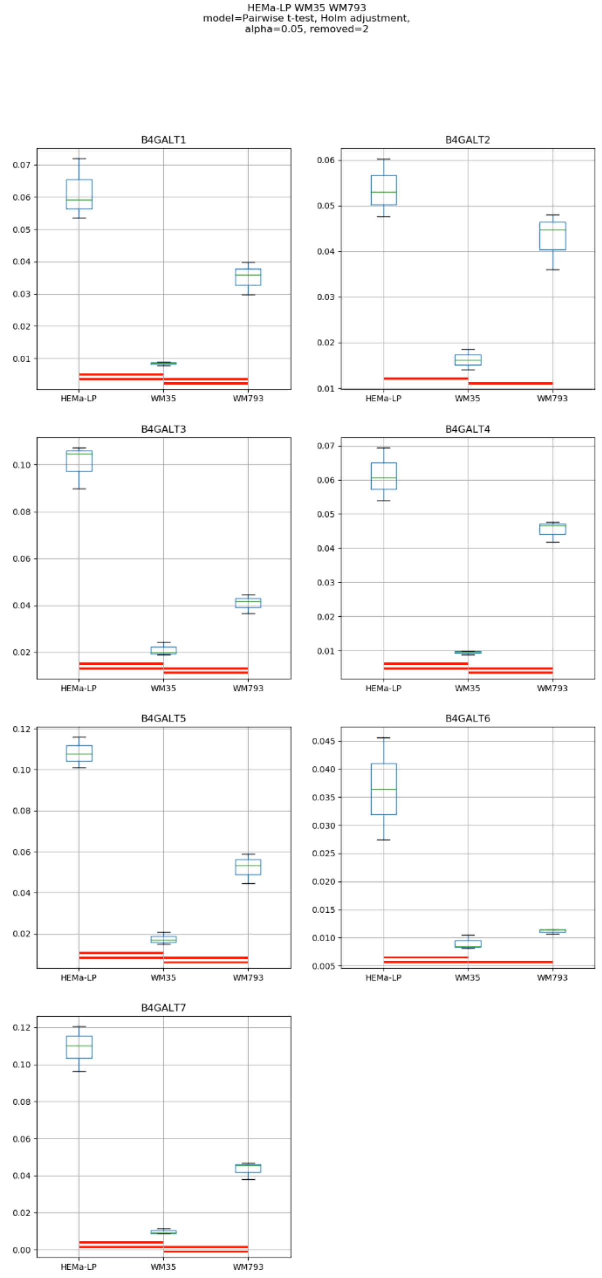
HEMa-LP WM35 WM793
model=Pairwise t-test, Holm adjustment,
alpha=0.05, removed=2
B4GALT1
B4GALT2
B4GALT3
B4GALT4
B4GALT5
B4GALT6
B4GALT7
HEMa-LP
WM35
WM793

### Model No. 15

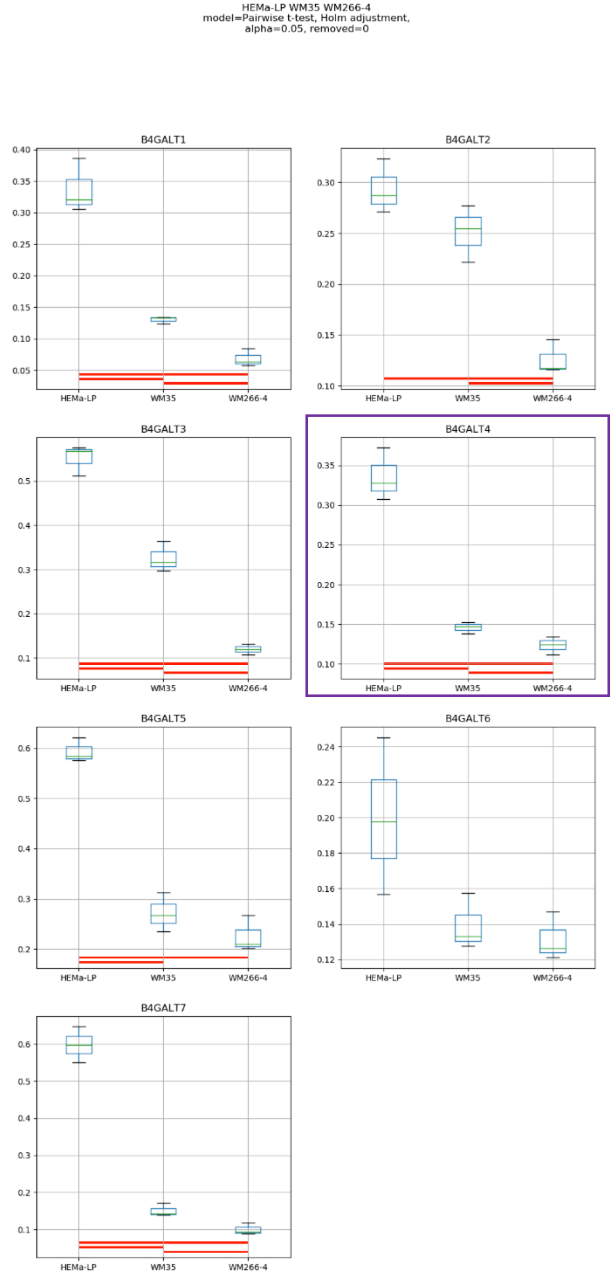
HEMa-LP WM35 WM266-4
model=Pairwise t-test, Holm adjustment,
alpha=0.05, removed=0
B4GALT1
B4GALT2
B4GALT3
B4GALT4
B4GALT5
B4GALT6
B4GALT7
HEMa-LP
WM35
WM266-4

### Model No. 16

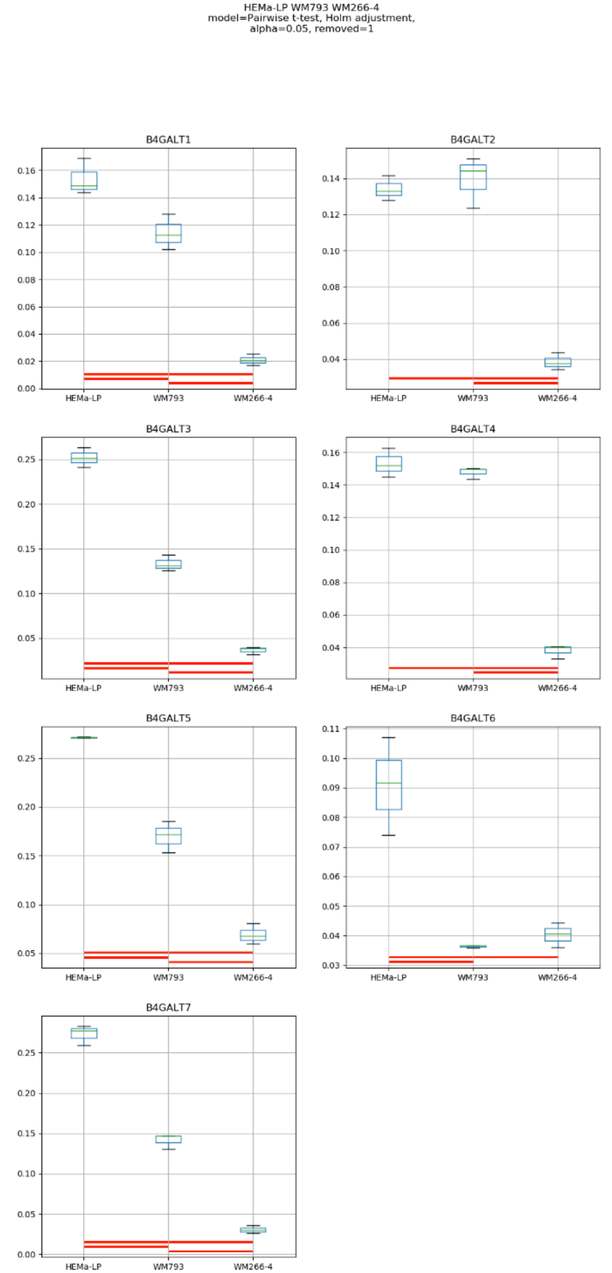
HEMa-LP WM793 WM266-4
model=Pairwise t-test, Holm adjustment,
alpha=0.05, removed=1
B4GALT1
B4GALT2
B4GALT3
B4GALT4
B4GALT5
B4GALT6
B4GALT7
HEMa-LP
WM793
WM266-4

Model No. 17

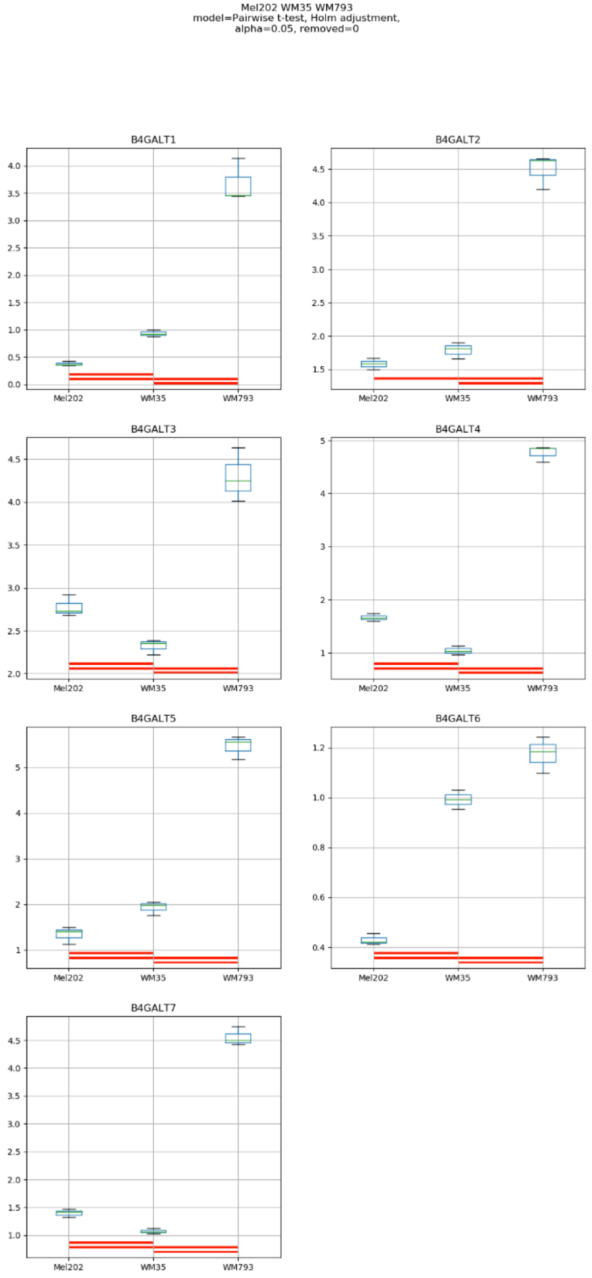
Mel202 WM35 WM793
model=Pairwise t-test, Holm adjustment,
alpha=0.05, removed=0
B4GALT1
B4GALT2
B4GALT3
B4GALT4
B4GALT5
B4GALT6
B4GALT7
Mel202
WM35
WM793

### Model No. 18

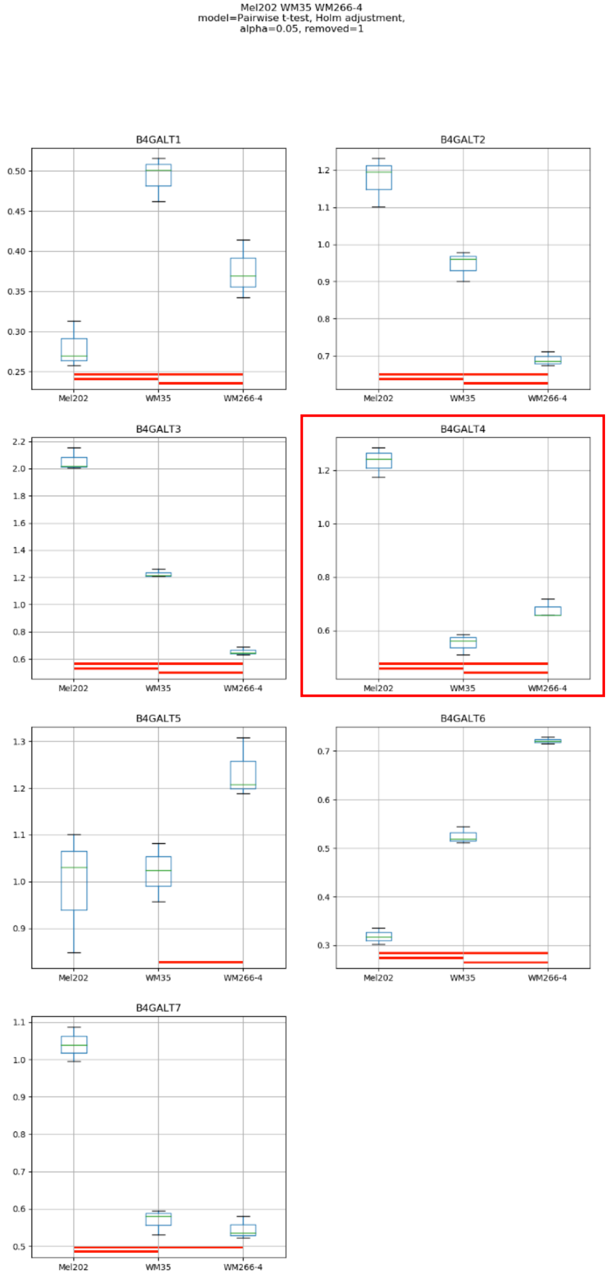

Mel202 WM35 WM266-4
model=Pairwise t-test, Holm adjustment,
alpha=0.05, removed=1
B4GALT1
B4GALT2
B4GALT3
B4GALT4
B4GALT5
B4GALT6
B4GALT7
Mel202
WM35
WM266-4

### Model No. 19

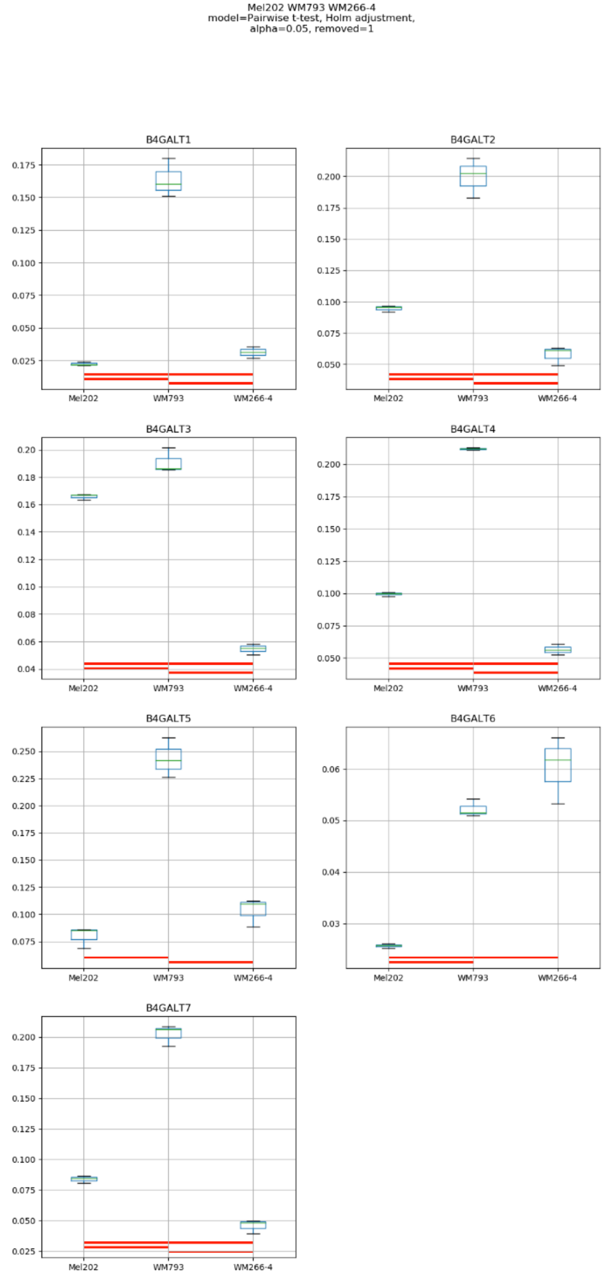
Mel202 WM793 WM266-4
model=Pairwise t-test, Holm adjustment,
alpha=0.05, removed=1
B4GALT1
B4GALT2
B4GALT3
B4GALT4
B4GALT5
B4GALT6
B4GALT7
Mel202
WM793
WM266-4

### Model No. 20

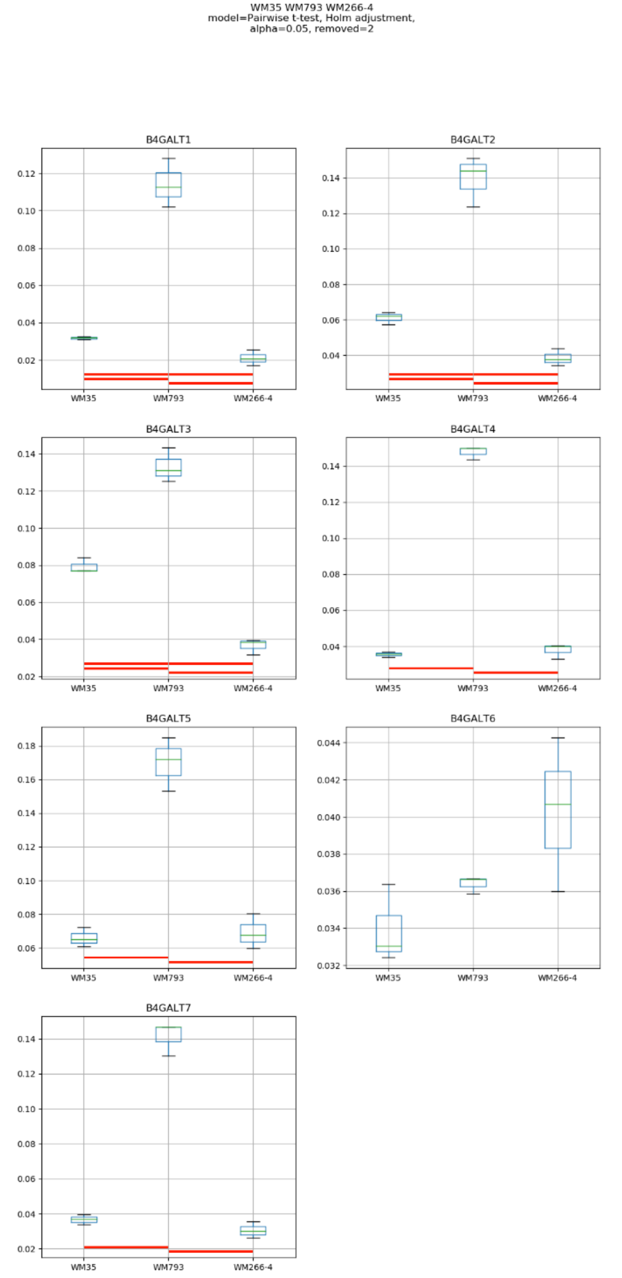
WM35 WM793 WM266-4
model=Pairwise t-test, Holm adjustment,
alpha=0.05, removed=2
B4GALT1
B4GALT2
B4GALT3
B4GALT4
B4GALT5
B4GALT6
B4GALT7
WM35
WM793
WM266-4

### Model No. 21

HEMa-LP Mel202 WM35 WM793
model=Pairwise t-test, Holm adjustment,
alpha=0.05, removed=2

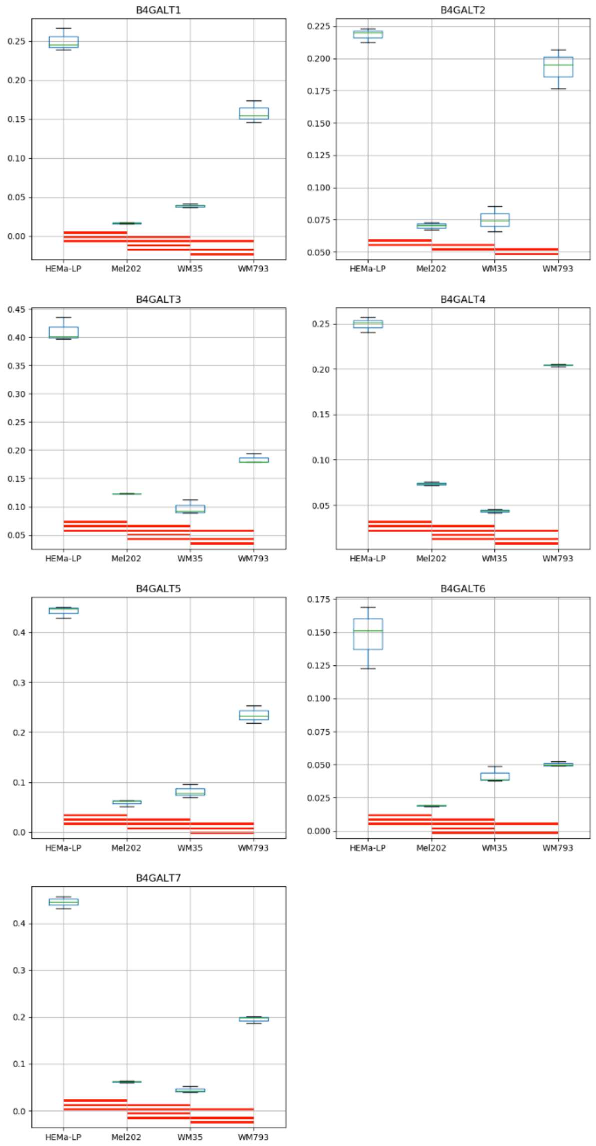

Model No. 22

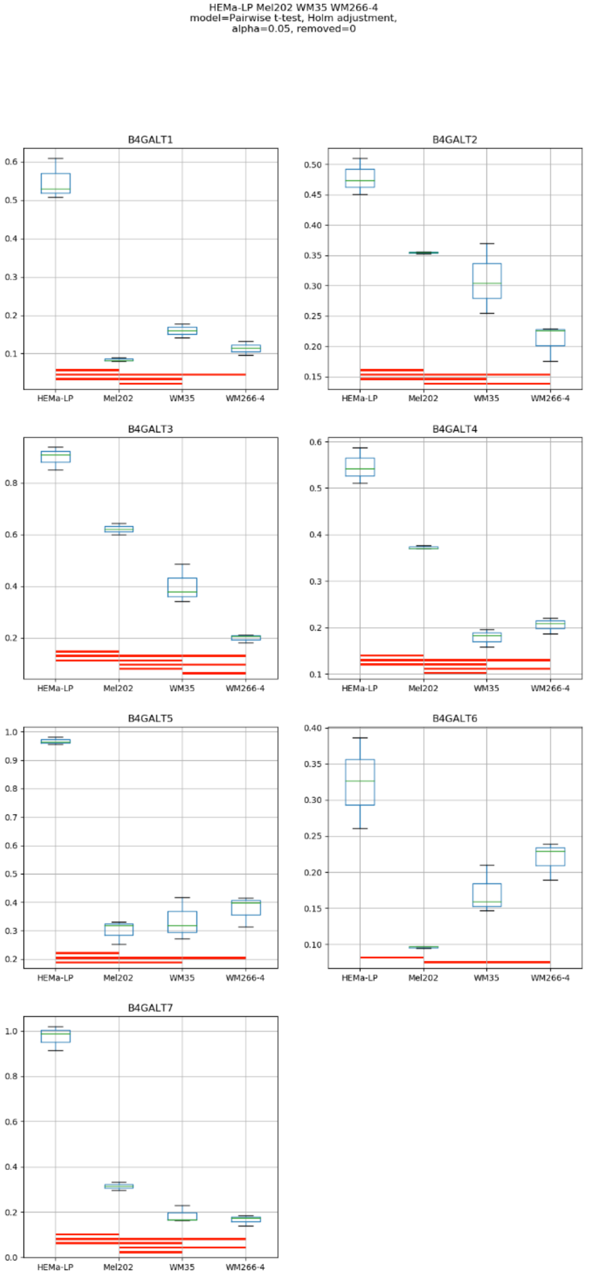
HEMa-LP Mel202 WM35 WM266-4
model=Pairwise t-test, Holm adjustment,
alpha=0.05, removed=0
B4GALT1
B4GALT2
B4GALT3
B4GALT4
B4GALT5
B4GALT6
B4GALT7
HEMa-LP
Mel202
WM35
WM266-4

### Model No. 23

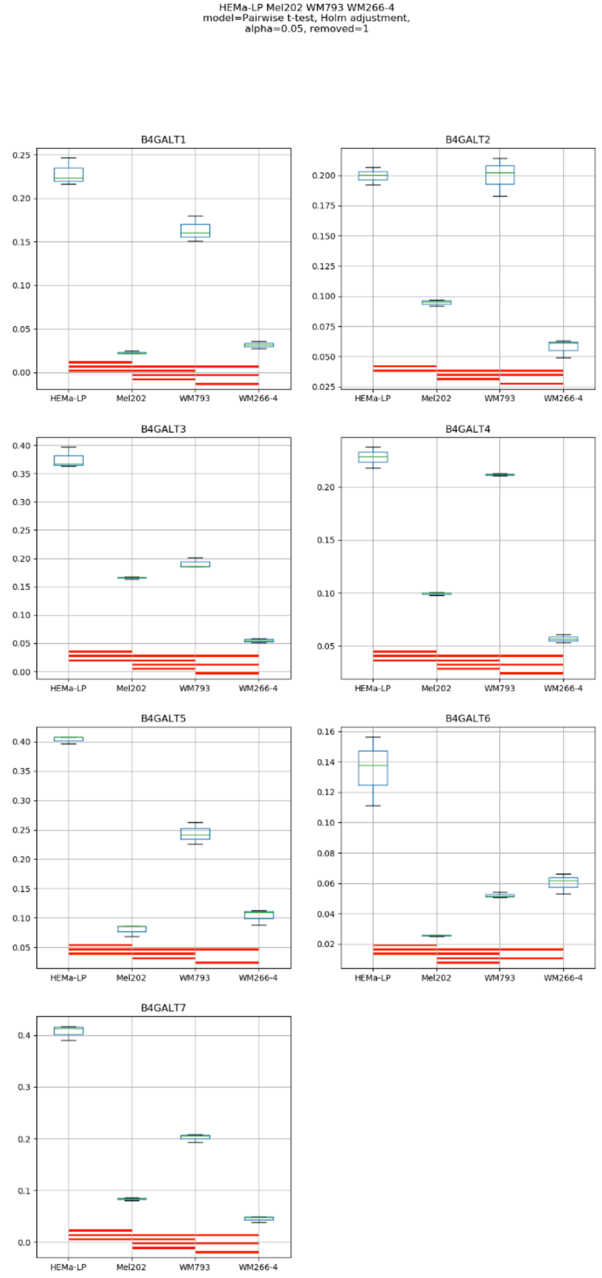
HEMa-LP Mel202 WM793 WM266-4
model=Pairwise t-test, Holm adjustment,
alpha=0.05, removed=1
B4GALT1
B4GALT2
B4GALT3
B4GALT4
B4GALT5
B4GALT6
B4GALT7
HEMa-LP
Mel202
WM793
WM266-4

Model No. 24

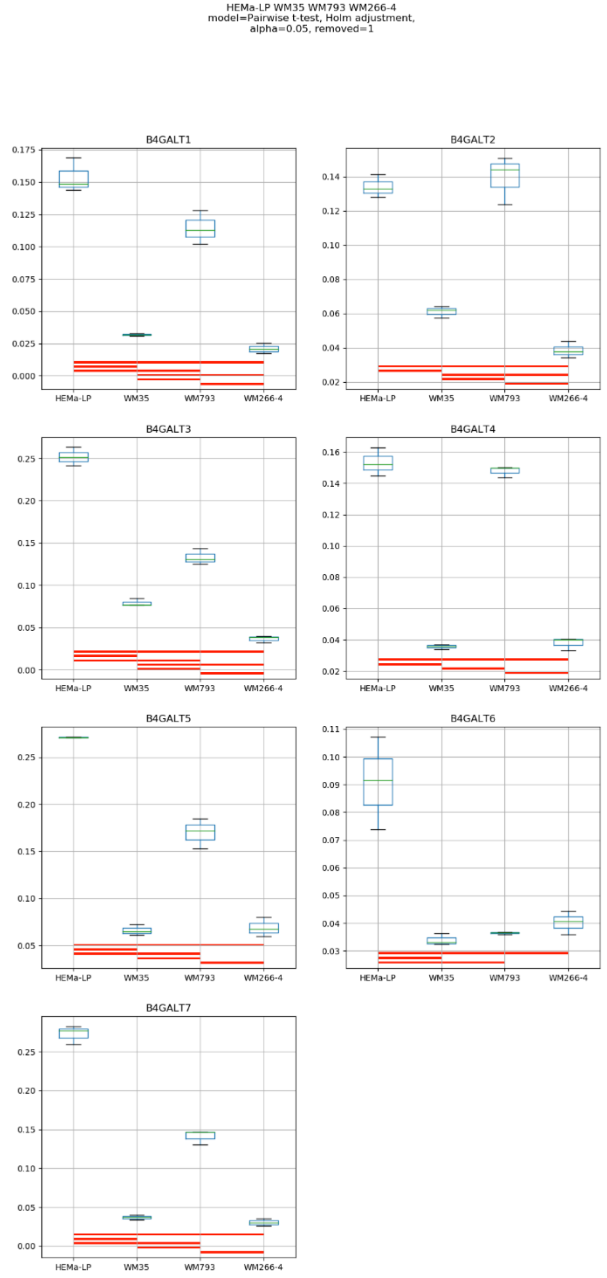
HEMa-LP WM35 WM793 WM266-4
model=Pairwise t-test, Holm adjustment,
alpha=0.05, removed=1
B4GALT1
B4GALT2
B4GALT3
B4GALT4
B4GALT5
B4GALT6
B4GALT7
HEMa-LP
WM35
WM793
WM266-4

### Model No. 25

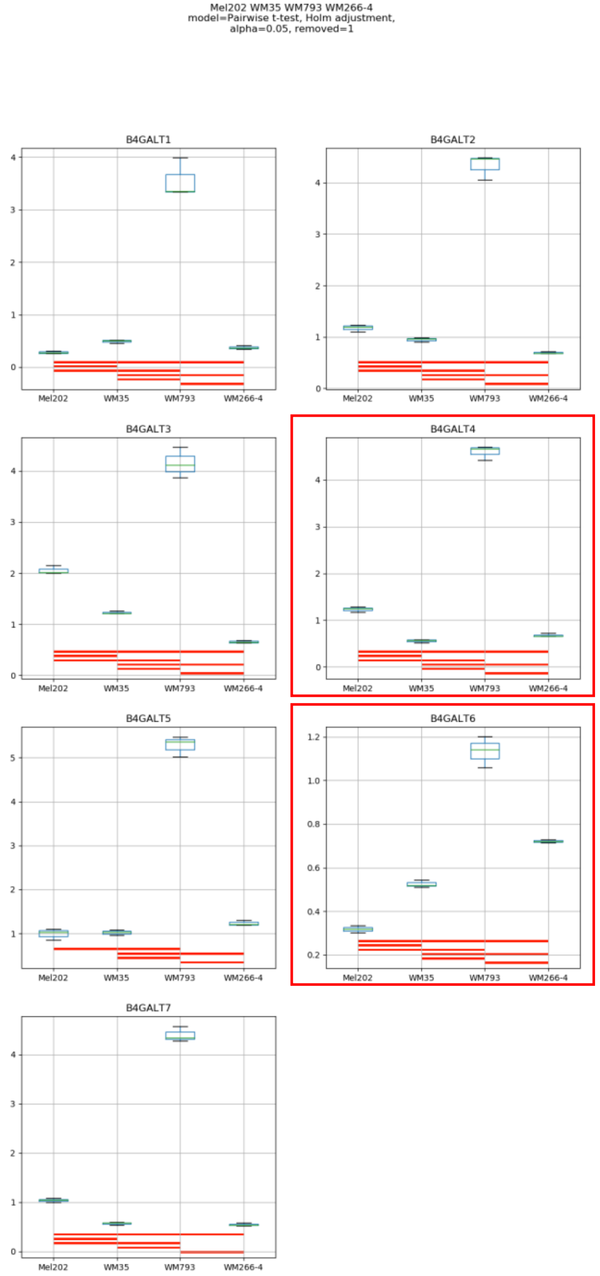

Mel202 WM35 WM793 WM266-4
model=Pairwise t-test, Holm adjustment,
alpha=0.05, removed=1
B4GALT1
B4GALT2
B4GALT3
B4GALT4
B4GALT5
B4GALT6
B4GALT7
Mel202
WM35
WM793
WM266-4

Model No. 26

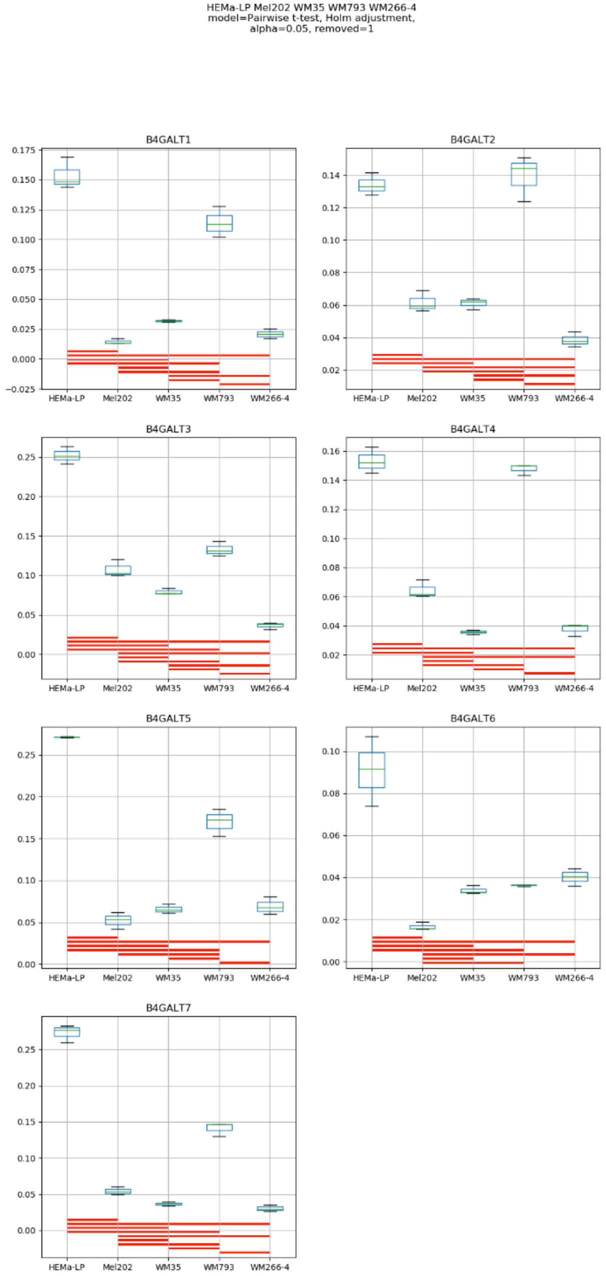
HEMa-LP Mel202 WM35 WM793 WM266-4
model=Pairwise t-test, Holm adjustment,
alpha=0.05, removed=1
B4GALT1
B4GALT2
B4GALT3
B4GALT4
B4GALT5
B4GALT6
B4GALT7
HEMa-LP
Mel202
WM35
WM793
WM266-4

**Supplementary Figure S10:** Medians of relative quantity (RQ) for target genes *B4GALT1–B4GALT7* normalized to the reference gene or pair of reference genes with the best stability values resulting from GenExpA analysis based on set of five candidate reference genes (*PGK1*, *RPS23*, *SNRPA*, *HPRT1*, *GUSB*) and remove repetition level 2 (see Table 1, part B). The statistical analyses used the pairwise t-test with Holm adjustment. Red line represents statistical significance, $P < 0.05$. Frames denote box-plots representing the medians of the RQ values for a given target gene without a repeatable pattern in the same cell lines under different models (the ‘Remove repetitions’ box set to 0 – violet frame, or to 1 – red frame, or to 2 – green frame). Detailed results are listed in Supplementary Table 9.

### Model No. 1

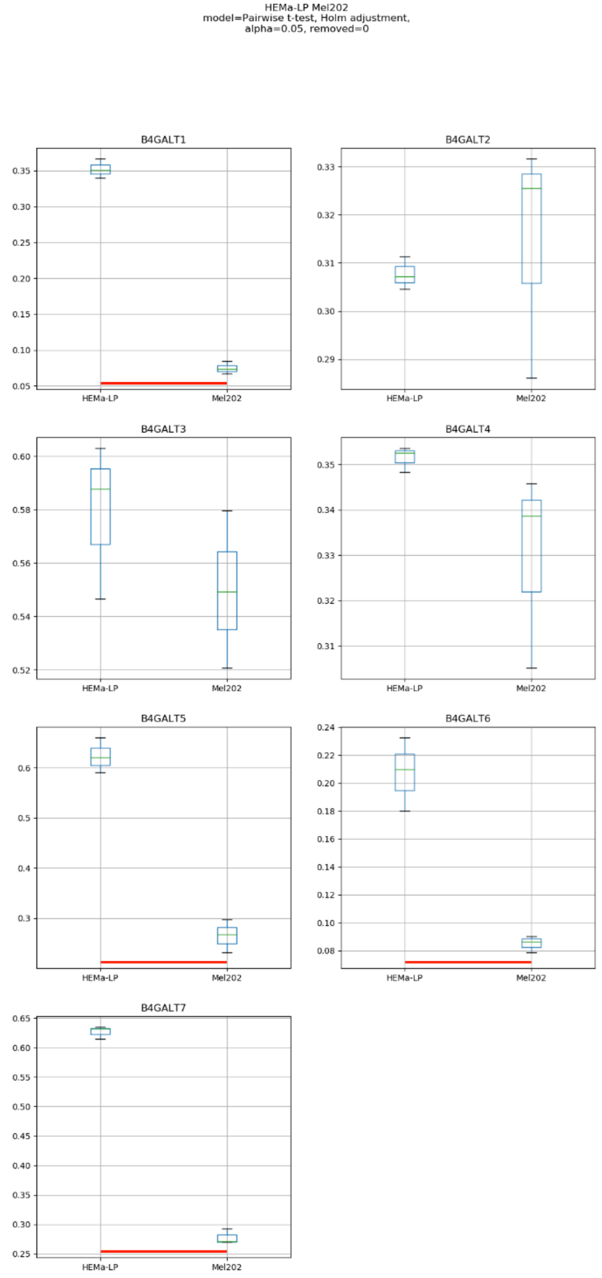
HEMa-LP Mel202
model=Pairwise t-test, Holm adjustment,
alpha=0.05, removed=0
B4GALT1
B4GALT2
B4GALT3
B4GALT4
B4GALT5
B4GALT6
B4GALT7
HEMa-LP
Mel202

### Model No. 2

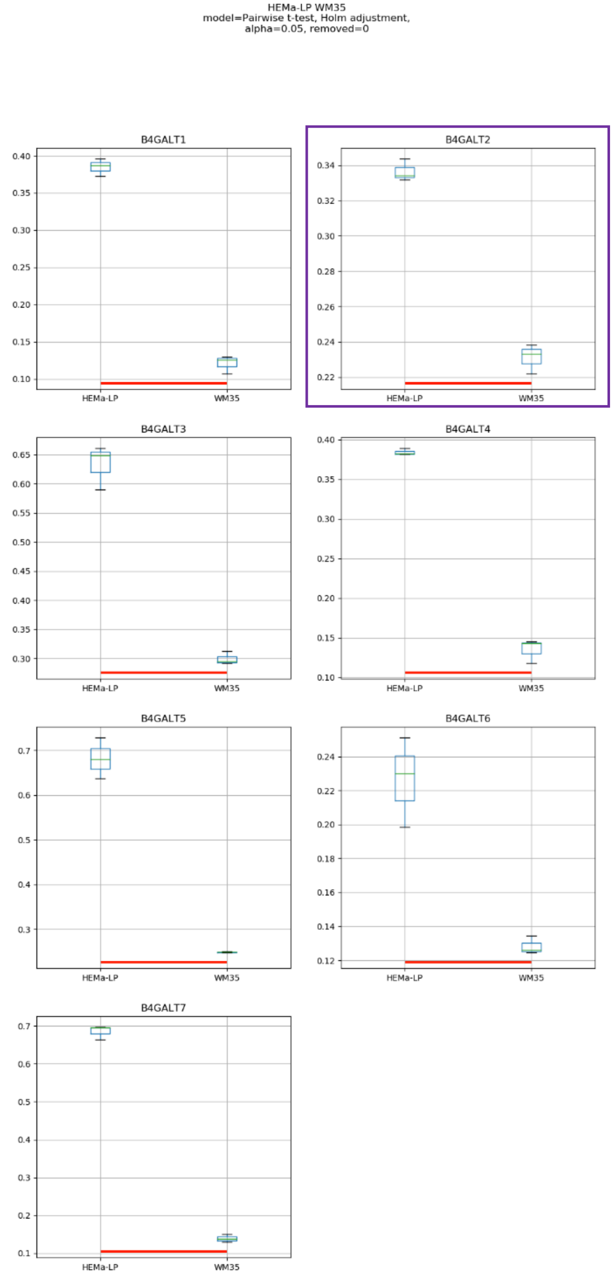

HEMa-LP WM35
model=Pairwise t-test, Holm adjustment,
alpha=0.05, removed=0
B4GALT1
B4GALT2
B4GALT3
B4GALT4
B4GALT5
B4GALT6
B4GALT7
HEMa-LP
WM35

Model No. 3

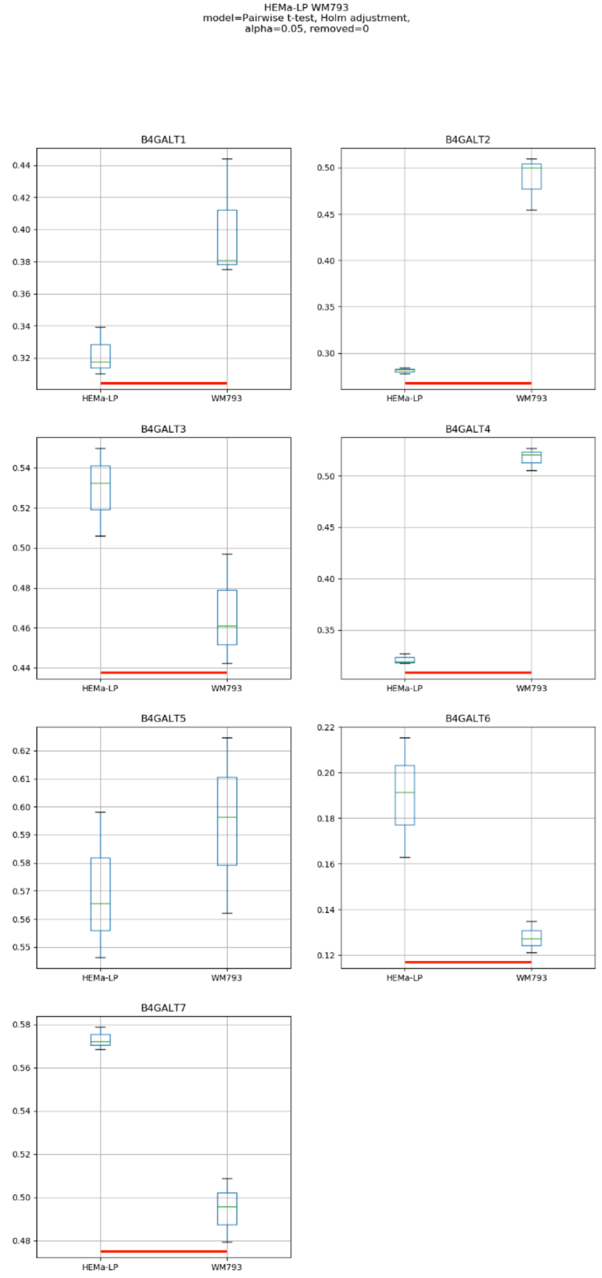
HEMa-LP WM793
model=Pairwise t-test, Holm adjustment,
alpha=0.05, removed=0
B4GALT1
B4GALT2
B4GALT3
B4GALT4
B4GALT5
B4GALT6
B4GALT7
HEMa-LP
WM793

Model No. 4

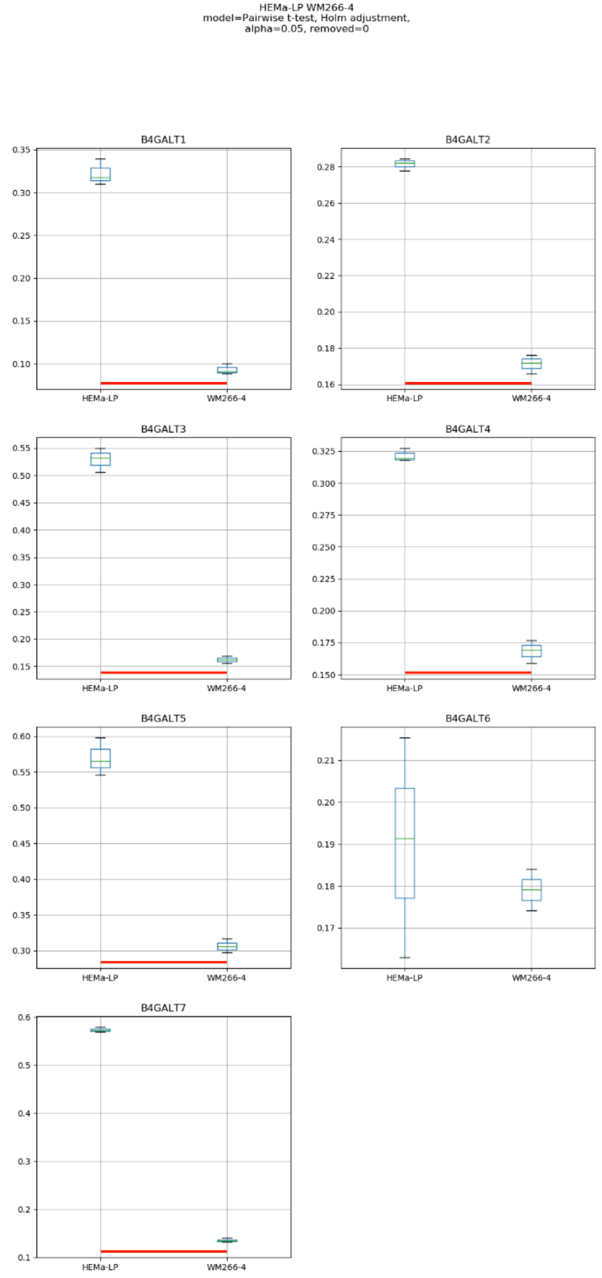
HEMa-LP WM266-4
model=Pairwise t-test, Holm adjustment,
alpha=0.05, removed=0
B4GALT1
B4GALT2
B4GALT3
B4GALT4
B4GALT5
B4GALT6
B4GALT7
HEMa-LP
WM266-4

### Model No. 5

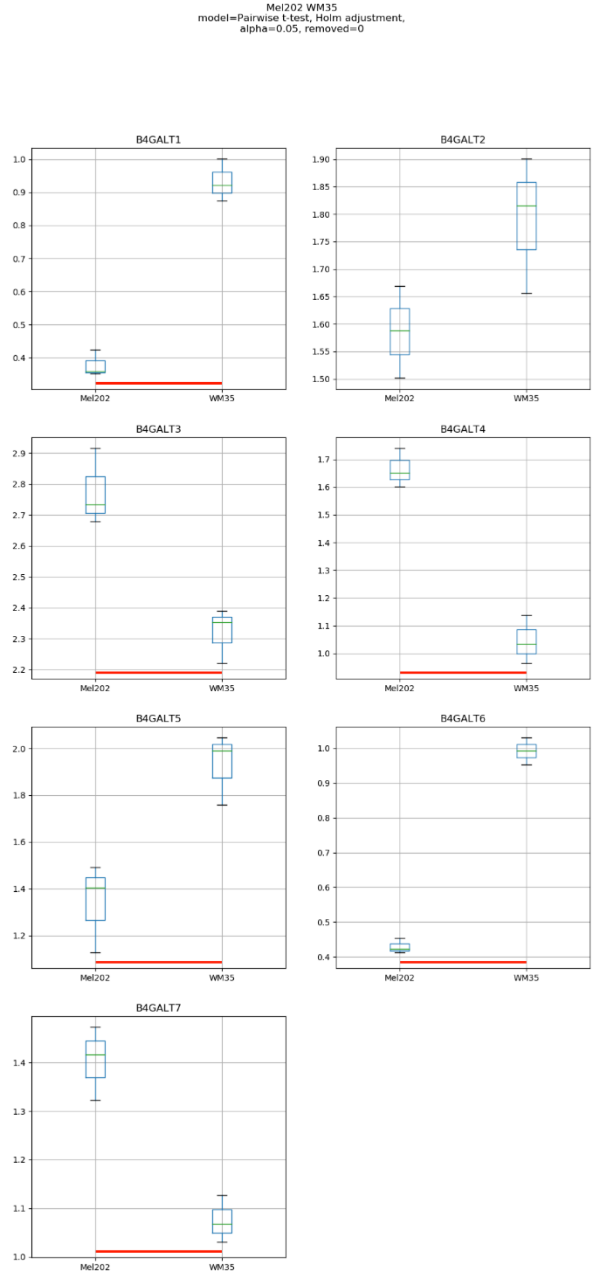
Mel202 WM35
model=Pairwise t-test, Holm adjustment,
alpha=0.05, removed=0
B4GALT1
B4GALT2
B4GALT3
B4GALT4
B4GALT5
B4GALT6
B4GALT7
Mel202
WM35

### Model No. 6

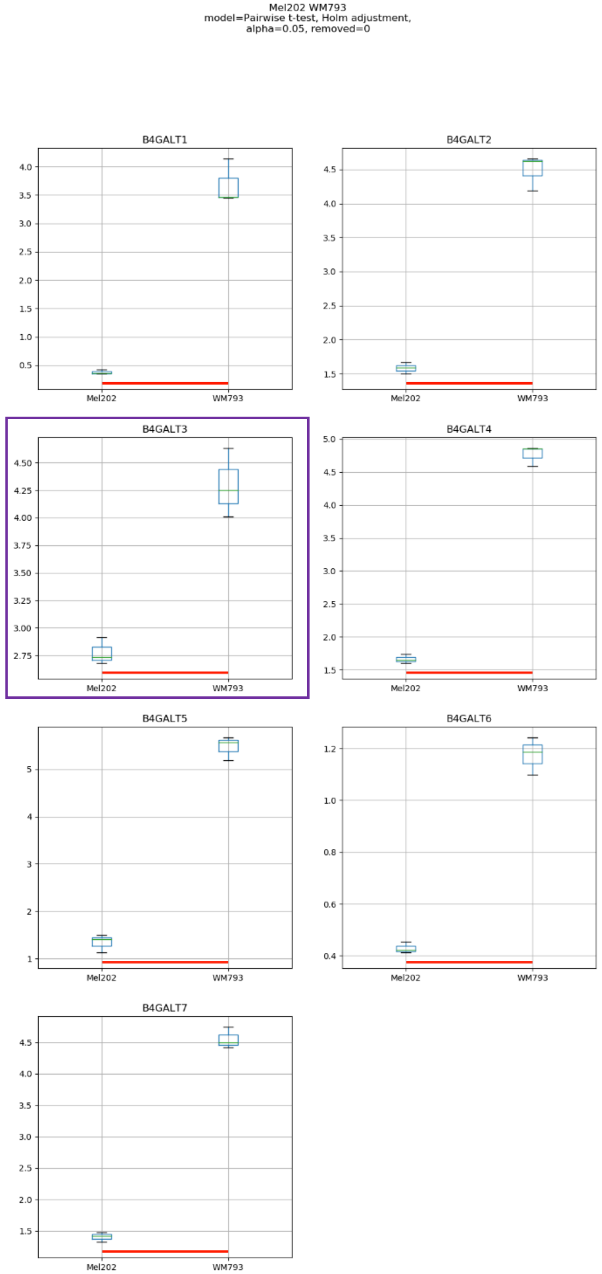

Mel202 WM793
model=Pairwise t-test, Holm adjustment,
alpha=0.05, removed=0
B4GALT1
B4GALT2
B4GALT3
B4GALT4
B4GALT5
B4GALT6
B4GALT7
Mel202
WM793

### Model No. 7

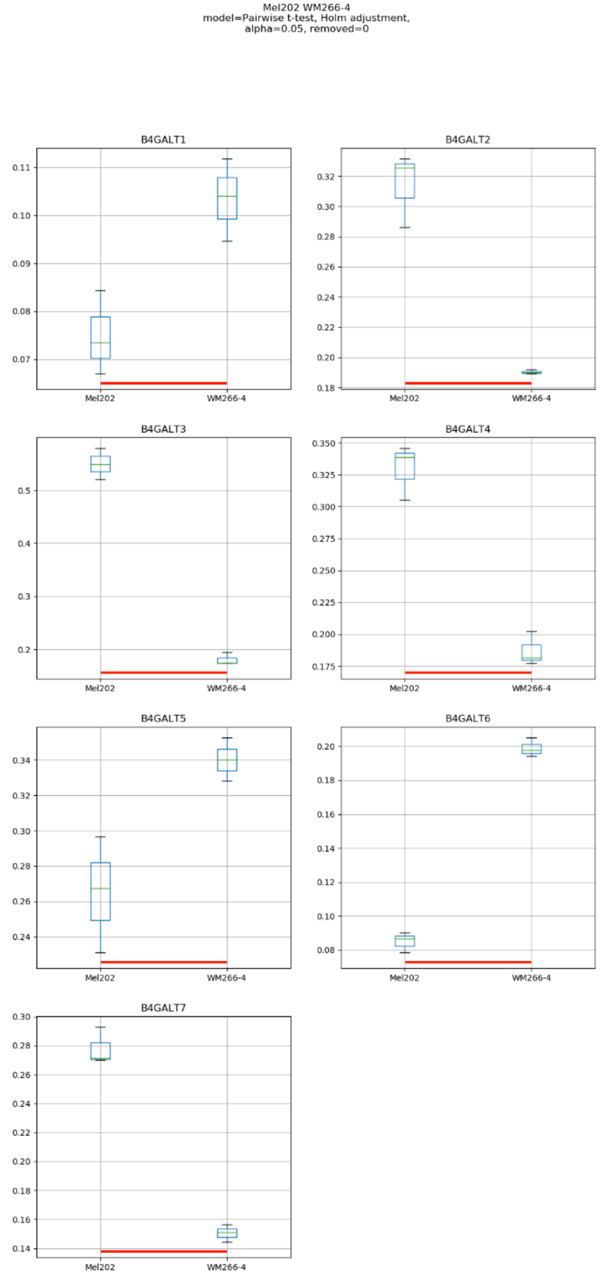
Mel202 WM266-4
model=Pairwise t-test, Holm adjustment,
alpha=0.05, removed=0
B4GALT1
B4GALT2
B4GALT3
B4GALT4
B4GALT5
B4GALT6
B4GALT7
Mel202
WM266-4

Model No. 8

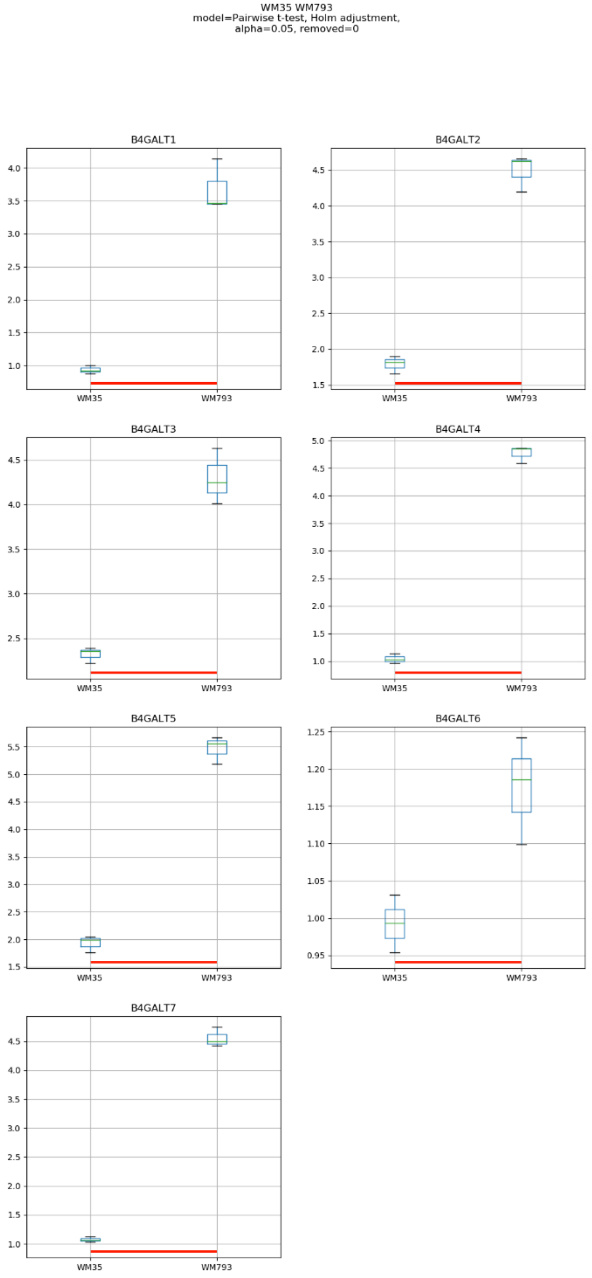
WM35 WM793
model=Pairwise t-test, Holm adjustment,
alpha=0.05, removed=0
B4GALT1
B4GALT2
B4GALT3
B4GALT4
B4GALT5
B4GALT6
B4GALT7
WM35
WM793

### Model No. 9

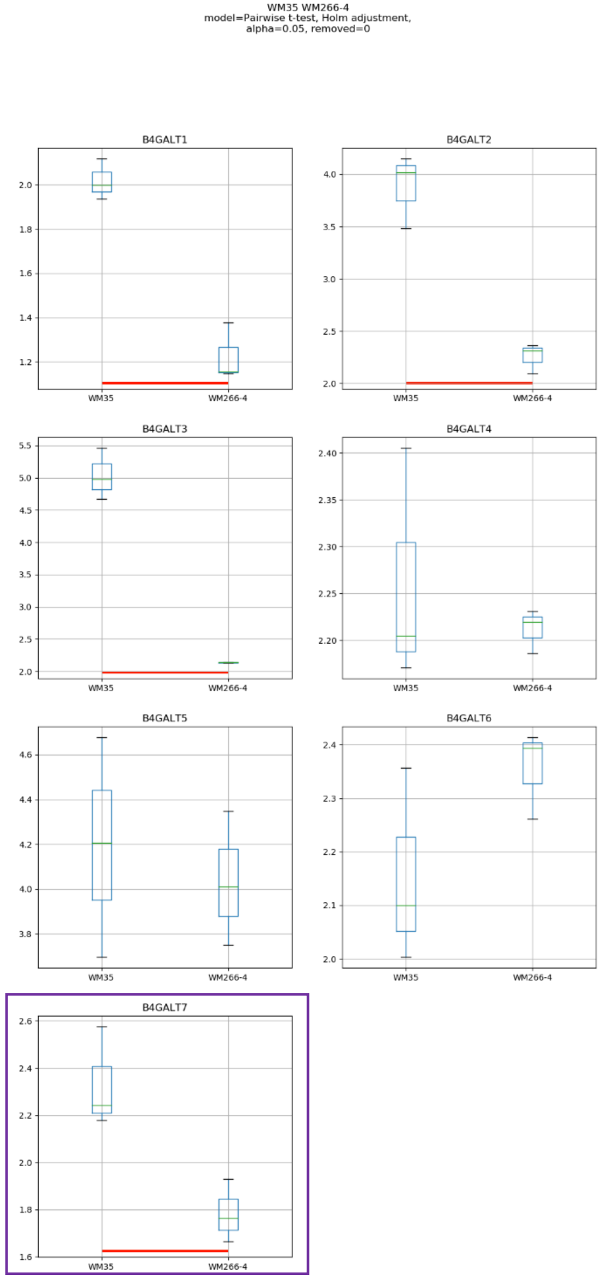
WM35 WM266-4
model=Pairwise t-test, Holm adjustment,
alpha=0.05, removed=0
B4GALT1
B4GALT2
B4GALT3
B4GALT4
B4GALT5
B4GALT6
B4GALT7
WM35
WM266-4

Model No. 10

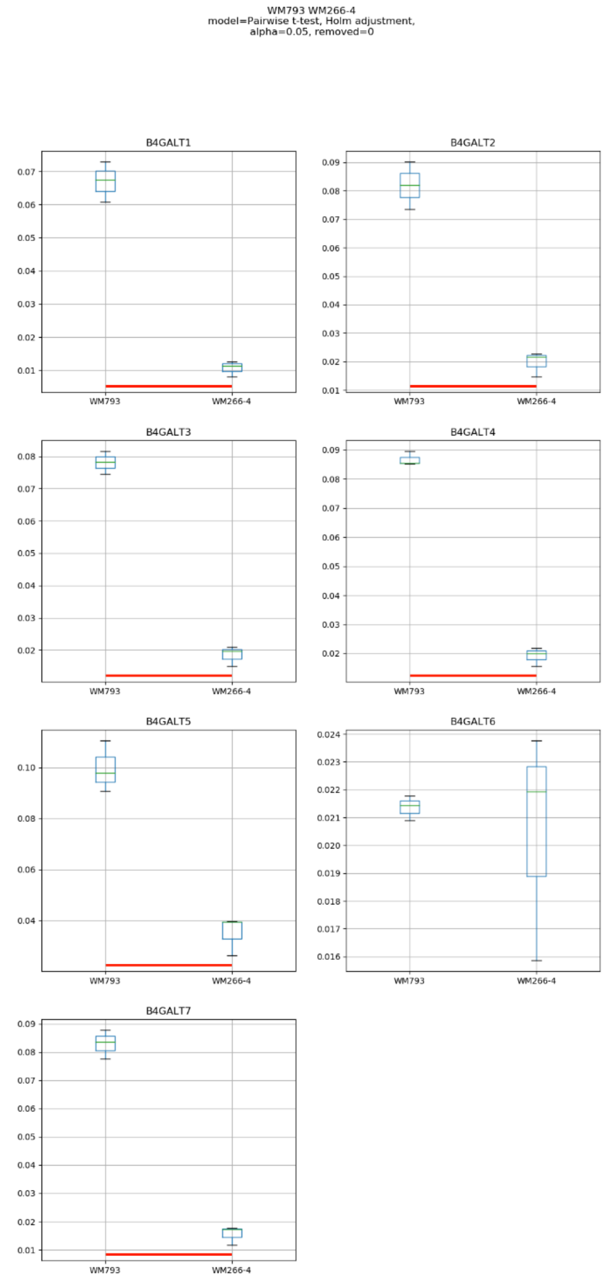
WM793 WM266-4
model=Pairwise t-test, Holm adjustment,
alpha=0.05, removed=0
B4GALT1
B4GALT2
B4GALT3
B4GALT4
B4GALT5
B4GALT6
B4GALT7
WM793
WM266-4

Model No. 11

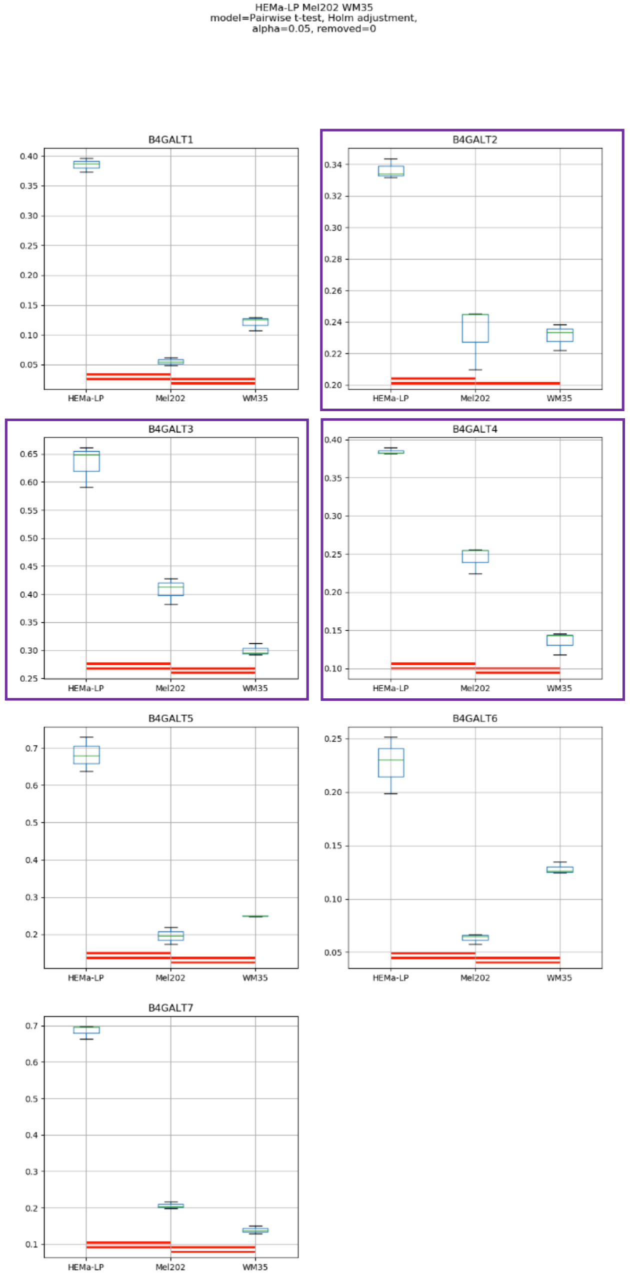
HEMa-LP Mel202 WM35
model=Pairwise t-test, Holm adjustment,
alpha=0.05, removed=0
B4GALT1
B4GALT2
B4GALT3
B4GALT4
B4GALT5
B4GALT6
B4GALT7
HEMa-LP
Mel202
WM35

### Model No. 12

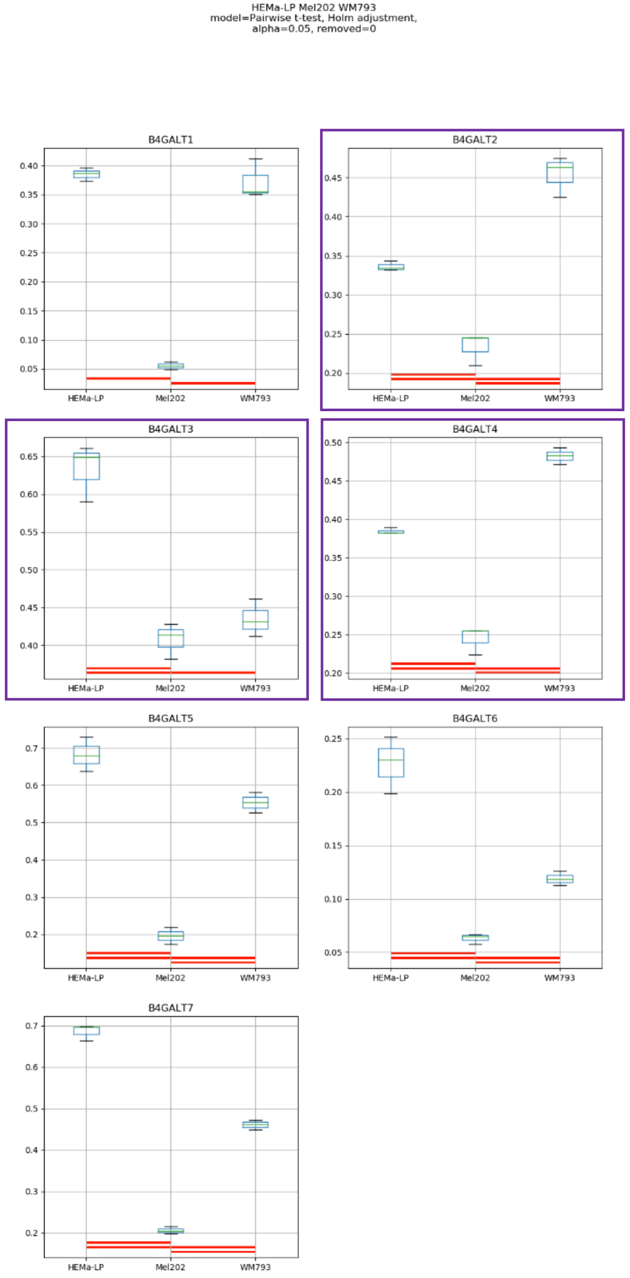
HEMa-LP Mel202 WM793
model=Pairwise t-test, Holm adjustment,
alpha=0.05, removed=0
B4GALT1
B4GALT2
B4GALT3
B4GALT4
B4GALT5
B4GALT6
B4GALT7
HEMa-LP
Mel202
WM793

Model No. 13

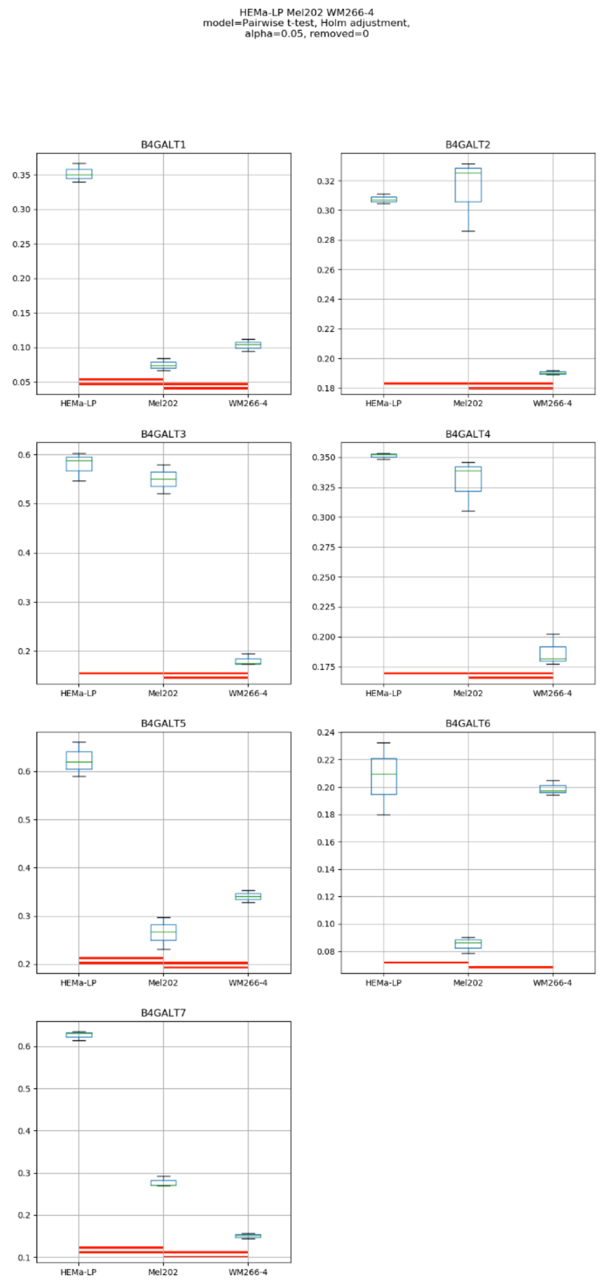
HEMa-LP Mel202 WM266-4
model=Pairwise t-test, Holm adjustment,
alpha=0.05, removed=0
B4GALT1
B4GALT2
B4GALT3
B4GALT4
B4GALT5
B4GALT6
B4GALT7
HEMa-LP
Mel202
WM266-4

### Model No. 14

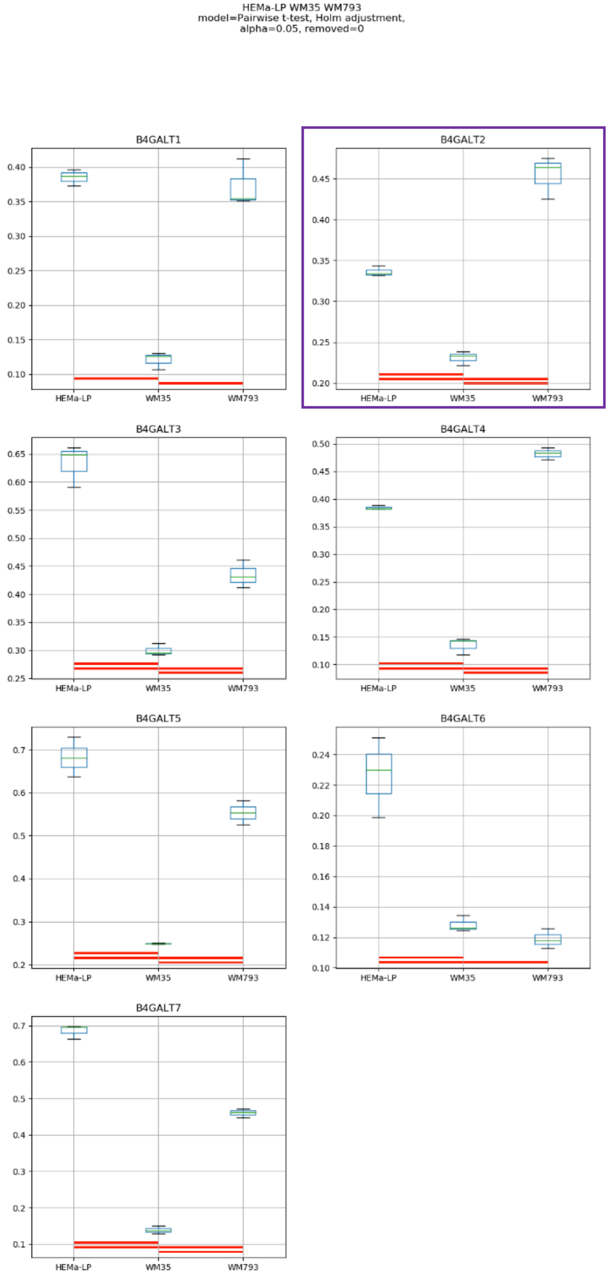
HEMa-LP WM35 WM793
model=Pairwise t-test, Holm adjustment,
alpha=0.05, removed=0
B4GALT1
B4GALT2
B4GALT3
B4GALT4
B4GALT5
B4GALT6
B4GALT7
HEMa-LP
WM35
WM793

Model No. 15

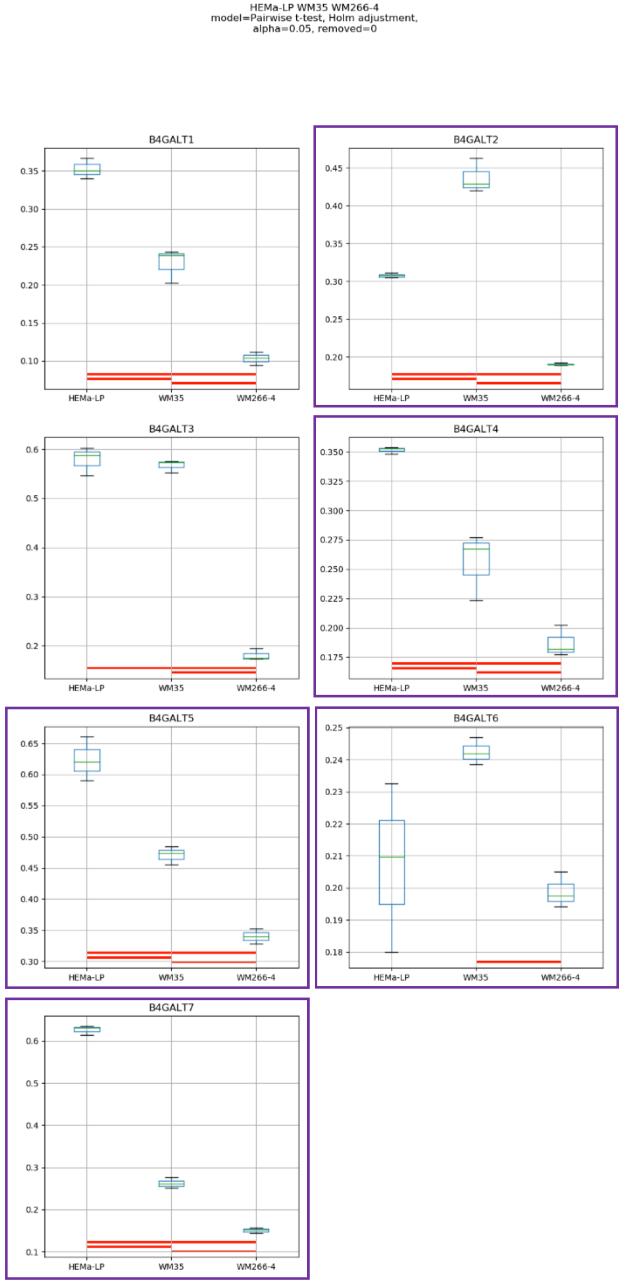
HEMa-LP WM35 WM266-4
model=Pairwise t-test, Holm adjustment,
alpha=0.05, removed=0
B4GALT1
B4GALT2
B4GALT3
B4GALT4
B4GALT5
B4GALT6
B4GALT7
HEMa-LP
WM35
WM266-4

### Model No. 16

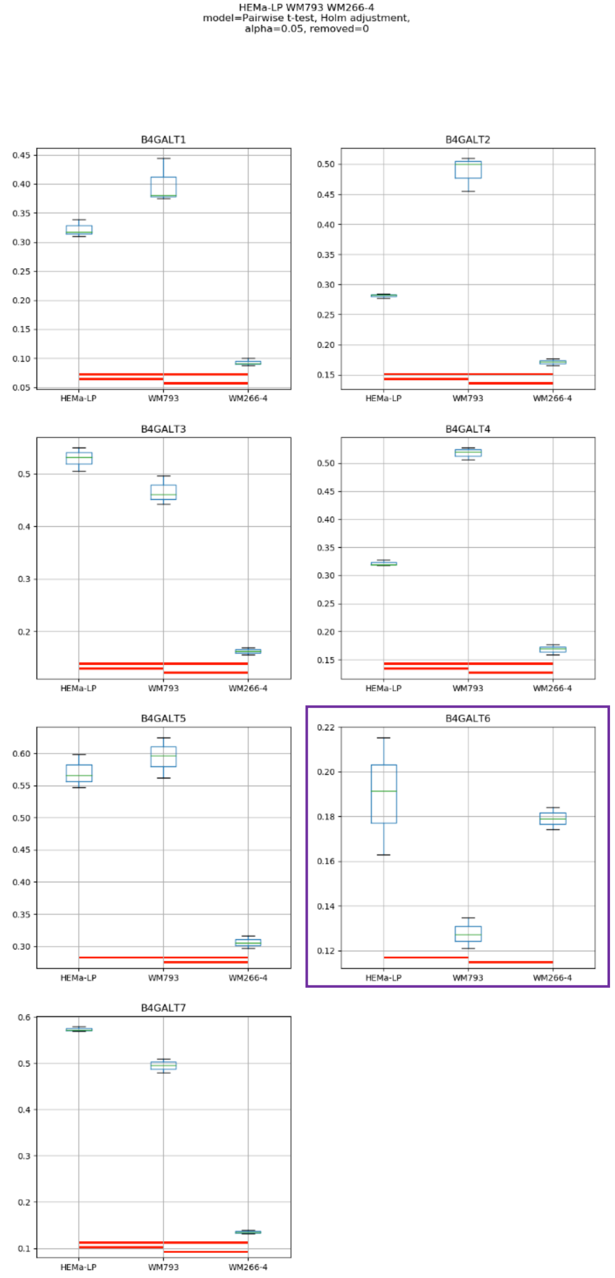
HEMa-LP WM793 WM266-4
model=Pairwise t-test, Holm adjustment,
alpha=0.05, removed=0
B4GALT1
B4GALT2
B4GALT3
B4GALT4
B4GALT5
B4GALT6
B4GALT7
HEMa-LP
WM793
WM266-4

### Model No. 17

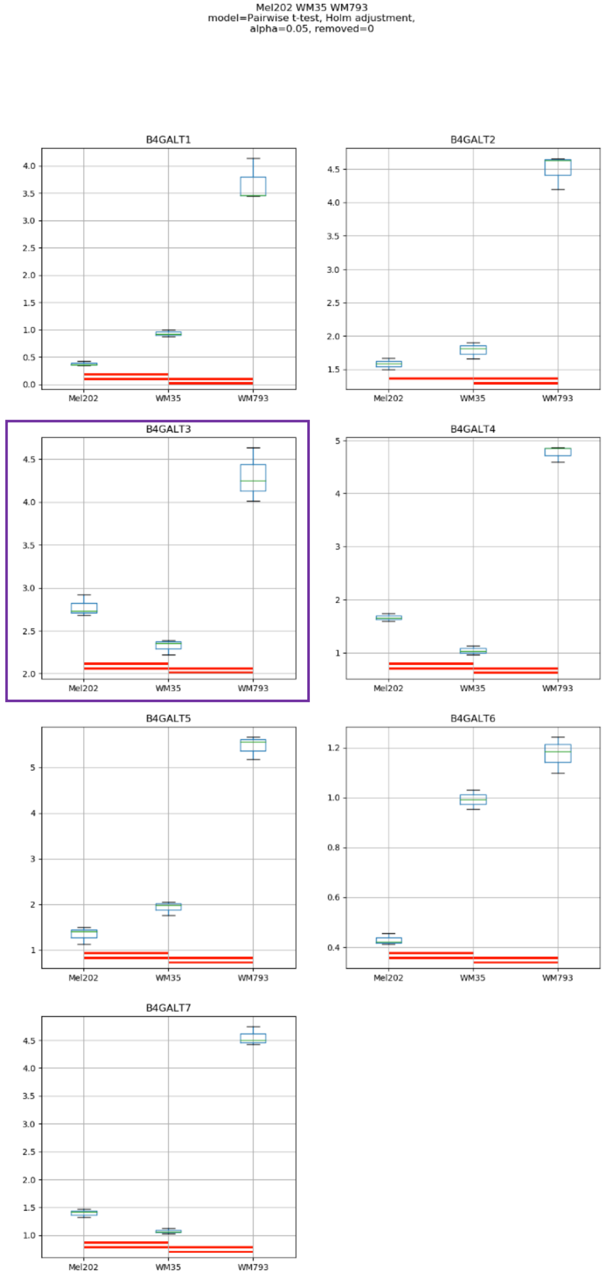
Mel202 WM35 WM793
model=Pairwise t-test, Holm adjustment,
alpha=0.05, removed=0
B4GALT1
B4GALT2
B4GALT3
B4GALT4
B4GALT5
B4GALT6
B4GALT7
Mel202
WM35
WM793

### Model No. 18

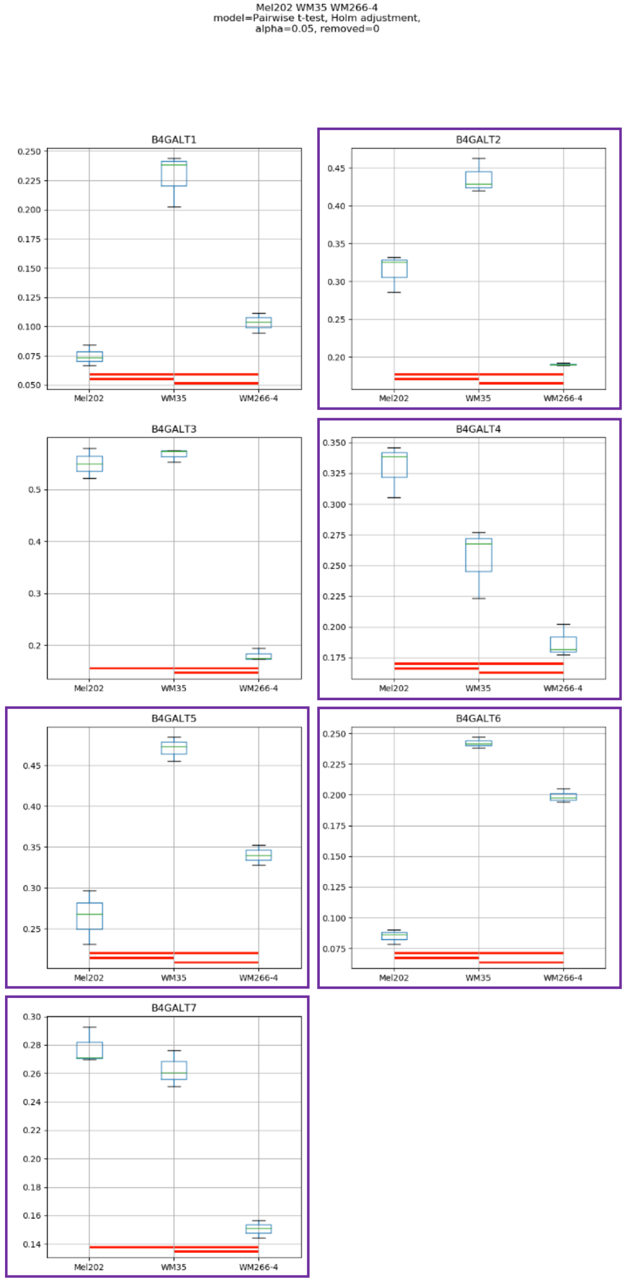

Mel202 WM35 WM266-4
model=Pairwise t-test, Holm adjustment,
alpha=0.05, removed=0
B4GALT1
B4GALT2
B4GALT3
B4GALT4
B4GALT5
B4GALT6
B4GALT7
Mel202
WM35
WM266-4

### Model No. 19

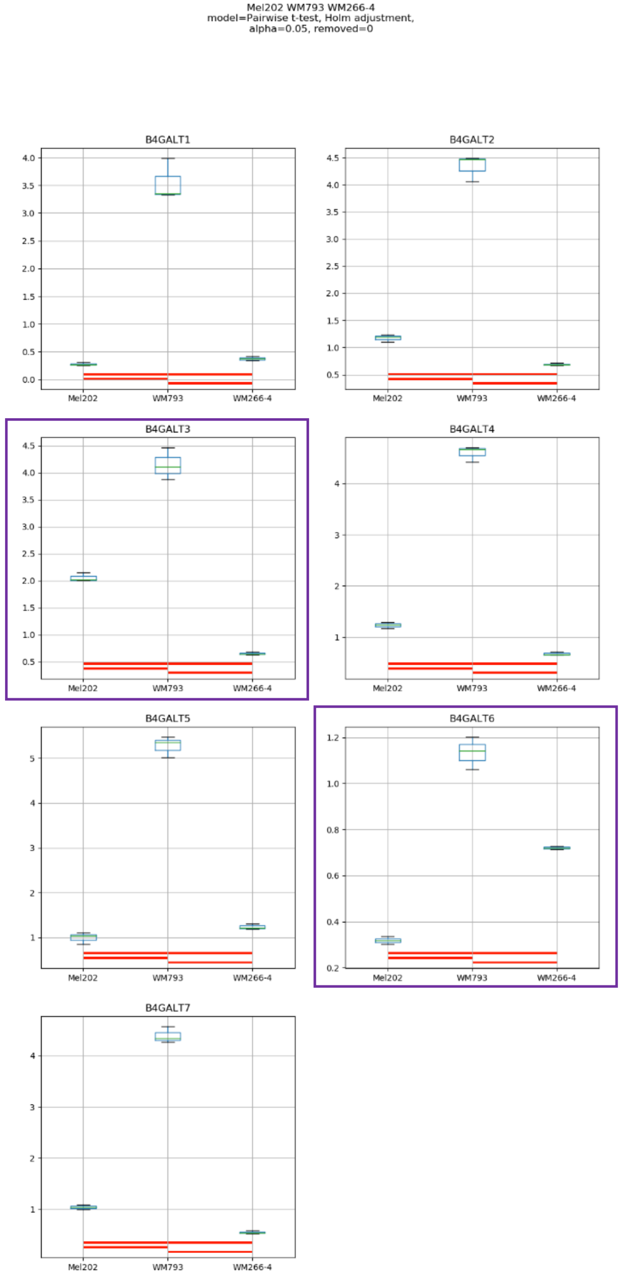
Mel202 WM793 WM266-4
model=Pairwise t-test, Holm adjustment,
alpha=0.05, removed=0
B4GALT1
B4GALT2
B4GALT3
B4GALT4
B4GALT5
B4GALT6
B4GALT7
Mel202
WM793
WM266-4

Model No. 20

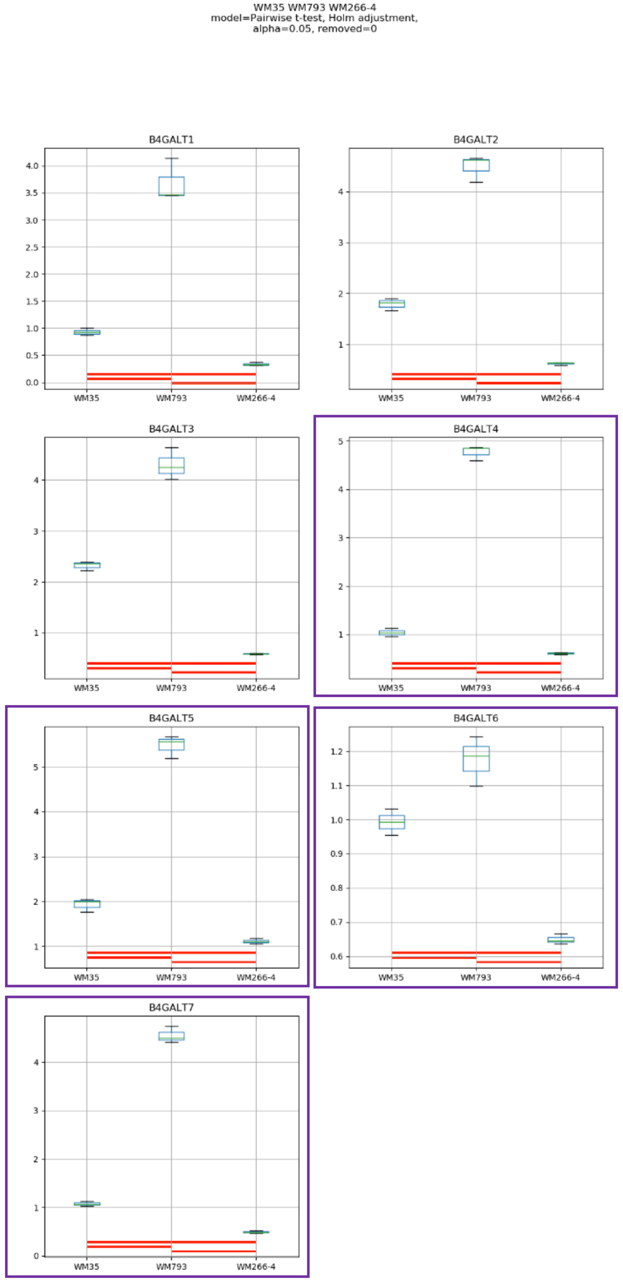
WM35 WM793 WM266-4
model=Pairwise t-test, Holm adjustment,
alpha=0.05, removed=0
B4GALT1
B4GALT2
B4GALT3
B4GALT4
B4GALT5
B4GALT6
B4GALT7
WM35
WM793
WM266-4

Model No. 21

HEMa-LP Mel202 WM35 WM793
model=Pairwise t-test, Holm adjustment,
alpha=0.05, removed=0

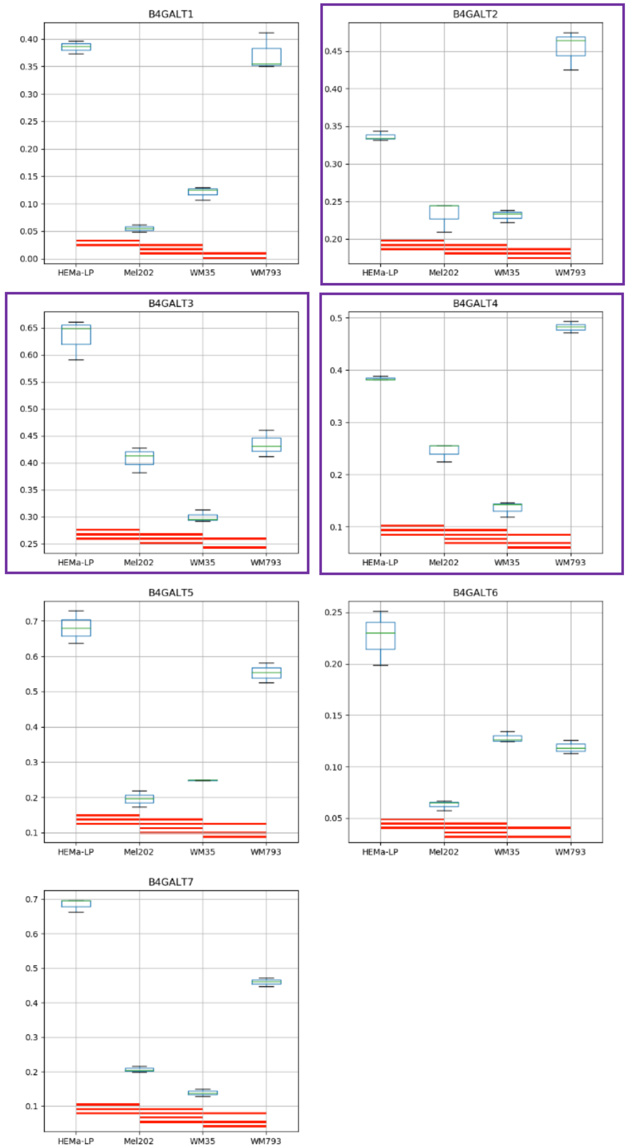

Model No. 22

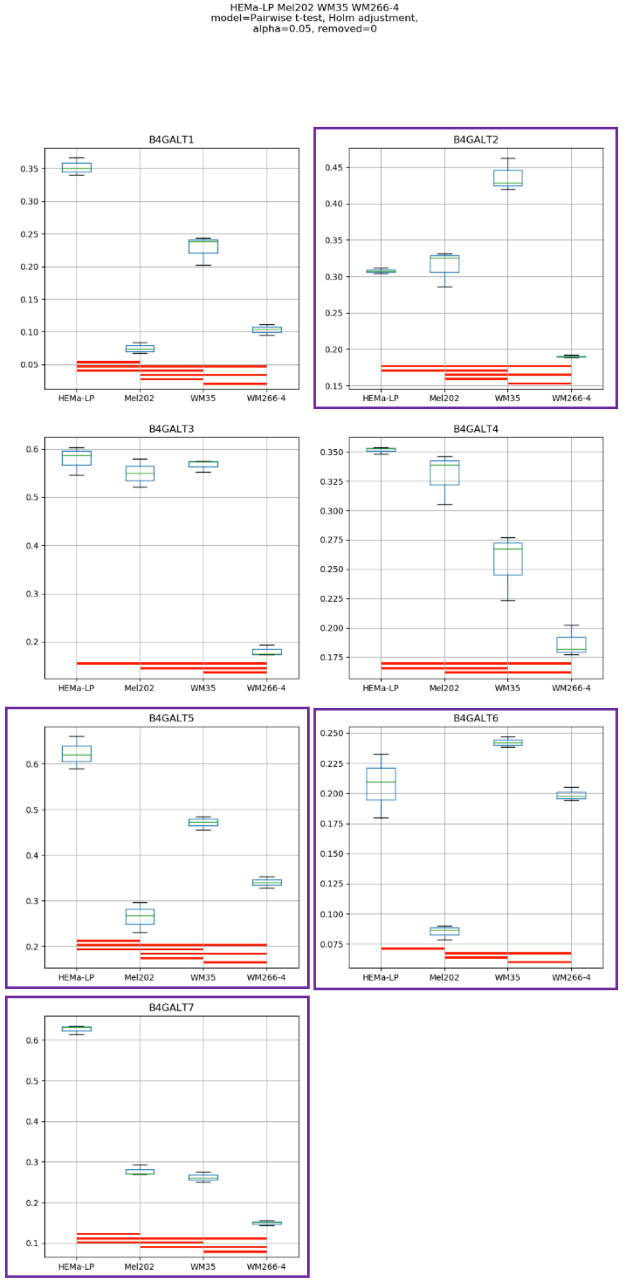
HEMa-LP Mel202 WM35 WM266-4
model=Pairwise t-test, Holm adjustment,
alpha=0.05, removed=0
B4GALT1
B4GALT2
B4GALT3
B4GALT4
B4GALT5
B4GALT6
B4GALT7
HEMa-LP
Mel202
WM35
WM266-4

Model No. 23

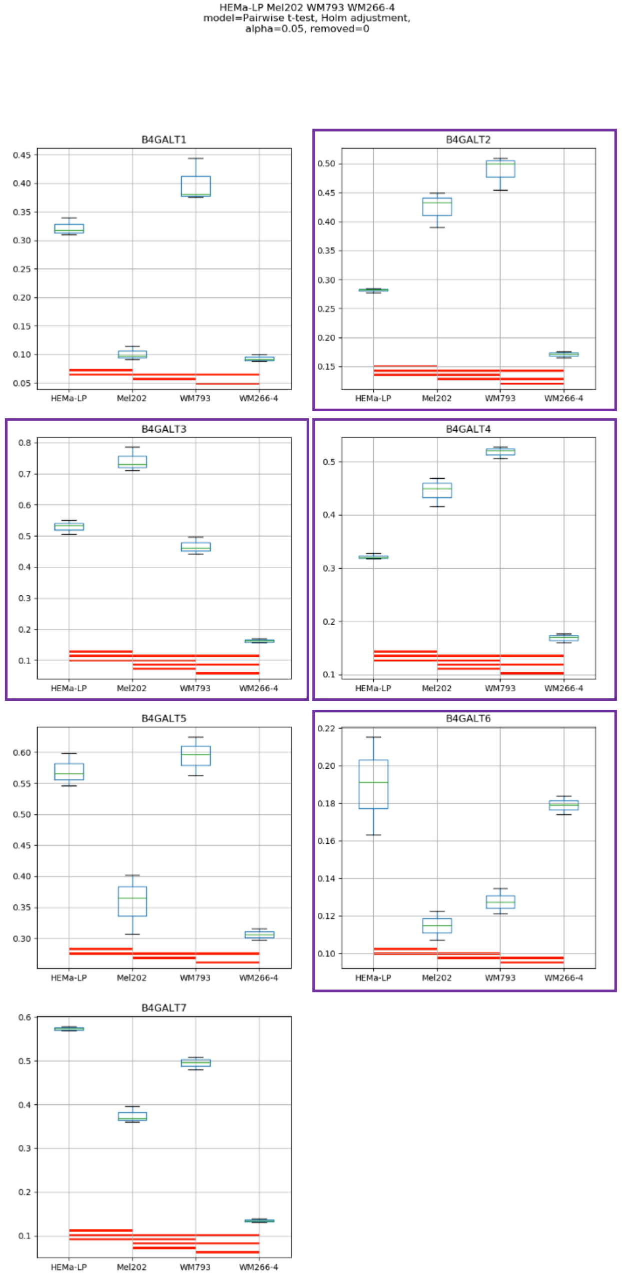
HEMa-LP Mel202 WM793 WM266-4
model=Pairwise t-test, Holm adjustment,
alpha=0.05, removed=0
B4GALT1
B4GALT2
B4GALT3
B4GALT4
B4GALT5
B4GALT6
B4GALT7
HEMa-LP
Mel202
WM793
WM266-4

### Model No. 24

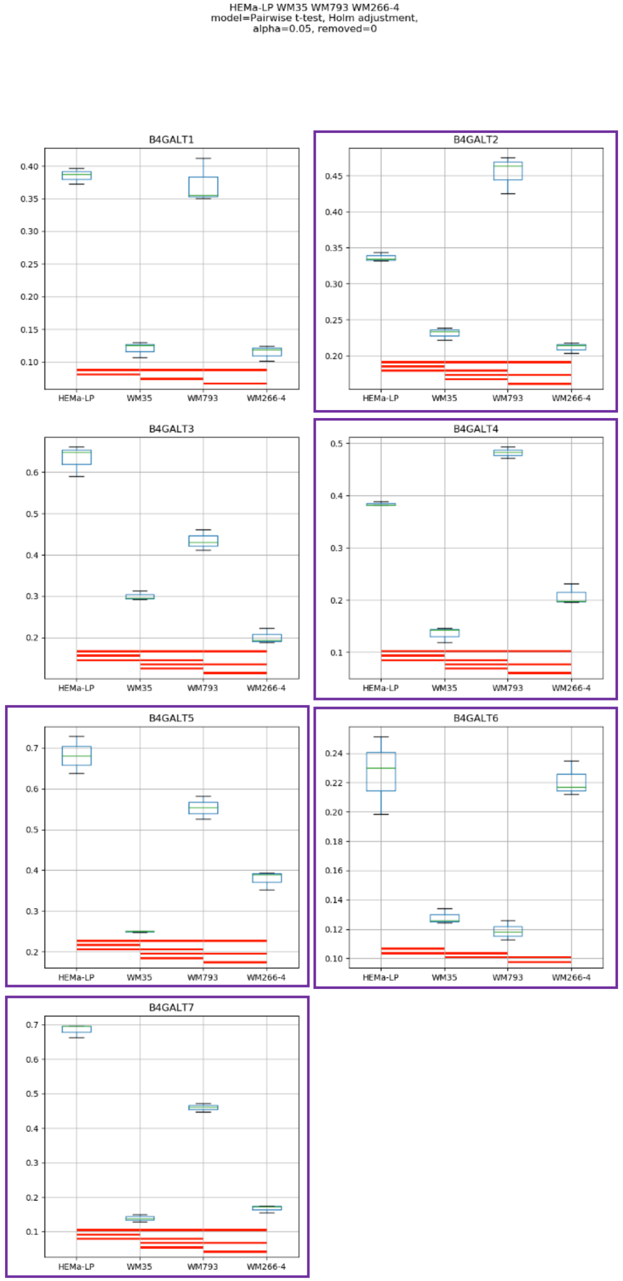

HEMa-LP WM35 WM793 WM266-4
model=Pairwise t-test, Holm adjustment,
alpha=0.05, removed=0
B4GALT1
B4GALT2
B4GALT3
B4GALT4
B4GALT5
B4GALT6
B4GALT7
HEMa-LP
WM35
WM793
WM266-4

### Model No. 25

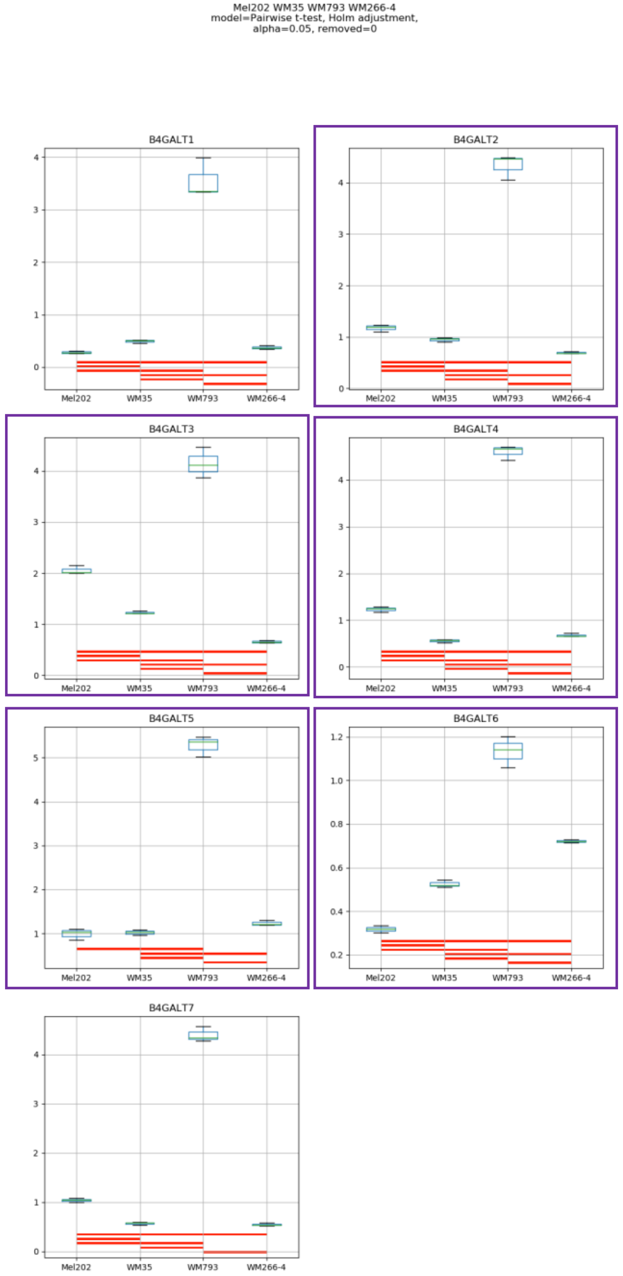
Mel202 WM35 WM793 WM266-4
model=Pairwise t-test, Holm adjustment,
alpha=0.05, removed=0
B4GALT1
B4GALT2
B4GALT3
B4GALT4
B4GALT5
B4GALT6
B4GALT7
Mel202
WM35
WM793
WM266-4

### Model No. 26

HEMa-LP Mel202 WM35 WM793 WM266-4
model=Pairwise t-test, Holm adjustment,
alpha=0.05, removed=0

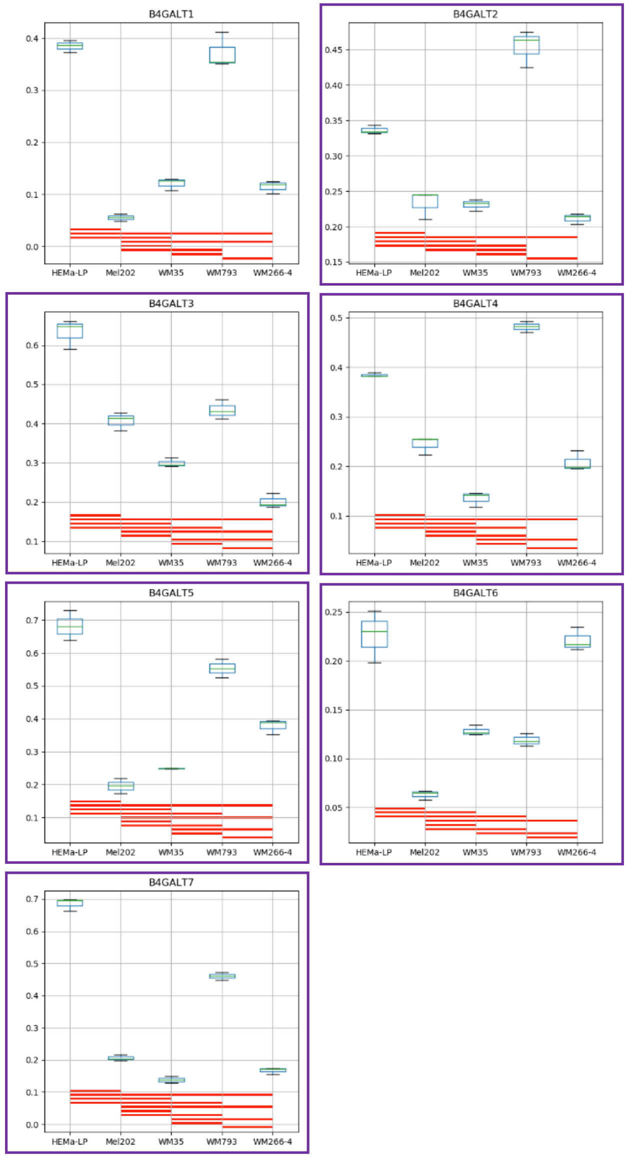

**Supplementary Figure S11:** Medians of relative quantity (RQ) for target genes *B4GALT1–B4GALT7* normalized to the reference gene or pair of reference genes with the best stability values resulting from GenExpA analysis based on set 2 of four candidate reference genes (*RPS23*, *SNRPA*, *HPRT1*, *GUSB*) and remove repetition level 0. The statistical analyses used the pairwise t-test with Holm adjustment. Red line represents statistical significance, $P < 0.05$. Frames denote box-plots representing the medians of the RQ values for a given target gene without a repeatable pattern in the same cell lines under different models (the 'Remove repetitions' box set to 0). Detailed results are listed in Supplementary Table 10.

### Model No. 1

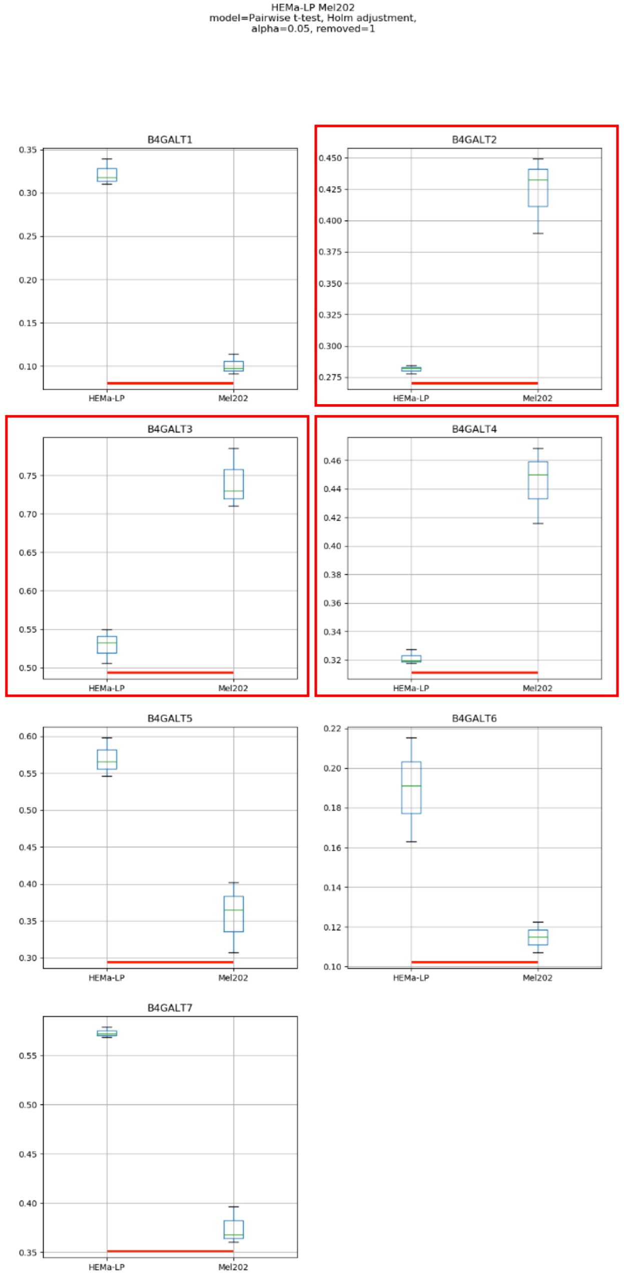

HEMa-LP Mel202
model=Pairwise t-test, Holm adjustment,
alpha=0.05, removed=1
B4GALT1
B4GALT2
B4GALT3
B4GALT4
B4GALT5
B4GALT6
B4GALT7
HEMa-LP
Mel202

Model No. 2

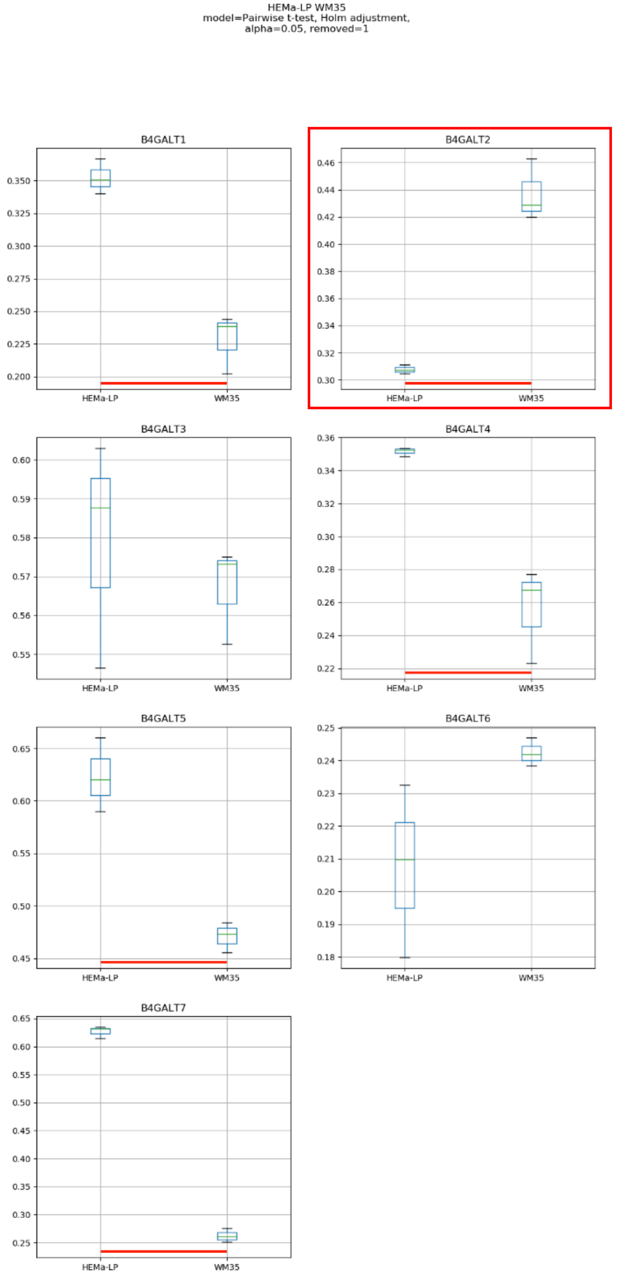
HEMa-LP WM35
model=Pairwise t-test, Holm adjustment,
alpha=0.05, removed=1
B4GALT1
B4GALT2
B4GALT3
B4GALT4
B4GALT5
B4GALT6
B4GALT7
HEMa-LP
WM35

Model No. 3

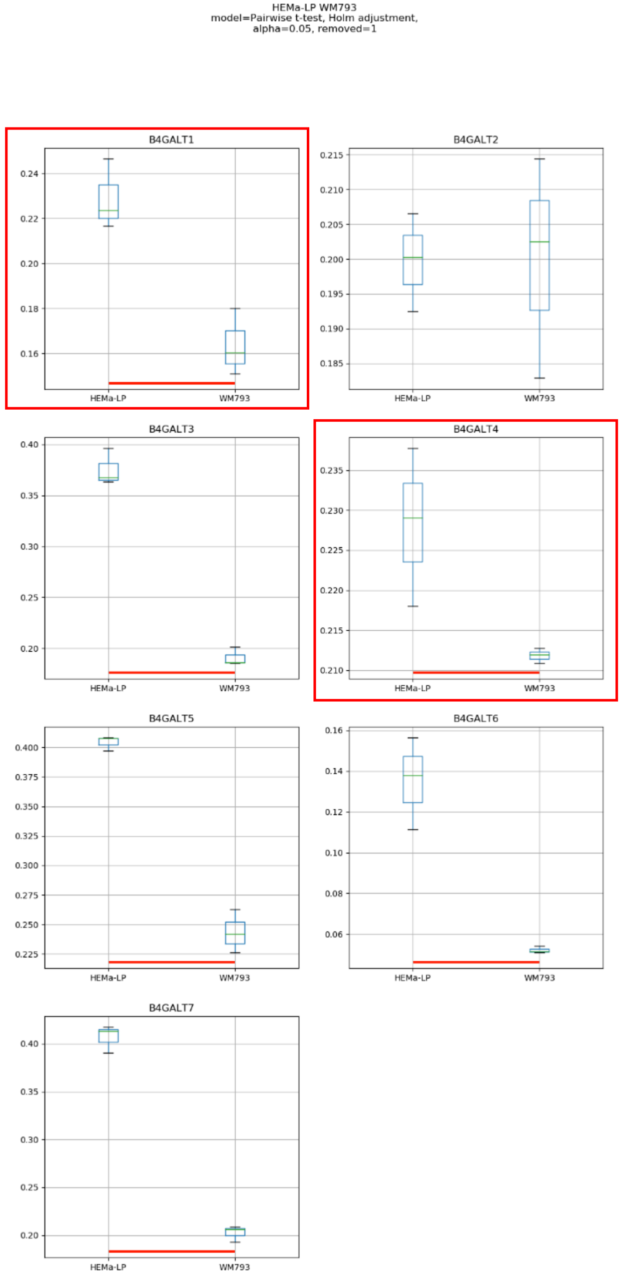
HEMa-LP WM793
model=Pairwise t-test, Holm adjustment,
alpha=0.05, removed=1
B4GALT1
B4GALT2
B4GALT3
B4GALT4
B4GALT5
B4GALT6
B4GALT7
HEMa-LP
WM793

### Model No. 4

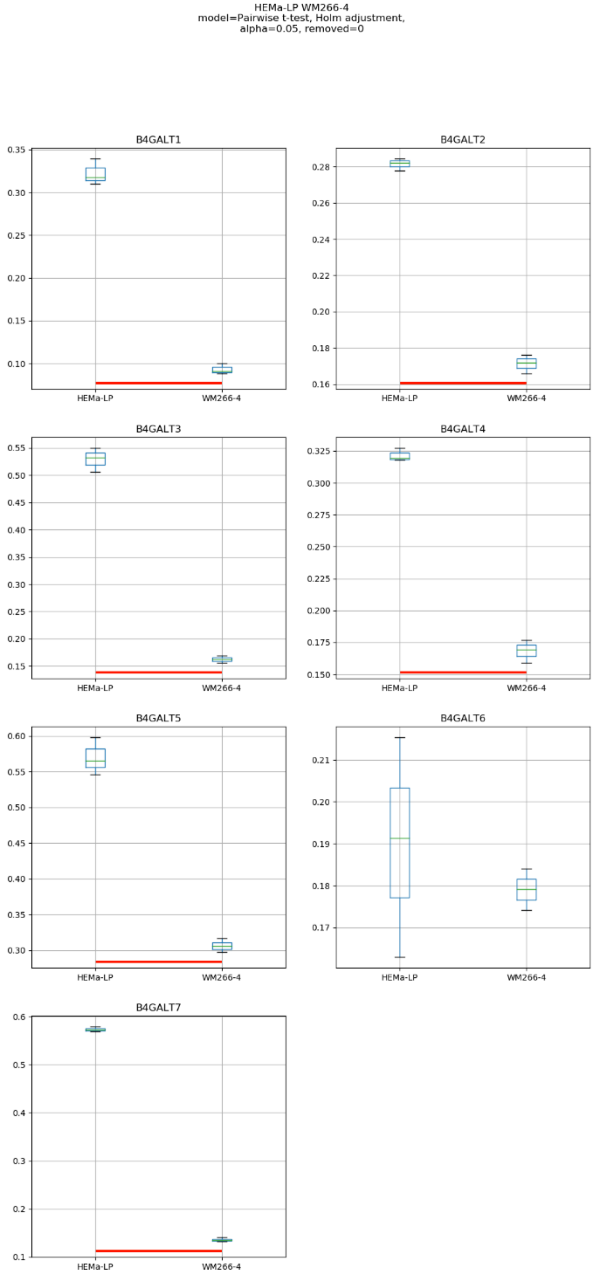
HEMa-LP WM266-4
model=Pairwise t-test, Holm adjustment,
alpha=0.05, removed=0
B4GALT1
B4GALT2
B4GALT3
B4GALT4
B4GALT5
B4GALT6
B4GALT7
HEMa-LP
WM266-4

### Model No. 5

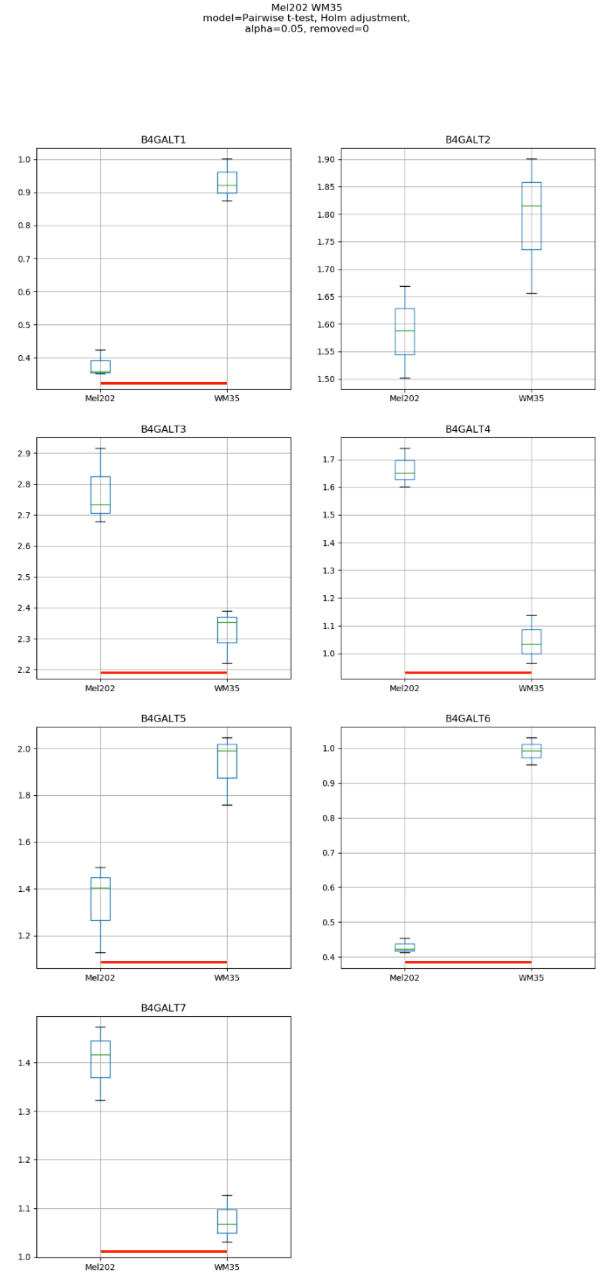

Mel202 WM35
model=Pairwise t-test, Holm adjustment,
alpha=0.05, removed=0
B4GALT1
B4GALT2
B4GALT3
B4GALT4
B4GALT5
B4GALT6
B4GALT7
Mel202
WM35

Model No. 6

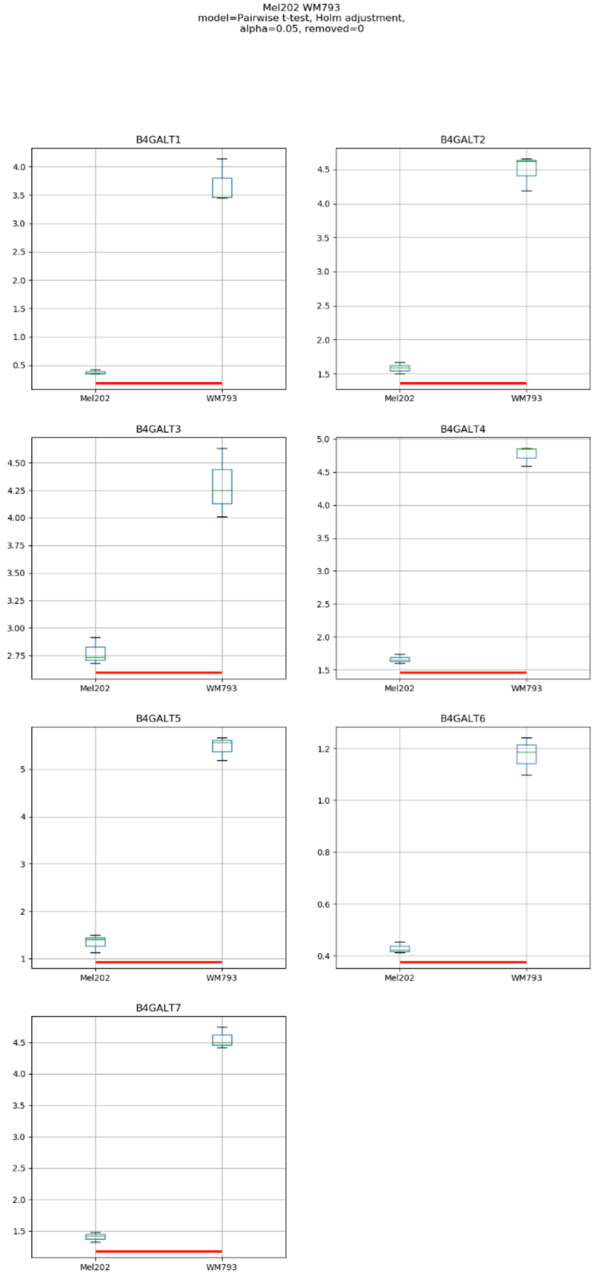
Mel202 WM793
model=Pairwise t-test, Holm adjustment,
alpha=0.05, removed=0
B4GALT1
B4GALT2
B4GALT3
B4GALT4
B4GALT5
B4GALT6
B4GALT7
Mel202
WM793

### Model No. 7

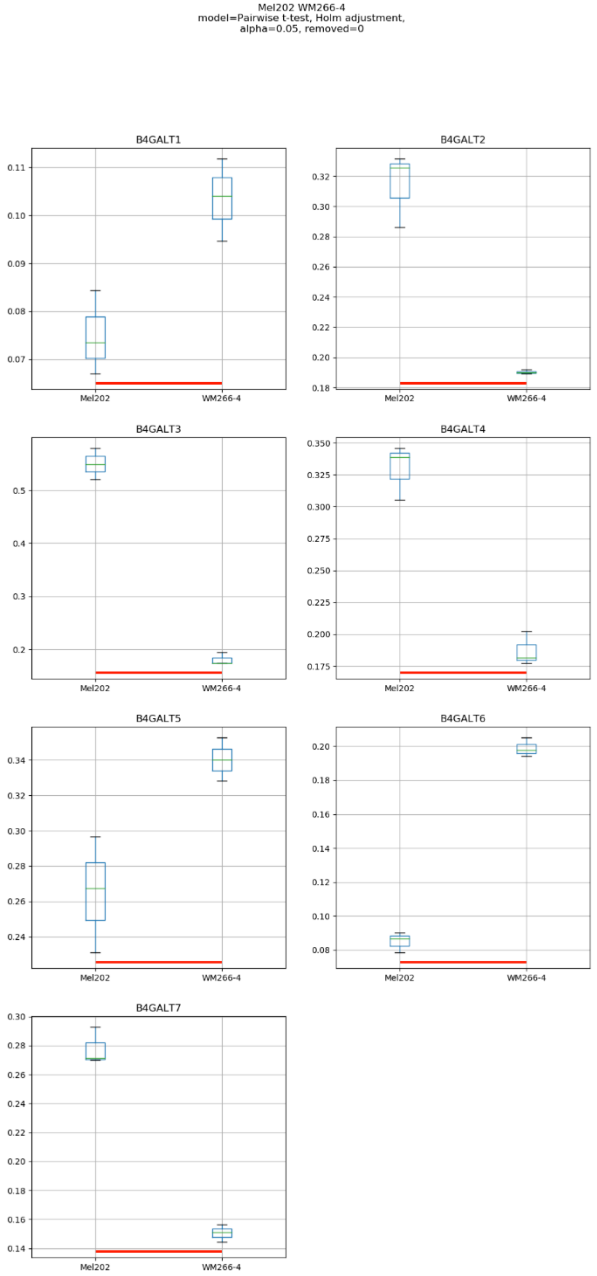
Mel202 WM266-4
model=Pairwise t-test, Holm adjustment,
alpha=0.05, removed=0
B4GALT1
B4GALT2
B4GALT3
B4GALT4
B4GALT5
B4GALT6
B4GALT7
Mel202
WM266-4

### Model No. 8

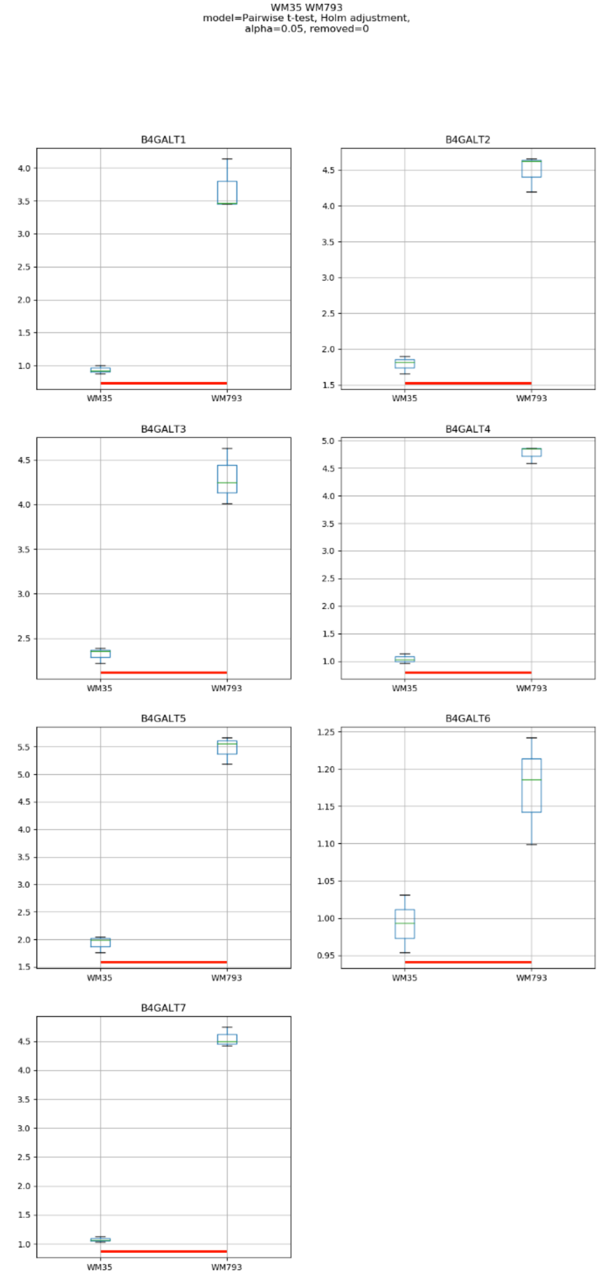
WM35 WM793
model=Pairwise t-test, Holm adjustment,
alpha=0.05, removed=0
B4GALT1
B4GALT2
B4GALT3
B4GALT4
B4GALT5
B4GALT6
B4GALT7
WM35
WM793

### Model No. 9

WM35 WM266-4
model=Pairwise t-test, Holm adjustment,
alpha=0.05, removed=0

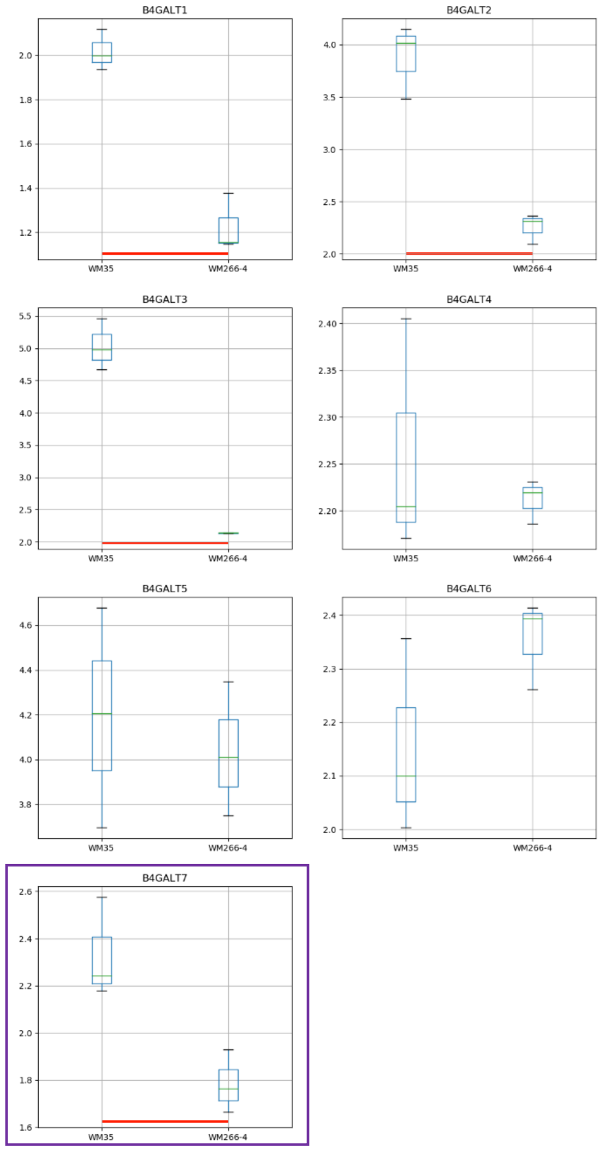

### Model No. 10

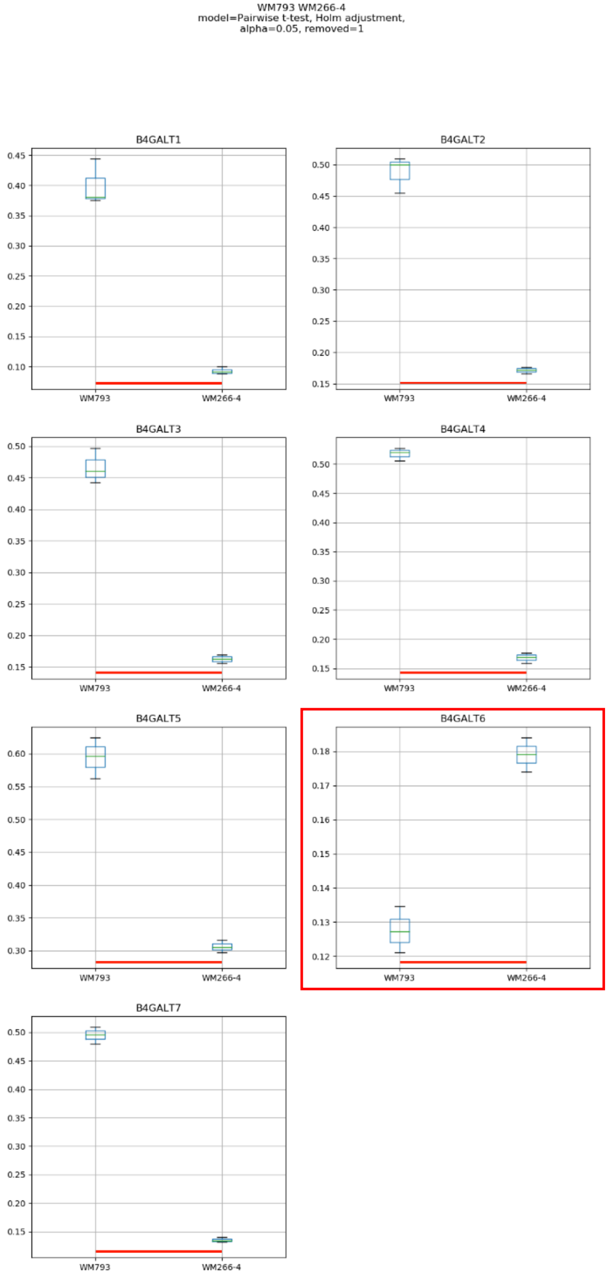

WM793 WM266-4
model=Pairwise t-test, Holm adjustment,
alpha=0.05, removed=1
B4GALT1
B4GALT2
B4GALT3
B4GALT4
B4GALT5
B4GALT6
B4GALT7
WM793
WM266-4

### Model No. 11

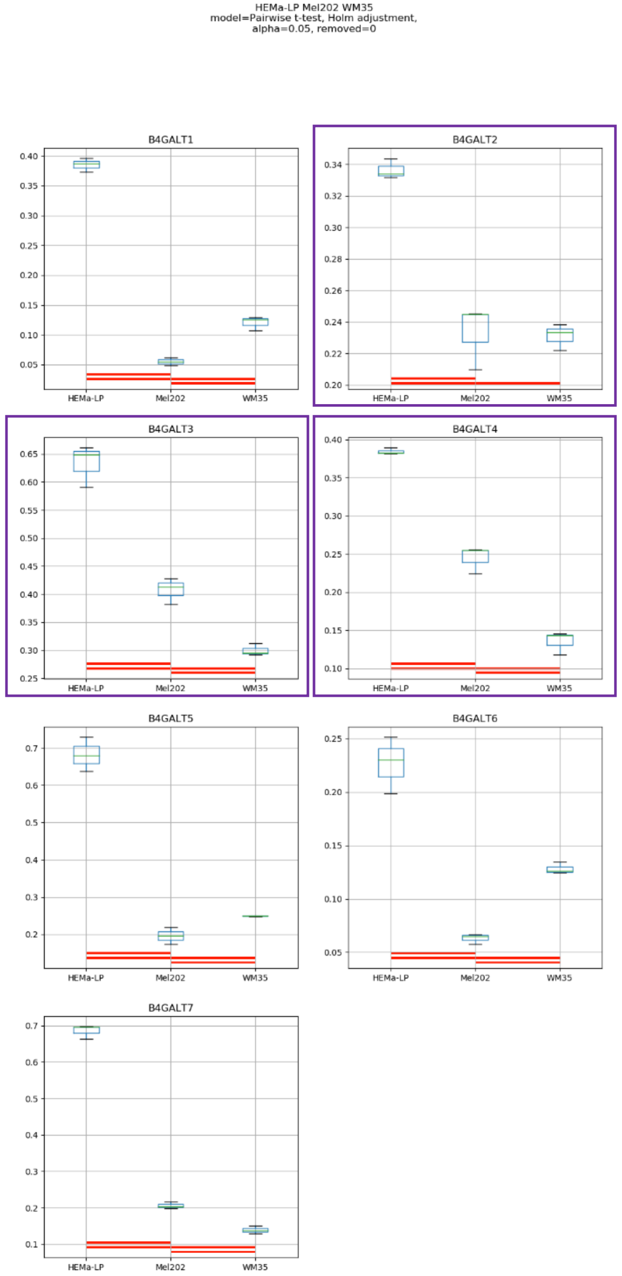
HEMa-LP Mel202 WM35
model=Pairwise t-test, Holm adjustment,
alpha=0.05, removed=0
B4GALT1
B4GALT2
B4GALT3
B4GALT4
B4GALT5
B4GALT6
B4GALT7
HEMa-LP
Mel202
WM35

### Model No. 12

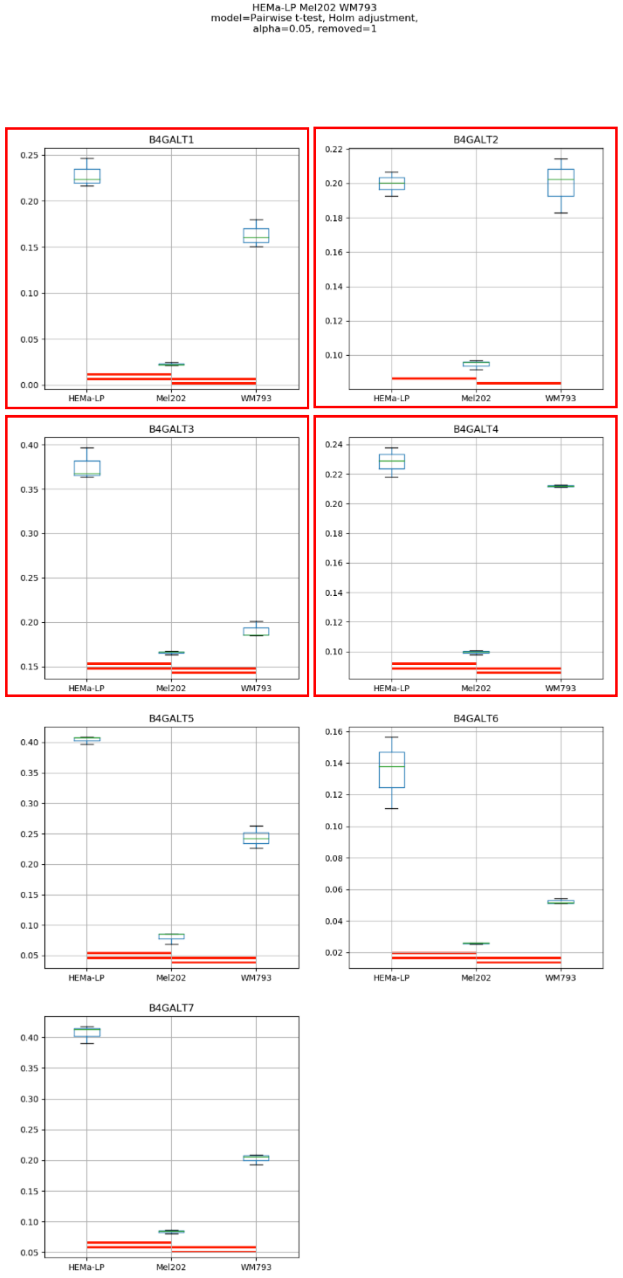
HEMa-LP Mel202 WM793
model=Pairwise t-test, Holm adjustment,
alpha=0.05, removed=1
B4GALT1
B4GALT2
B4GALT3
B4GALT4
B4GALT5
B4GALT6
B4GALT7
HEMa-LP
Mel202
WM793

### Model No. 13

HEMa-LP Mel202 WM266-4
model=Pairwise t-test, Holm adjustment,
alpha=0.05, removed=1

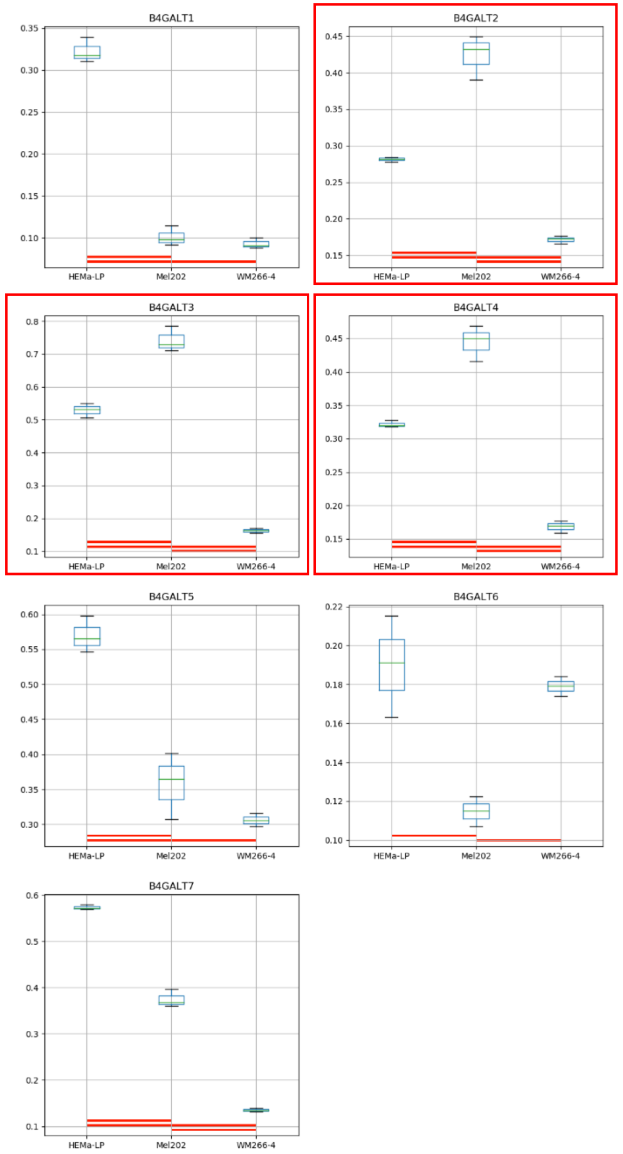

### Model No. 14

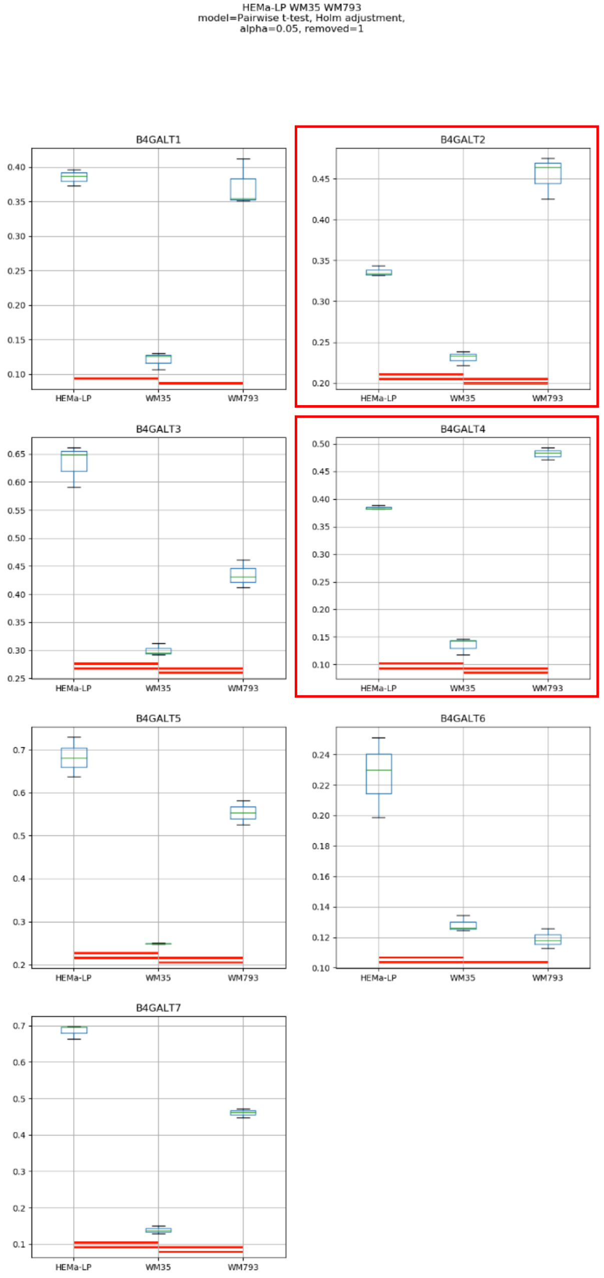
HEMa-LP WM35 WM793
model=Pairwise t-test, Holm adjustment,
alpha=0.05, removed=1
B4GALT1
B4GALT2
B4GALT3
B4GALT4
B4GALT5
B4GALT6
B4GALT7
HEMa-LP
WM35
WM793

### Model No. 15

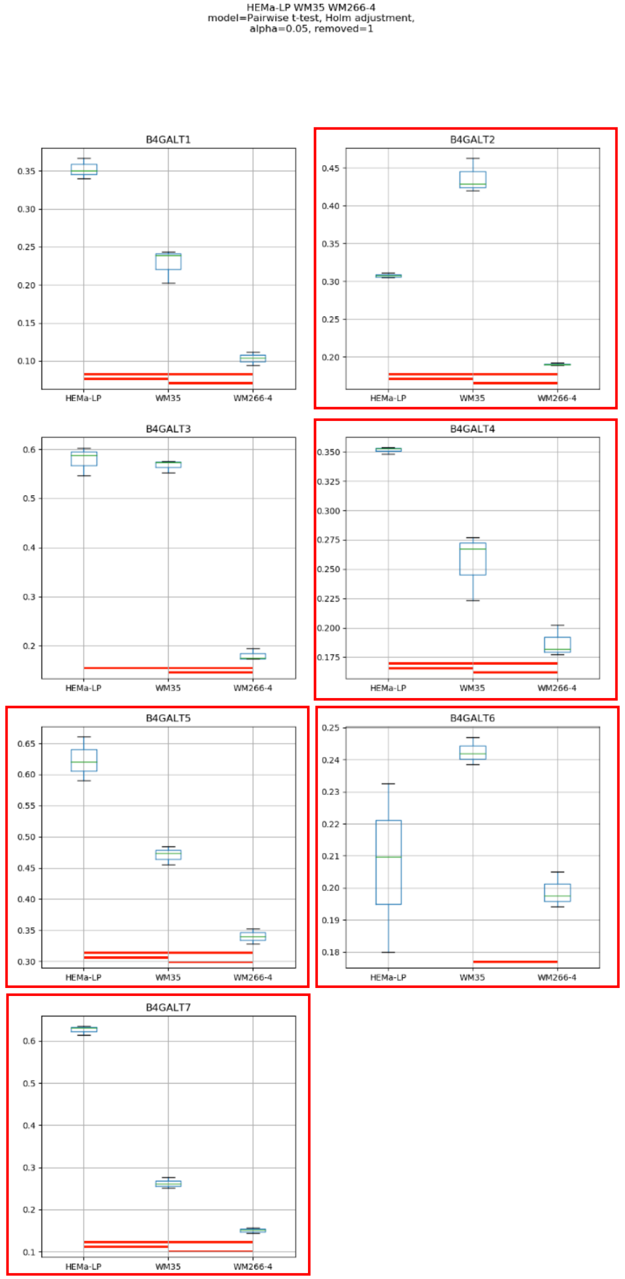

HEMa-LP WM35 WM266-4
model=Pairwise t-test, Holm adjustment,
alpha=0.05, removed=1
B4GALT1
B4GALT2
B4GALT3
B4GALT4
B4GALT5
B4GALT6
B4GALT7
HEMa-LP
WM35
WM266-4

### Model No. 16

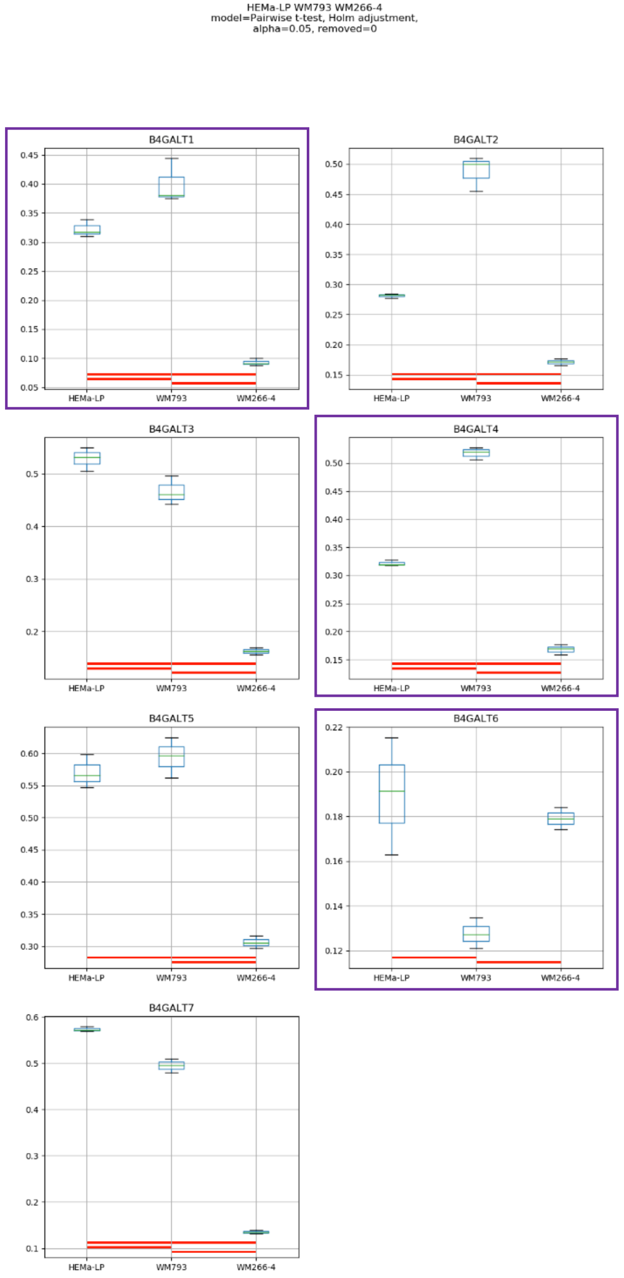
HEMa-LP WM793 WM266-4
model=Pairwise t-test, Holm adjustment,
alpha=0.05, removed=0
B4GALT1
B4GALT2
B4GALT3
B4GALT4
B4GALT5
B4GALT6
B4GALT7
HEMa-LP
WM793
WM266-4

Model No. 17

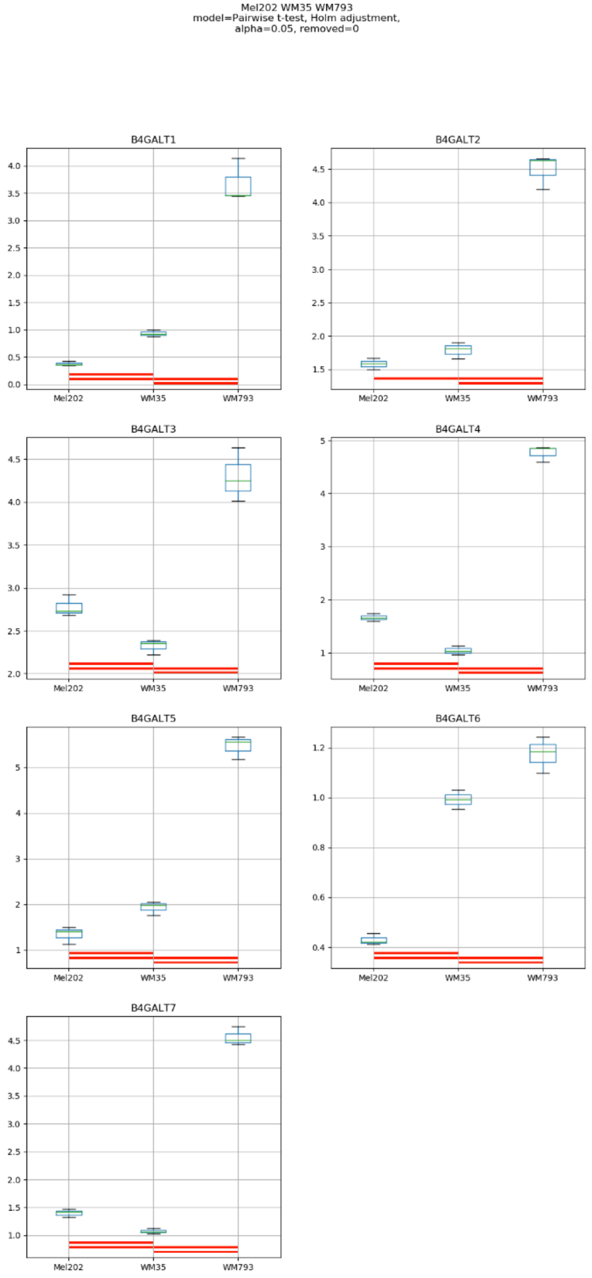
Mel202 WM35 WM793
model=Pairwise t-test, Holm adjustment,
alpha=0.05, removed=0
B4GALT1
B4GALT2
B4GALT3
B4GALT4
B4GALT5
B4GALT6
B4GALT7
Mel202
WM35
WM793

Model No. 18

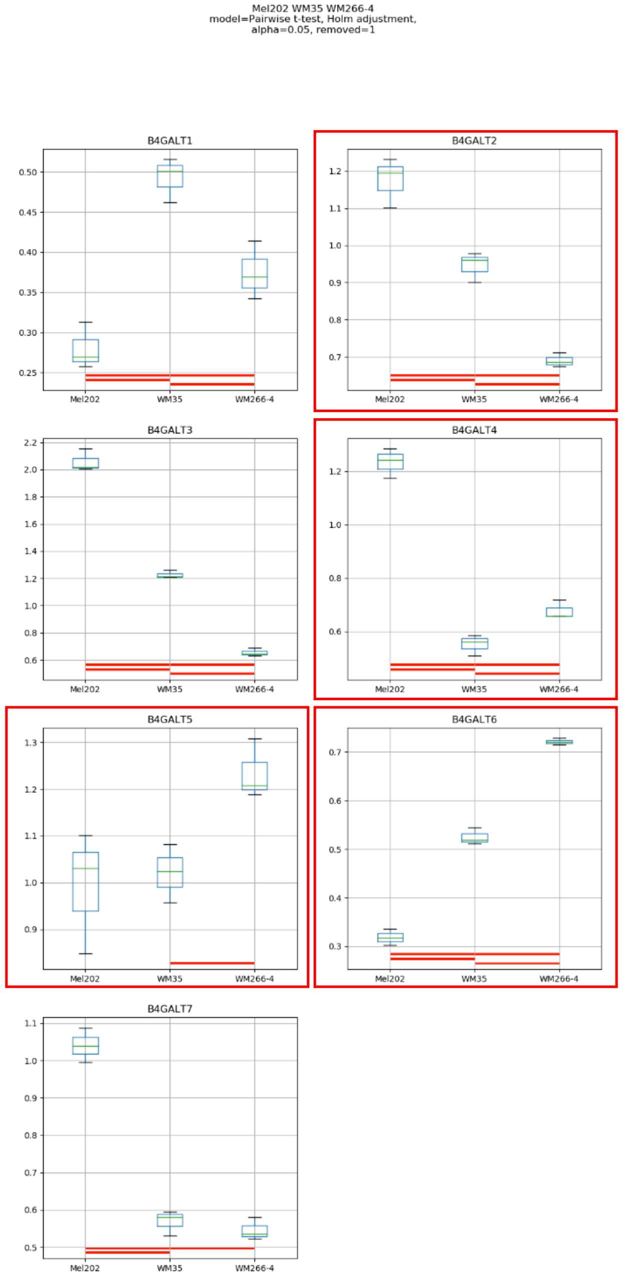
Mel202 WM35 WM266-4
model=Pairwise t-test, Holm adjustment,
alpha=0.05, removed=1
B4GALT1
B4GALT2
B4GALT3
B4GALT4
B4GALT5
B4GALT6
B4GALT7
Mel202
WM35
WM266-4

Model No. 19

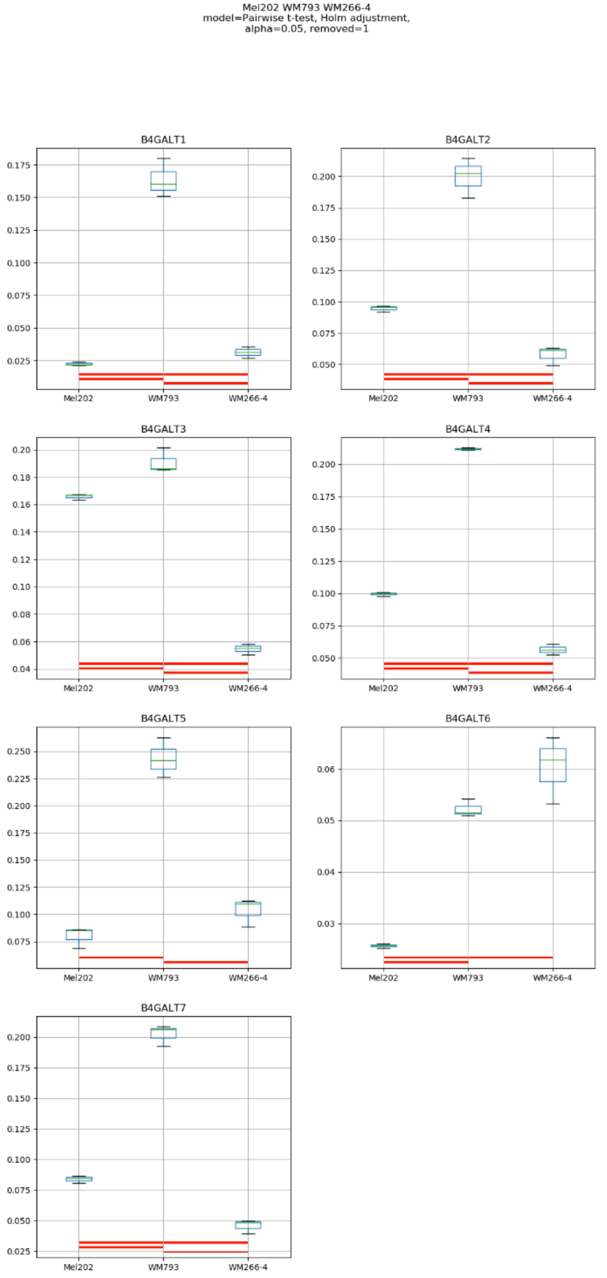

Mel202 WM793 WM266-4
model=Pairwise t-test, Holm adjustment,
alpha=0.05, removed=1
B4GALT1
B4GALT2
B4GALT3
B4GALT4
B4GALT5
B4GALT6
B4GALT7
Mel202
WM793
WM266-4

### Model No. 20

WM35 WM793 WM266-4
model=Pairwise t-test, Holm adjustment,
alpha=0.05, removed=1

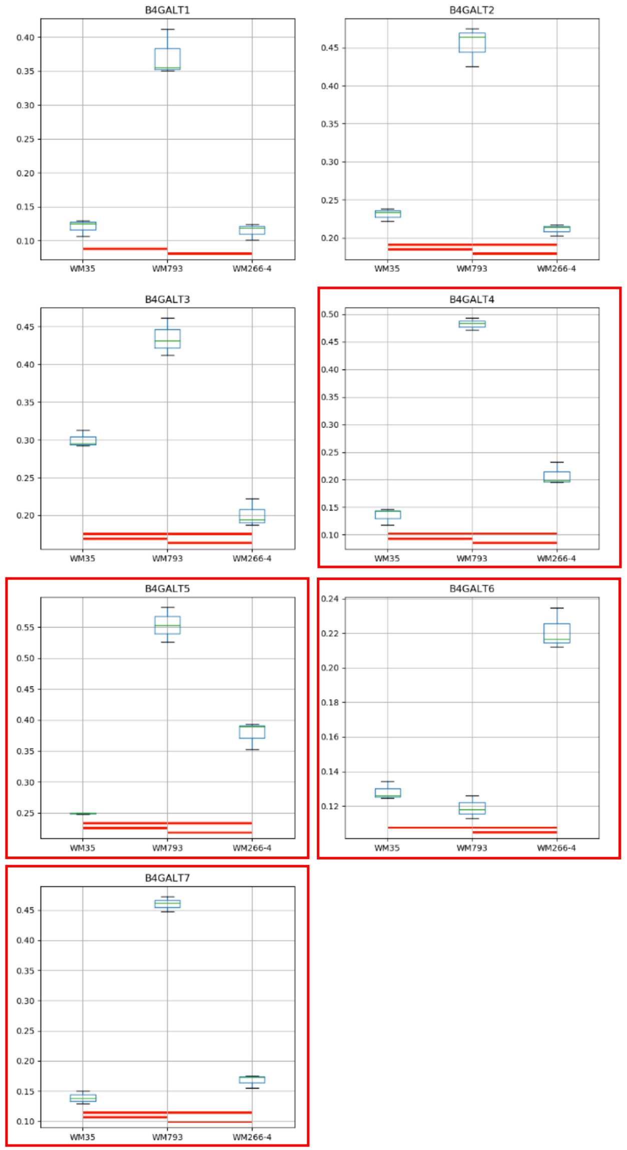

### Model No. 21

HEMa-LP Mel202 WM35 WM793
model=Pairwise t-test, Holm adjustment,
alpha=0.05, removed=1

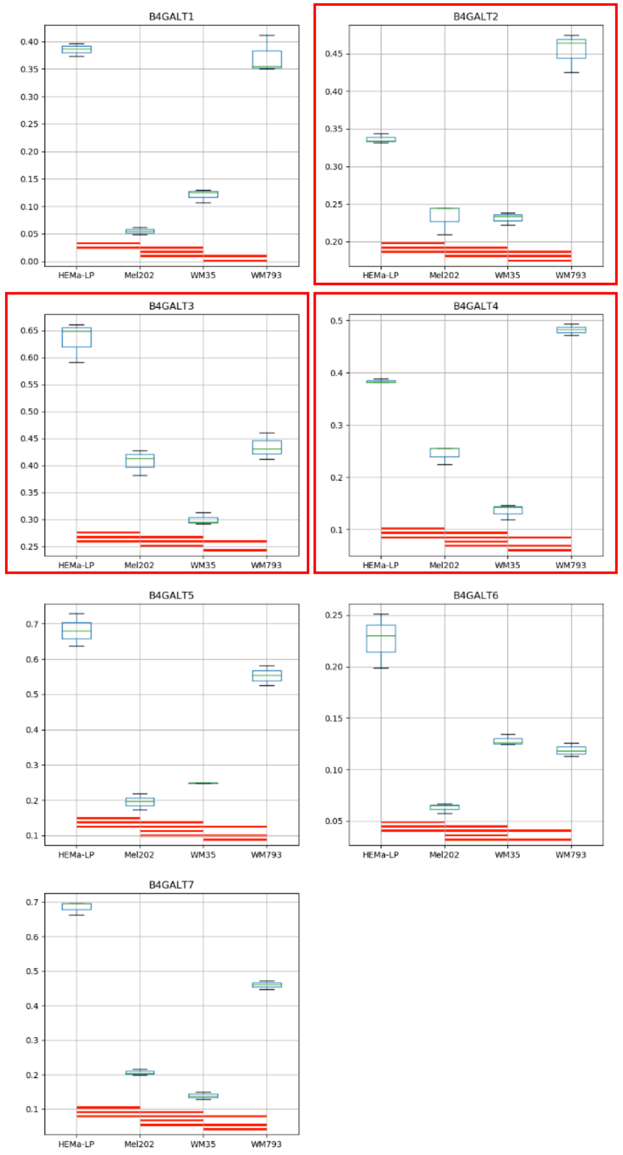

### Model No. 22

HEMa-LP Mel202 WM35 WM266-4
model=Pairwise t-test, Holm adjustment,
alpha=0.05, removed=0

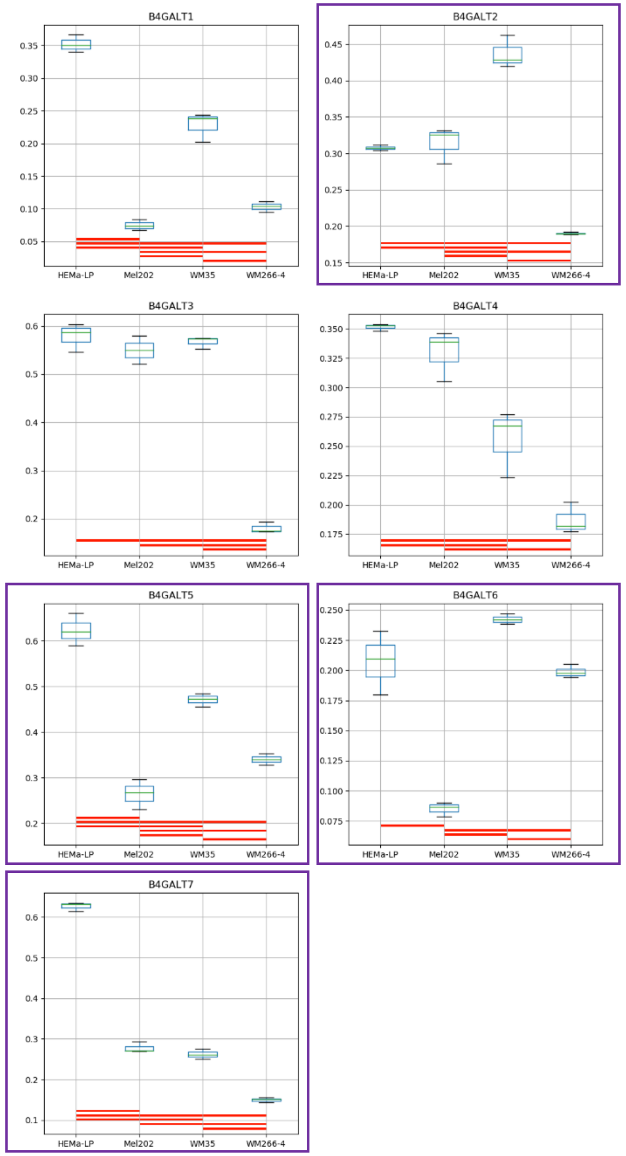

Model No. 23

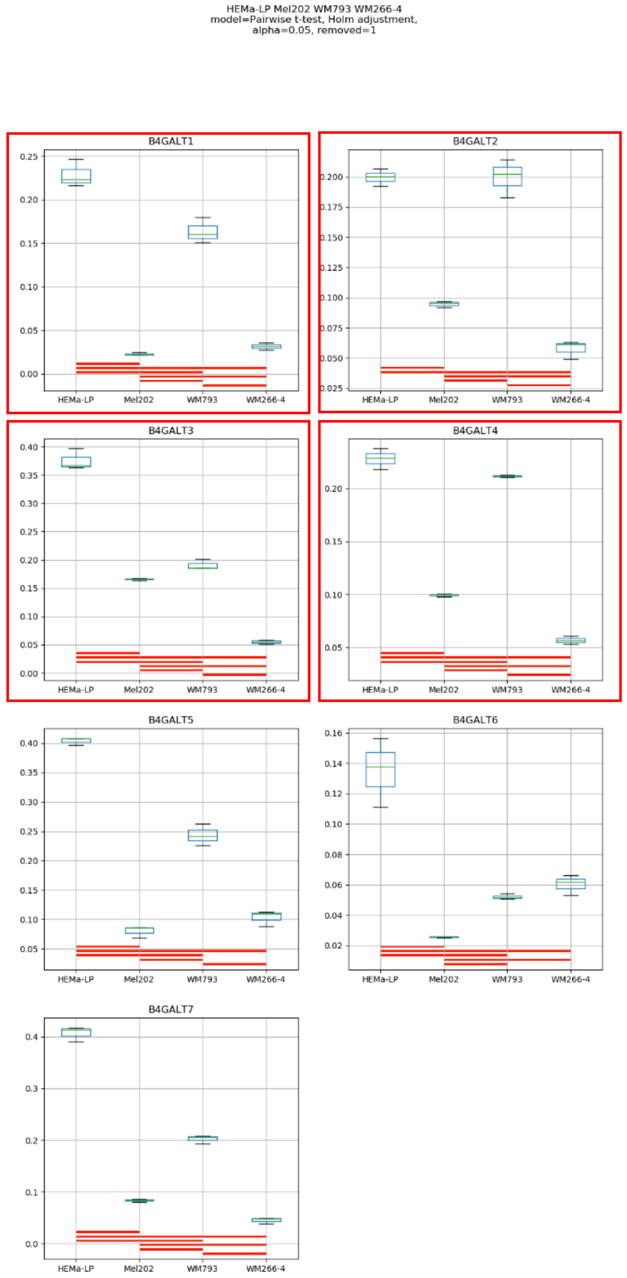
HEMa-LP Mel202 WM793 WM266-4
model=Pairwise t-test, Holm adjustment,
alpha=0.05, removed=1
B4GALT1
B4GALT2
B4GALT3
B4GALT4
B4GALT5
B4GALT6
B4GALT7
HEMa-LP
Mel202
WM793
WM266-4

Model No. 24

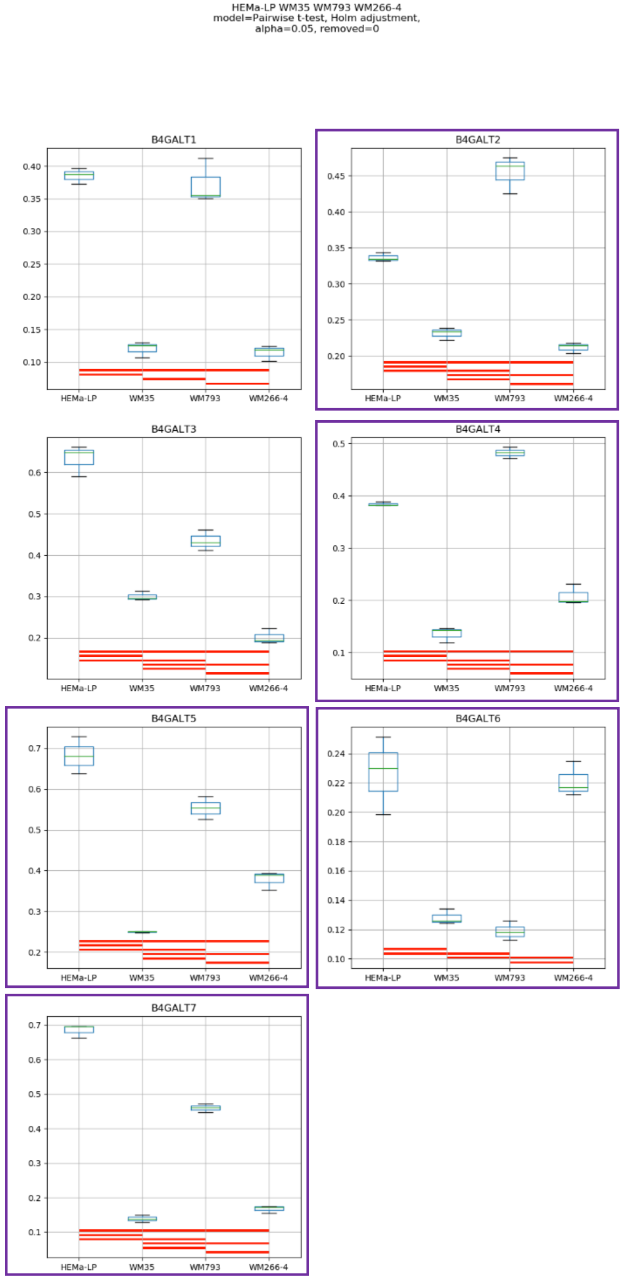
HEMa-LP WM35 WM793 WM266-4
model=Pairwise t-test, Holm adjustment,
alpha=0.05, removed=0
B4GALT1
B4GALT2
B4GALT3
B4GALT4
B4GALT5
B4GALT6
B4GALT7
HEMa-LP
WM35
WM793
WM266-4

### Model No. 25

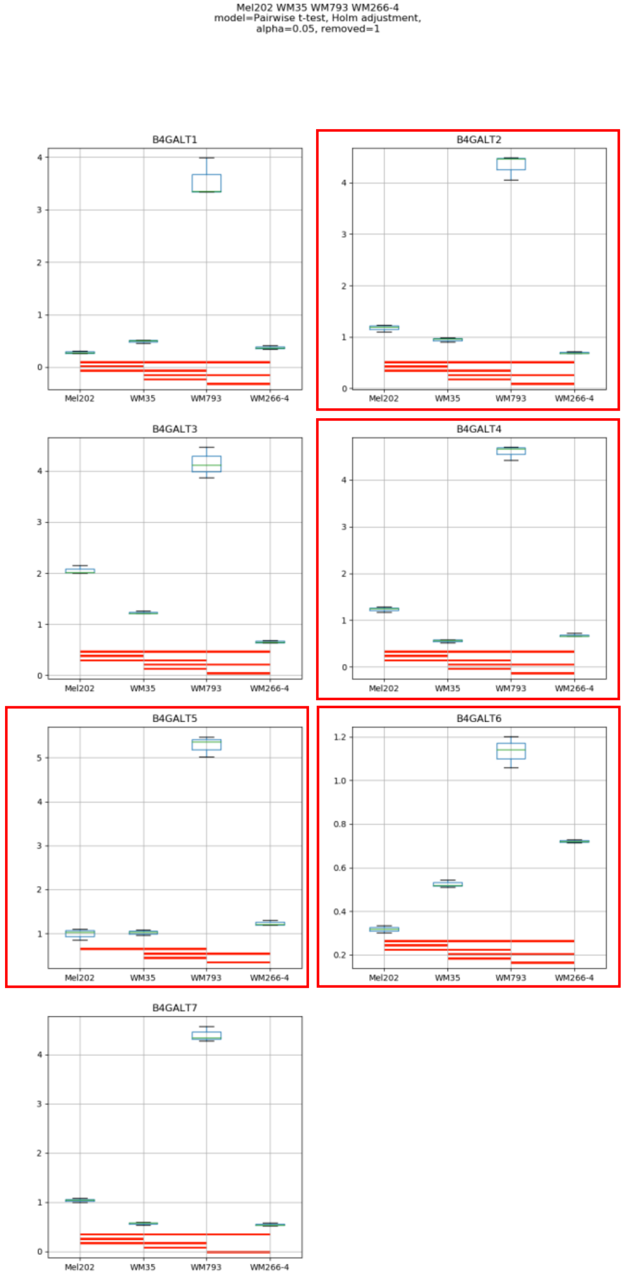
Mel202 WM35 WM793 WM266-4
model=Pairwise t-test, Holm adjustment,
alpha=0.05, removed=1
B4GALT1
B4GALT2
B4GALT3
B4GALT4
B4GALT5
B4GALT6
B4GALT7
Mel202
WM35
WM793
WM266-4

### Model No. 26

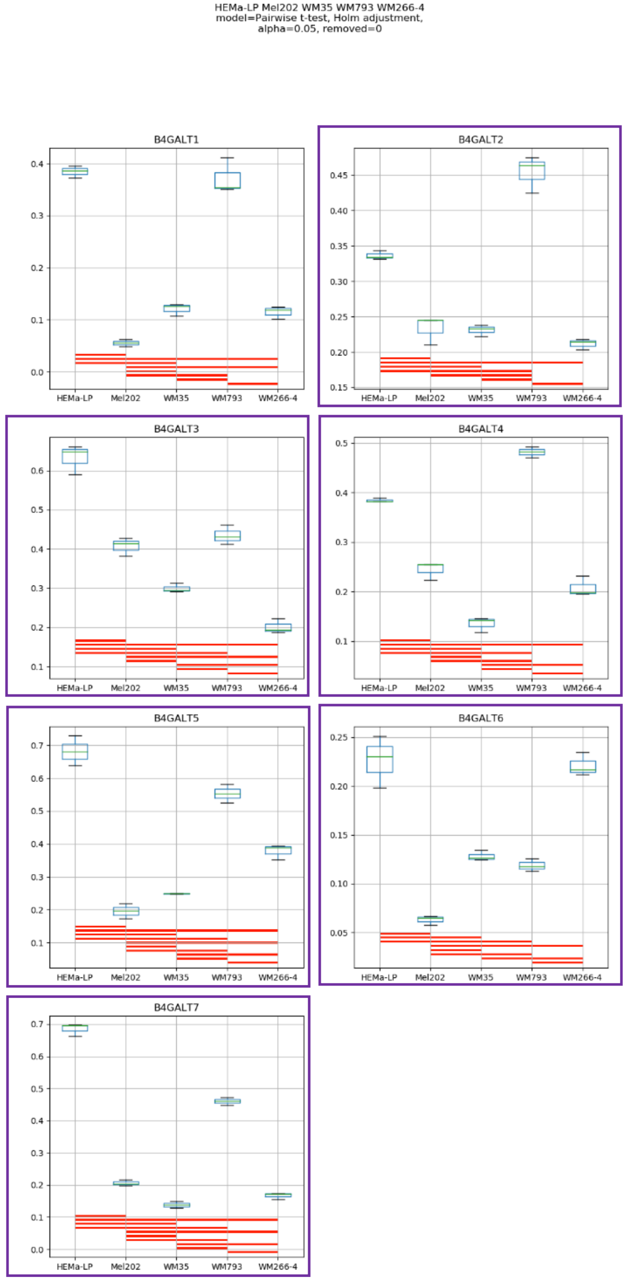
HEMa-LP Mel202 WM35 WM793 WM266-4
model=Pairwise t-test, Holm adjustment,
alpha=0.05, removed=0
B4GALT1
B4GALT2
B4GALT3
B4GALT4
B4GALT5
B4GALT6
B4GALT7
HEMa-LP
Mel202
WM35
WM793
WM266-4

**Supplementary Figure S12:** Medians of relative quantity (RQ) for target genes *B4GALT1–B4GALT7* normalized to the reference gene or pair of reference genes with the best stability values resulting from GenExpA analysis based on set 2 of four candidate reference genes (*RPS23*, *SNRPA*, *HPRT1*, *GUSB*) and remove repetition level 1. The statistical analyses used the pairwise t-test with Holm adjustment. Red line represents statistical significance, $P < 0.05$. Frames denote box-plots representing the medians of the RQ values for a given target gene without a repeatable pattern in the same cell lines under different models (the 'Remove repetitions' box set to 0 – violet frame,  or to 1 – red frame). Detailed results are listed in Supplementary Table 11.

Model No. 1

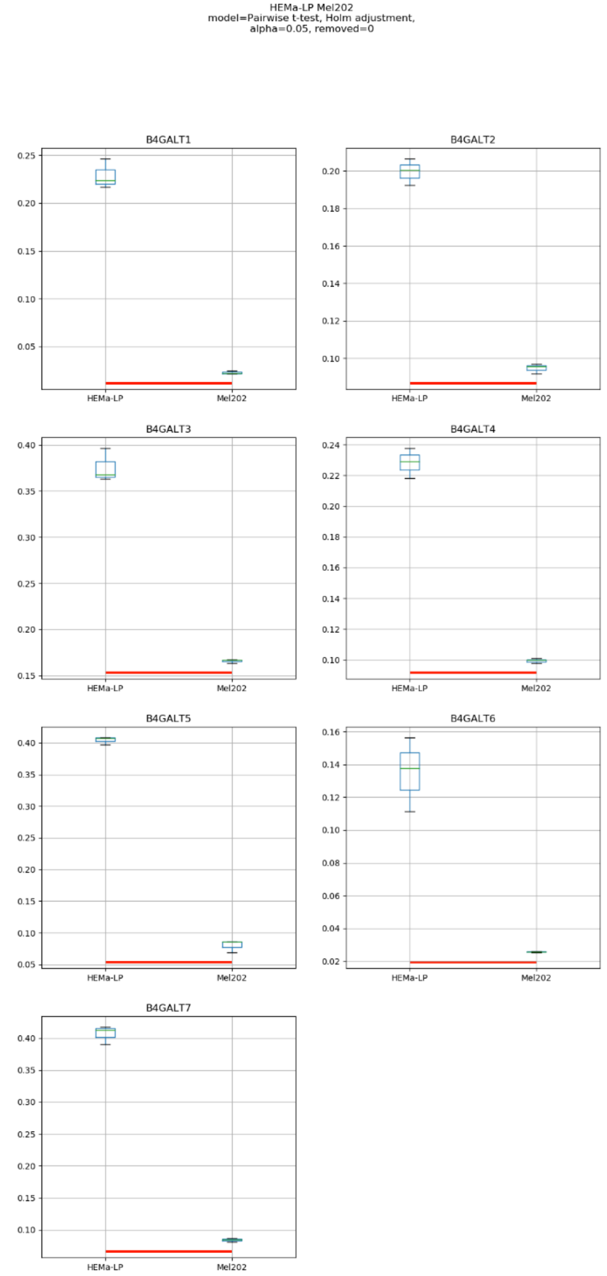

HEMa-LP Mel202
model=Pairwise t-test, Holm adjustment,
alpha=0.05, removed=0
B4GALT1
B4GALT2
B4GALT3
B4GALT4
B4GALT5
B4GALT6
B4GALT7
HEMa-LP
Mel202

Model No. 2

HEMa-LP WM35
model=Pairwise t-test, Holm adjustment,
alpha=0.05, removed=0

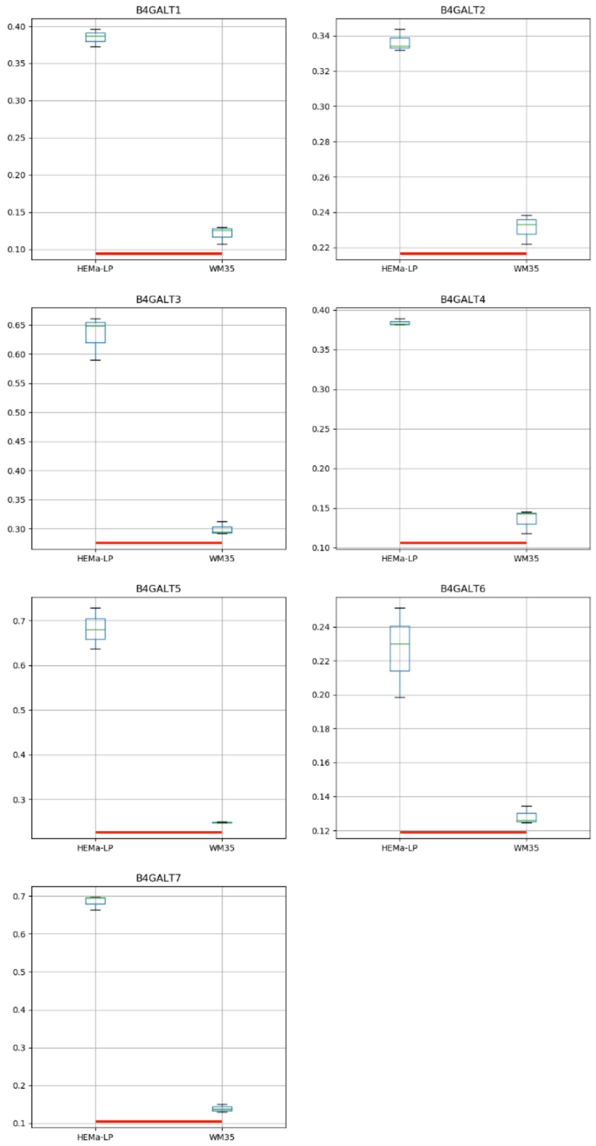

### Model No. 3

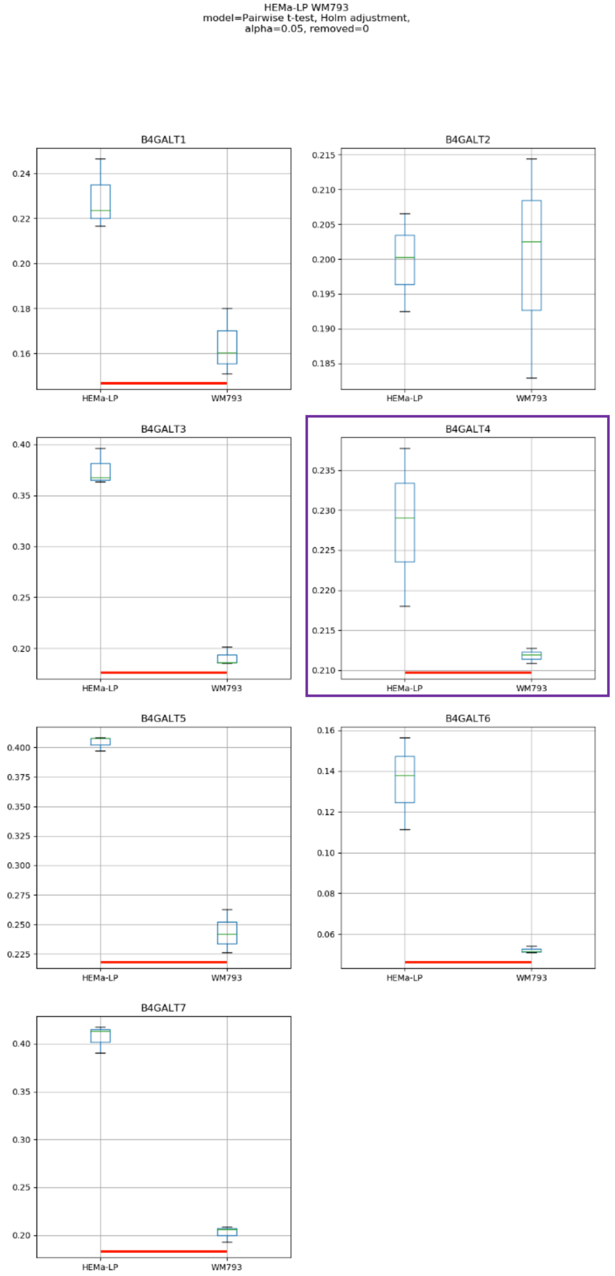
HEMa-LP WM793
model=Pairwise t-test, Holm adjustment,
alpha=0.05, removed=0
B4GALT1
B4GALT2
B4GALT3
B4GALT4
B4GALT5
B4GALT6
B4GALT7
HEMa-LP
WM793

Model No. 4

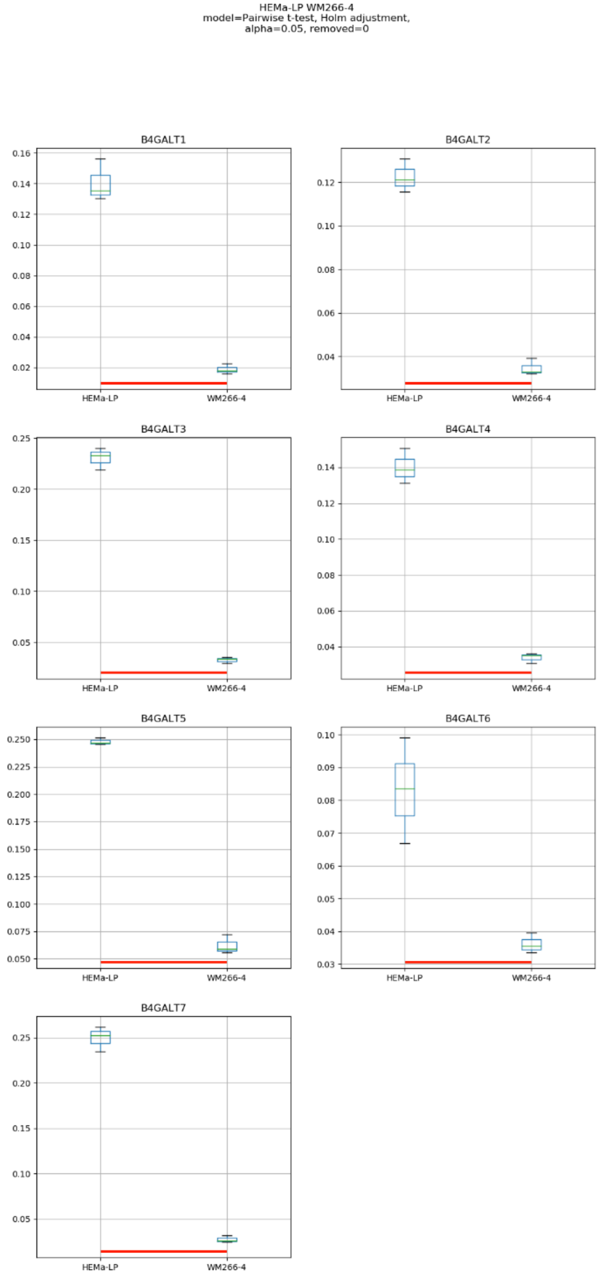
HEMa-LP WM266-4
model=Pairwise t-test, Holm adjustment,
alpha=0.05, removed=0
B4GALT1
B4GALT2
B4GALT3
B4GALT4
B4GALT5
B4GALT6
B4GALT7
HEMa-LP
WM266-4

### Model No. 5

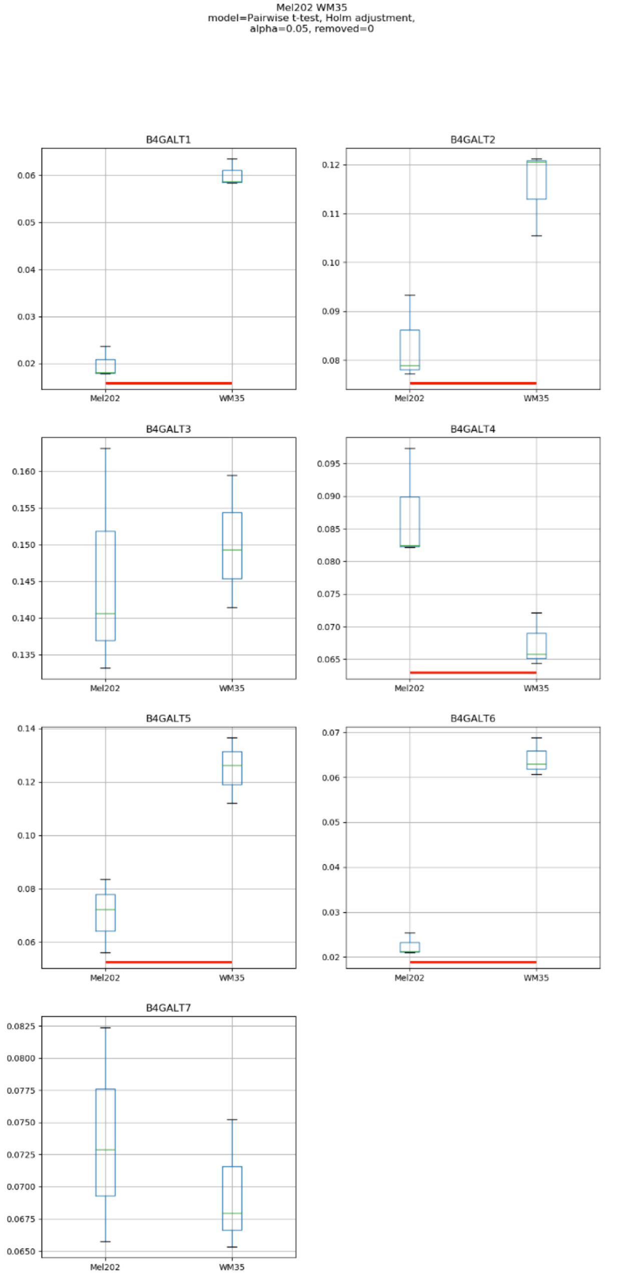
Mel202 WM35
model=Pairwise t-test, Holm adjustment,
alpha=0.05, removed=0
B4GALT1
B4GALT2
B4GALT3
B4GALT4
B4GALT5
B4GALT6
B4GALT7
Mel202
WM35

### Model No. 6

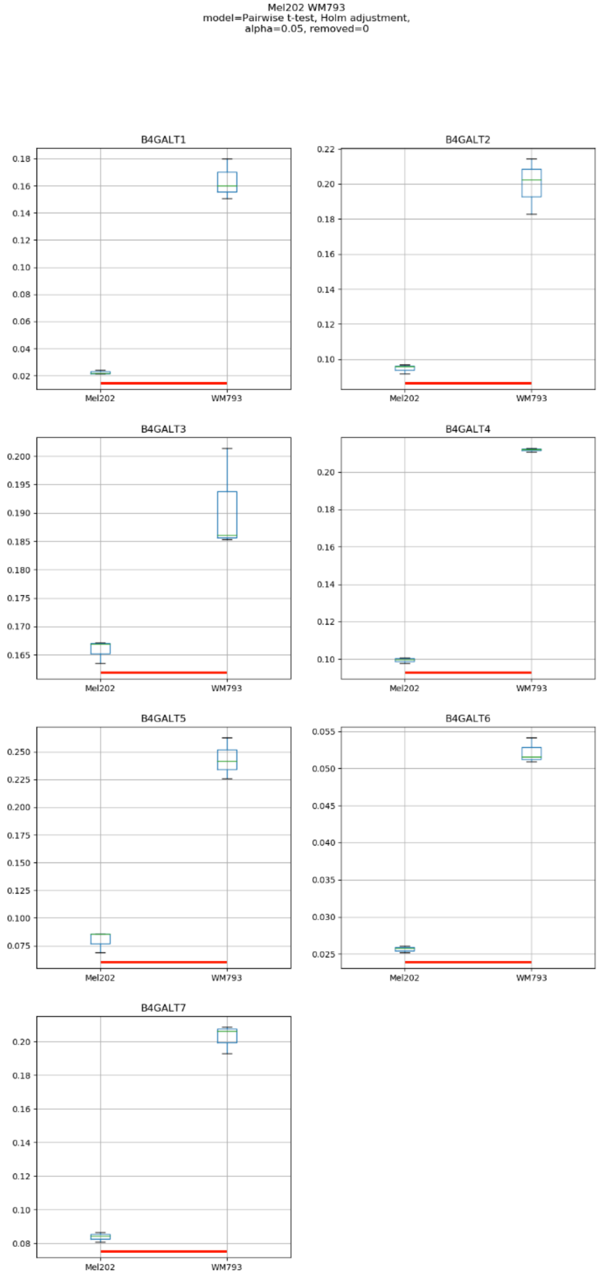
Mel202 WM793
model=Pairwise t-test, Holm adjustment,
alpha=0.05, removed=0
B4GALT1
B4GALT2
B4GALT3
B4GALT4
B4GALT5
B4GALT6
B4GALT7
Mel202
WM793

### Model No. 7

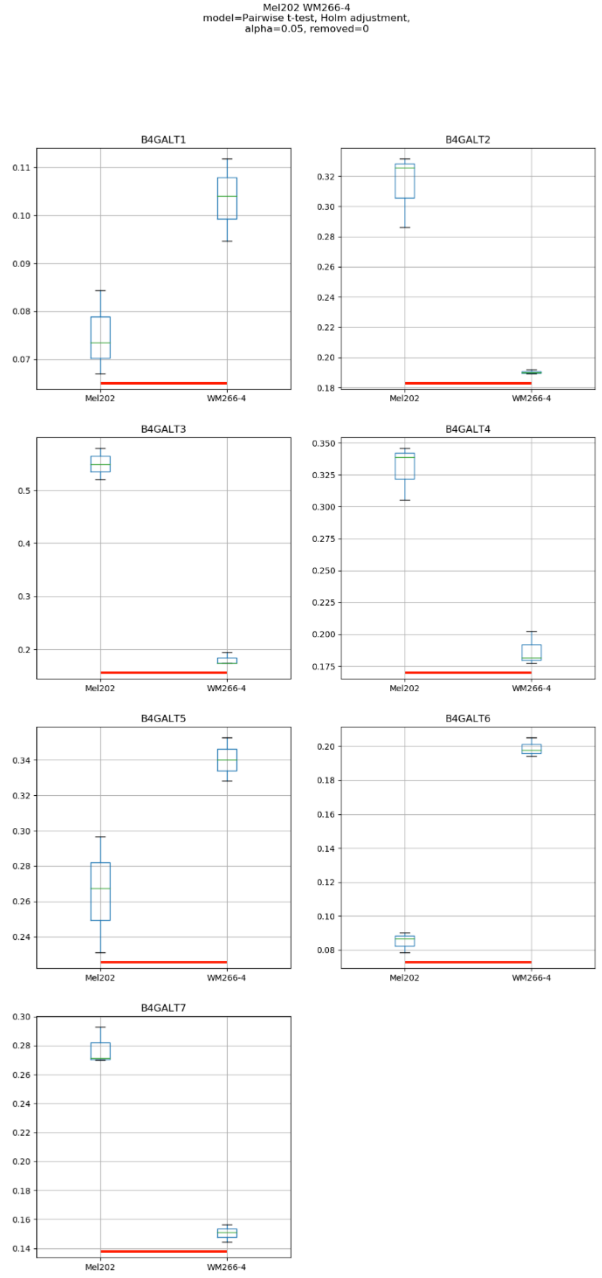

Mel202 WM266-4
model=Pairwise t-test, Holm adjustment,
alpha=0.05, removed=0
B4GALT1
B4GALT2
B4GALT3
B4GALT4
B4GALT5
B4GALT6
B4GALT7
Mel202
WM266-4

### Model No. 8

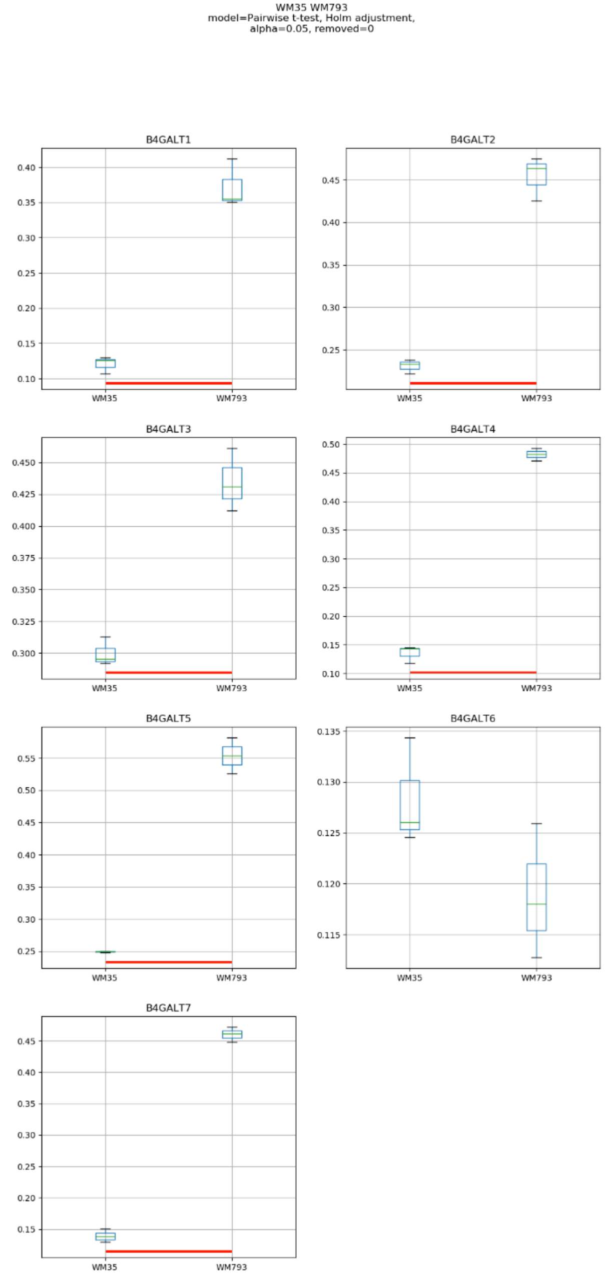
WM35 WM793
model=Pairwise t-test, Holm adjustment,
alpha=0.05, removed=0
B4GALT1
B4GALT2
B4GALT3
B4GALT4
B4GALT5
B4GALT6
B4GALT7
WM35
WM793

### Model No. 9

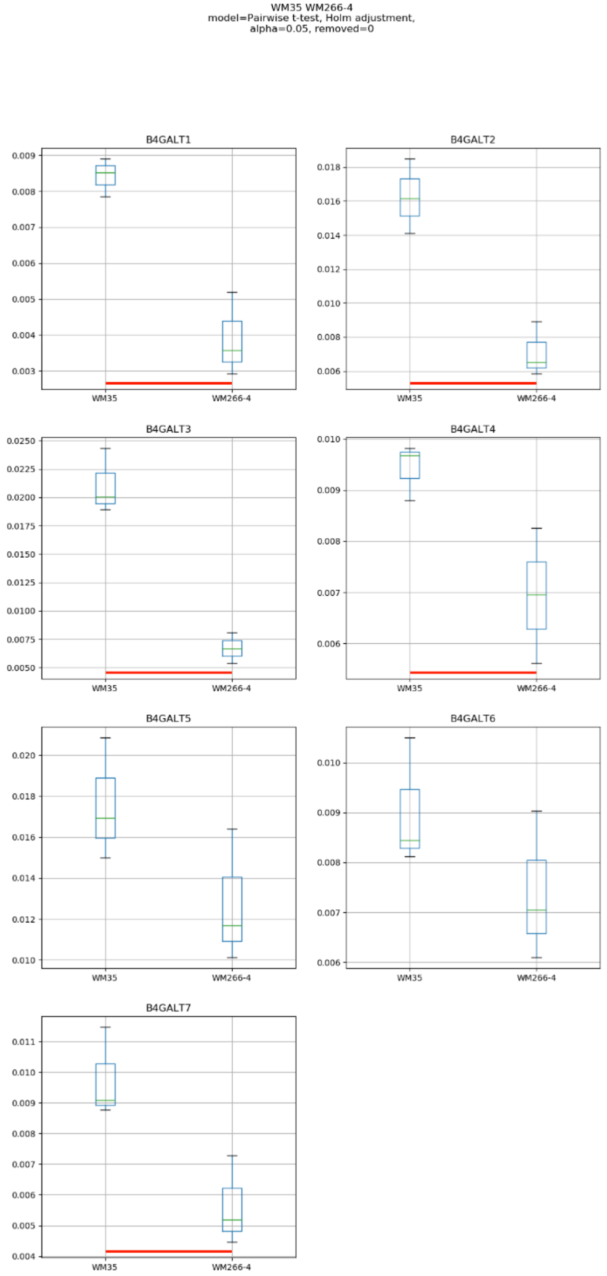
WM35 WM266-4
model=Pairwise t-test, Holm adjustment,
alpha=0.05, removed=0
B4GALT1
B4GALT2
B4GALT3
B4GALT4
B4GALT5
B4GALT6
B4GALT7
WM35
WM266-4

### Model No. 10

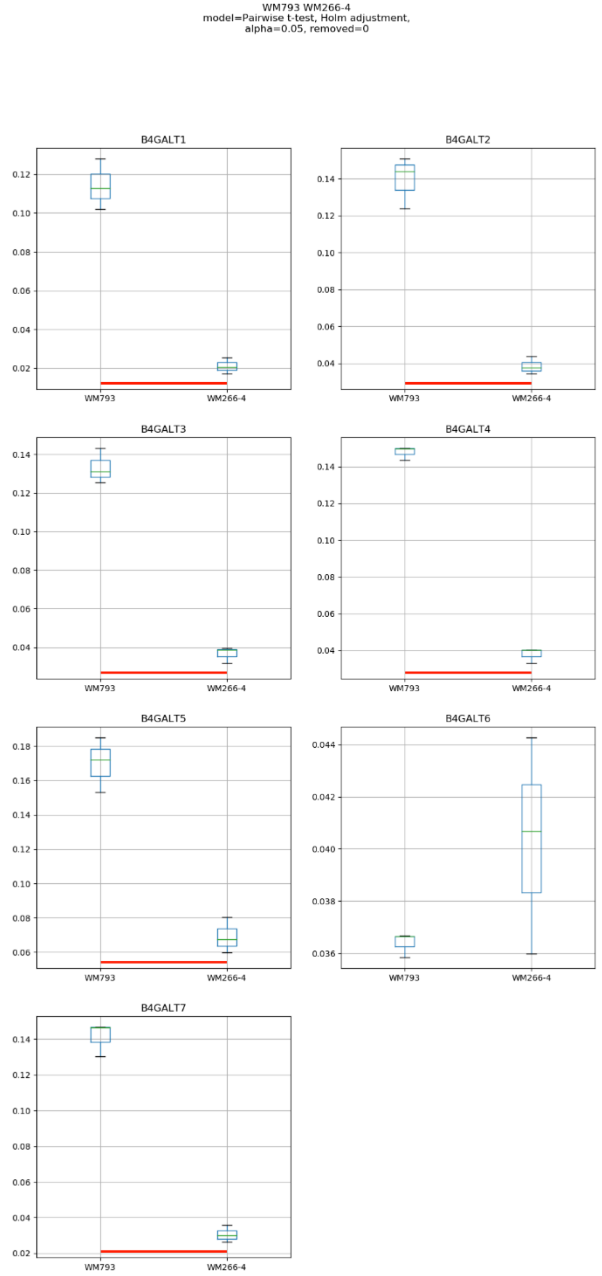
WM793 WM266-4
model=Pairwise t-test, Holm adjustment,
alpha=0.05, removed=0
B4GALT1
B4GALT2
B4GALT3
B4GALT4
B4GALT5
B4GALT6
B4GALT7
WM793
WM266-4

Model No. 11

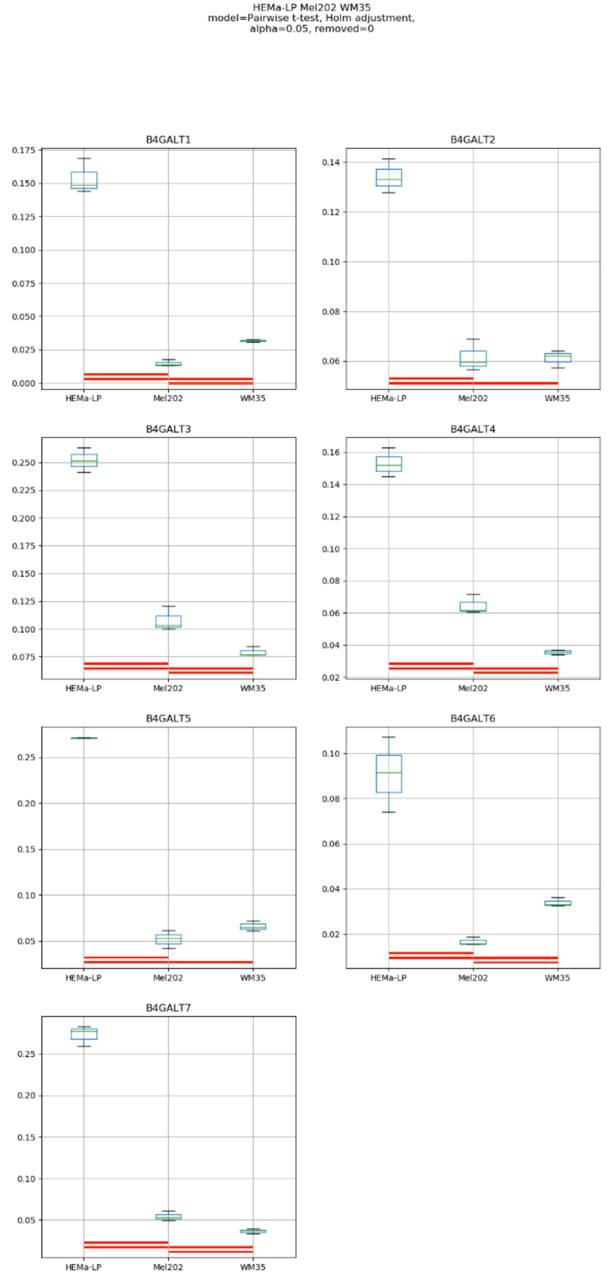
HEMa-LP Mel202 WM35
model=Pairwise t-test, Holm adjustment,
alpha=0.05, removed=0
B4GALT1
B4GALT2
B4GALT3
B4GALT4
B4GALT5
B4GALT6
B4GALT7
HEMa-LP
Mel202
WM35

### Model No. 12

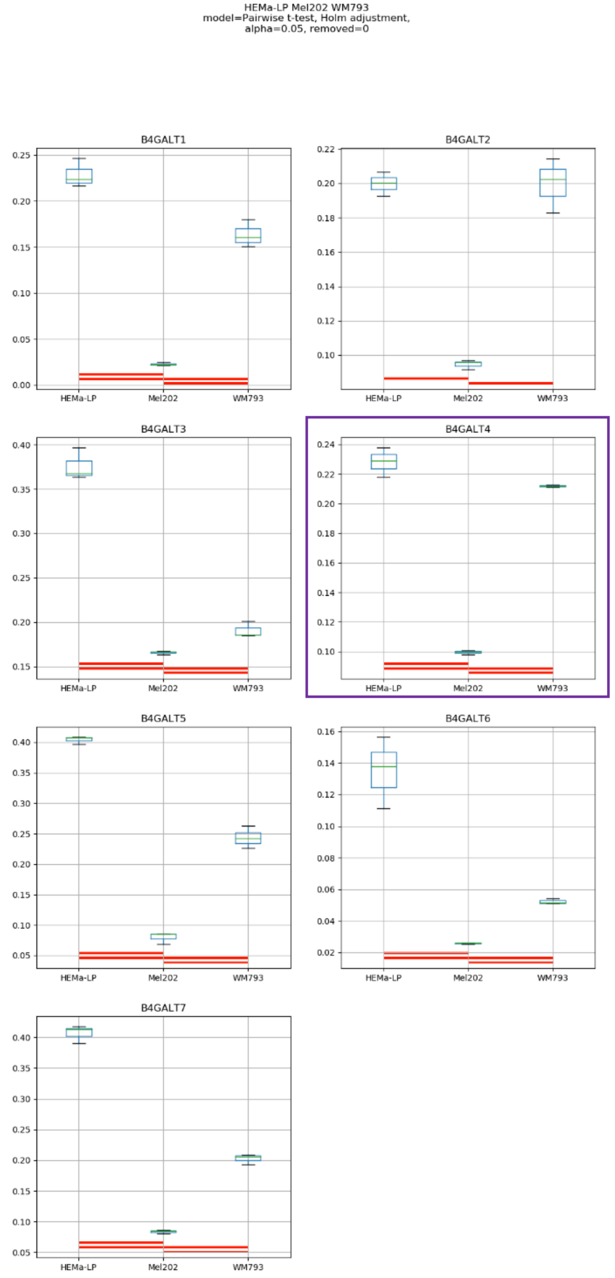

HEMa-LP Mel202 WM793
model=Pairwise t-test, Holm adjustment,
alpha=0.05, removed=0
B4GALT1
B4GALT2
B4GALT3
B4GALT4
B4GALT5
B4GALT6
B4GALT7
HEMa-LP
Mel202
WM793

Model No. 13

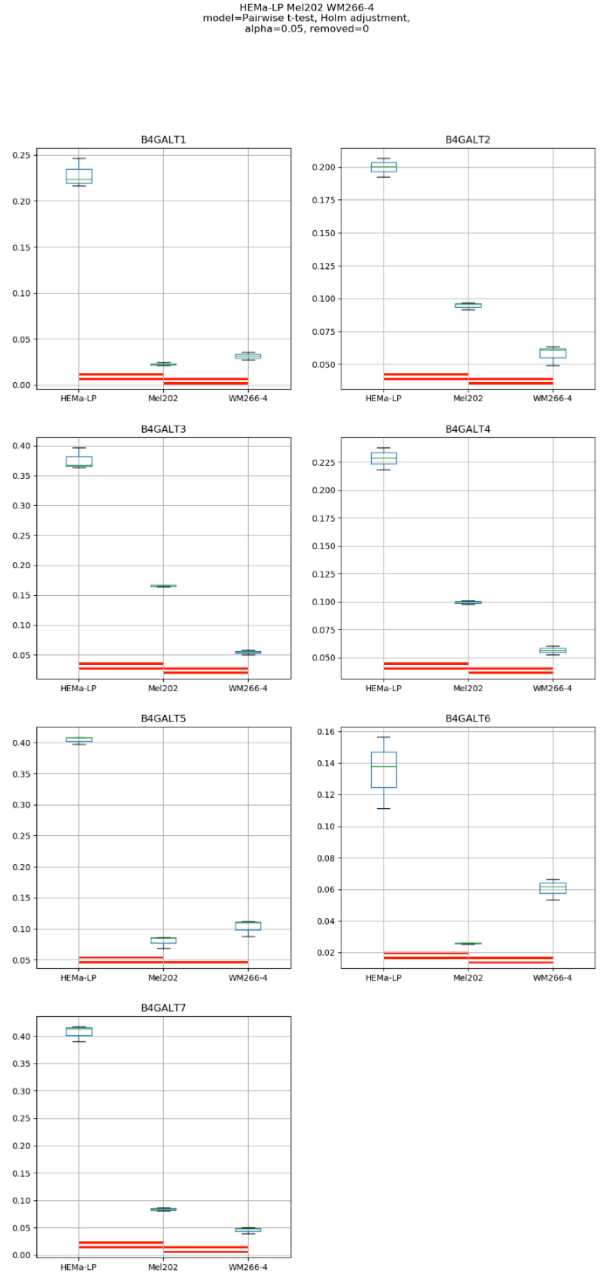
HEMa-LP Mel202 WM266-4
model=Pairwise t-test, Holm adjustment,
alpha=0.05, removed=0
B4GALT1
B4GALT2
B4GALT3
B4GALT4
B4GALT5
B4GALT6
B4GALT7
HEMa-LP
Mel202
WM266-4

### Model No. 14

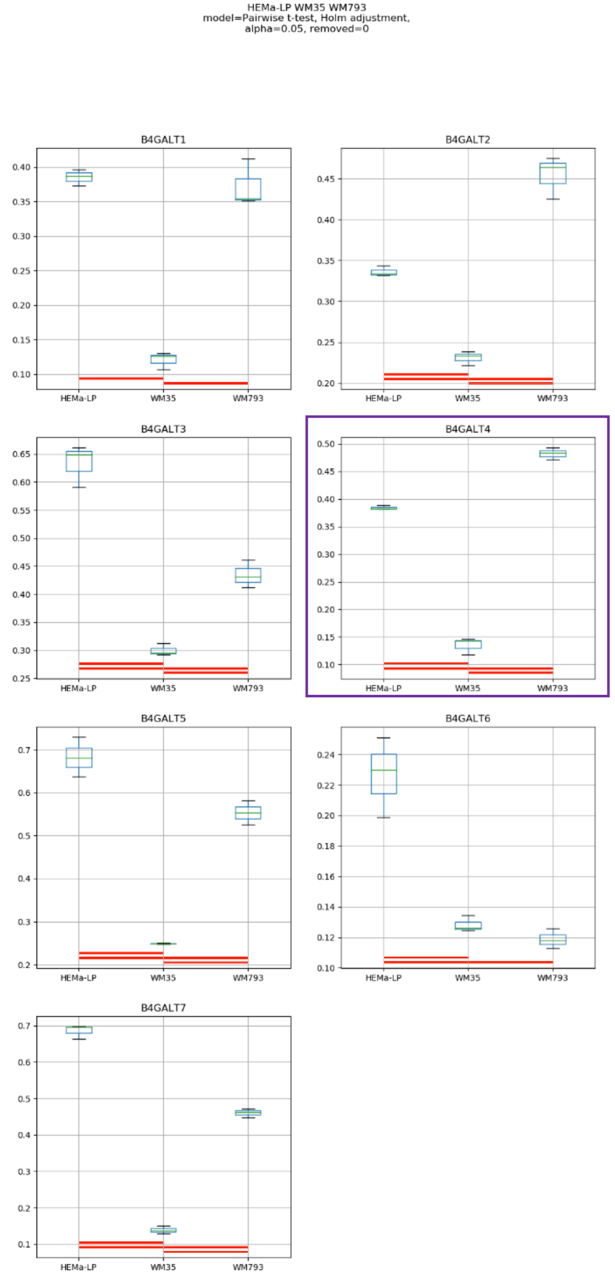
HEMa-LP WM35 WM793
model=Pairwise t-test, Holm adjustment,
alpha=0.05, removed=0
B4GALT1
B4GALT2
B4GALT3
B4GALT4
B4GALT5
B4GALT6
B4GALT7
HEMa-LP
WM35
WM793

Model No. 15

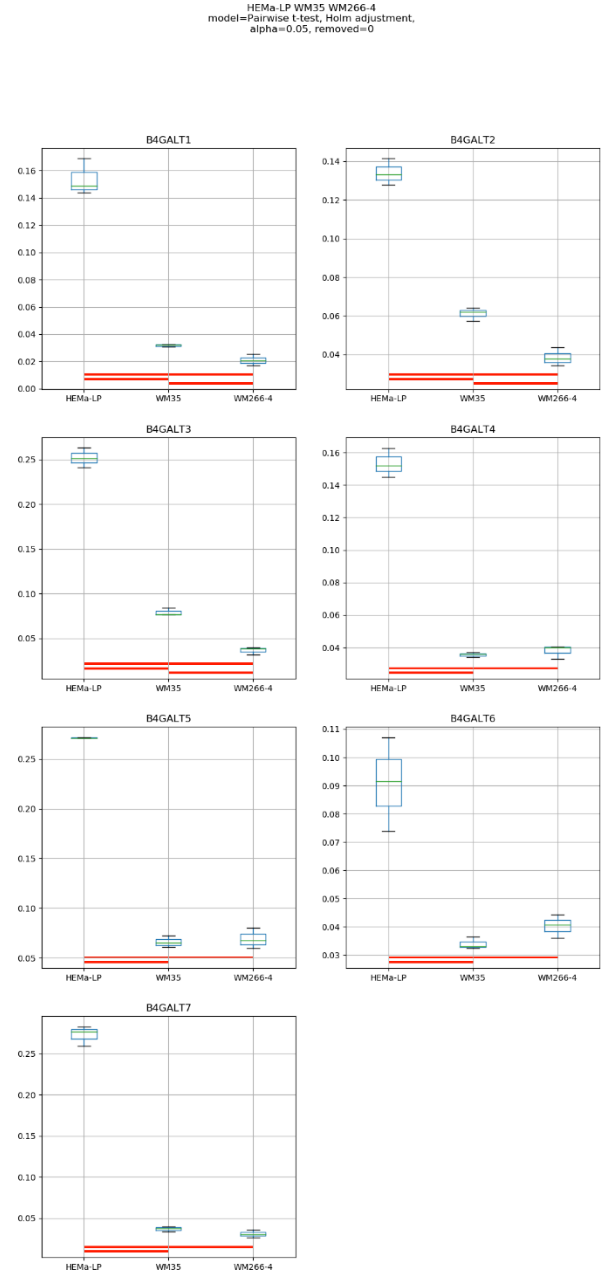
HEMa-LP WM35 WM266-4
model=Pairwise t-test, Holm adjustment,
alpha=0.05, removed=0
B4GALT1
B4GALT2
B4GALT3
B4GALT4
B4GALT5
B4GALT6
B4GALT7
HEMa-LP
WM35
WM266-4

Model No. 16

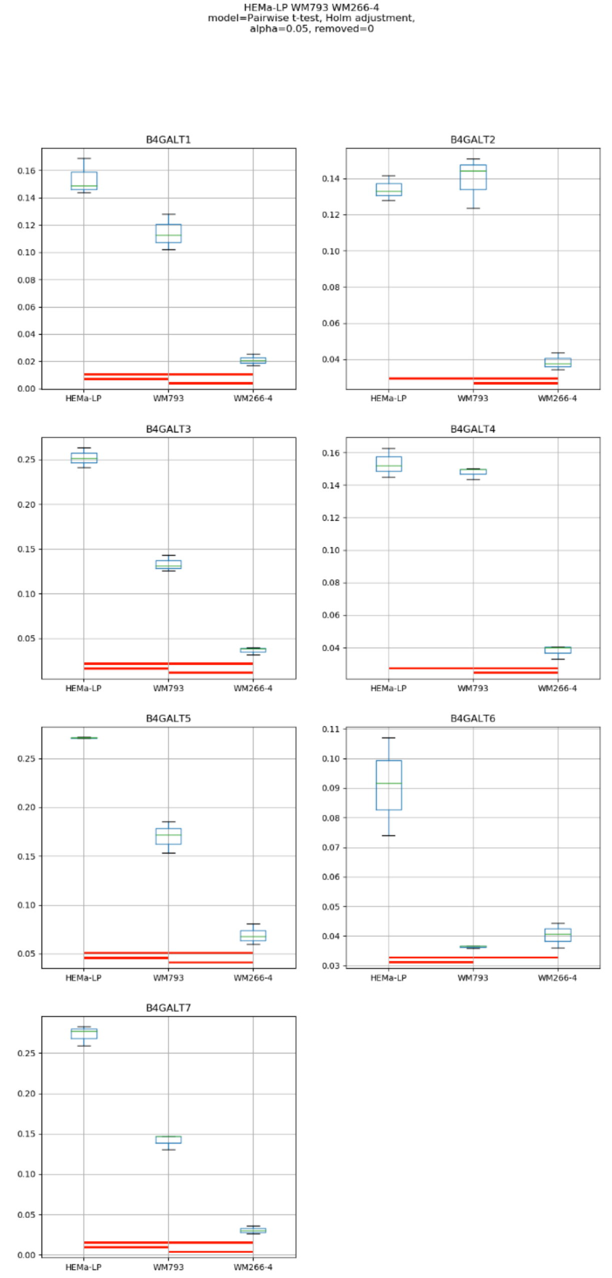
HEMa-LP WM793 WM266-4
model=Pairwise t-test, Holm adjustment,
alpha=0.05, removed=0
B4GALT1
B4GALT2
B4GALT3
B4GALT4
B4GALT5
B4GALT6
B4GALT7
HEMa-LP
WM793
WM266-4

Model No. 17

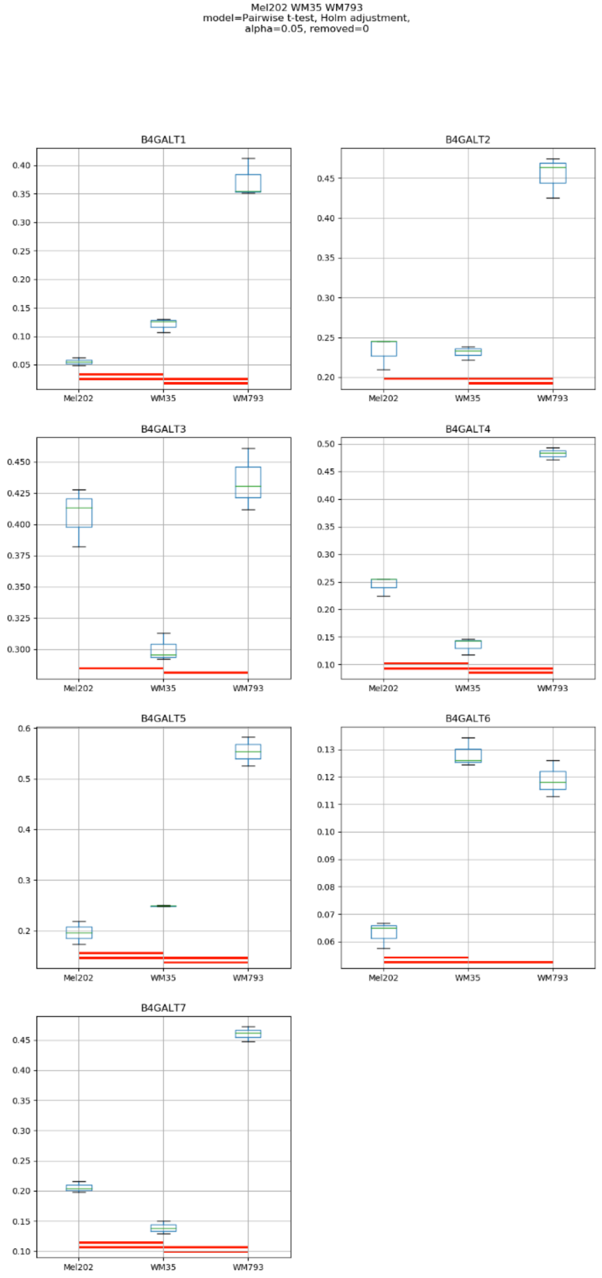
Mel202 WM35 WM793
model=Pairwise t-test, Holm adjustment,
alpha=0.05, removed=0
B4GALT1
B4GALT2
B4GALT3
B4GALT4
B4GALT5
B4GALT6
B4GALT7
Mel202
WM35
WM793

### Model No. 18

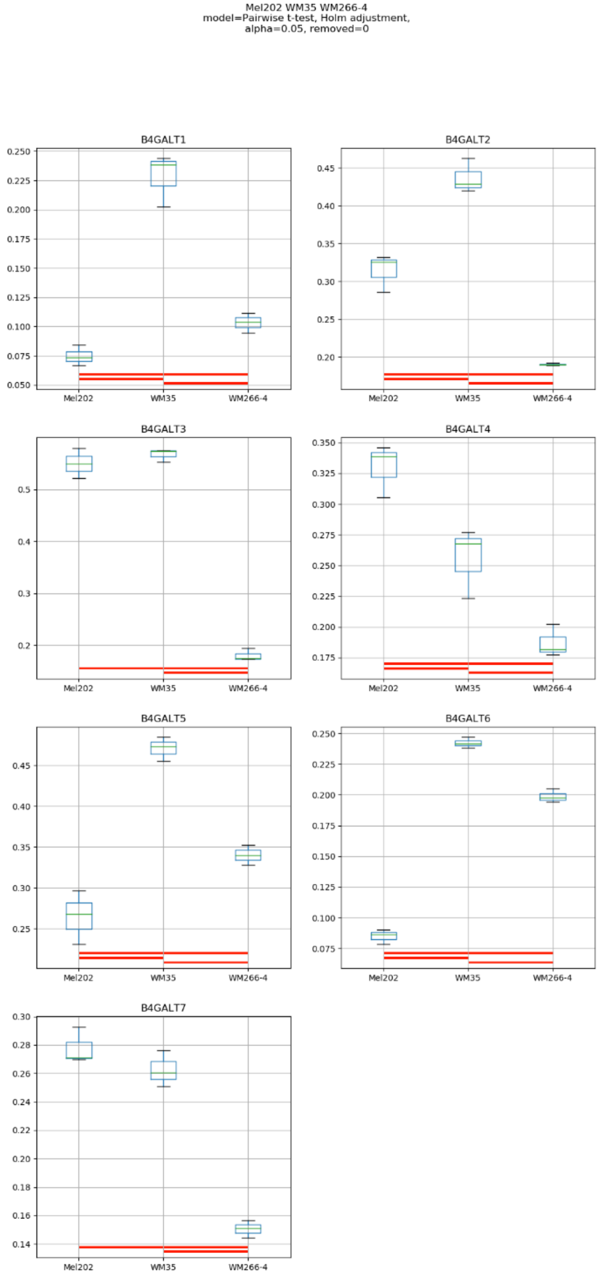
Mel202 WM35 WM266-4
model=Pairwise t-test, Holm adjustment,
alpha=0.05, removed=0
B4GALT1
B4GALT2
B4GALT3
B4GALT4
B4GALT5
B4GALT6
B4GALT7
Mel202
WM35
WM266-4

### Model No. 19

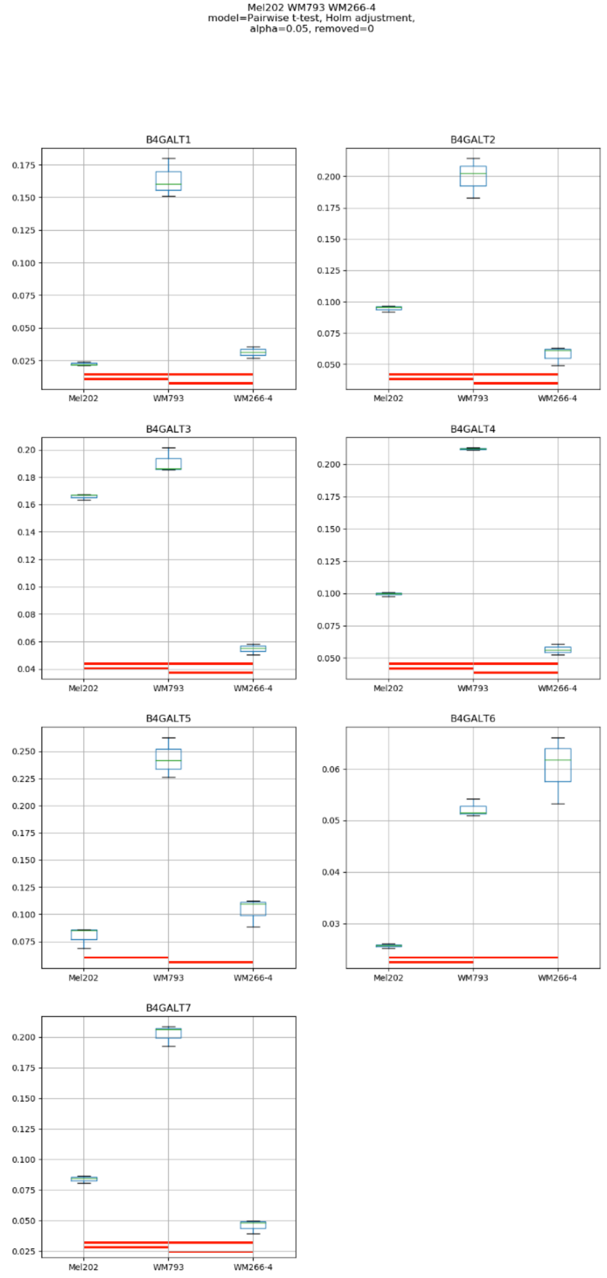

Mel202 WM793 WM266-4
model=Pairwise t-test, Holm adjustment,
alpha=0.05, removed=0
B4GALT1
B4GALT2
B4GALT3
B4GALT4
B4GALT5
B4GALT6
B4GALT7
Mel202
WM793
WM266-4

### Model No. 20

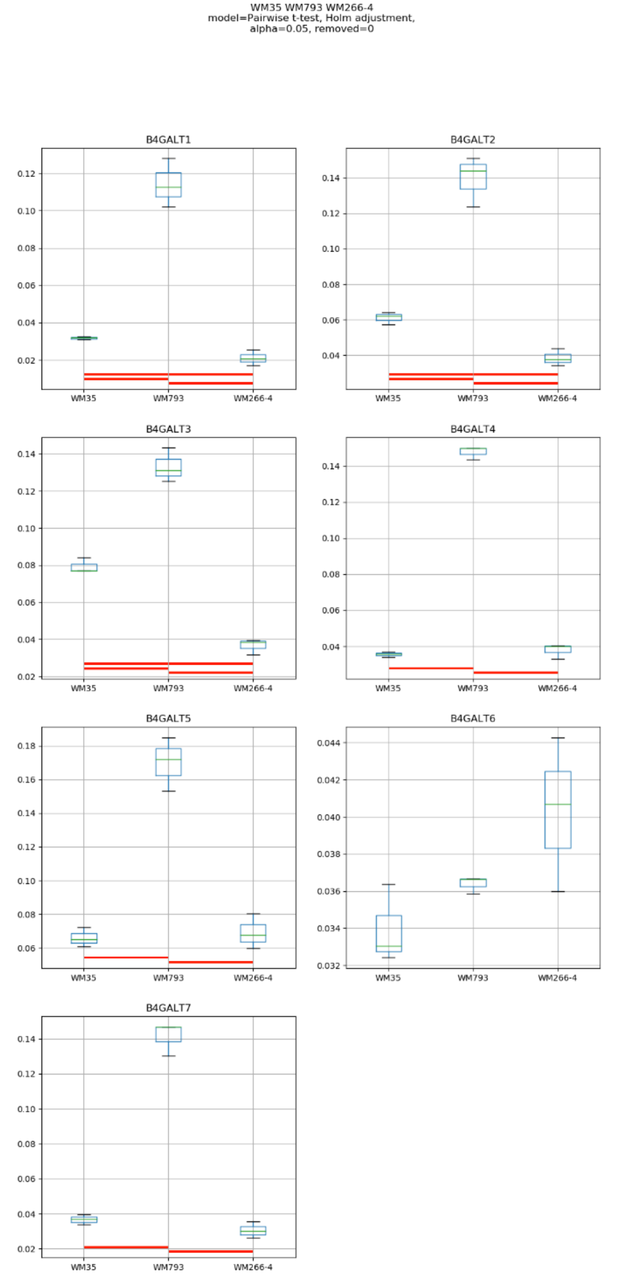
WM35 WM793 WM266-4
model=Pairwise t-test, Holm adjustment,
alpha=0.05, removed=0
B4GALT1
B4GALT2
B4GALT3
B4GALT4
B4GALT5
B4GALT6
B4GALT7
WM35
WM793
WM266-4

### Model No. 21

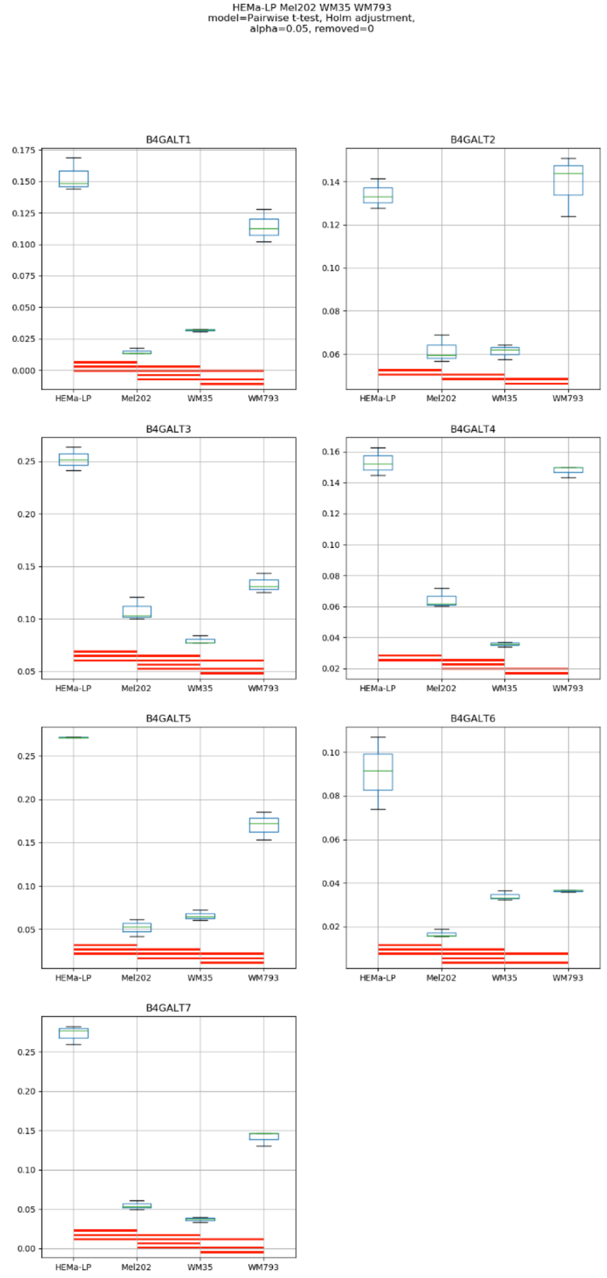
HEMa-LP Mel202 WM35 WM793
model=Pairwise t-test, Holm adjustment,
alpha=0.05, removed=0
B4GALT1
B4GALT2
B4GALT3
B4GALT4
B4GALT5
B4GALT6
B4GALT7
HEMa-LP
Mel202
WM35
WM793

### Model No. 22

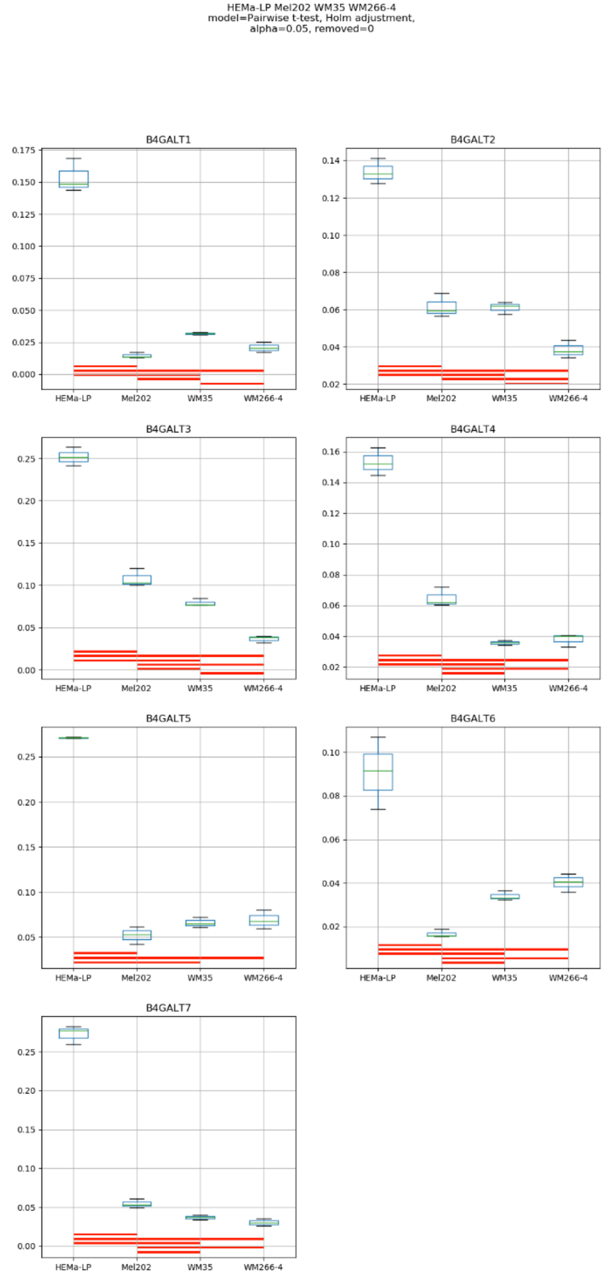
HEMa-LP Mel202 WM35 WM266-4
model=Pairwise t-test, Holm adjustment,
alpha=0.05, removed=0
B4GALT1
B4GALT2
B4GALT3
B4GALT4
B4GALT5
B4GALT6
B4GALT7
HEMa-LP
Mel202
WM35
WM266-4

### Model No. 23

HEMa-LP Mel202 WM793 WM266-4
model=Pairwise t-test, Holm adjustment,
alpha=0.05, removed=0

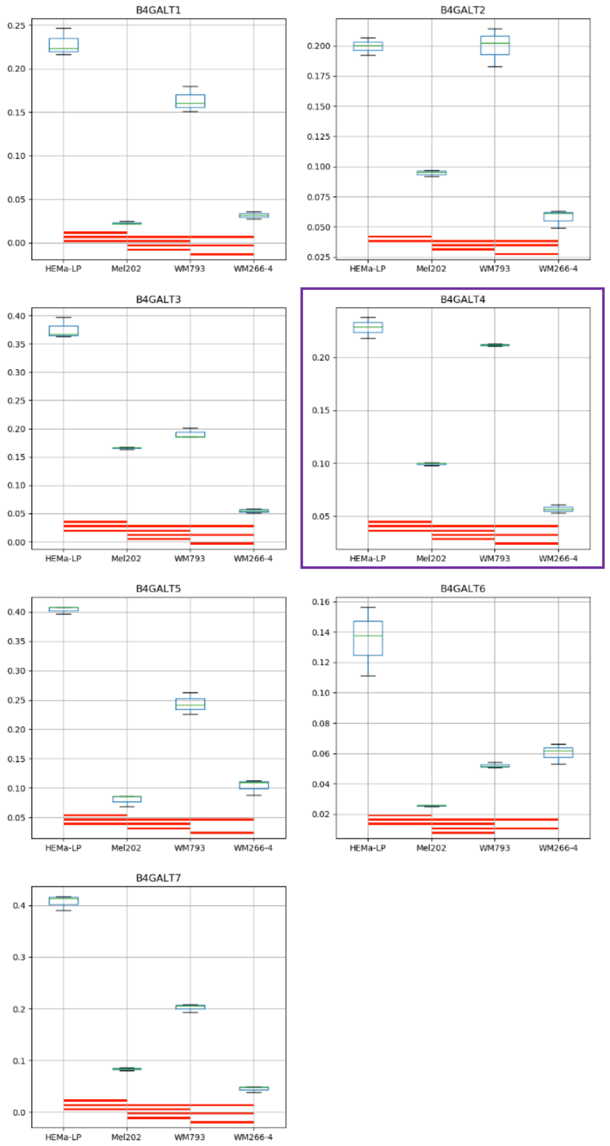

Model No. 24

HEMa-LP WM35 WM793 WM266-4
model=Pairwise t-test, Holm adjustment,
alpha=0.05, removed=0

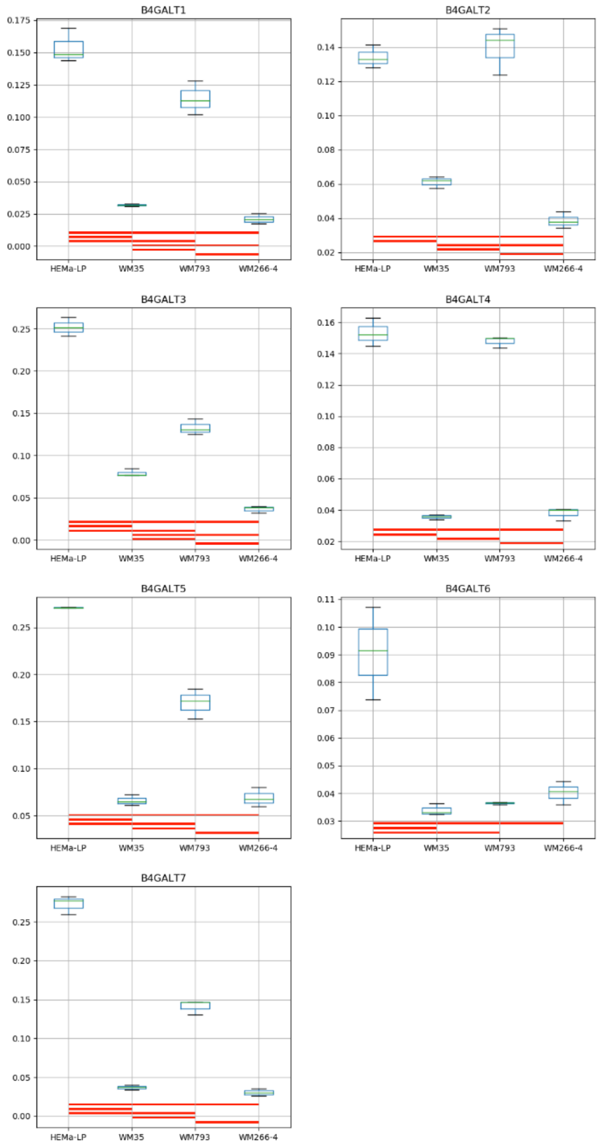

Model No. 25

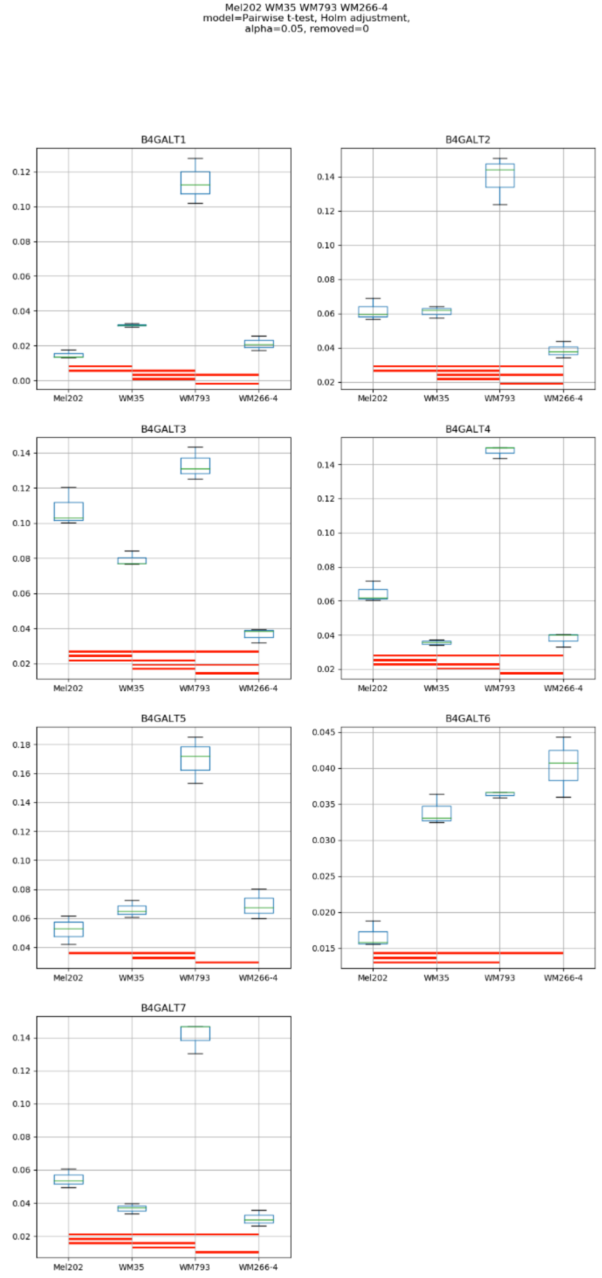
Mel202 WM35 WM793 WM266-4
model=Pairwise t-test, Holm adjustment,
alpha=0.05, removed=0
B4GALT1
B4GALT2
B4GALT3
B4GALT4
B4GALT5
B4GALT6
B4GALT7
Mel202
WM35
WM793
WM266-4

Model No. 26

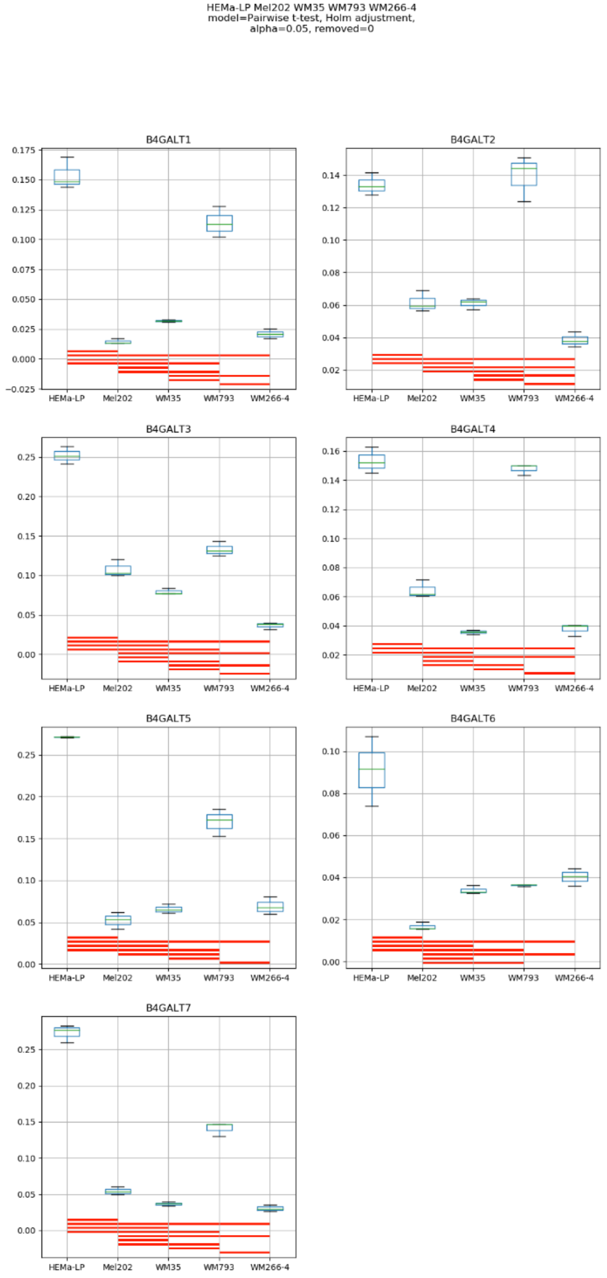

HEMa-LP Mel202 WM35 WM793 WM266-4
model=Pairwise t-test, Holm adjustment,
alpha=0.05, removed=0
B4GALT1
B4GALT2
B4GALT3
B4GALT4
B4GALT5
B4GALT6
B4GALT7
HEMa-LP
Mel202
WM35
WM793
WM266-4

**Supplementary Figure S13:** Medians of relative quantity (RQ) for target genes *B4GALT1–B4GALT7* normalized to the reference gene or pair of reference genes with the best stability values resulting from GenExpA analysis based on set 3 of four candidate reference genes (*SNRPA*, *HPRT1*, *GUSB*, *PGK1*) and remove repetition level 0. The statistical analyses used the pairwise t-test with Holm adjustment. Red line represents statistical significance, $P < 0.05$. Frames denote box-plots representing the medians of the RQ values for a given target gene without a repeatable pattern in the same cell lines under different models (the ‘Remove repetitions’ box set to 0). Detailed results are listed in Supplementary Table 12.

### Model No. 1

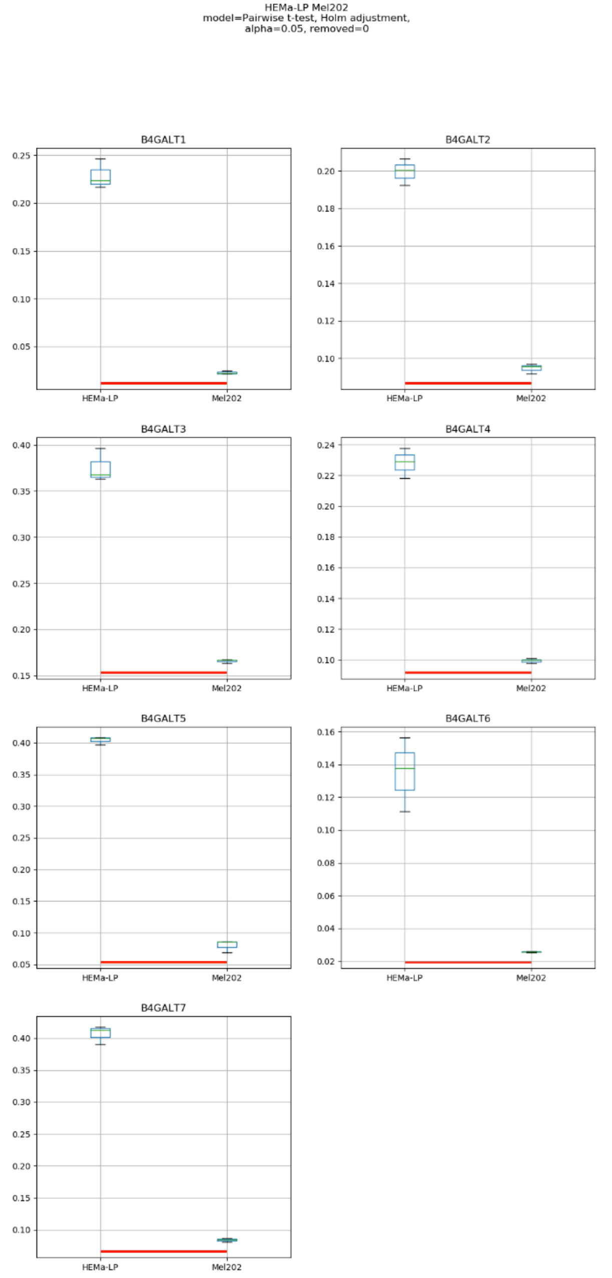
HEMa-LP Mel202
model=Pairwise t-test, Holm adjustment,
alpha=0.05, removed=0
B4GALT1
B4GALT2
B4GALT3
B4GALT4
B4GALT5
B4GALT6
B4GALT7
HEMa-LP
Mel202

### Model No. 2

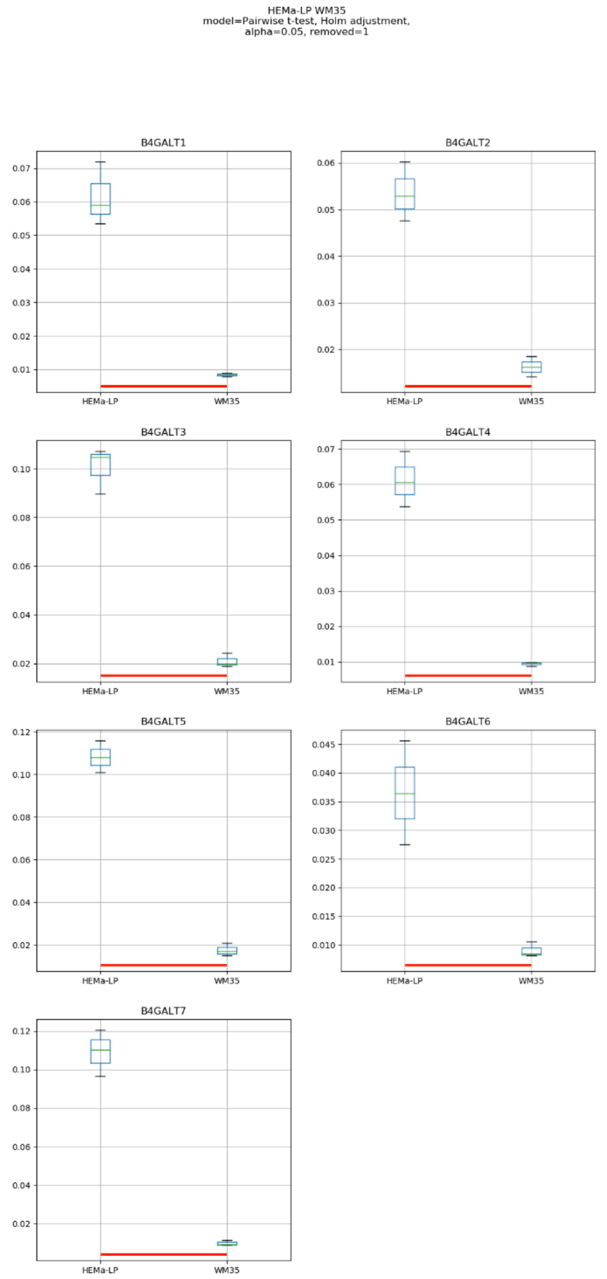
HEMa-LP WM35
model=Pairwise t-test, Holm adjustment,
alpha=0.05, removed=1
B4GALT1
B4GALT2
B4GALT3
B4GALT4
B4GALT5
B4GALT6
B4GALT7
HEMa-LP
WM35

### Model No. 3

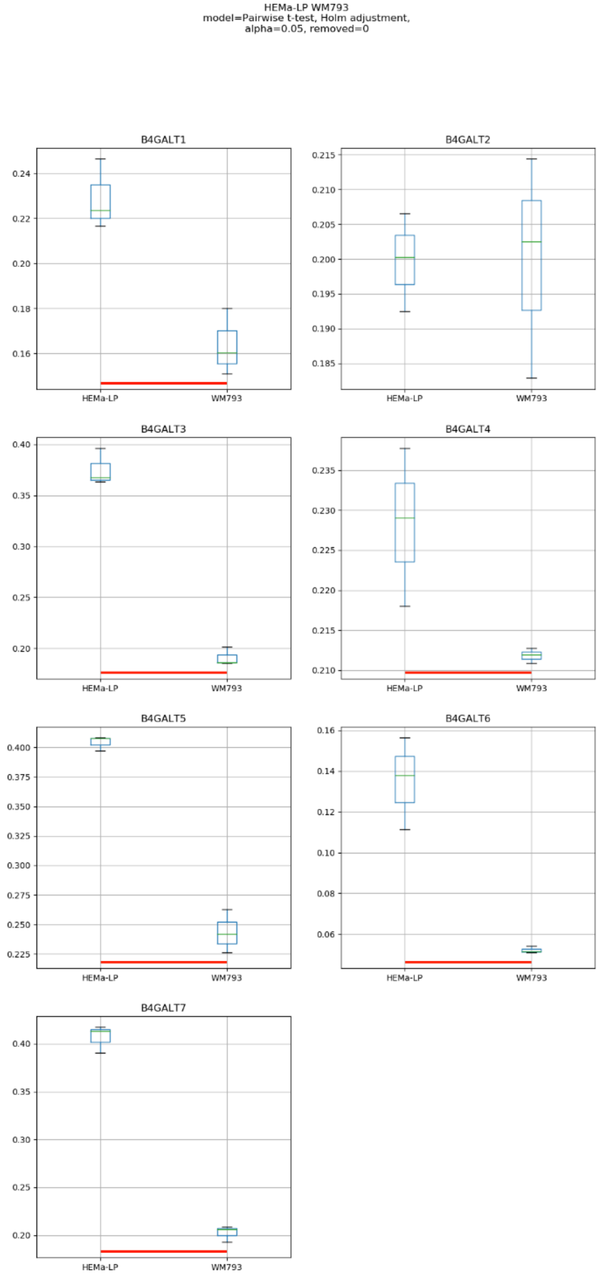
HEMa-LP WM793
model=Pairwise t-test, Holm adjustment,
alpha=0.05, removed=0
B4GALT1
B4GALT2
B4GALT3
B4GALT4
B4GALT5
B4GALT6
B4GALT7
HEMa-LP
WM793

### Model No. 4

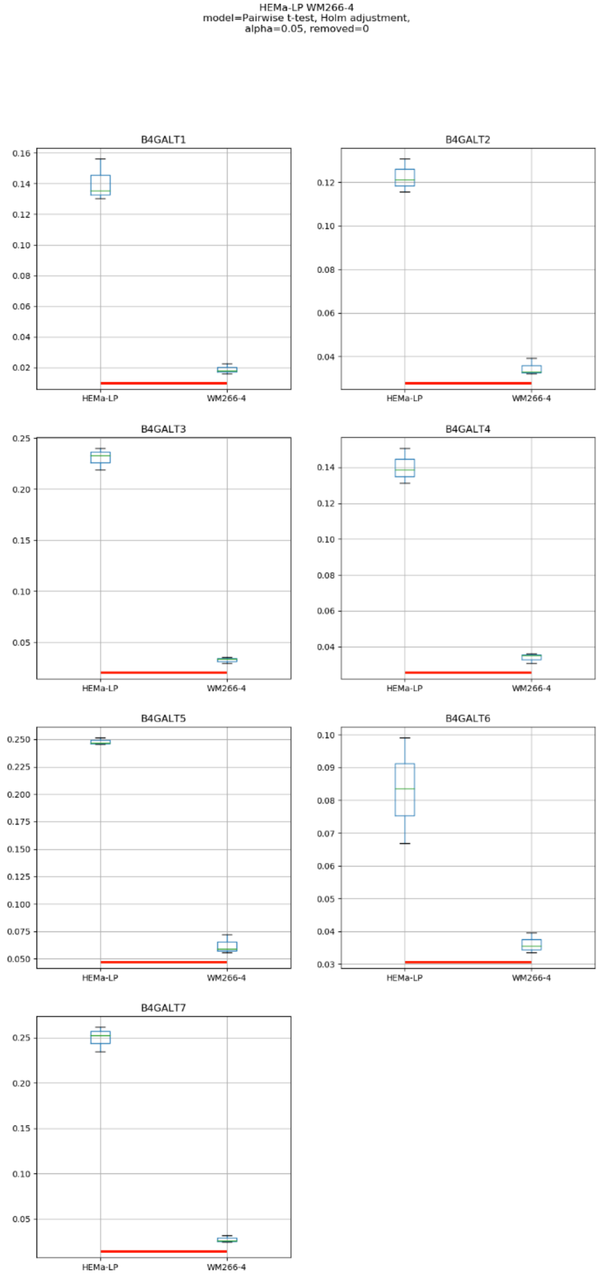
HEMa-LP WM266-4
model=Pairwise t-test, Holm adjustment,
alpha=0.05, removed=0
B4GALT1
B4GALT2
B4GALT3
B4GALT4
B4GALT5
B4GALT6
B4GALT7
HEMa-LP
WM266-4

### Model No. 5

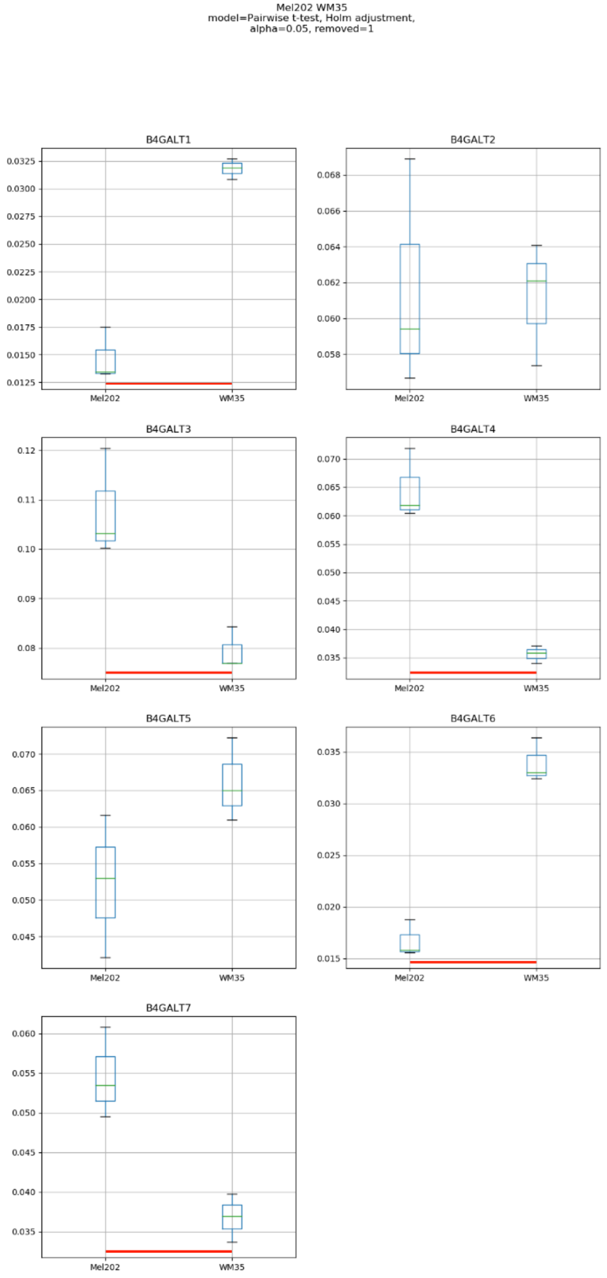
Mel202 WM35
model=Pairwise t-test, Holm adjustment,
alpha=0.05, removed=1
B4GALT1
B4GALT2
B4GALT3
B4GALT4
B4GALT5
B4GALT6
B4GALT7
Mel202
WM35

### Model No. 6

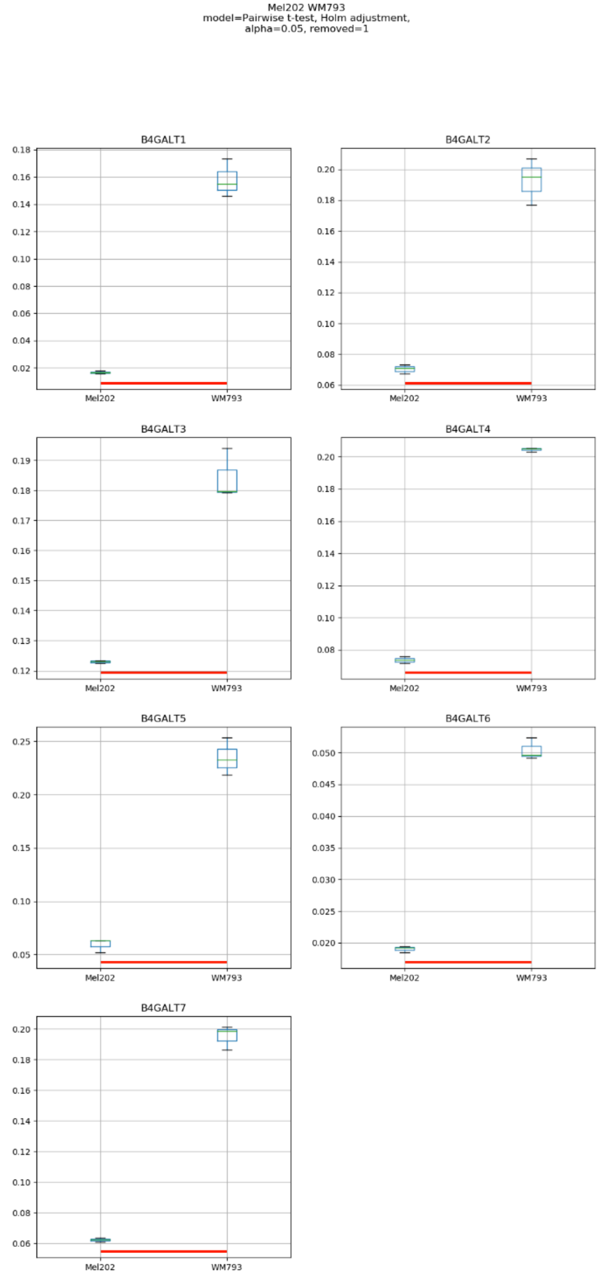

Mel202 WM793
model=Pairwise t-test, Holm adjustment,
alpha=0.05, removed=1
B4GALT1
B4GALT2
B4GALT3
B4GALT4
B4GALT5
B4GALT6
B4GALT7
Mel202
WM793

Model No. 7

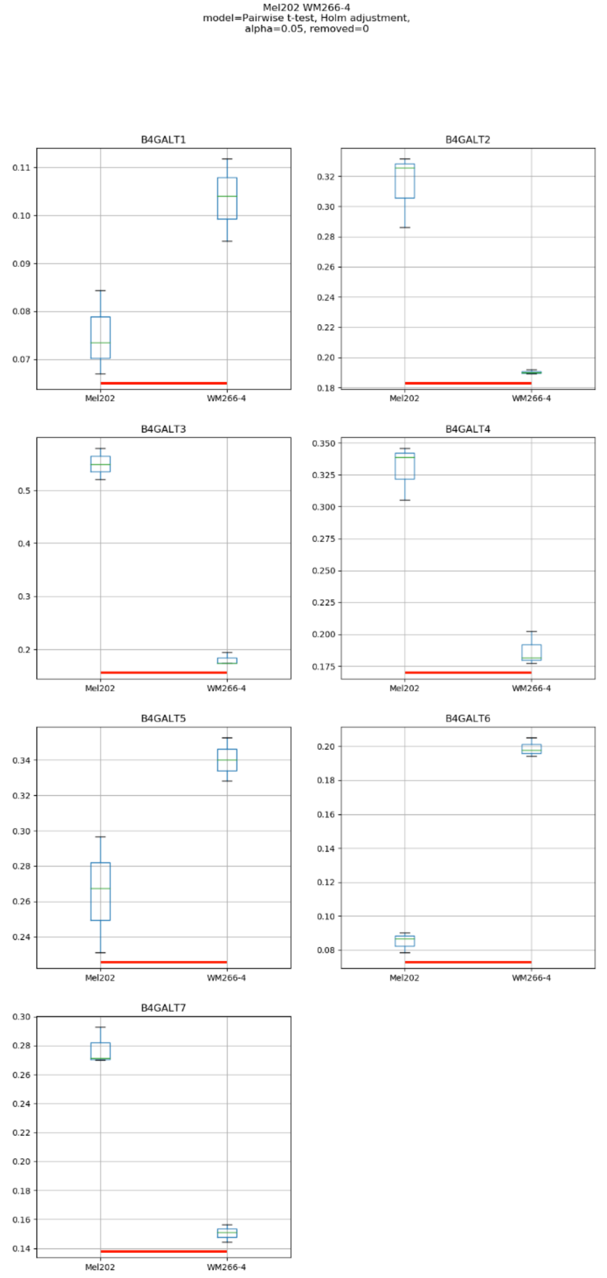
Mel202 WM266-4
model=Pairwise t-test, Holm adjustment,
alpha=0.05, removed=0
B4GALT1
B4GALT2
B4GALT3
B4GALT4
B4GALT5
B4GALT6
B4GALT7
Mel202
WM266-4

### Model No. 8

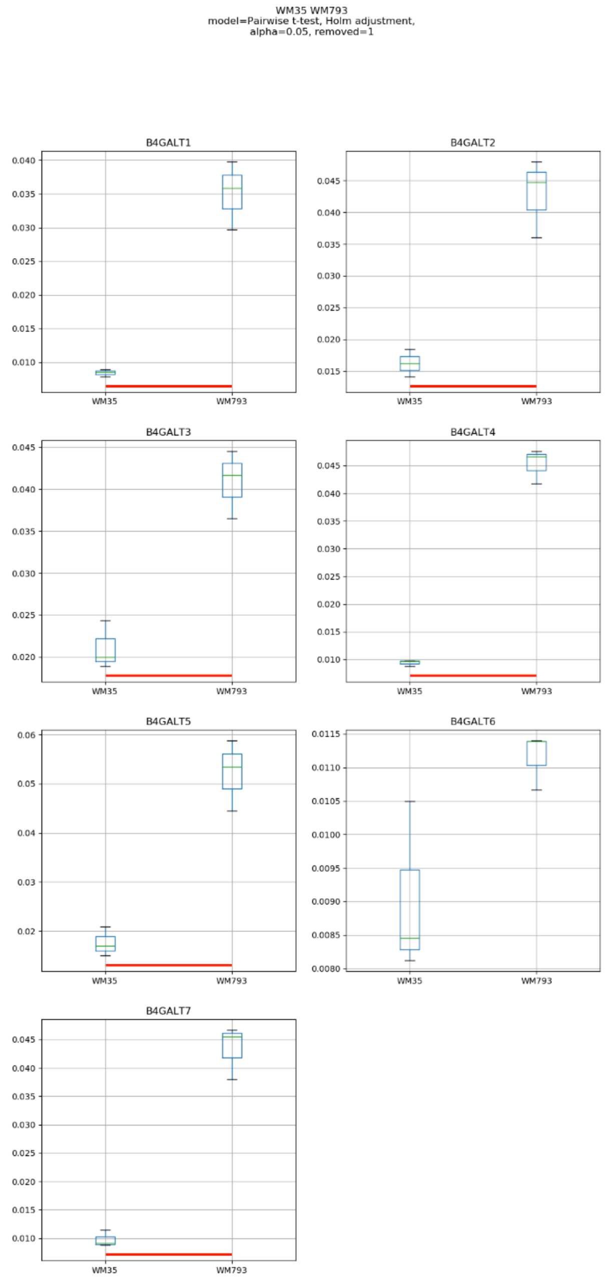
WM35 WM793
model=Pairwise t-test, Holm adjustment,
alpha=0.05, removed=1
B4GALT1
B4GALT2
B4GALT3
B4GALT4
B4GALT5
B4GALT6
B4GALT7
WM35
WM793

### Model No. 9

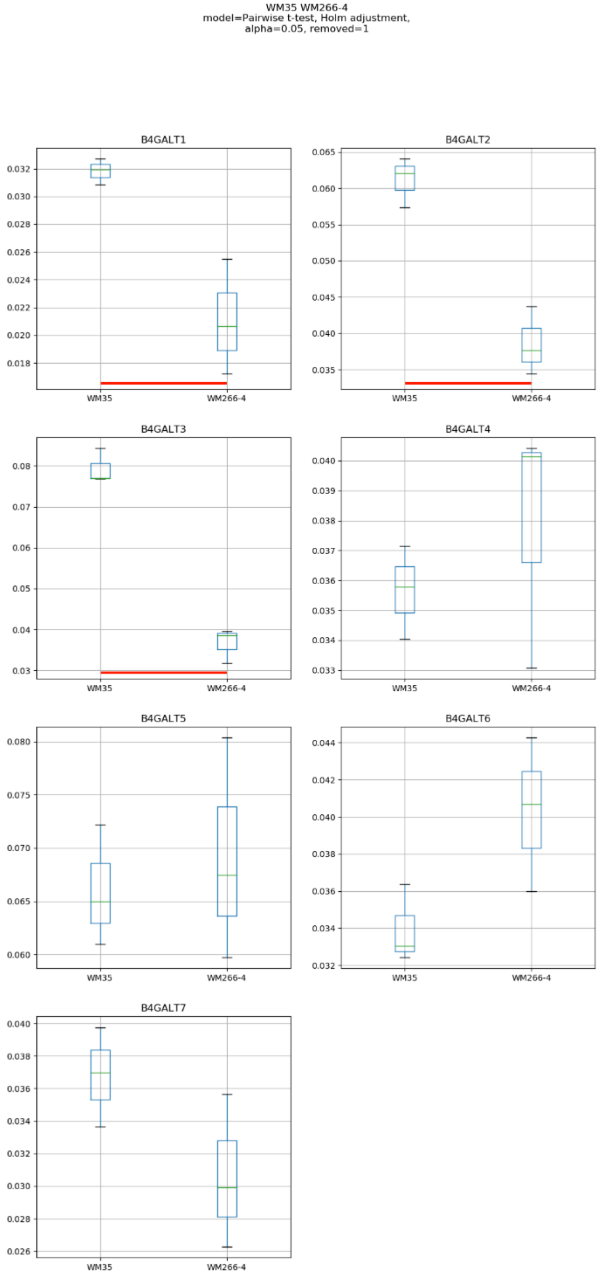
WM35 WM266-4
model=Pairwise t-test, Holm adjustment,
alpha=0.05, removed=1
B4GALT1
B4GALT2
B4GALT3
B4GALT4
B4GALT5
B4GALT6
B4GALT7
WM35
WM266-4

Model No. 10

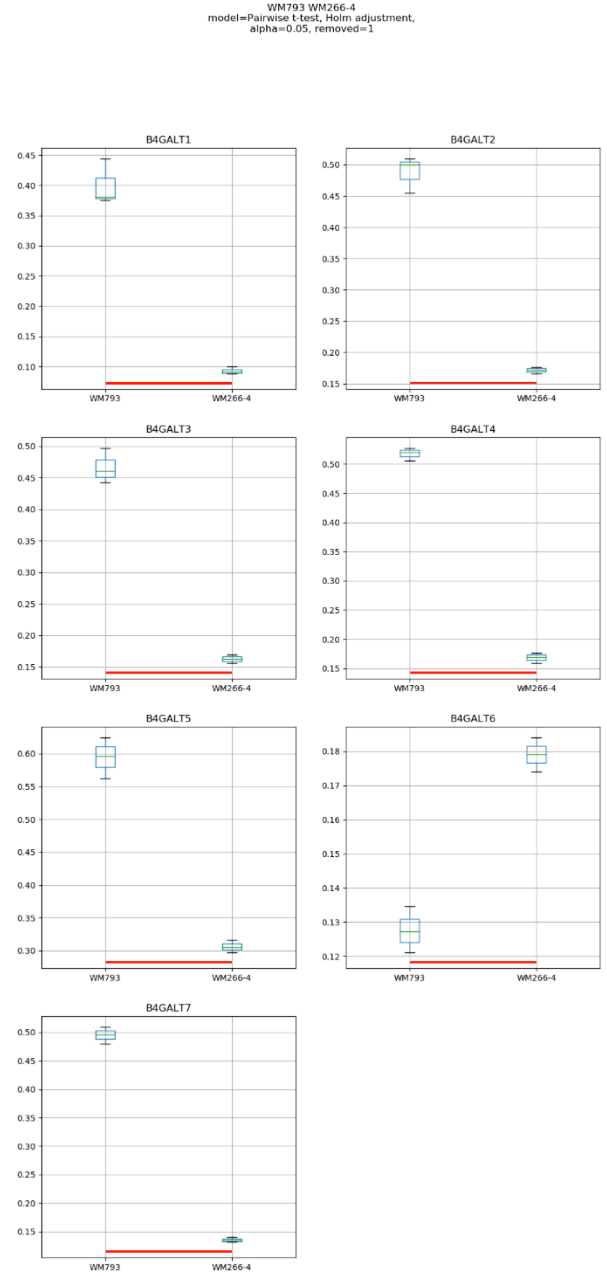
WM793 WM266-4
model=Pairwise t-test, Holm adjustment,
alpha=0.05, removed=1
B4GALT1
B4GALT2
B4GALT3
B4GALT4
B4GALT5
B4GALT6
B4GALT7
WM793
WM266-4

### Model No. 11

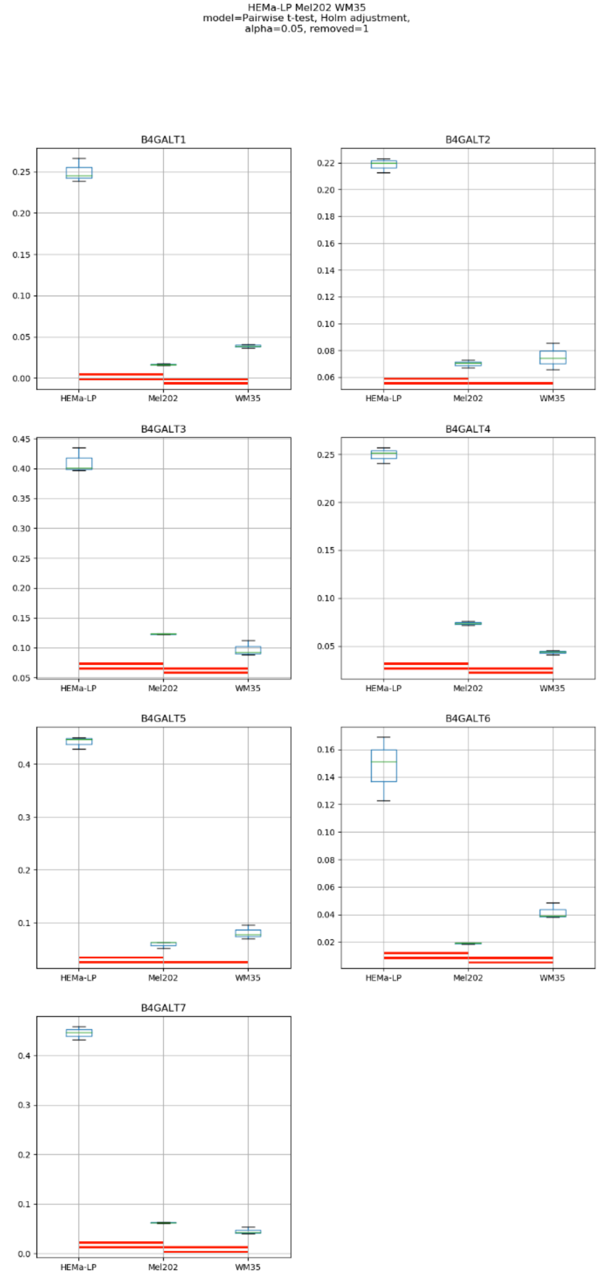
HEMa-LP Mel202 WM35
model=Pairwise t-test, Holm adjustment,
alpha=0.05, removed=1
B4GALT1
B4GALT2
B4GALT3
B4GALT4
B4GALT5
B4GALT6
B4GALT7
HEMa-LP
Mel202
WM35

### Model No. 12

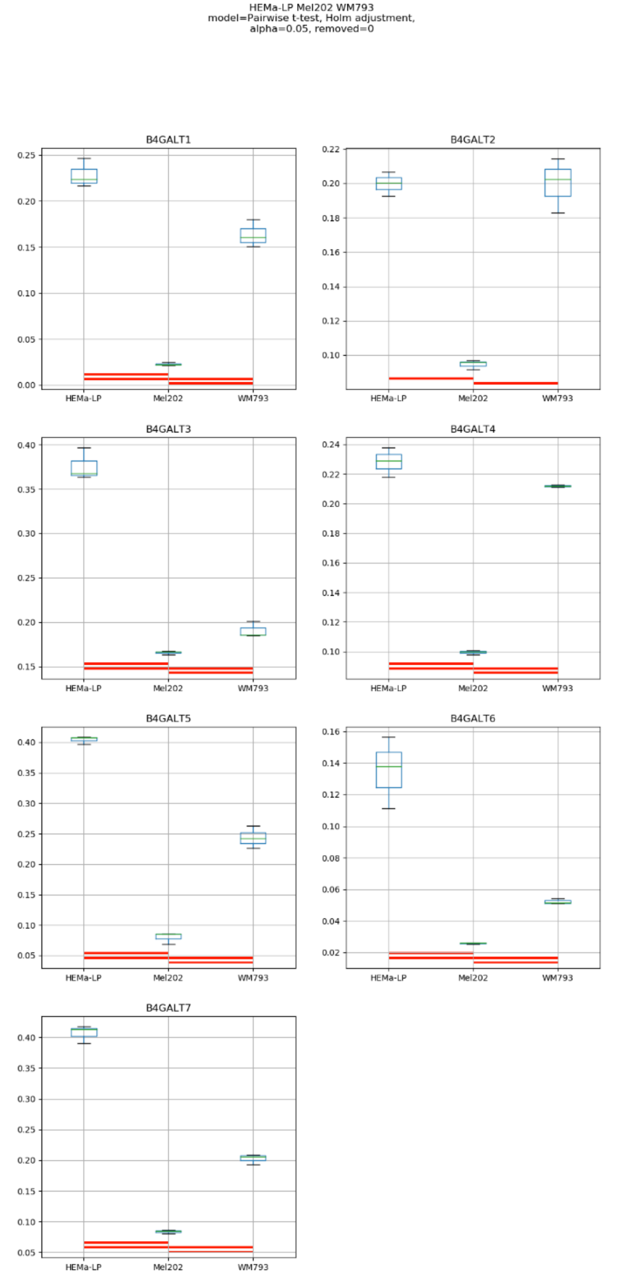
HEMa-LP Mel202 WM793
model=Pairwise t-test, Holm adjustment,
alpha=0.05, removed=0
B4GALT1
B4GALT2
B4GALT3
B4GALT4
B4GALT5
B4GALT6
B4GALT7
HEMa-LP
Mel202
WM793

### Model No. 13

HEMa-LP Mel202 WM266-4
model=Pairwise t-test, Holm adjustment,
alpha=0.05, removed=0

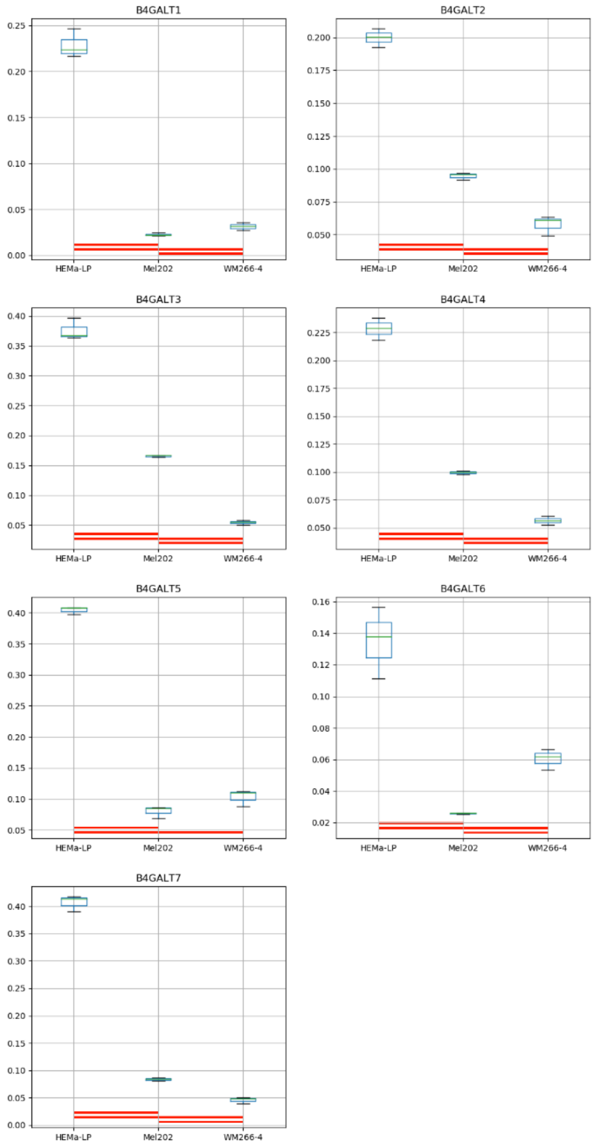

Model No. 14

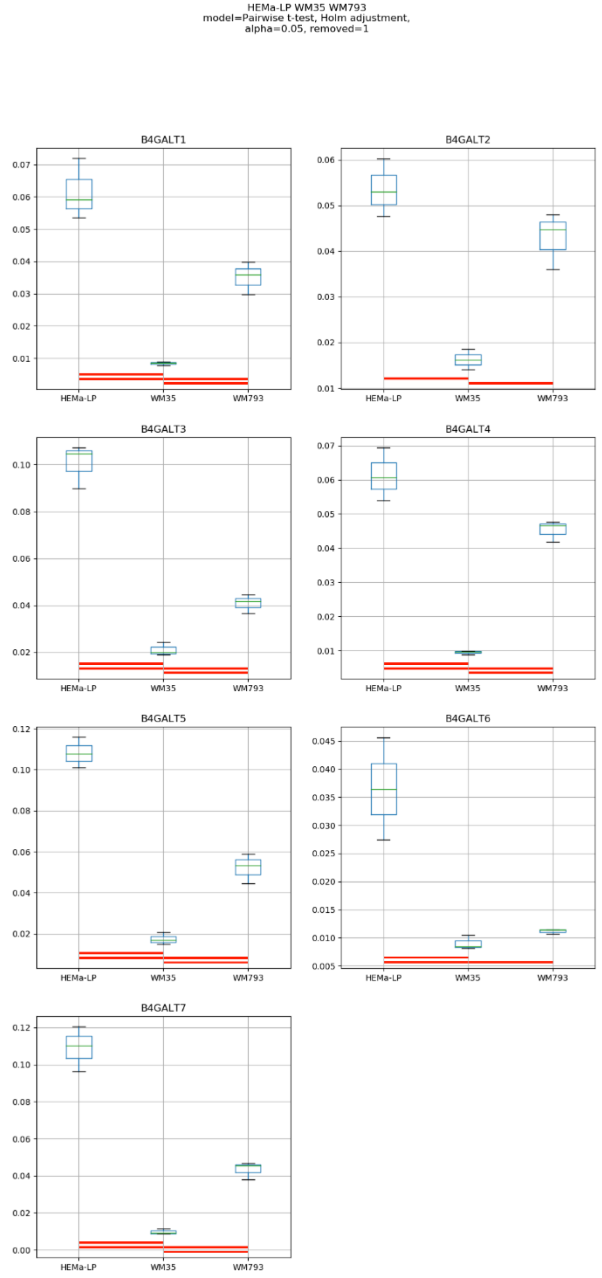
HEMa-LP WM35 WM793
model=Pairwise t-test, Holm adjustment,
alpha=0.05, removed=1
B4GALT1
B4GALT2
B4GALT3
B4GALT4
B4GALT5
B4GALT6
B4GALT7
HEMa-LP
WM35
WM793

Model No. 15

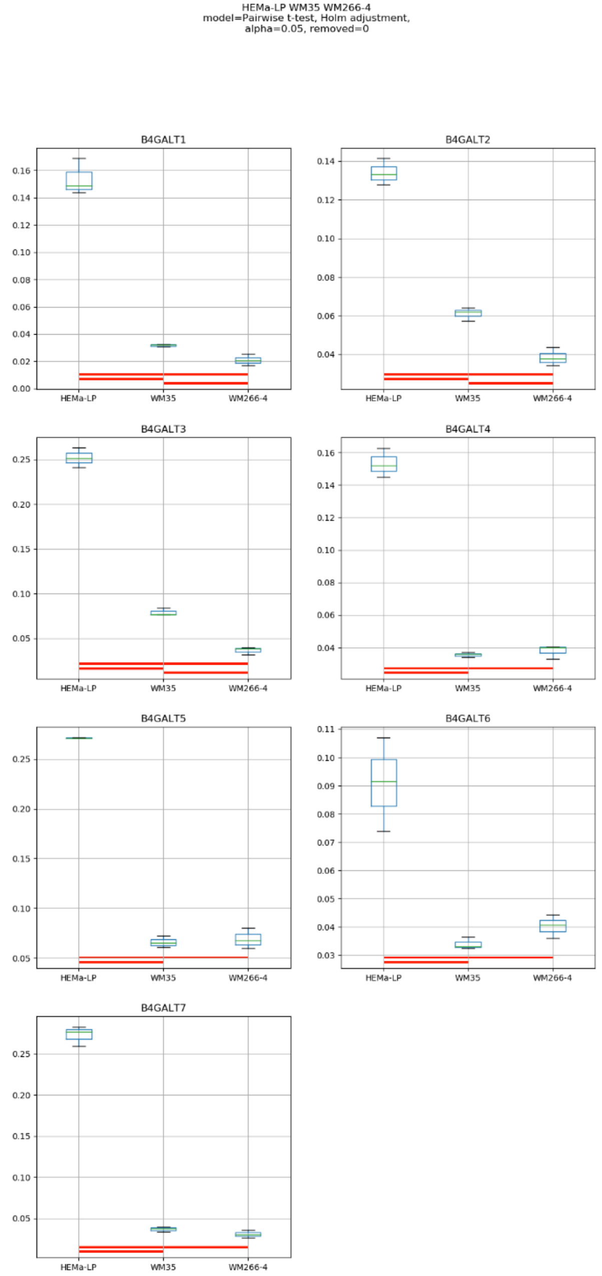
HEMa-LP WM35 WM266-4
model=Pairwise t-test, Holm adjustment,
alpha=0.05, removed=0
B4GALT1
B4GALT2
B4GALT3
B4GALT4
B4GALT5
B4GALT6
B4GALT7
HEMa-LP
WM35
WM266-4

### Model No. 16

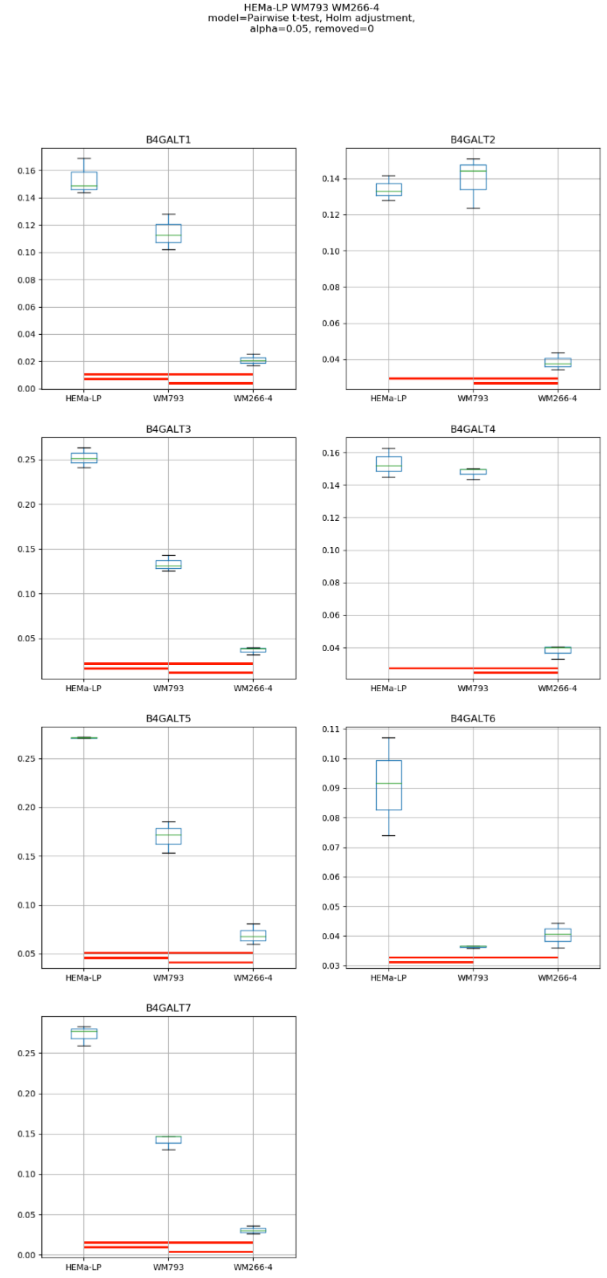
HEMa-LP WM793 WM266-4
model=Pairwise t-test, Holm adjustment,
alpha=0.05, removed=0
B4GALT1
B4GALT2
B4GALT3
B4GALT4
B4GALT5
B4GALT6
B4GALT7
HEMa-LP
WM793
WM266-4

### Model No. 17

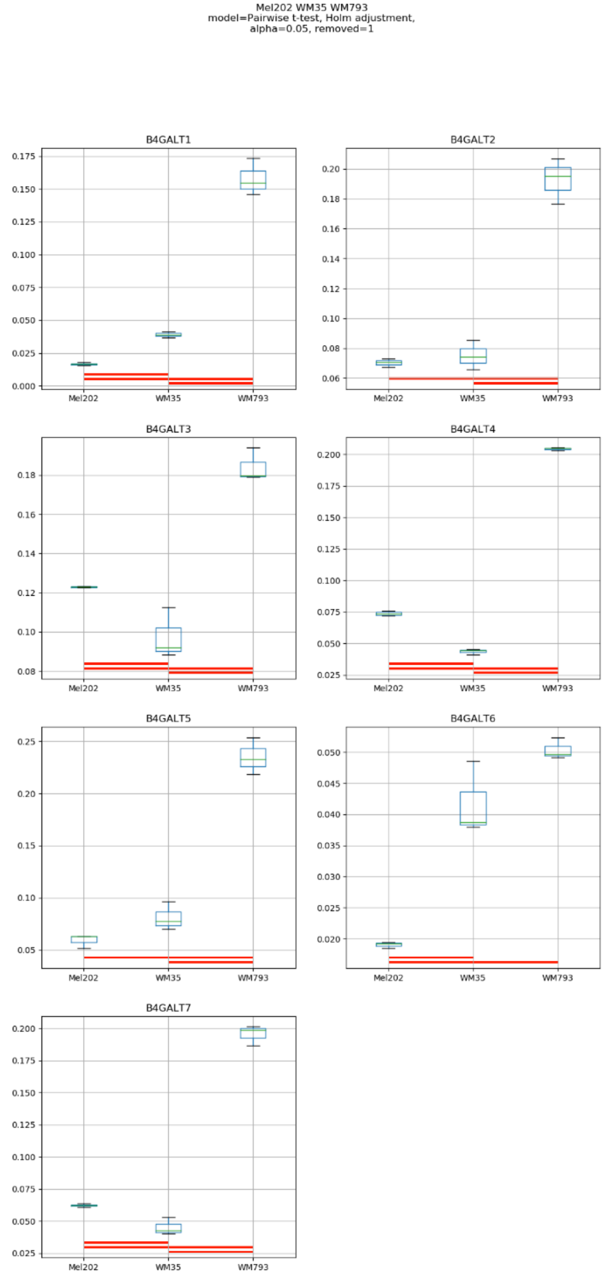
Mel202 WM35 WM793
model=Pairwise t-test, Holm adjustment,
alpha=0.05, removed=1
B4GALT1
B4GALT2
B4GALT3
B4GALT4
B4GALT5
B4GALT6
B4GALT7
Mel202
WM35
WM793

### Model No. 18

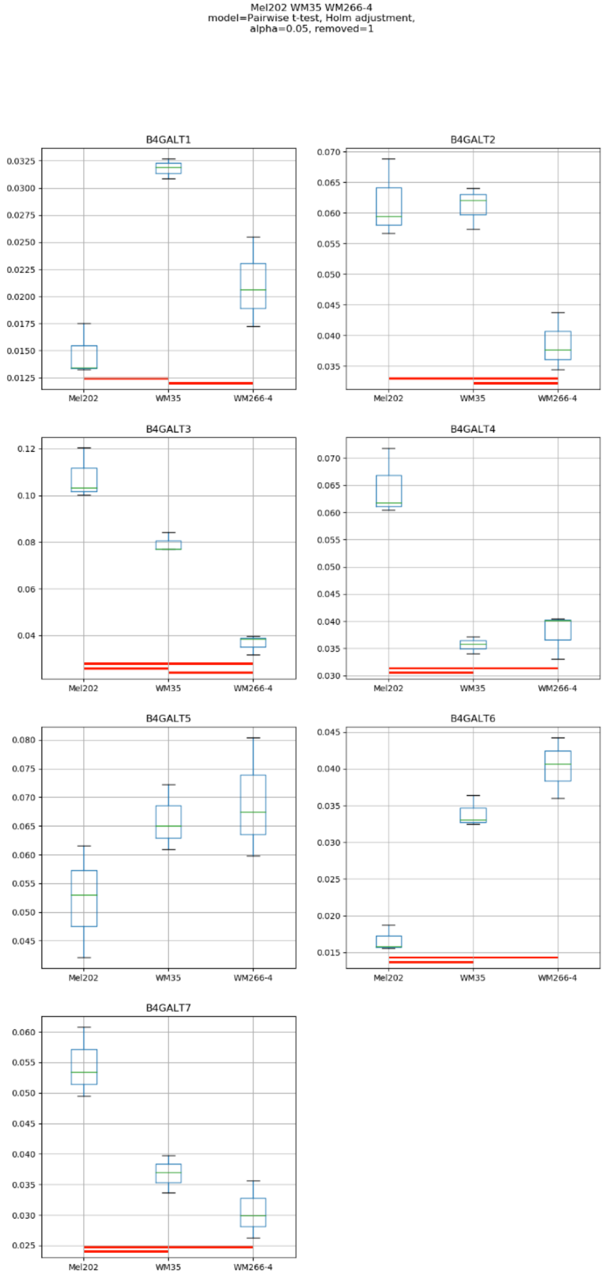
Mel202 WM35 WM266-4
model=Pairwise t-test, Holm adjustment,
alpha=0.05, removed=1
B4GALT1
B4GALT2
B4GALT3
B4GALT4
B4GALT5
B4GALT6
B4GALT7
Mel202
WM35
WM266-4

### Model No. 19

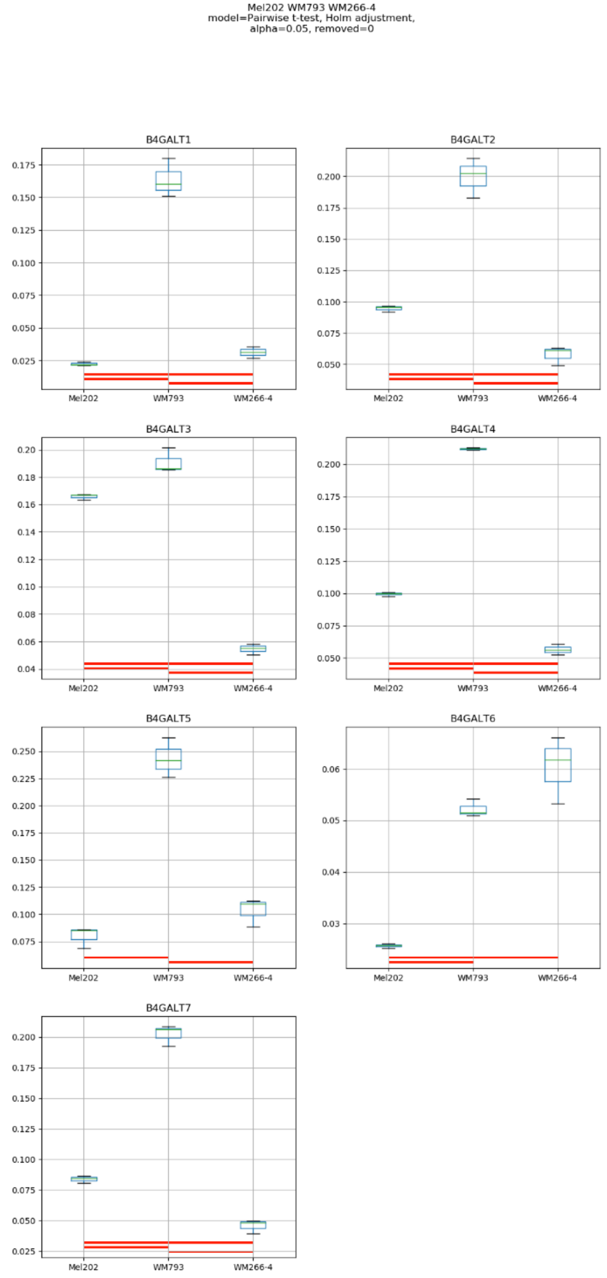
Mel202 WM793 WM266-4
model=Pairwise t-test, Holm adjustment,
alpha=0.05, removed=0
B4GALT1
B4GALT2
B4GALT3
B4GALT4
B4GALT5
B4GALT6
B4GALT7
Mel202
WM793
WM266-4

### Model No. 20

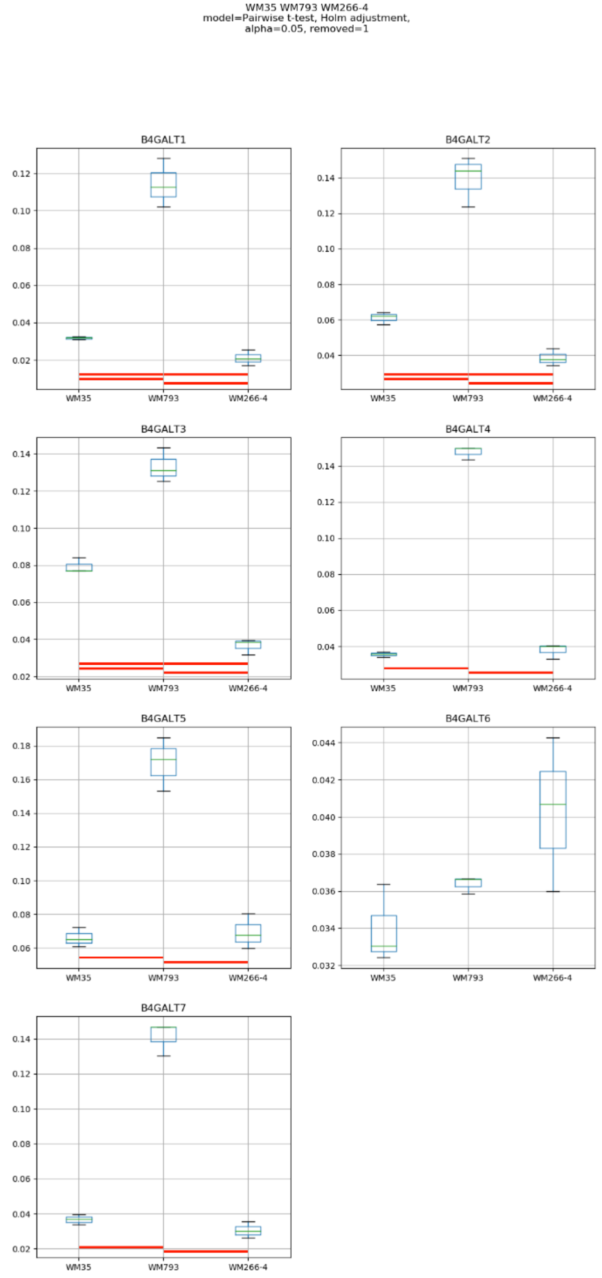
WM35 WM793 WM266-4
model=Pairwise t-test, Holm adjustment,
alpha=0.05, removed=1
B4GALT1
B4GALT2
B4GALT3
B4GALT4
B4GALT5
B4GALT6
B4GALT7
WM35
WM793
WM266-4

### Model No. 21

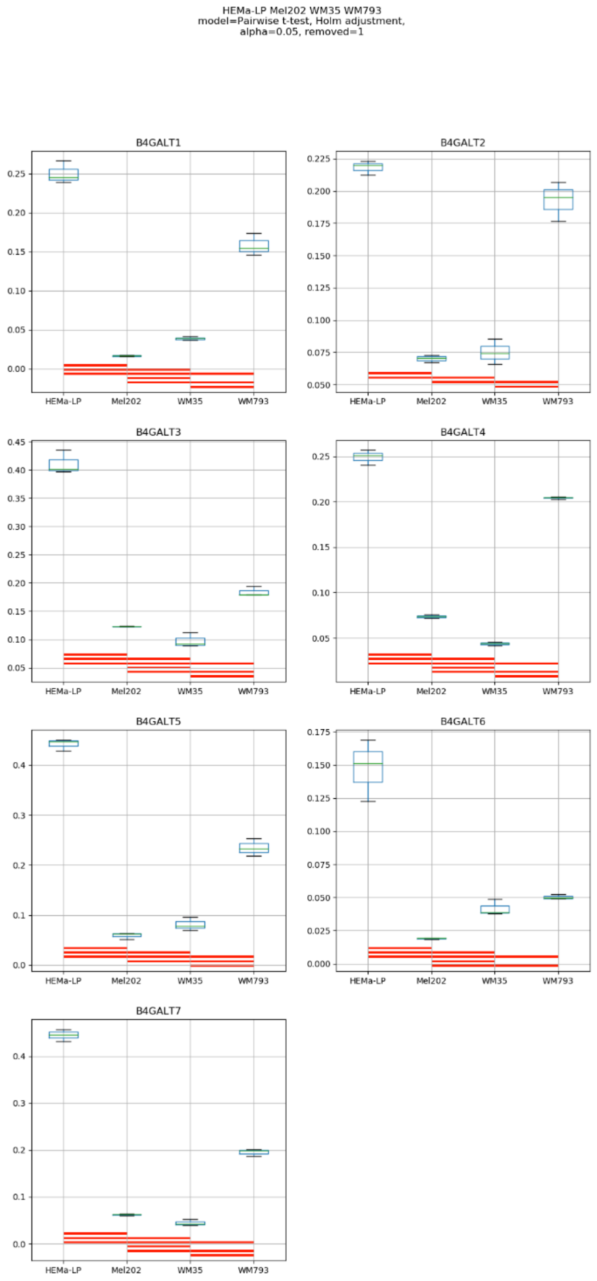
HEMa-LP Mel202 WM35 WM793
model=Pairwise t-test, Holm adjustment,
alpha=0.05, removed=1
B4GALT1
B4GALT2
B4GALT3
B4GALT4
B4GALT5
B4GALT6
B4GALT7
HEMa-LP
Mel202
WM35
WM793

### Model No. 22

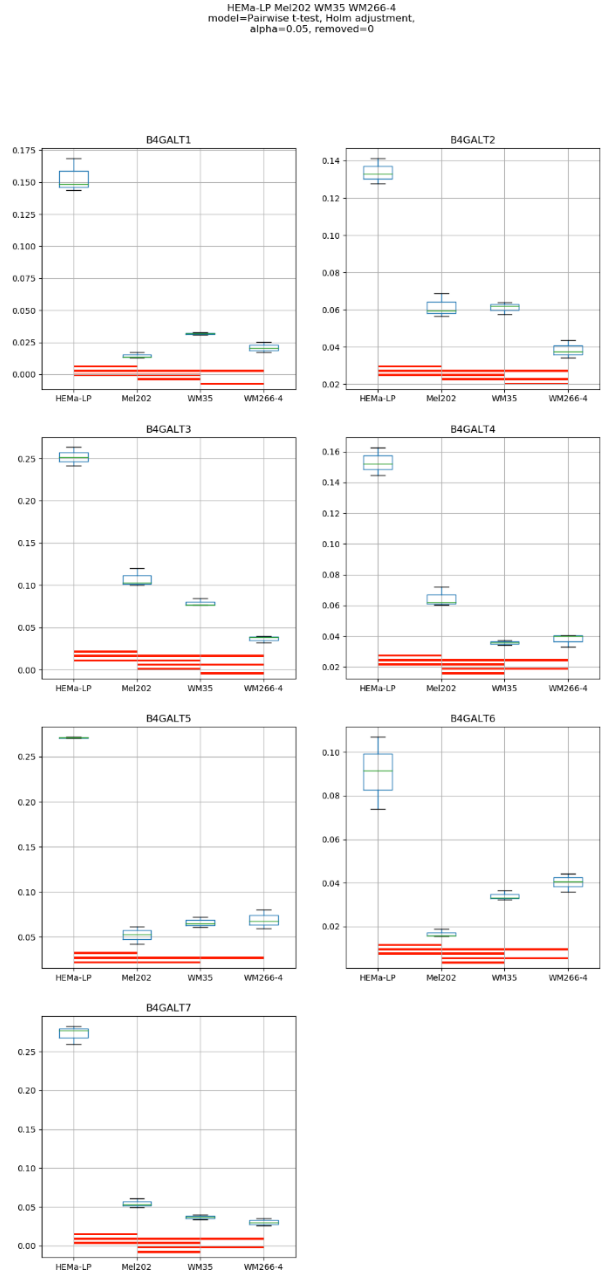
HEMa-LP Mel202 WM35 WM266-4
model=Pairwise t-test, Holm adjustment,
alpha=0.05, removed=0
B4GALT1
B4GALT2
B4GALT3
B4GALT4
B4GALT5
B4GALT6
B4GALT7
HEMa-LP
Mel202
WM35
WM266-4

### Model No. 23

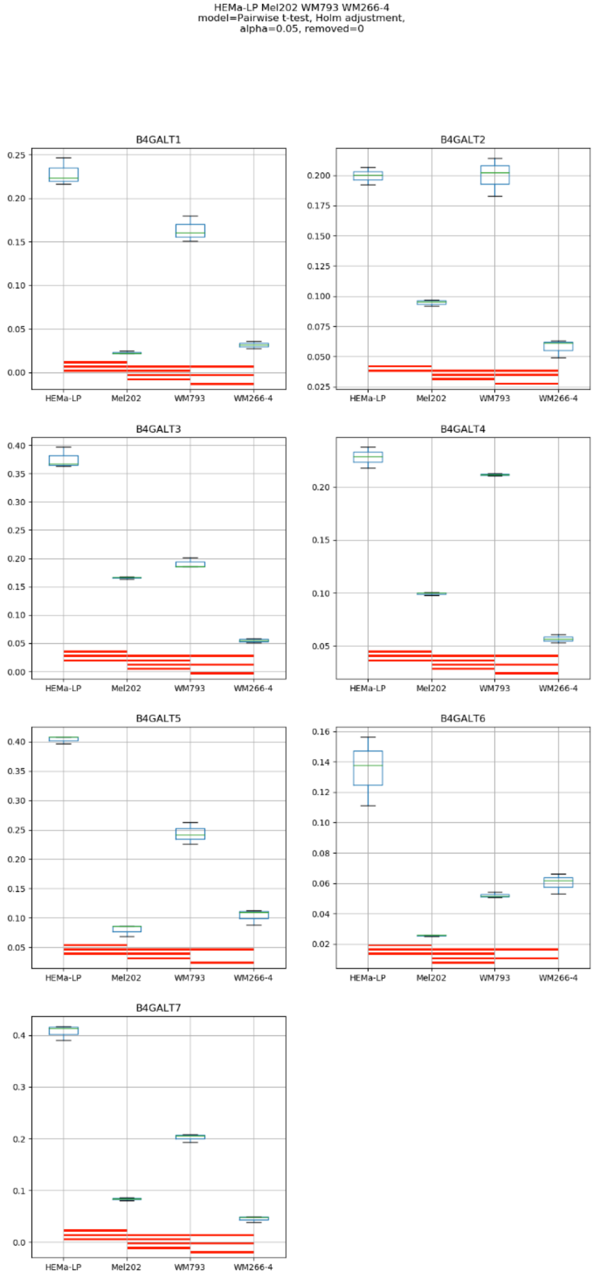
HEMa-LP Mel202 WM793 WM266-4
model=Pairwise t-test, Holm adjustment,
alpha=0.05, removed=0
B4GALT1
B4GALT2
B4GALT3
B4GALT4
B4GALT5
B4GALT6
B4GALT7
HEMa-LP
Mel202
WM793
WM266-4

### Model No. 24

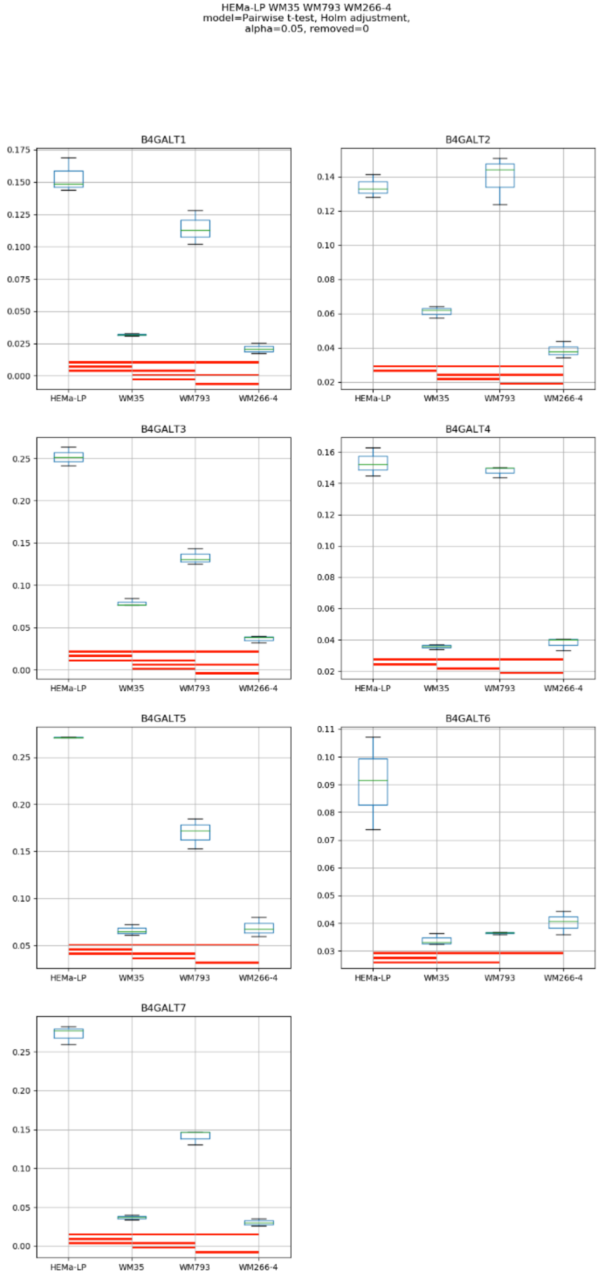
HEMa-LP WM35 WM793 WM266-4
model=Pairwise t-test, Holm adjustment,
alpha=0.05, removed=0
B4GALT1
B4GALT2
B4GALT3
B4GALT4
B4GALT5
B4GALT6
B4GALT7
HEMa-LP
WM35
WM793
WM266-4

### Model No. 25

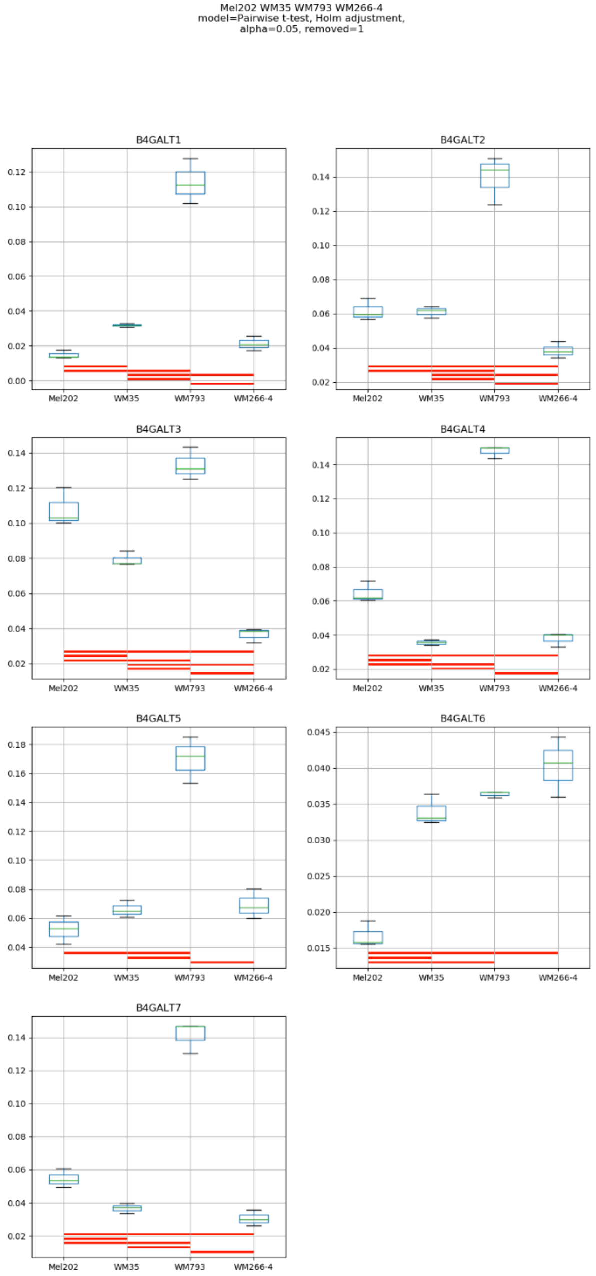
Mel202 WM35 WM793 WM266-4
model=Pairwise t-test, Holm adjustment,
alpha=0.05, removed=1
B4GALT1
B4GALT2
B4GALT3
B4GALT4
B4GALT5
B4GALT6
B4GALT7
Mel202
WM35
WM793
WM266-4

### Model No. 26

HEMa-LP Mel202 WM35 WM793 WM266-4
model=Pairwise t-test, Holm adjustment,
alpha=0.05, removed=0

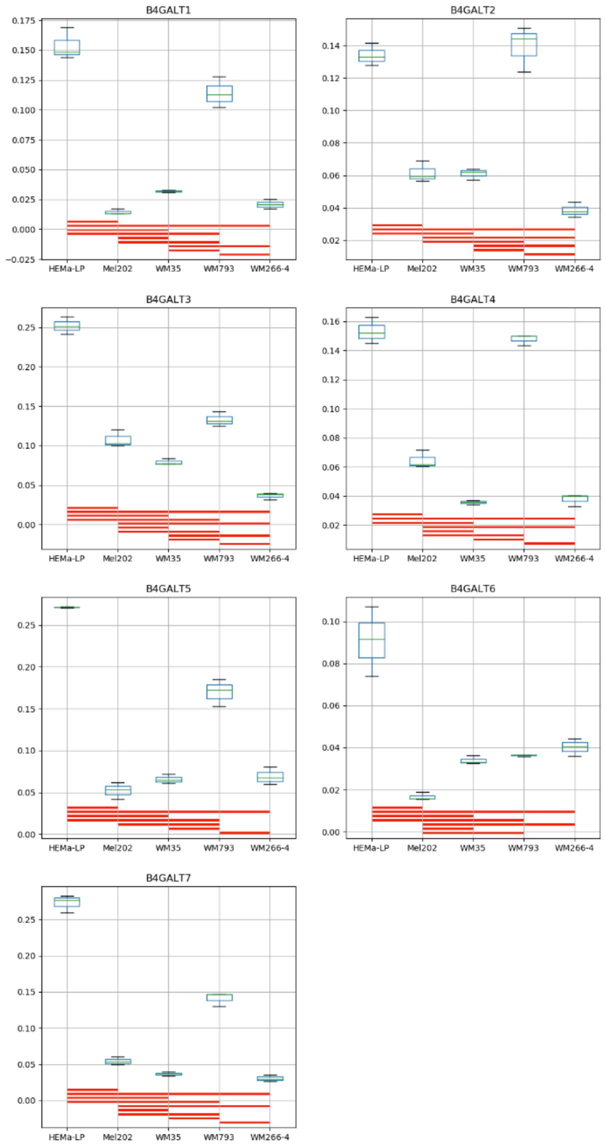

**Supplementary Figure S14:** Medians of relative quantity (RQ) for target genes *B4GALT1–B4GALT7* normalized to the reference gene or pair of reference genes with the best stability values resulting from GenExpA analysis based on set 3 of four candidate reference genes (*SNRPA*, *HPRT1*, *GUSB*, *PGK1*) and remove repetition level 1. The statistical analyses used the pairwise t-test with Holm adjustment. Red line represents statistical significance, $P < 0.05$. Detailed results are listed in Supplementary Table 13.

### Model No. 1

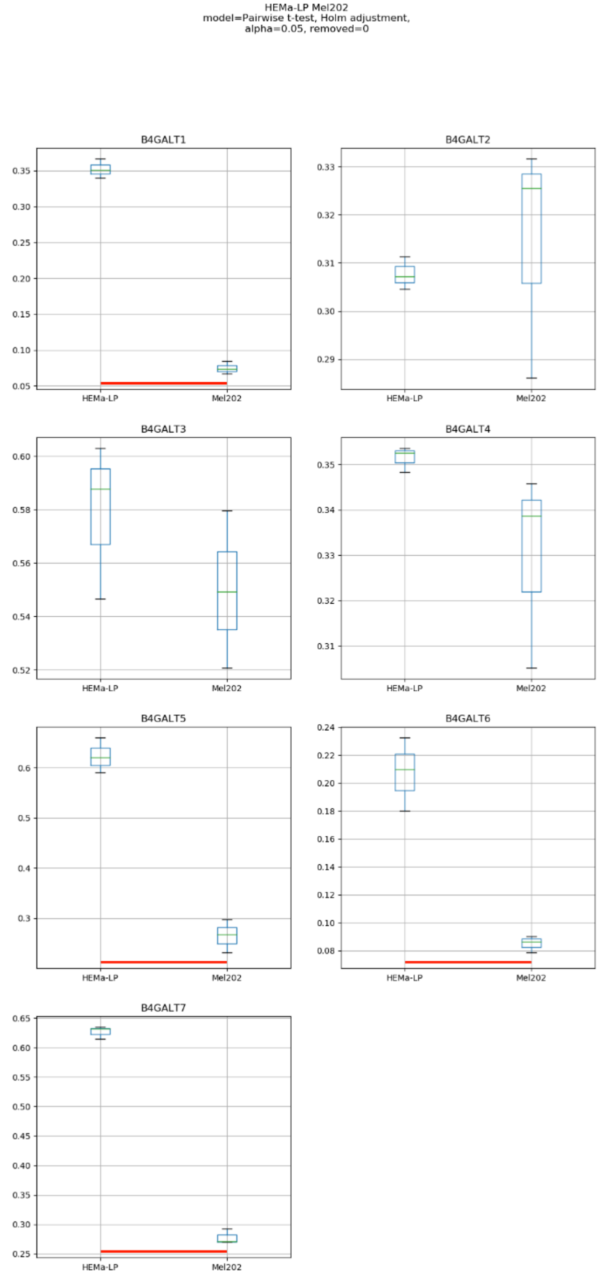
HEMa-LP Mel202
model=Pairwise t-test, Holm adjustment,
alpha=0.05, removed=0
B4GALT1
B4GALT2
B4GALT3
B4GALT4
B4GALT5
B4GALT6
B4GALT7
HEMa-LP
Mel202

### Model No. 2

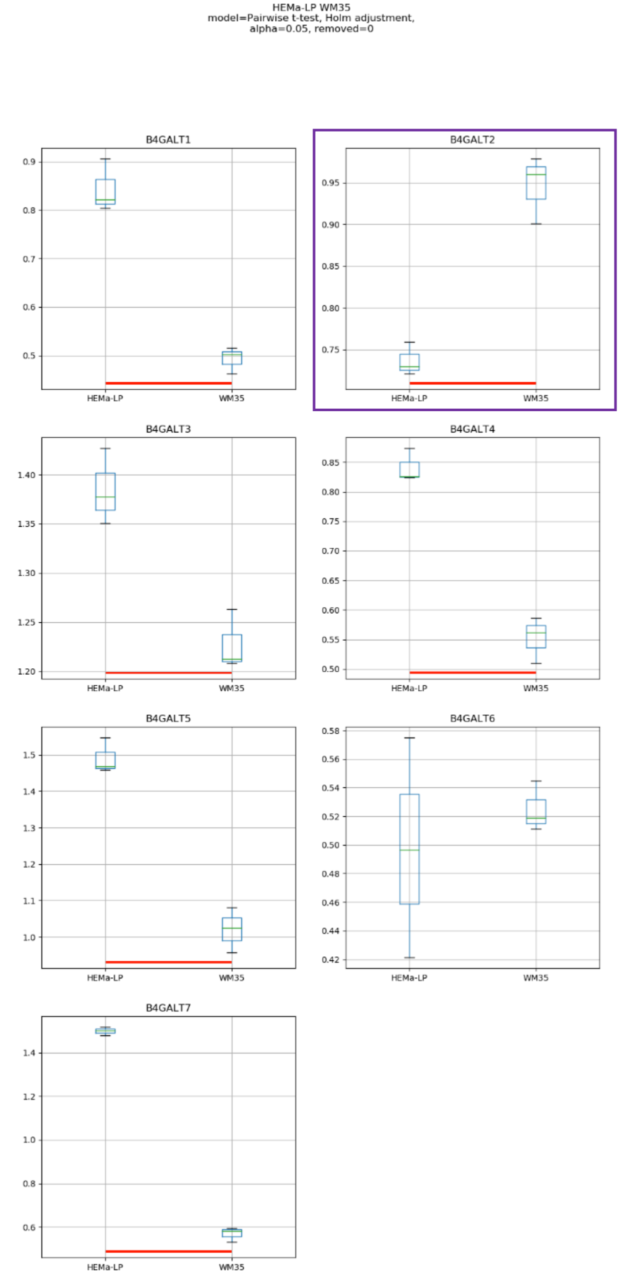

HEMa-LP WM35
model=Pairwise t-test, Holm adjustment,
alpha=0.05, removed=0
B4GALT1
B4GALT2
B4GALT3
B4GALT4
B4GALT5
B4GALT6
B4GALT7
HEMa-LP
WM35

### Model No. 3

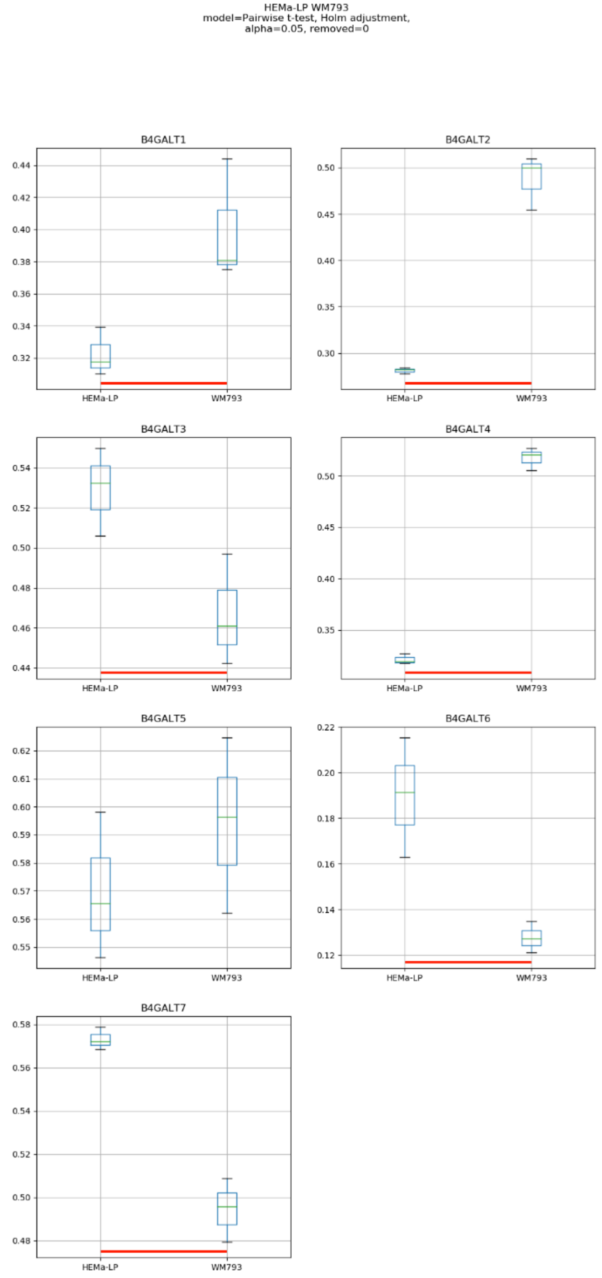
HEMa-LP WM793
model=Pairwise t-test, Holm adjustment,
alpha=0.05, removed=0
B4GALT1
B4GALT2
B4GALT3
B4GALT4
B4GALT5
B4GALT6
B4GALT7
HEMa-LP
WM793

### Model No. 4

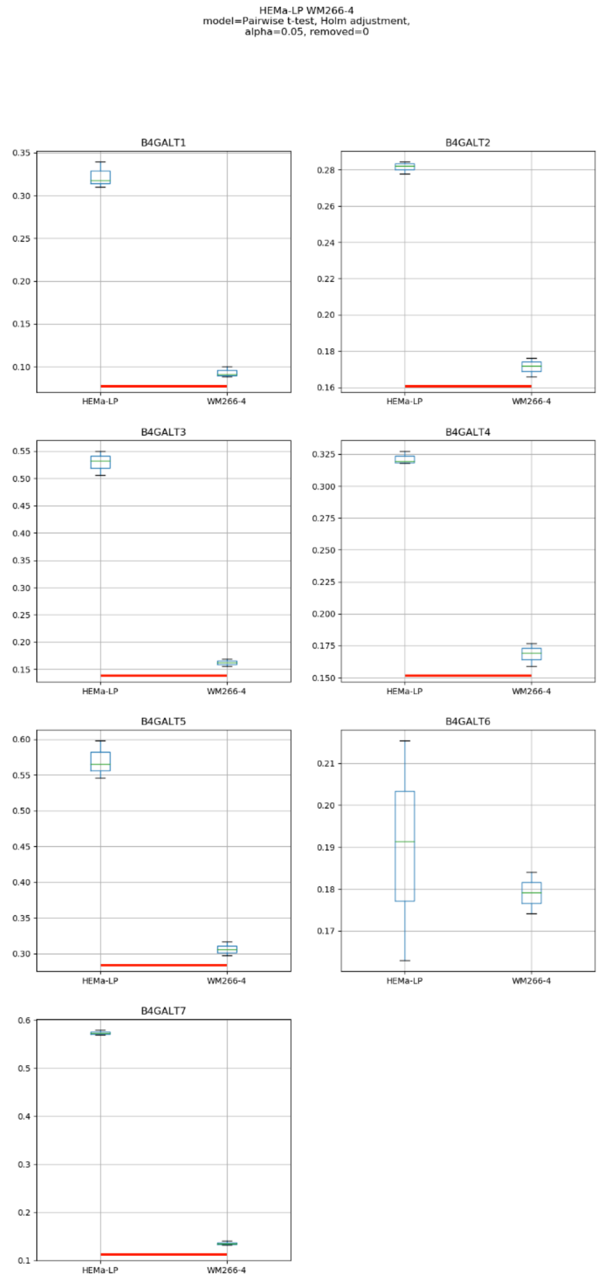
HEMa-LP WM266-4
model=Pairwise t-test, Holm adjustment,
alpha=0.05, removed=0
B4GALT1
B4GALT2
B4GALT3
B4GALT4
B4GALT5
B4GALT6
B4GALT7
HEMa-LP
WM266-4

### Model No. 5

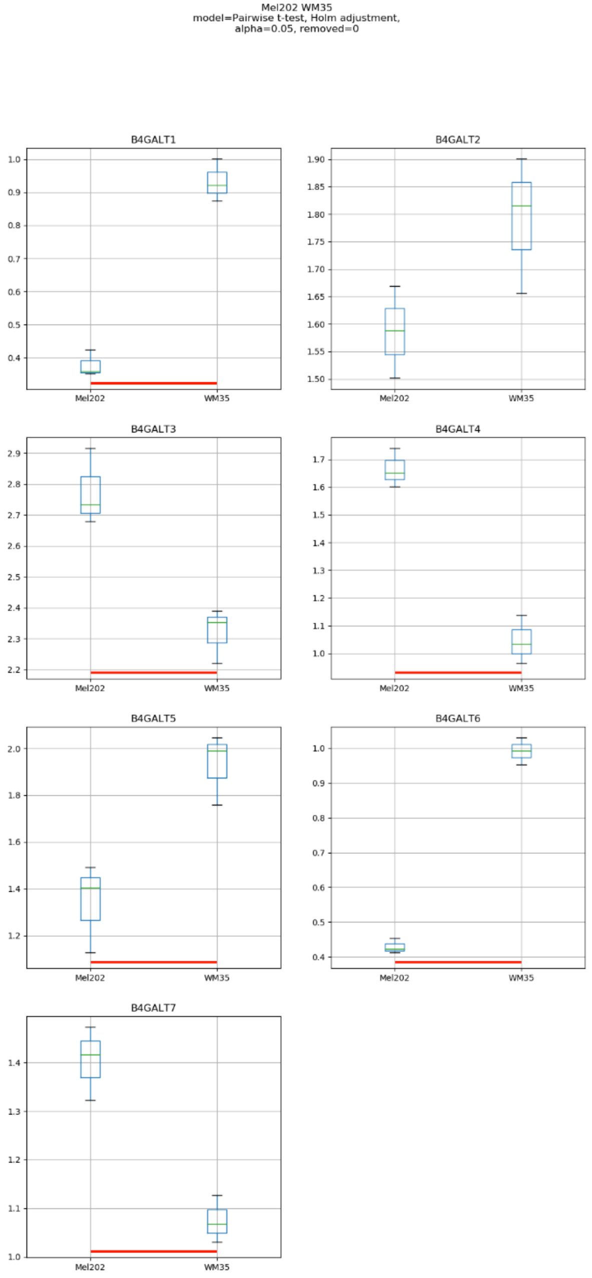
Mel202 WM35
model=Pairwise t-test, Holm adjustment,
alpha=0.05, removed=0
B4GALT1
B4GALT2
B4GALT3
B4GALT4
B4GALT5
B4GALT6
B4GALT7
Mel202
WM35

### Model No. 6

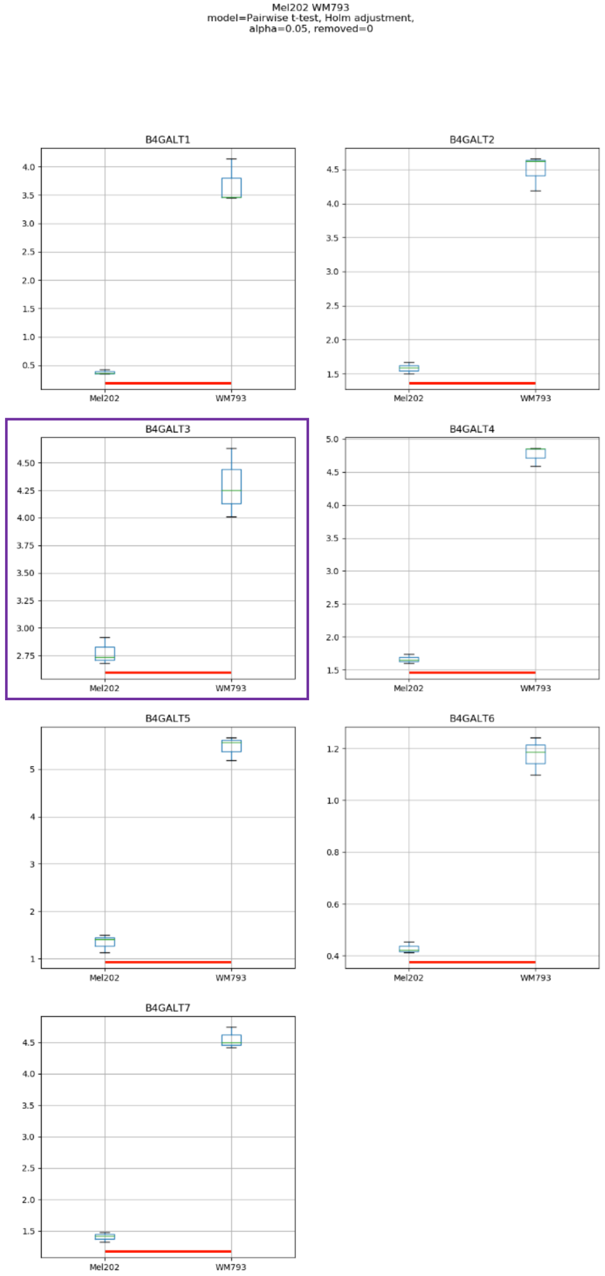
Mel202 WM793
model=Pairwise t-test, Holm adjustment,
alpha=0.05, removed=0
B4GALT1
B4GALT2
B4GALT3
B4GALT4
B4GALT5
B4GALT6
B4GALT7
Mel202
WM793

Model No. 7

Mel202 WM266-4
model=Pairwise t-test, Holm adjustment,
alpha=0.05, removed=0

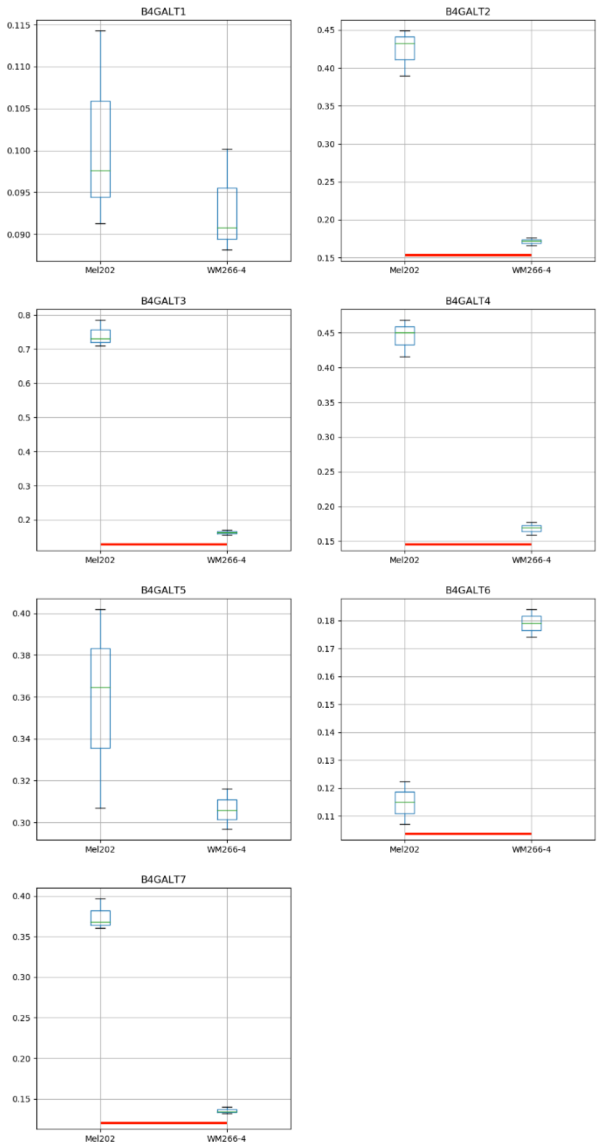

### Model No. 8

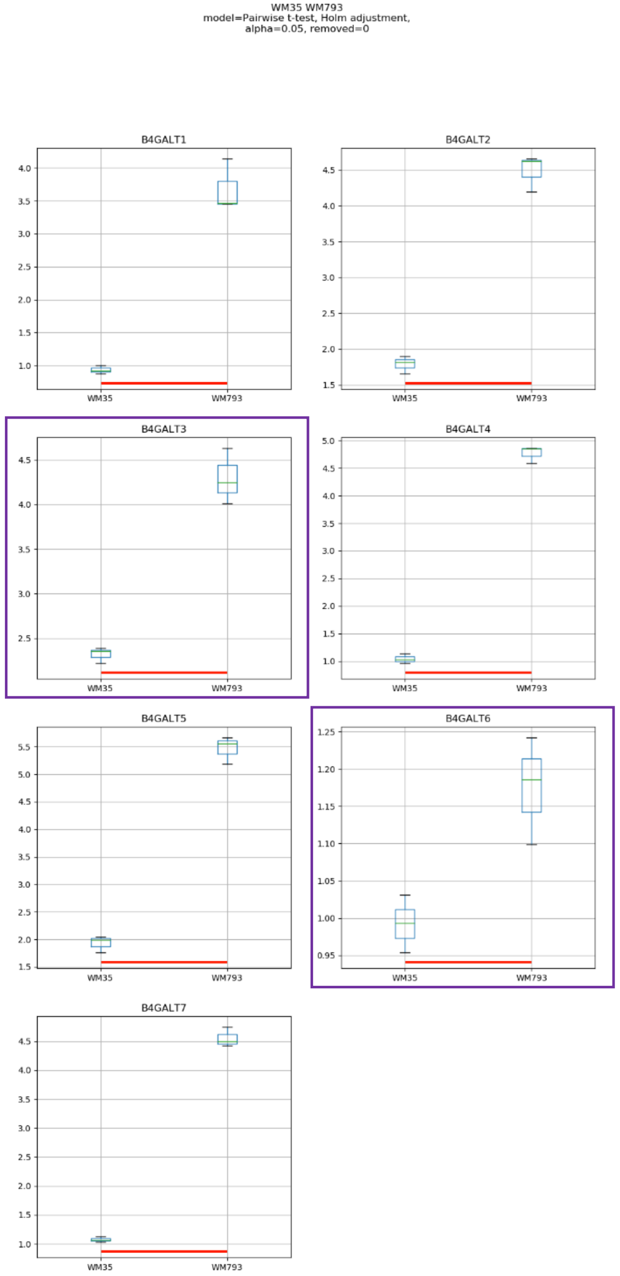
WM35 WM793
model=Pairwise t-test, Holm adjustment,
alpha=0.05, removed=0
B4GALT1
B4GALT2
B4GALT3
B4GALT4
B4GALT5
B4GALT6
B4GALT7
WM35
WM793

Model No. 9

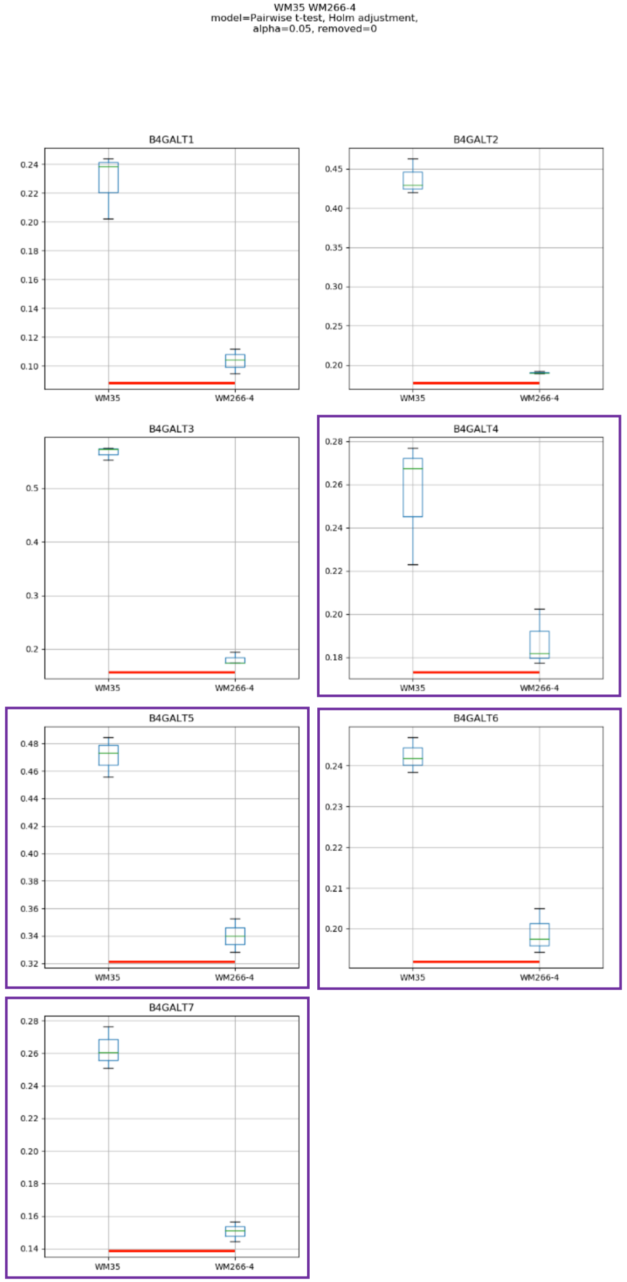
WM35 WM266-4
model=Pairwise t-test, Holm adjustment,
alpha=0.05, removed=0
B4GALT1
B4GALT2
B4GALT3
B4GALT4
B4GALT5
B4GALT6
B4GALT7
WM35
WM266-4

Model No. 10

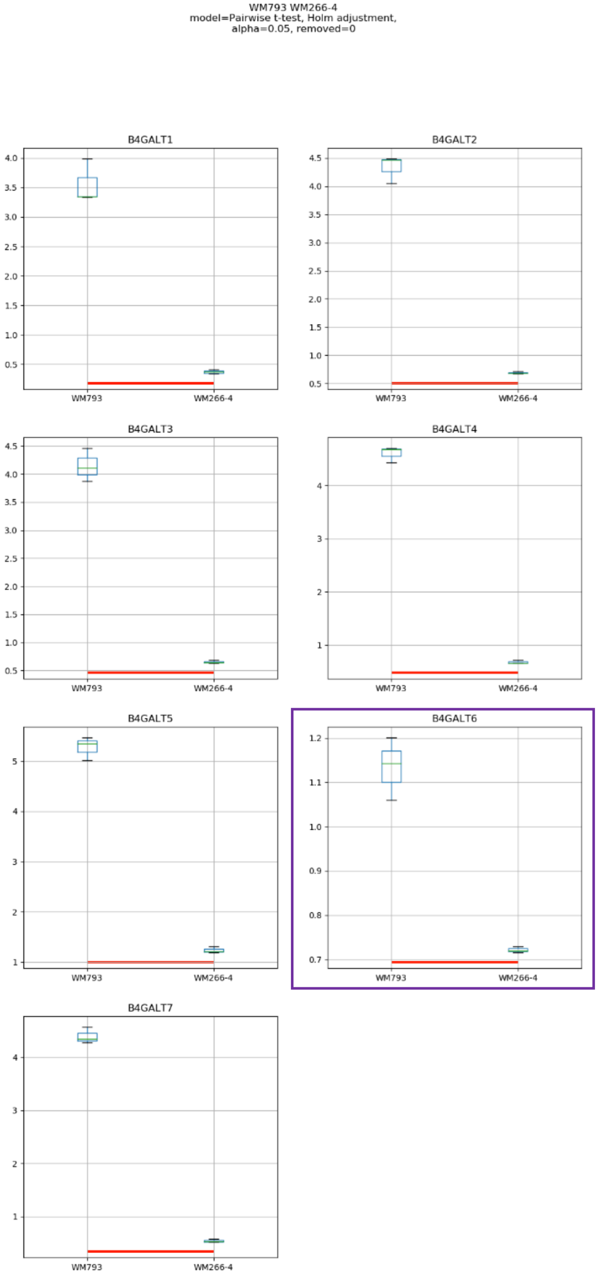

WM793 WM266-4
model=Pairwise t-test, Holm adjustment,
alpha=0.05, removed=0
B4GALT1
B4GALT2
B4GALT3
B4GALT4
B4GALT5
B4GALT6
B4GALT7
WM793
WM266-4

### Model No. 11

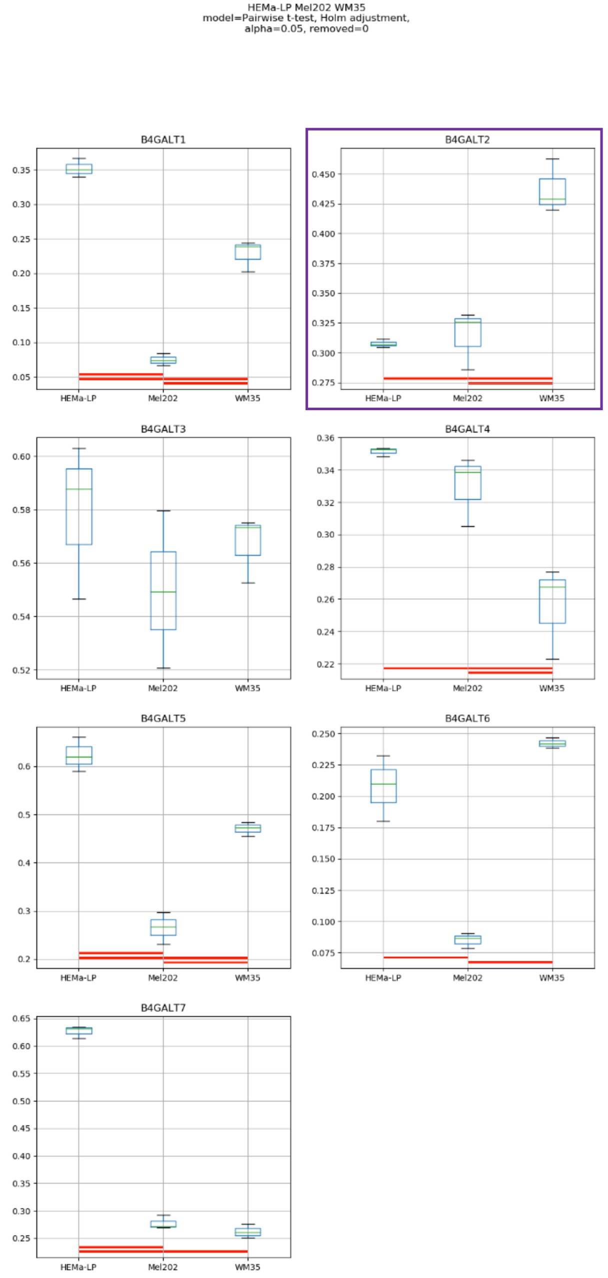

HEMa-LP Mel202 WM35
model=Pairwise t-test, Holm adjustment,
alpha=0.05, removed=0
B4GALT1
B4GALT2
B4GALT3
B4GALT4
B4GALT5
B4GALT6
B4GALT7
HEMa-LP
Mel202
WM35

### Model No. 12

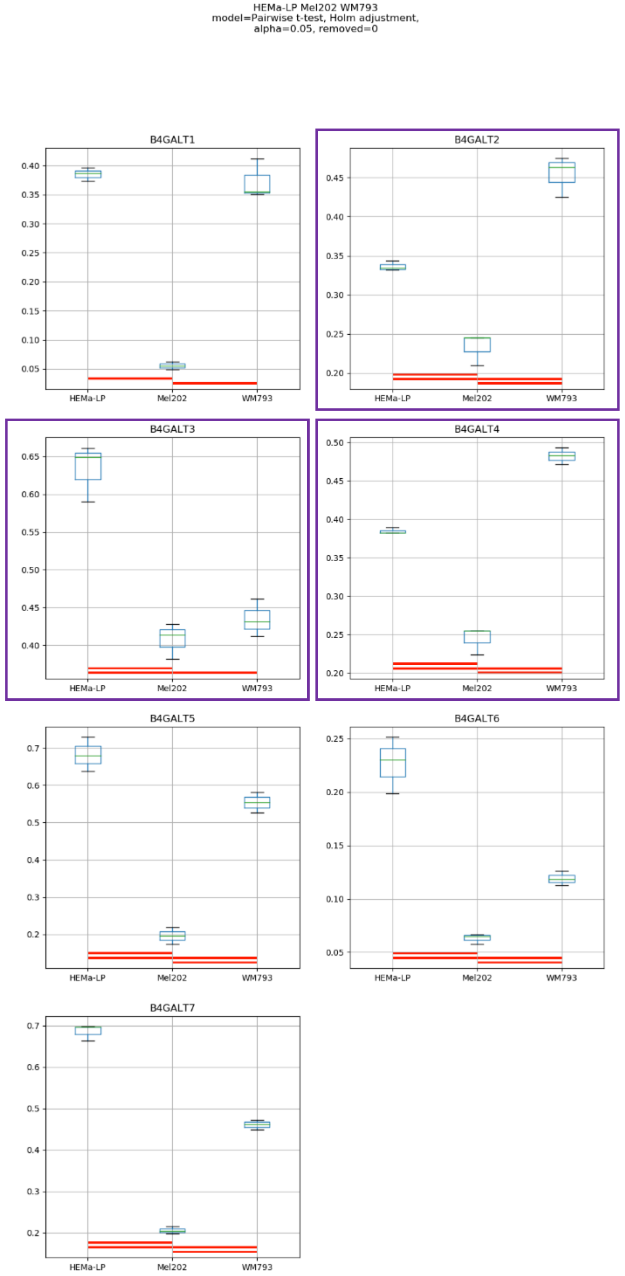

HEMa-LP Mel202 WM793
model=Pairwise t-test, Holm adjustment,
alpha=0.05, removed=0
B4GALT1
B4GALT2
B4GALT3
B4GALT4
B4GALT5
B4GALT6
B4GALT7
HEMa-LP
Mel202
WM793

### Model No. 13

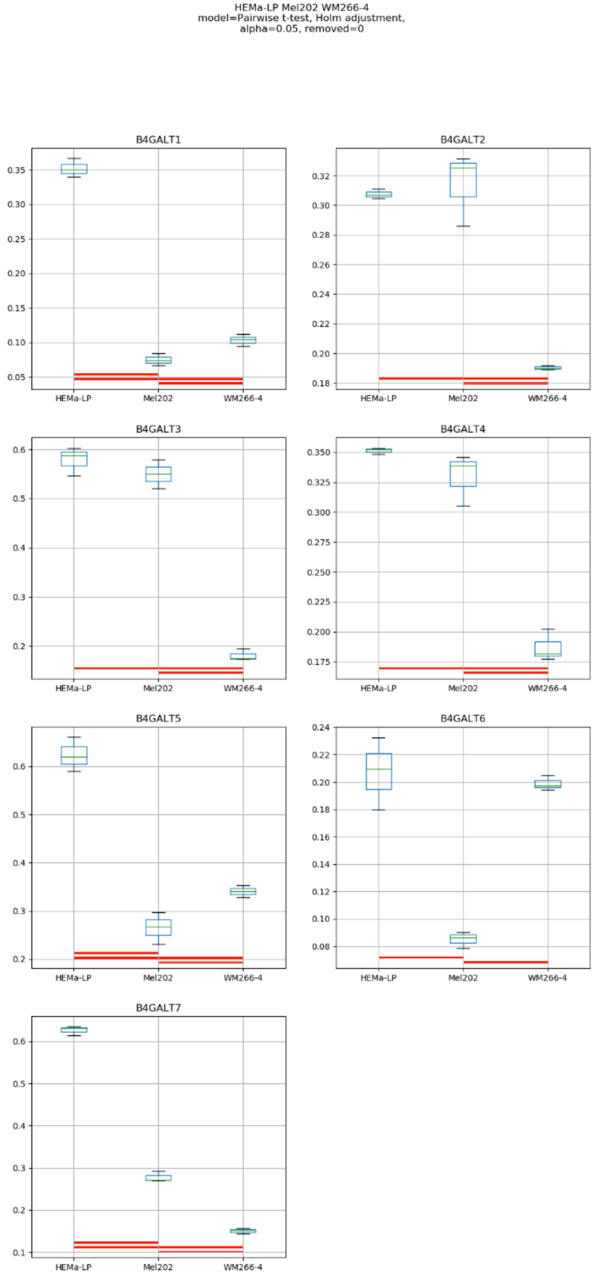
HEMa-LP Mel202 WM266-4
model=Pairwise t-test, Holm adjustment,
alpha=0.05, removed=0
B4GALT1
B4GALT2
B4GALT3
B4GALT4
B4GALT5
B4GALT6
B4GALT7
HEMa-LP
Mel202
WM266-4

### Model No. 14

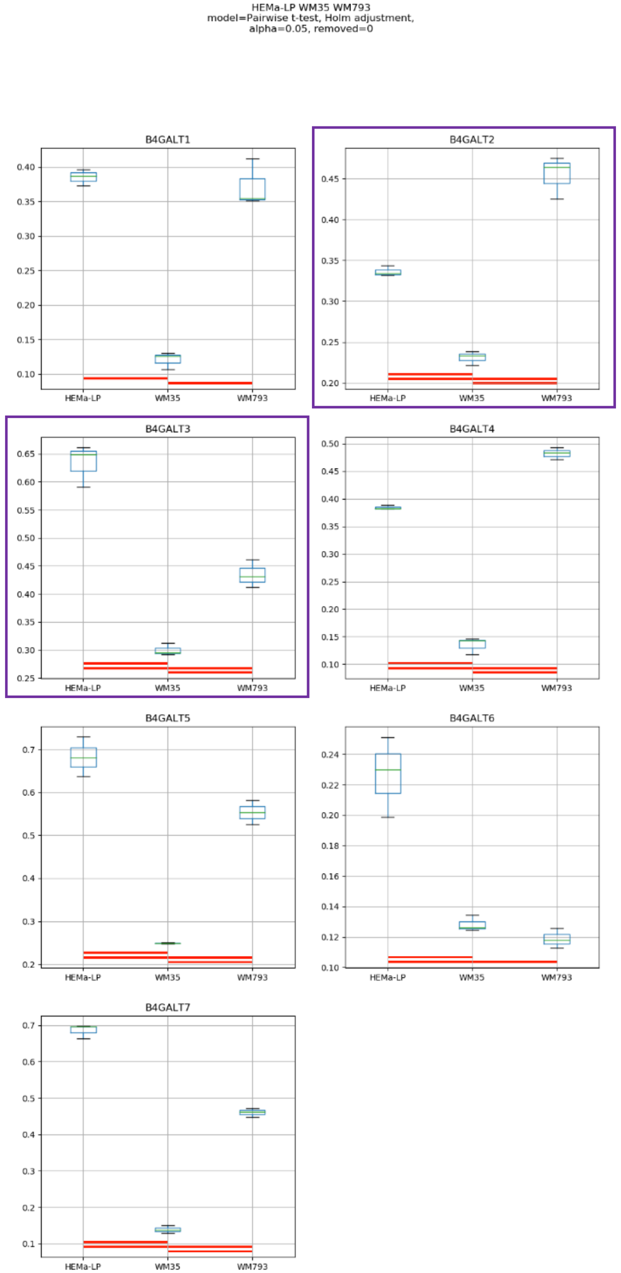
HEMa-LP WM35 WM793
model=Pairwise t-test, Holm adjustment,
alpha=0.05, removed=0
B4GALT1
B4GALT2
B4GALT3
B4GALT4
B4GALT5
B4GALT6
B4GALT7
HEMa-LP
WM35
WM793

Model No. 15

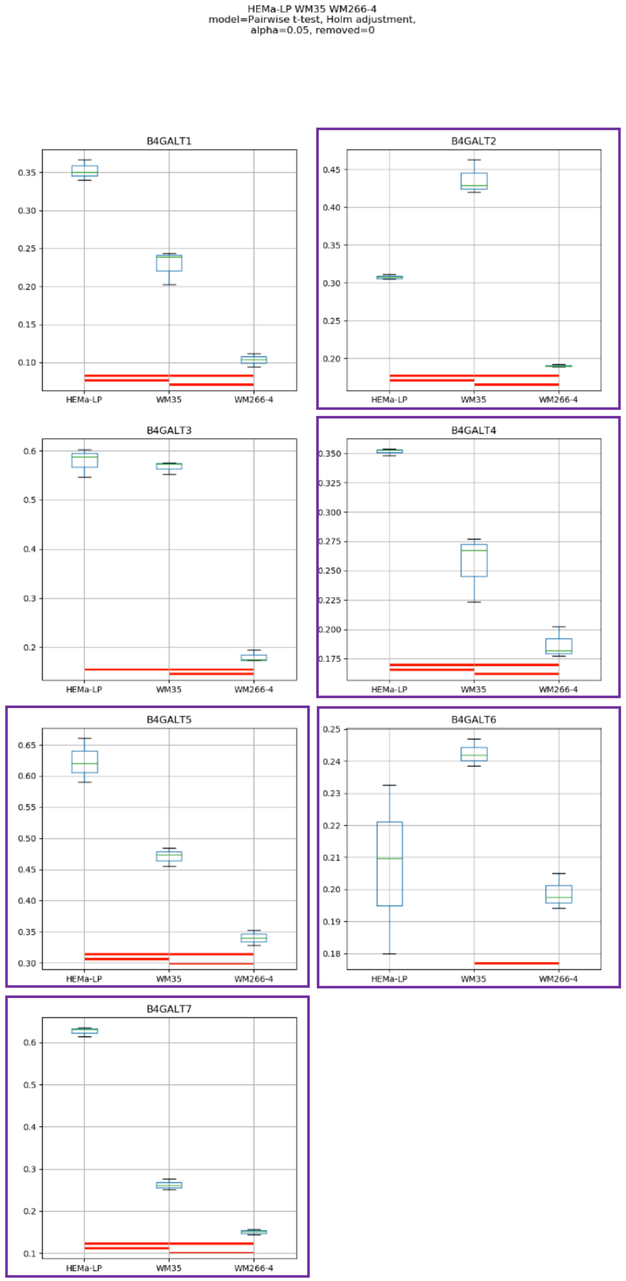

HEMa-LP WM35 WM266-4
model=Pairwise t-test, Holm adjustment,
alpha=0.05, removed=0
B4GALT1
B4GALT2
B4GALT3
B4GALT4
B4GALT5
B4GALT6
B4GALT7
HEMa-LP
WM35
WM266-4

### Model No. 16

HEMa-LP WM793 WM266-4
model=Pairwise t-test, Holm adjustment,
alpha=0.05, removed=0

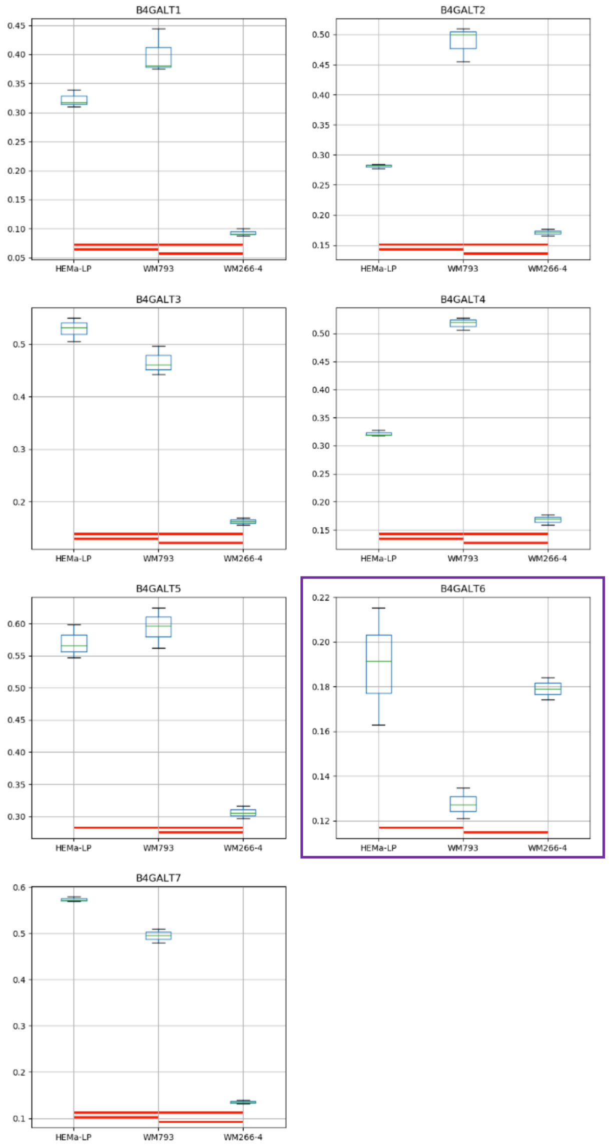

Model No. 17

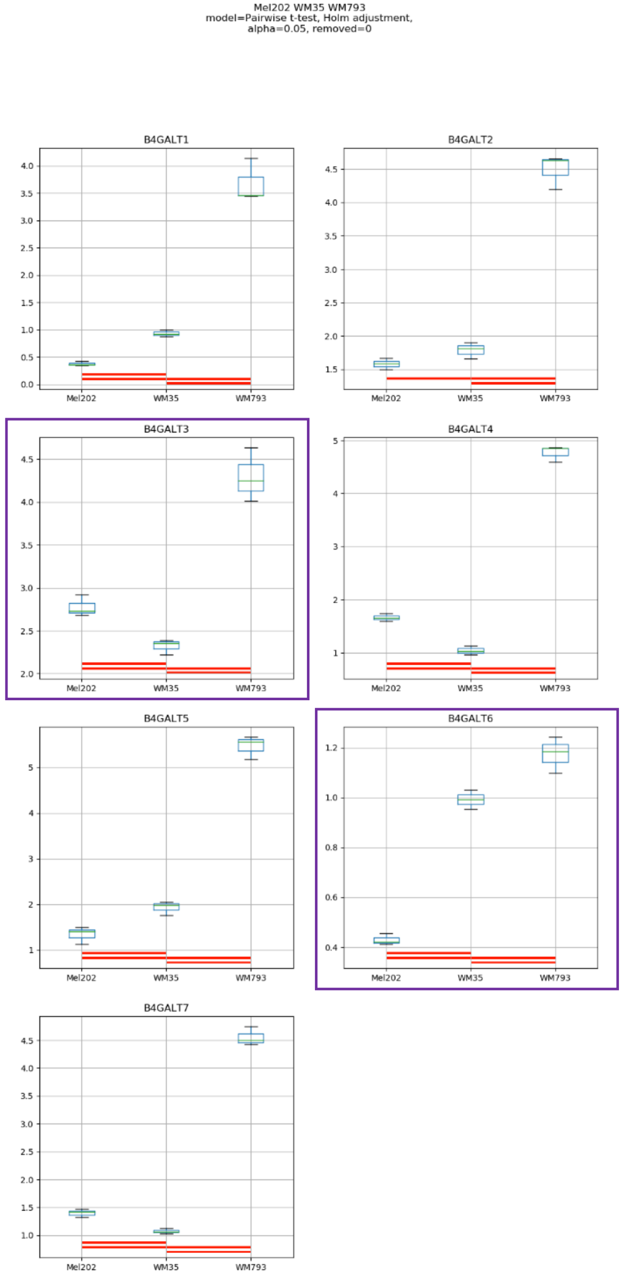
Mel202 WM35 WM793
model=Pairwise t-test, Holm adjustment,
alpha=0.05, removed=0
B4GALT1
B4GALT2
B4GALT3
B4GALT4
B4GALT5
B4GALT6
B4GALT7
Mel202
WM35
WM793

Model No. 18

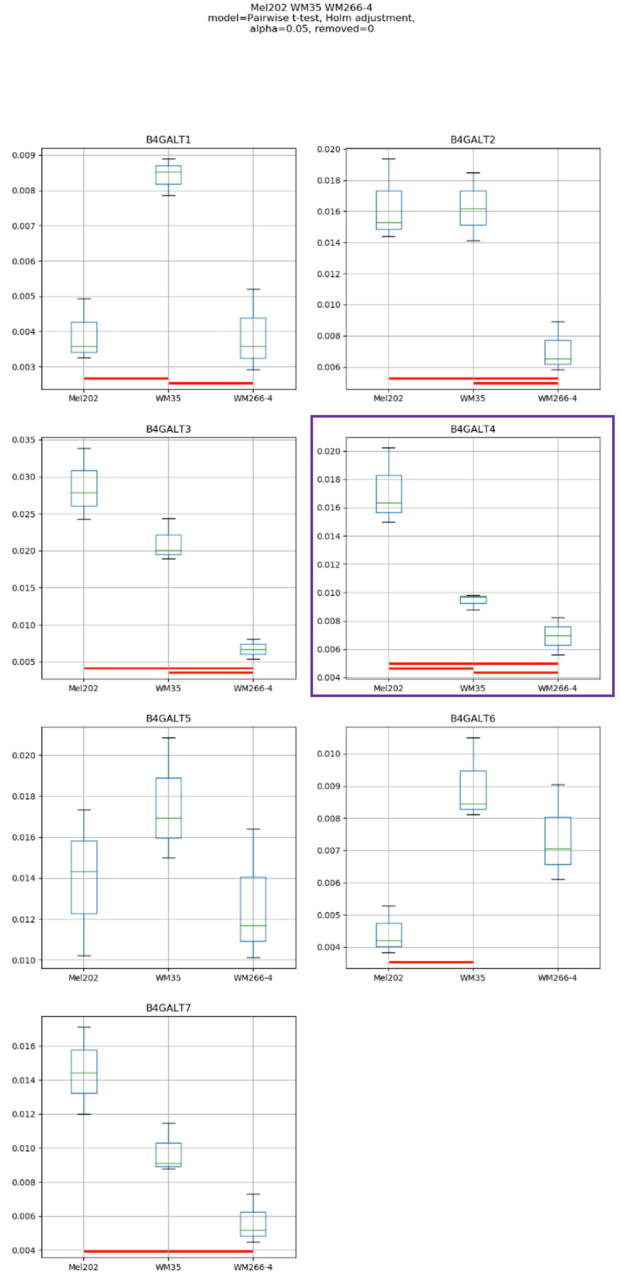
Mel202 WM35 WM266-4
model=Pairwise t-test, Holm adjustment,
alpha=0.05, removed=0
B4GALT1
B4GALT2
B4GALT3
B4GALT4
B4GALT5
B4GALT6
B4GALT7
Mel202
WM35
WM266-4

### Model No. 19

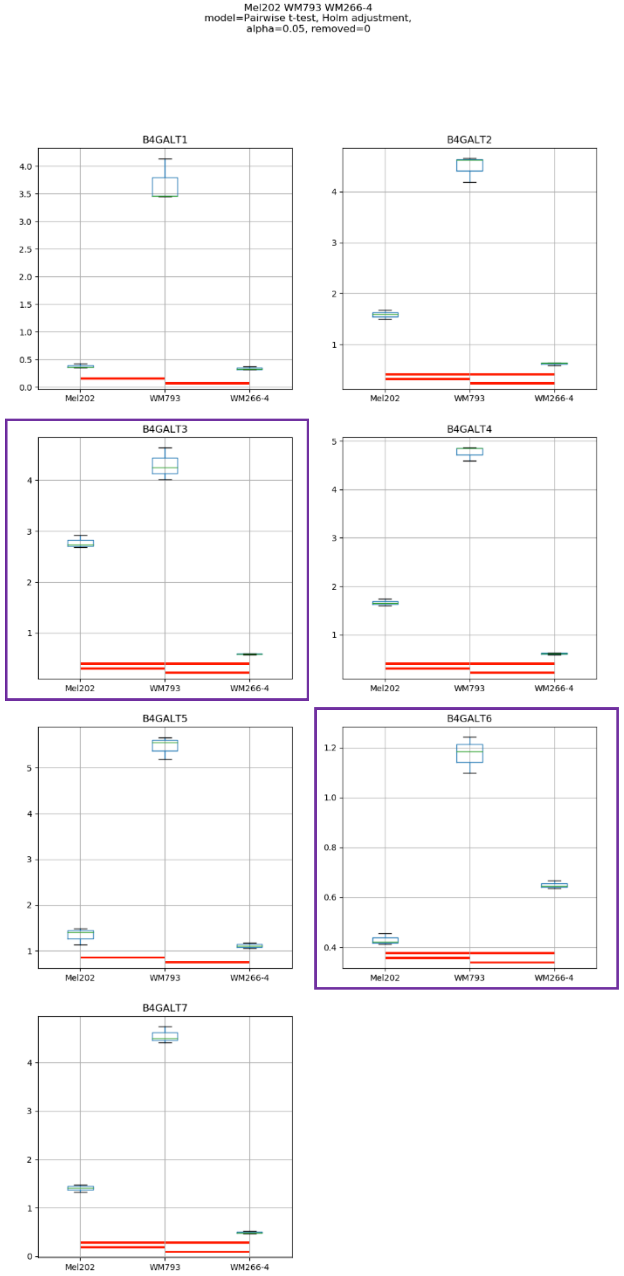

Mel202 WM793 WM266-4
model=Pairwise t-test, Holm adjustment,
alpha=0.05, removed=0
B4GALT1
B4GALT2
B4GALT3
B4GALT4
B4GALT5
B4GALT6
B4GALT7
Mel202
WM793
WM266-4

### Model No. 20

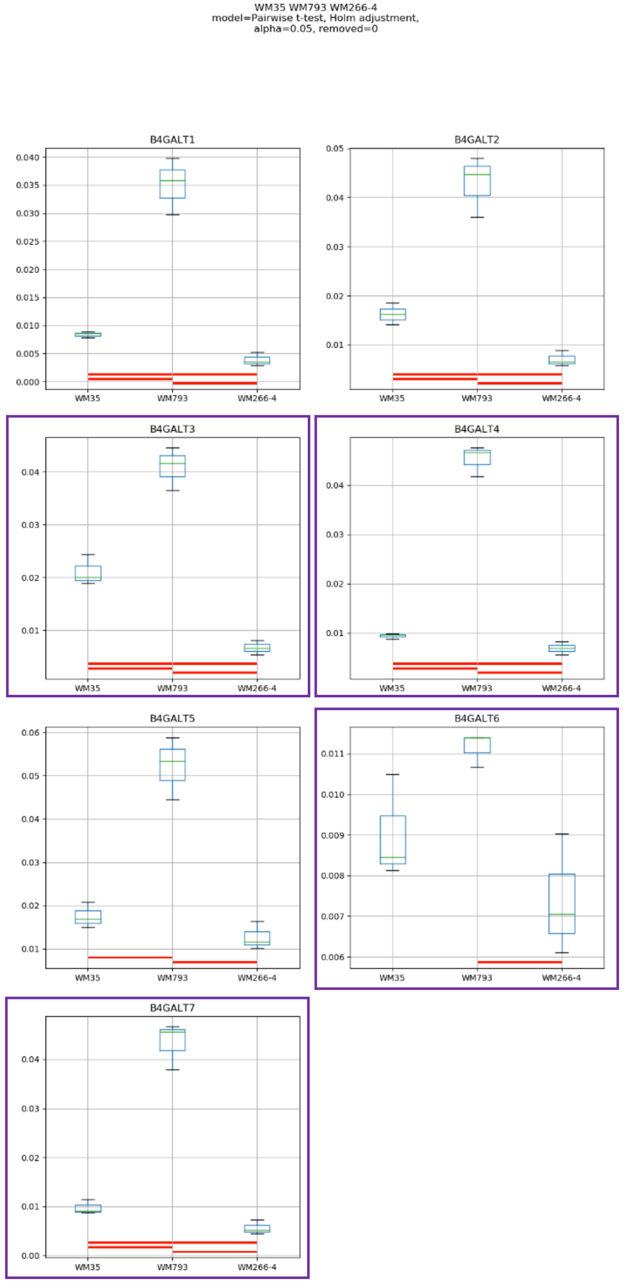
WM35 WM793 WM266-4
model=Pairwise t-test, Holm adjustment,
alpha=0.05, removed=0
B4GALT1
B4GALT2
B4GALT3
B4GALT4
B4GALT5
B4GALT6
B4GALT7
WM35
WM793
WM266-4

### Model No. 21

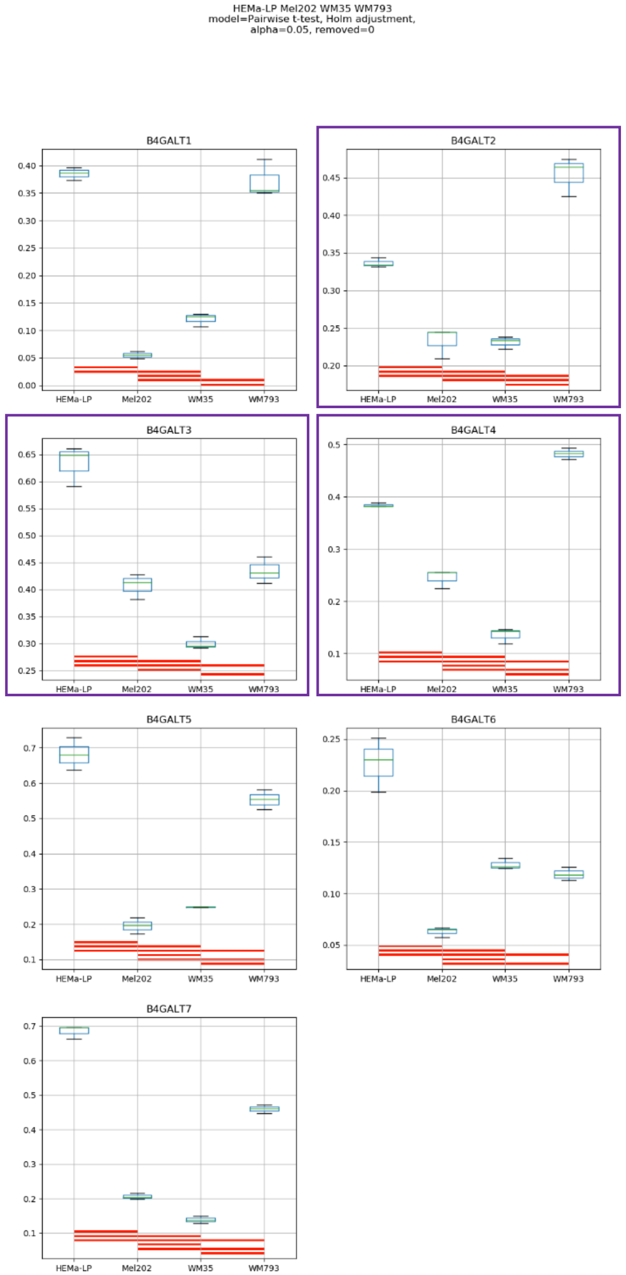
HEMa-LP Mel202 WM35 WM793
model=Pairwise t-test, Holm adjustment,
alpha=0.05, removed=0
B4GALT1
B4GALT2
B4GALT3
B4GALT4
B4GALT5
B4GALT6
B4GALT7
HEMa-LP
Mel202
WM35
WM793

### Model No. 22

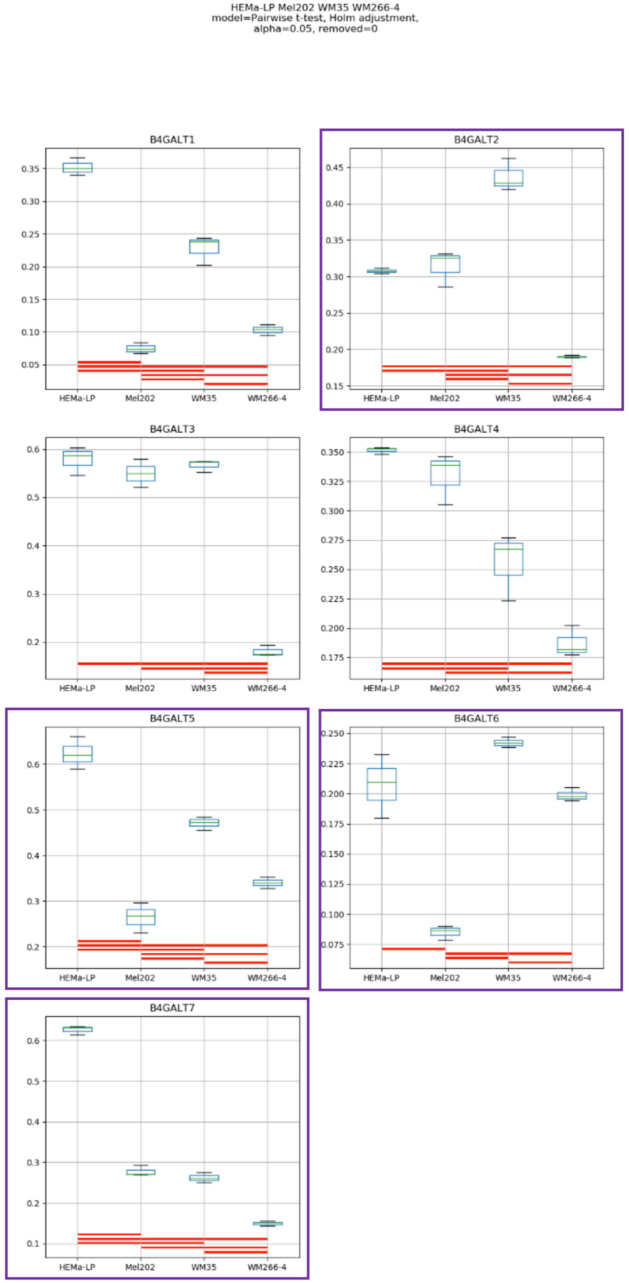
HEMa-LP Mel202 WM35 WM266-4
model=Pairwise t-test, Holm adjustment,
alpha=0.05, removed=0
B4GALT1
B4GALT2
B4GALT3
B4GALT4
B4GALT5
B4GALT6
B4GALT7
HEMa-LP
Mel202
WM35
WM266-4

### Model No. 23

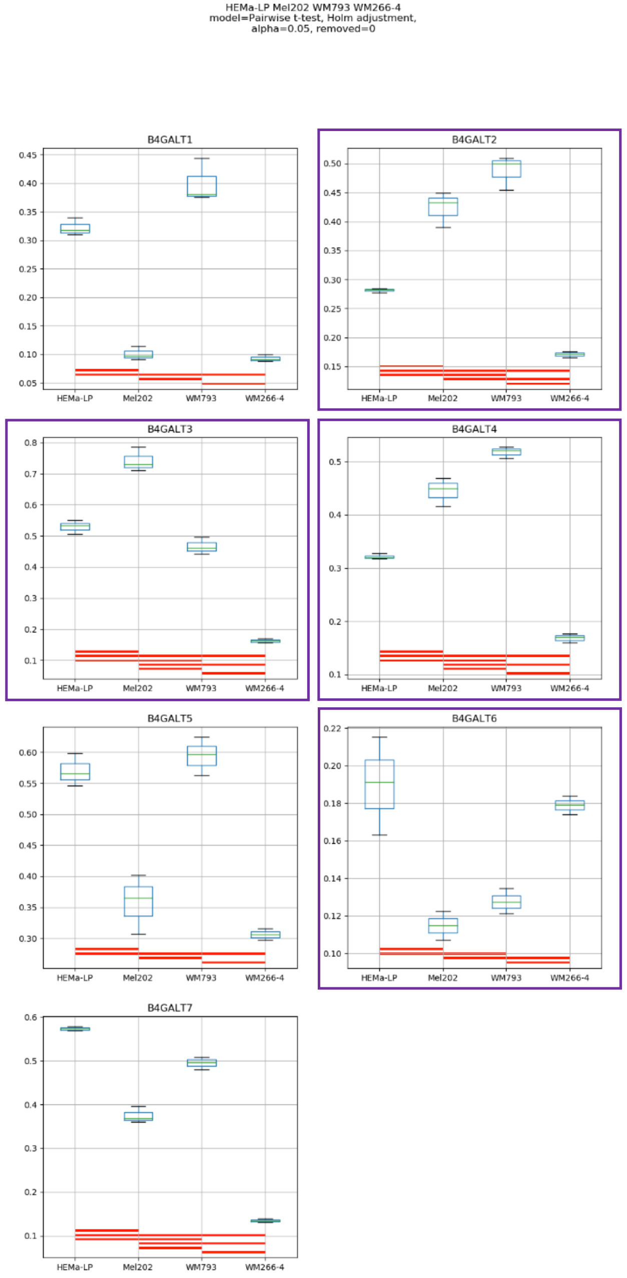
HEMa-LP Mel202 WM793 WM266-4
model=Pairwise t-test, Holm adjustment,
alpha=0.05, removed=0
B4GALT1
B4GALT2
B4GALT3
B4GALT4
B4GALT5
B4GALT6
B4GALT7
HEMa-LP
Mel202
WM793
WM266-4

### Model No. 24

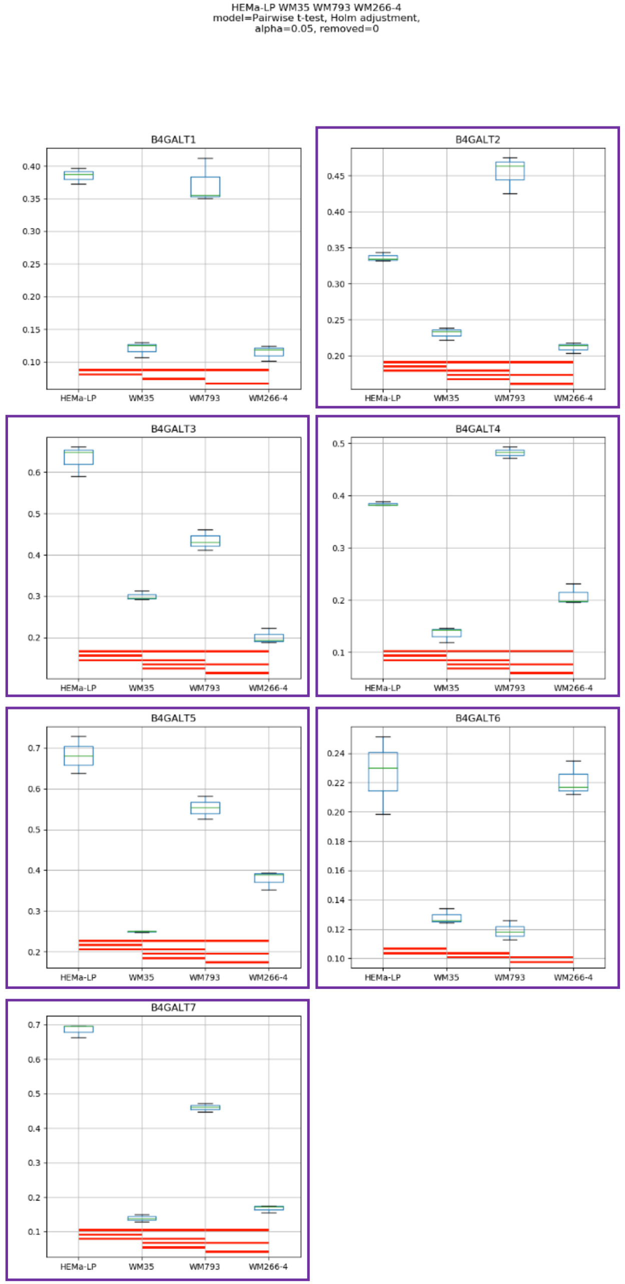
HEMa-LP WM35 WM793 WM266-4
model=Pairwise t-test, Holm adjustment,
alpha=0.05, removed=0
B4GALT1
B4GALT2
B4GALT3
B4GALT4
B4GALT5
B4GALT6
B4GALT7
HEMa-LP
WM35
WM793
WM266-4

Model No. 25

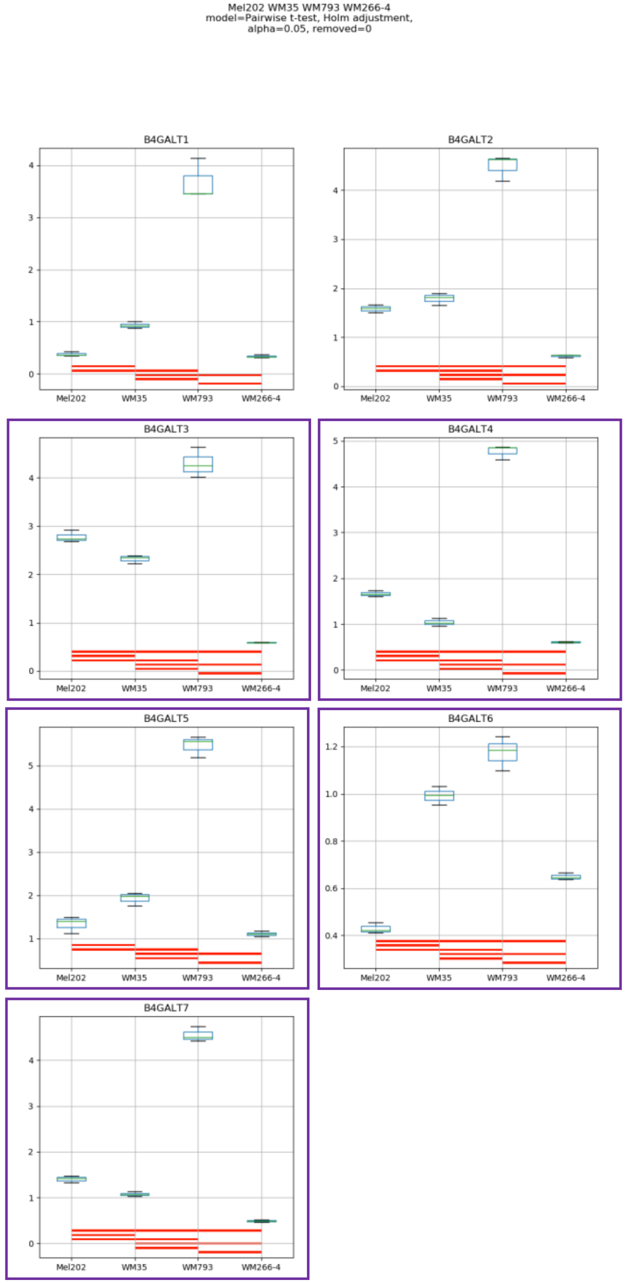
Mel202 WM35 WM793 WM266-4
model=Pairwise t-test, Holm adjustment,
alpha=0.05, removed=0
B4GALT1
B4GALT2
B4GALT3
B4GALT4
B4GALT5
B4GALT6
B4GALT7
Mel202
WM35
WM793
WM266-4

Model No. 26

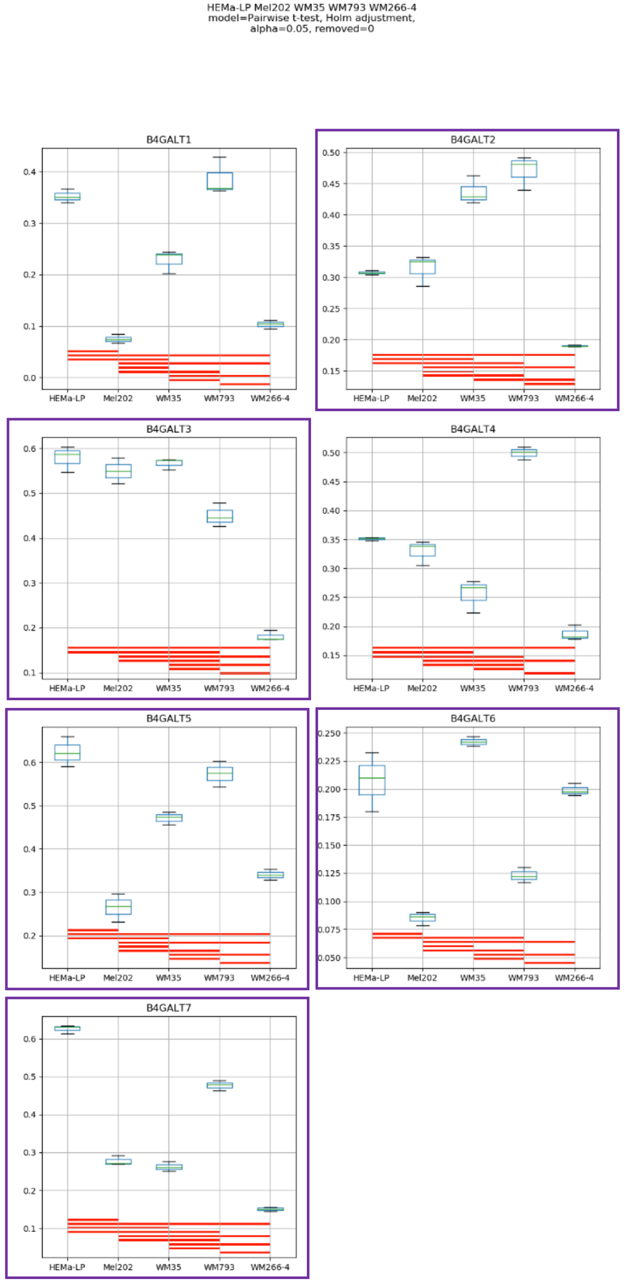
HEMa-LP Mel202 WM35 WM793 WM266-4
model=Pairwise t-test, Holm adjustment,
alpha=0.05, removed=0
B4GALT1
B4GALT2
B4GALT3
B4GALT4
B4GALT5
B4GALT6
B4GALT7
HEMa-LP
Mel202
WM35
WM793
WM266-4

**Supplementary Figure S15:** Medians of relative quantity (RQ) for target genes *B4GALT1–B4GALT7* normalized to the reference gene or pair of reference genes with the best stability values resulting from GenExpA analysis based on set 4 of four candidate reference genes (*HPRT1*, *GUSB*, *PGK1*, *RPS23*) and remove repetition level 0. The statistical analyses used the pairwise t-test with Holm adjustment. Red line represents statistical significance, $P < 0.05$. Frames denote box-plots representing the medians of the RQ values for a given target gene without a repeatable pattern in the same cell lines under different models (the 'Remove repetitions' box set to 0). Detailed results are listed in Supplementary Table 14.

Model No. 1

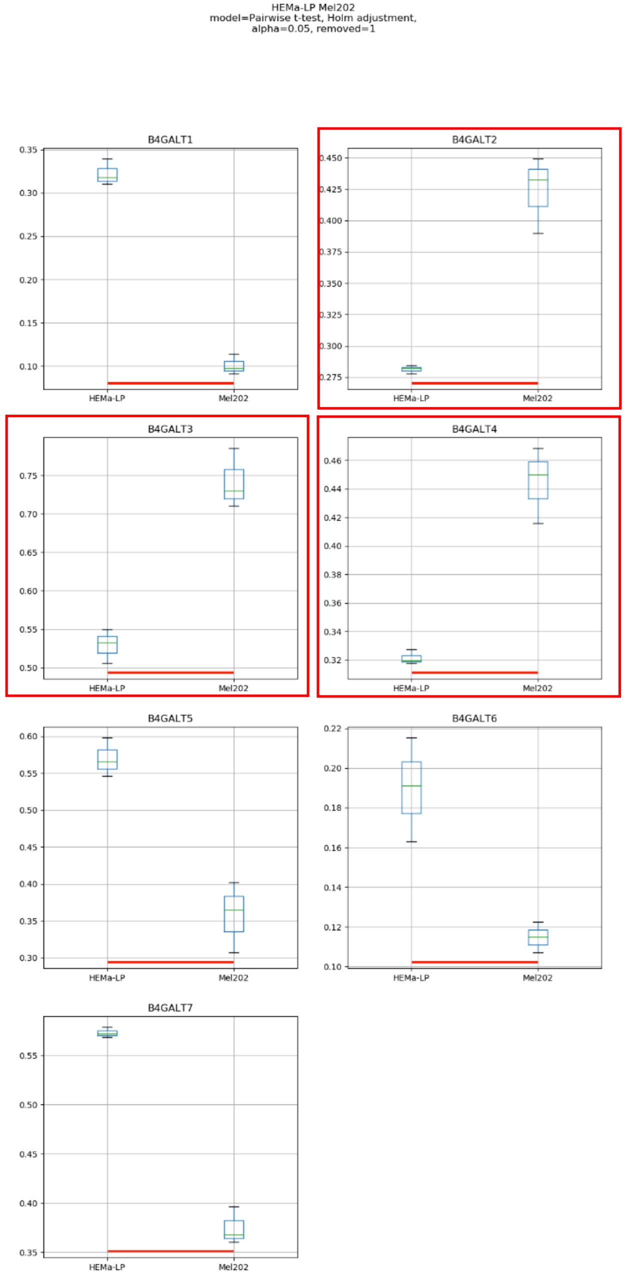
HEMa-LP Mel202
model=Pairwise t-test, Holm adjustment,
alpha=0.05, removed=1
B4GALT1
B4GALT2
B4GALT3
B4GALT4
B4GALT5
B4GALT6
B4GALT7
HEMa-LP
Mel202

### Model No. 2

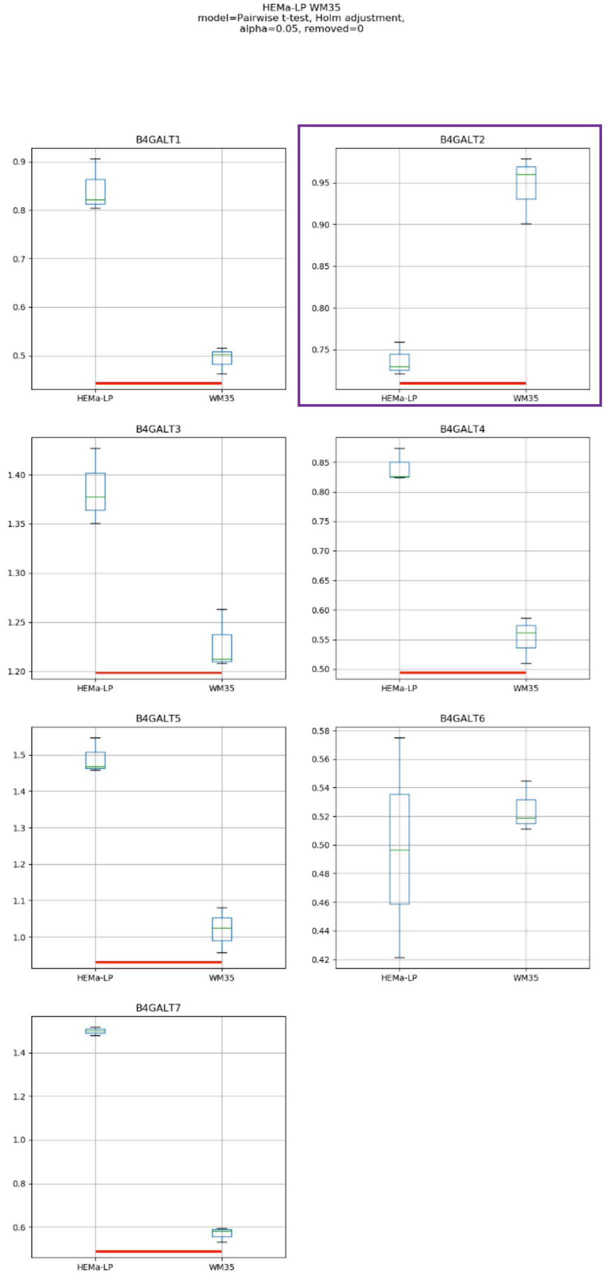
HEMa-LP WM35
model=Pairwise t-test, Holm adjustment,
alpha=0.05, removed=0
B4GALT1
B4GALT2
B4GALT3
B4GALT4
B4GALT5
B4GALT6
B4GALT7
HEMa-LP
WM35

### Model No. 3

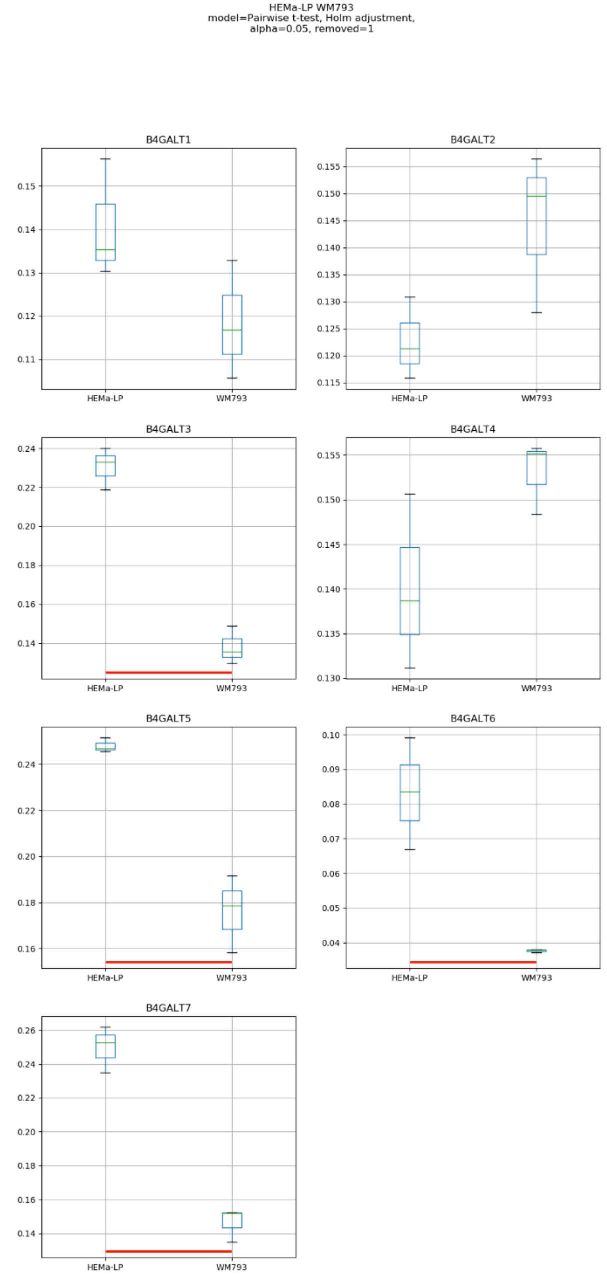
HEMa-LP WM793
model=Pairwise t-test, Holm adjustment,
alpha=0.05, removed=1
B4GALT1
B4GALT2
B4GALT3
B4GALT4
B4GALT5
B4GALT6
B4GALT7
HEMa-LP
WM793

### Model No. 4

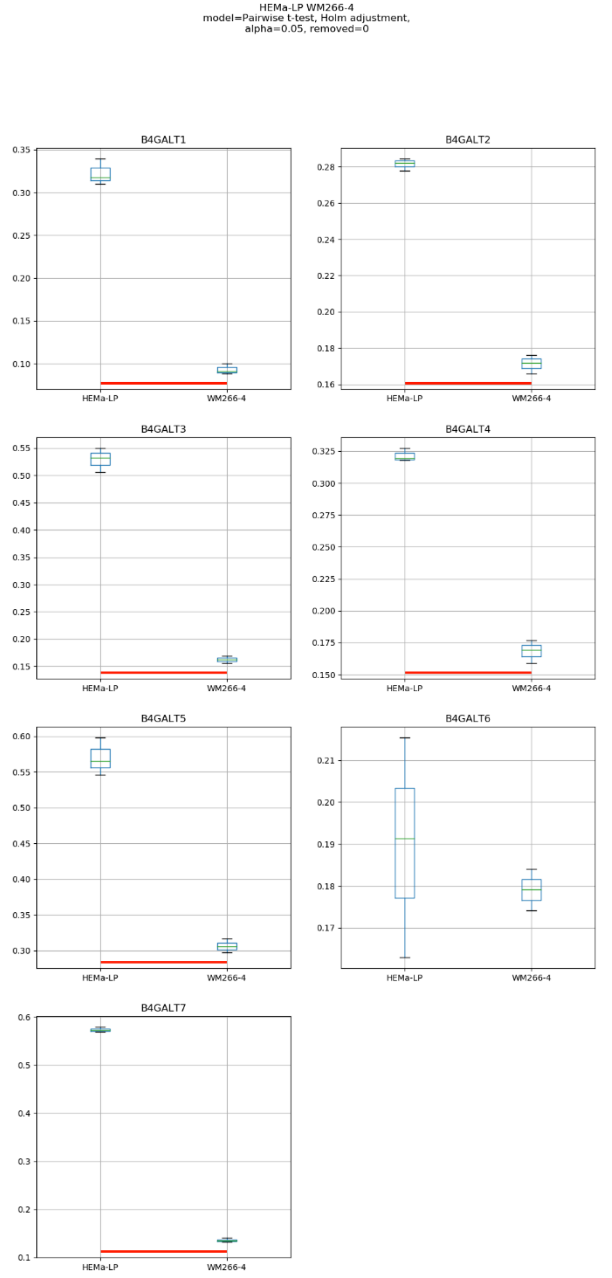
HEMa-LP WM266-4
model=Pairwise t-test, Holm adjustment,
alpha=0.05, removed=0
B4GALT1
B4GALT2
B4GALT3
B4GALT4
B4GALT5
B4GALT6
B4GALT7
HEMa-LP
WM266-4

### Model No. 5

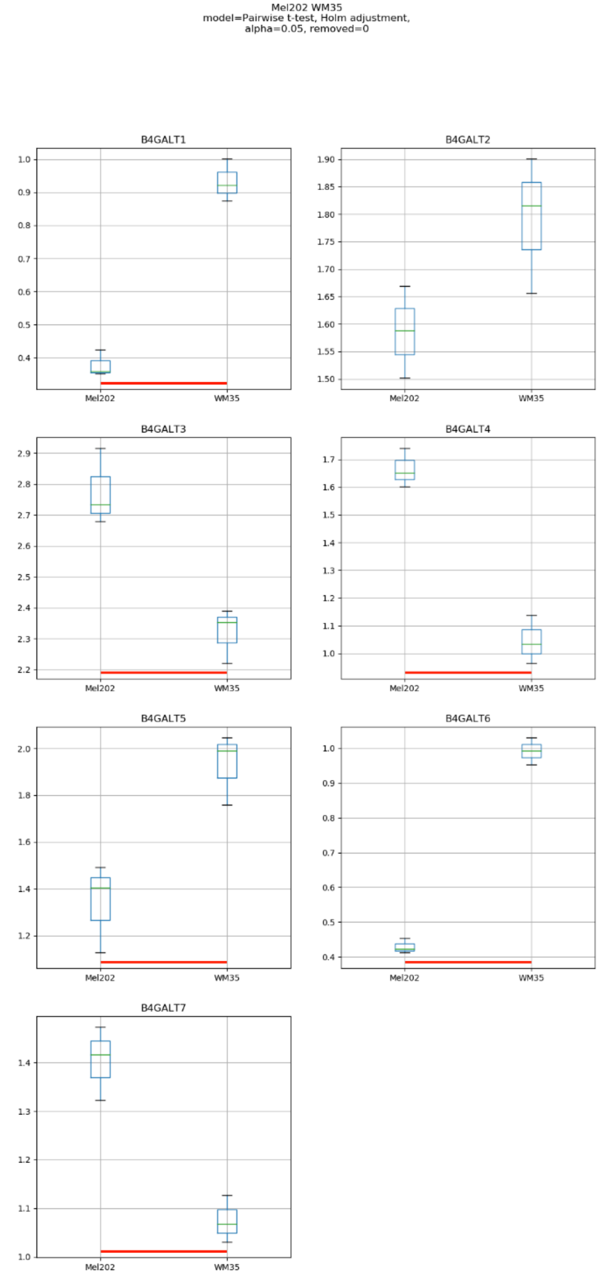
Mel202 WM35
model=Pairwise t-test, Holm adjustment,
alpha=0.05, removed=0
B4GALT1
B4GALT2
B4GALT3
B4GALT4
B4GALT5
B4GALT6
B4GALT7
Mel202
WM35

### Model No. 6

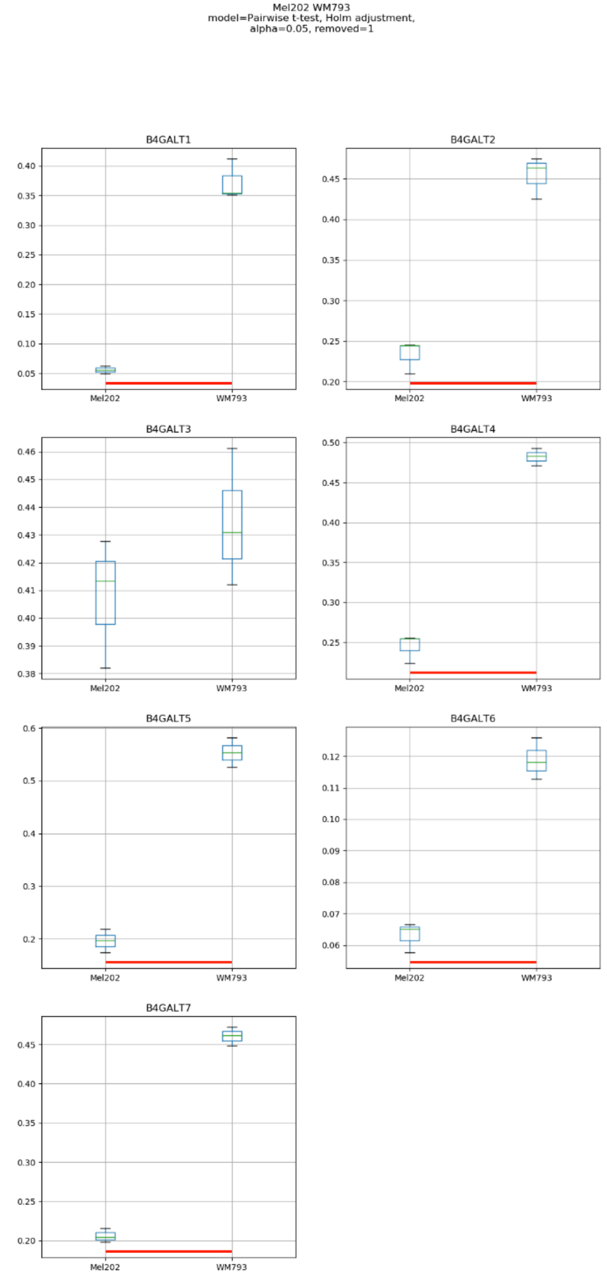
Mel202 WM793
model=Pairwise t-test, Holm adjustment,
alpha=0.05, removed=1
B4GALT1
B4GALT2
B4GALT3
B4GALT4
B4GALT5
B4GALT6
B4GALT7
Mel202
WM793

### Model No. 7

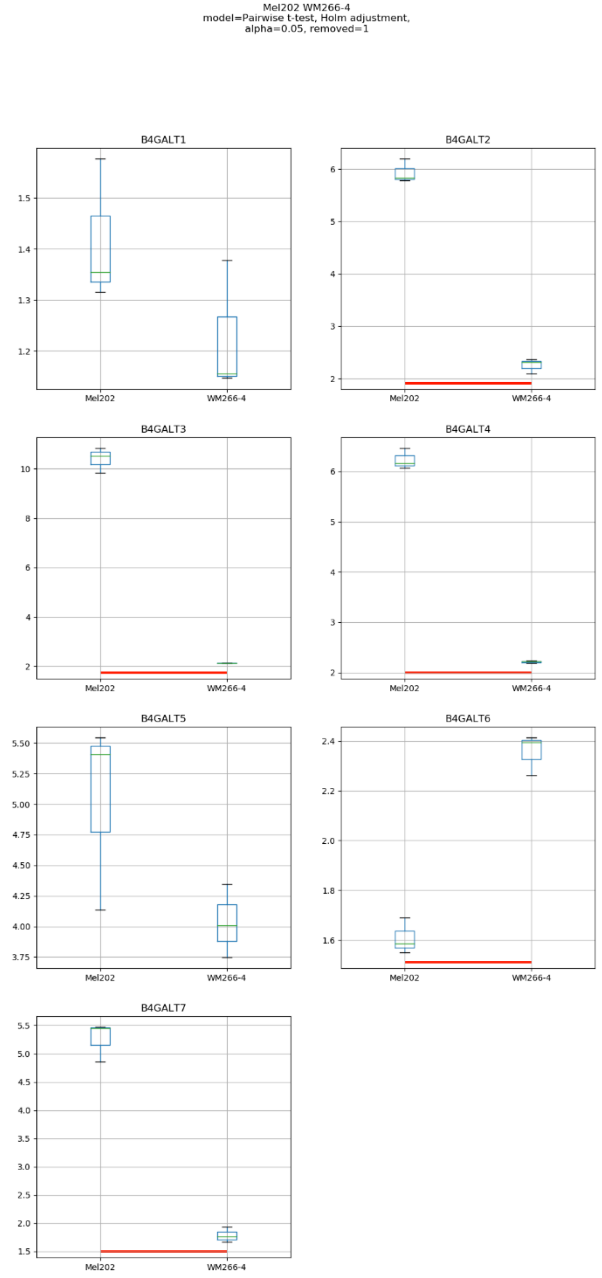
Mel202 WM266-4
model=Pairwise t-test, Holm adjustment,
alpha=0.05, removed=1
B4GALT1
B4GALT2
B4GALT3
B4GALT4
B4GALT5
B4GALT6
B4GALT7
Mel202
WM266-4

### Model No. 8

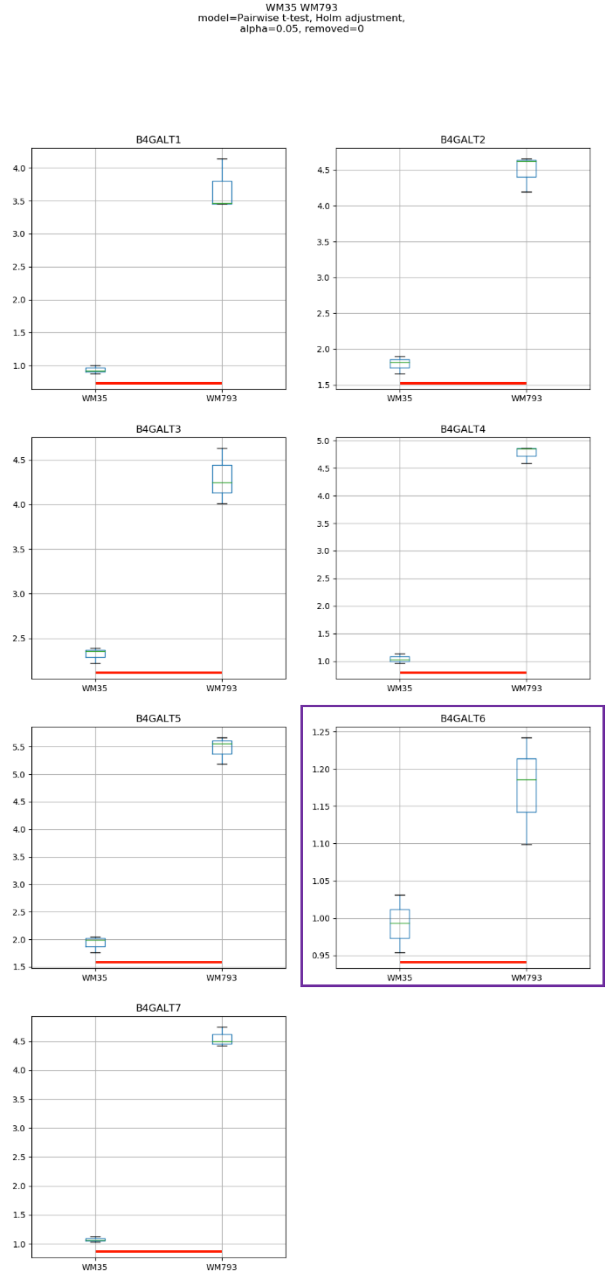
WM35 WM793
model=Pairwise t-test, Holm adjustment,
alpha=0.05, removed=0
B4GALT1
B4GALT2
B4GALT3
B4GALT4
B4GALT5
B4GALT6
B4GALT7
WM35
WM793

### Model No. 9

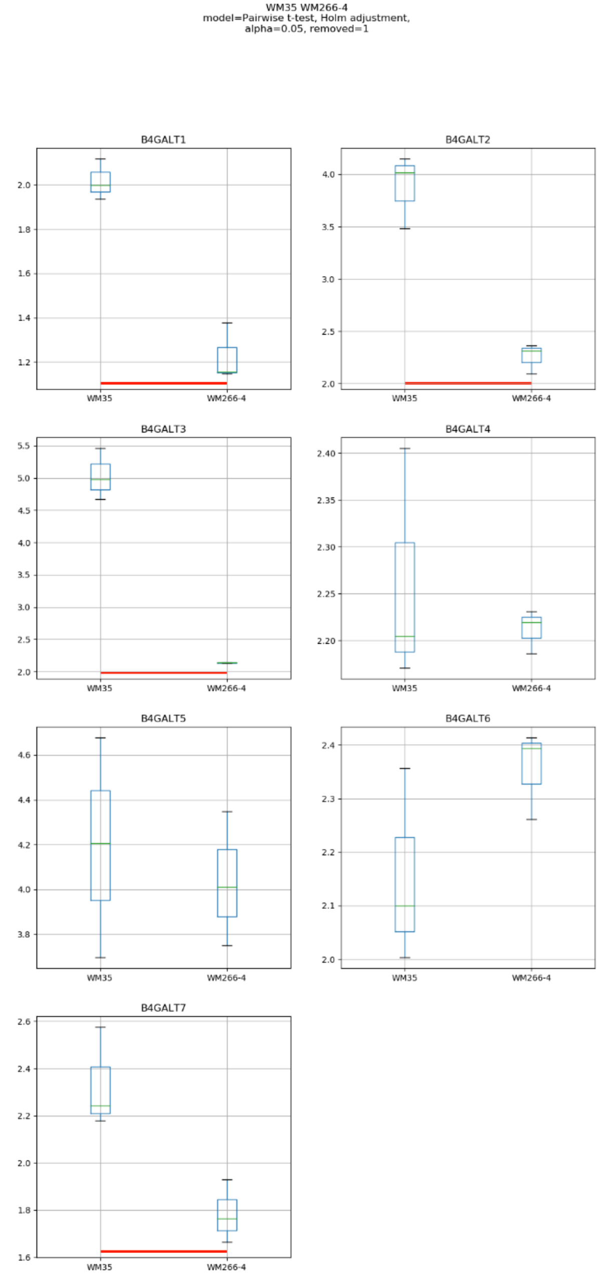

WM35 WM266-4
model=Pairwise t-test, Holm adjustment,
alpha=0.05, removed=1
B4GALT1
B4GALT2
B4GALT3
B4GALT4
B4GALT5
B4GALT6
B4GALT7
WM35
WM266-4

Model No. 10

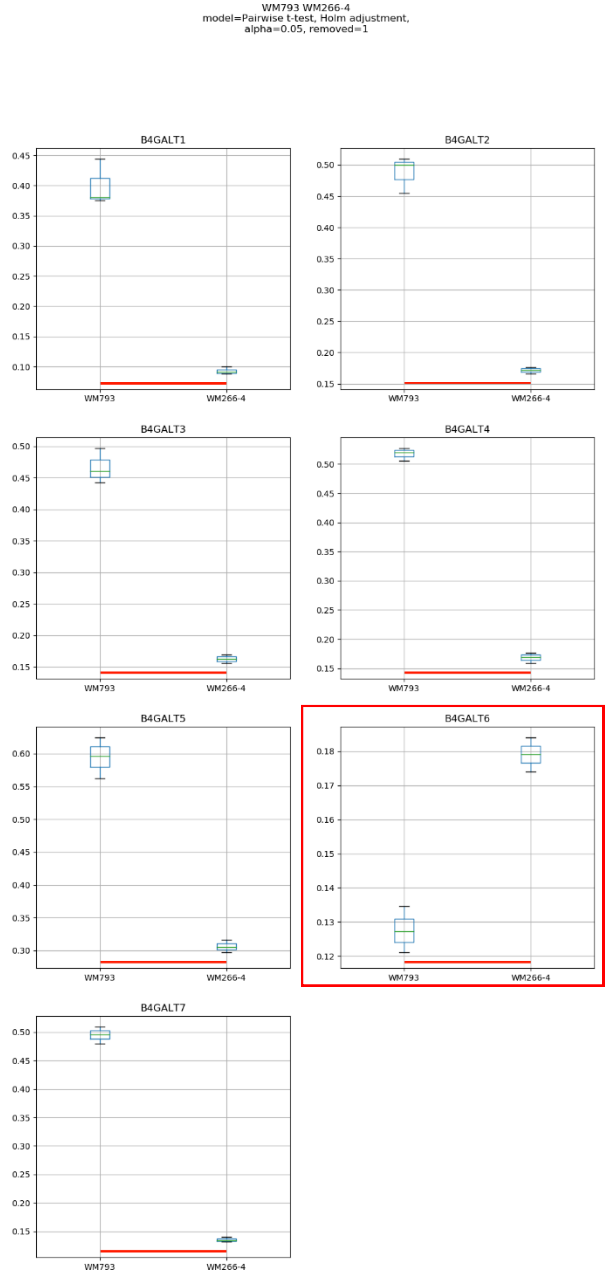

WM793 WM266-4
model=Pairwise t-test, Holm adjustment,
alpha=0.05, removed=1
B4GALT1
B4GALT2
B4GALT3
B4GALT4
B4GALT5
B4GALT6
B4GALT7
WM793
WM266-4

### Model No. 11

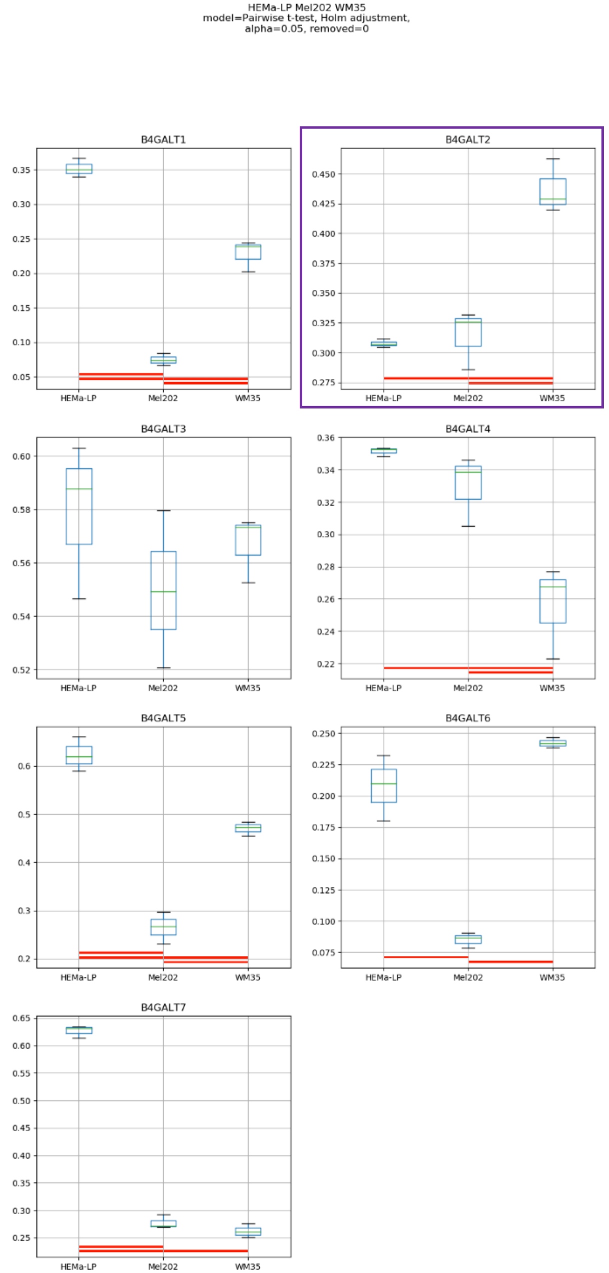
HEMa-LP Mel202 WM35
model=Pairwise t-test, Holm adjustment,
alpha=0.05, removed=0
B4GALT1
B4GALT2
B4GALT3
B4GALT4
B4GALT5
B4GALT6
B4GALT7
HEMa-LP
Mel202
WM35

### Model No. 12

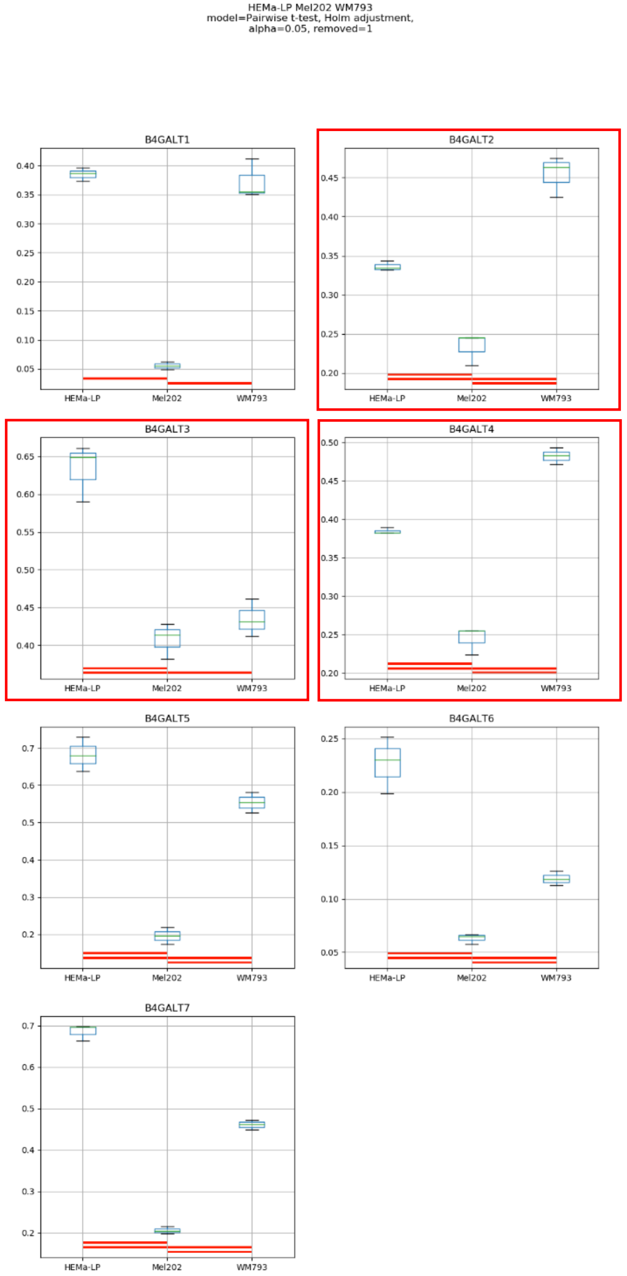
HEMa-LP Mel202 WM793
model=Pairwise t-test, Holm adjustment,
alpha=0.05, removed=1
B4GALT1
B4GALT2
B4GALT3
B4GALT4
B4GALT5
B4GALT6
B4GALT7
HEMa-LP
Mel202
WM793

Model No. 13

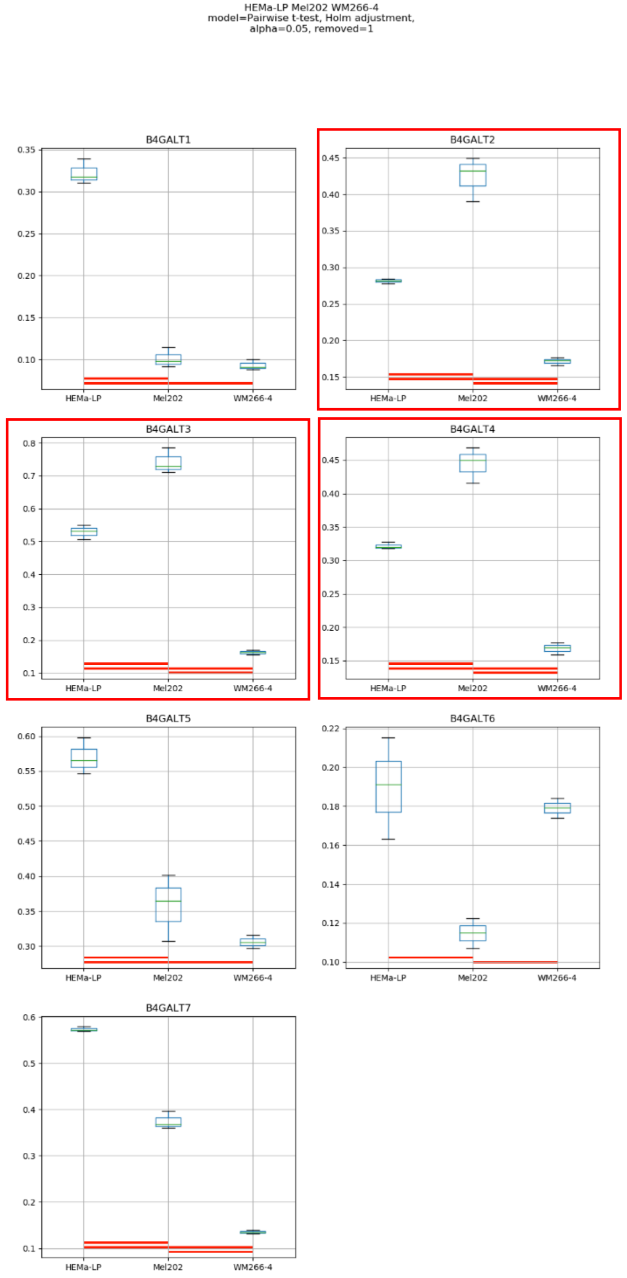
HEMa-LP Mel202 WM266-4
model=Pairwise t-test, Holm adjustment,
alpha=0.05, removed=1
B4GALT1
B4GALT2
B4GALT3
B4GALT4
B4GALT5
B4GALT6
B4GALT7
HEMa-LP
Mel202
WM266-4

### Model No. 14

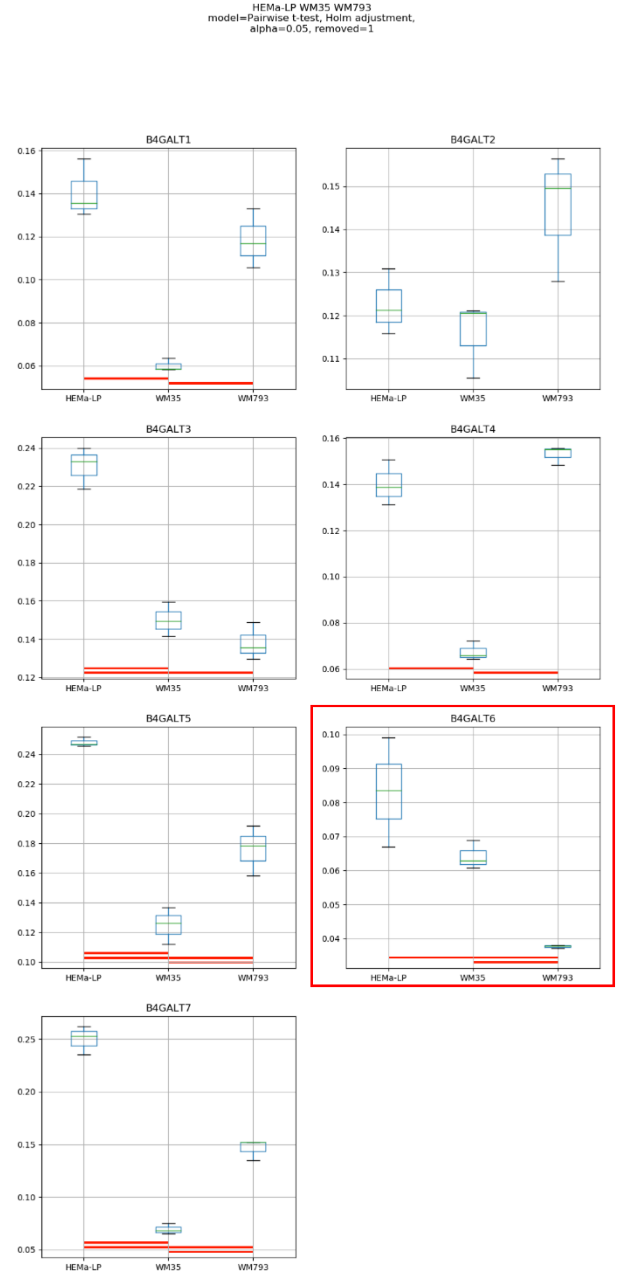
HEMa-LP WM35 WM793
model=Pairwise t-test, Holm adjustment,
alpha=0.05, removed=1
B4GALT1
B4GALT2
B4GALT3
B4GALT4
B4GALT5
B4GALT6
B4GALT7
HEMa-LP
WM35
WM793

Model No. 15

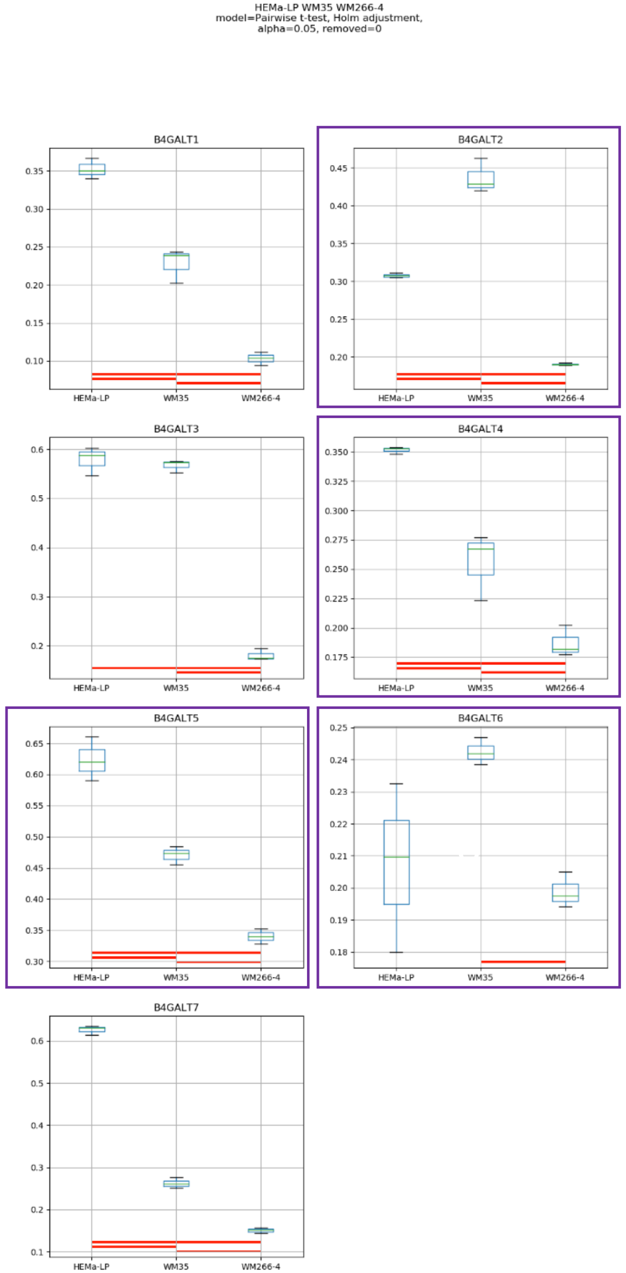
HEMa-LP WM35 WM266-4
model=Pairwise t-test, Holm adjustment,
alpha=0.05, removed=0
B4GALT1
B4GALT2
B4GALT3
B4GALT4
B4GALT5
B4GALT6
B4GALT7
HEMa-LP
WM35
WM266-4

Model No. 16

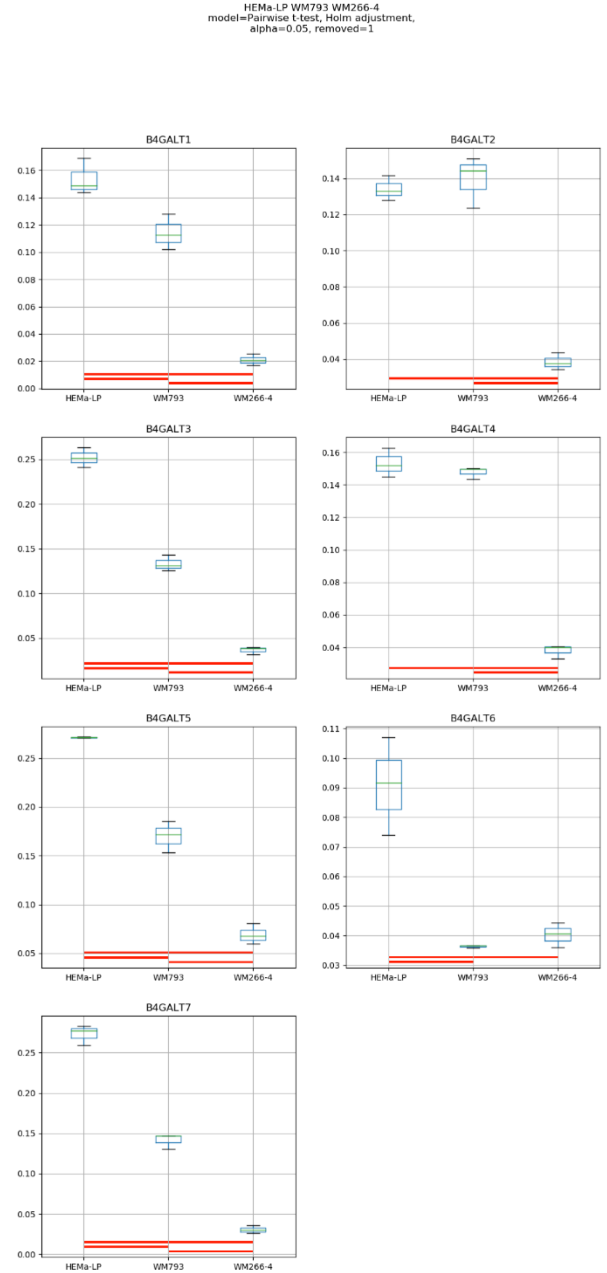
HEMa-LP WM793 WM266-4
model=Pairwise t-test, Holm adjustment,
alpha=0.05, removed=1
B4GALT1
B4GALT2
B4GALT3
B4GALT4
B4GALT5
B4GALT6
B4GALT7
HEMa-LP
WM793
WM266-4

### Model No. 17

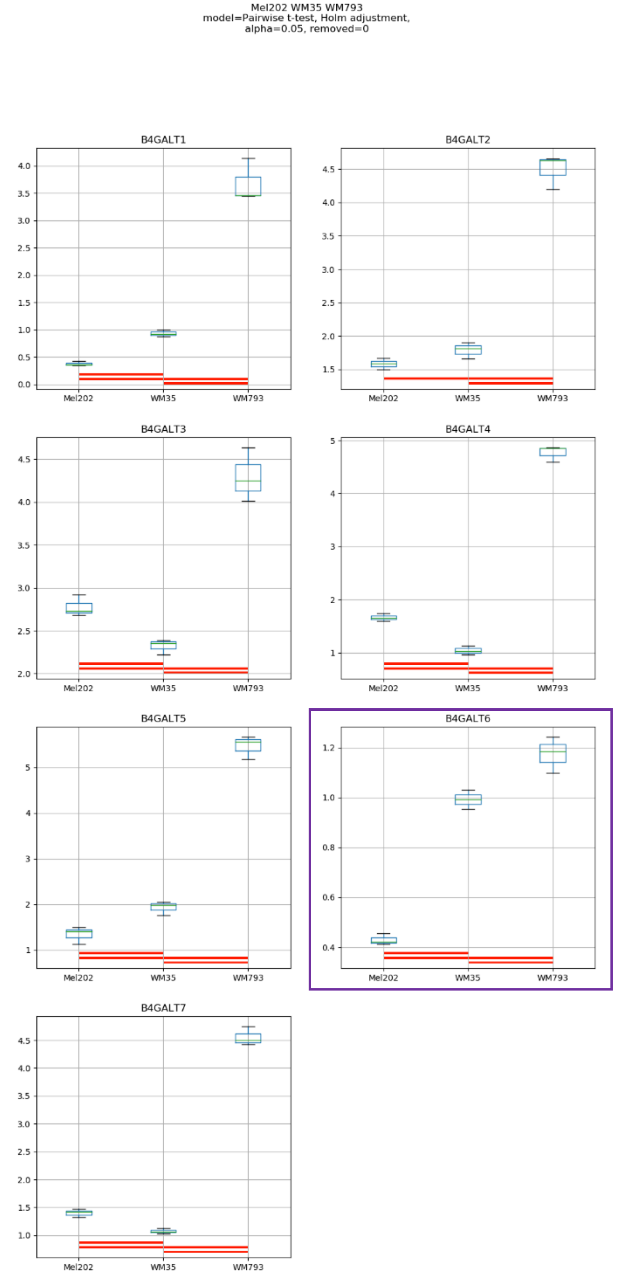
Mel202 WM35 WM793
model=Pairwise t-test, Holm adjustment,
alpha=0.05, removed=0
B4GALT1
B4GALT2
B4GALT3
B4GALT4
B4GALT5
B4GALT6
B4GALT7
Mel202
WM35
WM793

### Model No. 18

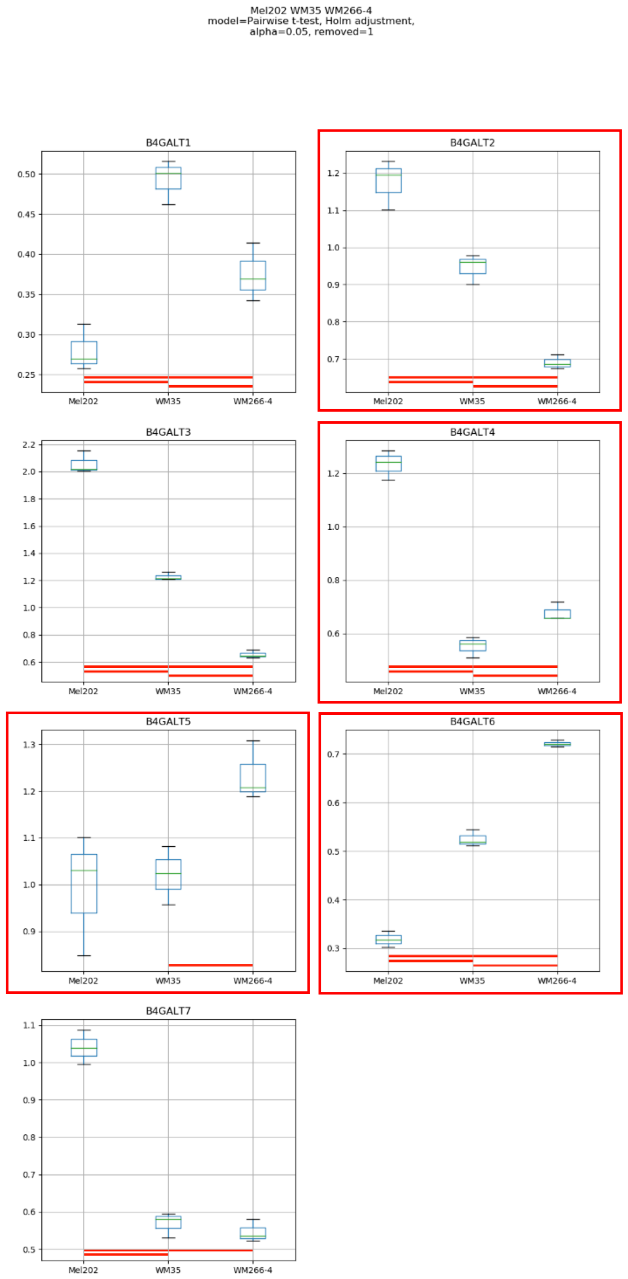

Mel202 WM35 WM266-4
model=Pairwise t-test, Holm adjustment,
alpha=0.05, removed=1
B4GALT1
B4GALT2
B4GALT3
B4GALT4
B4GALT5
B4GALT6
B4GALT7
Mel202
WM35
WM266-4

Model No. 19

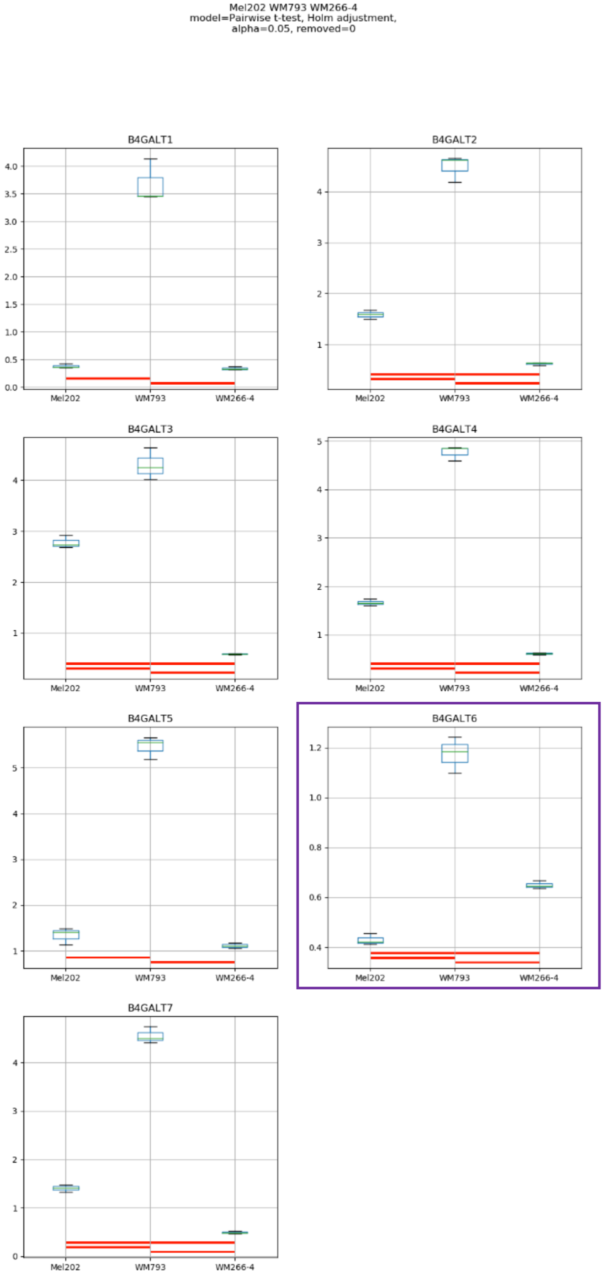
Mel202 WM793 WM266-4
model=Pairwise t-test, Holm adjustment,
alpha=0.05, removed=0
B4GALT1
B4GALT2
B4GALT3
B4GALT4
B4GALT5
B4GALT6
B4GALT7
Mel202
WM793
WM266-4

Model No. 20

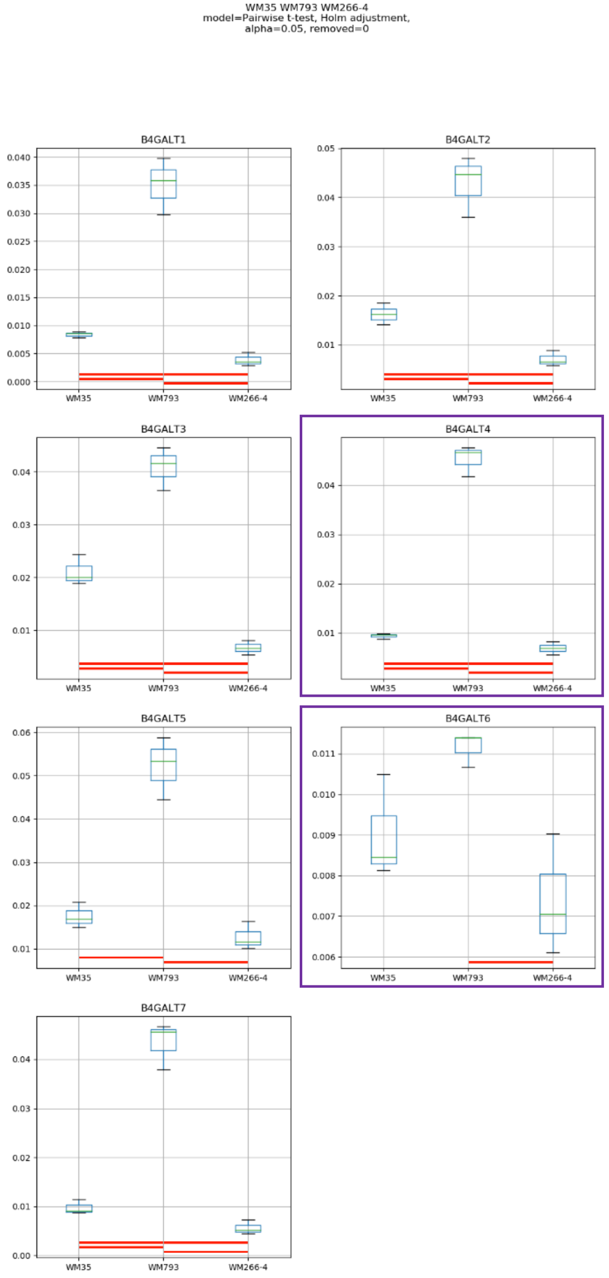
WM35 WM793 WM266-4
model=Pairwise t-test, Holm adjustment,
alpha=0.05, removed=0
B4GALT1
B4GALT2
B4GALT3
B4GALT4
B4GALT5
B4GALT6
B4GALT7
WM35
WM793
WM266-4

Model No. 21

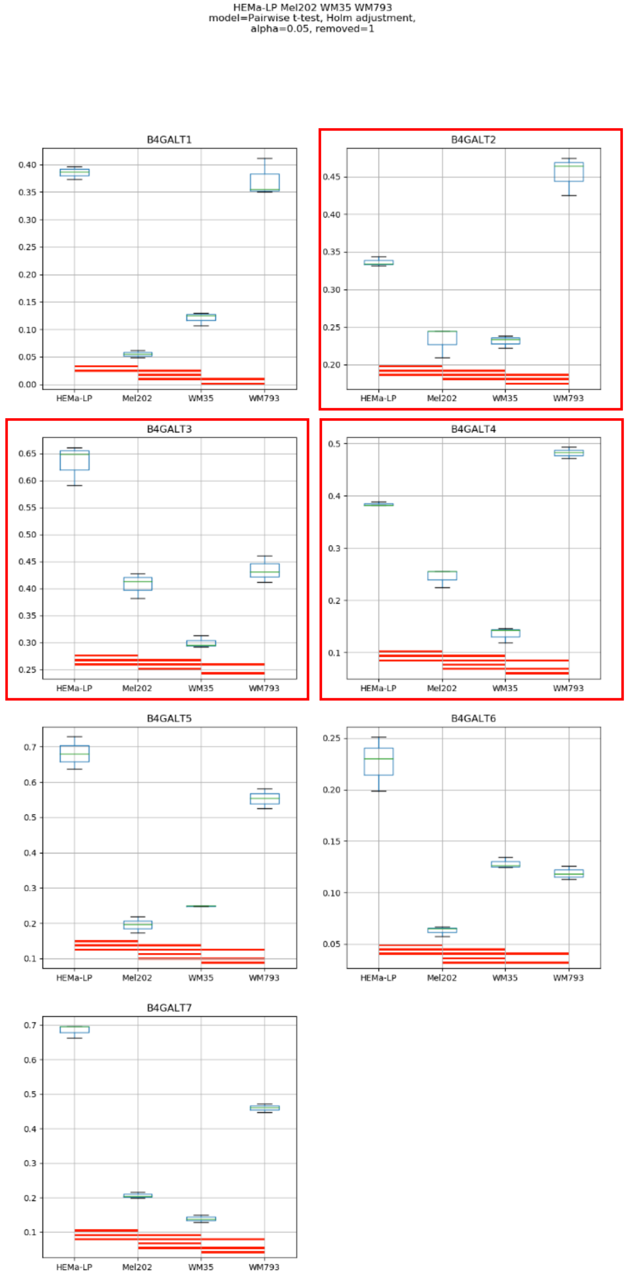

HEMa-LP Mel202 WM35 WM793
model=Pairwise t-test, Holm adjustment,
alpha=0.05, removed=1
B4GALT1
B4GALT2
B4GALT3
B4GALT4
B4GALT5
B4GALT6
B4GALT7
HEMa-LP
Mel202
WM35
WM793

### Model No. 22

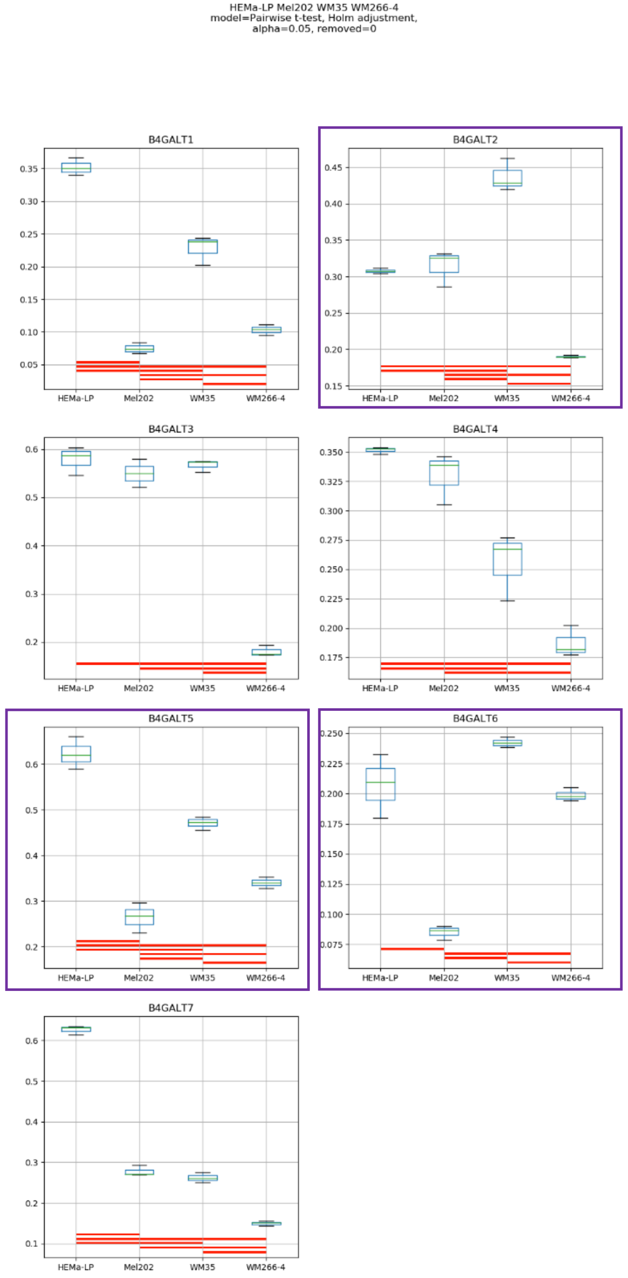
HEMa-LP Mel202 WM35 WM266-4
model=Pairwise t-test, Holm adjustment,
alpha=0.05, removed=0
B4GALT1
B4GALT2
B4GALT3
B4GALT4
B4GALT5
B4GALT6
B4GALT7
HEMa-LP
Mel202
WM35
WM266-4

Model No. 23

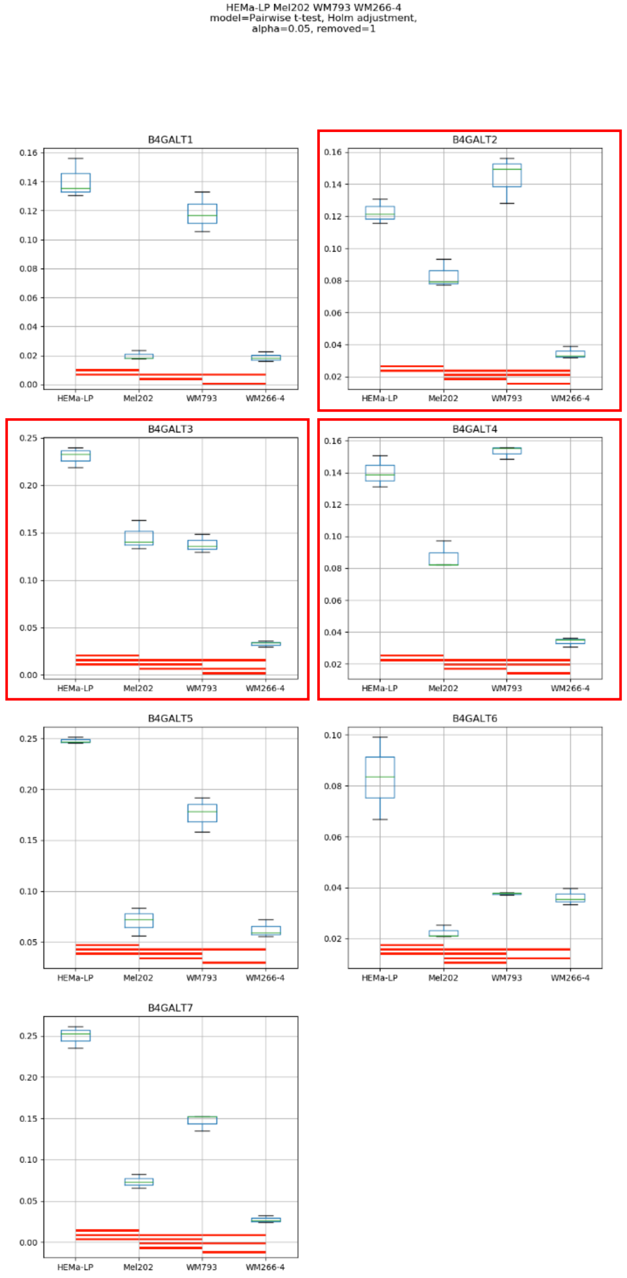

HEMa-LP Mel202 WM793 WM266-4
model=Pairwise t-test, Holm adjustment,
alpha=0.05, removed=1
B4GALT1
B4GALT2
B4GALT3
B4GALT4
B4GALT5
B4GALT6
B4GALT7
HEMa-LP
Mel202
WM793
WM266-4

### Model No. 24

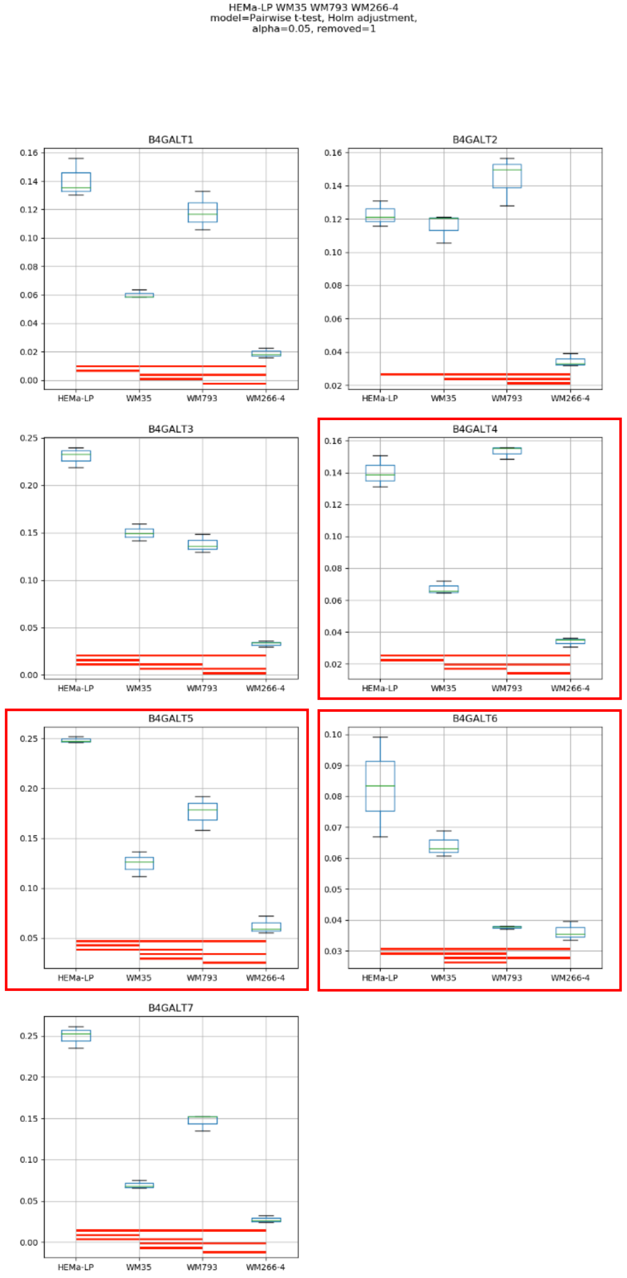
HEMa-LP WM35 WM793 WM266-4
model=Pairwise t-test, Holm adjustment,
alpha=0.05, removed=1
B4GALT1
B4GALT2
B4GALT3
B4GALT4
B4GALT5
B4GALT6
B4GALT7
HEMa-LP
WM35
WM793
WM266-4

### Model No. 25

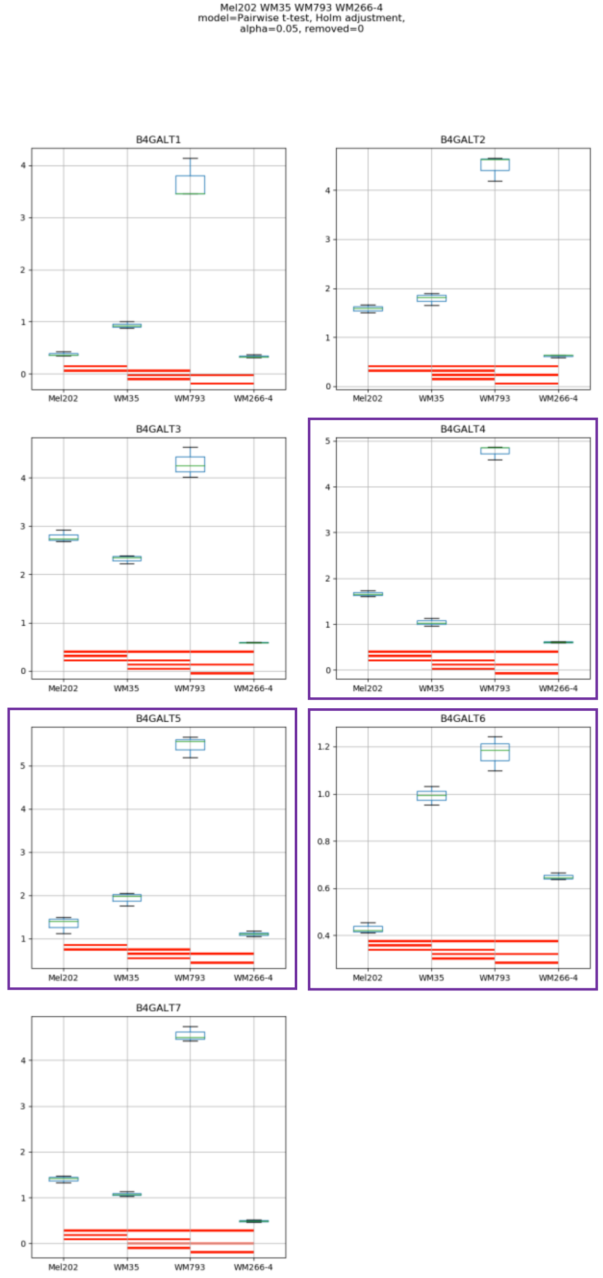
Mel202 WM35 WM793 WM266-4
model=Pairwise t-test, Holm adjustment,
alpha=0.05, removed=0
B4GALT1
B4GALT2
B4GALT3
B4GALT4
B4GALT5
B4GALT6
B4GALT7
Mel202
WM35
WM793
WM266-4

### Model No. 26

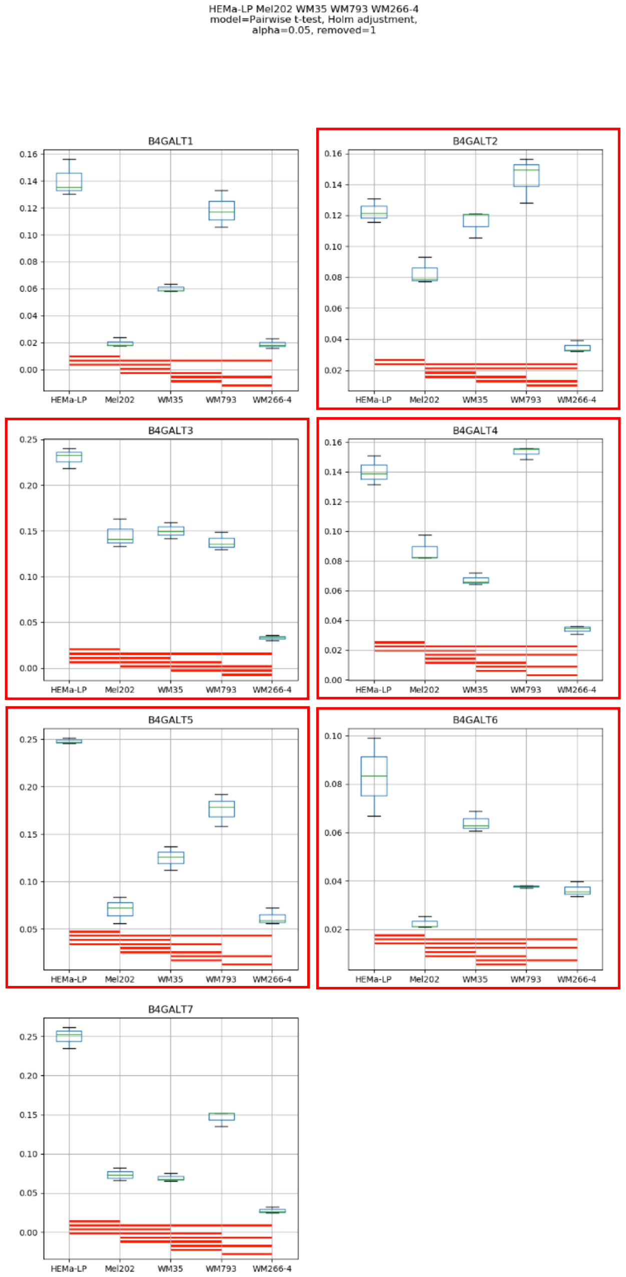

HEMa-LP Mel202 WM35 WM793 WM266-4
model=Pairwise t-test, Holm adjustment,
alpha=0.05, removed=1
B4GALT1
B4GALT2
B4GALT3
B4GALT4
B4GALT5
B4GALT6
B4GALT7
HEMa-LP
Mel202
WM35
WM793
WM266-4

**Supplementary Figure S16:** Medians of relative quantity (RQ) for target genes *B4GALT1–B4GALT7* normalized to the reference gene or pair of reference genes with the best stability values resulting from GenExpA analysis based on set 4 of four candidate reference genes (*HPRT1, GUSB, PGK1, RPS23*) and remove repetition level 1. The statistical analyses used the pairwise t-test with Holm adjustment. Red line represents statistical significance, $P < 0.05$. Frames denote box-plots representing the medians of the RQ values for a given target gene without a repeatable pattern in the same cell lines under different models (the 'Remove repetitions' box set to 0 – violet frame,  or to 1 – red frame). Detailed results are listed in Supplementary Table 15.

### Model No. 1

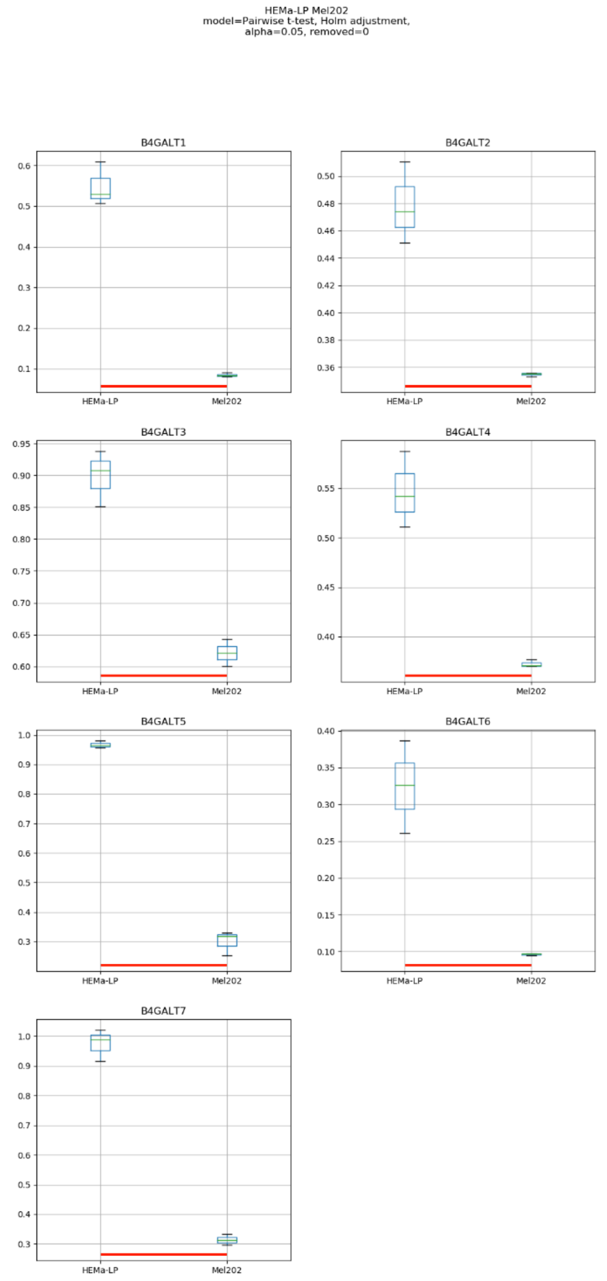
HEMa-LP Mel202
model=Pairwise t-test, Holm adjustment,
alpha=0.05, removed=0
B4GALT1
B4GALT2
B4GALT3
B4GALT4
B4GALT5
B4GALT6
B4GALT7
HEMa-LP
Mel202

Model No. 2

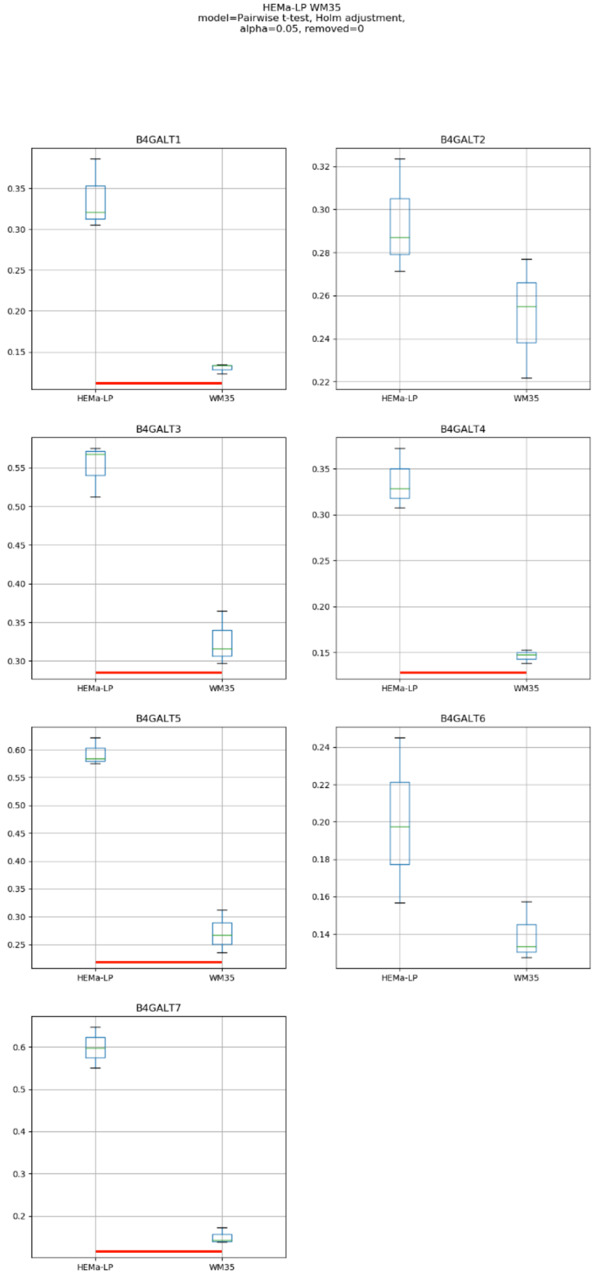

HEMa-LP WM35
model=Pairwise t-test, Holm adjustment,
alpha=0.05, removed=0
B4GALT1
B4GALT2
B4GALT3
B4GALT4
B4GALT5
B4GALT6
B4GALT7
HEMa-LP
WM35

### Model No. 3

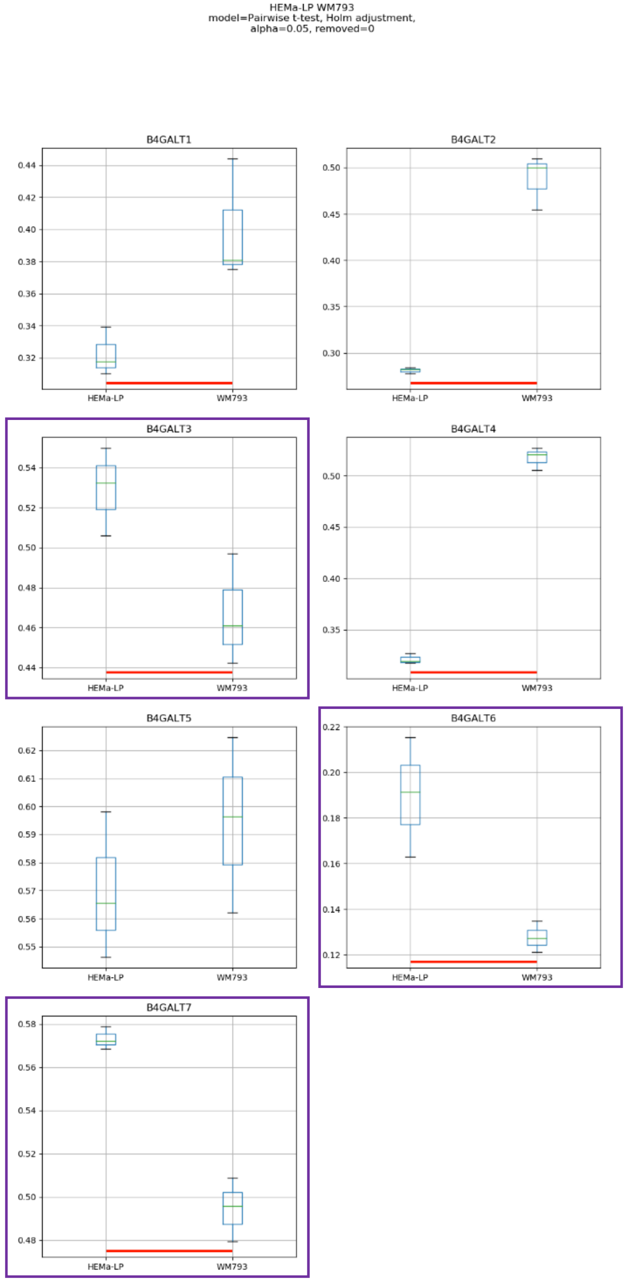
HEMa-LP WM793
model=Pairwise t-test, Holm adjustment,
alpha=0.05, removed=0
B4GALT1
B4GALT2
B4GALT3
B4GALT4
B4GALT5
B4GALT6
B4GALT7
HEMa-LP
WM793

### Model No. 4

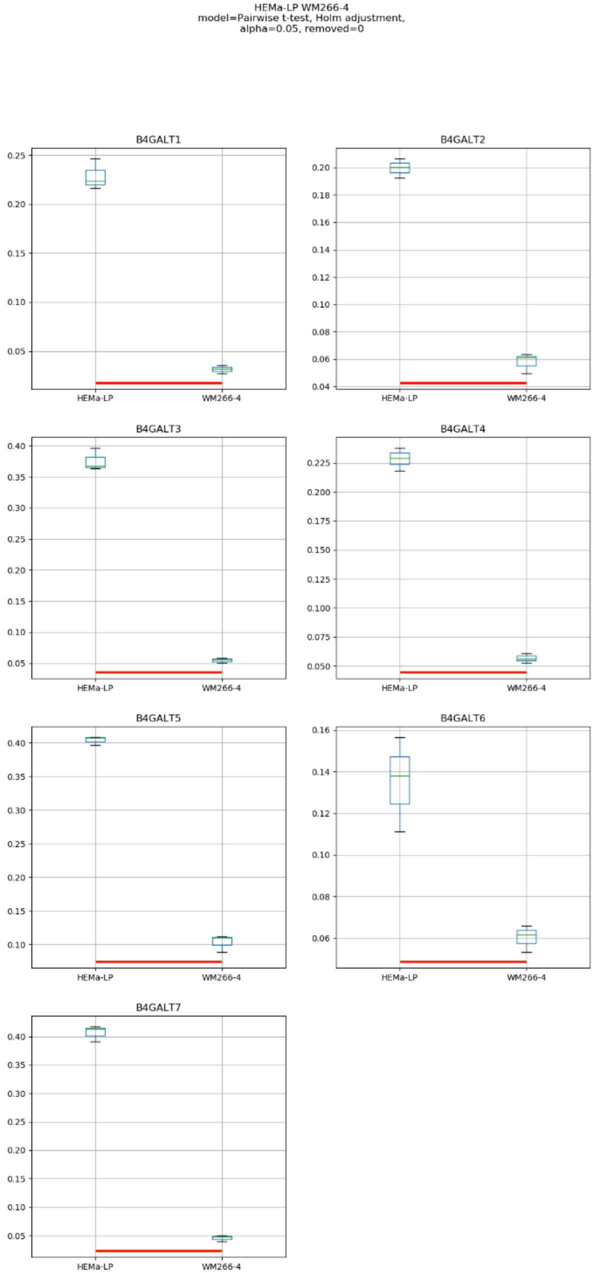

HEMa-LP WM266-4
model=Pairwise t-test, Holm adjustment,
alpha=0.05, removed=0
B4GALT1
B4GALT2
B4GALT3
B4GALT4
B4GALT5
B4GALT6
B4GALT7
HEMa-LP
WM266-4

### Model No. 5

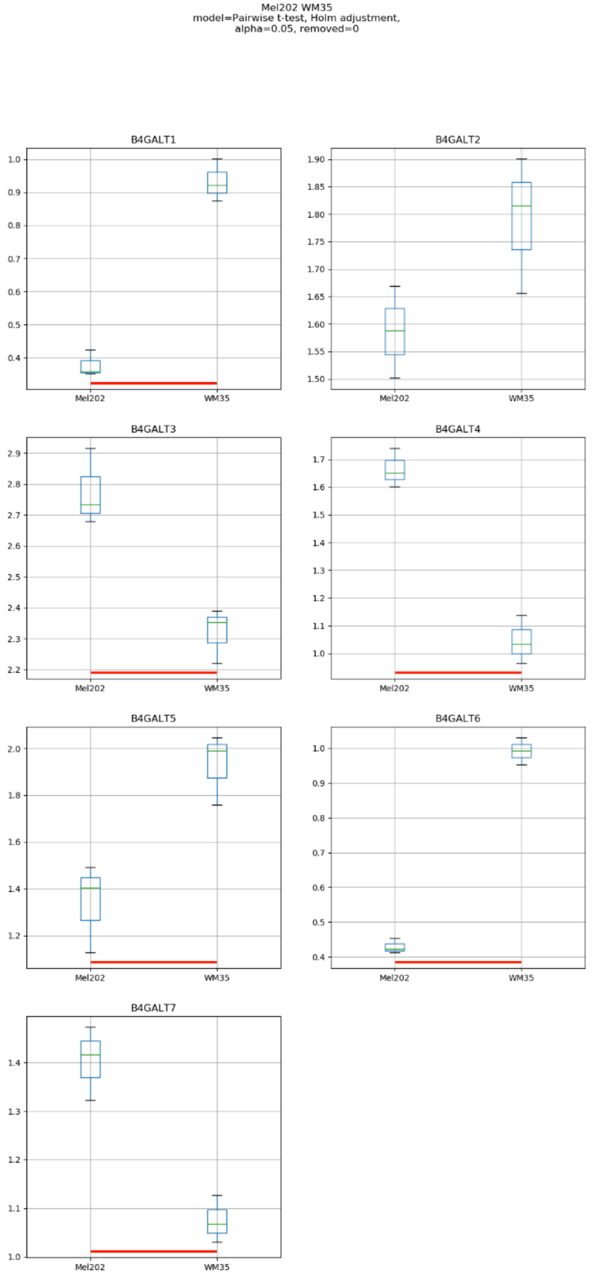
Mel202 WM35
model=Pairwise t-test, Holm adjustment,
alpha=0.05, removed=0
B4GALT1
B4GALT2
B4GALT3
B4GALT4
B4GALT5
B4GALT6
B4GALT7
Mel202
WM35

Model No. 6

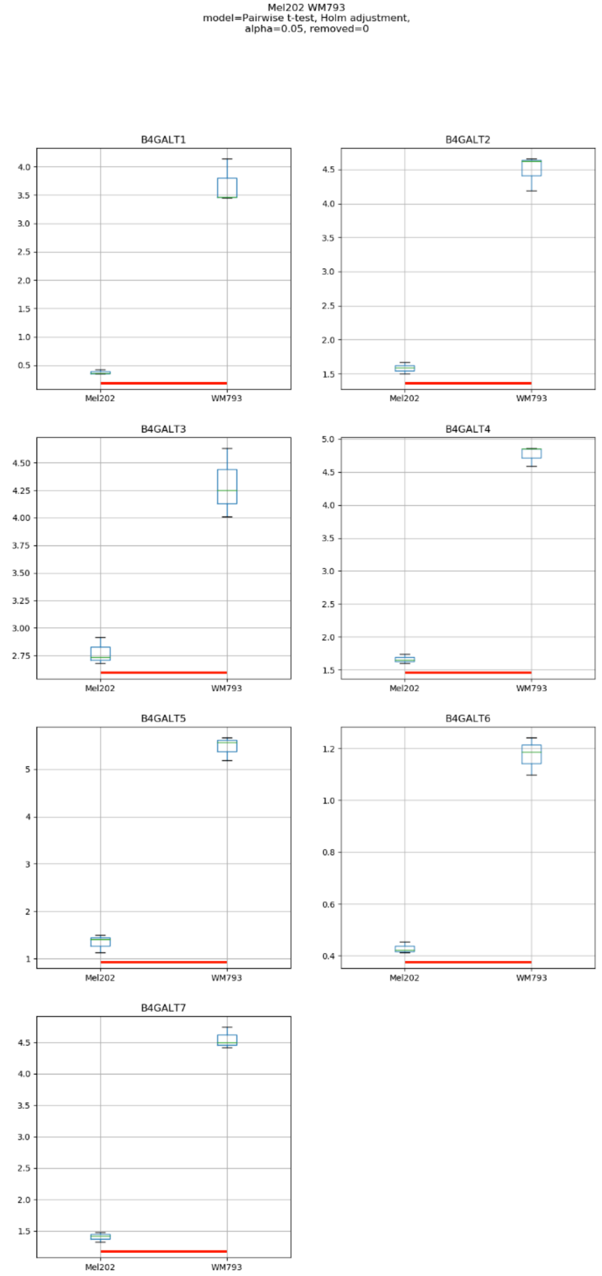
Mel202 WM793
model=Pairwise t-test, Holm adjustment,
alpha=0.05, removed=0
B4GALT1
B4GALT2
B4GALT3
B4GALT4
B4GALT5
B4GALT6
B4GALT7
Mel202
WM793

### Model No. 7

Mel202 WM266-4
model=Pairwise t-test, Holm adjustment,
alpha=0.05, removed=0

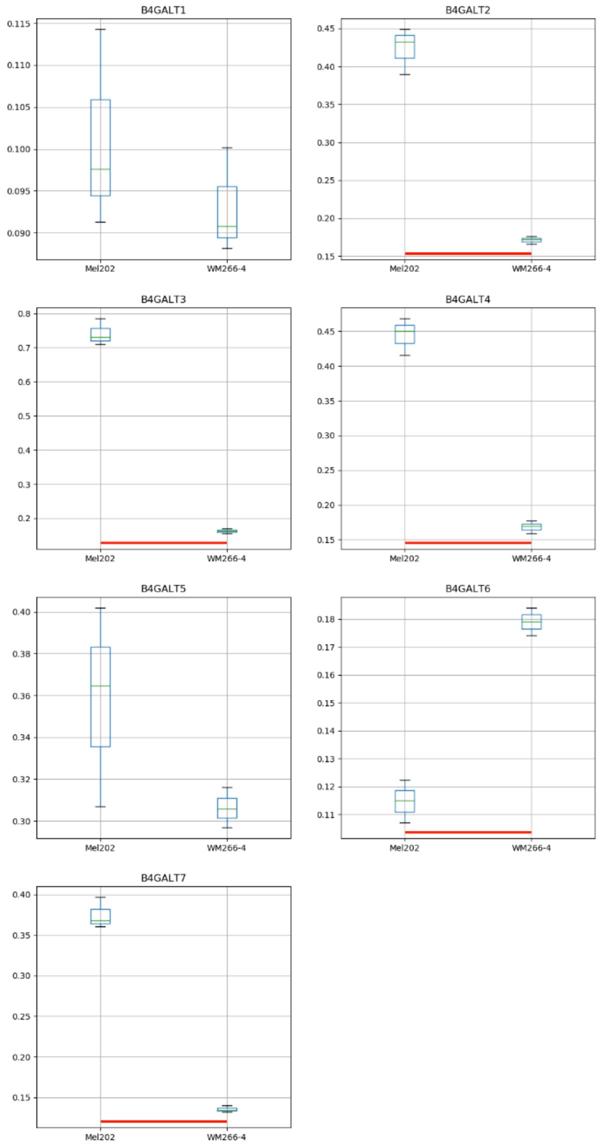

### Model No. 8

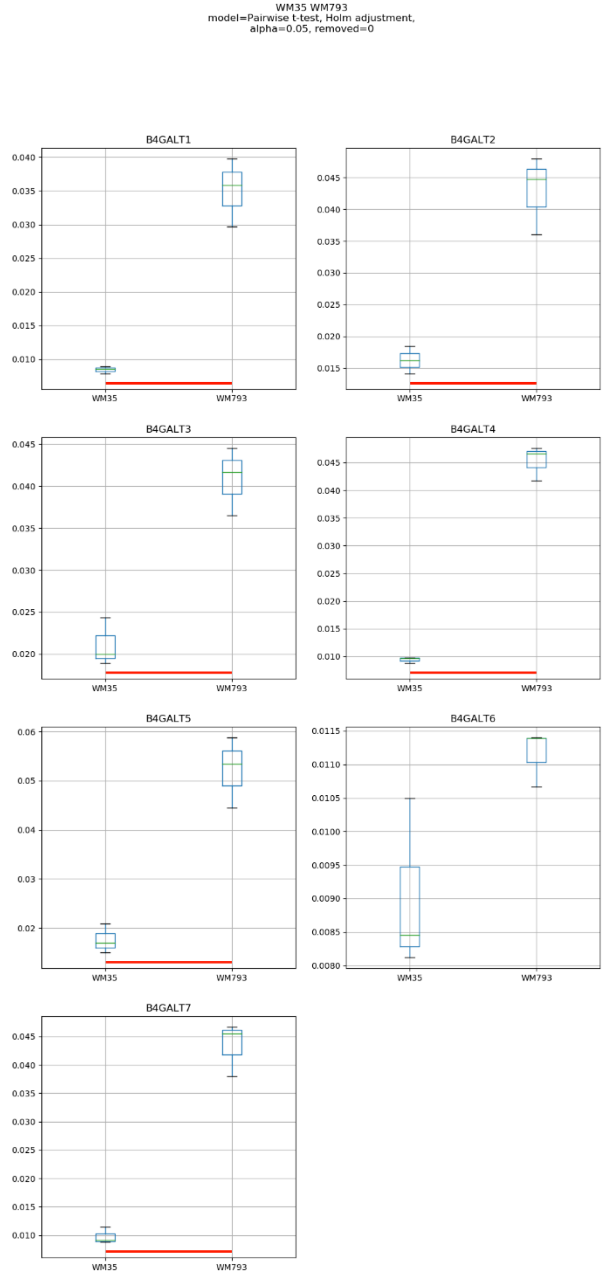
WM35 WM793
model=Pairwise t-test, Holm adjustment,
alpha=0.05, removed=0
B4GALT1
B4GALT2
B4GALT3
B4GALT4
B4GALT5
B4GALT6
B4GALT7
WM35
WM793

### Model No. 9

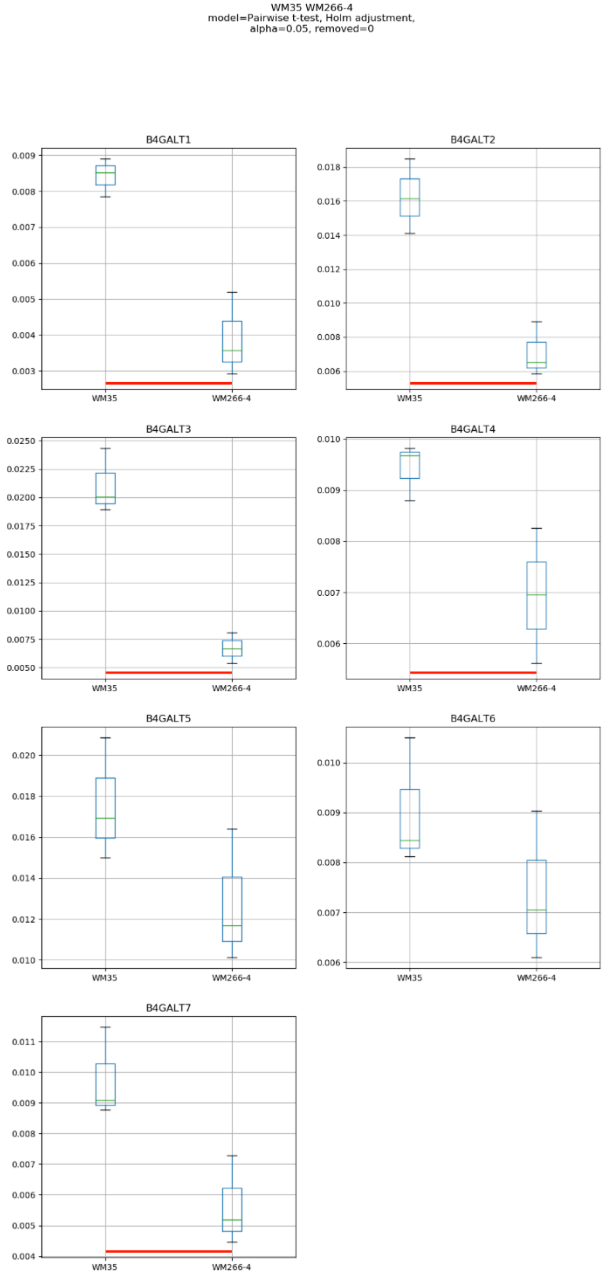
WM35 WM266-4
model=Pairwise t-test, Holm adjustment,
alpha=0.05, removed=0
B4GALT1
B4GALT2
B4GALT3
B4GALT4
B4GALT5
B4GALT6
B4GALT7
WM35
WM266-4

### Model No. 10

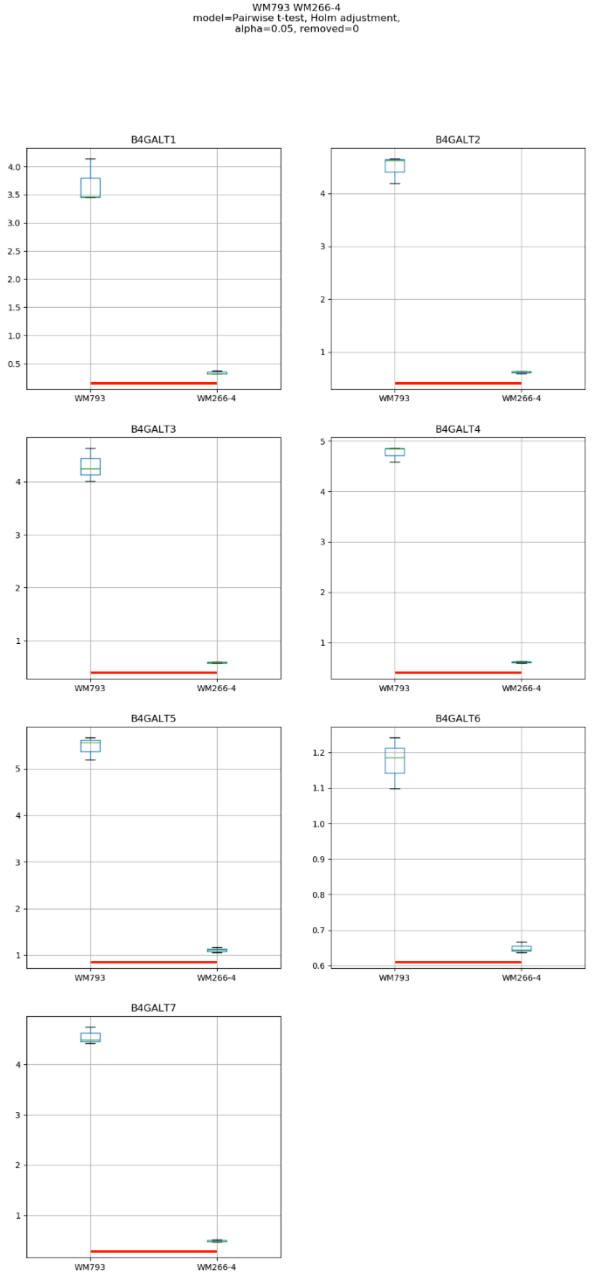
WM793 WM266-4
model=Pairwise t-test, Holm adjustment,
alpha=0.05, removed=0
B4GALT1
B4GALT2
B4GALT3
B4GALT4
B4GALT5
B4GALT6
B4GALT7
WM793
WM266-4

### Model No. 11

HEMa-LP Mel202 WM35
model=Pairwise t-test, Holm adjustment,
alpha=0.05, removed=0

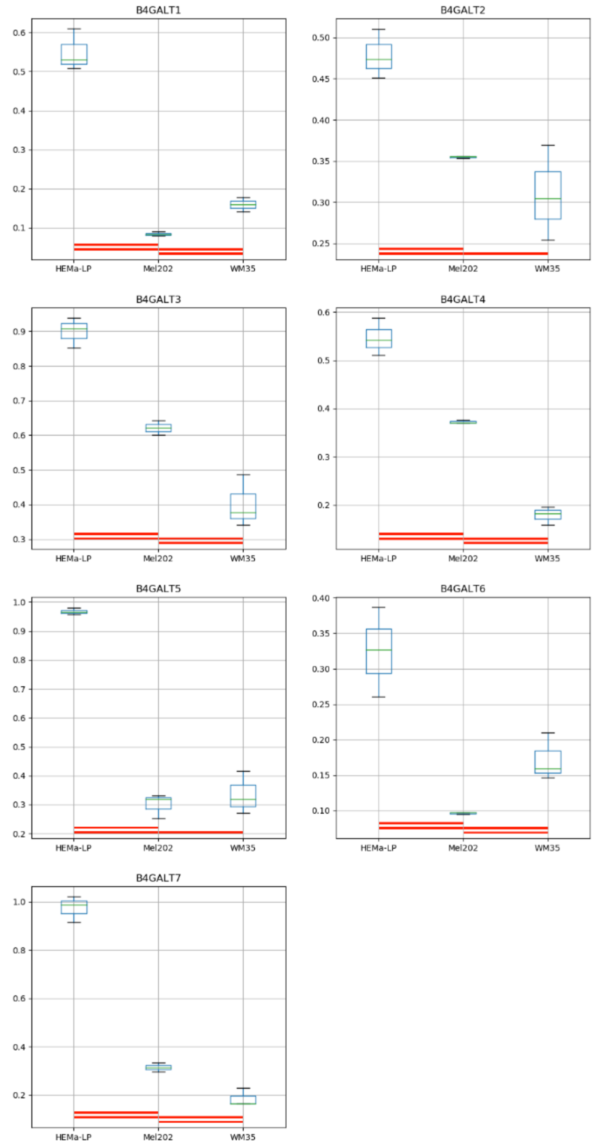

### Model No. 12

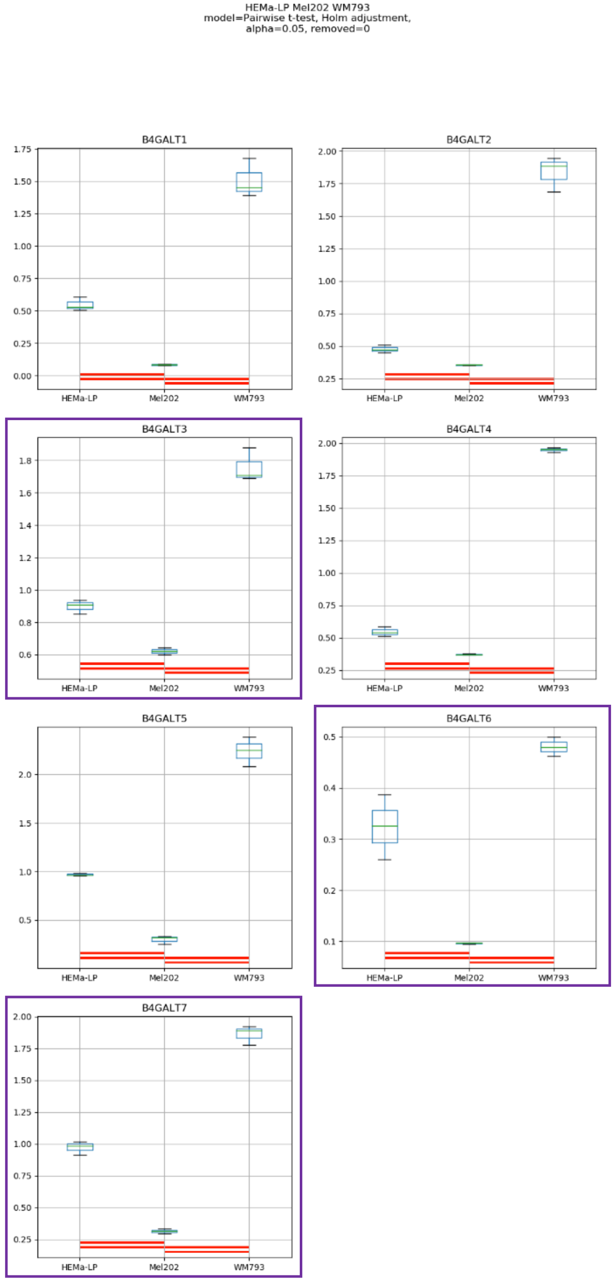
HEMa-LP Mel202 WM793
model=Pairwise t-test, Holm adjustment,
alpha=0.05, removed=0
B4GALT1
B4GALT2
B4GALT3
B4GALT4
B4GALT5
B4GALT6
B4GALT7
HEMa-LP
Mel202
WM793

### Model No. 13

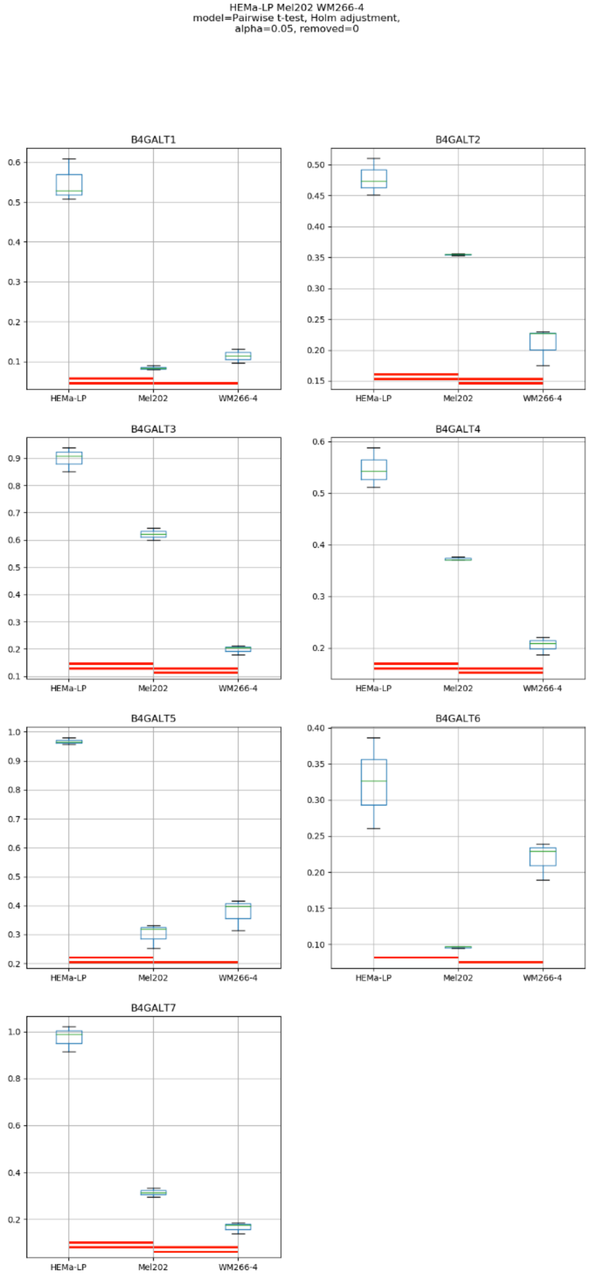
HEMa-LP Mel202 WM266-4
model=Pairwise t-test, Holm adjustment,
alpha=0.05, removed=0
B4GALT1
B4GALT2
B4GALT3
B4GALT4
B4GALT5
B4GALT6
B4GALT7
HEMa-LP
Mel202
WM266-4

Model No. 14

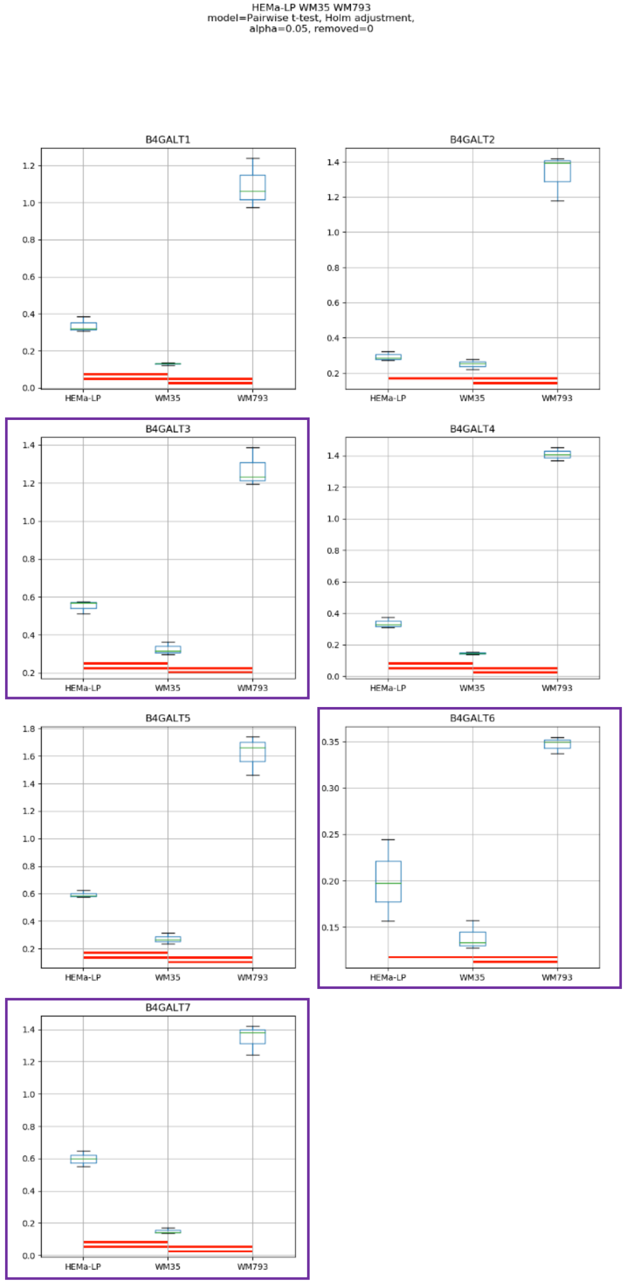
HEMa-LP WM35 WM793
model=Pairwise t-test, Holm adjustment,
alpha=0.05, removed=0
B4GALT1
B4GALT2
B4GALT3
B4GALT4
B4GALT5
B4GALT6
B4GALT7
HEMa-LP
WM35
WM793

Model No. 15

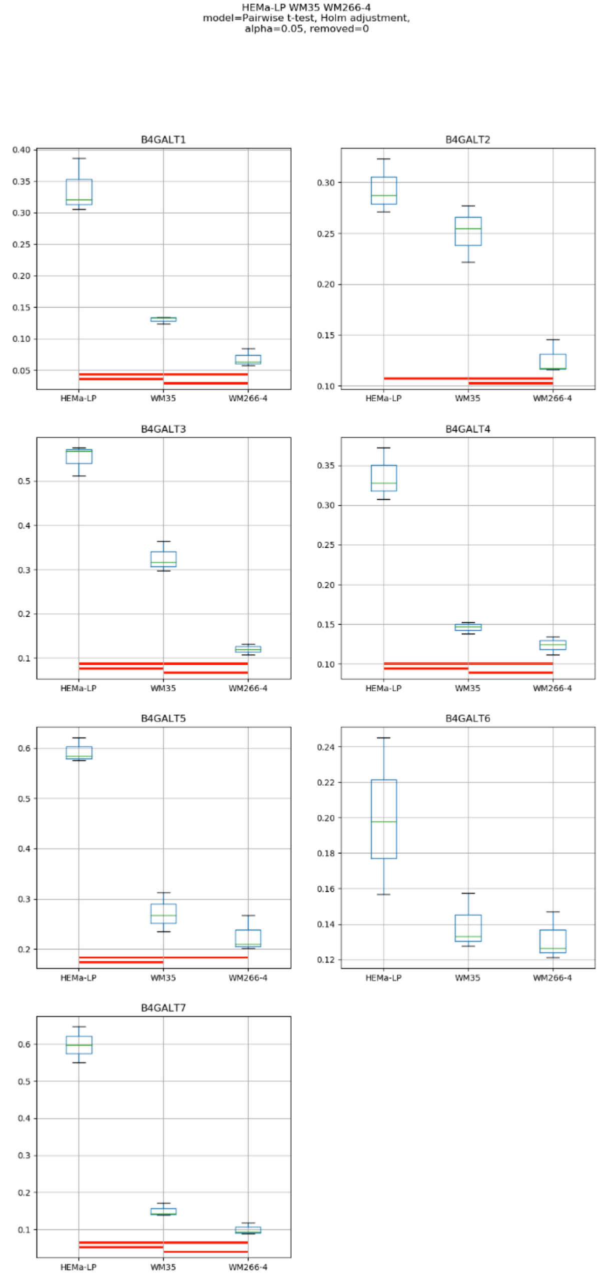
HEMa-LP WM35 WM266-4
model=Pairwise t-test, Holm adjustment,
alpha=0.05, removed=0
B4GALT1
B4GALT2
B4GALT3
B4GALT4
B4GALT5
B4GALT6
B4GALT7
HEMa-LP
WM35
WM266-4

### Model No. 16

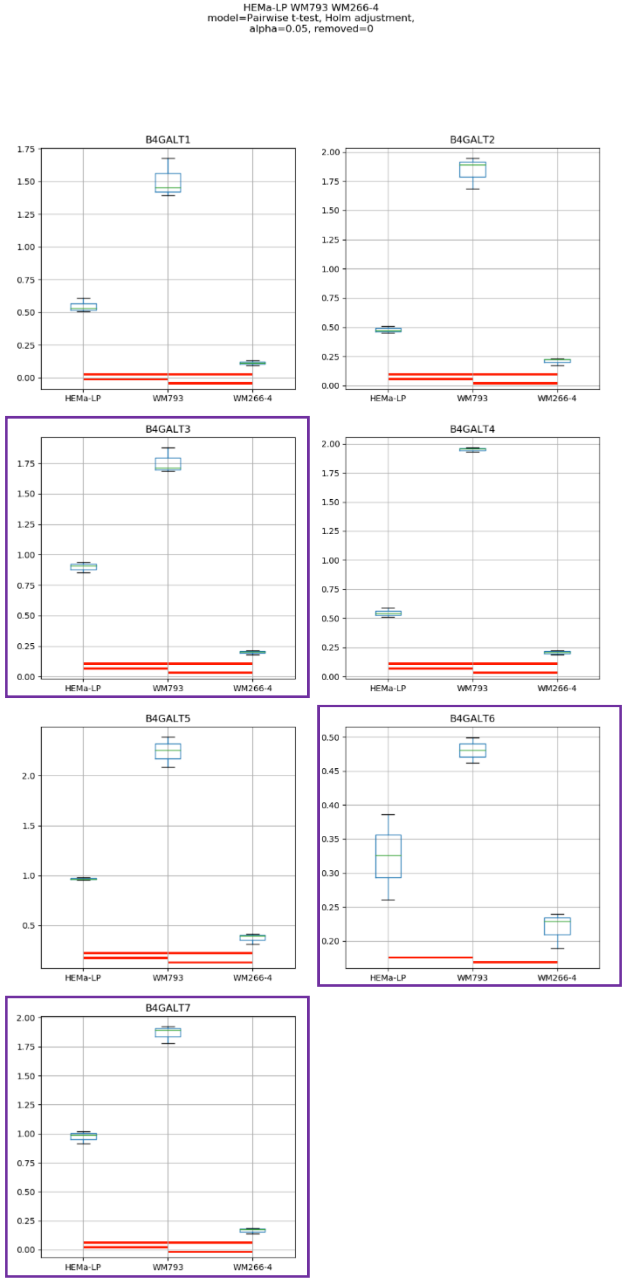
HEMa-LP WM793 WM266-4
model=Pairwise t-test, Holm adjustment,
alpha=0.05, removed=0
B4GALT1
B4GALT2
B4GALT3
B4GALT4
B4GALT5
B4GALT6
B4GALT7
HEMa-LP
WM793
WM266-4

### Model No. 17

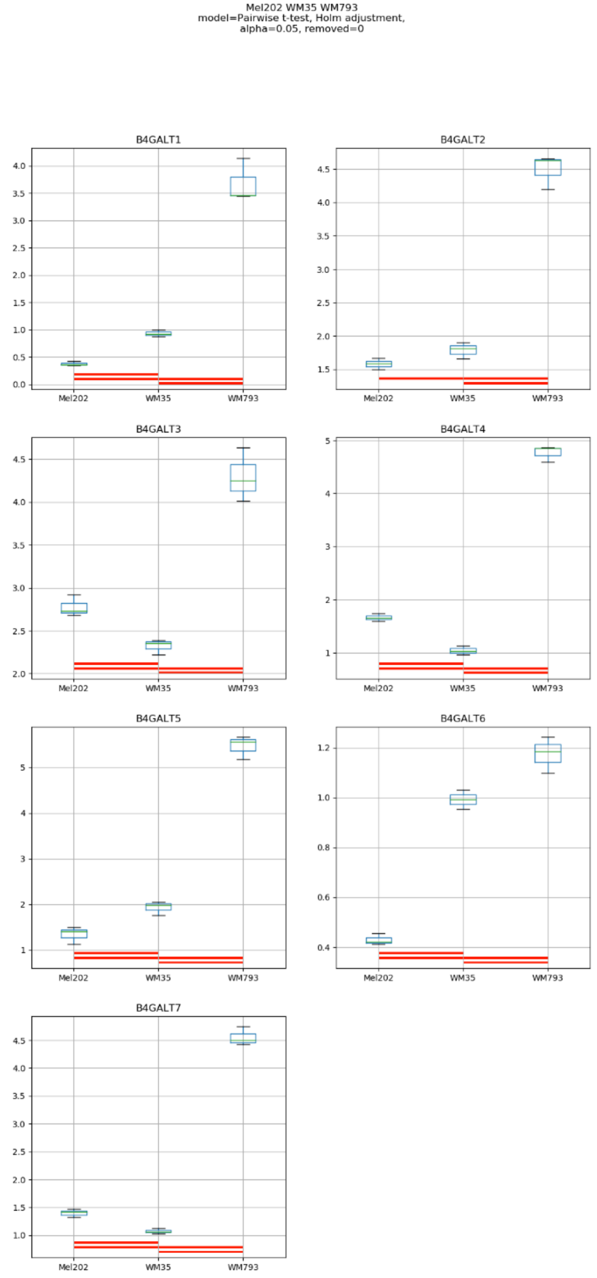
Mel202 WM35 WM793
model=Pairwise t-test, Holm adjustment,
alpha=0.05, removed=0
B4GALT1
B4GALT2
B4GALT3
B4GALT4
B4GALT5
B4GALT6
B4GALT7
Mel202
WM35
WM793

Model No. 18

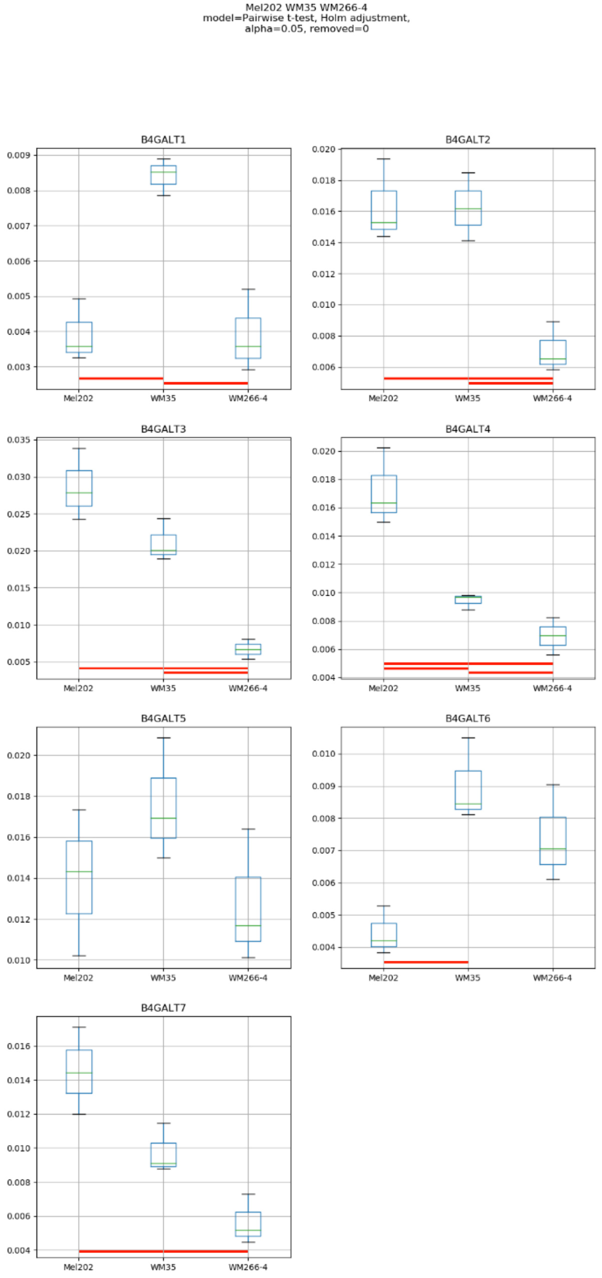

Mel202 WM35 WM266-4
model=Pairwise t-test, Holm adjustment,
alpha=0.05, removed=0
B4GALT1
B4GALT2
B4GALT3
B4GALT4
B4GALT5
B4GALT6
B4GALT7
Mel202
WM35
WM266-4

Model No. 19

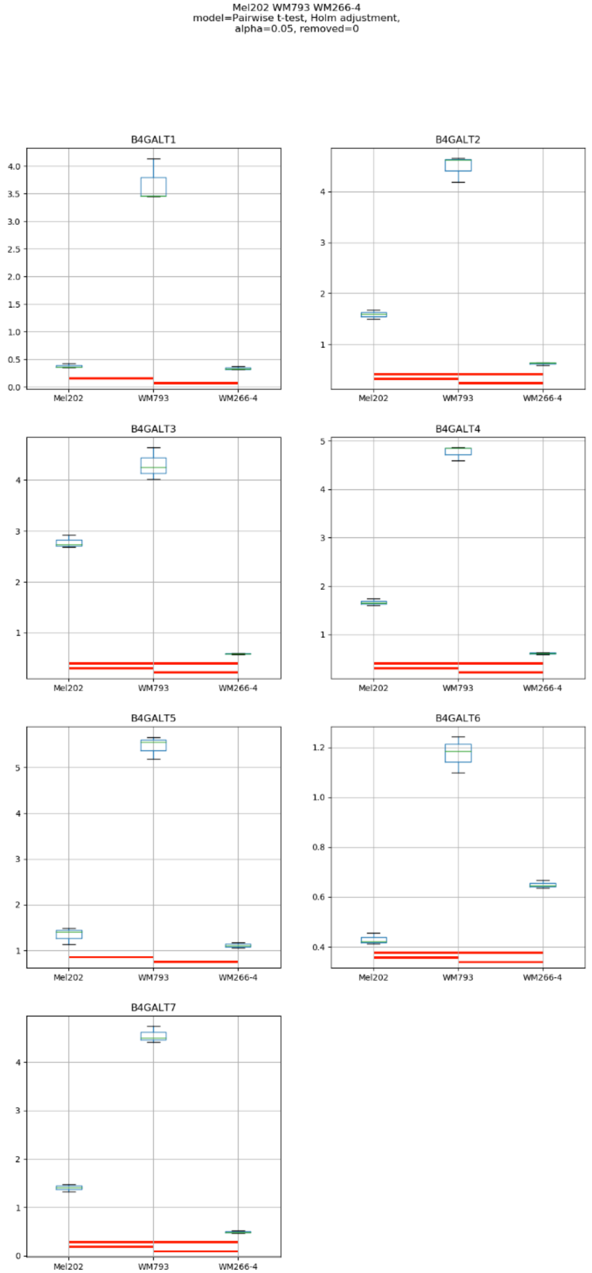
Mel202 WM793 WM266-4
model=Pairwise t-test, Holm adjustment,
alpha=0.05, removed=0
B4GALT1
B4GALT2
B4GALT3
B4GALT4
B4GALT5
B4GALT6
B4GALT7
Mel202
WM793
WM266-4

### Model No. 20

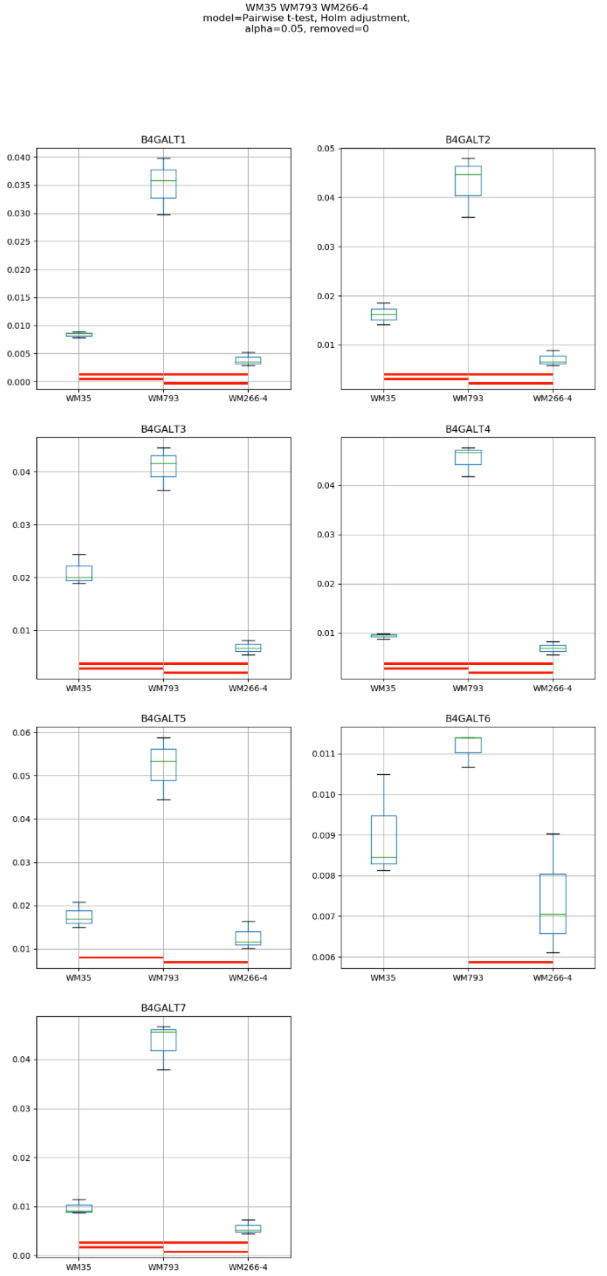
WM35 WM793 WM266-4
model=Pairwise t-test, Holm adjustment,
alpha=0.05, removed=0
B4GALT1
B4GALT2
B4GALT3
B4GALT4
B4GALT5
B4GALT6
B4GALT7
WM35
WM793
WM266-4

### Model No. 21

HEMa-LP Mel202 WM35 WM793
model=Pairwise t-test, Holm adjustment,
alpha=0.05, removed=0

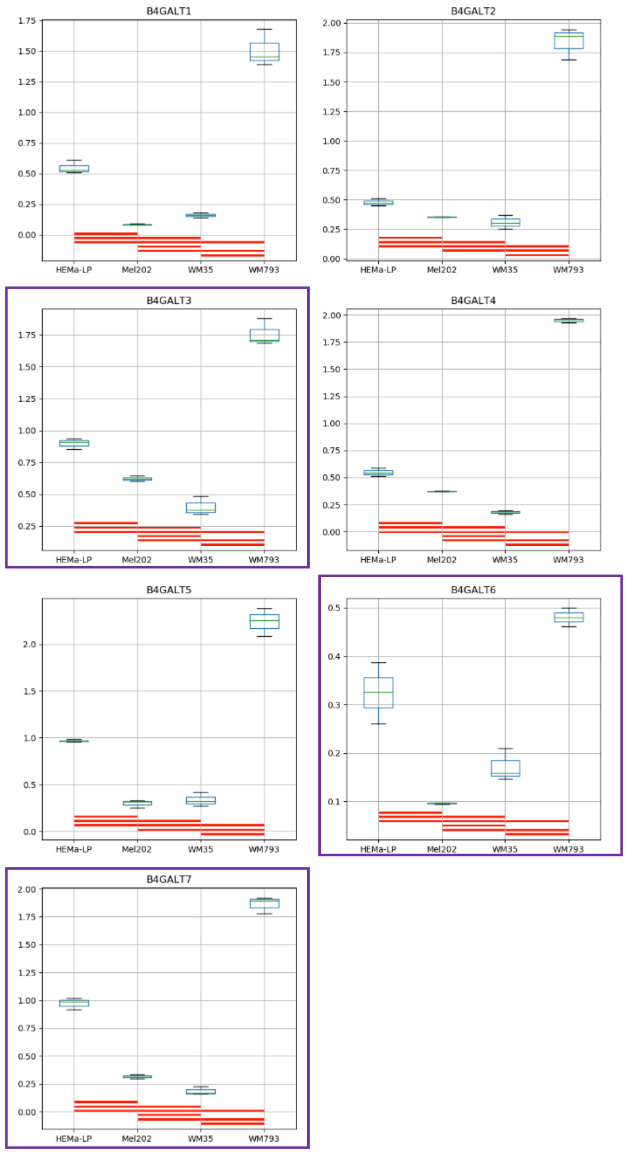

Model No. 22

HEMa-LP Mel202 WM35 WM266-4
model=Pairwise t-test, Holm adjustment,
alpha=0.05, removed=0

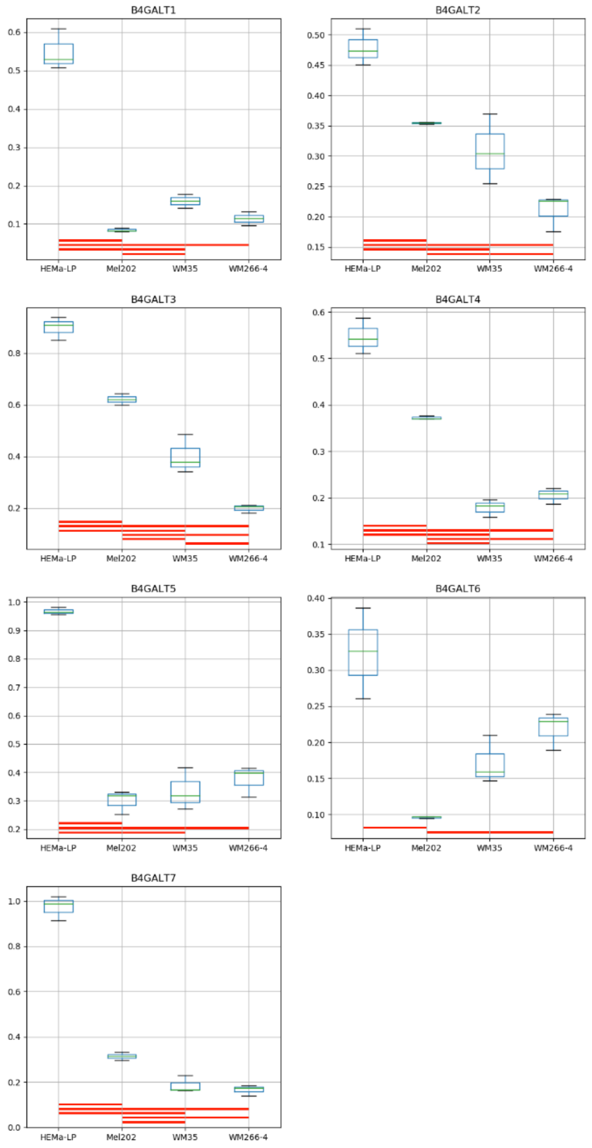

### Model No. 23

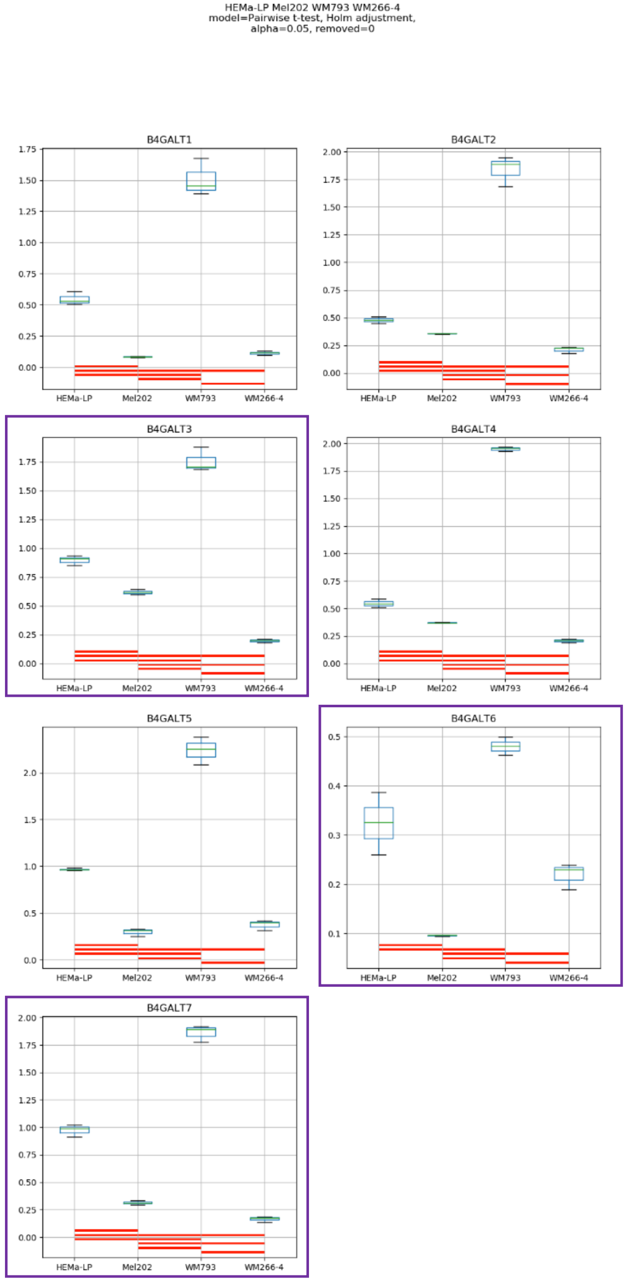
HEMa-LP Mel202 WM793 WM266-4
model=Pairwise t-test, Holm adjustment,
alpha=0.05, removed=0
B4GALT1
B4GALT2
B4GALT3
B4GALT4
B4GALT5
B4GALT6
B4GALT7
HEMa-LP
Mel202
WM793
WM266-4

Model No. 24

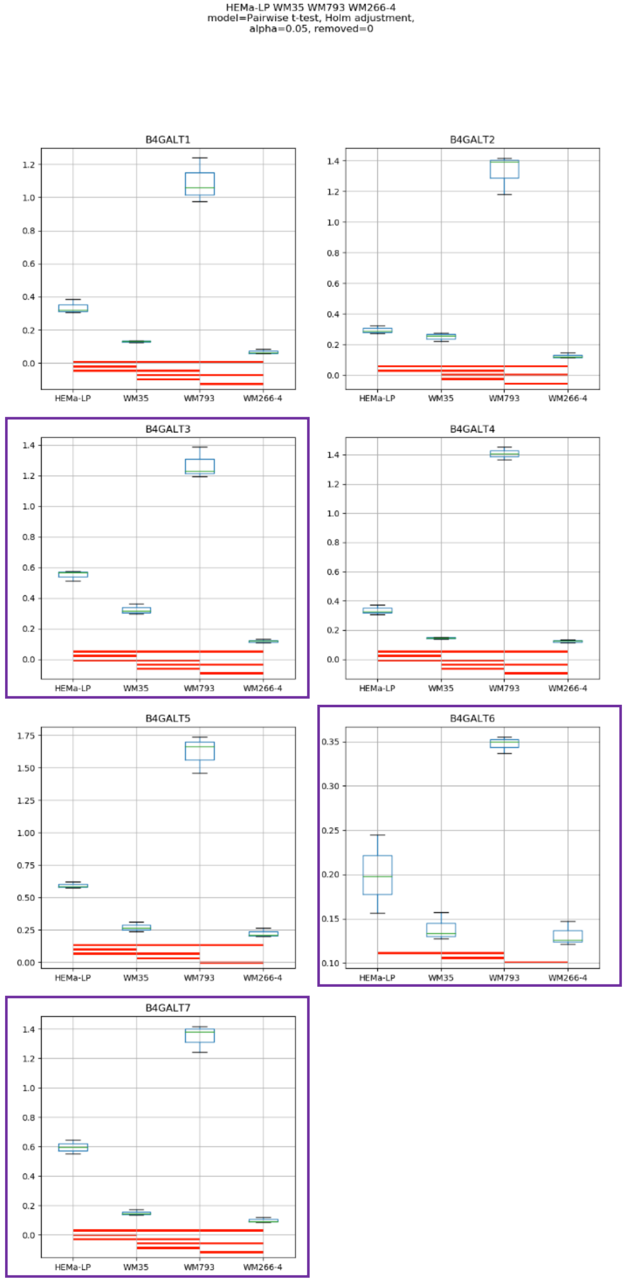
HEMa-LP WM35 WM793 WM266-4
model=Pairwise t-test, Holm adjustment,
alpha=0.05, removed=0
B4GALT1
B4GALT2
B4GALT3
B4GALT4
B4GALT5
B4GALT6
B4GALT7
HEMa-LP
WM35
WM793
WM266-4

### Model No. 25

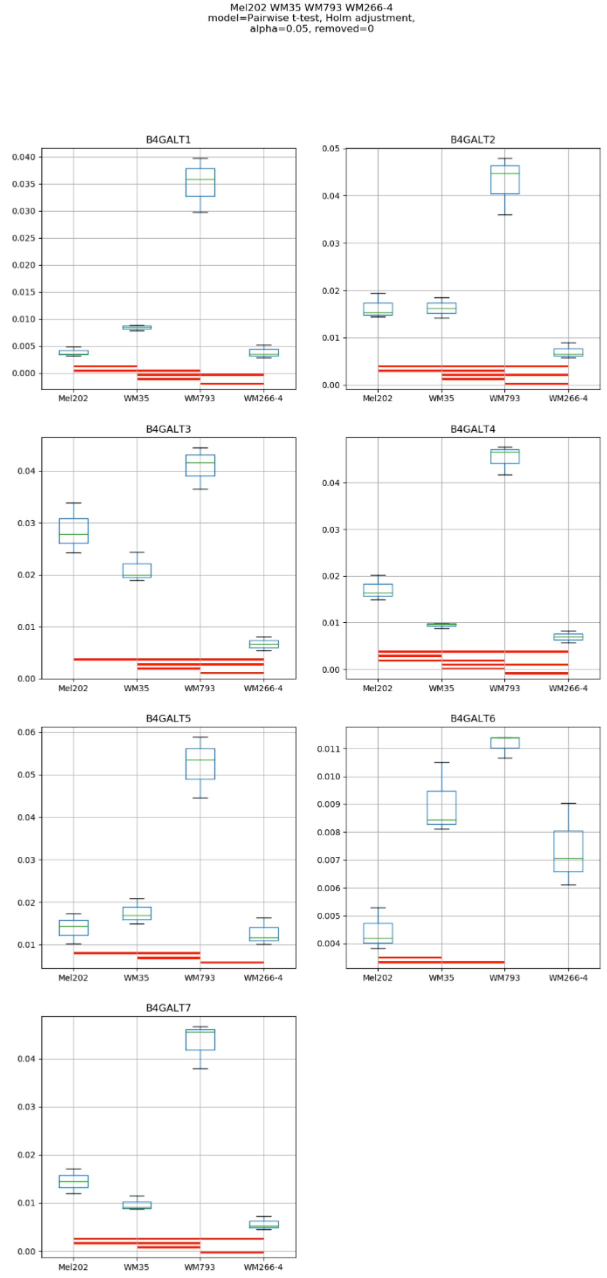
Mel202 WM35 WM793 WM266-4
model=Pairwise t-test, Holm adjustment,
alpha=0.05, removed=0
B4GALT1
B4GALT2
B4GALT3
B4GALT4
B4GALT5
B4GALT6
B4GALT7
Mel202
WM35
WM793
WM266-4

### Model No. 26

HEMa-LP Mel202 WM35 WM793 WM266-4
model=Pairwise t-test, Holm adjustment,
alpha=0.05, removed=0

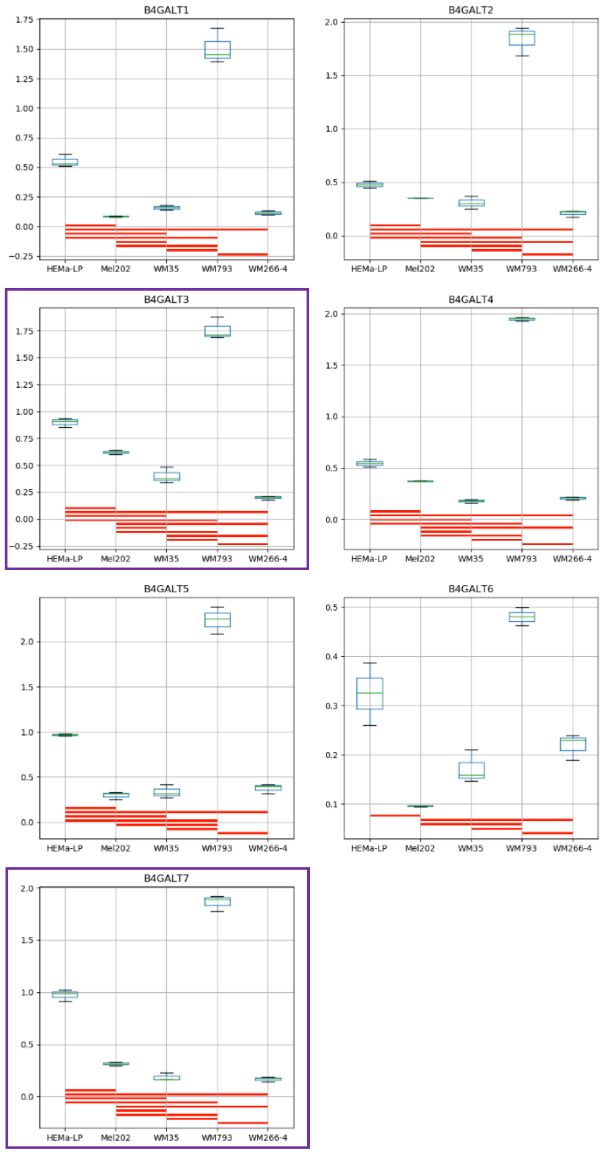

**Supplementary Figure S17:** Medians of relative quantity (RQ) for target genes *B4GALT1–B4GALT7* normalized to the reference gene or pair of reference genes with the best stability values resulting from GenExpA analysis based on set 5 of four candidate reference genes (*GUSB*, *PGK1*, *RPS23*, *HPRT1*) and remove repetition level 0. The statistical analyses used the pairwise t-test with Holm adjustment. Red line represents statistical significance, $P < 0.05$. Frames denote box-plots representing the medians of the RQ values for a given target gene without a repeatable pattern in the same cell lines under different models (the ‘Remove repetitions’ box set to 0). Detailed results are listed in Supplementary Table 16.

### Model No. 1

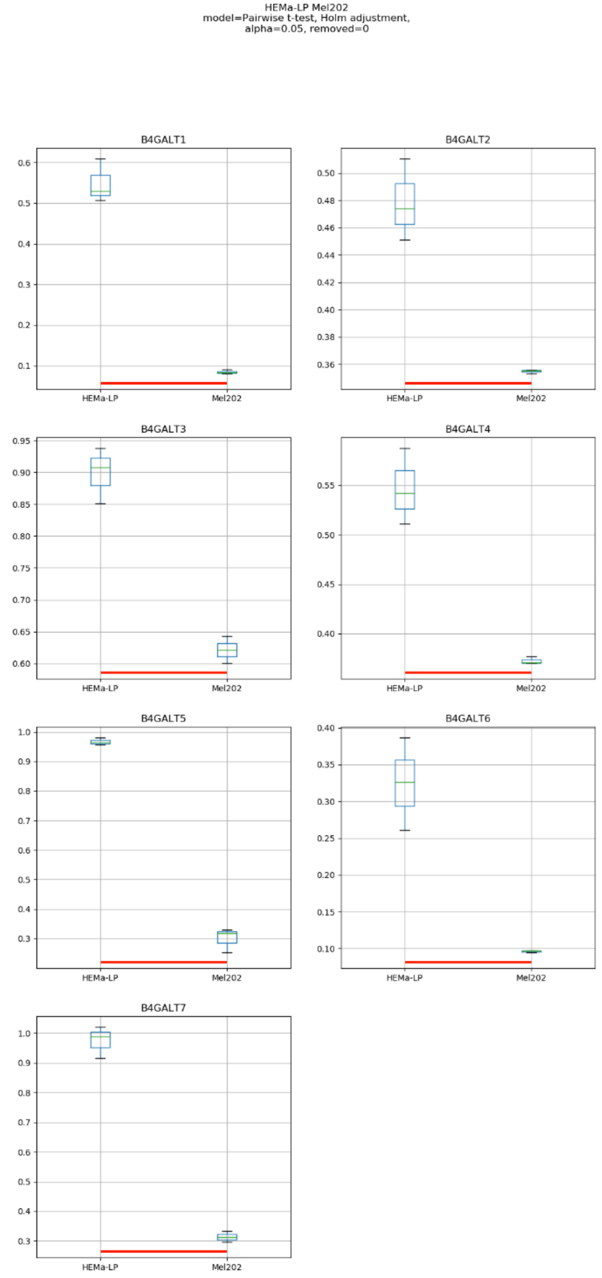
HEMa-LP Mel202
model=Pairwise t-test, Holm adjustment,
alpha=0.05, removed=0
B4GALT1
B4GALT2
B4GALT3
B4GALT4
B4GALT5
B4GALT6
B4GALT7
HEMa-LP
Mel202

Model No. 2

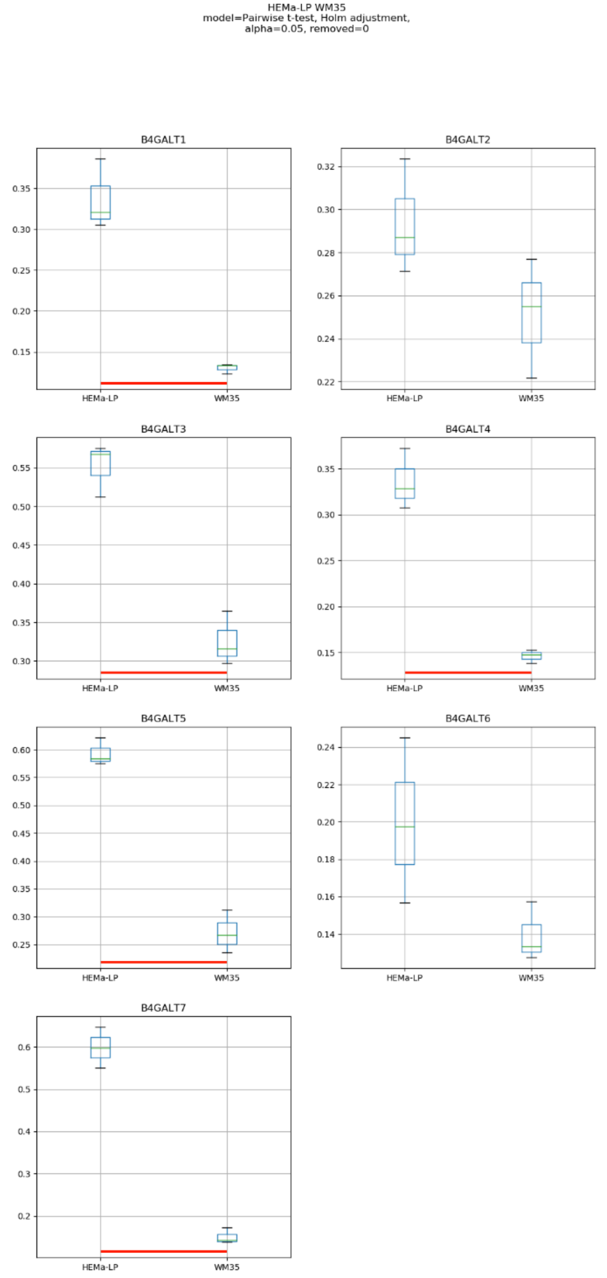
HEMa-LP WM35
model=Pairwise t-test, Holm adjustment,
alpha=0.05, removed=0
B4GALT1
B4GALT2
B4GALT3
B4GALT4
B4GALT5
B4GALT6
B4GALT7
HEMa-LP
WM35

### Model No. 3

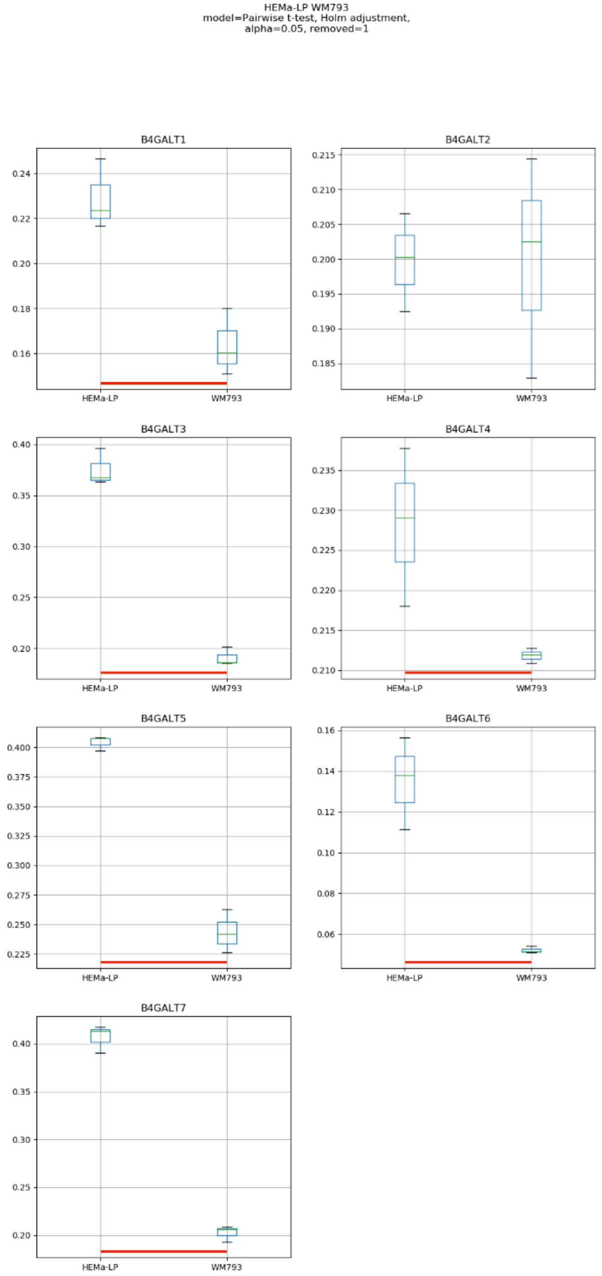
HEMa-LP WM793
model=Pairwise t-test, Holm adjustment,
alpha=0.05, removed=1
B4GALT1
B4GALT2
B4GALT3
B4GALT4
B4GALT5
B4GALT6
B4GALT7
HEMa-LP
WM793

#### Model No. 4

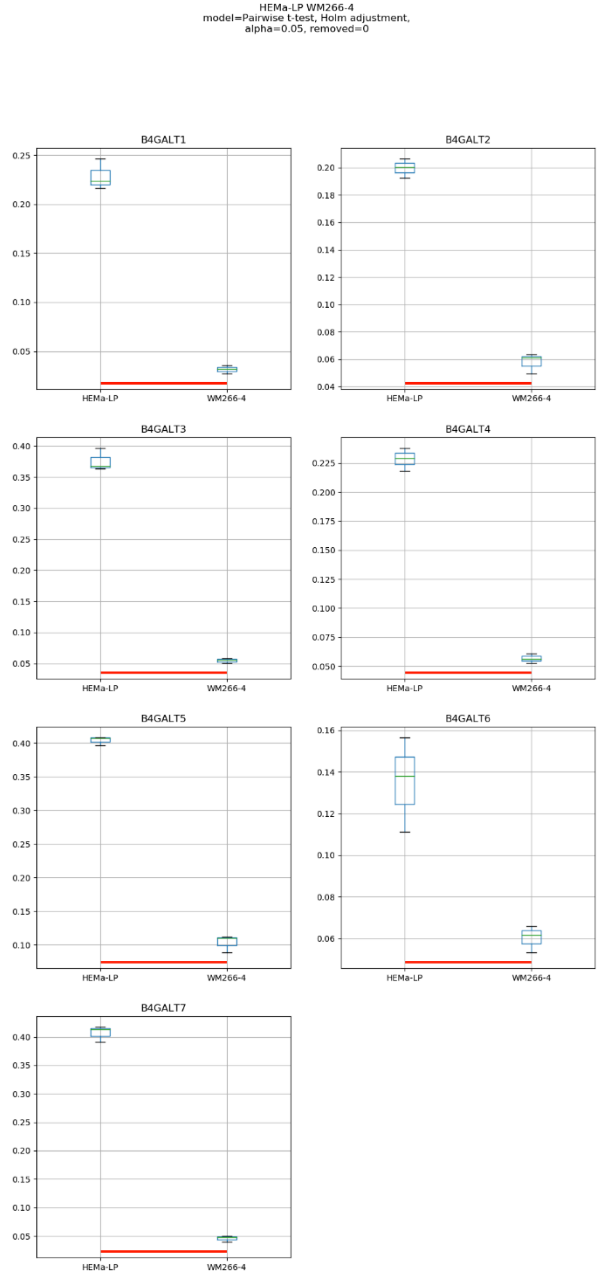

HEMa-LP WM266-4
model=Pairwise t-test, Holm adjustment,
alpha=0.05, removed=0
B4GALT1
0.25
0.20
0.15
0.10
0.05
HEMa-LP
WM266-4
B4GALT2
0.20
0.18
0.16
0.14
0.12
0.10
0.08
0.06
0.04
HEMa-LP
WM266-4
B4GALT3
0.40
0.35
0.30
0.25
0.20
0.15
0.10
0.05
HEMa-LP
WM266-4
B4GALT4
0.225
0.200
0.175
0.150
0.125
0.100
0.075
0.050
HEMa-LP
WM266-4
B4GALT5
0.40
0.35
0.30
0.25
0.20
0.15
0.10
HEMa-LP
WM266-4
B4GALT6
0.16
0.14
0.12
0.10
0.08
0.06
HEMa-LP
WM266-4
B4GALT7
0.40
0.35
0.30
0.25
0.20
0.15
0.10
0.05
HEMa-LP
WM266-4

### Model No. 5

Mel202 WM35
model=Pairwise t-test, Holm adjustment,
alpha=0.05, removed=0

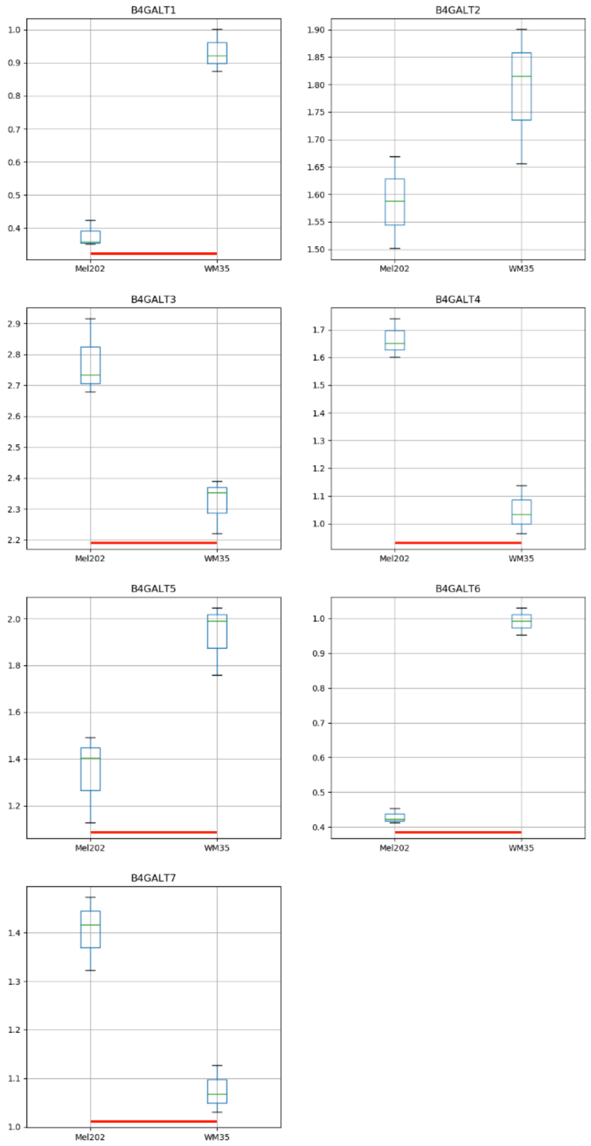

### Model No. 6

Mel202 WM793
model=Pairwise t-test, Holm adjustment,
alpha=0.05, removed=0

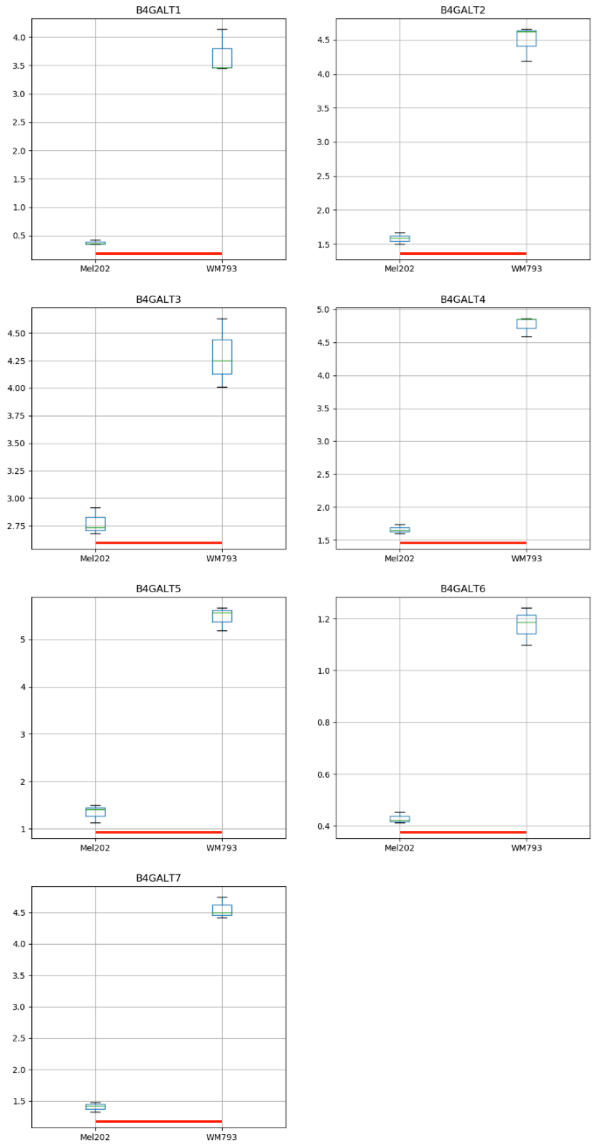

### Model No. 7

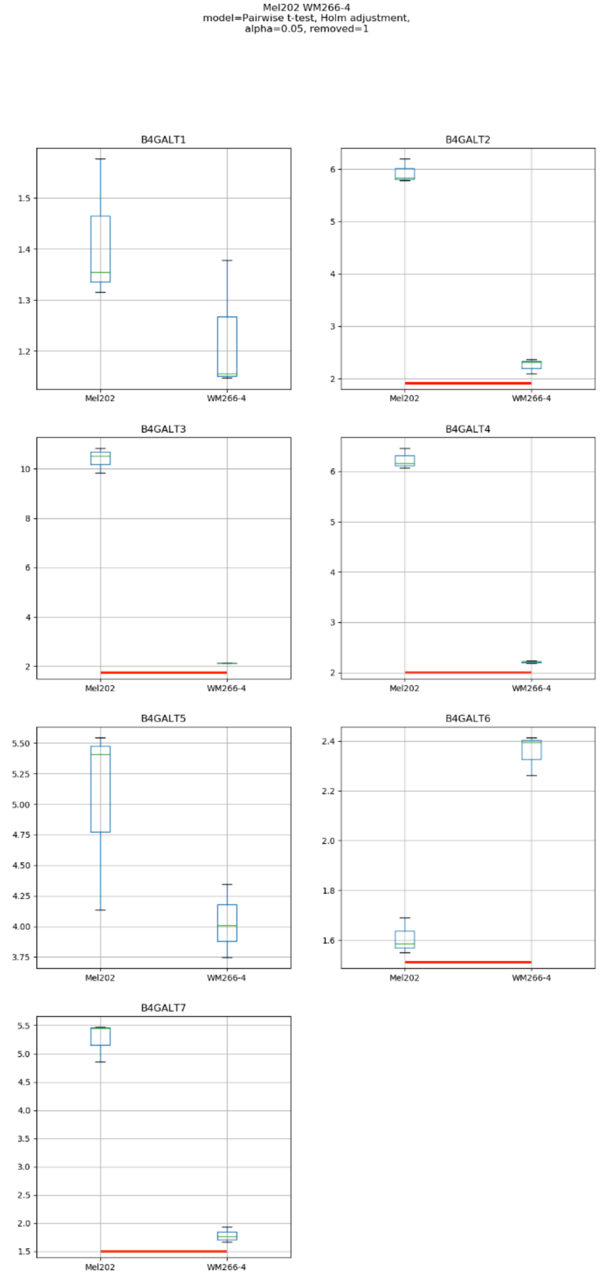

Mel202 WM266-4
model=Pairwise t-test, Holm adjustment,
alpha=0.05, removed=1
B4GALT1
B4GALT2
B4GALT3
B4GALT4
B4GALT5
B4GALT6
B4GALT7
Mel202
WM266-4

### Model No. 8

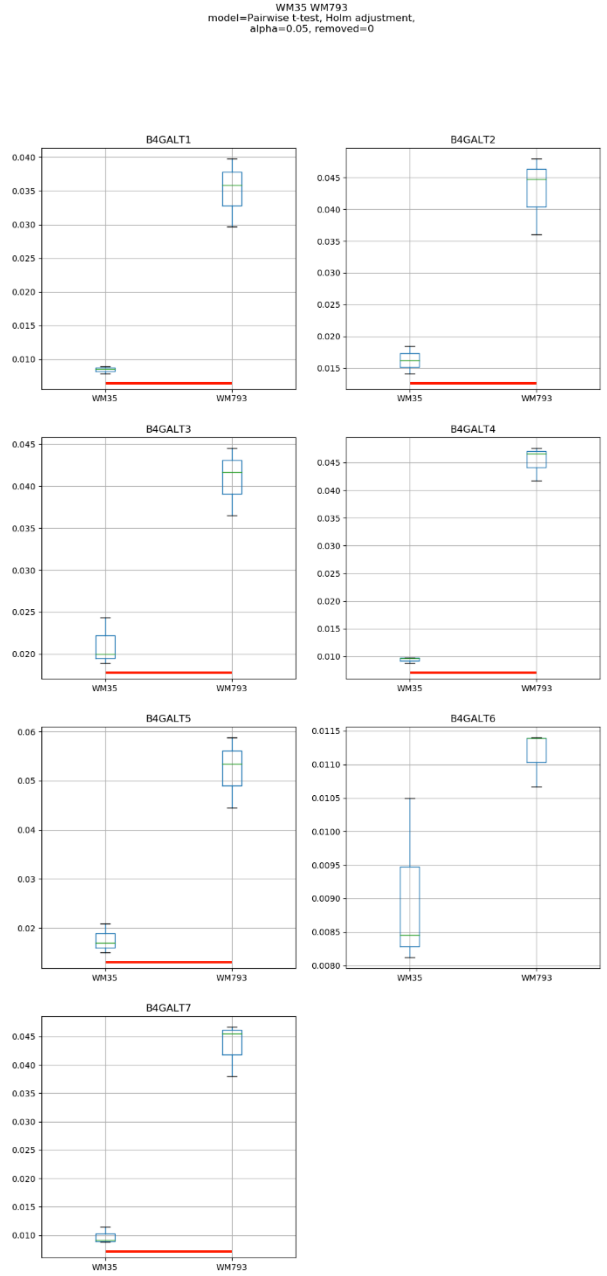
WM35 WM793
model=Pairwise t-test, Holm adjustment,
alpha=0.05, removed=0
B4GALT1
B4GALT2
B4GALT3
B4GALT4
B4GALT5
B4GALT6
B4GALT7
WM35
WM793

### Model No. 9

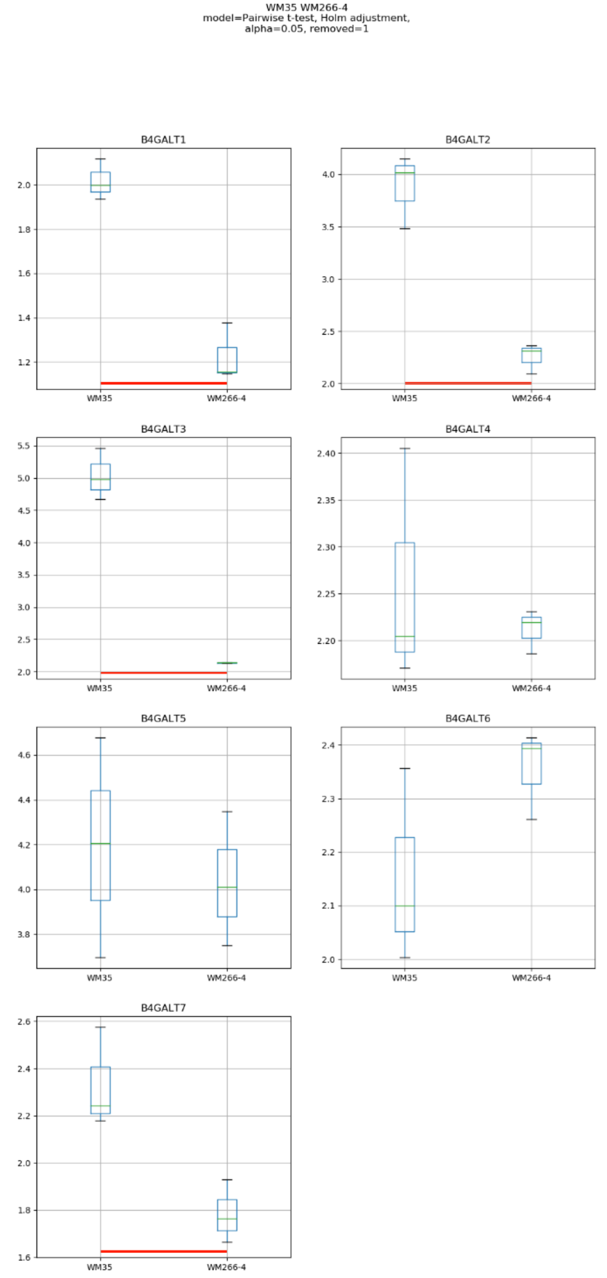
WM35 WM266-4
model=Pairwise t-test, Holm adjustment,
alpha=0.05, removed=1
B4GALT1
B4GALT2
B4GALT3
B4GALT4
B4GALT5
B4GALT6
B4GALT7
WM35
WM266-4

### Model No. 10

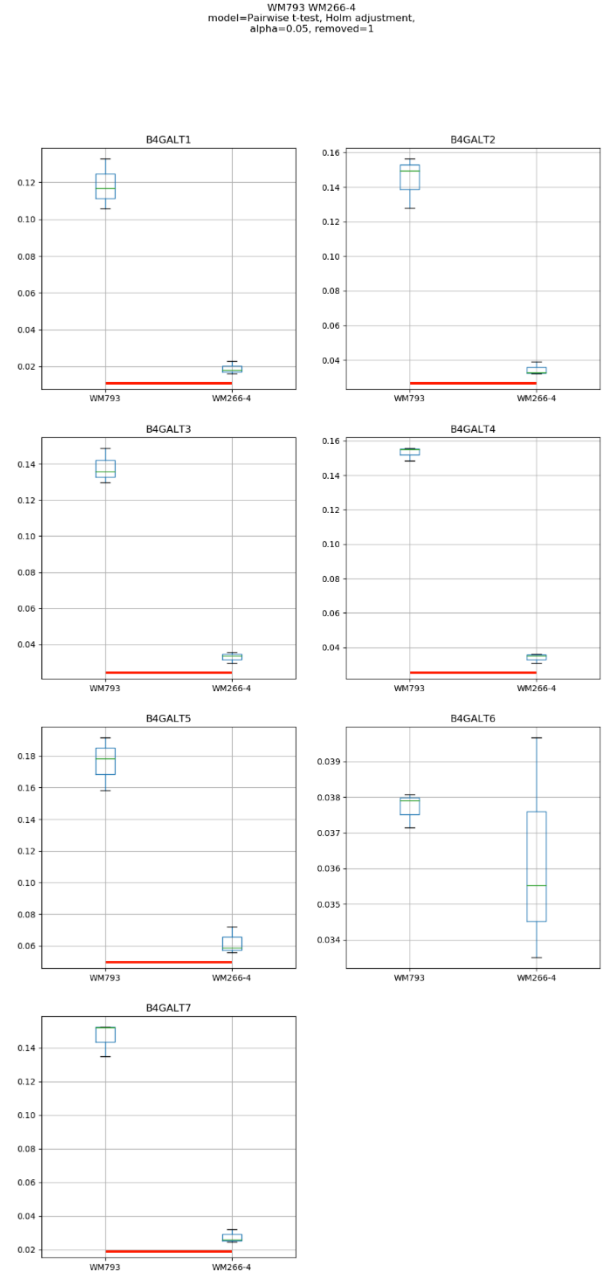
WM793 WM266-4
model=Pairwise t-test, Holm adjustment,
alpha=0.05, removed=1
B4GALT1
B4GALT2
B4GALT3
B4GALT4
B4GALT5
B4GALT6
B4GALT7
WM793
WM266-4

### Model No. 11

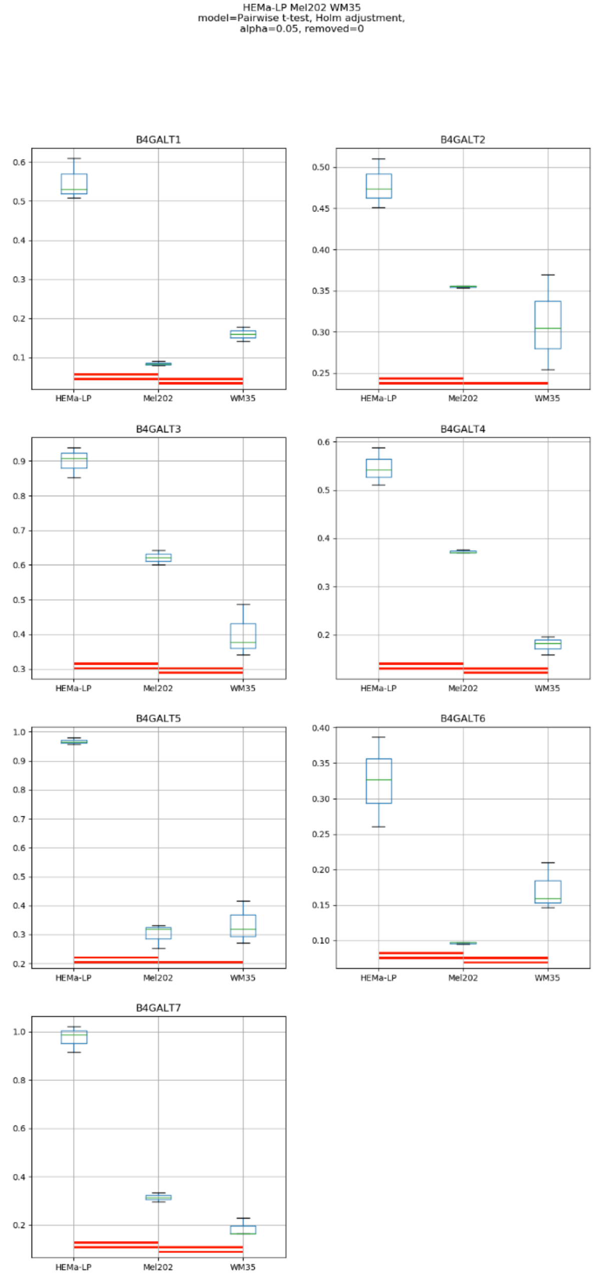
HEMa-LP Mel202 WM35
model=Pairwise t-test, Holm adjustment,
alpha=0.05, removed=0
B4GALT1
B4GALT2
B4GALT3
B4GALT4
B4GALT5
B4GALT6
B4GALT7
HEMa-LP
Mel202
WM35

### Model No. 12

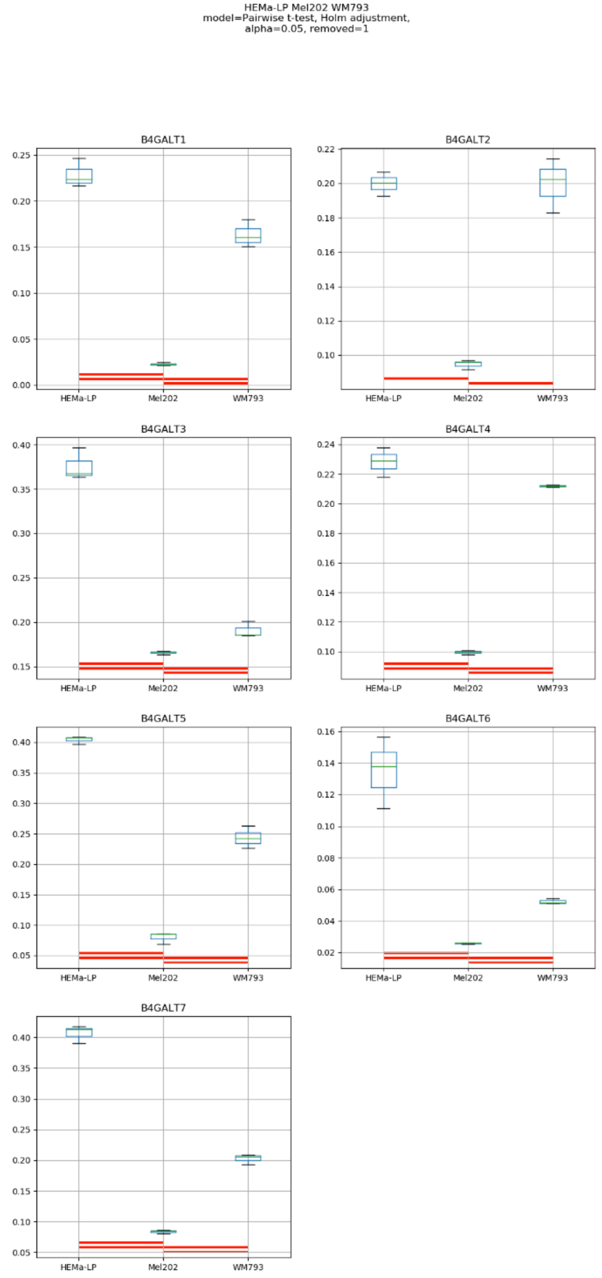
HEMa-LP Mel202 WM793
model=Pairwise t-test, Holm adjustment,
alpha=0.05, removed=1
B4GALT1
B4GALT2
B4GALT3
B4GALT4
B4GALT5
B4GALT6
B4GALT7
HEMa-LP
Mel202
WM793

### Model No. 13

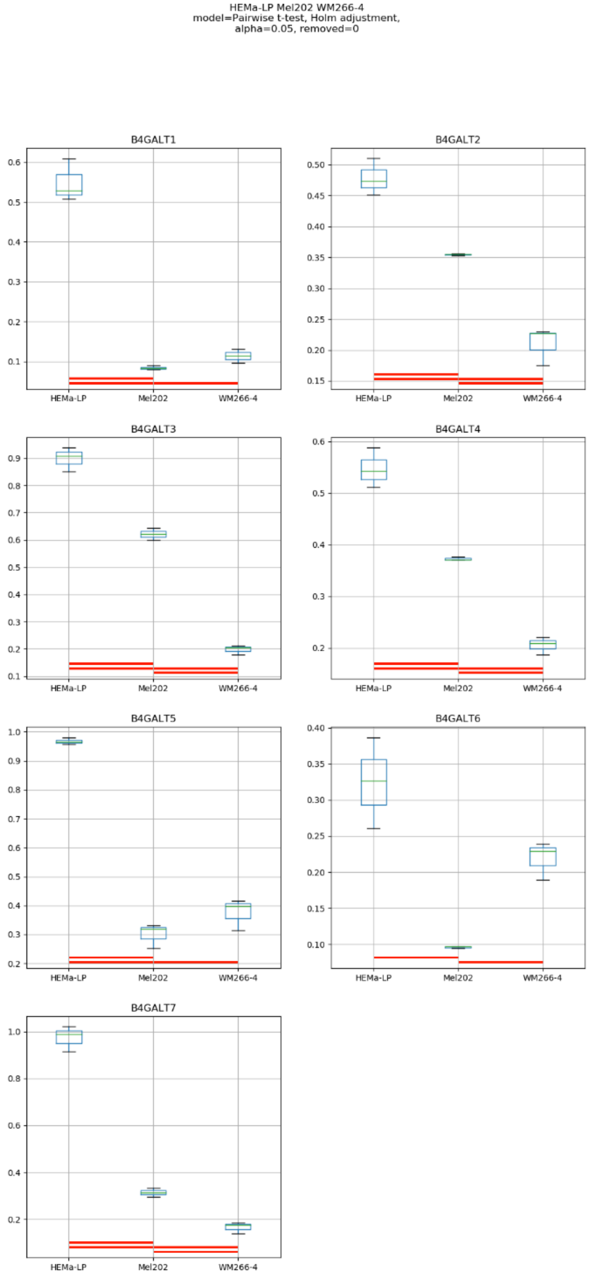
HEMa-LP Mel202 WM266-4
model=Pairwise t-test, Holm adjustment,
alpha=0.05, removed=0
B4GALT1
B4GALT2
B4GALT3
B4GALT4
B4GALT5
B4GALT6
B4GALT7
HEMa-LP
Mel202
WM266-4

### Model No. 14

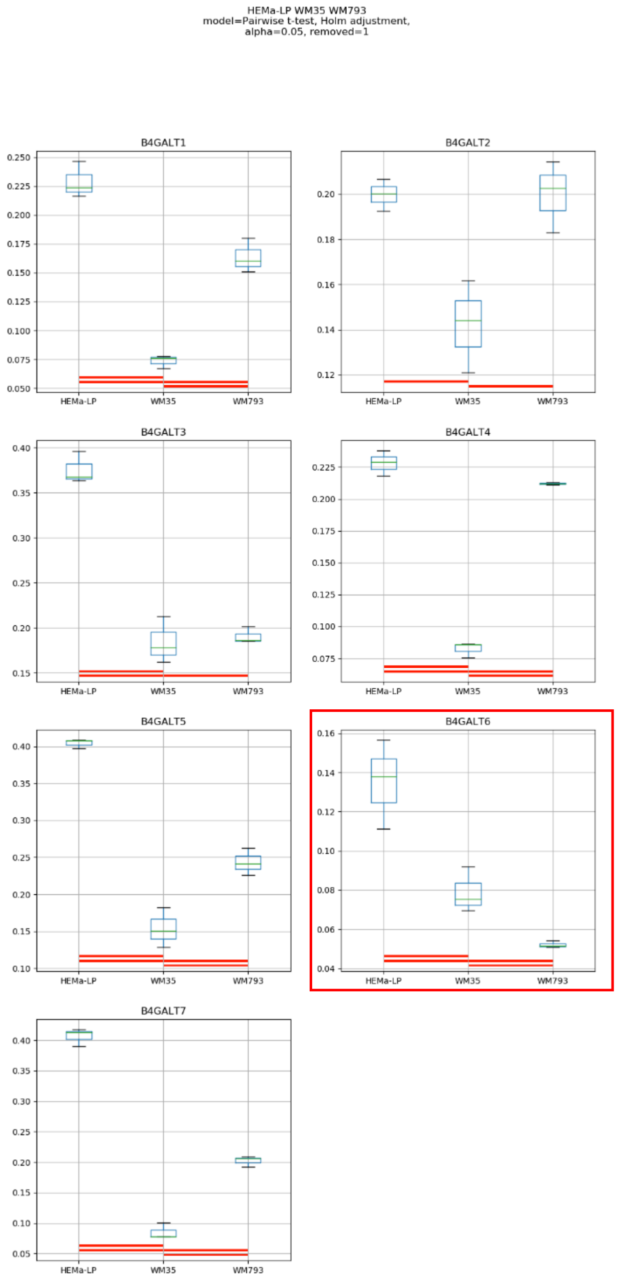
HEMa-LP WM35 WM793
model=Pairwise t-test, Holm adjustment,
alpha=0.05, removed=1
B4GALT1
B4GALT2
B4GALT3
B4GALT4
B4GALT5
B4GALT6
B4GALT7
HEMa-LP
WM35
WM793

### Model No. 15

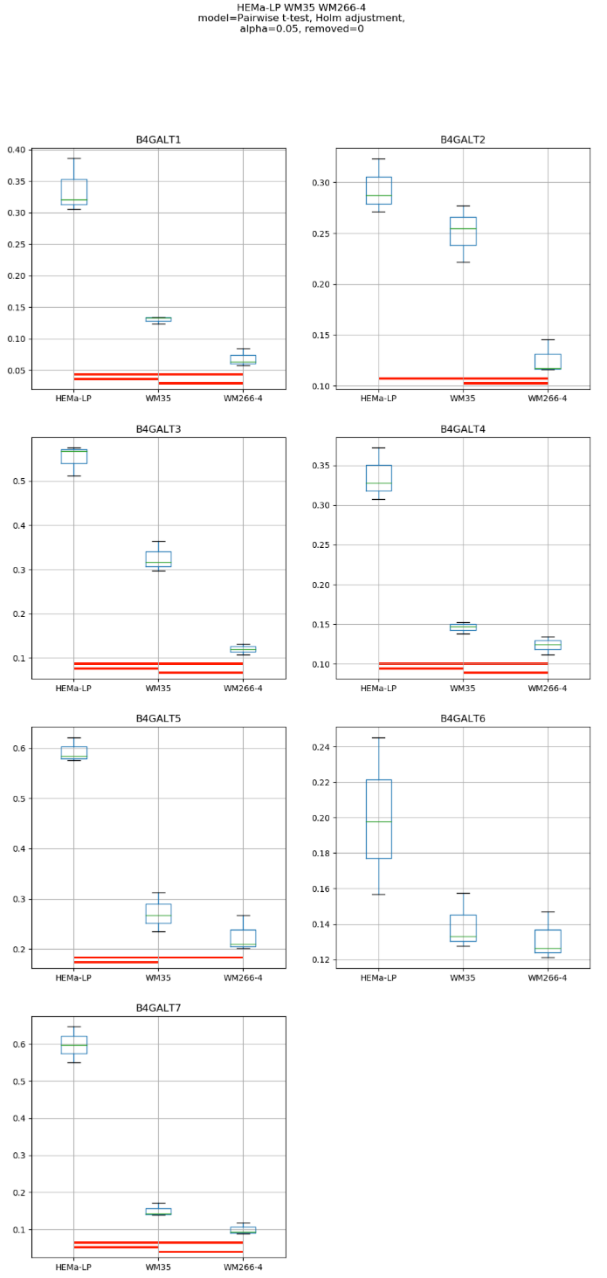
HEMa-LP WM35 WM266-4
model=Pairwise t-test, Holm adjustment,
alpha=0.05, removed=0
B4GALT1
B4GALT2
B4GALT3
B4GALT4
B4GALT5
B4GALT6
B4GALT7
HEMa-LP
WM35
WM266-4

### Model No. 16

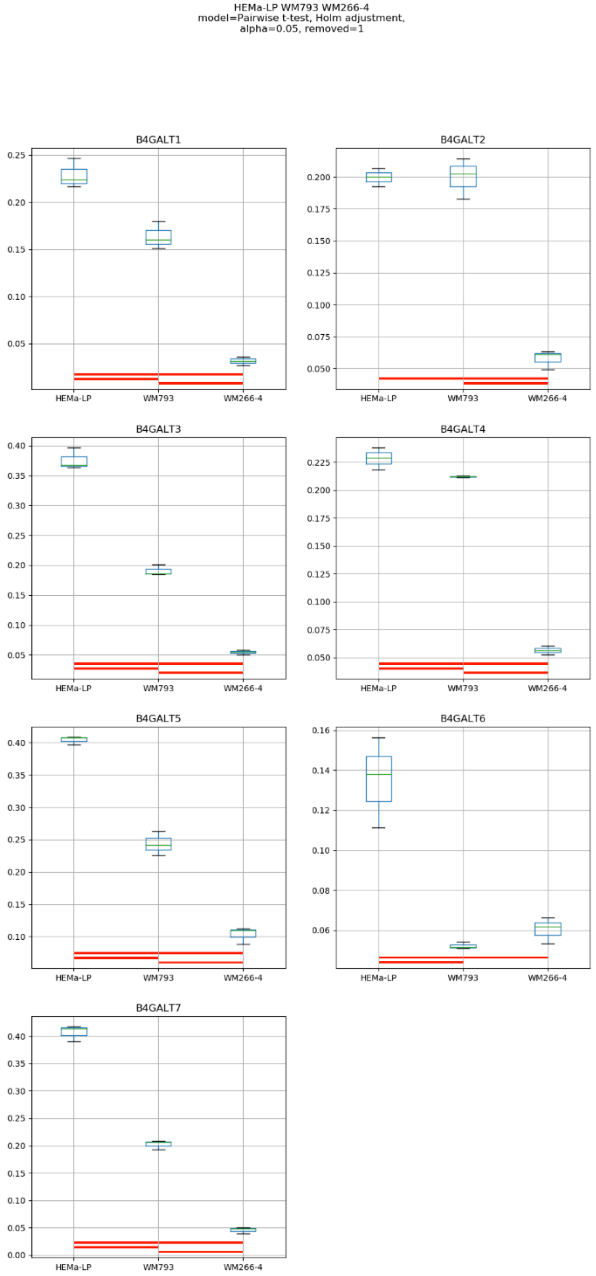
HEMa-LP WM793 WM266-4
model=Pairwise t-test, Holm adjustment,
alpha=0.05, removed=1
B4GALT1
B4GALT2
B4GALT3
B4GALT4
B4GALT5
B4GALT6
B4GALT7
HEMa-LP
WM793
WM266-4

### Model No. 17

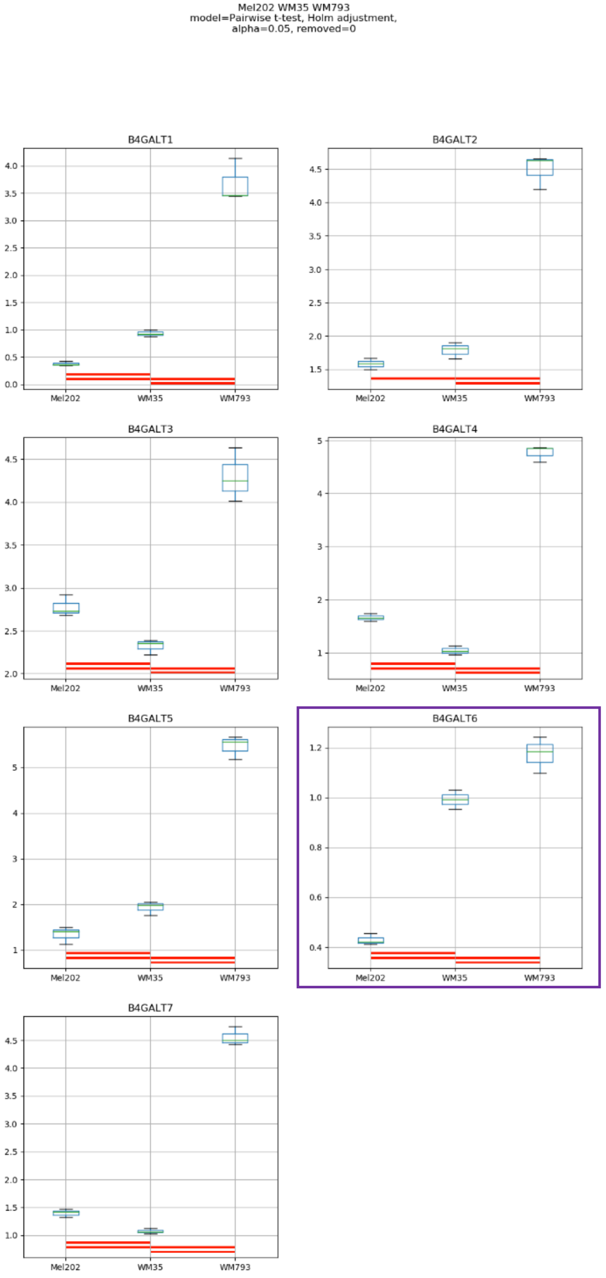
Mel202 WM35 WM793
model=Pairwise t-test, Holm adjustment,
alpha=0.05, removed=0
B4GALT1
B4GALT2
B4GALT3
B4GALT4
B4GALT5
B4GALT6
B4GALT7
Mel202
WM35
WM793

Model No. 18

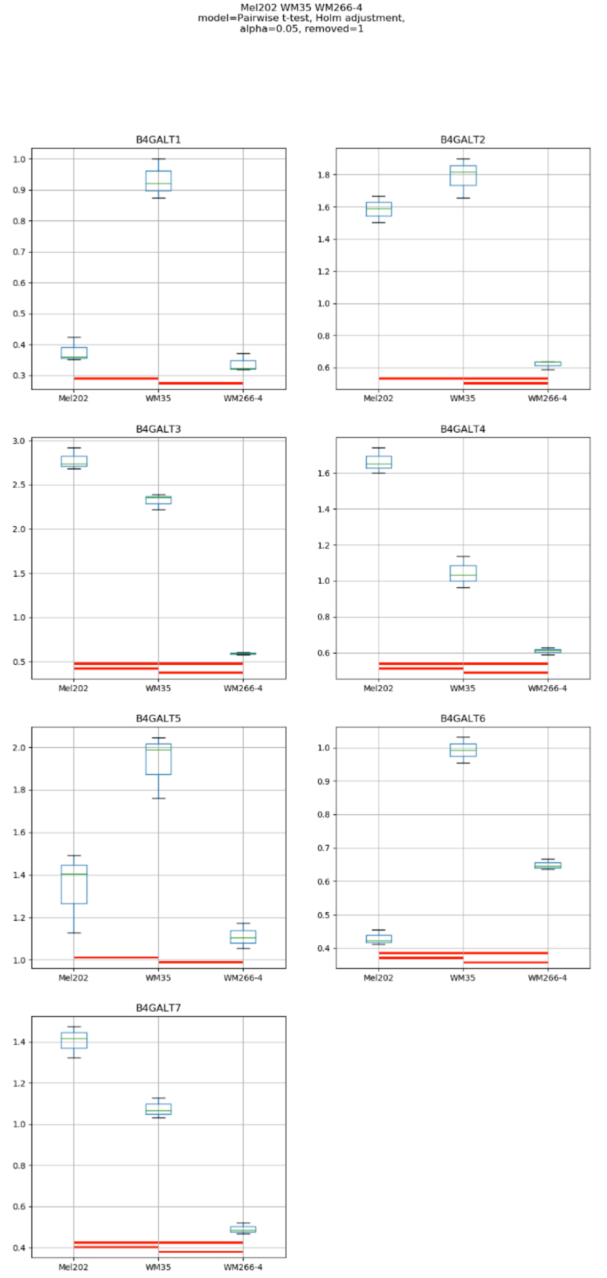
Mel202 WM35 WM266-4
model=Pairwise t-test, Holm adjustment,
alpha=0.05, removed=1
B4GALT1
B4GALT2
B4GALT3
B4GALT4
B4GALT5
B4GALT6
B4GALT7
Mel202
WM35
WM266-4

### Model No. 19

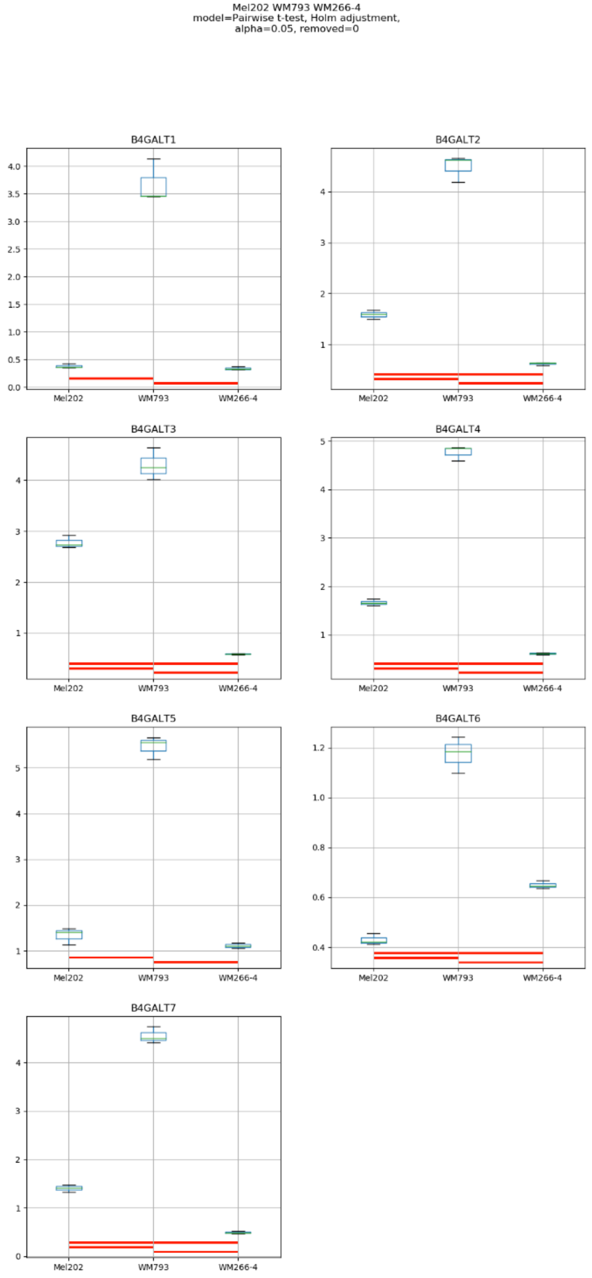
Mel202 WM793 WM266-4
model=Pairwise t-test, Holm adjustment,
alpha=0.05, removed=0
B4GALT1
B4GALT2
B4GALT3
B4GALT4
B4GALT5
B4GALT6
B4GALT7
Mel202
WM793
WM266-4

Model No. 20

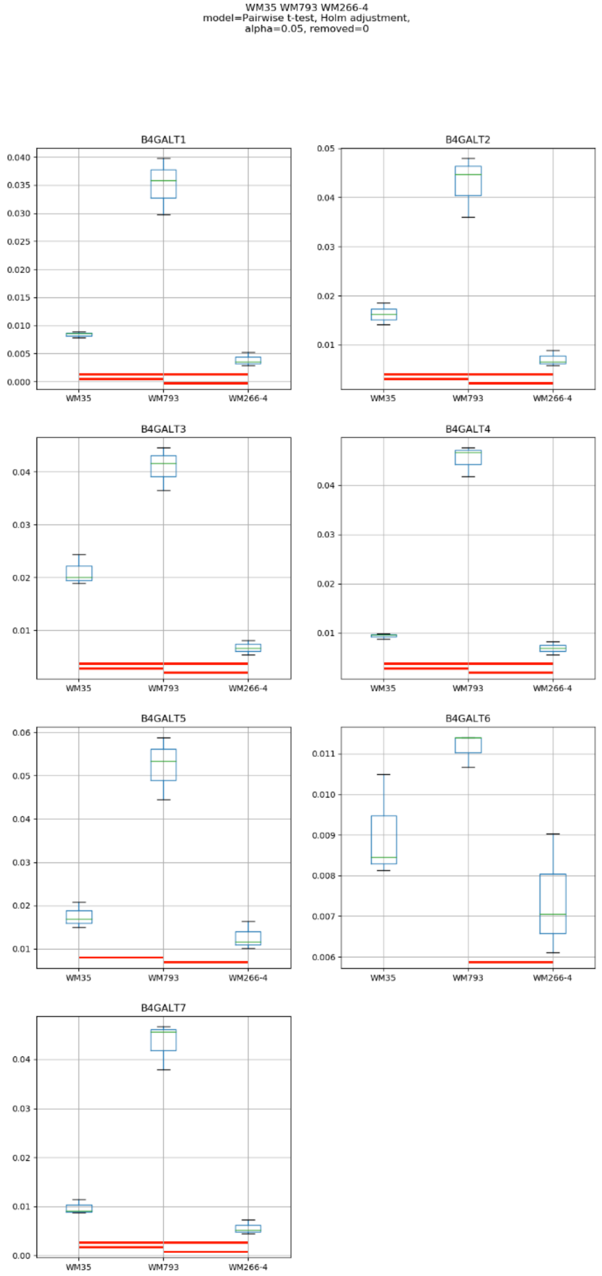
WM35 WM793 WM266-4
model=Pairwise t-test, Holm adjustment,
alpha=0.05, removed=0
B4GALT1
B4GALT2
B4GALT3
B4GALT4
B4GALT5
B4GALT6
B4GALT7
WM35
WM793
WM266-4

### Model No. 21

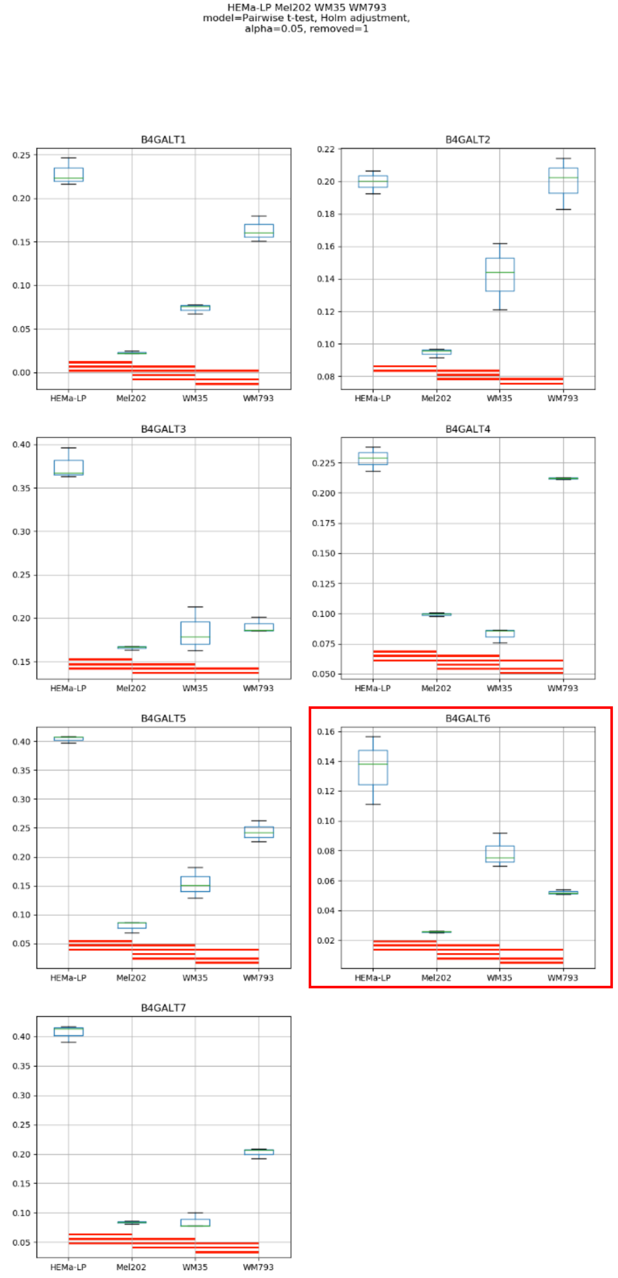
HEMa-LP Mel202 WM35 WM793
model=Pairwise t-test, Holm adjustment,
alpha=0.05, removed=1
B4GALT1
B4GALT2
B4GALT3
B4GALT4
B4GALT5
B4GALT6
B4GALT7
HEMa-LP
Mel202
WM35
WM793

### Model No. 22

HEMa-LP Mel202 WM35 WM266-4
model=Pairwise t-test, Holm adjustment,
alpha=0.05, removed=0

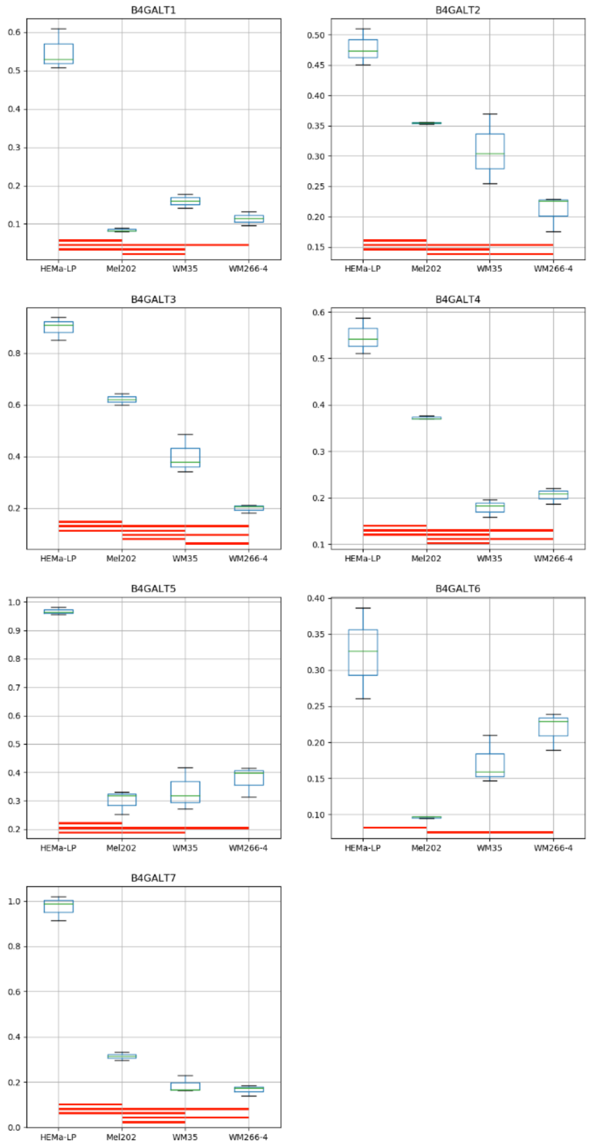

Model No. 23

HEMa-LP Mel202 WM793 WM266-4
model=Pairwise t-test, Holm adjustment,
alpha=0.05, removed=1

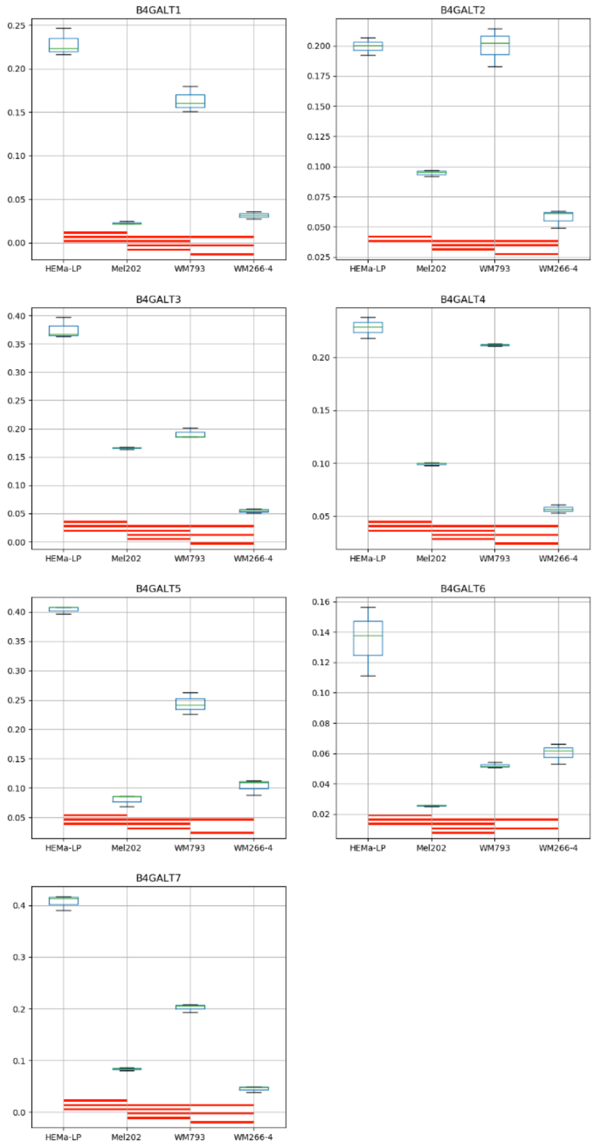

### Model No. 24

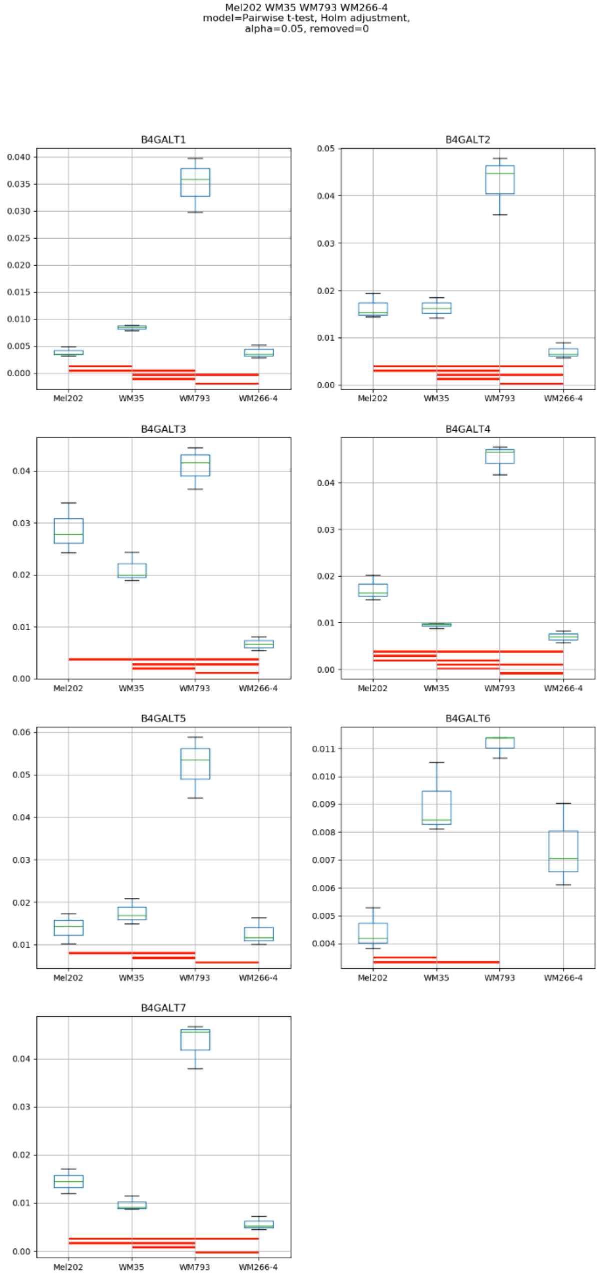
Mel202 WM35 WM793 WM266-4
model=Pairwise t-test, Holm adjustment,
alpha=0.05, removed=0
B4GALT1
B4GALT2
B4GALT3
B4GALT4
B4GALT5
B4GALT6
B4GALT7
Mel202
WM35
WM793
WM266-4

### Model No. 25

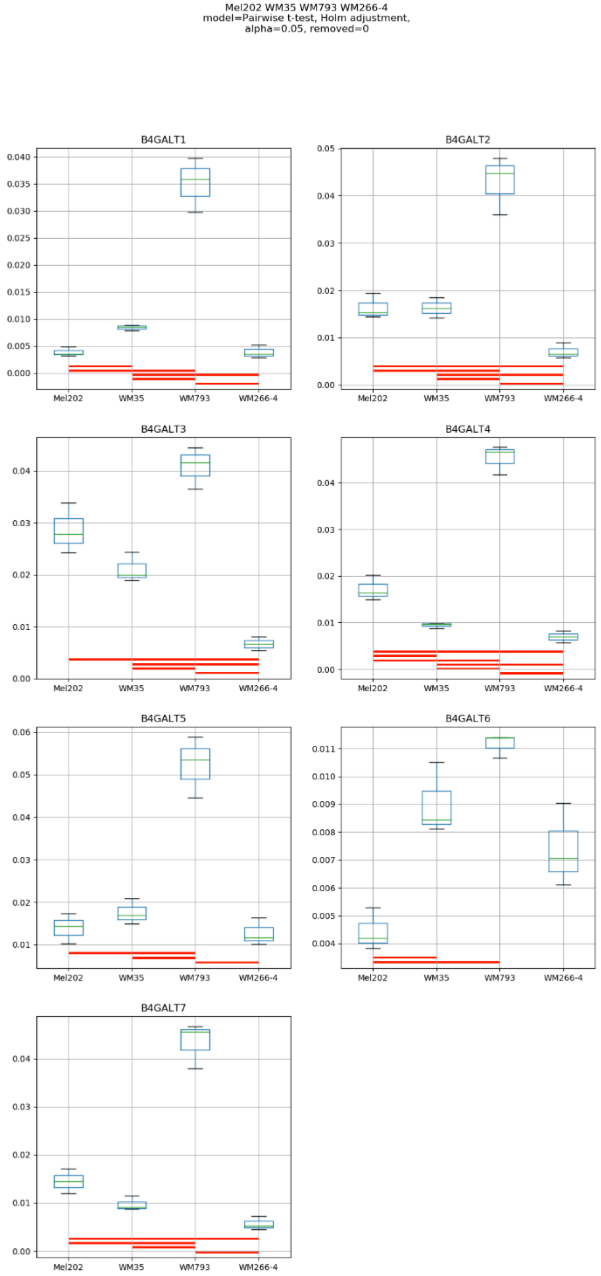

Mel202 WM35 WM793 WM266-4
model=Pairwise t-test, Holm adjustment,
alpha=0.05, removed=0
B4GALT1
B4GALT2
B4GALT3
B4GALT4
B4GALT5
B4GALT6
B4GALT7
Mel202
WM35
WM793
WM266-4

Model No. 26

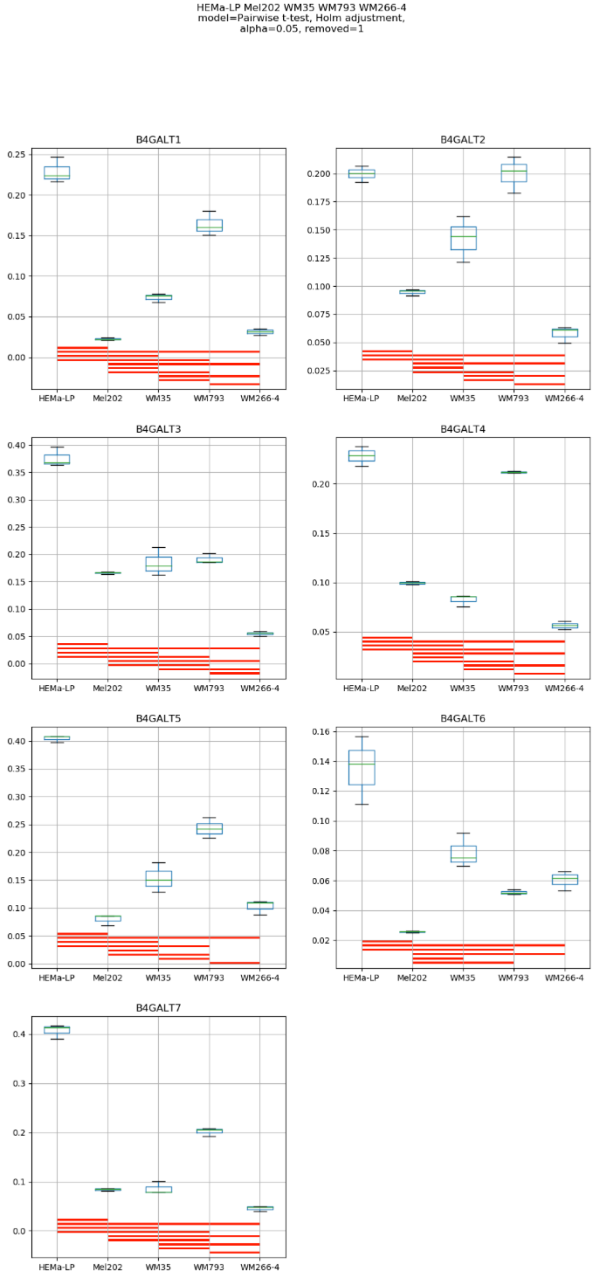
HEMa-LP Mel202 WM35 WM793 WM266-4
model=Pairwise t-test, Holm adjustment,
alpha=0.05, removed=1
B4GALT1
B4GALT2
B4GALT3
B4GALT4
B4GALT5
B4GALT6
B4GALT7
HEMa-LP
Mel202
WM35
WM793
WM266-4

**Supplementary Figure S18:** Medians of relative quantity (RQ) for target genes *B4GALT1–B4GALT7* normalized to the reference gene or pair of reference genes with the best stability values resulting from GenExpA analysis based on set 5 of four candidate reference genes (*GUSB*, *PGK1*, *RPS23, HPRT1*) and remove repetition level 1. The statistical analyses used the pairwise t-test with Holm adjustment. Red line represents statistical significance, $P < 0.05$. Frames denote box-plots representing the medians of the RQ values for a given target gene without a repeatable pattern in the same cell lines under different models (the ‘Remove repetitions’ box set to 0 – violet frame, or to 1 – red frame. Detailed results are listed in Supplementary Table 17.

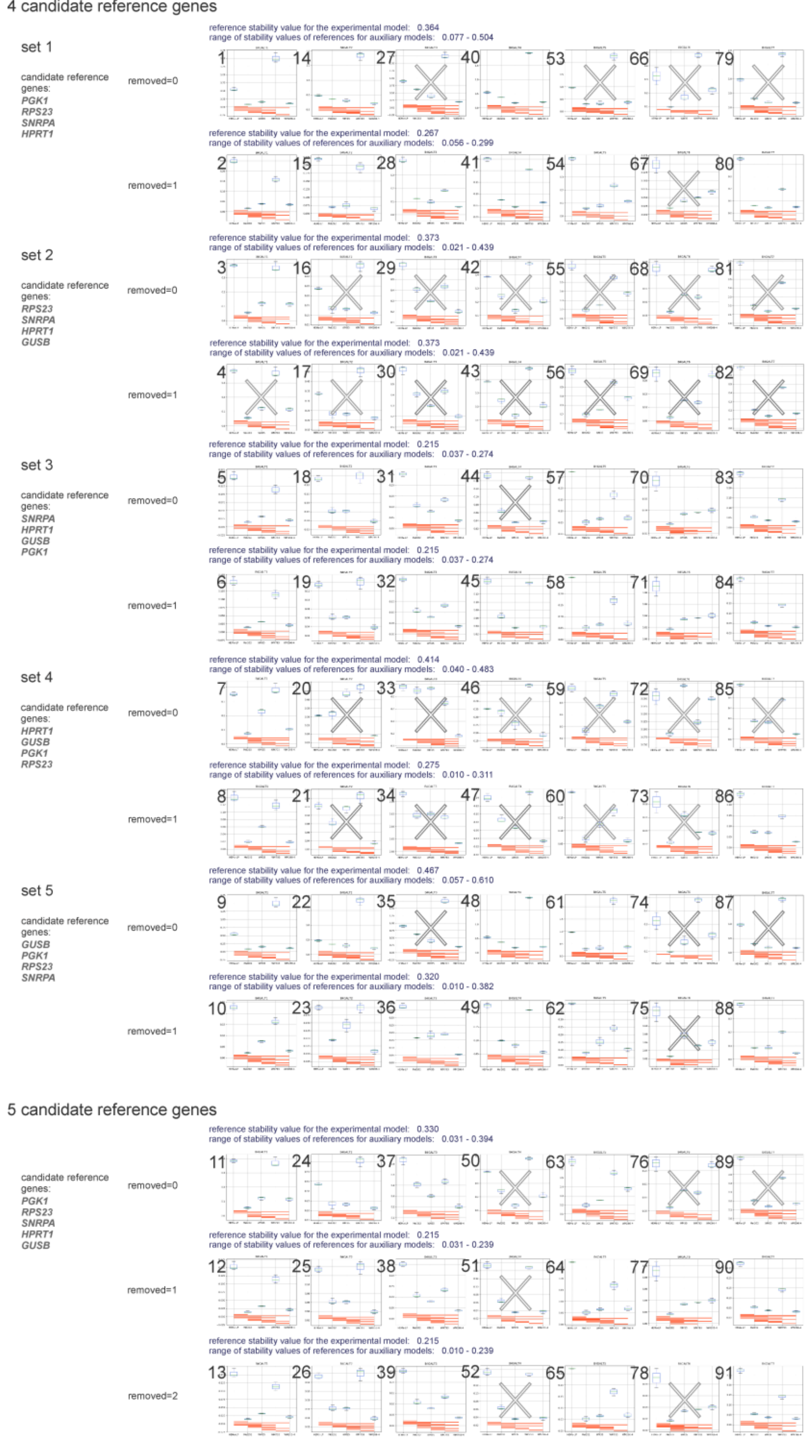


**Supplementary Figure S19:** Medians of relative quantity (RQ) for target genes *B4GALT1*–*B4GALT7* normalized to the reference gene or pair of reference genes with the best stability values in the experimental model of interest resulting from GenExpA analysis based on selection of references from four (sets 1–5; remove repetition levels 0 and 1) and five candidate reference genes (remove repetition levels 0, 1 and 2). A crossed-out box-plot represents unreliable normalization of a given target gene (CS <1). Red line represents statistical significance, P < 0.05.
